# Permuted 23S rRNA is integrated in 50S ribosome particles in *Thermococcus barophilus*

**DOI:** 10.64898/2026.03.23.713588

**Authors:** Christine Gaspin, Isabelle Canal, Régine Capeyrou, Violette Da Cunha, Gabrielle Bourgeois, Clément Madru, Emmanuelle Schmitt, Béatrice Clouet d’Orval, Marta Kwapisz, Marie Bouvier

## Abstract

In Archaea, a prevalent class of circular RNAs corresponds to 16S and 23S ribosomal RNA intermediates (circ-pre-rRNAs). A conserved bulge-helix-bulge (BHB) motif within the 16S and 23S rRNAs processing stems and adjacent to the circularization site in Euryarchaeota and TACK superphylum suggests that pre-rRNAs circularization is widely conserved across Archaea. Using genome-wide transcriptomic data obtained on total RNAs of wild-type *Thermococcus barophilus* cells, we recovered the canonical circularization junctions of the 16S and 23S circ-pre-rRNAs at the predicted BHB motifs. We also identified three alternatives 23S circular junctions introducing variability at the 3’ end of the mature rRNA. We investigated the different forms of 16S and 23S by performing primer extension and RACE experiments. We showed that while the 16S rRNA has standard 5’ and 3’ extremities, the main form of 23S rRNA is circularly permuted, with helix H99 now at its 5’ end. This permutation most probably emerged from the deletion of helix H98 from the 23S circ-pre-rRNA. Interestingly, we showed that the permuted 23S rRNA is incorporated into ribosome subunits and 70S monosomes. The significance of this event in generating functional 50S particles remains to be determined.

## INTRODUCTION

Ribosome biogenesis is a fundamental cellular process required for the translation of messenger RNAs (mRNAs) into functional proteins. Ribosomes are composed of ribosomal proteins and ribosomal RNAs (rRNAs). Archaeal ribosomes consist of a small 30S subunit containing the 16S rRNA, and of a large 50S subunit composed of 23S and 5S rRNAs. In archaea, the genomic organization of the rDNA genes is not conserved in the different archaeal phylum (1). In Euryarchaeota, the rDNA genes are typically found together in a single *rrn* operon, arranged in the order 16S-23S-5S. However, in the Thermococcales genus, the 5S genes are separated from the 16S-23S operons. In the TACK superphylum, the 5S rDNA is also usually dissociated from the 16S-23S operon. When in operons, the rDNA genes are separated by variable regions called internally transcribed spacers (ITS), while the regions that surround the operons are called external transcribed spacers (ETS). Another difference between the archaeal phylum is the copy number of the rDNA genes. According to the rrnDB database (2), in 2025, the number of 16S rDNA copies varies from one to five copies in the 490 archaeal genomes of the 5.10 version. The majority of archaeal genomes possess only one *rrn* operon, and only six genomes are reported with five 16S rDNA copies. Genomes with more than one copy of the 16S rDNA gene are mostly from the Euryarchaeota phylum. While the crenarchaeal *rrn* operons often do not encode transfer RNAs (tRNAs) in their intergenic sequences, tRNAs are often present between the 16S and 23S rDNAs in Euryarchaeota.

The ribosomal RNAs are transcribed in the form of precursors (pre-rRNA) that are processed to give rise of the mature functional rRNAs (3). In Archaea, these maturation steps are for the most part uncharacterized, and the enzymes responsible of the pre-rRNA processing remain to be identified. Nevertheless, in recent years, some progresses have been made in a handful of organisms. Long read nanopore sequencing of total RNAs from the Euryarchaeota *Haloferax volcanii* and *Pyrococcus furiosus*, and the Crenarchaeota *Sulfolobus acidocaldarius* determined the maturation stages of their 16S and 23S rRNAs (4). The archaea specific RNase W was also shown to be involved in the processing of the 16S pre-rRNA in *Haloferax volcanii* (5). Previously, the archaeal Rio proteins, as well as the endoribonuclease Nob1, have been implicated in the maturation of the 16S rRNA 3′end in archaea (6–8).

One intriguing step of archaeal rRNA maturation that is known about for a long time and that seems to be conserved in phylogenetically distant archaeal species is the circularization of the 16S and 23S pre-rRNAs (8–11). The 5’ leader and 3’ trailer sequences flanking the rRNAs form long stems that contain a characteristic Bulge-Helix-Bulge (BHB) motif that is recognized and cleaved by the conserved EndA RNA-splicing endonuclease. The generated pre-16S and pre-23S rRNAs are then ligated by a ligase (presumably RtcB ligase, like tRNA introns removal (8,12), to release a circular pre-rRNAs (circ-pre-rRNAs) molecules. A by-product is also formed, as the ETS and ITS, adjacent to the BHB motifs, were shown to be also ligated in several archaeal species (4,9,10). The BHB motif was shown to be well conserved within the 16S and 23S rRNAs stems in Euryarchaeota and TACK superphylum suggesting that pre-rRNAs circularization is widely conserved across Archaea (3). Some archaeal species are missing BHB motifs at the stem of their rRNAs, and their mature ends are generated by a second splicing-independent pathway (13,14). As a consequence to the circularization of rRNAs precursors, short reads RiboMethSeq and long reads Nanopore sequencing showed that the 23S rRNA of *P. furiosus* exists as a circularly permuted rRNA (4,15). This permutated rRNA does not seem to be conserved since it was not observed in *H. volcanii* and *S. acidocaldarius* (4).

In the present study, we examined the rRNA products of maturation using *Thermococcus barophilus* deep Illumina short-read sequencing on total RNAs. First, we identified a non-coding RNA in the 16S and 23S *rrn* operon, upstream of the 16S rRNA, that is conserved across all the Thermococcales genomes. With a particular focus on the 16S and 23S circ-pre-rRNAs, we extracted the most abundant circularization junctions, and not surprisingly, they correspond to the canonical circularization junctions of the 16S and 23S circ-pre-rRNAs at the predicted BHB motifs. Intriguingly, we also identified additional abundant circularization junctions corresponding to alternative forms of 23S circ-pre-rRNAs. Moreover, altogether the 23S junctions appeared to be more abundant than the 16S ones. Performing primer extension and RACE analyses, we provide the first evidence showing that the permuted 23S rRNAs which contain the BHB-produced junction are linear molecules incorporated in functional ribosome particles.

## MATERIAL AND METHODS

### Sequences of Thermococci genomes and rRNA operons

Based on the 2025-year release, we obtained 34 Thermococcales complete reference genomes and rRNA annotations from NCBI microbial genomes by using the taxa name Thermococcales and filters keeping reference and annotated complete genomes while excluding atypical genomes, MAGs and genomes from large multi-isolated projects (**Supplementary Table S1**). Conserved sequences of ribosomal operons were obtained by extracting and aligning sequences covering 16S and 23S between downstream and upstream CDS. Resulting sequences were aligned combining automatic sequence alignment using multalin (16) and manual finishing for local secondary structures alignment. The Tba19 H/ACA annotation was retrieved by submitting to RFAM 14 (17), the conserved nucleotide sequence downstream the 16S rDNA.

### Identification of circularization junctions in ribosomal operons

We used chimeric alignments provided by STAR aligner (18) to identify positions of circRNAs. In STAR, a chimeric alignment of a read consists of a read that aligns into two segments, each segment being non-chimeric on its own, but the segments are chimeric to each other (i.e. the segments belong to different chromosomes, or different strands, or are far from each other or don’t align in a linear way against the genome sequence). Setting the --chimOutType parameter to ‘Junctions’ allows generating chimeric alignments in outputs. All RNA-seq fastq files were submitted to STAR-2.7.5a for alignment with options --alignIntronMax 6000 -- alignMatesGapMax 300 --chimSegmentMin 15 --chimOutType Junctions. Circularization junctions were identified by filtering and counting chimeric alignments from the junction file output from STAR. We used strand and position columns to search all chimeric alignments for reads in which the two segments of the read align on the same region and on the same strand, with the 5′ segment aligning downstream of the 3′ segment. We kept those that aligned within the rRNA operon for further analysis.

### Cell culture

Strains were grown anaerobically at 85°C, at atmospheric pressure (0.1 MPa) in TRM medium (19). 2X TRM (46 g.L^-1^ NaCl, 10 g.L^-1^ MgCl □ ·6H □ O, 6.9 g.L^-1^ PIPES, 8 g.L^-1^ tryptone, 2 g.L^-1^ extrait de levure, 1.4 g.L^-1^ KCl, 1 g.L^-1^ (NH □) □ SO □, 0.1 g.L^-1^ NaBr, 0.02 g.L^-1^ SrCl □ ·6H □ O; pH 6.8) was sterilised (121 □ °C, 20 min), diluted to 1X and supplemented with 25 mg.L^-1^ K □ HPO □, 25 mg.L^-1^ KH □ PO □, 5 µM CaCl □ ·2H □ O, 125 µM FeCl □ ·6H □ O, 5 µM Na □ O □ W, 8 g.L^-1^ of sulfur S □ and 0.1 g.L^-1^ of Na □ S. Cells were collected in exponential (6h) and stationary (16h) phases by centrifugation at 7 500 g at 4°C for 15 minutes.

### Extraction of total RNAs

Total RNA was extracted as described previously (20). Cell pellet was dissolved in an equivalent volume of denaturing buffer (4 M guanidium thiocyanate, 42 mM sodium citrate, 0.83 % N-laurosylsarcosine pH 7, and 0.2 mM β-mercaptoethanol). 0.1 mL of 2 M sodium acetate pH 4, and 1 mL of phenol-chloroform-isoamyl alcohol (25:24:1) were added per 1 mL of resuspended cells. After 15 min of incubation on ice, the mixture was centrifuged for 20 min at 10 000 g at 4°C. The aqueous phase was recovered and RNA was precipitated for 1 hour at -20°C with an equal volume of isopropanol. The RNA was recovered after centrifugation for 20 min at 10 000 g and 4°C, then resuspended in 1.5 mL of 4M LiCl and pelleted by centrifugation for 20 min at 10 000 g at 4°C. The RNA was then resuspended in denaturing buffer, precipitated with 100% ethanol and washed twice with cold 75% ethanol. The ethanol was air dried and the RNA was dissolved in water. DNA potentially contained in the sample was degraded using 3U of Turbo DNase (TURBO DNA-free kit™; Invitrogen) according to the supplier’s protocol. The RNeasy MinElute Cleanup kit (Qiagen) was used to purify the RNA, which was then measured by the NanoDrop 1000 spectrophotometer (Thermo Fisher Scientific). Finally, the RNA was stored at - 80°C.

### Reverse Transcription and cDNA amplification

The cDNAs were generated by reverse transcription (RT) according to SuperScript™ III protocol (Invitrogen). The reactions were performed on 2 μg of DNA-free total RNA using random primers (hexamers, Promega™), in presence of RiboLock RNase Inhibitor (Thermo Fisher Scientific). Amplicons for the circular junction analyses were produced on cDNA template (1 μL of a 10-fold dilution of RT reaction) using primers specific for 16S (16S-262R & 16S-1Fj) and 23S (23-214R & 23S-1Fj). The amplicons were separated on a 1.5 % agarose gel, together with the 100 bp GeneRuler (Thermo Fisher Scientific). The nucleic acids were detected using SYBR™ Safe (Invitrogen).

### Primer extension

The 5’ ends of the 16S and 23S rRNAs were determined by primer extension analyses according to the SuperScript™ IV protocol (Invitrogen). The reverse transcriptions were performed on 1 μg of DNA-free total RNA extracted from cells in exponential or stationary phases, with 2 pmol of 5’-end radio-labeled oligonucleotide and 40 U of RiboLock RNase inhibitor. The cycling conditions were 25°C for 5 min and 55°C for 55 min, followed by a treatment with 2.5 U of RNase H (Thermo Fisher Scientific) at 37°C for 20 min. A radiolabeled ladder was also generated by first dephosphorylating 8.5 μg of GeneRuler 100 bp (Thermo Fisher Scientific) with 20 U of CIAP alkaline phosphatase (Invitrogen). A fourth of the reaction was then phosphorylated with 5 μCi of γ-^32^P ATP and 10 U of T4 polynucleotide kinase (Thermo Fisher Scientific) and diluted 10-fold with loading buffer II (LBII, Invitrogen). 2 μL of primer extension products, denatured with 5 μL of LBII, were separated, together with 2 μL of ladder, on a denaturing 6 % polyacrylamide (#1610144, Biorad) / 7 M urea sequencing gel. The gel was dried and signals were visualized on a phosphorimager (Cytivia™).

### Rapid Amplification of cDNA ends (RACE)

The 5’ and 3’ ends of the 16S and 23S rRNAs were precisely mapped performing 5’ and 3’ RACE analyses, respectively. These 5’ and 3’ ends found in databases can vary depending on the version of the genome annotation used (**Supplementary Table S2**). Here, we used the lastest NC_014804.1 annotation. The RACE experiments were performed on DNA-free total RNA extracted from cells in stationary phases, as described in (21).

#### 5’ RACE analysis

Since the 16S and 23S rRNAs are processed RNAs, their 5’ end are monophosphorylated. Therefore, the DNA-free total RNAs were used as is. The RNA-adapter ligation was performed overnight at 17 □C in the presence of 1 μg of RNAs, 21 pmol of RNA adapter (RNA A3), 10 U of T4 RNA ligase (Thermo Fisher Scientific), 1X RNA ligase buffer containing ATP, 15 % DMSO, and 40 U of RiboLock RNase inhibitor. After a phenol/chloroform RNA extraction, the reverse transcriptions were done using a fourth of the adapter-ligated RNAs and 16S and 23S gene-specific oligonucleotides (16S-190R and 23S-214R, respectively) as described above (Primer extension). After treatment with 1 U of RNase H, 2 μL of the cDNA samples were used as template for a PCR reaction using 1 U of Phusion™ High-Fidelity DNA polymerase (Invitrogen), 1X GC buffer, 0.2 mM dNTPs, 3% DMSO, and 1 μM of each oligonucleotide. While the forward oligo is specific to the adapter (B6), the reverse oligo is specific to the gene of interest. Here, we used the same that were used for the RT. Following separation on 3% agarose gels, PCR products were excised, purified, cloned in pCR™-Blunt II-TOPO (Zero Blunt TOPO PCR cloning kit, Invitrogen), and sequenced by Eurofins Genomics using M13R or M13F oligos.

#### 3’ RACE analysis

They were performed on RNAs with 5’-OH RNAs to avoid the adapter ligation at the RNA 5’ end. Briefly, 10 μg of DNA-free total RNAs were dephosphorylated using 40 U of CIAP alkaline phosphatase at 37°C for 45 min, in presence of 40 U of RiboLock RNase inhibitor. The RNA preparation was then cleaned by a phenol/chloroform extraction. The RNA-adapter ligation and RT reactions were performed as described above using the RNA E1 and the adapter-specific oligonucleotide E4. The PCR amplifications of the generated cDNA were done using forward gene-specific oligos for the 16S and 23S rRNAs (16S-263F and 23S-267F) and the adapter-specific oligo E4. The PCR products were analyzed as described above.

### The DNA and RNA oligonucleotides

The DNA oligonucleotides were generated by Eurofins Genomics. Their sequences are as follow: **16S-262R**: TAC CCC ACC AAC TAC CTA ATC G; **16S-1Fj**: GAT CAC CTC CTA TCG CCG GAA A; **16S-190R**: CTT CCA GTA CCC CAC ACC TAT G; **23S-214R**: CAT CGC TTT TGC TTT CTT TTC; **23S-1Fj**: CAG CCC GAG TTT CTT GCC CCT; **23S-136R**: CAA AAG CCG CGG GAG GTC; **o5’RACEforward (B6)**: GCG CGA ATT CCT GTA GA; **16S-263F (B24-08)**: CCC GCC CTC AGT TCG GAT CGC; **23S-267F (B24-11)**: CAG GGT GCC GCG GCC TCT GG; **o3’RACEreverse (E4)**: GGC CGC TAA GAA CAG TGA A; **M13Forward (B23-88)**: CAC ACA GGA AAC AGC TAT GAC; **M13Reverse (B23-89)**: GTT GTA AAA CGA CGG CCA GT.

The RNA oligonucleotides used as adapters in the RACE experiments were produced by Dharmacon™. **RNA adapter A3** for the 5’ RACE: 5’-AUA UGC GCG AAU UCC UGU AGA ACG AAC ACU AGA AGA AAG-3’. **RNA adapter E1** for the 3’ RACE: 5’-phosphate-UUC ACU GUU CUU AGC GGC CGC AUG CUC-idT-3’.

### Ribo Mega-SEC

A total of 250 mg of cell pellet was re-suspended in buffer SEC100 (20 mM HEPES pH 7.5, 100 mM NH4Cl, 20 mM Mg Acetate, 2.5 mM DTT) containing a cocktail of EDTA-free protease inhibitor (cOmplete TM, Roche) with a ratio of 1 ml of buffer for 450 mg of cell pellet. Whole-cell extracts were prepared by lysis with one volume of glass beads (Ref. Sigma G4649, <106µm) using a homogeizer (Precellys) (4 × 30 sec at 5500 rpm at 4°C). Glass beads were removed by centrifugation at 5,000 g for 5 min. Cell lysates were cleared by centrifugation at 16,000 g for 10 min at 4°C and filtered through 0.45 µm Ultrafree-MC HV centrifugal filter units (Millipore) by 10,000g for 3 min before injection on SEC columns. Proteins and RNA amounts were quantified by Bradford protein assay and Nanophotometer respectively.

The profile of ribosomes was analyzed by using two sequential columns Agilent Bio SEC-5, a 2,000 Å column and a 1,000 Å column, installed in an AKTA Pure 25M system (Cytiva). SEC columns were equilibrated with two column volumes of filtered SEC100 buffer (20 mM HEPES pH 7.5, 100 mM NH4Cl, 20 mM Mg Acetate, 2.5 mM DTT). 100 µL of cell lysate (1.5 mg) was injected onto the columns. The flow rate was 0.5 ml/min. The chromatogram was monitored by measuring UV absorbance at 280 and 254 nm. Fractions of 500µl were collected from 9 min to 22 min. 250 µl of each fraction was precipitated with 15% trichloroacetic acid (TCA) and used for Western blotting with antibodies against the ribosomal proteins S4e and L10e (diluted 10,000-fold, raised against *Pyrococcus abyssi* recombinant proteins, Eurogentec). 250 μL of fractions were used for TRIzol-based RNA extraction (TRI-reagent, Euromedex). The primer extensions were performed as described previously using, with halves of the RNAs extracted.

### Molecular modeling

The cryo-EM structure of the 50S ribosomal subunit (PDB 6SKF) was modified using Coot (22) in the *Thermococcus kodakarensis* 50S unsharpened map (EMD-10223). Minor adjustments were made to the original model: the 23S rRNA region spanning residues 2934-3036, which corresponds to helices H99-101, was renumbered 1-103, and U3011 was added to its 3’ end. The final model was locally refined using Coot’s real space refine zone and regularize tools. Figures were prepared using USFC ChimeraX (23). The 23S permuted rRNA of *T. kodakarensis* was designed based on sequence homology with the 23S rRNA of *T. barophilus*.

### Conservation of H98 in archaeal 23S rRNAs

The fasta sequence of the 23S rRNA gene of each archaeon was obtained using NCBI, Rfam databases. R2DT was used for the prediction of the 23S rRNA structure (24). R2DT is based on a library of 3,647 templates of known structured RNAs. In most cases, the structure was obtained using the PF_LSU_3D template. The two-dimensional editing of the predicted structure was done using RNAcanvas (25), that provides interactive editing features.

## RESULTS

### In Thermococcales, the ribosomal operons exhibit a conserved organization, with a box H/ACA non-coding RNA and BHB motifs

We selected 34 complete reference Thermococcales genomes, along with their annotations, and searched for ribosomal DNA (rDNA) operons and genes. Of the 34 genomes analyzed, 33 carried a single 16S-23S rDNA operon. While *Palaeococcus pacificus* was the only genome found to carry two. These operons are organized as previously described (3), with the 16S and 23S separated by a tRNA^Ala^. Theses genomes also all contain two 5S rRNA genes. In *T. barophilus*, one is monocistronic and one is dicistronic together with a tRNA^Asp^ (**Figure 1A**).

**Figure 1:**
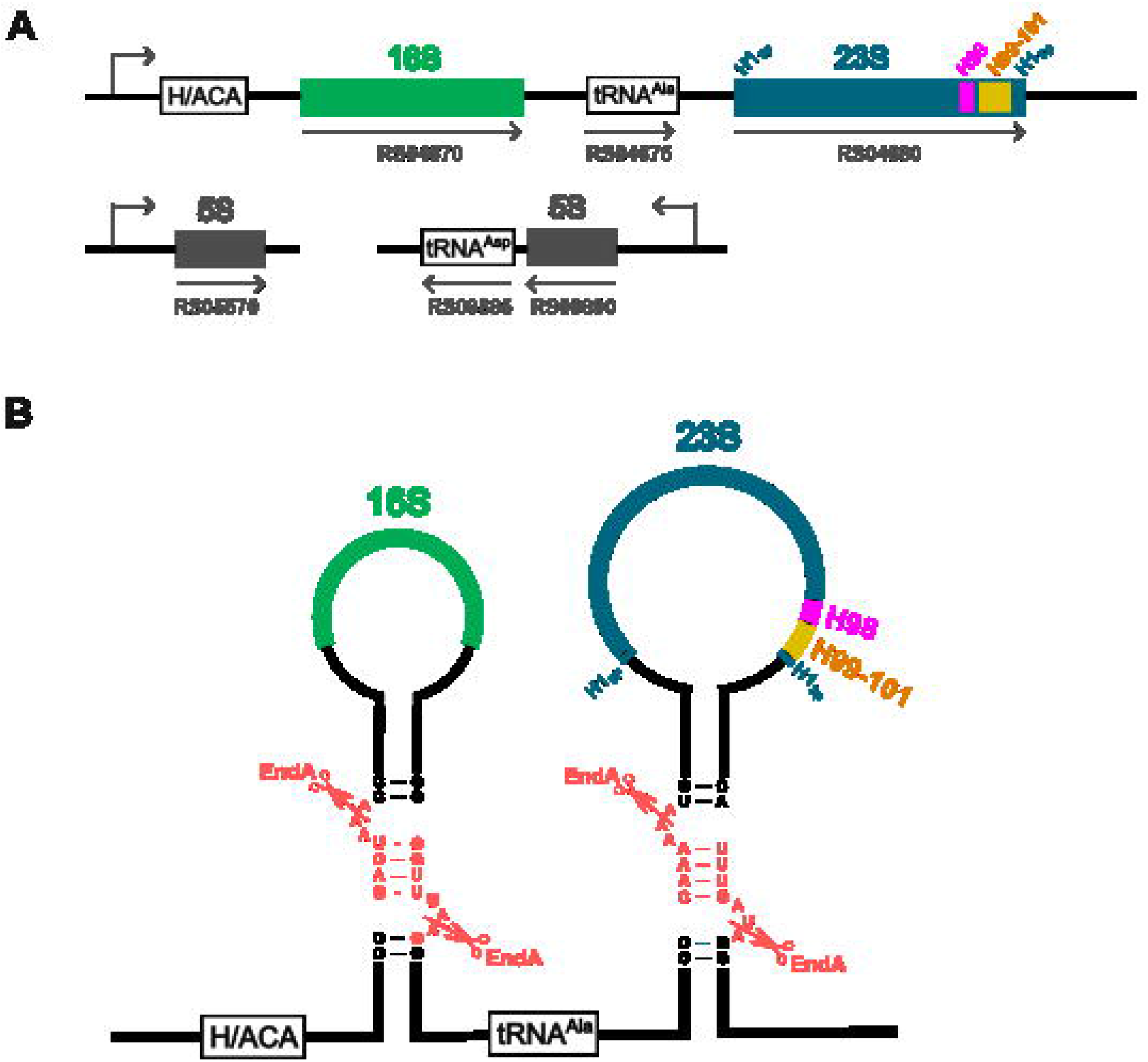
Ribosomal RNAs in *Thermococcus barophilus*. The organization and the number of rDNA genes are conserved across the Thermococcales order. Only *Palaeococcus pacificus* has two copies of the 16S-23S rRNA operon (*rrn*). **(A)** The *rrn* operon is composed of four genes coding for: an H/ACA guide RNA (Tba19), a 16S rRNA (RS04670/TERMP_022095, in green), a tRNA^Ala^ (RS04675/TERMP_022096) and a 23S rRNA (RS04680/TERMP_022097, in blue). There are two copies of 5S rDNA, one is monocistronic (RS05570/TERMP_022100) and one (RS09890/TERMP_022118) is in operon with a tRNA^Asp^ coding gene (RS09885/TERMP_022117). The orientation of the transcription units is given by the arrows. **(B)** The 16S and 23S rRNA processing stems contain a BHB motif (light red) that is recognized and cleaved by the endonuclease EndA (cleavage sites are marked by light red scissors). The rRNA molecules are then circularized by ligase RtcB to generate circular intermediates of the 16S and 23S rRNA (pre-circ-rRNA). On the 23S rRNA, the 5’ (H1_5p_) and 3’ (H1_3p_) stems of the helix H1, the helix H98 (pink) and the helices H99-101 (orange) are highlighted since they are of special interest in this study.

To go further, using the break in high conservation of nucleotide sequences of *P. pacificus* rDNA operons, we were able to determine the 5’ and 3’ extremities of a nucleotide (nt) sequence conserved in all 16S-23S rDNA operons of the 34 genomes. By searching against the Rfam library of covariance models, we found that a region of the 5’ ETS, located 200 nt upstream of the 16S rDNA gene, encodes Tba19, a homolog of the *Pyrococcus abyssi* Pab19 box H/ACA non-coding RNA (ncRNA) (**Supplementary Figure S1**). In a previous study, a systematic computational screen of H/ACA ncRNA genes in archaea identified and experimentally validated Pab19, which was predicted to target the U residue for conversion into pseudouridine at position U1017 of the *P. abyssi* (*Pab*) 16S rRNA (26). In *T. barophilus* (*Tba*), based on sequence complementarity, Tba19 should also pseudouridilate the 16S rRNA, but at the position U1002. Based on the 16S rRNA helix annotation found in (27) and their respective 2D structures in RNA central, both *Pab*-U1017 and *Tba*-U1002 were located in the helix h34 of their respective 16S rRNAs. It remains to understand why this specific H/ACA is part of the 16S-23S rRNA precursor.

Finally, we searched and identified the BHB motifs that are formed at the base of the long stems by the sequences surrounding the 16S and 23S rRNAs, in the 16S-23S rRNA precursor of *T. barophilus* (**Figure 1B**).

### Circular precursors of 23S and 16S rRNAs are the most abundant circular RNA species

Building on previous work, we were able to identify the EndA cleavage sites and predict the canonical circularization junctions of the circular 16S and 23S rRNA precursors **(Figure 2A)**. Using genome-wide transcriptomic data obtained on total RNAs of wild-type *T. barophilus* cells, in stationary and exponential phases, we searched for circular junctions by mapping the chimeric reads that align into two segments on the genome. Briefly, we performed deep sequencing of three paired-end replicates of total RNA, including the ribosomal RNA, and obtained over 60 million of paired-end reads for each dataset (**Supplementary Table S3**). After mapping the reads against *T. barophilus* using STAR (18), we extracted circularization junction candidates. Among them, seven different junctions have high read support (**Table 1, Supplementary Figures 2a-e and 3**). Five of these correspond to circular precursors of rRNAs (circ-pre-rRNAs): four candidate junctions are 23S circ-pre-rRNAs and one candidate junction is a 16S circ-pre-rRNA. We recovered the canonical circularization junctions of the 16S and 23S circ-pre-rRNAs formed at the predicted BHB motifs (**Figure 2A**), and three novel alternatives 23S circular junctions. The canonical junctions were confirmed by reverse transcription and PCR amplification using pairs of oligonucleotides specific of the junctions (**Figure 2B**).

**Figure 2:**
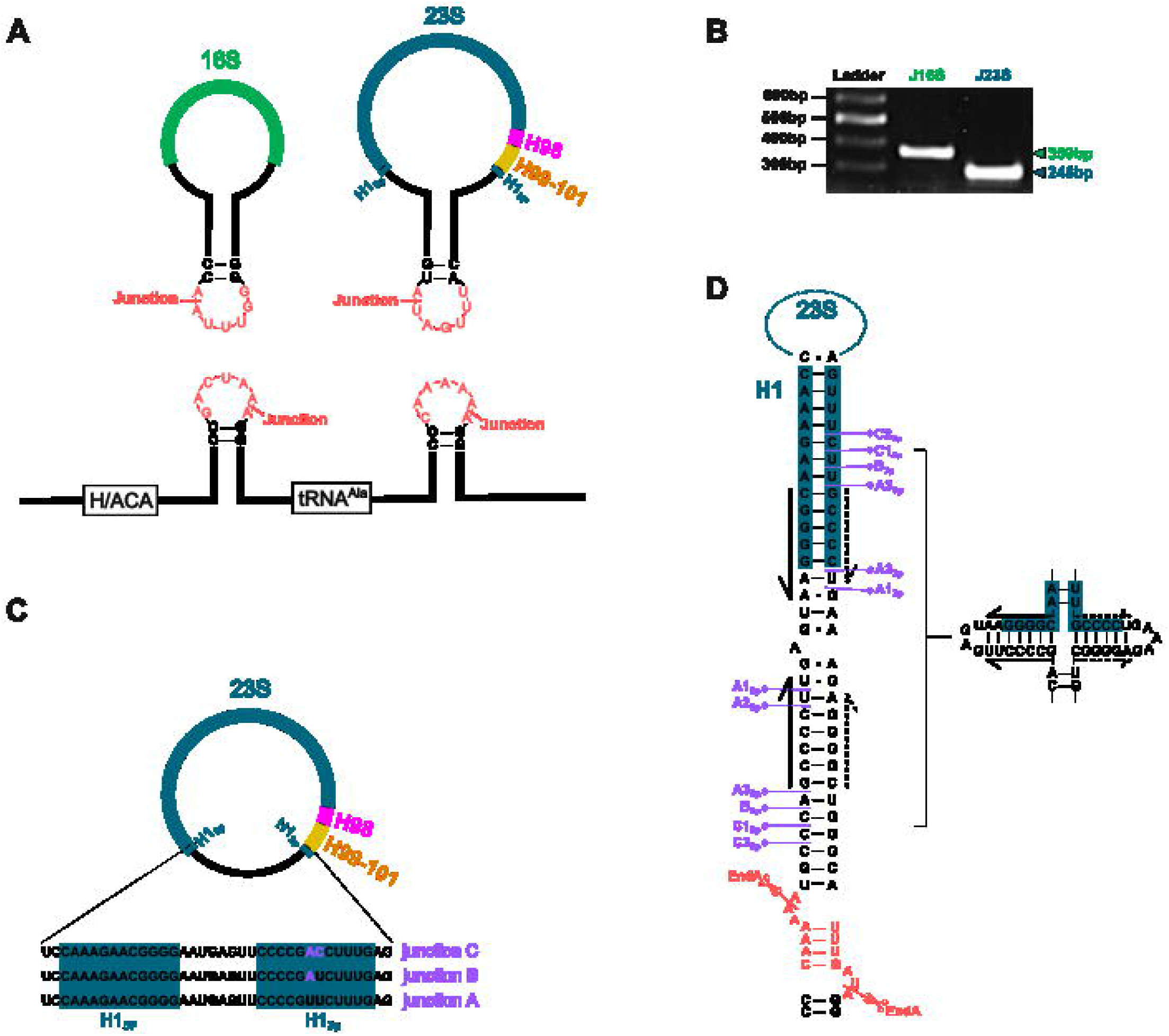
The circular forms of 16S and 23S. **(A)** Representation of the canonical 16S and 23S pre-circ-rRNAs, circularized at BHB motifs. The resulting linear product is shown at the bottom. **(B)** The circularization junctions of 16S and 23S rRNAs were amplified from cDNA by PCR using specific primers adjacent to the junctions (262R/1FJ for 16S and 214R/1FJ for 23S). **(C)** Three alternative 23S circular junctions were identified (A, B and C, in green, red and purple, respectively). Based on their sequences, they emerged from helix H1, that is formed by the 5’ (H1_5p_) and 3’ (H1_3p_) ends of the 23S rRNA. **(D)** The helix H1 is extended by the 5’ leader and 3’ trailer sequences surrounding the 23S rRNA. This extended helix that is composed of inverted repeats can adopt two distinct conformations. The putative cleavage sites are indicated by diamond arrows on each side of the stem. Junction B can only be generated by one cleavage site. For junctions A and C, three and two putative cleavage sites are possible, respectively. The BHB motif and EndA cleavage sites are also represented in light red.

**Table 1:**
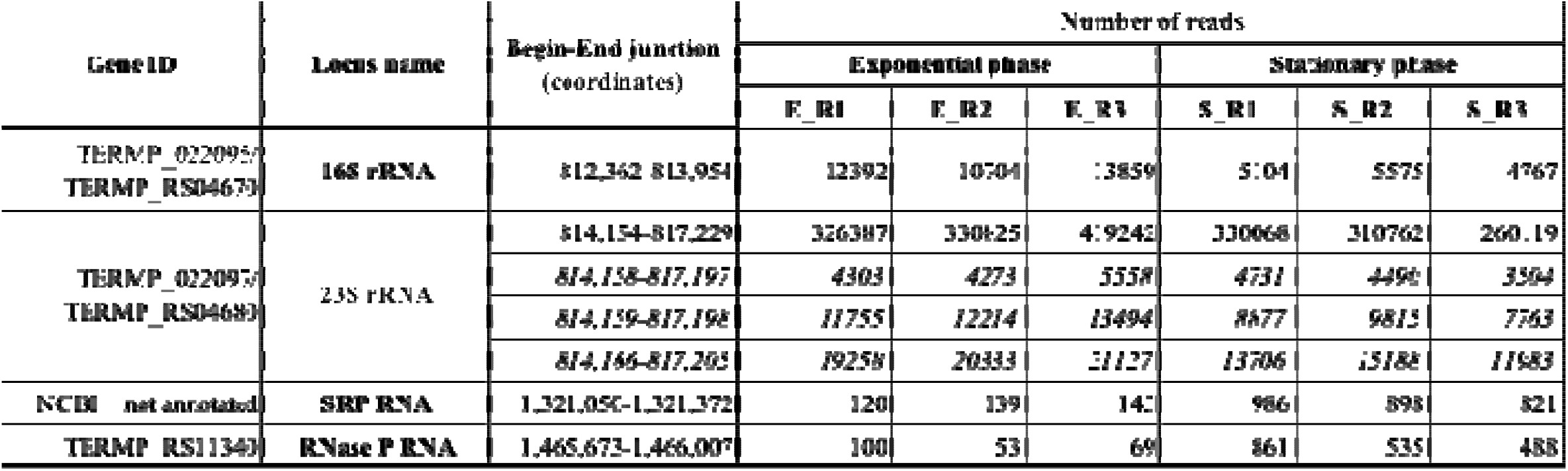
Junctions with high read counts identified using STAR aligner. The 23S rRNA alternative junctions are in italics.

Aside from the ribosomal RNAs, we also retrieved the junctions of the circular SRP and RNase P RNAs (10,28). As expected, these molecules are far less abundant than the circ-pre-rRNAs. To investigate differences in growth phases with respect to the quantity of junctions, we calculated their percentage of the total number of reads (**Table 1 and Supplementary Table S3**). This revealed 7- and 9-fold increases in circular SRP and RNase P RNAs, respectively, during the stationary phase. On the other hand, while the 16S junction appears to be a little less abundant in stationary phase (2-fold), the 23S canonical and alternative junctions seem to be equally present in both phases. Aside from the growth phase, the most striking difference is the one existing between the 16S and 23S junctions. Indeed, the reads covering the 23S canonical junction are 29 times more abundant than those corresponding to the 16S junction, despite both being expressed from an operon. The following experimental work will demonstrate that the 23S circularization junction is conserved in a linear permuted 23S rRNA molecule that is incorporated into ribosomal subunits

### Identification of three novel alternative 23S pre-circ-rRNAs

Three novel alternative circ-pre-23S rRNA were also identified, with closely related junctions (**Figure 2C**). Since these junctions only involve eight nucleotides outside the annotated rDNA, and that they are less prevalent than the canonical BHB-produced junction, molecular biology approaches were not appropriate to confirm their existence. Based on alternative junction sequences, we searched for cleavage/ligation sites (**Figure 2D**). While junction B can only occur at one site, junctions A and C can be produced at different sites. All are formed by cleavage/ligation in the helix H1 formed by the complementary 5’ and 3’ ends of the 23 rRNA, and extended by their respective leader/trailer sequences. Since it is composed of inverted repeats, two distinct structures can be adopted. Interestingly, junctions B and C introduce variability in the 23S 3’ end sequence (H1_3p_). This can impact the pairing of the 5’ and 3’ ends to form the terminal helix. These modified nucleotides in the 23S 5’ end sequence (H1_5p_) can be found in the 23S of other Thermococcales, but they are associated with complementary modifications in the H1_5P_ to keep the helix H1 intact (**Supplementary Figure 4**). Indeed, a common aspect of ribosome assembly, conserved across all domains of life, is the establishment of connections between the 5′ and 3′ ends of the large subunit rRNA to initiate rRNA domain compaction and subunit assembly (29).

### Specificities of the permuted 23S rRNA

To generate a mature rRNA from a circ-pre-rRNA, the circular molecule needs to be reopened by ribonucleolytic cleavages to generate the 5’ and 3’ ends. Depending on the reopening site and any further maturation steps, rRNA maturation is a complex and regulated process that generate mature linear rRNAs with defined ends. In our *T. barophilus* data set, we observe that only few reads cover the 23S helix H98 (**Figure 3**; **Supplementary Table S4**). Interestingly, many reads in yellow are found both upstream and downstream of what is expected to be the mature 23S rRNA, as annotated. When looking closer, it corresponds to sequence flanking the EndA cleavage sites that are then ligated in the 23S circ-pre-rRNA. Altogether, the data suggest that the 23S rRNA is also circularly permuted in *T. barophilus*. To go further, we chose to use established molecular biology techniques to map the precise extremities of the 16S and 23S rRNA, in total RNAs, but also in cellular fractions enriched in ribosome subunits.

**Figure 3:**
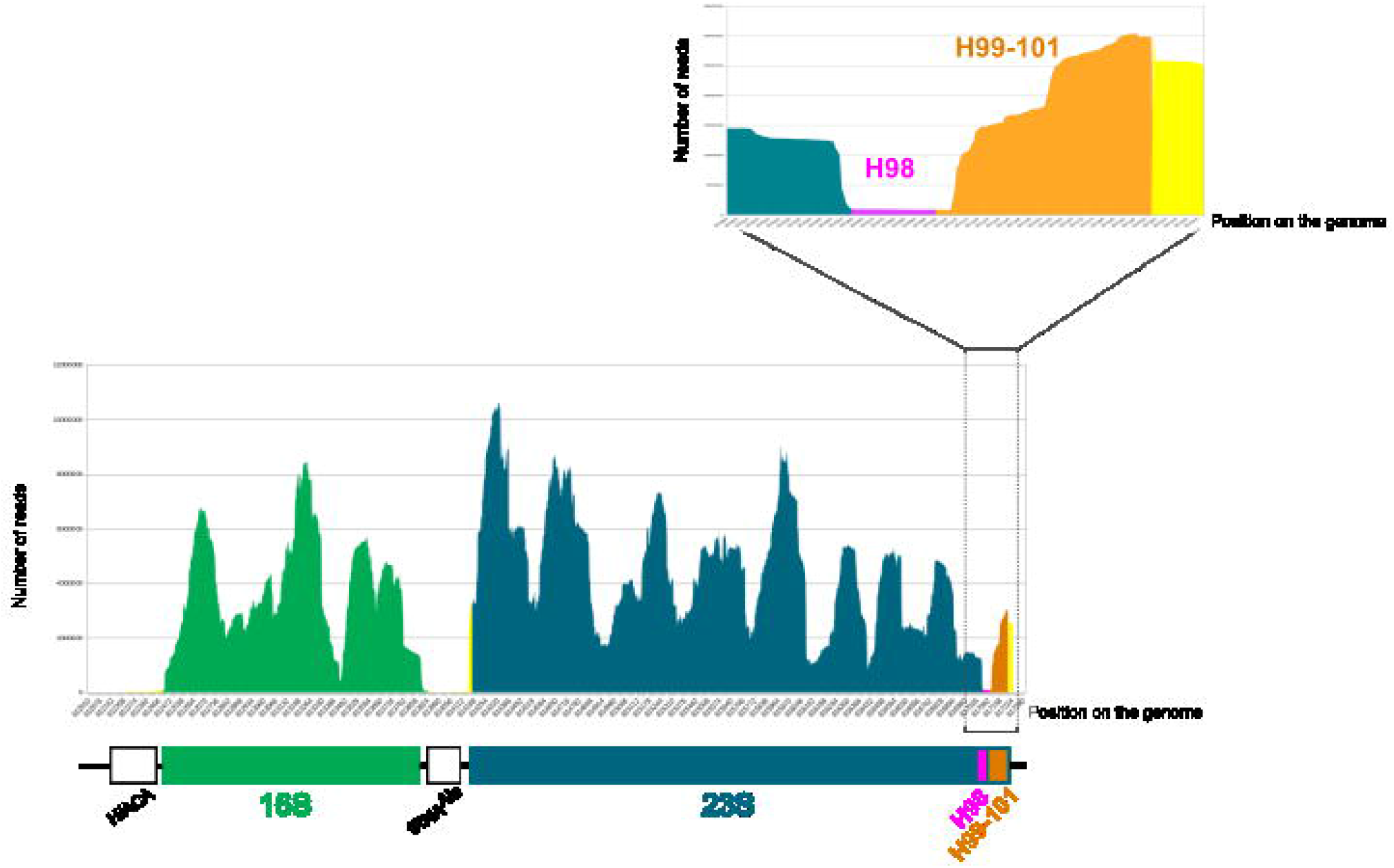
RNA-seq reads alignment on the 16S and 23S rDNAs. The reads from the total RNA transcriptome were aligned on the sequence of H/ACA-16S-tRNA-23S operon using BWA. The 16S reads are in green and the 23S are in blue. The coverage of the helices H98 and H99-101 of the 23S rRNA are in pink and orange, respectively. For the purpose of simplification, H1_3p_ is integrated in H99-H101. The rest of the operon is in yellow. A focus on the 3’ end of 23S is made. Only few reads align to the helix H98.

Of note, **in Figure 3**, it is also interesting to observe that almost no reads cover the Tba19 ncRNA and the tRNA^Ala^, while they are in operon with the 16S and 23S rRNAs. We also had difficulties to probe them by Northern blotting. Most likely, they were lost during the total RNA extraction process due to the pH used or were not effectively reverse transcribed due to their structures and/or modifications.

First, we performed primer extensions to compare the actual 5’ ends of the 16S and 23S rRNAs with those annotated in the *T. barophilus* genome (**Figure 4**). It should be noted that in the most recent NC_014804.1 annotation, the extremities of the 16S rRNA are very different from those found in the previous NC_014804 one, that is still used in some database to this day (**Supplementary Table S2**). For the 16S rRNA, its 5’ end is as expected, at around 190 nucleotides from the chosen primer (16S-190R, **Figure 4**). The upper band marked with an asterisk corresponds to an unspecific product (see below). Here, no significant difference is observed between the exponential and stationary growth phases. For the 23S rRNA, the primer extensions were performed with two oligonucleotides, with their 5’ ends hybridizing the positions +136 (23S-136R) and +214 (23S-214R) of the annotated 23S rDNA (**Figure 4**). While the primer extensions give rise to faint bands of 136 and 214 nucleotides, respectively, the main bands are located above and correspond to extended forms of 23S rRNA (around 302- and 380-nt, respectively). The profiles obtained for both oligos, are similar which is consistent with a permutation event that would generate a new 5’ end.

**Figure 4:**
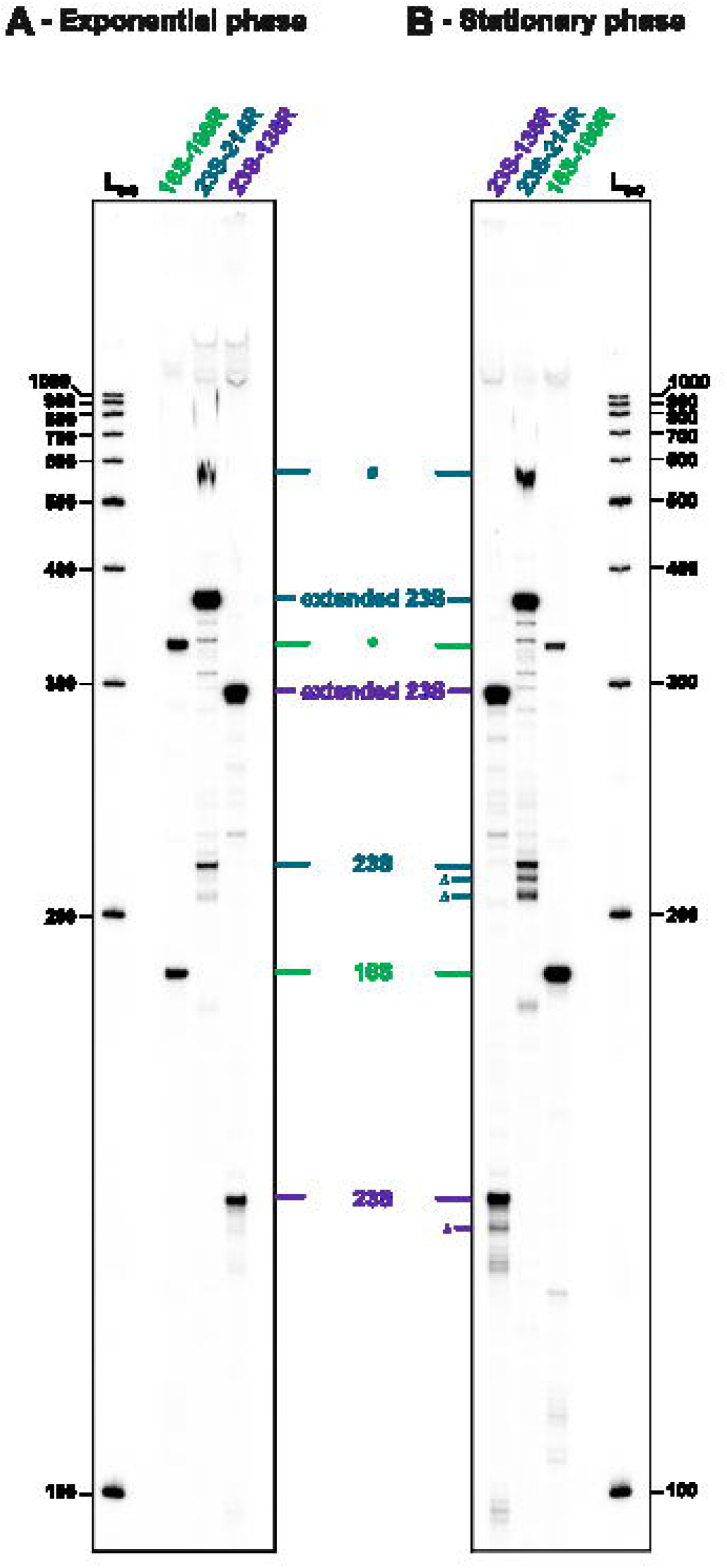
Mapping of the 16S and 23S 5’ ends. Primer extensions were performed on total RNAs extracted from *T. barophilus* cells grown in exponential **(A)** and stationary **(B)** phases. The primer extensions were performed with one primer specific for 16S (16S-190R; in green) and two primers specific for 23S (23S-136R and -214R; in purple and blue, respectively). The detected bands are annotated as follow: 16S and 23S are the bands corresponding to the expected 5’ end based on the genome annotation; Δ, several bands (+/-few nucleotides) that could correspond to the 23S annotated 5’ end; extended 23S, where the 5’ end is upstream to the one annotated on the genome; *, the 16S-190R primer amplifies a non-specific product that has the same 5’ end than the 16S band (see 5’ RACE results below; Figure 5); #, some reaction products are not fully denatured and do not separate correctly on the sequencing gel. L_(nt)_ is a radio-labeled DNA ladder prepared under denaturing conditions.

Secondly, to precisely map the 5’ end nucleotides of the 16S and 23S rRNAs, at the nucleotide level, we performed 5’ RACE experiments. Briefly, the 5’ end of the RNA molecules were ligated with an A3 adapter and reversed transcribed with oligonucleotides specific to the 16S or 23S rRNA. The cDNA was then amplified by PCR with oligonucleotide pairs specific to the adapter and the 16S or 23S rRNA (16S-190R or 23S-214R). The bands of interest are then extracted from the agarose gel, cloned and sequenced. For the 16S rRNA, the PCR product corresponding to the a1 band is at the excepted size around 225 base pairs (bp), the adapter adding 35-nt to the PCR product (**Figure 5A**). After removing the sequence corresponding to the A3 adapter, the sequencing data were aligned with the rDNA operon. All six sequences have the same 5’ end that corresponds to the annotated one (**Figure 5B**). On the RACE gel, an a2 band is also visible on the gel (**Figure 5A**) that seems to correspond the upper band labeled with an asterisk on the primer extension gel (**Figure 4**). This band was also extracted and sequenced to found out that a2 has the same 5’ end than a1 (**Figure 5B**). Upon further investigation, we have found that the 16S-190R oligo partially bind to a sequence located 298-nt downstream of 16S-rRNA. Therefore, it corresponds to an unspecific product.

**Figure 5:**
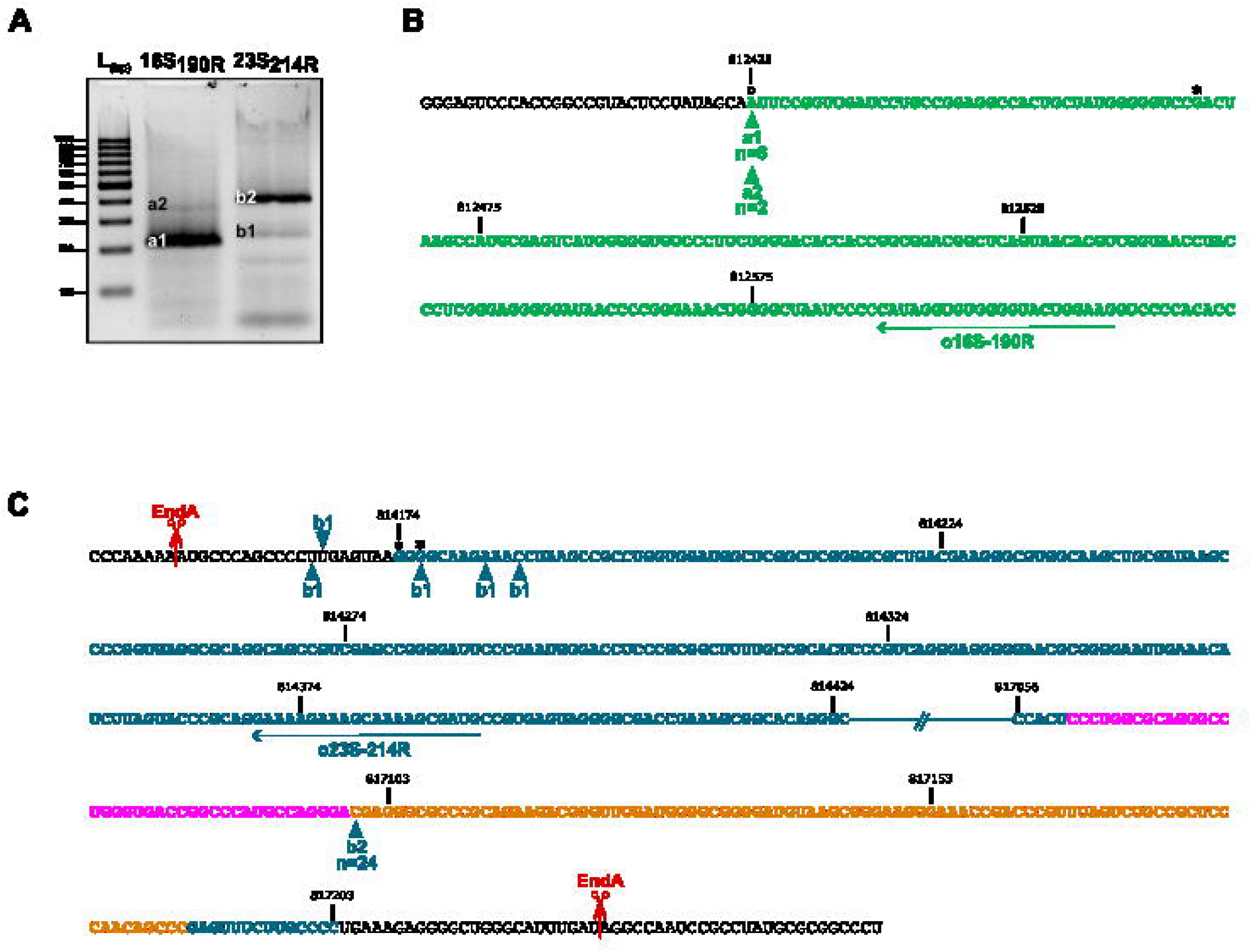
Precise mapping of the 16S and 23S 5’ ends by 5’ RACE. The cellular RNAs that carry a monophosphate at their 5’ end were ligated with an adapter, and reverse transcribed with an oligonucleotide specific for the 16S or 23S rRNAs (16S-190R and 23S-214R, respectively). The resulting cDNAs were amplified by PCR with an oligonucleotide specific for the 5’ end adapter and with 16S-190R or 23S-214R, respectively. The amplification products were separated by electrophoresis on a 3% agarose gel **(A)**. The annotated bands were extracted and cloned in the pCR Blunt II-TOPO vector. Several clones were sequenced with M13 reverse oligonucleotide. The sequences, corresponding to 16S **(B)** or 23S **(C)**, were extracted and aligned to their reference sequence. The obtained 5’ ends are marked by a triangle. n indicates the number of independent sequences. a2 that results from a partial pairing of 16S-190R with the 16S rRNA at positions 289-298 has the same 5’ end as a1. The *T. barophilus* genome was recently re-annotated, and the 5’ ends of 16S and 23S rRNAs are modified: ° is in accordance to the most recent annotation (NC_014804.1); * corresponds to the former annotation (NC_014804), and can still be used in some databases.

For the 23S rRNA, we chose to perform the RACE experiments with the 23S-214R oligonucleotide. Again, the main product that is amplified is just above 400-bp (b2) while a faint band is at the expected size, around 249-nt (b1) (**Figure 5A**). Both bands were sequenced. The b2 band has a defined 5’ end that corresponds to position 2926 of the annotated 23S rDNA, which is located just at the end of helix H98 (2888-2926) (**Figure 5C**). The b1 smearing band has various 5’ ends located around the annotated one. The RACE experiments were done using total RNA extracted from cells grown in stationary phase. Accordingly, the various positions (Δ) are observed on the primer extension gel (**Figure 4B**). Interestingly, this is not as evident using total RNA extracted from cells in exponential phase (**Figure 4A**). This could correspond to degradation products that are more prevalent in stationary phase.

The sequenced 5’ end of the 23S rRNA that is located downstream of the 23S-214R indicates that a permutation event occurs. This also raises the question of the nature of 23S rRNA 3’ end. To answer this, we also performed 3’ RACE experiments on the 16S and 23S rRNAs, using oligonucleotides 263 (16S-263F) and 267 (23S-267F) nucleotides upstream of their respective annotated 3’ ends (**Figure 6**). Briefly, the 3’ end of the RNA molecules were ligated with an E1 adapter and reversed transcribed with an oligonucleotide specific for this adapter. The cDNA was then amplified using oligonucleotide pairs specific to the adapter and the 16S or 23S rRNA (16S-263F or 23S-267F). The bands of interest are then extracted from the agarose gel, cloned and sequenced. For the 16S rRNA, one main a1 band is obtained at the expected size of around 282-bp, the adapter adding 19-nt to the PCR product (**Figure 6A**). After the removal of the adapter, the four sequenced clones have the same 3’ end that corresponds to the annotated one (**Figure 6B**). For the 23S rRNA, one main b1 band is obtained around 144-bp that is way shorter than the expected size at around 286-bp (**Figure 6A**). The sequenced 3’ end is located at the position 2888, just upstream of helix H98, and 142-nt prior the annotated 3’ end of the 23S rDNA (**Figure 6C**). To not miss alternative products, we also extracted two faint bands (#, **Figure 6A**), but upon sequencing, they do not correspond to 23S molecules and are therefore unspecific products.

**Figure 6:**
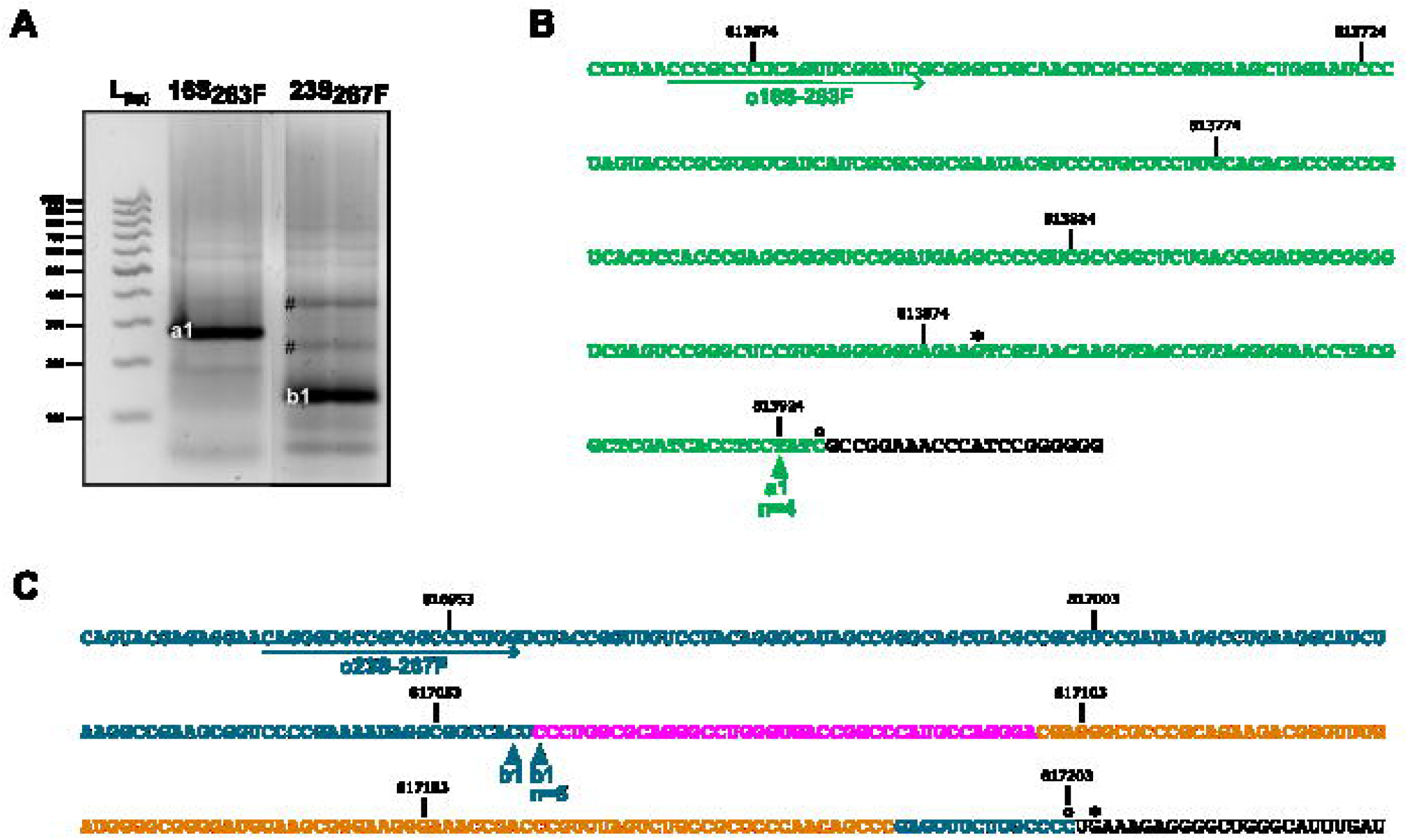
Precise mapping of the 16S and 23S 3’ ends by 3’ RACE. The cellular RNAs were dephosphorylated, ligated with an adapter phosphorylated at its 5’ end, and reverse transcribed with a primer specific for the adapter. The resulting cDNA was amplified by PCR with a primer specific for the 16S (o16S-263F) or the 23S (o23S-267F) rRNAs, and the primer specific for the 3’ end adapter. The amplification products were separated by electrophoresis on a 3% agarose gel **(A)**. The annotated bands were extracted and cloned in the pCR Blunt II-TOPO vector. Several clones were sequenced with M13 reverse primer. The sequences, corresponding to 16S **(B)** or 23S **(C)**, were extracted and aligned to their reference sequence. The obtained 3’ ends are marked by a triangle. n indicates the number of independent sequences. # corresponds to the amplification of unspecific products (6 clones were sequenced for each band). The *T. barophilus* genome was recently re-annotated, and the 3’ ends of 16S and 23S rRNAs are modified: ° is in accordance to the most recent annotation (NC_014804.1); * corresponds to the former annotation (NC_014804), and can still be used in some databases.

### The permutated 23S rRNA is incorporated into ribosome in *T. barophilus* cells

All the previous experiments were performed on total RNA. To determine if the permuted 23S rRNA is integrated in ribosomal particles, we did primer extension on cellular fractions that are enriched in ribosome particles. These fractions were obtained by size exclusion chromatography of very large intracellular structures (30) of whole-cell extracts of *T. barophilus* in stationary phase (**Figure 7A**). The presence of ribosomal particles in the fractions was determined by western blotting using antibodies specific to proteins of the small (S4e) and large (L10e) subunits (**Figure 7B**). In our conditions, we separate the 30S, 50S and 70S particles from low molecular weight particles but without defined peaks for each specie. In our conditions, no polysome fractions were observed. The RNAs from the fractions A12, B1-3 were extracted and used to perform primer extensions (**Figure 7C**). For both the 16S and 23S rRNAs, the profiles are similar to the one obtained in **Figure 4**. Therefore, the ribosomal particles, including translating 70S monosomes, contain conventional 16S rRNAs and permuted 23S rRNAs.

**Figure 7:**
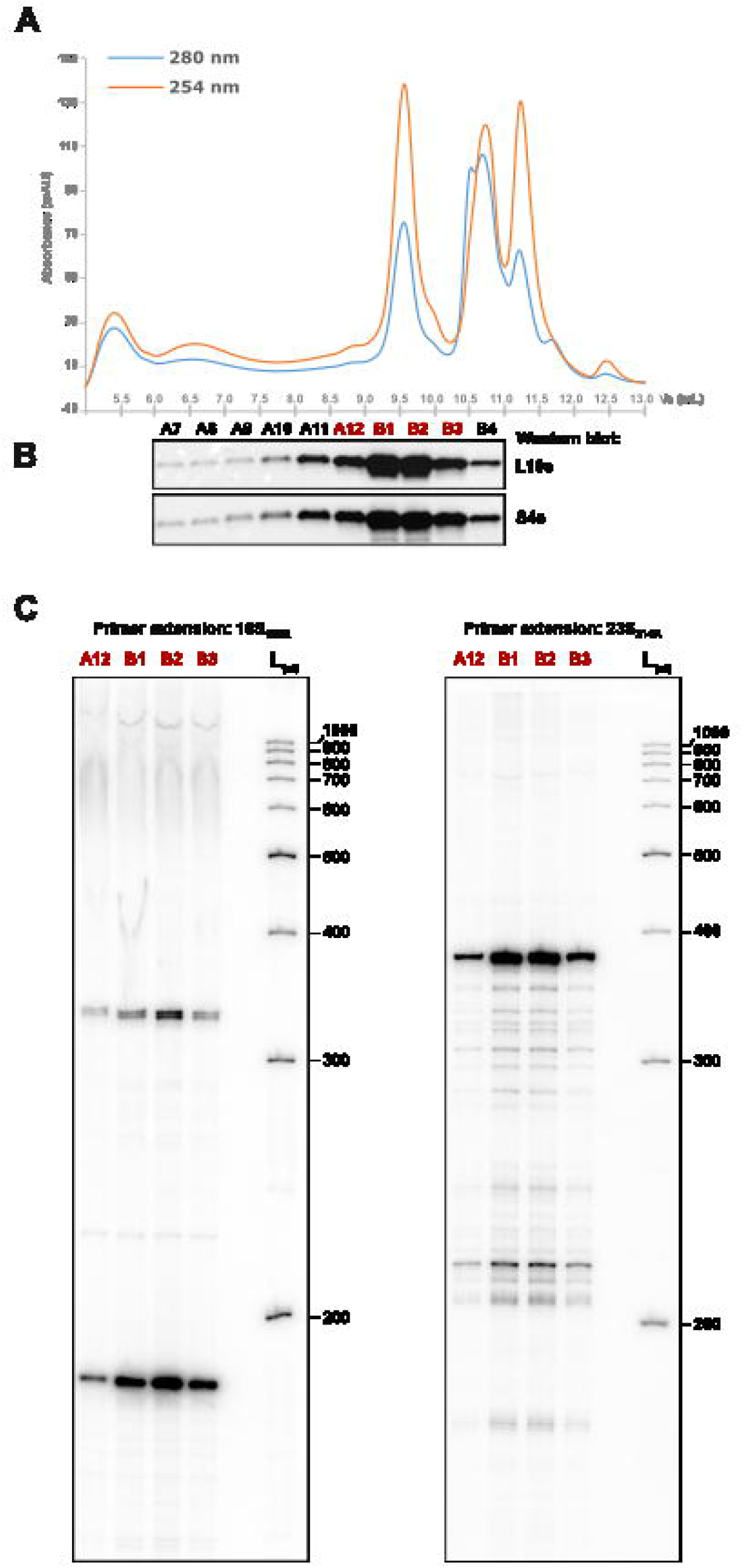
The permuted 23S rRNA is integrated in ribosome particles. **(A)** Clarified cellular extract from *T. barophilus* cells, at early stationary phase, were fractionated by Ribo Mega-SEC. The chromatogram profiles obtained by UV spectrophotometric detection at 254 nm and 280 nm are given. **(B)** Proteins from the identified fractions were precipitated and analyzed by Western blotting using antibodies specific for L10e and S4e, ribosomal proteins for the 50S and 30S particles, respectively. **(C)** The 5’ ends of the 16S and 23S rRNAs, assembled in ribosome particles, from fractions A12 and B1-3, were mapped by primer extension, as in Figure 4.

Altogether, our results indicate that the 23S permuted rRNA starts with helix H99 and ends with helix H97, helix H98 being deleted upon maturation (**Figure 8A**). It should also be noted that the permuted 23S contains an additional 44 nucleotides that corresponds to the BHB junction and that is not eliminated upon maturation. In order to give structural support to this model, we used the cryo-EM map of *Thermococcus kodakarensis* 50S (EMD-10223, PDB 6SKF). Consistent with our results, a permuted 23S rRNA can be modeled into the cryo-EM map. The resulting model is consistent with the absence of H98, with H99-101 located at the 5’ end and with the presence of the BHB junction, which fits into a previously unassigned part of the cryo-EM map (**Figure 8B**). While it is not clear whether or not the 23S rRNA permutation could affect, or not, the activity of the 70S ribosome, the BHB junction does not seem to form stable interactions with proteins. Also, depending on the organism, the length of this additional sequence varies, with 44-nt for *T. barophilus* and 22-nt for *T. kodakarensis*.

**Figure 8:**
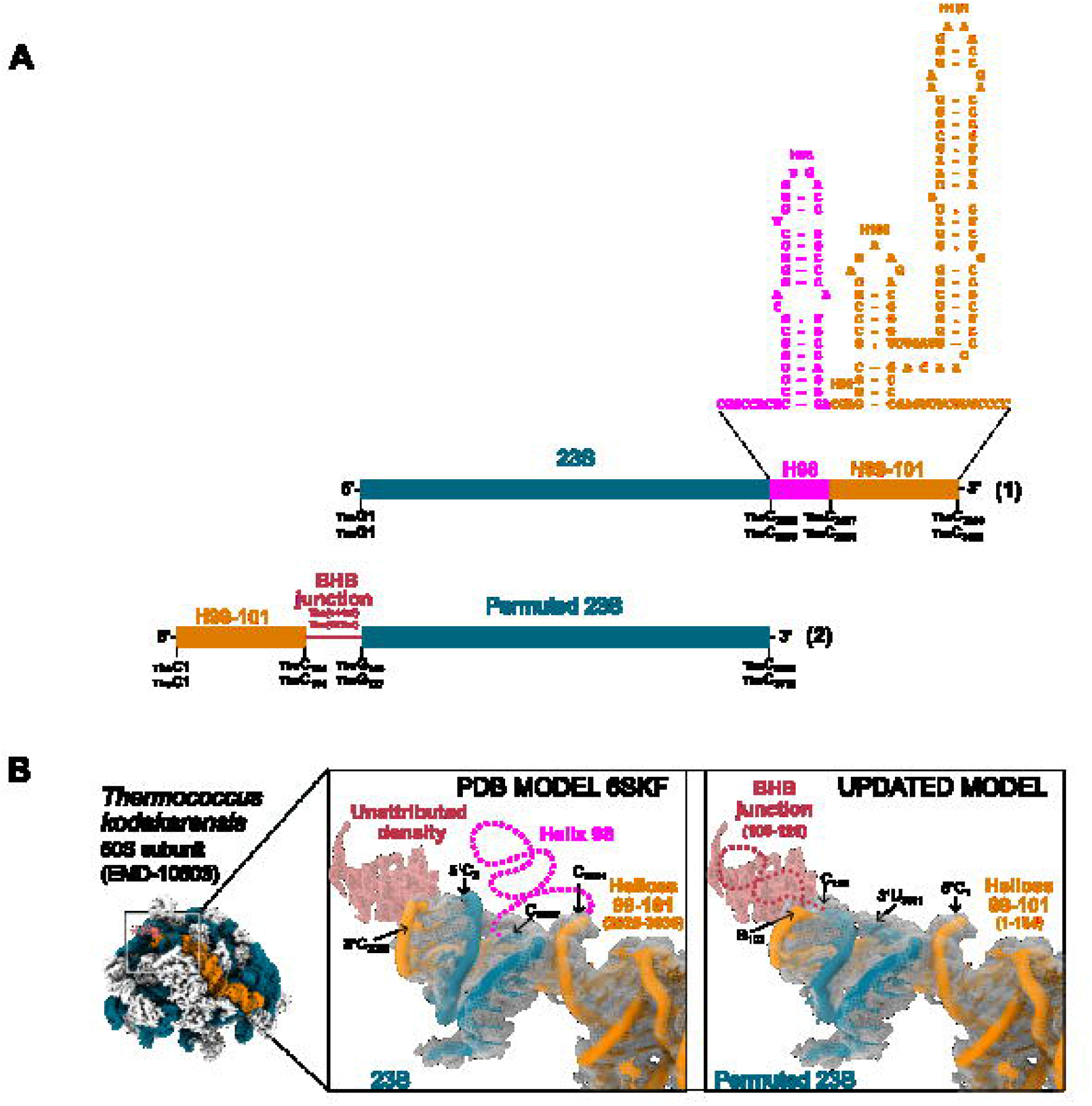
The 23S rRNA is circularly permuted, concomitantly with a deletion of helix H98. **(A)** The 5’ and 3’ ends of the 23S rRNAs inferred form mapping experiments. (1) represents the mature rRNA as encoded in the genome. (2) is the circularly permuted 23S rRNA that was generated concomitantly with the deletion of H98. The helices H99 to H101 (H99-101) are permuted to the 5’ end. The permuted rRNA also conserve the BHB junction that adds 44 nucleotides (nt) to the final molecule. The 3’ end is situated right on the edge of helix H98. In this simplified representation, the H1_3p_ is in orange, integrated within the helices H99-101. The sequence of H1_3p_ is written in italics. **(B)** Integration of the permuted 23S rRNA in the structure of the 50S ribosome subunit of *T. kodakarensis*. Helices H99-101(-H1_3p_) are shown in orange. Permuted 23S, lacking H98 and keeping the BHB junction (updated model, right panel), fits with unattributed density and explains the lack of density expected for H98 (left panel, PDN model 6SKF).

## DISCUSSION

We explored rRNA processing in *T. barophilus* by mining genome-wide transcriptomic data obtained on total RNAs of wild-type cells in exponential and stationary phases. We primarily focused on the recovery of circular junctions of the 16S and 23S rRDNAs that are the most abundant. We identified the canonical 16S and 23S BHB junctions, but also three alternative junctions for the 23S rRNA. We also recovered to a lesser extent the circularization junctions of the RNase P and SRP RNAs. Using molecular biology approaches, we further investigated the circular precursors and mature forms of the 16S and 23S rRNAs. While the mature 16S is as predicted in the Rfam database, the mature 23S is not. The 23S circ-pre-rRNA is opened within helix H98, by processing steps that remain to be characterized, to release a permuted 23S rRNA missing H98 and carrying an additional junction segment. The 50S subunit, alone or within the 70S, contain the permuted 23S rRNA. When a permuted 23S rRNA is modeled into a Cryo-EM structures of the 50S from *T. kodakarensis*, it allows to clarify the absence of density for H98 and to attribute a newly integrated RNA segment to previously unattributed density.

Circular RNAs are covalently closed RNA molecules that has been described in all domain of life. Advanced RNA sequencing analyses and computational studies, tailored towards detecting circRNA, have identified numerous circRNA classes. In archaea, one class stands out as it corresponds to circular rRNAs that are intermediated in the generation of the mature 16S and 23S rRNAs. The first processing step that takes place to generate the mature 16S and 23S rRNA involves the recognition, by endonuclease EndA, of a specific RNA structural motif named Bulge-Helix-Bulge (BHB) that is formed by sequences surrounding the rDNA genes. The generated precursors are then ligated by the RNA splicing ligase RtcB to generate 16S and 23S circular intermediates (circ-pre-rRNA). To this day, these intermediates were not identified in bacteria or eukaryotes. Other circRNA commonly found in archaea are the box C/D non-coding RNA, the RNA moieties of RNase P and of the signal recognition particle (SRP) (31).

While our total RNA deep sequencing analysis was not designed for circRNA identification, we were able to identify several classes of circular RNAs. To do so, we specifically extracted chimeric reads aligning to two genome segments using STAR. The circ-pre-rRNA class is the most abundant one and contains 16S and 23S circ-pre-rRNAs formed at canonical BHB motifs that are conserved in Thermococcales. Intriguingly, when investigating this difference in occurrence, even though they are expressed from an operon. Our results show that it is because the mature 23S rRNAs keep the circularization junction. This is due to a circular permutation that place the helices H99-101 at the 23S 5’ end. This is done by opening the 23S circ-pre-rRNAs at the extremities of helix H98. As a consequence, helix H98 is deleted in the process.

The deletion of helix H98 together with the circular permutation of the 23S rRNA was first observed in Pyrococcus furiosus using RiboMeth-Seq data (15). It was later confirmed using long-read nanopore sequencing data (4). Interestingly, this circular permutation does not seem to be conserved in *Haloferax volcanii* and in *Sulfolobus acidocaldarius* (4). In both organisms, the reopening of the 23S circ-pre-rRNA occurs in the circularization junction. Further maturation steps remove any sequence corresponding to the junction to leave the encoded 5’ and 3’ ends. Interestingly, helix H98 correspond to an expansion segment (ES) that give rise to ES39 in eukaryotic 28S rRNAs. While H98 is absent from most archaeal 23S rRNAs or restricted to few nucleotides, it is present in Thermococcales and Asgard archaea (**Figure 9**) (15,32). It remains to be seen if circular permutation is restricted to the Thermococcales and the presence of helix H98. If so, this could imply that the permutation is a way of removing H98 from the mature 23S rRNA as it does not fulfill a function there.

**Figure 9:**
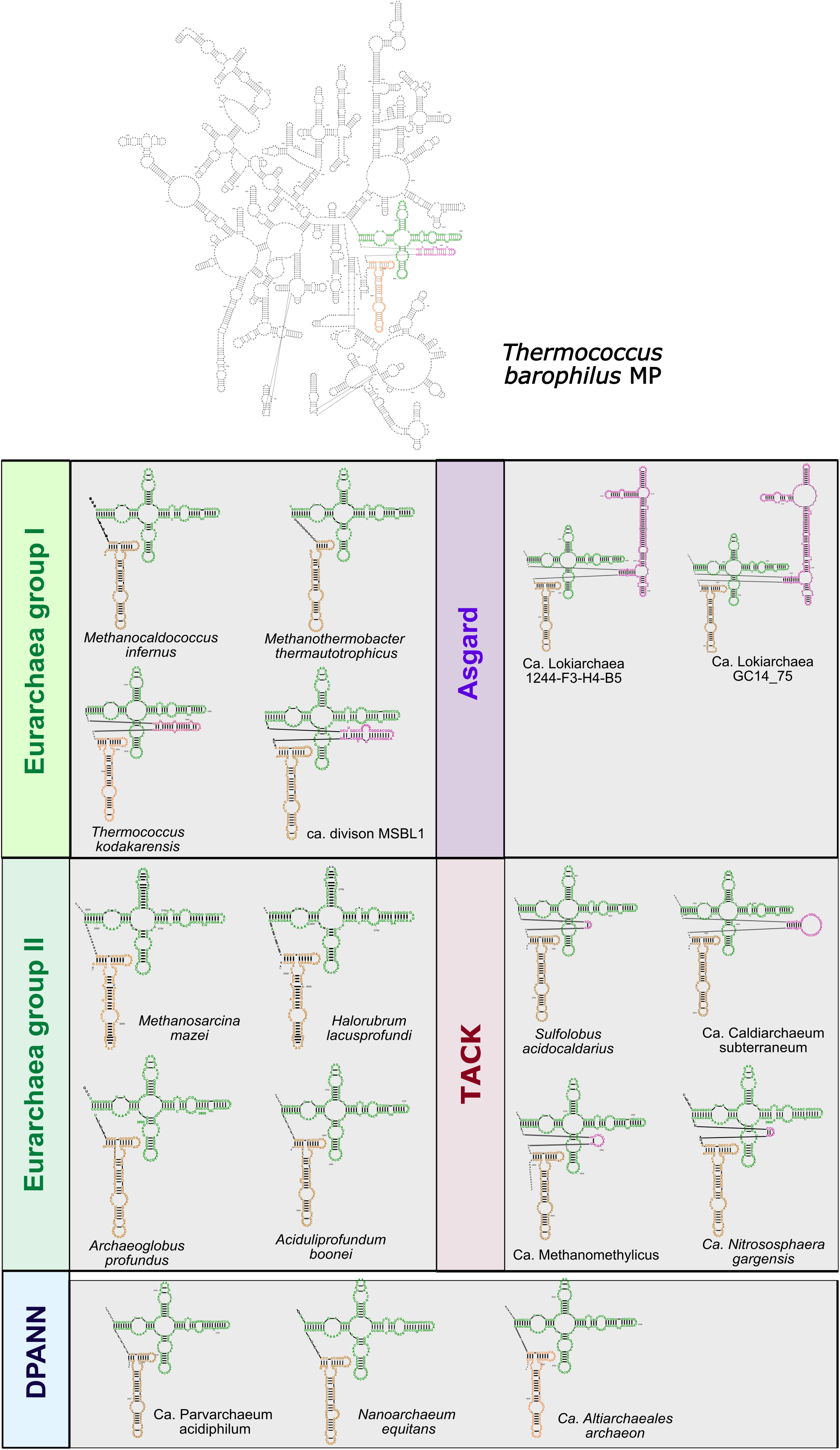
The deleted H98 helix is not conserved in archaeal genome. Representation of the 23S rRNA of *Thermococcus barophilus*, with helices H94-97, H98 and H99-101 represented in green, pink and orange, respectively. The sequences and structures of these helices were extracted from the 23S rRNA of representatives of different archaeal clades. H98 is absent in DPANN, TACK and Euryarchaea group I, and present in Euryarchaea group II and Asgard.

We also recovered few reads for other known circRNA such as some box C/D non-coding RNAs. However, we found out that while STAR can recover the long non-coding RNAs, it does not work that well with small ones (<150-nt).

While a 5S pre-circ-rRNA was identified as an intermediate of 5S in *Sulfolobus solfataricus* (10) and *Pyrococcus abyssi* (33), we did not find circularization junctions corresponding to the 5S rRNA of *T. barophilus*, a close relative of *P. abyssi*. Since they were found in low quantities, even using techniques that enrich circRNA, it is possible that our data set has not the depth needed. If 5S pre-circ-rRNA are confirmed, it would mean that circular RNA can be formed in absence of BHB motifs, as for the alternatives 23S circ-pre-rRNA.

We identified new 23S pre-circ-rRNA intermediates with alternative circularization junctions. They all occur at the same motif but have different cleavage sites: U/U, U/A, C/A. The new intermediates are abundant. The circularization junctions do not contain a canonical BHB motif. While it is not a BHB motif, it forms a structure. This structure is conserved in Thermococcales. If confirmed this means that circularization can occur by other mechanism. It would also mean that the maturation process of 23S rRNA is more complex than suspected and that several forms of 23S rRNA might exist in the cells. One advantages that could arise from these alternatives form is that the circularization junction that is kept during circular permutation is shorter, with only few nucleotides added in the sequence.

Recently, it was proposed that the 23S rRNA of *Methanosarcina acetivorans* remains circular in the 50S ribosomal particles (34). This is based on the analysis of sequencing of RNA isolated from *Methanosarcina acetivorans* ribosomes and the recovery of the circularization junction. As very little is known on the maturation of rRNAs and ribosome biogenesis in Archaea, it is possible that while 16S and 23S circularization seems to be conserved in many archaeal genomes, due to the presence of BHB motifs, the outcome of the maturation process is different depending on the rRNA nucleotide content and/or the maturation enzyme that are encoded (many of them being unknown). However, we would like to point out that in the *T. barophilus* and *P. furiosus* permuted 23 rRNA, the circularization junction is present in the mature 23S rRNA that is linear. Therefore, to be fully conclusive, the sole analysis of sequencing data needs to include samples treated with 3’-5’ exoribonuclease RNase R for confirming an enrichment of circular RNAs. A reliable and quick proof could also be determining the profiles of rRNAs by capillary migration, since circular RNA molecules will not separate based on their size.

## DATA AVAILIBILITY

The Transcriptomic data mass has been deposited via the GEO repository with the dataset identifier GSE229955 (https://www.ncbi.nlm.nih.gov/geo/query/acc.cgi?acc=GSE229955).

## LEGEND OF SUPPLEMENTARY FIGURES

**Supplementary Figure S1: An alignment of the sequences encoded upstream of the 16S rDNAs from Thermococcales genomes show that the H/ACA Tba19 is conserved**.

**(A)** Sequences alignment of the 200 nucleotides presents upstream of the 16S rDNA genes in 34 archaeal genomes belonging to the Thermococcales genus. The box H/ACA stems are boxed in grey. The nucleotides that are predicted to pair the 16S rRNA are in green. **(B)** 2D structure of box H/ACA Tba19, based on the predicted structure of Pab19 from *Pyrococcus abyssi* (26). The nucleotides involved in the pairing of 16S rRNA are circled in green. **(C)** Tba19 is complementary to the 16S rRNA (in green) and predicted to pseudouridilate U1017 (Ψ).

**Supplementary Figure S2a to S2e:** The chimeric reads mapping the 16S (2a) or 23S rRNA (2b-e) were extracted and aligned. One representative of each sequence with five occurrences or more was kept. The number N indicates the number of occurrences (e.g. 16SBHB_N).

**Supplementary Figure S3:** The linear by-products resulting from the excision of the 16S and 23S rRNAs (LB_rRNA) were shown to be ligated together (4,9,10). Few reads aligning on the LB_rRNA were found in our data set that support the existence of the LB_rRNA.

**Supplementary Figure S4:** The 5’ and 3’ ends of Thermococcales 23S rRNA were aligned. This shows that they are mostly conserved and form helix H1.

## Notes

### Competing Interest Statement

The authors have declared no competing interest.

### Funding

- Agence Nationale de la Recherche: ANR-22-CE12-0009

