## Supplementary Table S2 for "Permuted 23S rRNA is integrated in 50S ribosome particles in *Thermococcus barophilus*"

| **Name** | **Old Locus Tag** | **Locus Tag** | **KEGG**  NC_014804 | **UCSC Archaeal Genome Browser**  NC_014804 | **NCBI**  NC_014804.1  (2023_09_06) |
| --- | --- | --- | --- | --- | --- |
| **16S rRNA** | Termp_022095 | RS04670 | 812,466-813,878 | 812,466-813,878 | 812,425-813,927 |
| **23S rRNA** | Termp_022097 | RS04680 | 814,176-817,205 | 814,176-817,205 | 814,174-817,203 |

**Supplementary Table S2:** **Coordinates of 16S and 23S rDNAs in different databases**

The genome sequence of *Thermococcus barophilus* is referenced as CP002372.1 [(Vannier et al. 2011)](https://sciwheel.com/work/citation?ids=3994700&pre=&suf=&sa=0&dbf=0). Recently, the NCBI database integrates a genome that has been re-annotated (<https://www.ncbi.nlm.nih.gov/nuccore/NC_014804.1/>). On publication of this article, the coordinates of the 16S and 23S rDNAs in the databases KEGG (<https://www.genome.jp/entry/gn:T01377>) and UCSC Archaeal Genome Browser (<http://archaea.ucsc.edu/>) originate from the previous annotation (NC_014804).
