## Supplementary Table S3 for "Permuted 23S rRNA is integrated in 50S ribosome particles in *Thermococcus barophilus*"

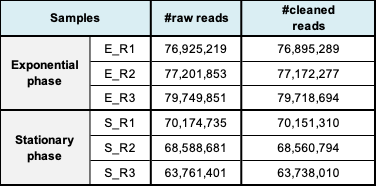

**Supplementary Table S3: Number of reads obtained for each dataset**

RNA extracted from wild-type cells grew in exponential (E) and stationary (S) phases was subjected to deep sequencing (paired-end, three replicates (R) for each condition). The RNA samples included ribosomal RNA. Over 60 million of paired-end reads were obtained for each dataset.
