## Supplementary Table S4 for "Permuted 23S rRNA is integrated in 50S ribosome particles in *Thermococcus barophilus*"

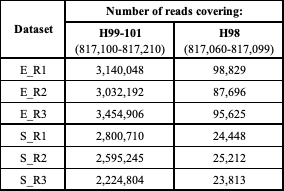

**Supplementary Table S4:** Number of reads covering the helices H98 and H99-101 from the 23S rRNA. The H98 reads correspond to precursors of the 23S rRNA, and among them to circular precursors.
