## Supplementary Figure S2a for "Permuted 23S rRNA is integrated in 50S ribosome particles in *Thermococcus barophilus*"

```
1                                                                                                                443
16SBHB_162 CCT ACGGCTCGAT CACCTCCTAT CGCCGGAAAC CCATCCGGGG GGTTTAACCC CGGATGGGAT TTTGGTACTC CCCTTTATGG GGAGTCCCAC CGGCCGTACT CCTATAGCAA TTCCGGTTGA TCCTGCCGGA GGCCACTGCT ATGGGGGTCC GACTAAG
16SBHB_52_ CCT ACGGCTCGAT CACCTCCTAT CGCCGGAAAC CCATCCGGGG GGTTTAACCC CGGATGGGAT TTTGGTACTC CCCTTTATGG GGAGTCCCAC CGGCCGTACT CCTATAGCAA TTCCGGTTGA TCCTGCCGGA GGCCACTGCT ATGGGGGTCC GACTAAGC
16SBHB_30_ CCT ACGGCTCGAT CACCTCCTAT CGCCGGAAAC CCATCCGGGG GGTTTAACCC CGGATGGGAT TTTGGTACTC CCCTTTATGG GGAGTCCCAC CGGCCGTACT CCTATAGCAA TTCCGGTTGA TCCTGCCGGA GGCCACTGCT ATGGGGGTCC GACTAAGCC
16SBHB_12_ CCT ACGGCTCGAT CACCTCCTAT CGCCGGAAAC CCATCCGGGG GGTTTAACCC CGGATGGGAT TTTGGTACTC CCCTTTATGG GGAGTCCCAC CGGCCGTACT CCTATAGCAA TTCCGGTTGA TCCTGCCGGA GGCCACTGCT ATGGGGGTCC GACTAAGCCG
16SBHB_3_9 CT ACGGCTCGAT CACCTCCTAT CGCCGGAAAC CCATCCGGGG GGTTTAACCC CGGATGGGAT TTTGGTACTC CCCTTTATGG GGAGTCCCAC CGGCCGTACT CCTATAGCAA TTCCGGTTGA TCCTGCCGGA GGCCACTGCT ATGGGGGTCC GACTAAGCCG
16SBHB_8_2 CT ACGGCTCGAT CACCTCCTAT CGCCGGAAAC CCATCCGGGG GGTTTAACCC CGGATGGGAT TTTGGTACTC CCCTTTATGG GGAGTCCCAC CGGCCGTACT CCTATAGCAA TTCCGGTTGA TCCTGCCGGA GGCCACTGCT ATGGGGGTCC GACTAAGCC
16SBHB_11_ CT ACGGCTCGAT CACCTCCTAT CGCCGGAAAC CCATCCGGGG GGTTTAACCC CGGATGGGAT TTTGGTACTC CCCTTTATGG GGAGTCCCAC CGGCCGTACT CCTATAGCAA TTCCGGTTGA TCCTGCCGGA GGCCACTGCT ATGGGGGTCC GACTAAGCCA
16SBHB_109 CGT AGGGGAACCT ACGGCTCGAT CACCTCCTAT CGCCGGAAAC CCATCCGGGG GGTTTAACCC CGGATGGGAT TTTGGTACTC CCCTTTATGG GGAGTCCCAC CGGCCGTACT CCTATAGCAA TTCCGGTTGA TCCTGCCGGA GGCCACTGCT ATGGGGGTCC GACTAAG
16SBHB_50_ CGT AGGGGAACCT ACGGCTCGAT CACCTCCTAT CGCCGGAAAC CCATCCGGGG GGTTTAACCC CGGATGGGAT TTTGGTACTC CCCTTTATGG GGAGTCCCAC CGGCCGTACT CCTATAGCAA TTCCGGTTGA TCCTGCCGGA GGCCACTGCT ATGGGGGTCC GACTAAGC
16SBHB_30_ CGT AGGGGAACCT ACGGCTCGAT CACCTCCTAT CGCCGGAAAC CCATCCGGGG GGTTTAACCC CGGATGGGAT TTTGGTACTC CCCTTTATGG GGAGTCCCAC CGGCCGTACT CCTATAGCAA TTCCGGTTGA TCCTGCCGGA GGCCACTGCT ATGGGGGTCC GACTAAGCC
16SBHB_4_6 CGT AGGGGAACCT ACGGCTCGAT CACCTCCTAT CGCCGGAAAC CCATCCGGGG GGTTTAACCC CGGATGGGAT TTTGGTACTC CCCTTTATGG GGAGTCCCAC CGGCCGTACT CCTATAGCAA TTCCGGTTGA TCCTGCCGGA GGCCACTGCT ATGGGGGTCC GACTAAGCCC G
16SBHB_13_ CGT AGGGGAACCT ACGGCTCGAT CACCTCCTAT CGCCGGAAAC CCATCCGGGG GGTTTAACCC CGGATGGGAT TTTGGTACTC CCCTTTATGG GGAGTCCCAC CGGCCGTACT CCTATAGCAA TTCCGGTTGA TCCTGCCGGA GGCCACTGCT ATGGGGGTCC GACTAAGCCA
16SBHB_10_ GGTAGCCGT AGGGGAACCT ACGGCTCGAT CACCTCCTAT CGCCGGAAAC CCATCCGGGG GGTTTAACCC CGGATGGGAT TTTGGTACTC CCCTTTATGG GGAGTCCCAC CGGCCGTACT CCTATAGCAA TTCCGGTTGA TCCTGCCGGA GGCCACTGCT ATGGGGGTCC GACTAAG
16SBHB_7_3 GGTAGCCGT AGGGGAACCT ACGGCTCGAT CACCTCCTAT CGCCGGAAAC CCATCCGGGG GGTTTAACCC CGGATGGGAT TTTGGTACTC CCCTTTATGG GGAGTCCCAC CGGCCGTACT CCTATAGCAA TTCCGGTTGA TCCTGCCGGA GGCCACTGCT ATGGGGGTCC GACTAAGC
16SBHB_7_3 GTAGCCGT AGGGGAACCT ACGGCTCGAT CACCTCCTAT CGCCGGAAAC CCATCCGGGG GGTTTAACCC CGGATGGGAT TTTGGTACTC CCCTTTATGG GGAGTCCCAC CGGCCGTACT CCTATAGCAA TTCCGGTTGA TCCTGCCGGA GGCCACTGCT ATGGGGGTCC GACTAAGCC
16SBHB_3_7 TAGCCGT AGGGGAACCT ACGGCTCGAT CACCTCCTAT CGCCGGAAAC CCATCCGGGG GGTTTAACCC CGGATGGGAT TTTGGTACTC CCCTTTATGG GGAGTCCCAC CGGCCGTACT CCTATAGCAA TTCCGGTTGA TCCTGCCGGA GGCCACTGCT ATGGGGGTCC GACTAAGCCA
16SBHB_8_2 GTAGCCGT AGGGGAACCT ACGGCTCGAT CACCTCCTAT CGCCGGAAAC CCATCCGGGG GGTTTAACCC CGGATGGGAT TTTGGTACTC CCCTTTATGG GGAGTCCCAC CGGCCGTACT CCTATAGCAA TTCCGGTTGA TCCTGCCGGA GGCCACTGCT ATGGGGGTCC GACTAAG
16SBHB_6_4 CCGT AGGGGAACCT ACGGCTCGAT CACCTCCTAT CGCCGGAAAC CCATCCGGGG GGTTTAACCC CGGATGGGAT TTTGGTACTC CCCTTTATGG GGAGTCCCAC CGGCCGTACT CCTATAGCAA TTCCGGTTGA TCCTGCCGGA GGCCACTGCT ATGGGGGTCC GACTAAGCC
16SBHB_9_2 CCGT AGGGGAACCT ACGGCTCGAT CACCTCCTAT CGCCGGAAAC CCATCCGGGG GGTTTAACCC CGGATGGGAT TTTGGTACTC CCCTTTATGG GGAGTCCCAC CGGCCGTACT CCTATAGCAA TTCCGGTTGA TCCTGCCGGA GGCCACTGCT ATGGGGGTCC GACTAAGC
16SBHB_4_5 GT AGGGGAACCT ACGGCTCGAT CACCTCCTAT CGCCGGAAAC CCATCCGGGG GGTTTAACCC CGGATGGGAT TTTGGTACTC CCCTTTATGG GGAGTCCCAC CGGCCGTACT CCTATAGCAA TTCCGGTTGA TCCTGCCGGA GGCCACTGCT ATGGGGGTCC GACTAAGCCG
16SBHB_3_8 GT AGGGGAACCT ACGGCTCGAT CACCTCCTAT CGCCGGAAAC CCATCCGGGG GGTTTAACCC CGGATGGGAT TTTGGTACTC CCCTTTATGG GGAGTCCCAC CGGCCGTACT CCTATAGCAA TTCCGGTTGA TCCTGCCGGA GGCCACTGCT ATGGGGGTCC GACTAAGCC
16SBHB_3_8 GT AGGGGAACCT ACGGCTCGAT CACCTCCTAT CGCCGGAAAC CCATCCGGGG GGTTTAACCC CGGATGGGAT TTTGGTACTC CCCTTTATGG GGAGTCCCAC CGGCCGTACT CCTATAGCAA TTCCGGTTGA TCCTGCCGGA GGCCACTGCT ATGGGGGTCC GACTAAGCCA
16SBHB_7_3 CCGT AGGGGAACCT ACGGCTCGAT CACCTCCTAT CGCCGGAAAC CCATCCGGGG GGTTTAACCC CGGATGGGAT TTTGGTACTC CCCTTTATGG GGAGTCCCAC CGGCCGTACT CCTATAGCAA TTCCGGTTGA TCCTGCCGGA GGCCACTGCT ATGGGGGTCC GACTAAG
16SBHB_5_4 GT AGGGGAACCT ACGGCTCGAT CACCTCCTAT CGCCGGAAAC CCATCCGGGG GGTTTAACCC CGGATGGGAT TTTGGTACTC CCCTTTATGG GGAGTCCCAC CGGCCGTACT CCTATAGCAA TTCCGGTTGA TCCTGCCGGA GGCCACTGCT ATGGGGGTCC GACTAAGC
16SBHB_7_3 AGGGGAACCT ACGGCTCGAT CACCTCCTAT CGCCGGAAAC CCATCCGGGG GGTTTAACCC CGGATGGGAT TTTGGTACTC CCCTTTATGG GGAGTCCCAC CGGCCGTACT CCTATAGCAA TTCCGGTTGA TCCTGCCGGA GGCCACTGCT ATGGGGGTCC GACTAAGCC
16SBHB_5_5 AGGGGAACCT ACGGCTCGAT CACCTCCTAT CGCCGGAAAC CCATCCGGGG GGTTTAACCC CGGATGGGAT TTTGGTACTC CCCTTTATGG GGAGTCCCAC CGGCCGTACT CCTATAGCAA TTCCGGTTGA TCCTGCCGGA GGCCACTGCT ATGGGGGTCC GACTAAGCCA G
16SBHB_3_9 CGT AGGGGAACCT ACGGCTCGAT CACCTCCTAT CGCCGGAAAC CCATCCGGGG GGTTTAACCC CGGATGGGAT TTTGGTACTC CCCTTTATGG GGAGTCCCAC CGGCCGTACT CCTATAGCAA TTCCGGTTGA TCCTGCCGGA GGCCACTGCT ATGGGGGTCC GACCAAG
16SBHB_7_3 GT AGGGGAACCT ACGGCTCGAT CACCTCCTAT CGCCGGAAAC CCATCCGGGG GGTTTAACCC CGGATGGGAT TTTGGTACTC CCCTTTATGG GGAGTCCCAC CGGCCGTACT CCTATAGCAA TTCCGGTTGA TCCTGCCGGA GGCCACTGCT ATGGGGGTCC GACTAAG
16SBHB_59_ CCT ACGGCTCGAT CACCTCCTAT CGCCGGAAAC CCATCCGGGG GGTTTAACCC CGGATGGGAT TTTGGTACTC CCCTTTATGG GGAGTCCCAC CGGCCGTACT CCTATAGCAA TTCCGGTTGA TCCTGCCGGA GGCCACTGCT ATGGGGGTCC GACTAAGCCA TGCGAGTCAT GGGGGTGTCC CTTCTGGGAC ACCACCGGCG GACGGCTCAG TAACACGTCG
16SBHB_45_ CCT ACGGCTCGAT CACCTCCTAT CGCCGGAAAC CCATCCGGGG GGTTTAACCC CGGATGGGAT TTTGGTACTC CCCTTTATGG GGAGTCCCAC CGGCCGTACT CCTATAGCAA TTCCGGTTGA TCCTGCCGGA GGCCACTGCT ATGGGGGTCC GACTAAGCCA TGCGAGTCAT GGGGGTGTCC CTTCTGGGAC ACCACCGGCG GACGGCTCAG TAACACGTCG G
16SBHB_9_2 CT ACGGCTCGAT CACCTCCTAT CGCCGGAAAC CCATCCGGGG GGTTTAACCC CGGATGGGAT TTTGGTACTC CCCTTTATGG GGAGTCCCAC CGGCCGTACT CCTATAGCAA TTCCGGTTGA TCCTGCCGGA GGCCACTGCT ATGGGGGTCC GACTAAGCCA TGCGAGTCAT GGGGGTGTCC CTTCTGGGAC ACCACCGGCG GACGGCTCAG TAACACGTCG G
16SBHB_4_6 CT ACGGCTCGAT CACCTCCTAT CGCCGGAAAC CCATCCGGGG GGTTTAACCC CGGATGGGAT TTTGGTACTC CCCTTTATGG GGAGTCCCAC CGGCCGTACT CCTATAGCAA TTCCGGTTGA TCCTGCCGGA GGCCACTGCT ATGGGGGTCC GACTAAGCCA TGCGAGTCAT GGGGGTGTCC CTTCTGGGAC ACCACCGGCG GACGGCTCAG TAACACGTCG GTG
16SBHB_20_ CT ACGGCTCGAT CACCTCCTAT CGCCGGAAAC CCATCCGGGG GGTTTAACCC CGGATGGGAT TTTGGTACTC CCCTTTATGG GGAGTCCCAC CGGCCGTACT CCTATAGCAA TTCCGGTTGA TCCTGCCGGA GGCCACTGCT ATGGGGGTCC GACTAAGCCA TGCGAGTCAT GGGGGTGTCC CTTCTGGGAC ACCACCGGCG GACGGCTCAG TAACACGTCG
16SBHB_3_7 T ACGGCTCGAT CACCTCCTAT CGCCGGAAAC CCATCCGGGG GGTTTAACCC CGGATGGGAT TTTGGTACTC CCCTTTATGG GGAGTCCCAC CGGCCGTACT CCTATAGCAA TTCCGGTTGA TCCTGCCGGA GGCCACTGCT ATGGGGGTCC GACTAAGCCA TGCGAGTCAT GGGGGTGTCC CTTCTGGGAC ACCACCGGCG GACGGCTCAG TAACACGTCG G
16SBHB_6_4 CCT ACGGCTCGAT CACCTCCTAT CGCCGGAAAC CCATCCGGGG GGTTTAACCC CGGATGGGAT TTTGGTACTC CCCTTTATGG GGAGTCCCAC CGGCCGTACT CCTATAGCAA TTCCGGTTGA TCCTGCCGGA GGCCACTGCT ATGGGGGTCC GACTAAGCCA TGCGAGTCAT GGGGGTGTCC CTTCTGGGAC ACCACCGGCG GACGGCTCAG TAACACGTC
16SBHB_7_3 T ACGGCTCGAT CACCTCCTAT CGCCGGAAAC CCATCCGGGG GGTTTAACCC CGGATGGGAT TTTGGTACTC CCCTTTATGG GGAGTCCCAC CGGCCGTACT CCTATAGCAA TTCCGGTTGA TCCTGCCGGA GGCCACTGCT ATGGGGGTCC GACTAAGCCA TGCGAGTCAT GGGGGTGTCC CTTCTGGGAC ACCACCGGCG GACGGCTCAG TAACACGTCG
16SBHB_46_ CGT AGGGGAACCT ACGGCTCGAT CACCTCCTAT CGCCGGAAAC CCATCCGGGG GGTTTAACCC CGGATGGGAT TTTGGTACTC CCCTTTATGG GGAGTCCCAC CGGCCGTACT CCTATAGCAA TTCCGGTTGA TCCTGCCGGA GGCCACTGCT ATGGGGGTCC GACTAAGCCA TGCGAGTCAT GGGGGTGTCC CTTCTGGGAC ACCACCGGCG GACGGCTCAG TAACACGTCG
16SBHB_19_ CGT AGGGGAACCT ACGGCTCGAT CACCTCCTAT CGCCGGAAAC CCATCCGGGG GGTTTAACCC CGGATGGGAT TTTGGTACTC CCCTTTATGG GGAGTCCCAC CGGCCGTACT CCTATAGCAA TTCCGGTTGA TCCTGCCGGA GGCCACTGCT ATGGGGGTCC GACTAAGCCA TGCGAGTCAT GGGGGTGTCC CTTCTGGGAC ACCACCGGCG GACGGCTCAG TAACACGTCG G
16SBHB_3_9 CGT AGGGGAACCT ACGGCTCGAT CACCTCCTAT CGCCGGAAAC CCATCCGGGG GGTTTAACCC CGGATGGGAT TTTGGTACTC CCCTTTATGG GGAGTCCCAC CGGCCGTACT CCTATAGCAA TTCCGGTTGA TCCTGCCGGA GGCCACTGCT ATGGGGGTCC GACTAAGCCA TGCGAGTCAT GGGGGTGTCC CTTCTGGGAC ACCACCGGCG GACGGCTCAG TAACACGTCG GT
16SBHB_5_5 CCGT AGGGGAACCT ACGGCTCGAT CACCTCCTAT CGCCGGAAAC CCATCCGGGG GGTTTAACCC CGGATGGGAT TTTGGTACTC CCCTTTATGG GGAGTCCCAC CGGCCGTACT CCTATAGCAA TTCCGGTTGA TCCTGCCGGA GGCCACTGCT ATGGGGGTCC GACTAAGCCA TGCGAGTCAT GGGGGTGTCC CTTCTGGGAC ACCACCGGCG GACGGCTCAG TAACACGTCG G
16SBHB_3_9 CGT AGGGGAACCT ACGGCTCGAT CACCTCCTAT CGCCGGAAAC CCATCCGGGG GGTTTAACCC CGGATGGGAT TTTGGTACTC CCCTTTATGG GGAGTCCCAC CGGCCGTACT CCTATAGCAA TTCCGGTTGA TCCTGCCGGA GGCCACTGCT ATGGGGGTCC GACTAAGCCA TGCGAGTCAT GGGGGTGTCC CTTCTGGGAC ACCACCGGCG GACGGCTCAG TAACACGTC
16SBHB_3_8 GT AGGGGAACCT ACGGCTCGAT CACCTCCTAT CGCCGGAAAC CCATCCGGGG GGTTTAACCC CGGATGGGAT TTTGGTACTC CCCTTTATGG GGAGTCCCAC CGGCCGTACT CCTATAGCAA TTCCGGTTGA TCCTGCCGGA GGCCACTGCT ATGGGGGTCC GACTAAGCCA TGCGAGTCAT GGGGGTGTCC CTTCTGGGAC ACCACCGGCG GACGGCTCAG TAACACGTCG
16SBHB_27_ GGGGAACCT ACGGCTCGAT CACCTCCTAT CGCCGGAAAC CCATCCGGGG GGTTTAACCC CGGATGGGAT TTTGGTACTC CCCTTTATGG GGAGTCCCAC CGGCCGTACT CCTATAGCAA TTCCGGTTGA TCCTGCCGGA GGCCACTGCT ATGGGGGTCC GACTAAGCCA TGCGAGTCAT GGGGGTGTCC CTTCTGGGAC ACCACCGGCG GACGGCTCAG TAACACGTCG G
16SBHB_22_ AGGGGAACCT ACGGCTCGAT CACCTCCTAT CGCCGGAAAC CCATCCGGGG GGTTTAACCC CGGATGGGAT TTTGGTACTC CCCTTTATGG GGAGTCCCAC CGGCCGTACT CCTATAGCAA TTCCGGTTGA TCCTGCCGGA GGCCACTGCT ATGGGGGTCC GACTAAGCCA TGCGAGTCAT GGGGGTGTCC CTTCTGGGAC ACCACCGGCG GACGGCTCAG TAACACGTCG
16SBHB_3_8 GGGGAACCT ACGGCTCGAT CACCTCCTAT CGCCGGAAAC CCATCCGGGG GGTTTAACCC CGGATGGGAT TTTGGTACTC CCCTTTATGG GGAGTCCCAC CGGCCGTACT CCTATAGCAA TTCCGGTTGA TCCTGCCGGA GGCCACTGCT ATGGGGGTCC GACTAAGCCA TGCGAGTCAT GGGGGTGTCC CTTCTGGGAC ACCACCGGCG GACGGCTCAG TAACACGTCG GT
16SBHB_23_ GGGGAACCT ACGGCTCGAT CACCTCCTAT CGCCGGAAAC CCATCCGGGG GGTTTAACCC CGGATGGGAT TTTGGTACTC CCCTTTATGG GGAGTCCCAC CGGCCGTACT CCTATAGCAA TTCCGGTTGA TCCTGCCGGA GGCCACTGCT ATGGGGGTCC GACTAAGCCA TGCGAGTCAT GGGGGTGTCC CTTCTGGGAC ACCACCGGCG GACGGCTCAG TAACACGTCG
16SBHB_3_9 CGT AGGGGAACCT ACGGCTCGAT CACCTCCTAT CGCCGGAAAC CCATCCGGGG GGTTTAACCC CGGATGGGAT TTTGGTACTC CCCTTTATGG GGAGTCCCAC CGGCCGTACT CCTATAGCAA TTCCGGTTGA TCCTGCCGGA GGCCACTGCT ATGGGGGTCC GACTAAGCCA TGCGAGTCAT GGGGGTGTCC CTTCTGGGAC ACCACCGGCG GACGGCTCAG TAACACG
16SBHB_5_4 GGGAACCT ACGGCTCGAT CACCTCCTAT CGCCGGAAAC CCATCCGGGG GGTTTAACCC CGGATGGGAT TTTGGTACTC CCCTTTATGG GGAGTCCCAC CGGCCGTACT CCTATAGCAA TTCCGGTTGA TCCTGCCGGA GGCCACTGCT ATGGGGGTCC GACTAAGCCA TGCGAGTCAT GGGGGTGTCC CTTCTGGGAC ACCACCGGCG GACGGCTCAG TAACACGTCG G
16SBHB_25_ CCT ACGGCTCGAT CACCTCCTAT CGCCGGAAAC CCATCCGGGG GGTTTAACCC CGGATGGGAT TTTGGTACTC CCCTTTATGG GGAGTCCCAC CGGCCGTACT CCTATAGCAA TTCCGGTTGA TCCTGCCGGA GGCCACTGCT ATGGGGGTCC GACTAAGCCA TGCGAGTCAT GGGGGTGTCC CTTCTGGGAC ACCACCGGCG GACGGCTCAG TAACACGTCG GTAACCTACC CTCGGG
16SBHB_7_3 CCT ACGGCTCGAT CACCTCCTAT CGCCGGAAAC CCATCCGGGG GGTTTAACCC CGGATGGGAT TTTGGTACTC CCCTTTATGG GGAGTCCCAC CGGCCGTACT CCTATAGCAA TTCCGGTTGA TCCTGCCGGA GGCCACTGCT ATGGGGGTCC GACTAAGCCA TGCGAGTCAT GGGGGTGTCC CTTCTGGGAC ACCACCGGCG GACGGCTCAG TAACACGTCG GTAACCTACC CTCGGGAG
16SBHB_20_ GGGGAACCT ACGGCTCGAT CACCTCCTAT CGCCGGAAAC CCATCCGGGG GGTTTAACCC CGGATGGGAT TTTGGTACTC CCCTTTATGG GGAGTCCCAC CGGCCGTACT CCTATAGCAA TTCCGGTTGA TCCTGCCGGA GGCCACTGCT ATGGGGGTCC GACTAAGCCA TGCGAGTCAT GGGGGTGTCC CTTCTGGGAC ACCACCGGCG GACGGCTCAG TAACACGTCG GTAACCTACC CTCGGG
16SBHB_19_ CGT AGGGGAACCT ACGGCTCGAT CACCTCCTAT CGCCGGAAAC CCATCCGGGG GGTTTAACCC CGGATGGGAT TTTGGTACTC CCCTTTATGG GGAGTCCCAC CGGCCGTACT CCTATAGCAA TTCCGGTTGA TCCTGCCGGA GGCCACTGCT ATGGGGGTCC GACTAAGCCA TGCGAGTCAT GGGGGTGTCC CTTCTGGGAC ACCACCGGCG GACGGCTCAG TAACACGTCG GTAACCTACC CTCGGG
16SBHB_4_6 CGT AGGGGAACCT ACGGCTCGAT CACCTCCTAT CGCCGGAAAC CCATCCGGGG GGTTTAACCC CGGATGGGAT TTTGGTACTC CCCTTTATGG GGAGTCCCAC CGGCCGTACT CCTATAGCAA TTCCGGTTGA TCCTGCCGGA GGCCACTGCT ATGGGGGTCC GACTAAGCCA TGCGAGTCAT GGGGGTGTCC CTTCTGGGAC ACCACCGGCG GACGGCTCAG TAACACGTCG GTAACCTACC CTCGG
16SBHB_4_6 CGT AGGGGAACCT ACGGCTCGAT CACCTCCTAT CGCCGGAAAC CCATCCGGGG GGTTTAACCC CGGATGGGAT TTTGGTACTC CCCTTTATGG GGAGTCCCAC CGGCCGTACT CCTATAGCAA TTCCGGTTGA TCCTGCCGGA GGCCACTGCT ATGGGGGTCC GACTAAGCCA TGCGAGTCAT GGGGGTGTCC CTTCTGGGAC ACCACCGGCG GACGGCTCAG TAACACGTCG GTAACCTACC CCCGGG
16SBHB_6_3 GGGAACCT ACGGCTCGAT CACCTCCTAT CGCCGGAAAC CCATCCGGGG GGTTTAACCC CGGATGGGAT TTTGGTACTC CCCTTTATGG GGAGTCCCAC CGGCCGTACT CCTATAGCAA TTCCGGTTGA TCCTGCCGGA GGCCACTGCT ATGGGGGTCC GACTAAGCCA TGCGAGTCAT GGGGGTGTCC CTTCTGGGAC ACCACCGGCG GACGGCTCAG TAACACGTCG GTAACCTACC CTCGGG
16SBHB_14_ GGAACCT ACGGCTCGAT CACCTCCTAT CGCCGGAAAC CCATCCGGGG GGTTTAACCC CGGATGGGAT TTTGGTACTC CCCTTTATGG GGAGTCCCAC CGGCCGTACT CCTATAGCAA TTCCGGTTGA TCCTGCCGGA GGCCACTGCT ATGGGGGTCC GACTAAGCCA TGCGAGTCAT GGGGGTGTCC CTTCTGGGAC ACCACCGGCG GACGGCTCAG TAACACGTCG GTAACCTACC CTCGGG
16SBHB_12_ CGT AGGGGAACCT ACGGCTCGAT CACCTCCTAT CGCCGGAAAC CCATCCGGGG GGTTTAACCC CGGATGGGAT TTTGGTACTC CCCTTTATGG GGAGTCCCAC CGGCCGTACT CCTATAGCAA TTCCGGTTGA TCCTGCCGGA GGCCACTGCT ATGGGGGTCC GACTAAGCCA TGCGAGTCAT GGGGGTGTCC CTTCTGGGAC ACCACCGGCG GACGGCTCAG TAACACGTCG GTAACCTACC C
16SBHB_3_9 CGT AGGGGAACCT ACGGCTCGAT CACCTCCTAT CGCCGGAAAC CCATCCGGGG GGTTTAACCC CGGATGGGAT TTTGGTACTC CCCTTTATGG GGAGTCCCAC CGGCCGTACT CCTATAGCAA TTCCGGTTGA TCCTGCCGGA GGCCACTGCT ATGGGGGTCC GACTAAGCCA TGCGAGTCAT GGGGGTGTCC CTTCTGGGAC ACCACCGGCG GACGGCTCAG TAACACGTCG GTAACCTACC CT
16SBHB_7_3 CGT AGGGGAACCT ACGGCTCGAT CACCTCCTAT CGCCGGAAAC CCATCCGGGG GGTTTAACCC CGGATGGGAT TTTGGTACTC CCCTTTATGG GGAGTCCCAC CGGCCGTACT CCTATAGCAA TTCCGGTTGA TCCTGCCGGA GGCCACTGCT ATGGGGGTCC GACTAAGCCA TGCGAGTCAT GGGGGTGTCC CTTCTGGGAC ACCACCGGCG GACGGCTCAG TAACACGTCG GTAACCTACC CTCGGGAGGG GGATAACCCC GG
16SBHB_6_4 CGT AGGGGAACCT ACGGCTCGAT CACCTCCTAT CGCCGGAAAC CCATCCGGGG GGTTTAACCC CGGATGGGAT TTTGGTACTC CCCTTTATGG GGAGTCCCAC CGGCCGTACT CCTATAGCAA TTCCGGTTGA TCCTGCCGGA GGCCACTGCT ATGGGGGTCC GACTAAGCCA TGCGAGTCAT GGGGGTGTCC CTTCTGGGAC ACCACCGGCG GACGGCTCAG TAACACGTCG GTAACCTACC CTCGGGAGGG GGATAACCCC GGG
16SBHB_4_5 GTCGTAACA AGGTAGCCGT AGGGGAACCT ACGGCTCGAT CACCTCCTAT CGCCGGAAAC CCATCCGGGG GGTTTAACCC CGGATGGGAT TTTGGTACTC CCCTTTATGG GGAGTCCCAC CGGCCGTACT CCTATAGCAA TTCCGGTTGA TCCTGCCGGA GGCCACTGCT ATGGGGGTCC GACTAAGCCA TGCGAGTCAT GGGGGTGTCC CTTCTGGGAC ACCACCGGCG GACGGCTCAG TAACACGTCG GTAACCTACC CTCGGGAGGG GGATAACCCC GGG
16SBHB_3_9 CGT AGGGGAACCT ACGGCTCGAT CACCTCCTAT CGCCGGAAAC CCATCCGGGG GGTTTAACCC CGGATGGGAT TTTGGTACTC CCCTTTATGG GGAGTCCCAC CGGCCGTACT CCTATAGCAA TTCCGGTTGA TCCTGCCGGA GGCCACTGCT ATGGGGGTCC GACTAAGCCA TGCGAGTCAT GGGGGTGTCC CTTCTGGGAC ACCACCGGCG GACGGCTCAG TAACACGTCG GTAACCTACC CTCGGGAGGG GGATAACCCC G
16SBHB_6_3 GGGGAACCT ACGGCTCGAT CACCTCCTAT CGCCGGAAAC CCATCCGGGG GGTTTAACCC CGGATGGGAT TTTGGTACTC CCCTTTATGG GGAGTCCCAC CGGCCGTACT CCTATAGCAA TTCCGGTTGA TCCTGCCGGA GGCCACTGCT ATGGGGGTCC GACTAAGCCA TGCGAGTCAT GGGGGTGTCC CTTCTGGGAC ACCACCGGCG GACGGCTCAG TAACACGTCG GTAACCTACC CTCGGGAGGG GGATAACCCC GGG
16SBHB_3_8 GGGGAACCT ACGGCTCGAT CACCTCCTAT CGCCGGAAAC CCATCCGGGG GGTTTAACCC CGGATGGGAT TTTGGTACTC CCCTTTATGG GGAGTCCCAC CGGCCGTACT CCTATAGCAA TTCCGGTTGA TCCTGCCGGA GGCCACTGCT ATGGGGGTCC GACTAAGCCA TGCGAGTCAT GGGGGTGTCC CTTCTGGGAC ACCACCGGCG GACGGCTCAG TAACACGTCG GTAACCTACC CTCGGGAGGG GGATAACCCC GG
16SBHB_5_4 GGAACCT ACGGCTCGAT CACCTCCTAT CGCCGGAAAC CCATCCGGGG GGTTTAACCC CGGATGGGAT TTTGGTACTC CCCTTTATGG GGAGTCCCAC CGGCCGTACT CCTATAGCAA TTCCGGTTGA TCCTGCCGGA GGCCACTGCT ATGGGGGTCC GACTAAGCCA TGCGAGTCAT GGGGGTGTCC CTTCTGGGAC ACCACCGGCG GACGGCTCAG TAACACGTCG GTAACCTACC CTCGGGAGGG GGATAACCCC GGG
16SBHB_3_8 GGGAACCT ACGGCTCGAT CACCTCCTAT CGCCGGAAAC CCATCCGGGG GGTTTAACCC CGGATGGGAT TTTGGTACTC CCCTTTATGG GGAGTCCCAC CGGCCGTACT CCTATAGCAA TTCCGGTTGA TCCTGCCGGA GGCCACTGCT ATGGGGGTCC GACTAAGCCA TGCGAGTCAT GGGGGTGTCC CTTCTGGGAC ACCACCGGCG GACGGCTCAG TAACACGTCG GTAACCTACC CTCGGGAGGG GGATAACCCC GG
16SBHB_6_4 GAACCT ACGGCTCGAT CACCTCCTAT CGCCGGAAAC CCATCCGGGG GGTTTAACCC CGGATGGGAT TTTGGTACTC CCCTTTATGG GGAGTCCCAC CGGCCGTACT CCTATAGCAA TTCCGGTTGA TCCTGCCGGA GGCCACTGCT ATGGGGGTCC GACTAAGCCA TGCGAGTCAT GGGGGTGTCC CTTCTGGGAC ACCACCGGCG GACGGCTCAG TAACACGTCG GTAACCTACC CTCGGGAGGG GGATAACCCC GGG
16SBHB_6_4 CCT ACGGCTCGAT CACCTCCTAT CGCCGGAAAC CCATCCGGGG GGTTTAACCC CGGATGGGAT TTTGGTACTC CCCTTTATGG GGAGTCCCAC CGGCCGTACT CCTATAGCAA TTCCGGTTGA TCCTGCCGGA GGCCACTGCT ATGGGGGTCC GACTAAGCCA TGCGAGTCAT GGGGGTGTCC CTTCTGGGAC ACCACCGGCG GACGGCTCAG TAACACGTCG GTAACCTACC CTCGGGAGGG GGATAACCCC GGG
16SBHB_4_6 CT ACGGCTCGAT CACCTCCTAT CGCCGGAAAC CCATCCGGGG GGTTTAACCC CGGATGGGAT TTTGGTACTC CCCTTTATGG GGAGTCCCAC CGGCCGTACT CCTATAGCAA TTCCGGTTGA TCCTGCCGGA GGCCACTGCT ATGGGGGTCC GACTAAGCCA TGCGAGTCAT GGGGGTGTCC CTTCTGGGAC ACCACCGGCG GACGGCTCAG TAACACGTCG GTAACCTACC CTCGGGAGGG GGATAACCCC GGG
16SBHB_4_7 CCT ACGGCTCGAT CACCTCCTAT CGCCGGAAAC CCATCCGGGG GGTTTAACCC CGGATGGGAT TTTGGTACTC CCCTTTATGG GGAGTCCCAC CGGCCGTACT CCTATAGCAA TTCCGGTTGA TCCTGCCGGA GGCCACTGCT ATGGGGGTCC GACTAAGCCA TGCGAGTCAT GGGGGTGTCC CTTCTGGGAC ACCACCGGCG GACGGCTCAG TAACACGTCG GTAACCTACC CTCGGGAGGG GGATAACCCC GG
16SBHB_6_4 CCT ACGGCTCGAT CACCTCCTAT CGCCGGAAAC CCATCCGGGG GGTTTAACCC CGGATGGGAT TTTGGTACTC CCCTTTATGG GGAGTCCCAC CGGCCGTACT CCTATAGCAA TTCCGGTTGA TCCTGCCGGA GGCCACTGCT ATGGGGGTCC GACTAAGCCA TGCGAGTCAT GGGGGTGTCC CTTCTGGGAC ACCACCGGCG GACGGCTCAG TAACACGTCG GTAACCTACC CTCGGGAGGG GGATAACCCC G
16SBHB_4_6 CT ACGGCTCGAT CACCTCCTAT CGCCGGAAAC CCATCCGGGG GGTTTAACCC CGGATGGGAT TTTGGTACTC CCCTTTATGG GGAGTCCCAC CGGCCGTACT CCTATAGCAA TTCCGGTTGA TCCTGCCGGA GGCCACTGCT ATGGGGGTCC GACTAAGCCA TGCGAGTCAT GGGGGTGTCC CTTCTGGGAC ACCACCGGCG GACGGCTCAG TAACACGTCG GTAACCTACC CTCGGGAGGG GGATAACCCC GG
16SBHB_3_9 CGGCTCGAT CACCTCCTAT CGCCGGAAAC CCATCCGGGG GGTTTAACCC CGGATGGGAT TTTGGTACTC CCCTTTATGG GGAGTCCCAC CGGCCGTACT CCTATAGCAA TTCCGGTTGA TCCTGCCGGA GGCCACTGCT ATGGGGGTCC GACTAAGCCA TGCGAGTCAT GGGGGTGTCC CTTCTGGGAC ACCACCGGCG GACGGCTCAG TAACACGTCG GTAACCTACC CTCGGGAGGG GGATAACCCC GG
16SBHB_4_6 CGAT CACCTCCTAT CGCCGGAAAC CCATCCGGGG GGTTTAACCC CGGATGGGAT TTTGGTACTC CCCTTTATGG GGAGTCCCAC CGGCCGTACT CCTATAGCAA TTCCGGTTGA TCCTGCCGGA GGCCACTGCT ATGGGGGTCC GACTAAGCCA TGCGAGTCAT GGGGGTGTCC CTTCTGGGAC ACCACCGGCG GACGGCTCAG TAACACGTCG GTAACCTACC CTCGGGAGGG GGATAACCCC GGG
16SBHB_3_9 CGGCTCGAT CACCTCCTAT CGCCGGAAAC CCATCCGGGG GGTTTAACCC CGGATGGGAT TTTGGTACTC CCCTTTATGG GGAGTCCCAC CGGCCGTACT CCTATAGCAA TTCCGGTTGA TCCTGCCGGA GGCCACTGCT ATGGGGGTCC GACTAAGCCA TGCGAGTCAT GGGGGTGTCC CTTCTGGGAC ACCACCGGCG GACGGCTCAG TAACACGTCG GTAACCTACC CTCGGGAGGG G
16SBHB_8_2 CT ACGGCTCGAT CACCTCCTAT CGCCGGAAAC CCATCCGGGG GGTTTAACCC CGGATGGGAT TTTGGTACTC CCCTTTATGG GGAGTCCCAC CGGCCGTACT CCTATAGCAA TTCCGGTTGA TCCTGCCGGA GGCCACTGCT ATGGGGGTCC GACTAAGCCA TGCGAGTCAT GGGGGTGTCC CTTCTGGGAC ACCACCGGCG GACGGCTCAG TAACACGTCG GTAACCTACC CTCGGG
16SBHB_14_ CCT ACGGCTCGAT CACCTCCTAT CGCCGGAAAC CCATCCGGGG GGTTTAACCC CGGATGGGAT TTTGGTACTC CCCTTTATGG GGAGTCCCAC CGGCCGTACT CCTATAGCAA TTCCGGTTGA TCCTGCCGGA GGCCACTGCT ATGGGGGTCC GACTAAGCCA TGCGAGTCAT GGGGGTGTCC CTTCTGGGAC ACCACCGGCG GACGGCTCAG TAACACGTCG GTAACCTACC CTCGG
16SBHB_8_2 T ACGGCTCGAT CACCTCCTAT CGCCGGAAAC CCATCCGGGG GGTTTAACCC CGGATGGGAT TTTGGTACTC CCCTTTATGG GGAGTCCCAC CGGCCGTACT CCTATAGCAA TTCCGGTTGA TCCTGCCGGA GGCCACTGCT ATGGGGGTCC GACTAAGCCA TGCGAGTCAT GGGGGTGTCC CTTCTGGGAC ACCACCGGCG GACGGCTCAG TAACACGTCG GTAACCTACC CTCGGG
16SBHB_6_3 GGGGAACCT ACGGCTCGAT CACCTCCTAT CGCCGGAAAC CCATCCGGGG GGTTTAACCC CGGATGGGAT TTTGGTACTC CCCTTTATGG GGAGTCCCAC CGGCCGTACT CCTATAGCAA TTCCGGTTGA TCCTGCCGGA GGCCACTGCT ATGGGGGTCC GACTAAGCCA TGCGAGTCAT GGGGGTGTCC CTTCTGGGAC ACCACCGGCG GACGGCTCAG TAACACGTCG GTAACCTACC C
16SBHB_3_8 GGGAACCT ACGGCTCGAT CACCTCCTAT CGCCGGAAAC CCATCCGGGG GGTTTAACCC CGGATGGGAT TTTGGTACTC CCCTTTATGG GGAGTCCCAC CGGCCGTACT CCTATAGCAA TTCCGGTTGA TCCTGCCGGA GGCCACTGCT ATGGGGGTCC GACTAAGCCA TGCGAGTCAT GGGGGTGTCC CTTCTGGGAC ACCACCGGCG GACGGCTCAG TAACACGTCG GTAACCTACC C
16SBHB_5_4 GGAACCT ACGGCTCGAT CACCTCCTAT CGCCGGAAAC CCATCCGGGG GGTTTAACCC CGGATGGGAT TTTGGTACTC CCCTTTATGG GGAGTCCCAC CGGCCGTACT CCTATAGCAA TTCCGGTTGA TCCTGCCGGA GGCCACTGCT ATGGGGGTCC GACTAAGCCA TGCGAGTCAT GGGGGTGTCC CTTCTGGGAC ACCACCGGCG GACGGCTCAG TAACACGTCG GTAACCTACC C
16SBHB_4_6 CGT AGGGGAACCT ACGGCTCGAT CACCTCCTAT CGCCGGAAAC CCATCCGGGG GGTTTAACCC CGGATGGGAT TTTGGTACTC CCCTTTATGG GGAGTCCCAC CGGCCGTACT CCTATAGCAA TTCCGGTTGA TCCTGCCGGA GGCCACTGCT ATGGGGGTCC GACTAAGCCA TGCGAGTCAT GGGGGTGTCC CTTCTGGGAC ACCACCGGCG GACGGCTCAG TAACACGTCG GTAACCTAC
16SBHB_10_ CCT ACGGCTCGAT CACCTCCTAT CGCCGGAAAC CCATCCGGGG GGTTTAACCC CGGATGGGAT TTTGGTACTC CCCTTTATGG GGAGTCCCAC CGGCCGTACT CCTATAGCAA TTCCGGTTGA TCCTGCCGGA GGCCACTGCT ATGGGGGTCC GACTAAGCCA TGCGAGTCAT GGGGGTGTCC CTTCTGGGAC ACCACCGGCG GACGGCTCAG TAACACGTCG GTAACCTACC CTCG
16SBHB_14_ CGGCTCGAT CACCTCCTAT CGCCGGAAAC CCATCCGGGG GGTTTAACCC CGGATGGGAT TTTGGTACTC CCCTTTATGG GGAGTCCCAC CGGCCGTACT CCTATAGCAA TTCCGGTTGA TCCTGCCGGA GGCCACTGCT ATGGGGGTCC GACTAAGCCA TGCGAGTCAT GGGGGTGTCC CTTCTGGGAC ACCACCGGCG GACGGCTCAG TAACACGTCG GTAACCTACC CTCGGG
16SBHB_3_8 GGAACCT ACGGCTCGAT CACCTCCTAT CGCCGGAAAC CCATCCGGGG GGTTTAACCC CGGATGGGAT TTTGGTACTC CCCTTTATGG GGAGTCCCAC CGGCCGTACT CCTATAGCAA TTCCGGTTGA TCCTGCCGGA GGCCACTGCT ATGGGGGTCC GACTAAGCCA TGCGAGTCAT GGGGGTGTCC CTTCTGGGAC ACCACCGGCG GACGGCTCAG TAACACGTCG GTAACCTAC
16SBHB_4_6 GGCTCGAT CACCTCCTAT CGCCGGAAAC CCATCCGGGG GGTTTAACCC CGGATGGGAT TTTGGTACTC CCCTTTATGG GGAGTCCCAC CGGCCGTACT CCTATAGCAA TTCCGGTTGA TCCTGCCGGA GGCCACTGCT ATGGGGGTCC GACTAAGCCA TGCGAGTCAT GGGGGTGTCC CTTCTGGGAC ACCACCGGCG GACGGCTCAG TAACACGTCG GTAACCTACC CTCGGG
16SBHB_5_5 CCT ACGGCTCGAT CACCTCCTAT CGCCGGAAAC CCATCCGGGG GGTTTAACCC CGGATGGGAT TTTGGTACTC CCCTTTATGG GGAGTCCCAC CGGCCGTACT CCTATAGCAA TTCCGGTTGA TCCTGCCGGA GGCCACTGCT ATGGGGGTCC GACTAAGCCA TGCGAGTCAT GGGGGTGTCC CTTCTGGGAC ACCACCGGCG GACGGCTCAG TAACACGTCG GTAACCTACC C
16SBHB_3_9 CT ACGGCTCGAT CACCTCCTAT CGCCGGAAAC CCATCCGGGG GGTTTAACCC CGGATGGGAT TTTGGTACTC CCCTTTATGG GGAGTCCCAC CGGCCGTACT CCTATAGCAA TTCCGGTTGA TCCTGCCGGA GGCCACTGCT ATGGGGGTCC GACTAAGCCA TGCGAGTCAT GGGGGTGTCC CTTCTGGGAC ACCACCGGCG GACGGCTCAG TAACACGTCG GTAACCTACC
16SBHB_5_5 CCT ACGGCTCGAT CACCTCCTAT CGCCGGAAAC CCATCCGGGG GGTTTAACCC CGGATGGGAT TTTGGTACTC CCCTTTATGG GGAGTCCCAC CGGCCGTACT CCTATAGCAA TTCCGGTTGA TCCTGCCGGA GGCCACTGCT ATGGGGGTCC GACTAAGCCA TGCGAGTCAT GGGGGTGTCC CTTCTGGGAC ACCACCGGCG GACGGCTCAG TAACACGTCG GTAACCTACC
16SBHB_3_9 CTCGAT CACCTCCTAT CGCCGGAAAC CCATCCGGGG GGTTTAACCC CGGATGGGAT TTTGGTACTC CCCTTTATGG GGAGTCCCAC CGGCCGTACT CCTATAGCAA TTCCGGTTGA TCCTGCCGGA GGCCACTGCT ATGGGGGTCC GACTAAGCCA TGCGAGTCAT GGGGGTGTCC CTTCTGGGAC ACCACCGGCG GACGGCTCAG TAACACGTCG GTAACCTACC CTCGGG
16SBHB_3_8 GGGGAACCT ACGGCTCGAT CACCTCCTAT CGCCGGAAAC CCATCCGGGG GGTTTAACCC CGGATGGGAT TTTGGTACTC CCCTTTATGG GGAGTCCCAC CGGCCGTACT CCTATAGCAA TTCCGGTTGA TCCTGCCGGA GGCCACTGCT ATGGGGGTCC GACTAAGCCA TGCGAGTCAT GGGGGTGTCC CTTCTGGGAC ACCACCGGCG GACGGCTCAG TAACACGTCG GTAAC
16SBHB_4_7 CCT ACGGCTCGAT CACCTCCTAT CGCCGGAAAC CCATCCGGGG GGTTTAACCC CGGATGGGAT TTTGGTACTC CCCTTTATGG GGAGTCCCAC CGGCCGTACT CCTATAGCAA TTCCGGTTGA TCCTGCCGGA GGCCACTGCT ATGGGGGTCC GACTAAGCCA TGCGAGTCAT GGGGGTGTCC CTTCTGGGAC ACCACCGGCG GACGGCTCAG TAACACGTCG GTAACCTAC
16SBHB_13_ CGAT CACCTCCTAT CGCCGGAAAC CCATCCGGGG GGTTTAACCC CGGATGGGAT TTTGGTACTC CCCTTTATGG GGAGTCCCAC CGGCCGTACT CCTATAGCAA TTCCGGTTGA TCCTGCCGGA GGCCACTGCT ATGGGGGTCC GACTAAGCCA TGCGAGTCAT GGGGGTGTCC CTTCTGGGAC ACCACCGGCG GACGGCTCAG TAACACGTCG GTAACCTACC CTCGGG
16SBHB_10_ CCTAT CGCCGGAAAC CCATCCGGGG GGTTTAACCC CGGATGGGAT TTTGGTACTC CCCTTTATGG GGAGTCCCAC CGGCCGTACT CCTATAGCAA TTCCGGTTGA TCCTGCCGGA GGCCACTGCT ATGGGGGTCC GACTAAGCCA TGCGAGTCAT GGGGGTGTCC CTTCTGGGAC ACCACCGGCG GACGGCTCAG TAACACGTCG GTAACCTACC CTCGGGAGGG GGATAACCCC GGG
16SBHB_9_2 CTCCTAT CGCCGGAAAC CCATCCGGGG GGTTTAACCC CGGATGGGAT TTTGGTACTC CCCTTTATGG GGAGTCCCAC CGGCCGTACT CCTATAGCAA TTCCGGTTGA TCCTGCCGGA GGCCACTGCT ATGGGGGTCC GACTAAGCCA TGCGAGTCAT GGGGGTGTCC CTTCTGGGAC ACCACCGGCG GACGGCTCAG TAACACGTCG GTAACCTACC CTCGGGAGGG GGATAACCCC GGG
16SBHB_4_7 CCTAT CGCCGGAAAC CCATCCGGGG GGTTTAACCC CGGATGGGAT TTTGGTACTC CCCTTTATGG GGAGTCCCAC CGGCCGTACT CCTATAGCAA TTCCGGTTGA TCCTGCCGGA GGCCACTGCT ATGGGGGTCC GACTAAGCCA TGCGAGTCAT GGGGGTGTCC CTTCTGGGAC ACCACCGGCG GACGGCTCAG TAACACGTCG GTAACCTACC CTCGGGAGGG GGATAACCCC G
16SBHB_4_6 CGAT CACCTCCTAT CGCCGGAAAC CCATCCGGGG GGTTTAACCC CGGATGGGAT TTTGGTACTC CCCTTTATGG GGAGTCCCAC CGGCCGTACT CCTATAGCAA TTCCGGTTGA TCCTGCCGGA GGCCACTGCT ATGGGGGTCC GACTAAGCCA TGCGAGTCAT GGGGGTGTCC CTTCTGGGAC ACCACCGGCG GACGGCTCAG TAACACGTCG GTAACCTACC CTCGGGAG
16SBHB_3_9 CT ACGGCTCGAT CACCTCCTAT CGCCGGAAAC CCATCCGGGG GGTTTAACCC CGGATGGGAT TTTGGTACTC CCCTTTATGG GGAGTCCCAC CGGCCGTACT CCTATAGCAA TTCCGGTTGA TCCTGCCGGA GGCCACTGCT ATGGGGGTCC GACTAAGCCA TGCGAGTCAT GGGGGTGTCC CTTCTGGGAC ACCACCGGCG GACGGCTCAG TAACACGTCG GTAACCTA
16SBHB_3_8 GAT CACCTCCTAT CGCCGGAAAC CCATCCGGGG GGTTTAACCC CGGATGGGAT TTTGGTACTC CCCTTTATGG GGAGTCCCAC CGGCCGTACT CCTATAGCAA TTCCGGTTGA TCCTGCCGGA GGCCACTGCT ATGGGGGTCC GACTAAGCCA TGCGAGTCAT GGGGGTGTCC CTTCTGGGAC ACCACCGGCG GACGGCTCAG TAACACGTCG GTAACCTACC CTCGGG
16SBHB_5_4 CGGCTCGAT CACCTCCTAT CGCCGGAAAC CCATCCGGGG GGTTTAACCC CGGATGGGAT TTTGGTACTC CCCTTTATGG GGAGTCCCAC CGGCCGTACT CCTATAGCAA TTCCGGTTGA TCCTGCCGGA GGCCACTGCT ATGGGGGTCC GACTAAGCCA TGCGAGTCAT GGGGGTGTCC CTTCTGGGAC ACCACCGGCG GACGGCTCAG TAACACGTCG GTAACCTACC C
16SBHB_4_6 GGAACCT ACGGCTCGAT CACCTCCTAT CGCCGGAAAC CCATCCGGGG GGTTTAACCC CGGATGGGAT TTTGGTACTC CCCTTTATGG GGAGTCCCAC CGGCCGTACT CCTATAGCAA TTCCGGTTGA TCCTGCCGGA GGCCACTGCT ATGGGGGTCC GACTAAGCCA TGCGAGTCAT GGGGGTGTCC CTTCTGGGAC ACCACCGGCG GACGGCTCAG TAACACGTCG GTAG
16SBHB_22_ GGAACCT ACGGCTCGAT CACCTCCTAT CGCCGGAAAC CCATCCGGGG GGTTTAACCC CGGATGGGAT TTTGGTACTC CCCTTTATGG GGAGTCCCAC CGGCCGTACT CCTATAGCAA TTCCGGTTGA TCCTGCCGGA GGCCACTGCT ATGGGGGTCC GACTAAGCCA TGCGAGTCAT GGGGGTGTCC CTTCTGGGAC ACCACCGGCG GACGGCTCAG TAACACGTCG
16SBHB_19_ GGAACCT ACGGCTCGAT CACCTCCTAT CGCCGGAAAC CCATCCGGGG GGTTTAACCC CGGATGGGAT TTTGGTACTC CCCTTTATGG GGAGTCCCAC CGGCCGTACT CCTATAGCAA TTCCGGTTGA TCCTGCCGGA GGCCACTGCT ATGGGGGTCC GACTAAGCCA TGCGAGTCAT GGGGGTGTCC CTTCTGGGAC ACCACCGGCG GACGGCTCAG TAACACGTCG G
16SBHB_3_8 GGGAACCT ACGGCTCGAT CACCTCCTAT CGCCGGAAAC CCATCCGGGG GGTTTAACCC CGGATGGGAT TTTGGTACTC CCCTTTATGG GGAGTCCCAC CGGCCGTACT CCTATAGCAA TTCCGGTTGA TCCTGCCGGA GGCCACTGCT ATGGGGGTCC GACTAAGCCA TGCGAGTCAT GGGGGTGTCC CTTCTGGGAC ACCACCGGCG GACGGCTCAG TAACACGCCG G
16SBHB_3_9 GAACCT ACGGCTCGAT CACCTCCTAT CGCCGGAAAC CCATCCGGGG GGTTTAACCC CGGATGGGAT TTTGGTACTC CCCTTTATGG GGAGTCCCAC CGGCCGTACT CCTATAGCAA TTCCGGTTGA TCCTGCCGGA GGCCACTGCT ATGGGGGTCC GACTAAGCCA TGCGAGTCAT GGGGGTGTCC CTTCTGGGAC ACCACCGGCG GACGGCTCAG TAACACGTCG G
16SBHB_5_4 GGGAACCT ACGGCTCGAT CACCTCCTAT CGCCGGAAAC CCATCCGGGG GGTTTAACCC CGGATGGGAT TTTGGTACTC CCCTTTATGG GGAGTCCCAC CGGCCGTACT CCTATAGCAA TTCCGGTTGA TCCTGCCGGA GGCCACTGCT ATGGGGGTCC GACTAAGCCA TGCGAGTCAT GGGGGTGTCC CTTCTGGGAC ACCACCGGCG GACGGCTCAG TAACACGTCG
16SBHB_3_9 GAACCT ACGGCTCGAT CACCTCCTAT CGCCGGAAAC CCATCCGGGG GGTTTAACCC CGGATGGGAT TTTGGTACTC CCCTTTATGG GGAGTCCCAC CGGCCGTACT CCTATAGCAA TTCCGGTTGA TCCTGCCGGA GGCCACTGCT ATGGGGGTCC GACTAAGCCA TGCGAGTCAT GGGGGTGTCC CTTCTGGGAC ACCACCGGCG GACGGCTCAG TAACACGTCG
16SBHB_20_ GTCGTAACA AGGTAGCCGT AGGGGAACCT ACGGCTCGAT CACCTCCTAT CGCCGGAAAC CCATCCGGGG GGTTTAACCC CGGATGGGAT TTTGGTACTC CCCTTTATGG GGAGTCCCAC CGGCCGTACT CCTATAGCAA TTCCGGTTGA TCCTGCCGGA GGCCACTGCT ATGGGGGTCC GACTAAGCCA TGCGAGTCAT GGGGGTGTCC CTTCTGGGAC ACCACCGGCG GACGGCTCAG TAACACGTCG G
16SBHB_18_ GTCGTAACA AGGTAGCCGT AGGGGAACCT ACGGCTCGAT CACCTCCTAT CGCCGGAAAC CCATCCGGGG GGTTTAACCC CGGATGGGAT TTTGGTACTC CCCTTTATGG GGAGTCCCAC CGGCCGTACT CCTATAGCAA TTCCGGTTGA TCCTGCCGGA GGCCACTGCT ATGGGGGTCC GACTAAGCCA TGCGAGTCAT GGGGGTGTCC CTTCTGGGAC ACCACCGGCG GACGGCTCAG TAACACGTCG
16SBHB_8_2 GTCGTAACA AGGTAGCCGT AGGGGAACCT ACGGCTCGAT CACCTCCTAT CGCCGGAAAC CCATCCGGGG GGTTTAACCC CGGATGGGAT TTTGGTACTC CCCTTTATGG GGAGTCCCAC CGGCCGTACT CCTATAGCAA TTCCGGTTGA TCCTGCCGGA GGCCACTGCT ATGGGGGTCC GACTAAGCCA TGCGAGTCAT GGGGGTGTCC CTTCTGGGAC ACCACCGGCG GACGGCTCAG TAACACGTCG GTAACCTACC CTCGGG
16SBHB_7_3 GTCGTAACA AGGTAGCCGT AGGGGAACCT ACGGCTCGAT CACCTCCTAT CGCCGGAAAC CCATCCGGGG GGTTTAACCC CGGATGGGAT TTTGGTACTC CCCTTTATGG GGAGTCCCAC CGGCCGTACT CCTATAGCAA TTCCGGTTGA TCCTGCCGGA GGCCACTGCT ATGGGGGTCC GACTAAGCCA TGCGAGTCAT GGGGGTGTCC CTTCTGGGAC ACCACCGGCG GACGGCTCAG TAACACGTCG GTAACCTACC CTCGG
16SBHB_7_3 GTCGTAACA AGGTAGCCGT AGGGGAACCT ACGGCTCGAT CACCTCCTAT CGCCGGAAAC CCATCCGGGG GGTTTAACCC CGGATGGGAT TTTGGTACTC CCCTTTATGG GGAGTCCCAC CGGCCGTACT CCTATAGCAA TTCCGGTTGA TCCTGCCGGA GGCCACTGCT ATGGGGGTCC GACTAAGCCA TGCGAGTCAT GGGGGTGTCC CTTCTGGGAC ACCACCGGCG GACGGCTCAG TAACACGTCG GTAACCTACC C
16SBHB_4_7 CA AGGTAGCCGT AGGGGAACCT ACGGCTCGAT CACCTCCTAT CGCCGGAAAC CCATCCGGGG GGTTTAACCC CGGATGGGAT TTTGGTACTC CCCTTTATGG GGAGTCCCAC CGGCCGTACT CCTATAGCAA TTCCGGTTGA TCCTGCCGGA GGCCACTGCT ATGGGGGTCC GACTAAGCCA TGCGAGTCAT GGGGGTGTCC CTTCTGGGAC ACCACCGGCG GACGGCTCAG TAACACGTCG GTAACCTACC C
16SBHB_3_8 GTCGTAACA AGGTAGCCGT AGGGGAACCT ACGGCTCGAT CACCTCCTAT CGCCGGAAAC CCATCCGGGG GGTTTAACCC CGGATGGGAT TTTGGTACTC CCCTTTATGG GGAGTCCCAC CGGCCGTACT CCTATAGCAA TTCCGGTTGA TCCTGCCGGA GGCCACTGCT ATGGGGGTCC GACTAAGCCA TGCGAGTCAT GGGGGTGTCC CTTCTGGGAC ACCACCGGCG GACGGCTCAG TAACACGCCG
16SBHB_4_6 CGTAACA AGGTAGCCGT AGGGGAACCT ACGGCTCGAT CACCTCCTAT CGCCGGAAAC CCATCCGGGG GGTTTAACCC CGGATGGGAT TTTGGTACTC CCCTTTATGG GGAGTCCCAC CGGCCGTACT CCTATAGCAA TTCCGGTTGA TCCTGCCGGA GGCCACTGCT ATGGGGGTCC GACTAAGCCA TGCGAGTCAT GGGGGTGTCC CTTCTGGGAC ACCACCGGCG GACGGCTCAG TAACACGTCG G
16SBHB_7_3 GTCGTAACA AGGTAGCCGT AGGGGAACCT ACGGCTCGAT CACCTCCTAT CGCCGGAAAC CCATCCGGGG GGTTTAACCC CGGATGGGAT TTTGGTACTC CCCTTTATGG GGAGTCCCAC CGGCCGTACT CCTATAGCAA TTCCGGTTGA TCCTGCCGGA GGCCACTGCT ATGGGGGTCC GACTAAGCCA TGCGAGTCAT GGGGGTGTCC CTTCTGGGAC ACCACCGGCG GACGGCTCAG TAACAC
16SBHB_3_8 TAACA AGGTAGCCGT AGGGGAACCT ACGGCTCGAT CACCTCCTAT CGCCGGAAAC CCATCCGGGG GGTTTAACCC CGGATGGGAT TTTGGTACTC CCCTTTATGG GGAGTCCCAC CGGCCGTACT CCTATAGCAA TTCCGGTTGA TCCTGCCGGA GGCCACTGCT ATGGGGGTCC GACTAAGCCA TGCGAGTCAT GGGGGTGTCC CTTCTGGGAC ACCACCGGCG GACGGCTCAG TAACACGTCG
16SBHB_3_8 GTAGCCGT AGGGGAACCT ACGGCTCGAT CACCTCCTAT CGCCGGAAAC CCATCCGGGG GGTTTAACCC CGGATGGGAT TTTGGTACTC CCCTTTATGG GGAGTCCCAC CGGCCGTACT CCTATAGCAA TTCCGGTTGA TCCTGCCGGA GGCCACTGCT ATGGGGGTCC GACTAAGCCA TGCGAGTCAT GGGGGTGTCC CTTCTGGGAC ACCACCGGCG GACGGCTCAG TAACACGTCG GTAACCTAC
16SBHB_5_4 GTCGTAACA AGGTAGCCGT AGGGGAACCT ACGGCTCGAT CACCTCCTAT CGCCGGAAAC CCATCCGGGG GGTTTAACCC CGGATGGGAT TTTGGTACTC CCCTTTATGG GGAGTCCCAC CGGCCGTACT CCTATAGCAA TTCCGGTTGA TCCTGCCGGA GGCCACTGCT ATGGGGGTCC GACTAAGCCA TGCGAGTCAT GGGGGTGTCC CTTCTGGGAC ACCACCGGCG GACGGCTCAG TAAC
16SBHB_8_2 GGTAGCCGT AGGGGAACCT ACGGCTCGAT CACCTCCTAT CGCCGGAAAC CCATCCGGGG GGTTTAACCC CGGATGGGAT TTTGGTACTC CCCTTTATGG GGAGTCCCAC CGGCCGTACT CCTATAGCAA TTCCGGTTGA TCCTGCCGGA GGCCACTGCT ATGGGGGTCC GACTAAGCCA TGCGAGTCAT GGGGGTGTCC CTTCTGGGAC ACCACCGGCG GACGGCTCAG TAACACGTCG
16SBHB_3_8 GTAGCCGT AGGGGAACCT ACGGCTCGAT CACCTCCTAT CGCCGGAAAC CCATCCGGGG GGTTTAACCC CGGATGGGAT TTTGGTACTC CCCTTTATGG GGAGTCCCAC CGGCCGTACT CCTATAGCAA TTCCGGTTGA TCCTGCCGGA GGCCACTGCT ATGGGGGTCC GACTAAGCCA TGCGAGTCAT GGGGGTGTCC CTTCTGGGAC ACCACCGGCG GACGGCTCAG TAACACGTCG G
16SBHB_6_3 GTAGCCGT AGGGGAACCT ACGGCTCGAT CACCTCCTAT CGCCGGAAAC CCATCCGGGG GGTTTAACCC CGGATGGGAT TTTGGTACTC CCCTTTATGG GGAGTCCCAC CGGCCGTACT CCTATAGCAA TTCCGGTTGA TCCTGCCGGA GGCCACTGCT ATGGGGGTCC GACTAAGCCA TGCGAGTCAT GGGGGTGTCC CTTCTGGGAC ACCACCGGCG GACGGCTCAG TAACACGTCG
16SBHB_5_4 GTCC GGGCTCCGTG AGGGGGGAGA AGTCGTAACA AGGTAGCCGT AGGGGAACCT ACGGCTCGAT CACCTCCTAT CGCCGGAAAC CCATCCGGGG GGTTTAACCC CGGATGGGAT TTTGGTACTC CCCTTTATGG GGAGTCCCAC CGGCCGTACT CCTATAGCAA TTCCGGTTGA TCCTGCCGGA GGCCACTGCT ATGGGGGTCC GACTAAGCCA TGCGAGTCAT GGGGGTGTCC CTTCTGGGAC ACCACCGGCG GACGGCTCAG TAACACGTCG
16SBHB_5_5 CGAGTCC GGGCTCCGTG AGGGGGGAGA AGTCGTAACA AGGTAGCCGT AGGGGAACCT ACGGCTCGAT CACCTCCTAT CGCCGGAAAC CCATCCGGGG GGTTTAACCC CGGATGGGAT TTTGGTACTC CCCTTTATGG GGAGTCCCAC CGGCCGTACT CCTATAGCAA TTCCGGTTGA TCCTGCCGGA GGCCACTGCT ATGGGGGTCC GACTAAGCCA TGCGAGTCAT GGGGGTGTCC CTTCTGGGAC ACCACCGGCG GACGGCTCAG TAACACGTCG
16SBHB_4_7 CC GGGCTCCGTG AGGGGGGAGA AGTCGTAACA AGGTAGCCGT AGGGGAACCT ACGGCTCGAT CACCTCCTAT CGCCGGAAAC CCATCCGGGG GGTTTAACCC CGGATGGGAT TTTGGTACTC CCCTTTATGG GGAGTCCCAC CGGCCGTACT CCTATAGCAA TTCCGGTTGA TCCTGCCGGA GGCCACTGCT ATGGGGGTCC GACTAAGCCA TGCGAGTCAT GGGGGTGTCC CTTCTGGGAC ACCACCGGCG GACGGCTCAG TAACACGTCG
16SBHB_4_6 GCTCCGTG AGGGGGGAGA AGTCGTAACA AGGTAGCCGT AGGGGAACCT ACGGCTCGAT CACCTCCTAT CGCCGGAAAC CCATCCGGGG GGTTTAACCC CGGATGGGAT TTTGGTACTC CCCTTTATGG GGAGTCCCAC CGGCCGTACT CCTATAGCAA TTCCGGTTGA TCCTGCCGGA GGCCACTGCT ATGGGGGTCC GACTAAGCCA TGCGAGTCAT GGGGGTGTCC CTTCTGGGAC ACCACCGGCG GACGGCTCAG TAACACGTCG
16SBHB_3_8 GTG AGGGGGGAGA AGTCGTAACA AGGTAGCCGT AGGGGAACCT ACGGCTCGAT CACCTCCTAT CGCCGGAAAC CCATCCGGGG GGTTTAACCC CGGATGGGAT TTTGGTACTC CCCTTTATGG GGAGTCCCAC CGGCCGTACT CCTATAGCAA TTCCGGTTGA TCCTGCCGGA GGCCACTGCT ATGGGGGTCC GACTAAGCCA TGCGAGTCAT GGGGGTGTCC CTTCTGGGAC ACCACCGGCG GACGGCTCAG TAACACGTCG
16SBHB_4_6 GGCTCCGTG AGGGGGGAGA AGTCGTAACA AGGTAGCCGT AGGGGAACCT ACGGCTCGAT CACCTCCTAT CGCCGGAAAC CCATCCGGGG GGTTTAACCC CGGATGGGAT TTTGGTACTC CCCTTTATGG GGAGTCCCAC CGGCCGTACT CCTATAGCAA TTCCGGTTGA TCCTGCCGGA GGCCACTGCT ATGGGGGTCC GACTAAGCCA TGCGAGTCAT GGGGGTGTCC CTTCTGGGAC ACCACCGGCG GACGGCTCAG
16SBHB_3_8 GCTCCGTG AGGGGGGAGA AGTCGTAACA AGGTAGCCGT AGGGGAACCT ACGGCTCGAT CACCTCCTAT CGCCGGAAAC CCATCCGGGG GGTTTAACCC CGGATGGGAT TTTGGTACTC CCCTTTATGG GGAGTCCCAC CGGCCGTACT CCTATAGCAA TTCCGGTTGA TCCTGCCGGA GGCCACTGCT ATGGGGGTCC GACTAAGCCA TGCGAGTCAT GGGGGTGTCC CTTCTGGGAC ACCACCGGCG GACGGCTCAG
16SBHB_4_6 GGAGA AGTCGTAACA AGGTAGCCGT AGGGGAACCT ACGGCTCGAT CACCTCCTAT CGCCGGAAAC CCATCCGGGG GGTTTAACCC CGGATGGGAT TTTGGTACTC CCCTTTATGG GGAGTCCCAC CGGCCGTACT CCTATAGCAA TTCCGGTTGA TCCTGCCGGA GGCCACTGCT ATGGGGGTCC GACTAAGCCA TGCGAGTCAT GGGGGTGTCC CTTCTGGGAC ACCACCGGCG GACGGCTCAG TAACACGTCG
16SBHB_3_8 GGGGAGA AGTCGTAACA AGGTAGCCGT AGGGGAACCT ACGGCTCGAT CACCTCCTAT CGCCGGAAAC CCATCCGGGG GGTTTAACCC CGGATGGGAT TTTGGTACTC CCCTTTATGG GGAGTCCCAC CGGCCGTACT CCTATAGCAA TTCCGGTTGA TCCTGCCGGA GGCCACTGCT ATGGGGGTCC GACTAAGCCA TGCGAGTCAT GGGGGTGTCC CTTCTGGGAC ACCACCGGCG GACGGCTCAG TAACACGTCG G
16SBHB_4_5 GTCC GGGCTCCGTG AGGGGGGAGA AGTCGTAACA AGGTAGCCGT AGGGGAACCT ACGGCTCGAT CACCTCCTAT CGCCGGAAAC CCATCCGGGG GGTTTAACCC CGGATGGGAT TTTGGTACTC CCCTTTATGG GGAGTCCCAC CGGCCGTACT CCTATAGCAA TTCCGGTTGA TCCTGCCGGA GGCCACTGCT ATGGGGGTCC GACTAAGCCA TGCGAGTCAT GGGGGTGTCC CTTCTGGGAC ACCACCGGCG
16SBHB_4_6 CGG GGTCGAGTCC GGGCTCCGTG AGGGGGGAGA AGTCGTAACA AGGTAGCCGT AGGGGAACCT ACGGCTCGAT CACCTCCTAT CGCCGGAAAC CCATCCGGGG GGTTTAACCC CGGATGGGAT TTTGGTACTC CCCTTTATGG GGAGTCCCAC CGGCCGTACT CCTATAGCAA TTCCGGTTGA TCCTGCCGGA GGCCACTGCT ATGGGGGTCC GACTAAGCCA TGCGAGTCAT GGGGGTGTCC CTTCTGGGAC ACCACCGGCG GG
16SBHB_3_9 CGTCGC CGGCTCTGAC CGGATGGCGG GGTCGAGTCC GGGCTCCGTG AGGGGGGAGA AGTCGTAACA AGGTAGCCGT AGGGGAACCT ACGGCTCGAT CACCTCCTAT CGCCGGAAAC CCATCCGGGG GGTTTAACCC CGGATGGGAT TTTGGTACTC CCCTTTATGG GGAGTCCCAC CGGCCGTACT CCTATAGCAA TTCCGGTTGA TCCTGCCGGA GGCCACTGCT ATGGGGGTCC GACTAAGCCA TGCGAGTCAT GGGGGTGTCC CTTCTGGGAC ACCACCGGCG
16SBHB_3_9 CGG GGTCGAGTCC GGGCTCCGTG AGGGGGGAGA AGTCGTAACA AGGTAGCCGT AGGGGAACCT ACGGCTCGAT CACCTCCTAT CGCCGGAAAC CCATCCGGGG GGTTTAACCC CGGATGGGAT TTTGGTACTC CCCTTTATGG GGAGTCCCAC CGGCCGTACT CCTATAGCAA TTCCGGTTGA TCCTGCCGGA GGCCACTGCT ATGGGGGTCC GACTAAGCCA TGCGAGTCAT GGGGGTGTCC CTTCTGGGAC ACCACCGGCG
16SBHB_4_7 CC GGGCTCCGTG AGGGGGGAGA AGTCGTAACA AGGTAGCCGT AGGGGAACCT ACGGCTCGAT CACCTCCTAT CGCCGGAAAC CCATCCGGGG GGTTTAACCC CGGATGGGAT TTTGGTACTC CCCTTTATGG GGAGTCCCAC CGGCCGTACT CCTATAGCAA TTCCGGTTGA TCCTGCCGGA GGCCACTGCT ATGGGGGTCC GACTAAGCCA TGCGAGTCAT GGGGGTGTCC CTTCTGGGAC ACCACCGGCG
16SBHB_4_5 GTCGTAACA AGGTAGCCGT AGGGGAACCT ACGGCTCGAT CACCTCCTAT CGCCGGAAAC CCATCCGGGG GGTTTAACCC CGGATGGGAT TTTGGTACTC CCCTTTATGG GGAGTCCCAC CGGCCGTACT CCTATAGCAA TTCCGGTTGA TCCTGCCGGA GGCCACTGCT ATGGGGGTCC GACTAAGCCA TGCGAGTCAT GGGGGTGTCC CTTCTGGGAC ACCACCGGCG GACGGCTCAG
16SBHB_3_8 GTCGTAACA AGGTAGCCGT AGGGGAACCT ACGGCTCGAT CACCTCCTAT CGCCGGAAAC CCATCCGGGG GGTTTAACCC CGGATGGGAT TTTGGTACTC CCCTTTATGG GGAGTCCCAC CGGCCGTACT CCTATAGCAA TTCCGGTTGA TCCTGCCGGA GGCCACTGCT ATGGGGGTCC GACTAAGCCA TGCGAGTCAT GGGGGTGTCC CTTCTGGGAC ACCACCGGCG GACGGC
16SBHB_3_8 GGGGAGA AGTCGTAACA AGGTAGCCGT AGGGGAACCT ACGGCTCGAT CACCTCCTAT CGCCGGAAAC CCATCCGGGG GGTTTAACCC CGGATGGGAT TTTGGTACTC CCCTTTATGG GGAGTCCCAC CGGCCGTACT CCTATAGCAA TTCCGGTTGA TCCTGCCGGA GGCCACTGCT ATGGGGGTCC GACTAAGCCA TGCGAGTCAT GGGGGTGTCC CTTCTGGGAC ACCACCGGCG
16SBHB_7_3 GTCGTAACA AGGTAGCCGT AGGGGAACCT ACGGCTCGAT CACCTCCTAT CGCCGGAAAC CCATCCGGGG GGTTTAACCC CGGATGGGAT TTTGGTACTC CCCTTTATGG GGAGTCCCAC CGGCCGTACT CCTATAGCAA TTCCGGTTGA TCCTGCCGGA GGCCACTGCT ATGGGGGTCC GACTAAGCCA TGCGAGTCAT GGGGGTGTCC CTTCTGGGAC ACCACCGGCG GAC
```

```
16SBHB_7_3 GTCGTAACA AGGTAGCCGT AGGGGAACCT ACGGCTCGAT CACCTCCTAT CGCCGGAAAC CCATCGGGG GGTTTAACCC CGGATGGGAT TTTGGTACTC CCCTTTATGG GGAGTCCCAC CGGCCGTACT CCTATAGCAA TTCCGGTTGA TCCTGCCGGA GGCCACTGCT ATGGGGGTCC GACTAAGCCA TGCGAGTCAT GGGGGTGTCC CTTCTGGGAC ACCACCGGCG G
16SBHB_10_ GTCGTAACA AGGTAGCCGT AGGGGAACCT ACGGCTCGAT CACCTCCTAT CGCCGGAAAC CCATCGGGG GGTTTAACCC CGGATGGGAT TTTGGTACTC CCCTTTATGG GGAGTCCCAC CGGCCGTACT CCTATAGCAA TTCCGGTTGA TCCTGCCGGA GGCCACTGCT ATGGGGGTCC GACTAAGCCA TGCGAGTCAT GGGGGTGTCC CTTCTGGGAC ACCACCGGCG
16SBHB_3_8 GTCGTAACA AGGTAGCCGT AGGGGAACCT ACGGCTCGAT CACCTCCTAT CGCCGGAAAC CCATCGGGG GGTTTAACCC CGGATGGGAT TTTGGTACTC CCCTTTATGG GGAGTCCCAC CGGCCGTACT CCTATAGCAA TTCCGGTTGA TCCTGCCGGA GGCCACTGCT ATGGGGGTCC GACTAAGCCA TGCGAGTCAT GGGGGTGTCC CTTCTGGGAC ACCACCGGC
16SBHB_3_8 TAACA AGGTAGCCGT AGGGGAACCT ACGGCTCGAT CACCTCCTAT CGCCGGAAAC CCATCGGGG GGTTTAACCC CGGATGGGAT TTTGGTACTC CCCTTTATGG GGAGTCCCAC CGGCCGTACT CCTATAGCAA TTCCGGTTGA TCCTGCCGGA GGCCACTGCT ATGGGGGTCC GACTAAGCCA TGCGAGTCAT GGGGGTGTCC CTTCTGGGAC ACCACCGGCG GAC
16SBHB_14_ CGT AGGGGAACCT ACGGCTCGAT CACCTCCTAT CGCCGGAAAC CCATCGGGG GGTTTAACCC CGGATGGGAT TTTGGTACTC CCCTTTATGG GGAGTCCCAC CGGCCGTACT CCTATAGCAA TTCCGGTTGA TCCTGCCGGA GGCCACTGCT ATGGGGGTCC GACTAAGCCA TGCGAGTCAT GGGGGTGTCC CTTCTGGGAC ACCACCGGCG GACGGCTCAG TAACAC
16SBHB_11_ CGT AGGGGAACCT ACGGCTCGAT CACCTCCTAT CGCCGGAAAC CCATCGGGG GGTTTAACCC CGGATGGGAT TTTGGTACTC CCCTTTATGG GGAGTCCCAC CGGCCGTACT CCTATAGCAA TTCCGGTTGA TCCTGCCGGA GGCCACTGCT ATGGGGGTCC GACTAAGCCA TGCGAGTCAT GGGGGTGTCC CTTCTGGGAC ACCACCGGCG GACGGCTCAG TAAC
16SBHB_15_ CGT AGGGGAACCT ACGGCTCGAT CACCTCCTAT CGCCGGAAAC CCATCGGGG GGTTTAACCC CGGATGGGAT TTTGGTACTC CCCTTTATGG GGAGTCCCAC CGGCCGTACT CCTATAGCAA TTCCGGTTGA TCCTGCCGGA GGCCACTGCT ATGGGGGTCC GACTAAGCCA TGCGAGTCAT GGGGGTGTCC CTTCTGGGAC ACCACCGGCG GACGGCTCAG
16SBHB_4_7 CCGT AGGGGAACCT ACGGCTCGAT CACCTCCTAT CGCCGGAAAC CCATCGGGG GGTTTAACCC CGGATGGGAT TTTGGTACTC CCCTTTATGG GGAGTCCCAC CGGCCGTACT CCTATAGCAA TTCCGGTTGA TCCTGCCGGA GGCCACTGCT ATGGGGGTCC GACTAAGCCA TGCGAGTCAT GGGGGTGTCC CTTCTGGGAC ACCACCGGCG GACGGCTCAG
16SBHB_6_3 GGGGAACCT ACGGCTCGAT CACCTCCTAT CGCCGGAAAC CCATCGGGG GGTTTAACCC CGGATGGGAT TTTGGTACTC CCCTTTATGG GGAGTCCCAC CGGCCGTACT CCTATAGCAA TTCCGGTTGA TCCTGCCGGA GGCCACTGCT ATGGGGGTCC GACTAAGCCA TGCGAGTCAT GGGGGTGTCC CTTCTGGGAC ACCACCGGCG GACGGCTCAG TAAC
16SBHB_4_7 AGGGGAACCT ACGGCTCGAT CACCTCCTAT CGCCGGAAAC CCATCGGGG GGTTTAACCC CGGATGGGAT TTTGGTACTC CCCTTTATGG GGAGTCCCAC CGGCCGTACT CCTATAGCAA TTCCGGTTGA TCCTGCCGGA GGCCACTGCT ATGGGGGTCC GACTAAGCCA TGCGAGTCAT GGGGGTGTCC CTTCTGGGAC ACCACCGGCG GACGGCTCAG TAAC
16SBHB_3_8 GGGAACCT ACGGCTCGAT CACCTCCTAT CGCCGGAAAC CCATCGGGG GGTTTAACCC CGGATGGGAT TTTGGTACTC CCCTTTATGG GGAGTCCCAC CGGCCGTACT CCTATAGCAA TTCCGGTTGA TCCTGCCGGA GGCCACTGCT ATGGGGGTCC GACTAAGCCA TGCGAGTCAT GGGGGTGTCC CTTCTGGGAC ACCACCGGCG GACGGCTCAG TAAC
16SBHB_4_6 GGGGAACCT ACGGCTCGAT CACCTCCTAT CGCCGGAAAC CCATCGGGG GGTTTAACCC CGGATGGGAT TTTGGTACTC CCCTTTATGG GGAGTCCCAC CGGCCGTACT CCTATAGCAA TTCCGGTTGA TCCTGCCGGA GGCCACTGCT ATGGGGGTCC GACTAAGCCA TGCGAGTCAT GGGGGTGTCC CTTCTGGGAC ACCACCGGCG GACGGCTCAG TAACAC
16SBHB_3_8 GGAACCT ACGGCTCGAT CACCTCCTAT CGCCGGAAAC CCATCGGGG GGTTTAACCC CGGATGGGAT TTTGGTACTC CCCTTTATGG GGAGTCCCAC CGGCCGTACT CCTATAGCAA TTCCGGTTGA TCCTGCCGGA GGCCACTGCT ATGGGGGTCC GACTAAGCCA TGCGAGTCAT GGGGGTGTCC CTTCTGGGAC ACCACCGGCG GACGGCTCAG TAACAC
16SBHB_3_8 GGAACCT ACGGCTCGAT CACCTCCTAT CGCCGGAAAC CCATCGGGG GGTTTAACCC CGGATGGGAT TTTGGTACTC CCCTTTATGG GGAGTCCCAC CGGCCGTACT CCTATAGCAA TTCCGGTTGA TCCTGCCGGA GGCCACTGCT ATGGGGGTCC GACTAAGCCA TGCGAGTCAT GGGGGTGTCC CTTCTGGGAC ACCACCGGCG GACGGCTCAG TAAC
16SBHB_4_6 GAACCT ACGGCTCGAT CACCTCCTAT CGCCGGAAAC CCATCGGGG GGTTTAACCC CGGATGGGAT TTTGGTACTC CCCTTTATGG GGAGTCCCAC CGGCCGTACT CCTATAGCAA TTCCGGTTGA TCCTGCCGGA GGCCACTGCT ATGGGGGTCC GACTAAGCCA TGCGAGTCAT GGGGGTGTCC CTTCTGGGAC ACCACCGGCG GACGGCTCAG TAACAC
16SBHB_24_ CGGCTCGAT CACCTCCTAT CGCCGGAAAC CCATCGGGG GGTTTAACCC CGGATGGGAT TTTGGTACTC CCCTTTATGG GGAGTCCCAC CGGCCGTACT CCTATAGCAA TTCCGGTTGA TCCTGCCGGA GGCCACTGCT ATGGGGGTCC GACTAAGCCA TGCGAGTCAT GGGGGTGTCC CTTCTGGGAC ACCACCGGCG GACGGCTCAG TAACACGTCG
16SBHB_7_3 CGGCTCGAT CACCTCCTAT CGCCGGAAAC CCATCGGGG GGTTTAACCC CGGATGGGAT TTTGGTACTC CCCTTTATGG GGAGTCCCAC CGGCCGTACT CCTATAGCAA TTCCGGTTGA TCCTGCCGGA GGCCACTGCT ATGGGGGTCC GACTAAGCCA TGCGAGTCAT GGGGGTGTCC CTTCTGGGAC ACCACCGGCG GACGGCTCAG TAACACGTCG G
16SBHB_5_4 CGGCTCGAT CACCTCCTAT CGCCGGAAAC CCATCGGGG GGTTTAACCC CGGATGGGAT TTTGGTACTC CCCTTTATGG GGAGTCCCAC CGGCCGTACT CCTATAGCAA TTCCGGTTGA TCCTGCCGGA GGCCACTGCT ATGGGGGTCC GACTAAGCCA TGCGAGTCAT GGGGGTGTCC CTTCTGGGAC ACCACCGGCG GACGGCTCAG TAACACGCCG G
16SBHB_4_6 GGCTCGAT CACCTCCTAT CGCCGGAAAC CCATCGGGG GGTTTAACCC CGGATGGGAT TTTGGTACTC CCCTTTATGG GGAGTCCCAC CGGCCGTACT CCTATAGCAA TTCCGGTTGA TCCTGCCGGA GGCCACTGCT ATGGGGGTCC GACTAAGCCA TGCGAGTCAT GGGGGTGTCC CTTCTGGGAC ACCACCGGCG GACGGCTCAG TAACACGTCG G
16SBHB_3_9 CGGCTCGAT CACCTCCTAT CGCCGGAAAC CCATCGGGG GGTTTAACCC CGGATGGGAT TTTGGTACTC CCCTTTATGG GGAGTCCCAC CGGCCGTACT CCTATAGCAA TTCCGGTTGA TCCTGCCGGA GGCCACTGCT ATGGGGGTCC GACTAAGCCA TGCGAGTCAT GGGGGTGTCC CTTCTGGGAC ACCACCGGCG GACGGCTCAG TAACACGCCG
16SBHB_7_3 GGGAACCT ACGGCTCGAT CACCTCCTAT CGCCGGAAAC CCATCGGGG GGTTTAACCC CGGATGGGAT TTTGGTACTC CCCTTTATGG GGAGTCCCAC CGGCCGTACT CCTATAGCAA TTCCGGTTGA TCCTGCCGGA GGCCACTGCT ATGGGGGTCC GACTAAGCCA TGCGAGTCAT GGGGGTGTCC CTTCTGGGAC ACCACCGGCG GACGGCTCAG
16SBHB_6_3 GGGGAACCT ACGGCTCGAT CACCTCCTAT CGCCGGAAAC CCATCGGGG GGTTTAACCC CGGATGGGAT TTTGGTACTC CCCTTTATGG GGAGTCCCAC CGGCCGTACT CCTATAGCAA TTCCGGTTGA TCCTGCCGGA GGCCACTGCT ATGGGGGTCC GACTAAGCCA TGCGAGTCAT GGGGGTGTCC CTTCTGGGAC ACCACCGGCG GACGGCTCAG
16SBHB_7_3 CCT ACGGCTCGAT CACCTCCTAT CGCCGGAAAC CCATCGGGG GGTTTAACCC CGGATGGGAT TTTGGTACTC CCCTTTATGG GGAGTCCCAC CGGCCGTACT CCTATAGCAA TTCCGGTTGA TCCTGCCGGA GGCCACTGCT ATGGGGGTCC GACTAAGCCA TGCGAGTCAT GGGGGTGTCC CTTCTGGGAC ACCACCGGCG GACGGCTCAG TAACAC
16SBHB_6_4 CCT ACGGCTCGAT CACCTCCTAT CGCCGGAAAC CCATCGGGG GGTTTAACCC CGGATGGGAT TTTGGTACTC CCCTTTATGG GGAGTCCCAC CGGCCGTACT CCTATAGCAA TTCCGGTTGA TCCTGCCGGA GGCCACTGCT ATGGGGGTCC GACTAAGCCA TGCGAGTCAT GGGGGTGTCC CTTCTGGGAC ACCACCGGCG GACGGCTCAG TAACACG
16SBHB_4_6 GGAACCT ACGGCTCGAT CACCTCCTAT CGCCGGAAAC CCATCGGGG GGTTTAACCC CGGATGGGAT TTTGGTACTC CCCTTTATGG GGAGTCCCAC CGGCCGTACT CCTATAGCAA TTCCGGTTGA TCCTGCCGGA GGCCACTGCT ATGGGGGTCC GACTAAGCCA TGCGAGTCAT GGGGGTGTCC CTTCTGGGAC ACCACCGGCG GACGGCTCAG
16SBHB_17_ CGT AGGGGAACCT ACGGCTCGAT CACCTCCTAT CGCCGGAAAC CCATCGGGG GGTTTAACCC CGGATGGGAT TTTGGTACTC CCCTTTATGG GGAGTCCCAC CGGCCGTACT CCTATAGCAA TTCCGGTTGA TCCTGCCGGA GGCCACTGCT ATGGGGGTCC GACTAAGCCA TGCGAGTCAT GGGGGTGTCC CTTCTGGGAC ACCACCGGCG GAC
16SBHB_6_4 CGT AGGGGAACCT ACGGCTCGAT CACCTCCTAT CGCCGGAAAC CCATCGGGG GGTTTAACCC CGGATGGGAT TTTGGTACTC CCCTTTATGG GGAGTCCCAC CGGCCGTACT CCTATAGCAA TTCCGGTTGA TCCTGCCGGA GGCCACTGCT ATGGGGGTCC GACTAAGCCA TGCGAGTCAT GGGGGTGTCC CTTCTGGGAC ACCACCGGCG GACGGC
16SBHB_4_6 CGT AGGGGAACCT ACGGCTCGAT CACCTCCTAT CGCCGGAAAC CCATCGGGG GGTTTAACCC CGGATGGGAT TTTGGTACTC CCCTTTATGG GGAGTCCCAC CGGCCGTACT CCTATAGCAA TTCCGGTTGA TCCTGCCGGA GGCCACTGCT ATGGGGGTCC GACTAAGCCA TGCGAGTCAT GGGGGTGTCC CTTCTGGGAC ACCACCGGCG GACG
16SBHB_6_3 GT AGGGGAACCT ACGGCTCGAT CACCTCCTAT CGCCGGAAAC CCATCGGGG GGTTTAACCC CGGATGGGAT TTTGGTACTC CCCTTTATGG GGAGTCCCAC CGGCCGTACT CCTATAGCAA TTCCGGTTGA TCCTGCCGGA GGCCACTGCT ATGGGGGTCC GACTAAGCCA TGCGAGTCAT GGGGGTGTCC CTTCTGGGAC ACCACCGGCG GAC
16SBHB_14_ CCT ACGGCTCGAT CACCTCCTAT CGCCGGAAAC CCATCGGGG GGTTTAACCC CGGATGGGAT TTTGGTACTC CCCTTTATGG GGAGTCCCAC CGGCCGTACT CCTATAGCAA TTCCGGTTGA TCCTGCCGGA GGCCACTGCT ATGGGGGTCC GACTAAGCCA TGCGAGTCAT GGGGGTGTCC CTTCTGGGAC ACCACCGGCG GACGGCTCAG TAAC
16SBHB_3_9 CT ACGGCTCGAT CACCTCCTAT CGCCGGAAAC CCATCGGGG GGTTTAACCC CGGATGGGAT TTTGGTACTC CCCTTTATGG GGAGTCCCAC CGGCCGTACT CCTATAGCAA TTCCGGTTGA TCCTGCCGGA GGCCACTGCT ATGGGGGTCC GACTAAGCCA TGCGAGTCAT GGGGGTGTCC CTTCTGGGAC ACCACCGGCG GACGGCTCAG TAAC
16SBHB_4_5 T ACGGCTCGAT CACCTCCTAT CGCCGGAAAC CCATCGGGG GGTTTAACCC CGGATGGGAT TTTGGTACTC CCCTTTATGG GGAGTCCCAC CGGCCGTACT CCTATAGCAA TTCCGGTTGA TCCTGCCGGA GGCCACTGCT ATGGGGGTCC GACTAAGCCA TGCGAGTCAT GGGGGTGTCC CTTCTGGGAC ACCACCGGCG GACGGCTCAG TAAC
16SBHB_5_4 CGGCTCGAT CACCTCCTAT CGCCGGAAAC CCATCGGGG GGTTTAACCC CGGATGGGAT TTTGGTACTC CCCTTTATGG GGAGTCCCAC CGGCCGTACT CCTATAGCAA TTCCGGTTGA TCCTGCCGGA GGCCACTGCT ATGGGGGTCC GACTAAGCCA TGCGAGTCAT GGGGGTGTCC CTTCTGGGAC ACCACCGGCG GACGGCTCAG TAACAC
16SBHB_4_6 CGGCTCGAT CACCTCCTAT CGCCGGAAAC CCATCGGGG GGTTTAACCC CGGATGGGAT TTTGGTACTC CCCTTTATGG GGAGTCCCAC CGGCCGTACT CCTATAGCAA TTCCGGTTGA TCCTGCCGGA GGCCACTGCT ATGGGGGTCC GACTAAGCCA TGCGAGTCAT GGGGGTGTCC CTTCTGGGAC ACCACCGGCG GACGGCTCAG TAACACG
16SBHB_27_ CGAT CACCTCCTAT CGCCGGAAAC CCATCGGGG GGTTTAACCC CGGATGGGAT TTTGGTACTC CCCTTTATGG GGAGTCCCAC CGGCCGTACT CCTATAGCAA TTCCGGTTGA TCCTGCCGGA GGCCACTGCT ATGGGGGTCC GACTAAGCCA TGCGAGTCAT GGGGGTGTCC CTTCTGGGAC ACCACCGGCG GACGGCTCAG TAACACGTCG
16SBHB_4_6 CGAT CACCTCCTAT CGCCGGAAAC CCATCGGGG GGTTTAACCC CGGATGGGAT TTTGGTACTC CCCTTTATGG GGAGTCCCAC CGGCCGTACT CCTATAGCAA TTCCGGTTGA TCCTGCCGGA GGCCACTGCT ATGGGGGTCC GACTAAGCCA TGCGAGTCAT GGGGGTGTCC CTTCTGGGAC ACCACCGGCG GACGGCTCAG TAACACGCCG
16SBHB_23_ CGT AGGGGAACCT ACGGCTCGAT CACCTCCTAT CGCCGGAAAC CCATCGGGG GGTTTAACCC CGGATGGGAT TTTGGTACTC CCCTTTATGG GGAGTCCCAC CGGCCGTACT CCTATAGCAA TTCCGGTTGA TCCTGCCGGA GGCCACTGCT ATGGGGGTCC GACTAAGCCA TGCGAGTCAT GGGGGTGTCC CTTCTGGGAC ACCACCGGCG
16SBHB_13_ CGT AGGGGAACCT ACGGCTCGAT CACCTCCTAT CGCCGGAAAC CCATCGGGG GGTTTAACCC CGGATGGGAT TTTGGTACTC CCCTTTATGG GGAGTCCCAC CGGCCGTACT CCTATAGCAA TTCCGGTTGA TCCTGCCGGA GGCCACTGCT ATGGGGGTCC GACTAAGCCA TGCGAGTCAT GGGGGTGTCC CTTCTGGGAC ACCACCGGCG G
16SBHB_4_6 GGTAGCCGT AGGGGAACCT ACGGCTCGAT CACCTCCTAT CGCCGGAAAC CCATCGGGG GGTTTAACCC CGGATGGGAT TTTGGTACTC CCCTTTATGG GGAGTCCCAC CGGCCGTACT CCTATAGCAA TTCCGGTTGA TCCTGCCGGA GGCCACTGCT ATGGGGGTCC GACTAAGCCA TGCGAGTCAT GGGGGTGTCC CTTCTGGGAC ACCACCGGCG GAC
16SBHB_4_7 A AGGTAGCCGT AGGGGAACCT ACGGCTCGAT CACCTCCTAT CGCCGGAAAC CCATCGGGG GGTTTAACCC CGGATGGGAT TTTGGTACTC CCCTTTATGG GGAGTCCCAC CGGCCGTACT CCTATAGCAA TTCCGGTTGA TCCTGCCGGA GGCCACTGCT ATGGGGGTCC GACTAAGCCA TGCGAGTCAT GGGGGTGTCC CTTCTGGGAC ACCACCGGCG G
16SBHB_3_8 GTAGCCGT AGGGGAACCT ACGGCTCGAT CACCTCCTAT CGCCGGAAAC CCATCGGGG GGTTTAACCC CGGATGGGAT TTTGGTACTC CCCTTTATGG GGAGTCCCAC CGGCCGTACT CCTATAGCAA TTCCGGTTGA TCCTGCCGGA GGCCACTGCT ATGGGGGTCC GACTAAGCCA TGCGAGTCAT GGGGGTGTCC CTTCTGGGAC ACCACCGGCG
16SBHB_5_5 CCGT AGGGGAACCT ACGGCTCGAT CACCTCCTAT CGCCGGAAAC CCATCGGGG GGTTTAACCC CGGATGGGAT TTTGGTACTC CCCTTTATGG GGAGTCCCAC CGGCCGTACT CCTATAGCAA TTCCGGTTGA TCCTGCCGGA GGCCACTGCT ATGGGGGTCC GACTAAGCCA TGCGAGTCAT GGGGGTGTCC CTTCTGGGAC ACCACCGGCG
16SBHB_17_ CCT ACGGCTCGAT CACCTCCTAT CGCCGGAAAC CCATCGGGG GGTTTAACCC CGGATGGGAT TTTGGTACTC CCCTTTATGG GGAGTCCCAC CGGCCGTACT CCTATAGCAA TTCCGGTTGA TCCTGCCGGA GGCCACTGCT ATGGGGGTCC GACTAAGCCA TGCGAGTCAT GGGGGTGTCC CTTCTGGGAC ACCACCGGCG GACGGCTCAG
16SBHB_3_9 CT ACGGCTCGAT CACCTCCTAT CGCCGGAAAC CCATCGGGG GGTTTAACCC CGGATGGGAT TTTGGTACTC CCCTTTATGG GGAGTCCCAC CGGCCGTACT CCTATAGCAA TTCCGGTTGA TCCTGCCGGA GGCCACTGCT ATGGGGGTCC GACTAAGCCA TGCGAGTCAT GGGGGTGTCC CTTCTGGGAC ACCACCGGCG GACGGCTCAG
16SBHB_16_ CTCCTAT CGCCGGAAAC CCATCGGGG GGTTTAACCC CGGATGGGAT TTTGGTACTC CCCTTTATGG GGAGTCCCAC CGGCCGTACT CCTATAGCAA TTCCGGTTGA TCCTGCCGGA GGCCACTGCT ATGGGGGTCC GACTAAGCCA TGCGAGTCAT GGGGGTGTCC CTTCTGGGAC ACCACCGGCG GACGGCTCAG TAACACGTCG GTAACCTACC CTCGGG
16SBHB_10_ CACCTCCTAT CGCCGGAAAC CCATCGGGG GGTTTAACCC CGGATGGGAT TTTGGTACTC CCCTTTATGG GGAGTCCCAC CGGCCGTACT CCTATAGCAA TTCCGGTTGA TCCTGCCGGA GGCCACTGCT ATGGGGGTCC GACTAAGCCA TGCGAGTCAT GGGGGTGTCC CTTCTGGGAC ACCACCGGCG GACGGCTCAG TAACACGTCG GTAACCTACC CTCGGG
16SBHB_6_4 AT CACCTCCTAT CGCCGGAAAC CCATCGGGG GGTTTAACCC CGGATGGGAT TTTGGTACTC CCCTTTATGG GGAGTCCCAC CGGCCGTACT CCTATAGCAA TTCCGGTTGA TCCTGCCGGA GGCCACTGCT ATGGGGGTCC GACTAAGCCA TGCGAGTCAT GGGGGTGTCC CTTCTGGGAC ACCACCGGCG GACGGCTCAG TAACACGTCG GTAACCTACC CTCGGG
16SBHB_8_3 CACCTCCTAT CGCCGGAAAC CCATCGGGG GGTTTAACCC CGGATGGGAT TTTGGTACTC CCCTTTATGG GGAGTCCCAC CGGCCGTACT CCTATAGCAA TTCCGGTTGA TCCTGCCGGA GGCCACTGCT ATGGGGGTCC GACTAAGCCA TGCGAGTCAT GGGGGTGTCC CTTCTGGGAC ACCACCGGCG GACGGCTCAG TAACACGTCG GTAACCTACC CTCGG
16SBHB_7_3 CTCCTAT CGCCGGAAAC CCATCGGGG GGTTTAACCC CGGATGGGAT TTTGGTACTC CCCTTTATGG GGAGTCCCAC CGGCCGTACT CCTATAGCAA TTCCGGTTGA TCCTGCCGGA GGCCACTGCT ATGGGGGTCC GACTAAGCCA TGCGAGTCAT GGGGGTGTCC CTTCTGGGAC ACCACCGGCG GACGGCTCAG TAACACGTCG GTAACCTACC CTCGG
16SBHB_15_ CCTAT CGCCGGAAAC CCATCGGGG GGTTTAACCC CGGATGGGAT TTTGGTACTC CCCTTTATGG GGAGTCCCAC CGGCCGTACT CCTATAGCAA TTCCGGTTGA TCCTGCCGGA GGCCACTGCT ATGGGGGTCC GACTAAGCCA TGCGAGTCAT GGGGGTGTCC CTTCTGGGAC ACCACCGGCG GACGGCTCAG TAACACGTCG GTAACCTACC CTCGGG
16SBHB_6_4 CTCCTAT CGCCGGAAAC CCATCGGGG GGTTTAACCC CGGATGGGAT TTTGGTACTC CCCTTTATGG GGAGTCCCAC CGGCCGTACT CCTATAGCAA TTCCGGTTGA TCCTGCCGGA GGCCACTGCT ATGGGGGTCC GACTAAGCCA TGCGAGTCAT GGGGGTGTCC CTTCTGGGAC ACCACCGGCG GACGGCTCAG TAACACGTCG GTAACCTACC CTCGGGAGGG G
16SBHB_3_7 TAT CGCCGGAAAC CCATCGGGG GGTTTAACCC CGGATGGGAT TTTGGTACTC CCCTTTATGG GGAGTCCCAC CGGCCGTACT CCTATAGCAA TTCCGGTTGA TCCTGCCGGA GGCCACTGCT ATGGGGGTCC GACTAAGCCA TGCGAGTCAT GGGGGTGTCC CTTCTGGGAC ACCACCGGCG GACGGCTCAG TAACACGTCG GTAACCTACC CTCGGGAGGG G
16SBHB_7_3 CTAT CGCCGGAAAC CCATCGGGG GGTTTAACCC CGGATGGGAT TTTGGTACTC CCCTTTATGG GGAGTCCCAC CGGCCGTACT CCTATAGCAA TTCCGGTTGA TCCTGCCGGA GGCCACTGCT ATGGGGGTCC GACTAAGCCA TGCGAGTCAT GGGGGTGTCC CTTCTGGGAC ACCACCGGCG GACGGCTCAG TAACACGTCG GTAACCTACC CTCGGG
16SBHB_3_9 CTCCTAT CGCCGGAAAC CCATCGGGG GGTTTAACCC CGGATGGGAT TTTGGTACTC CCCTTTATGG GGAGTCCCAC CGGCCGTACT CCTATAGCAA TTCCGGTTGA TCCTGCCGGA GGCCACTGCT ATGGGGGTCC GACTAAGCCA TGCGAGTCAT GGGGGTGTCC CTTCTGGGAC ACCACCGGCG GACGGCTCAG TAACACGTCG GTAACCTACC C
16SBHB_6_4 CCTAT CGCCGGAAAC CCATCGGGG GGTTTAACCC CGGATGGGAT TTTGGTACTC CCCTTTATGG GGAGTCCCAC CGGCCGTACT CCTATAGCAA TTCCGGTTGA TCCTGCCGGA GGCCACTGCT ATGGGGGTCC GACTAAGCCA TGCGAGTCAT GGGGGTGTCC CTTCTGGGAC ACCACCGGCG GACGGCTCAG TAACACGTCG GTAACCTACC C
16SBHB_3_9 CTCCTAT CGCCGGAAAC CCATCGGGG GGTTTAACCC CGGATGGGAT TTTGGTACTC CCCTTTATGG GGAGTCCCAC CGGCCGTACT CCTATAGCAA TTCCGGTTGA TCCTGCCGGA GGCCACTGCT ATGGGGGTCC GACTAAGCCA TGCGAGTCAT GGGGGTGTCC CTTCTGGGAC ACCACCGGCG GACGGCTCAG TAACACGTCG GTAACCTAC
16SBHB_4_6 CTCCTAT CGCCGGAAAC CCATCGGGG GGTTTAACCC CGGATGGGAT TTTGGTACTC CCCTTTATGG GGAGTCCCAC CGGCCGTACT CCTATAGCAA TTCCGGTTGA TCCTGCCGGA GGCCACTGCT ATGGGGGTCC GACTAAGCCA TGCGAGTCAT GGGGGTGTCC CTTCTGGGAC ACCACCGGCG GACGGCTCAG TAACACGTCG GTAAC
16SBHB_7_3 CGGCTCGAT CACCTCCTAT CGCCGGAAAC CCATCGGGG GGTTTAACCC CGGATGGGAT TTTGGTACTC CCCTTTATGG GGAGTCCCAC CGGCCGTACT CCTATAGCAA TTCCGGTTGA TCCTGCCGGA GGCCACTGCT ATGGGGGTCC GACTAAGCCA TGCGAGTCAT GGGGGTGTCC CTTCTGGGAC ACCACCGGCG GACGGCTCAG TAAC
16SBHB_3_8 GGCTCGAT CACCTCCTAT CGCCGGAAAC CCATCGGGG GGTTTAACCC CGGATGGGAT TTTGGTACTC CCCTTTATGG GGAGTCCCAC CGGCCGTACT CCTATAGCAA TTCCGGTTGA TCCTGCCGGA GGCCACTGCT ATGGGGGTCC GACTAAGCCA TGCGAGTCAT GGGGGTGTCC CTTCTGGGAC ACCGCCGGCG GACGGCTCAG TAACA
16SBHB_4_6 GGAACCT ACGGCTCGAT CACCTCCTAT CGCCGGAAAC CCATCGGGG GGTTTAACCC CGGATGGGAT TTTGGTACTC CCCTTTATGG GGAGTCCCAC CGGCCGTACT CCTATAGCAA TTCCGGTTGA TCCTGCCGGA GGCCACTGCT ATGGGGGTCC GACTAAGCCA TGCGAGTCAT GGGGGTGTCC CTTCTGGGAC ACCACCGGCG GACGGC
16SBHB_4_6 CT ACGGCTCGAT CACCTCCTAT CGCCGGAAAC CCATCGGGG GGTTTAACCC CGGATGGGAT TTTGGTACTC CCCTTTATGG GGAGTCCCAC CGGCCGTACT CCTATAGCAA TTCCGGTTGA TCCTGCCGGA GGCCACTGCT ATGGGGGTCC GACTAAGCCA TGCGAGTCAT GGGGGTGTCC CTTCTGGGAC ACCACCGGCG GACGGCTC
16SBHB_13_ CACCTCCTAT CGCCGGAAAC CCATCGGGG GGTTTAACCC CGGATGGGAT TTTGGTACTC CCCTTTATGG GGAGTCCCAC CGGCCGTACT CCTATAGCAA TTCCGGTTGA TCCTGCCGGA GGCCACTGCT ATGGGGGTCC GACTAAGCCA TGCGAGTCAT GGGGGTGTCC CTTCTGGGAC ACCACCGGCG GACGGCTCAG TAACACGTCG
16SBHB_9_2 AT CACCTCCTAT CGCCGGAAAC CCATCGGGG GGTTTAACCC CGGATGGGAT TTTGGTACTC CCCTTTATGG GGAGTCCCAC CGGCCGTACT CCTATAGCAA TTCCGGTTGA TCCTGCCGGA GGCCACTGCT ATGGGGGTCC GACTAAGCCA TGCGAGTCAT GGGGGTGTCC CTTCTGGGAC ACCACCGGCG GACGGCTCAG TAACACGTCG
16SBHB_8_3 CACCTCCTAT CGCCGGAAAC CCATCGGGG GGTTTAACCC CGGATGGGAT TTTGGTACTC CCCTTTATGG GGAGTCCCAC CGGCCGTACT CCTATAGCAA TTCCGGTTGA TCCTGCCGGA GGCCACTGCT ATGGGGGTCC GACTAAGCCA TGCGAGTCAT GGGGGTGTCC CTTCTGGGAC ACCACCGGCG GACGGCTCAG TAACACGTCG G
16SBHB_20_ GGGGAACCT ACGGCTCGAT CACCTCCTAT CGCCGGAAAC CCATCGGGG GGTTTAACCC CGGATGGGAT TTTGGTACTC CCCTTTATGG GGAGTCCCAC CGGCCGTACT CCTATAGCAA TTCCGGTTGA TCCTGCCGGA GGCCACTGCT ATGGGGGTCC GACTAAGCCA TGCGAGTCAT GGGGGTGTCC CTTCTGGGAC ACCACCGGCG
16SBHB_9_2 GGGGAACCT ACGGCTCGAT CACCTCCTAT CGCCGGAAAC CCATCGGGG GGTTTAACCC CGGATGGGAT TTTGGTACTC CCCTTTATGG GGAGTCCCAC CGGCCGTACT CCTATAGCAA TTCCGGTTGA TCCTGCCGGA GGCCACTGCT ATGGGGGTCC GACTAAGCCA TGCGAGTCAT GGGGGTGTCC CTTCTGGGAC ACCACCGGCG GAC
16SBHB_6_3 GGGAACCT ACGGCTCGAT CACCTCCTAT CGCCGGAAAC CCATCGGGG GGTTTAACCC CGGATGGGAT TTTGGTACTC CCCTTTATGG GGAGTCCCAC CGGCCGTACT CCTATAGCAA TTCCGGTTGA TCCTGCCGGA GGCCACTGCT ATGGGGGTCC GACTAAGCCA TGCGAGTCAT GGGGGTGTCC CTTCTGGGAC ACCACCGGCG GAC
16SBHB_5_4 GGGGAACCT ACGGCTCGAT CACCTCCTAT CGCCGGAAAC CCATCGGGG GGTTTAACCC CGGATGGGAT TTTGGTACTC CCCTTTATGG GGAGTCCCAC CGGCCGTACT CCTATAGCAA TTCCGGTTGA TCCTGCCGGA GGCCACTGCT ATGGGGGTCC GACTAAGCCA TGCGAGTCAT GGGGGTGTCC CTTCTGGGAC ACCACCGGCG G
16SBHB_5_4 GGAACCT ACGGCTCGAT CACCTCCTAT CGCCGGAAAC CCATCGGGG GGTTTAACCC CGGATGGGAT TTTGGTACTC CCCTTTATGG GGAGTCCCAC CGGCCGTACT CCTATAGCAA TTCCGGTTGA TCCTGCCGGA GGCCACTGCT ATGGGGGTCC GACTAAGCCA TGCGAGTCAT GGGGGTGTCC CTTCTGGGAC ACCACCGGCG GAC
16SBHB_11_ GGGAACCT ACGGCTCGAT CACCTCCTAT CGCCGGAAAC CCATCGGGG GGTTTAACCC CGGATGGGAT TTTGGTACTC CCCTTTATGG GGAGTCCCAC CGGCCGTACT CCTATAGCAA TTCCGGTTGA TCCTGCCGGA GGCCACTGCT ATGGGGGTCC GACTAAGCCA TGCGAGTCAT GGGGGTGTCC CTTCTGGGAC ACCACCGGCG
16SBHB_11_ GGAACCT ACGGCTCGAT CACCTCCTAT CGCCGGAAAC CCATCGGGG GGTTTAACCC CGGATGGGAT TTTGGTACTC CCCTTTATGG GGAGTCCCAC CGGCCGTACT CCTATAGCAA TTCCGGTTGA TCCTGCCGGA GGCCACTGCT ATGGGGGTCC GACTAAGCCA TGCGAGTCAT GGGGGTGTCC CTTCTGGGAC ACCACCGGCG G
16SBHB_3_9 GAACCT ACGGCTCGAT CACCTCCTAT CGCCGGAAAC CCATCGGGG GGTTTAACCC CGGATGGGAT TTTGGTACTC CCCTTTATGG GGAGTCCCAC CGGCCGTACT CCTATAGCAA TTCCGGTTGA TCCTGCCGGA GGCCACTGCT ATGGGGGTCC GACTAAGCCA TGCGAGTCAT GGGGGTGTCC CTTCTGGGAC ACCACCGGCG GAC
16SBHB_10_ CGAT CACCTCCTAT CGCCGGAAAC CCATCGGGG GGTTTAACCC CGGATGGGAT TTTGGTACTC CCCTTTATGG GGAGTCCCAC CGGCCGTACT CCTATAGCAA TTCCGGTTGA TCCTGCCGGA GGCCACTGCT ATGGGGGTCC GACTAAGCCA TGCGAGTCAT GGGGGTGTCC CTTCTGGGAC ACCACCGGCG GACGGCTCAG TAACAC
16SBHB_7_3 CGAT CACCTCCTAT CGCCGGAAAC CCATCGGGG GGTTTAACCC CGGATGGGAT TTTGGTACTC CCCTTTATGG GGAGTCCCAC CGGCCGTACT CCTATAGCAA TTCCGGTTGA TCCTGCCGGA GGCCACTGCT ATGGGGGTCC GACTAAGCCA TGCGAGTCAT GGGGGTGTCC CTTCTGGGAC ACCACCGGCG GACGGCTCAG TAAC
16SBHB_5_5 CGGCTCGAT CACCTCCTAT CGCCGGAAAC CCATCGGGG GGTTTAACCC CGGATGGGAT TTTGGTACTC CCCTTTATGG GGAGTCCCAC CGGCCGTACT CCTATAGCAA TTCCGGTTGA TCCTGCCGGA GGCCACTGCT ATGGGGGTCC GACTAAGCCA TGCGAGTCAT GGGGGTGTCC CTTCTGGGAC ACCACCGGCG GACGGCTCAG
16SBHB_8_3 CCT ACGGCTCGAT CACCTCCTAT CGCCGGAAAC CCATCGGGG GGTTTAACCC CGGATGGGAT TTTGGTACTC CCCTTTATGG GGAGTCCCAC CGGCCGTACT CCTATAGCAA TTCCGGTTGA TCCTGCCGGA GGCCACTGCT ATGGGGGTCC GACTAAGCCA TGCGAGTCAT GGGGGTGTCC CTTCTGGGAC ACCACCGGCG GACGGC
16SBHB_4_7 CCT ACGGCTCGAT CACCTCCTAT CGCCGGAAAC CCATCGGGG GGTTTAACCC CGGATGGGAT TTTGGTACTC CCCTTTATGG GGAGTCCCAC CGGCCGTACT CCTATAGCAA TTCCGGTTGA TCCTGCCGGA GGCCACTGCT ATGGGGGTCC GACTAAGCCA TGCGAGTCAT GGGGGTGTCC CTTCTGGGAC ACCACCGGCG GACG
16SBHB_4_6 CGT AGGGGAACCT ACGGCTCGAT CACCTCCTAT CGCCGGAAAC CCATCGGGG GGTTTAACCC CGGATGGGAT TTTGGTACTC CCCTTTATGG GGAGTCCCAC CGGCCGTACT CCTATAGCAA TTCCGGTTGA TCCTGCCGGA GGCCACTGCT ATGGGGGTCC GACTAAGCCA TGCGAGTCAT GGGGGTGTCC CTTCTGGGAC ACCACCGGC
16SBHB_3_9 CGT AGGGGAACCT ACGGCTCGAT CACCTCCTAT CGCCGGAAAC CCATCGGGG GGTTTAACCC CGGATGGGAT TTTGGTACTC CCCTTTATGG GGAGTCCCAC CGGCCGTACT CCTATAGCAA TTCCGGTTGA TCCTGCCGGA GGCCACTGCT ATGGGGGTCC GACTAAGCCA TGCGAGTCAT GGGGGTGTCC CTTCTGGGAC ACCACCG
16SBHB_9_2 GGAACCT ACGGCTCGAT CACCTCCTAT CGCCGGAAAC CCATCGGGG GGTTTAACCC CGGATGGGAT TTTGGTACTC CCCTTTATGG GGAGTCCCAC CGGCCGTACT CCTATAGCAA TTCCGGTTGA TCCTGCCGGA GGCCACTGCT ATGGGGGTCC GACTAAGCCA TGCGAGTCAT GGGGGTGTCC CTTCTGGGAC ACCACCGGCG
16SBHB_3_8 GGAACCT ACGGCTCGAT CACCTCCTAT CGCCGGAAAC CCATCGGGG GGTTTAACCC CGGATGGGAT TTTGGTACTC CCCTTTATGG GGAGTCCCAC CGGCCGTACT CCTATAGCAA TTCCGGTTGA TCCTGCCGGA GGCCACTGCT ATGGGGGTCC GACTAAGCCA TGCGAGTCAT GGGGGTGTCC CTTCTGGGAC ACCACCGGC
16SBHB_28_ CCT ACGGCTCGAT CACCTCCTAT CGCCGGAAAC CCATCGGGG GGTTTAACCC CGGATGGGAT TTTGGTACTC CCCTTTATGG GGAGTCCCAC CGGCCGTACT CCTATAGCAA TTCCGGTTGA TCCTGCCGGA GGCCACTGCT ATGGGGGTCC GACTAAGCCA TGCGAGTCAT GGGGGTGTCC CTTCTGGGAC ACCACCGGCG GAC
16SBHB_9_2 CT ACGGCTCGAT CACCTCCTAT CGCCGGAAAC CCATCGGGG GGTTTAACCC CGGATGGGAT TTTGGTACTC CCCTTTATGG GGAGTCCCAC CGGCCGTACT CCTATAGCAA TTCCGGTTGA TCCTGCCGGA GGCCACTGCT ATGGGGGTCC GACTAAGCCA TGCGAGTCAT GGGGGTGTCC CTTCTGGGAC ACCACCGGCG GAC
16SBHB_3_9 CT ACGGCTCGAT CACCTCCTAT CGCCGGAAAC CCATCGGGG GGTTTAACCC CGGATGGGAT TTTGGTACTC CCCTTTATGG GGAGTCCCAC CGGCCGTACT CCTATAGCAA TTCCGGTTGA TCCTGCCGGA GGCCACTGCT ATGGGGGTCC GACTAAGCCA TGCGAGTCAT GGGGGTGTCC CTTCTGGGAC ACCACCGGCG GACG
16SBHB_6_3 T CACCTCCTAT CGCCGGAAAC CCATCGGGG GGTTTAACCC CGGATGGGAT TTTGGTACTC CCCTTTATGG GGAGTCCCAC CGGCCGTACT CCTATAGCAA TTCCGGTTGA TCCTGCCGGA GGCCACTGCT ATGGGGGTCC GACTAAGCCA TGCGAGTCAT GGGGGTGTCC CTTCTGGGAC ACCACCGGCG GACGGCTCAG TAACAC
16SBHB_6_4 CACCTCCTAT CGCCGGAAAC CCATCGGGG GGTTTAACCC CGGATGGGAT TTTGGTACTC CCCTTTATGG GGAGTCCCAC CGGCCGTACT CCTATAGCAA TTCCGGTTGA TCCTGCCGGA GGCCACTGCT ATGGGGGTCC GACTAAGCCA TGCGAGTCAT GGGGGTGTCC CTTCTGGGAC ACCACCGGCG GACGGCTCAG TAACA
16SBHB_40_ CTCCTAT CGCCGGAAAC CCATCGGGG GGTTTAACCC CGGATGGGAT TTTGGTACTC CCCTTTATGG GGAGTCCCAC CGGCCGTACT CCTATAGCAA TTCCGGTTGA TCCTGCCGGA GGCCACTGCT ATGGGGGTCC GACTAAGCCA TGCGAGTCAT GGGGGTGTCC CTTCTGGGAC ACCACCGGCG GACGGCTCAG TAACACGTCG G
16SBHB_35_ CTCCTAT CGCCGGAAAC CCATCGGGG GGTTTAACCC CGGATGGGAT TTTGGTACTC CCCTTTATGG GGAGTCCCAC CGGCCGTACT CCTATAGCAA TTCCGGTTGA TCCTGCCGGA GGCCACTGCT ATGGGGGTCC GACTAAGCCA TGCGAGTCAT GGGGGTGTCC CTTCTGGGAC ACCACCGGCG GACGGCTCAG TAACACGTCG
16SBHB_3_9 CTCCTAT CGCCGGAAAC CCATCGGGG GGTTTAACCC CGGATGGGAT TTTGGTACTC CCCTTTATGG GGAGTCCCAC CGGCCGTACT CCTATAGCAA TTCCGGTTGA TCCTGCCGGA GGCCACTGCT ATGGGGGTCC GACTAAGCCA TGCGAGTCAT GGGGGTGTCC CTTCTGGGAC ACCACCGGCG GACGGCTCAG TAACACGCCG G
16SBHB_7_3 CCTAT CGCCGGAAAC CCATCGGGG GGTTTAACCC CGGATGGGAT TTTGGTACTC CCCTTTATGG GGAGTCCCAC CGGCCGTACT CCTATAGCAA TTCCGGTTGA TCCTGCCGGA GGCCACTGCT ATGGGGGTCC GACTAAGCCA TGCGAGTCAT GGGGGTGTCC CTTCTGGGAC ACCACCGGCG GACGGCTCAG TAACACGTCG G
16SBHB_17_ CCTAT CGCCGGAAAC CCATCGGGG GGTTTAACCC CGGATGGGAT TTTGGTACTC CCCTTTATGG GGAGTCCCAC CGGCCGTACT CCTATAGCAA TTCCGGTTGA TCCTGCCGGA GGCCACTGCT ATGGGGGTCC GACTAAGCCA TGCGAGTCAT GGGGGTGTCC CTTCTGGGAC ACCACCGGCG GACGGCTCAG TAACACGTCG
16SBHB_3_9 GAACCT ACGGCTCGAT CACCTCCTAT CGCCGGAAAC CCATCGGGG GGTTTAACCC CGGATGGGAT TTTGGTACTC CCCTTTATGG GGAGTCCCAC CGGCCGTACT CCTATAGCAA TTCCGGTTGA TCCTGCCGGA GGCCACTGCT ATGGGGGTCC GACTAAGCCA TGCGAGTCAT GGGGGTGTCC CTTCTGGGAC ACCACCGGC
16SBHB_12_ CCT ACGGCTCGAT CACCTCCTAT CGCCGGAAAC CCATCGGGG GGTTTAACCC CGGATGGGAT TTTGGTACTC CCCTTTATGG GGAGTCCCAC CGGCCGTACT CCTATAGCAA TTCCGGTTGA TCCTGCCGGA GGCCACTGCT ATGGGGGTCC GACTAAGCCA TGCGAGTCAT GGGGGTGTCC CTTCTGGGAC ACCACCGGCG G
16SBHB_4_7 ACCT ACGGCTCGAT CACCTCCTAT CGCCGGAAAC CCATCGGGG GGTTTAACCC CGGATGGGAT TTTGGTACTC CCCTTTATGG GGAGTCCCAC CGGCCGTACT CCTATAGCAA TTCCGGTTGA TCCTGCCGGA GGCCACTGCT ATGGGGGTCC GACTAAGCCA TGCGAGTCAT GGGGGTGTCC CTTCTGGGAC ACCACCGGCG G
16SBHB_8_2 CGAT CACCTCCTAT CGCCGGAAAC CCATCGGGG GGTTTAACCC CGGATGGGAT TTTGGTACTC CCCTTTATGG GGAGTCCCAC CGGCCGTACT CCTATAGCAA TTCCGGTTGA TCCTGCCGGA GGCCACTGCT ATGGGGGTCC GACTAAGCCA TGCGAGTCAT GGGGGTGTCC CTTCTGGGAC ACCACCGGCG GACGGCTCAG
16SBHB_3_7 T ACGGCTCGAT CACCTCCTAT CGCCGGAAAC CCATCGGGG GGTTTAACCC CGGATGGGAT TTTGGTACTC CCCTTTATGG GGAGTCCCAC CGGCCGTACT CCTATAGCAA TTCCGGTTGA TCCTGCCGGA GGCCACTGCT ATGGGGGTCC GACTAAGCCA TGCGAGTCAT GGGGGTGTCC CTTCTGGGAC ACCACCGGCG GAC
16SBHB_3_8 GGGAACCT ACGGCTCGAT CACCTCCTAT CGCCGGAAAC CCATCGGGG GGTTTAACCC CGGATGGGAT TTTGGTACTC CCCTTTATGG GGAGTCCCAC CGGCCGTACT CCTATAGCAA TTCCGGTTGA TCCTGCCGGA GGCCACTGCT ATGGGGGTCC GACTAAGCCA TGCGAGTCAT GGGGGTGTCC CTTCTGGGAC ACCACCG
16SBHB_5_5 AT CACCTCCTAT CGCCGGAAAC CCATCGGGG GGTTTAACCC CGGATGGGAT TTTGGTACTC CCCTTTATGG GGAGTCCCAC CGGCCGTACT CCTATAGCAA TTCCGGTTGA TCCTGCCGGA GGCCACTGCT ATGGGGGTCC GACTAAGCCA TGCGAGTCAT GGGGGTGTCC CTTCTGGGAC ACCACCGGCG GACGGCTCAG
16SBHB_23_ CCT ACGGCTCGAT CACCTCCTAT CGCCGGAAAC CCATCGGGG GGTTTAACCC CGGATGGGAT TTTGGTACTC CCCTTTATGG GGAGTCCCAC CGGCCGTACT CCTATAGCAA TTCCGGTTGA TCCTGCCGGA GGCCACTGCT ATGGGGGTCC GACTAAGCCA TGCGAGTCAT GGGGGTGTCC CTTCTGGGAC ACCACCGGCG
16SBHB_3_9 CT ACGGCTCGAT CACCTCCTAT CGCCGGAAAC CCATCGGGG GGTTTAACCC CGGATGGGAT TTTGGTACTC CCCTTTATGG GGAGTCCCAC CGGCCGTACT CCTATAGCAA TTCCGGTTGA TCCTGCCGGA GGCCACTGCT ATGGGGGTCC GACTAAGCCA TGCGAGTCAT GGGGGTGTCC CTTCTGGGAC ACCACCGGCG
16SBHB_9_2 CT ACGGCTCGAT CACCTCCTAT CGCCGGAAAC CCATCGGGG GGTTTAACCC CGGATGGGAT TTTGGTACTC CCCTTTATGG GGAGTCCCAC CGGCCGTACT CCTATAGCAA TTCCGGTTGA TCCTGCCGGA GGCCACTGCT ATGGGGGTCC GACTAAGCCA TGCGAGTCAT GGGGGTGTCC CTTCTGGGAC ACCACCGGCG G
16SBHB_18_ CTCCTAT CGCCGGAAAC CCATCGGGG GGTTTAACCC CGGATGGGAT TTTGGTACTC CCCTTTATGG GGAGTCCCAC CGGCCGTACT CCTATAGCAA TTCCGGTTGA TCCTGCCGGA GGCCACTGCT ATGGGGGTCC GACTAAGCCA TGCGAGTCAT GGGGGTGTCC CTTCTGGGAC ACCACCGGCG GACGGCTCAG TAACAC
16SBHB_13_ CTCCTAT CGCCGGAAAC CCATCGGGG GGTTTAACCC CGGATGGGAT TTTGGTACTC CCCTTTATGG GGAGTCCCAC CGGCCGTACT CCTATAGCAA TTCCGGTTGA TCCTGCCGGA GGCCACTGCT ATGGGGGTCC GACTAAGCCA TGCGAGTCAT GGGGGTGTCC CTTCTGGGAC ACCACCGGCG GACGGCTCAG TAAC
16SBHB_11_ CACCTCCTAT CGCCGGAAAC CCATCGGGG GGTTTAACCC CGGATGGGAT TTTGGTACTC CCCTTTATGG GGAGTCCCAC CGGCCGTACT CCTATAGCAA TTCCGGTTGA TCCTGCCGGA GGCCACTGCT ATGGGGGTCC GACTAAGCCA TGCGAGTCAT GGGGGTGTCC CTTCTGGGAC ACCACCGGCG GACGGCTCAG
16SBHB_9_2 GGCTCGAT CACCTCCTAT CGCCGGAAAC CCATCGGGG GGTTTAACCC CGGATGGGAT TTTGGTACTC CCCTTTATGG GGAGTCCCAC CGGCCGTACT CCTATAGCAA TTCCGGTTGA TCCTGCCGGA GGCCACTGCT ATGGGGGTCC GACTAAGCCA TGCGAGTCAT GGGGGTGTCC CTTCTGGGAC ACCACCGGCG GAC
16SBHB_5_5 CGGCTCGAT CACCTCCTAT CGCCGGAAAC CCATCGGGG GGTTTAACCC CGGATGGGAT TTTGGTACTC CCCTTTATGG GGAGTCCCAC CGGCCGTACT CCTATAGCAA TTCCGGTTGA TCCTGCCGGA GGCCACTGCT ATGGGGGTCC GACTAAGCCA TGCGAGTCAT GGGGGTGTCC CTTCTGGGAC ACCACCGGCG GAC
16SBHB_4_5 TAT CGCCGGAAAC CCATCGGGG GGTTTAACCC CGGATGGGAT TTTGGTACTC CCCTTTATGG GGAGTCCCAC CGGCCGTACT CCTATAGCAA TTCCGGTTGA TCCTGCCGGA GGCCACTGCT ATGGGGGTCC GACTAAGCCA TGCGAGTCAT GGGGGTGTCC CTTCTGGGAC ACCACCGGCG GACGGCTCAG TAACACGTCG
16SBHB_3_8 T ACGGCTCGAT CACCTCCTAT CGCCGGAAAC CCATCGGGG GGTTTAACCC CGGATGGGAT TTTGGTACTC CCCTTTATGG GGAGTCCCAC CGGCCGTACT CCTATAGCAA TTCCGGTTGA TCCTGCCGGA GGCCACTGCT ATGGGGGTCC GACTAAGCCA TGCGAGTCAT GGGGGTGTCC CTTCTGGGAC ACCACCGGCG
16SBHB_13_ CCTAT CGCCGGAAAC CCATCGGGG GGTTTAACCC CGGATGGGAT TTTGGTACTC CCCTTTATGG GGAGTCCCAC CGGCCGTACT CCTATAGCAA TTCCGGTTGA TCCTGCCGGA GGCCACTGCT ATGGGGGTCC GACTAAGCCA TGCGAGTCAT GGGGGTGTCC CTTCTGGGAC ACCACCGGCG GACGGCTCAG TAACAC
16SBHB_6_4 CCTAT CGCCGGAAAC CCATCGGGG GGTTTAACCC CGGATGGGAT TTTGGTACTC CCCTTTATGG GGAGTCCCAC CGGCCGTACT CCTATAGCAA TTCCGGTTGA TCCTGCCGGA GGCCACTGCT ATGGGGGTCC GACTAAGCCA TGCGAGTCAT GGGGGTGTCC CTTCTGGGAC ACCACCGGCG GACGGCTCAG TAAC
16SBHB_19_ CTCCTAT CGCCGGAAAC CCATCGGGG GGTTTAACCC CGGATGGGAT TTTGGTACTC CCCTTTATGG GGAGTCCCAC CGGCCGTACT CCTATAGCAA TTCCGGTTGA TCCTGCCGGA GGCCACTGCT ATGGGGGTCC GACTAAGCCA TGCGAGTCAT GGGGGTGTCC CTTCTGGGAC ACCACCGGCG GACGGCTCAG
16SBHB_10_ CGGCTCGAT CACCTCCTAT CGCCGGAAAC CCATCGGGG GGTTTAACCC CGGATGGGAT TTTGGTACTC CCCTTTATGG GGAGTCCCAC CGGCCGTACT CCTATAGCAA TTCCGGTTGA TCCTGCCGGA GGCCACTGCT ATGGGGGTCC GACTAAGCCA TGCGAGTCAT GGGGGTGTCC CTTCTGGGAC ACCACCGGCG
16SBHB_3_8 T ACGGCTCGAT CACCTCCTAT CGCCGGAAAC CCATCGGGG GGTTTAACCC CGGATGGGAT TTTGGTACTC CCCTTTATGG GGAGTCCCAC CGGCCGTACT CCTATAGCAA TTCCGGTTGA TCCTGCCGGA GGCCACTGCT ATGGGGGTCC GACTAAGCCA TGCGAGTCAT GGGGGTGTCC CTTCTGGGAC ACCACCGG
16SBHB_4_6 CGAT CACCTCCTAT CGCCGGAAAC CCATCGGGG GGTTTAACCC CGGATGGGAT TTTGGTACTC CCCTTTATGG GGAGTCCCAC CGGCCGTACT CCTATAGCAA TTCCGGTTGA TCCTGCCGGA GGCCACTGCT ATGGGGGTCC GACTAAGCCA TGCGAGTCAT GGGGGTGTCC CTTCTGGGAC ACCACCGGCG GAC
16SBHB_3_9 CTCGAT CACCTCCTAT CGCCGGAAAC CCATCGGGG GGTTTAACCC CGGATGGGAT TTTGGTACTC CCCTTTATGG GGAGTCCCAC CGGCCGTACT CCTATAGCAA TTCCGGTTGA TCCTGCCGGA GGCCACTGCT ATGGGGGTCC GACTAAGCCA TGCGAGTCAT GGGGGTGTCC CTTCTGGGAC ACCACCGGCG G
16SBHB_4_6 CTCCTAT CGCCGGAAAC CCATCGGGG GGTTTAACCC CGGATGGGAT TTTGGTACTC CCCTTTATGG GGAGTCCCAC CGGCCGTACT CCTATAGCAA TTCCGGTTGA TCCTGCCGGA GGCCACTGCT ATGGGGGTCC GACTAAGCCA TGCGAGTCAT GGGGGTGTCC CTTCTGGGAC ACCACCGGCG GACGGCTC
16SBHB_4_7 CACCTCCTAT CGCCGGAAAC CCATCGGGG GGTTTAACCC CGGATGGGAT TTTGGTACTC CCCTTTATGG GGAGTCCCAC CGGCCGTACT CCTATAGCAA TTCCGGTTGA TCCTGCCGGA GGCCACTGCT ATGGGGGTCC GACTAAGCCA TGCGAGTCAT GGGGGTGTCC CTTCTGGGAC ACCACCGGCG GACGG
16SBHB_4_7 CCTAT CGCCGGAAAC CCATCGGGG GGTTTAACCC CGGATGGGAT TTTGGTACTC CCCTTTATGG GGAGTCCCAC CGGCCGTACT CCTATAGCAA TTCCGGTTGA TCCTGCCGGA GGCCACTGCT ATGGGGGTCC GACTAAGCCA TGCGAGTCAT GGGGGTGTCC CTTCTGGGAC ACCACCGGCG GACGGCTCAG
16SBHB_4_6 CTAT CGCCGGAAAC CCATCGGGG GGTTTAACCC CGGATGGGAT TTTGGTACTC CCCTTTATGG GGAGTCCCAC CGGCCGTACT CCTATAGCAA TTCCGGTTGA TCCTGCCGGA GGCCACTGCT ATGGGGGTCC GACTAAGCCA TGCGAGTCAT GGGGGTGTCC CTTCTGGGAC ACCACCGGCG GACGGCTCAG
16SBHB_8_2 CGAT CACCTCCTAT CGCCGGAAAC CCATCGGGG GGTTTAACCC CGGATGGGAT TTTGGTACTC CCCTTTATGG GGAGTCCCAC CGGCCGTACT CCTATAGCAA TTCCGGTTGA TCCTGCCGGA GGCCACTGCT ATGGGGGTCC GACTAAGCCA TGCGAGTCAT GGGGGTGTCC CTTCTGGGAC ACCACCGGCG G
16SBHB_15_ CACCTCCTAT CGCCGGAAAC CCATCGGGG GGTTTAACCC CGGATGGGAT TTTGGTACTC CCCTTTATGG GGAGTCCCAC CGGCCGTACT CCTATAGCAA TTCCGGTTGA TCCTGCCGGA GGCCACTGCT ATGGGGGTCC GACTAAGCCA TGCGAGTCAT GGGGGTGTCC CTTCTGGGAC ACCACCGGCG GAC
16SBHB_13_ CGAT CACCTCCTAT CGCCGGAAAC CCATCGGGG GGTTTAACCC CGGATGGGAT TTTGGTACTC CCCTTTATGG GGAGTCCCAC CGGCCGTACT CCTATAGCAA TTCCGGTTGA TCCTGCCGGA GGCCACTGCT ATGGGGGTCC GACTAAGCCA TGCGAGTCAT GGGGGTGTCC CTTCTGGGAC ACCACCGGCG
16SBHB_10_ CTCCTAT CGCCGGAAAC CCATCGGGG GGTTTAACCC CGGATGGGAT TTTGGTACTC CCCTTTATGG GGAGTCCCAC CGGCCGTACT CCTATAGCAA TTCCGGTTGA TCCTGCCGGA GGCCACTGCT ATGGGGGTCC GACTAAGCCA TGCGAGTCAT GGGGGTGTCC CTTCTGGGAC ACCACCGGCG GACGGC
16SBHB_5_5 CGAT CACCTCCTAT CGCCGGAAAC CCATCGGGG GGTTTAACCC CGGATGGGAT TTTGGTACTC CCCTTTATGG GGAGTCCCAC CGGCCGTACT CCTATAGCAA TTCCGGTTGA TCCTGCCGGA GGCCACTGCT ATGGGGGTCC GACTAAGCCA TGCGAGTCAT GGGGGTGTCC CTTCTGGGAC ACCACCGG
16SBHB_7_3 CACCTCCTAT CGCCGGAAAC CCATCGGGG GGTTTAACCC CGGATGGGAT TTTGGTACTC CCCTTTATGG GGAGTCCCAC CGGCCGTACT CCTATAGCAA TTCCGGTTGA TCCTGCCGGA GGCCACTGCT ATGGGGGTCC GACTAAGCCA TGCGAGTCAT GGGGGTGTCC CTTCTGGGAC ACCACCGGCG G
16SBHB_5_4 CTCCTAT CGCCGGAAAC CCATCGGGG GGTTTAACCC CGGATGGGAT TTTGGTACTC CCCTTTATGG GGAGTCCCAC CGGCCGTACT CCTATAGCAA TTCCGGTTGA TCCTGCCGGA GGCCACTGCT ATGGGGGTCC GACTAAGCCA TGCGAGTCAT GGGGGTGTCC CTTCTGGGAC ACCACCGGCG GACG
16SBHB_21_ CTCCTAT CGCCGGAAAC CCATCGGGG GGTTTAACCC CGGATGGGAT TTTGGTACTC CCCTTTATGG GGAGTCCCAC CGGCCGTACT CCTATAGCAA TTCCGGTTGA TCCTGCCGGA GGCCACTGCT ATGGGGGTCC GACTAAGCCA TGCGAGTCAT GGGGGTGTCC CTTCTGGGAC ACCACCGGCG GAC
16SBHB_10_ CACCTCCTAT CGCCGGAAAC CCATCGGGG GGTTTAACCC CGGATGGGAT TTTGGTACTC CCCTTTATGG GGAGTCCCAC CGGCCGTACT CCTATAGCAA TTCCGGTTGA TCCTGCCGGA GGCCACTGCT ATGGGGGTCC GACTAAGCCA TGCGAGTCAT GGGGGTGTCC CTTCTGGGAC ACCACCGGCG
16SBHB_4_7 CCTCCTAT CGCCGGAAAC CCATCGGGG GGTTTAACCC CGGATGGGAT TTTGGTACTC CCCTTTATGG GGAGTCCCAC CGGCCGTACT CCTATAGCAA TTCCGGTTGA TCCTGCCGGA GGCCACTGCT ATGGGGGTCC GACTAAGCCA TGCGAGTCAT GGGGGTGTCC CTTCTGGGAC ACCACCGGCG G
16SBHB_14_ CCTAT CGCCGGAAAC CCATCGGGG GGTTTAACCC CGGATGGGAT TTTGGTACTC CCCTTTATGG GGAGTCCCAC CGGCCGTACT CCTATAGCAA TTCCGGTTGA TCCTGCCGGA GGCCACTGCT ATGGGGGTCC GACTAAGCCA TGCGAGTCAT GGGGGTGTCC CTTCTGGGAC ACCACCGGCG GAC
16SBHB_13_ CTCCTAT CGCCGGAAAC CCATCGGGG GGTTTAACCC CGGATGGGAT TTTGGTACTC CCCTTTATGG GGAGTCCCAC CGGCCGTACT CCTATAGCAA TTCCGGTTGA TCCTGCCGGA GGCCACTGCT ATGGGGGTCC GACTAAGCCA TGCGAGTCAT GGGGGTGTCC CTTCTGGGAC ACCACCGGCG G
16SBHB_5_5 CCTCCTAT CGCCGGAAAC CCATCGGGG GGTTTAACCC CGGATGGGAT TTTGGTACTC CCCTTTATGG GGAGTCCCAC CGGCCGTACT CCTATAGCAA TTCCGGTTGA TCCTGCCGGA GGCCACTGCT ATGGGGGTCC GACTAAGCCA TGCGAGTCAT GGGGGTGTCC CTTCTGGGAC ACCACCGGCG
16SBHB_20_ CTCCTAT CGCCGGAAAC CCATCGGGG GGTTTAACCC CGGATGGGAT TTTGGTACTC CCCTTTATGG GGAGTCCCAC CGGCCGTACT CCTATAGCAA TTCCGGTTGA TCCTGCCGGA GGCCACTGCT ATGGGGGTCC GACTAAGCCA TGCGAGTCAT GGGGGTGTCC CTTCTGGGAC ACCACCGGCG
16SBHB_4_6 CTAT CGCCGGAAAC CCATCGGGG GGTTTAACCC CGGATGGGAT TTTGGTACTC CCCTTTATGG GGAGTCCCAC CGGCCGTACT CCTATAGCAA TTCCGGTTGA TCCTGCCGGA GGCCACTGCT ATGGGGGTCC GACTAAGCCA TGCGAGTCAT GGGGGTGTCC CTTCTGGGAC ACCACCGGCG GAC
16SBHB_3_7 TAT CGCCGGAAAC CCATCGGGG GGTTTAACCC CGGATGGGAT TTTGGTACTC CCCTTTATGG GGAGTCCCAC CGGCCGTACT CCTATAGCAA TTCCGGTTGA TCCTGCCGGA GGCCACTGCT ATGGGGGTCC GACTAAGCCA TGCGAGTCAT GGGGGTGTCC CTTCTGGGAC ACCACCGGCG GAC
16SBHB_3_9 CTCCTAT CGCCGGAAAC CCATCGGGG GGTTTAACCC CGGATGGGAT TTTGGTACTC CCCTTTATGG GGAGTCCCAC CGGCCGTACT CCTATAGCAA TTCCGGTTGA TCCTGCCGGA GGCCACTGCT ATGGGGGTCC GACTAAGCCA TGCGAGTCAT GGGGGTGTCC CTTCTGGGAC ACCACCGGC
16SBHB_7_3 CCTAT CGCCGGAAAC CCATCGGGG GGTTTAACCC CGGATGGGAT TTTGGTACTC CCCTTTATGG GGAGTCCCAC CGGCCGTACT CCTATAGCAA TTCCGGTTGA TCCTGCCGGA GGCCACTGCT ATGGGGGTCC GACTAAGCCA TGCGAGTCAT GGGGGTGTCC CTTCTGGGAC ACCACCGGCG G
16SBHB_12_ CCTAT CGCCGGAAAC CCATCGGGG GGTTTAACCC CGGATGGGAT TTTGGTACTC CCCTTTATGG GGAGTCCCAC CGGCCGTACT CCTATAGCAA TTCCGGTTGA TCCTGCCGGA GGCCACTGCT ATGGGGGTCC GACTAAGCCA TGCGAGTCAT GGGGGTGTCC CTTCTGGGAC ACCACCGGCG
16SBHB_5_5 CCTAT CGCCGGAAAC CCATCGGGG GGTTTAACCC CGGATGGGAT TTTGGTACTC CCCTTTATGG GGAGTCCCAC CGGCCGTACT CCTATAGCAA TTCCGGTTGA TCCTGCCGGA GGCCACTGCT ATGGGGGTCC GACTAAGCCA TGCGAGTCAT GGGGGTGTCC CTTCTGGGAC ACCACCGGC
16SBHB_3_9 CTCCTAT CGCCGGAAAC CCATCGGGG GGTTTAACCC CGGATGGGAT TTTGGTACTC CCCTTTATGG GGAGTCCCAC CGGCCGTACT CC-ATAGCAA TTCCGGTTGA TCCTGCCGGA GGCCACTGCT ATGGGGGTCC GACTAAGCCA TGCGAGTCAT GGGGGTGTCC CTTCTGGGAC ACCACCGGCG
16SBHB_10_ GGGGAACCT ACGGCTCGAT CACCTCCTAT CGCCGGAAAC CCATCGGGG GGTTTAACCC CGGATGGGAT TTTGGTACTC CCCTTTATGG GGAGTCCCAC CGGCCGTACT CCTATAGCAA TTCCGGTTGA TCCTGCCGGA GGCCACTGCT ATGGGGGTCC GACTAAGCCA TGCGAGTCAT GGGG
16SBHB_3_9 CGT AGGGGAACCT ACGGCTCGAT CACCTCCTAT CGCCGGAAAC CCATCGGGG GGTTTAACCC CGGATGGGAT TTTGGTACTC CCCTTTATGG GGAGTCCCAC CGGCCGTACT CCTATAGCAA TTCCGGTTGA TCCTGCCGGA GGCCACTGCT ATGGGGGTCC GACTAAGCCA TGCGAGTCAT GGG
```

```
16SBHB_47_ GTCGTAACA AGGTAGCCGT AGGGGAACCT ACGGCTCGAT CACCTCCTAT CGCCGGAAAC CCATCCGGGG GGTTTAACCC CGGATGGGAT TTTGGTACTC CCCTTTATGG GGAGTCCCAC CGGCCGTACT CCTATAGCAA TTCCGGTTGA TCCTGCCGGA GGCCACTGCT ATGGGGGTCC GACTAAG
16SBHB_27_ GTCGTAACA AGGTAGCCGT AGGGGAACCT ACGGCTCGAT CACCTCCTAT CGCCGGAAAC CCATCCGGGG GGTTTAACCC CGGATGGGAT TTTGGTACTC CCCTTTATGG GGAGTCCCAC CGGCCGTACT CCTATAGCAA TTCCGGTTGA TCCTGCCGGA GGCCACTGCT ATGGGGGTCC GACTAAGC
16SBHB_25_ GTCGTAACA AGGTAGCCGT AGGGGAACCT ACGGCTCGAT CACCTCCTAT CGCCGGAAAC CCATCCGGGG GGTTTAACCC CGGATGGGAT TTTGGTACTC CCCTTTATGG GGAGTCCCAC CGGCCGTACT CCTATAGCAA TTCCGGTTGA TCCTGCCGGA GGCCACTGCT ATGGGGGTCC GACTAAGCC
16SBHB_16_ GTCGTAACA AGGTAGCCGT AGGGGAACCT ACGGCTCGAT CACCTCCTAT CGCCGGAAAC CCATCCGGGG GGTTTAACCC CGGATGGGAT TTTGGTACTC CCCTTTATGG GGAGTCCCAC CGGCCGTACT CCTATAGCAA TTCCGGTTGA TCCTGCCGGA GGCCACTGCT ATGGGGGTCC GACTAAGCCA TGCG
16SBHB_11_ GTCGTAACA AGGTAGCCGT AGGGGAACCT ACGGCTCGAT CACCTCCTAT CGCCGGAAAC CCATCCGGGG GGTTTAACCC CGGATGGGAT TTTGGTACTC CCCTTTATGG GGAGTCCCAC CGGCCGTACT CCTATAGCAA TTCCGGTTGA TCCTGCCGGA GGCCACTGCT ATGGGGGTCC GACTAAGCCA TGC
16SBHB_6_3 GTCGTAACA AGGTAGCCGT AGGGGAACCT ACGGCTCGAT CACCTCCTAT CGCCGGAAAC CCATCCGGGG GGTTTAACCC CGGATGGGAT TTTGGTACTC CCCTTTATGG GGAGTCCCAC CGGCCGTACT CCTATAGCAA TTCCGGTTGA TCCTGCCGGA GGCCACTGCT ATGGGGGTCC GACTAAGCCA TGCGAG
16SBHB_3_8 GTCGTAACA AGGTAGCCGT AGGGGAACCT ACGGCTCGAT CACCTCCTAT CGCCGGAAAC CCATCCGGGG GGTTTAACCC CGGATGGGAT TTTGGTACTC CCCTTTATGG GGAGTCCCAC CGGCCGTACT CCTATAGCAA TTCCGGTTGA TCCTGCCGGA GGCCACTGCT ATGGGGGTCC GACTAAGCCA TGCGAGTCAT GGG
16SBHB_3_9 CGTAACA AGGTAGCCGT AGGGGAACCT ACGGCTCGAT CACCTCCTAT CGCCGGAAAC CCATCCGGGG GGTTTAACCC CGGATGGGAT TTTGGTACTC CCCTTTATGG GGAGTCCCAC CGGCCGTACT CCTATAGCAA TTCCGGTTGA TCCTGCCGGA GGCCACTGCT ATGGGGGTCC GACTAAGCCA TGCGAGTCAT GGGGGTGTCC CTTCTGGG
16SBHB_7_3 GTCC GGGCTCCGTG AGGGGGGAGA AGTCGTAACA AGGTAGCCGT AGGGGAACCT ACGGCTCGAT CACCTCCTAT CGCCGGAAAC CCATCCGGGG GGTTTAACCC CGGATGGGAT TTTGGTACTC CCCTTTATGG GGAGTCCCAC CGGCCGTACT CCTATAGCAA TTCCGGTTGA TCCTGCCGGA GGCCACTGCT ATGGGGGTCC GACTAAGCCA TGCG
16SBHB_4_6 GG GGTCGAGTCC GGGCTCCGTG AGGGGGGAGA AGTCGTAACA AGGTAGCCGT AGGGGAACCT ACGGCTCGAT CACCTCCTAT CGCCGGAAAC CCATCCGGGG GGTTTAACCC CGGATGGGAT TTTGGTACTC CCCTTTATGG GGAGTCCCAC CGGCCGTACT CCTATAGCAA TTCCGGTTGA TCCTGCCGGA GGCCACTGCT ATGGGGGTCC GACTAAGCCA TGCGAGTCAT GGGG
16SBHB_4_6 CGAGTCC GGGCTCCGTG AGGGGGGAGA AGTCGTAACA AGGTAGCCGT AGGGGAACCT ACGGCTCGAT CACCTCCTAT CGCCGGAAAC CCATCCGGGG GGTTTAACCC CGGATGGGAT TTTGGTACTC CCCTTTATGG GGAGTCCCAC CGGCCGTACT CCTATAGCAA TTCCGGTTGA TCCTGCCGGA GGCCACTGCT ATGGGGGTCC GACTAAGCCA TGCGAGTCAT GGG
16SBHB_3_8 GTCC GGGCTCCGTG AGGGGGGAGA AGTCGTAACA AGGTAGCCGT AGGGGAACCT ACGGCTCGAT CACCTCCTAT CGCCGGAAAC CCATCCGGGG GGTTTAACCC CGGATGGGAT TTTGGTACTC CCCTTTATGG GGAGTCCCAC CGGCCGTACT CCTATAGCAA TTCCGGTTGA TCCTGCCGGA GGCCACTGCT ATGGGGGTCC GACTAAGCCA TGCGAGTCAT GGGG
16SBHB_3_8 GTCGAGTCC GGGCTCCGTG AGGGGGGAGA AGTCGTAACA AGGTAGCCGT AGGGGAACCT ACGGCTCGAT CACCTCCTAT CGCCGGAAAC CCATCCGGGG GGTTTAACCC CGGATGGGAT TTTGGTACTC CCCTTTATGG GGAGTCCCAC CGGCCGTACT CCTATAGCAA TTCCGGTTGA TCCTGCCGGA GGCCACTGCT ATGGGGGTCC GACTAAGCCA TG
16SBHB_7_3 CC GGGCTCCGTG AGGGGGGAGA AGTCGTAACA AGGTAGCCGT AGGGGAACCT ACGGCTCGAT CACCTCCTAT CGCCGGAAAC CCATCCGGGG GGTTTAACCC CGGATGGGAT TTTGGTACTC CCCTTTATGG GGAGTCCCAC CGGCCGTACT CCTATAGCAA TTCCGGTTGA TCCTGCCGGA GGCCACTGCT ATGGGGGTCC GACTAAGCCA TGCG
16SBHB_6_3 GTCGAGTCC GGGCTCCGTG AGGGGGGAGA AGTCGTAACA AGGTAGCCGT AGGGGAACCT ACGGCTCGAT CACCTCCTAT CGCCGGAAAC CCATCCGGGG GGTTTAACCC CGGATGGGAT TTTGGTACTC CCCTTTATGG GGAGTCCCAC CGGCCGTACT CCTATAGCAA TTCCGGTTGA TCCTGCCGGA GGCCACTGCT ATGGGGGTCC GACTAAGC
16SBHB_4_6 CGAGTCC GGGCTCCGTG AGGGGGGAGA AGTCGTAACA AGGTAGCCGT AGGGGAACCT ACGGCTCGAT CACCTCCTAT CGCCGGAAAC CCATCCGGGG GGTTTAACCC CGGATGGGAT TTTGGTACTC CCCTTTATGG GGAGTCCCAC CGGCCGTACT CCTATAGCAA TTCCGGTTGA TCCTGCCGGA GGCCACTGCT ATGGGGGTCC GACTAAGC
16SBHB_4_6 CGAGTCC GGGCTCCGTG AGGGGGGAGA AGTCGTAACA AGGTAGCCGT AGGGGAACCT ACGGCTCGAT CACCTCCTAT CGCCGGAAAC CCATCCGGGG GGTTTAACCC CGGATGGGAT TTTGGTACTC CCCTTTATGG GGAGTCCCAC CGGCCGTACT CCTATAGCAA TTCCGGTTGA TCCTGCCGGA GGCCACTGCT ATGGGGGTCC GACTAAG
16SBHB_3_8 GTCC GGGCTCCGTG AGGGGGGAGA AGTCGTAACA AGGTAGCCGT AGGGGAACCT ACGGCTCGAT CACCTCCTAT CGCCGGAAAC CCATCCGGGG GGTTTAACCC CGGATGGGAT TTTGGTACTC CCCTTTATGG GGAGTCCCAC CGGCCGTACT CCTATAGCAA TTCCGGTTGA TCCTGCCGGA GGCCACTGCT ATGGGGGTCC GACTAAGC
16SBHB_4_5 GTCC GGGCTCCGTG AGGGGGGAGA AGTCGTAACA AGGTAGCCGT AGGGGAACCT ACGGCTCGAT CACCTCCTAT CGCCGGAAAC CCATCCGGGG GGTTTAACCC CGGATGGGAT TTTGGTACTC CCCTTTATGG GGAGTCCCAC CGGCCGTACT CCTATAGCAA TTCCGGTTGA TCCTGCCGGA GGCCACTGCT ATGGGGGTCC GACTAAG
16SBHB_5_5 CC GGGCTCCGTG AGGGGGGAGA AGTCGTAACA AGGTAGCCGT AGGGGAACCT ACGGCTCGAT CACCTCCTAT CGCCGGAAAC CCATCCGGGG GGTTTAACCC CGGATGGGAT TTTGGTACTC CCCTTTATGG GGAGTCCCAC CGGCCGTACT CCTATAGCAA TTCCGGTTGA TCCTGCCGGA GGCCACTGCT ATGGGGGTCC GACTAAGC
16SBHB_5_5 CC GGGCTCCGTG AGGGGGGAGA AGTCGTAACA AGGTAGCCGT AGGGGAACCT ACGGCTCGAT CACCTCCTAT CGCCGGAAAC CCATCCGGGG GGTTTAACCC CGGATGGGAT TTTGGTACTC CCCTTTATGG GGAGTCCCAC CGGCCGTACT CCTATAGCAA TTCCGGTTGA TCCTGCCGGA GGCCACTGCT ATGGGGGTCC GACTAAG
16SBHB_5_4 GGCTCCGTG AGGGGGGAGA AGTCGTAACA AGGTAGCCGT AGGGGAACCT ACGGCTCGAT CACCTCCTAT CGCCGGAAAC CCATCCGGGG GGTTTAACCC CGGATGGGAT TTTGGTACTC CCCTTTATGG GGAGTCCCAC CGGCCGTACT CCTATAGCAA TTCCGGTTGA TCCTGCCGGA GGCCACTGCT ATGGGGGTCC GACTAAG
16SBHB_3_7 TG AGGGGGGAGA AGTCGTAACA AGGTAGCCGT AGGGGAACCT ACGGCTCGAT CACCTCCTAT CGCCGGAAAC CCATCCGGGG GGTTTAACCC CGGATGGGAT TTTGGTACTC CCCTTTATGG GGAGTCCCAC CGGCCGTACT CCTATAGCAA TTCCGGTTGA TCCTGCCGGA GGCCACTGCT ATGGGGGTCC GACTAAGCCA TGAC
16SBHB_6_4 CCGTG AGGGGGGAGA AGTCGTAACA AGGTAGCCGT AGGGGAACCT ACGGCTCGAT CACCTCCTAT CGCCGGAAAC CCATCCGGGG GGTTTAACCC CGGATGGGAT TTTGGTACTC CCCTTTATGG GGAGTCCCAC CGGCCGTACT CCTATAGCAA TTCCGGTTGA TCCTGCCGGA GGCCACTGCT ATGGGGGTCC GACTAAGC
16SBHB_3_8 GTG AGGGGGGAGA AGTCGTAACA AGGTAGCCGT AGGGGAACCT ACGGCTCGAT CACCTCCTAT CGCCGGAAAC CCATCCGGGG GGTTTAACCC CGGATGGGAT TTTGGTACTC CCCTTTATGG GGAGTCCCAC CGGCCGTACT CCTATAGCAA TTCCGGTTGA TCCTGCCGGA GGCCACTGCT ATGGGGGTCC GACTAAGC
16SBHB_5_4 GTG AGGGGGGAGA AGTCGTAACA AGGTAGCCGT AGGGGAACCT ACGGCTCGAT CACCTCCTAT CGCCGGAAAC CCATCCGGGG GGTTTAACCC CGGATGGGAT TTTGGTACTC CCCTTTATGG GGAGTCCCAC CGGCCGTACT CCTATAGCAA TTCCGGTTGA TCCTGCCGGA GGCCACTGCT ATGGGGGTCC GACTAAG
16SBHB_4_6 GGGGGGAGA AGTCGTAACA AGGTAGCCGT AGGGGAACCT ACGGCTCGAT CACCTCCTAT CGCCGGAAAC CCATCCGGGG GGTTTAACCC CGGATGGGAT TTTGGTACTC CCCTTTATGG GGAGTCCCAC CGGCCGTACT CCTATAGCAA TTCCGGTTGA TCCTGCCGGA GGCCACTGCT ATGGGGGTCC GACTAAGCCG
16SBHB_4_6 GGAGA AGTCGTAACA AGGTAGCCGT AGGGGAACCT ACGGCTCGAT CACCTCCTAT CGCCGGAAAC CCATCCGGGG GGTTTAACCC CGGATGGGAT TTTGGTACTC CCCTTTATGG GGAGTCCCAC CGGCCGTACT CCTATAGCAA TTCCGGTTGA TCCTGCCGGA GGCCACTGCT ATGGGGGTCC GACTAAGCCA T
16SBHB_3_8 GGGAGA AGTCGTAACA AGGTAGCCGT AGGGGAACCT ACGGCTCGAT CACCTCCTAT CGCCGGAAAC CCATCCGGGG GGTTTAACCC CGGATGGGAT TTTGGTACTC CCCTTTATGG GGAGTCCCAC CGGCCGTACT CCTATAGCAA TTCCGGTTGA TCCTGCCGGA GGCCACTGCT ATGGGGGTCC GACTAAGCCA
16SBHB_3_8 GGGGAGA AGTCGTAACA AGGTAGCCGT AGGGGAACCT ACGGCTCGAT CACCTCCTAT CGCCGGAAAC CCATCCGGGG GGTTTAACCC CGGATGGGAT TTTGGTACTC CCCTTTATGG GGAGTCCCAC CGGCCGTACT CCTATAGCAA TTCCGGTTGA TCCTGCCGGA GGCCACTGCT ATGGGGGTCC GACTAAG
16SBHB_4_7 AGA AGTCGTAACA AGGTAGCCGT AGGGGAACCT ACGGCTCGAT CACCTCCTAT CGCCGGAAAC CCATCCGGGG GGTTTAACCC CGGATGGGAT TTTGGTACTC CCCTTTATGG GGAGTCCCAC CGGCCGTACT CCTATAGCAA TTCCGGTTGA TCCTGCCGGA GGCCACTGCT ATGGGGGTCC GACTAAGCCA T
16SBHB_12_ GTCGTAACA AGGTAGCCGT AGGGGAACCT ACGGCTCGAT CACCTCCTAT CGCCGGAAAC CCATCCGGGG GGTTTAACCC CGGATGGGAT TTTGGTACTC CCCTTTATGG GGAGTCCCAC CGGCCGTACT CCTATAGCAA TTCCGGTTGA TCCTGCCGGA GGCCACTGCT ATGGGGGTCC GACTAAGCCA TG
16SBHB_5_4 GTCGTAACA AGGTAGCCGT AGGGGAACCT ACGGCTCGAT CACCTCCTAT CGCCGGAAAC CCATCCGGGG GGTTTAACCC CGGATGGGAT TTTGGTACTC CCCTTTATGG GGAGTCCCAC CGGCCGTACT CCTATAGCAA TTCCGGTTGA TCCTGCCGGA GGCCACTGCT ATGGGGGTCC GACTAAGCCA T
16SBHB_6_4 AGA AGTCGTAACA AGGTAGCCGT AGGGGAACCT ACGGCTCGAT CACCTCCTAT CGCCGGAAAC CCATCCGGGG GGTTTAACCC CGGATGGGAT TTTGGTACTC CCCTTTATGG GGAGTCCCAC CGGCCGTACT CCTATAGCAA TTCCGGTTGA TCCTGCCGGA GGCCACTGCT ATGGGGGTCC GACTAAGC
16SBHB_4_7 AGA AGTCGTAACA AGGTAGCCGT AGGGGAACCT ACGGCTCGAT CACCTCCTAT CGCCGGAAAC CCATCCGGGG GGTTTAACCC CGGATGGGAT TTTGGTACTC CCCTTTATGG GGAGTCCCAC CGGCCGTACT CCTATAGCAA TTCCGGTTGA TCCTGCCGGA GGCCACTGCT ATGGGGGTCC GACTAAG
16SBHB_7_3 GTCGTAACA AGGTAGCCGT AGGGGAACCT ACGGCTCGAT CACCTCCTAT CGCCGGAAAC CCATCCGGGG GGTTTAACCC CGGATGGGAT TTTGGTACTC CCCTTTATGG GGAGTCCCAC CGGCCGTACT CCTATAGCAA TTCCGGTTGA TCCTGCCGGA GGCCACTGCT ATGGGGGTCC GACTAAGCCA
16SBHB_3_8 GTAACA AGGTAGCCGT AGGGGAACCT ACGGCTCGAT CACCTCCTAT CGCCGGAAAC CCATCCGGGG GGTTTAACCC CGGATGGGAT TTTGGTACTC CCCTTTATGG GGAGTCCCAC CGGCCGTACT CCTATAGCAA TTCCGGTTGA TCCTGCCGGA GGCCACTGCT ATGGGGGTCC GACTAAGCCA TG
16SBHB_3_8 GTAACA AGGTAGCCGT AGGGGAACCT ACGGCTCGAT CACCTCCTAT CGCCGGAAAC CCATCCGGGG GGTTTAACCC CGGATGGGAT TTTGGTACTC CCCTTTATGG GGAGTCCCAC CGGCCGTACT CCTATAGCAA TTCCGGTTGA TCCTGCCGGA GGCCACTGCT ATGGGGGTCC GACTAAGCCA AG
16SBHB_4_6 CGTAACA AGGTAGCCGT AGGGGAACCT ACGGCTCGAT CACCTCCTAT CGCCGGAAAC CCATCCGGGG GGTTTAACCC CGGATGGGAT TTTGGTACTC CCCTTTATGG GGAGTCCCAC CGGCCGTACT CCTATAGCAA TTCCGGTTGA TCCTGCCGGA GGCCACTGCT ATGGGGGTCC GACTAAGCC
16SBHB_4_7 AACA AGGTAGCCGT AGGGGAACCT ACGGCTCGAT CACCTCCTAT CGCCGGAAAC CCATCCGGGG GGTTTAACCC CGGATGGGAT TTTGGTACTC CCCTTTATGG GGAGTCCCAC CGGCCGTACT CCTATAGCAA TTCCGGTTGA TCCTGCCGGA GGCCACTGCT ATGGGGGTCC GACTAAGCCA TGCG
16SBHB_4_5 GTAACA AGGTAGCCGT AGGGGAACCT ACGGCTCGAT CACCTCCTAT CGCCGGAAAC CCATCCGGGG GGTTTAACCC CGGATGGGAT TTTGGTACTC CCCTTTATGG GGAGTCCCAC CGGCCGTACT CCTATAGCAA TTCCGGTTGA TCCTGCCGGA GGCCACTGCT ATGGGGGTCC GACTAAGCC
16SBHB_3_9 CGTAACA AGGTAGCCGT AGGGGAACCT ACGGCTCGAT CACCTCCTAT CGCCGGAAAC CCATCCGGGG GGTTTAACCC CGGATGGGAT TTTGGTACTC CCCTTTATGG GGAGTCCCAC CGGCCGTACT CCTATAGCAA TTCCGGTTGA TCCTGCCGGA GGCCACTGCT ATGGGGGTCC GACTAAGC
16SBHB_5_5 CA AGGTAGCCGT AGGGGAACCT ACGGCTCGAT CACCTCCTAT CGCCGGAAAC CCATCCGGGG GGTTTAACCC CGGATGGGAT TTTGGTACTC CCCTTTATGG GGAGTCCCAC CGGCCGTACT CCTATAGCAA TTCCGGTTGA TCCTGCCGGA GGCCACTGCT ATGGGGGTCC GACTAAGCCA TGCG
16SBHB_3_9 CGTAACA AGGTAGCCGT AGGGGAACCT ACGGCTCGAT CACCTCCTAT CGCCGGAAAC CCATCCGGGG GGTTTAACCC CGGATGGGAT TTTGGTACTC CCCTTTATGG GGAGTCCCAC CGGCCGTACT CCTATAGCAA TTCCGGTTGA TCCTGCCGGA GGCCACTGCT ATGGGGGTCC GACTAAG
16SBHB_3_8 GTAACA AGGTAGCCGT AGGGGAACCT ACGGCTCGAT CACCTCCTAT CGCCGGAAAC CCATCCGGGG GGTTTAACCC CGGATGGGAT TTTGGTACTC CCCTTTATGG GGAGTCCCAC CGGCCGTACT CCTATAGCAA TTCCGGTTGA TCCTGCCGGA GGCCACTGCT ATGGGGGTCC GACTAAGC
16SBHB_5_4 TAACA AGGTAGCCGT AGGGGAACCT ACGGCTCGAT CACCTCCTAT CGCCGGAAAC CCATCCGGGG GGTTTAACCC CGGATGGGAT TTTGGTACTC CCCTTTATGG GGAGTCCCAC CGGCCGTACT CCTATAGCAA TTCCGGTTGA TCCTGCCGGA GGCCACTGCT ATGGGGGTCC GACTAAGCC
16SBHB_5_5 AACA AGGTAGCCGT AGGGGAACCT ACGGCTCGAT CACCTCCTAT CGCCGGAAAC CCATCCGGGG GGTTTAACCC CGGATGGGAT TTTGGTACTC CCCTTTATGG GGAGTCCCAC CGGCCGTACT CCTATAGCAA TTCCGGTTGA TCCTGCCGGA GGCCACTGCT ATGGGGGTCC GACTAAGCC
16SBHB_4_6 GTAACA AGGTAGCCGT AGGGGAACCT ACGGCTCGAT CACCTCCTAT CGCCGGAAAC CCATCCGGGG GGTTTAACCC CGGATGGGAT TTTGGTACTC CCCTTTATGG GGAGTCCCAC CGGCCGTACT CCTATAGCAA TTCCGGTTGA TCCTGCCGGA GGCCACTGCT ATGGGGGTCC GACTAAG
16SBHB_4_5 TAACA AGGTAGCCGT AGGGGAACCT ACGGCTCGAT CACCTCCTAT CGCCGGAAAC CCATCCGGGG GGTTTAACCC CGGATGGGAT TTTGGTACTC CCCTTTATGG GGAGTCCCAC CGGCCGTACT CCTATAGCAA TTCCGGTTGA TCCTGCCGGA GGCCACTGCT ATGGGGGTCC GACTAAG
16SBHB_3_8 GGTAGCCGT AGGGGAACCT ACGGCTCGAT CACCTCCTAT CGCCGGAAAC CCATCCGGGG GGTTTAACCC CGGATGGGAT TTTGGTACTC CCCTTTATGG GGAGTCCCAC CGGCCGTACT CCTATAGCAA TTCCGGTTGA TCCTGCCGGA GGCCACTGCT ATGGGGGTCC GACTAAGCCA TGCG
16SBHB_6_4 AACA AGGTAGCCGT AGGGGAACCT ACGGCTCGAT CACCTCCTAT CGCCGGAAAC CCATCCGGGG GGTTTAACCC CGGATGGGAT TTTGGTACTC CCCTTTATGG GGAGTCCCAC CGGCCGTACT CCTATAGCAA TTCCGGTTGA TCCTGCCGGA GGCCACTGCT ATGGGGGTCC GACTAAG
16SBHB_5_4 GGTAGCCGT AGGGGAACCT ACGGCTCGAT CACCTCCTAT CGCCGGAAAC CCATCCGGGG GGTTTAACCC CGGATGGGAT TTTGGTACTC CCCTTTATGG GGAGTCCCAC CGGCCGTACT CCTATAGCAA TTCCGGTTGA TCCTGCCGGA GGCCACTGCT ATGGGGGTCC GACTAAGCCA TG
16SBHB_4_7 CA AGGTAGCCGT AGGGGAACCT ACGGCTCGAT CACCTCCTAT CGCCGGAAAC CCATCCGGGG GGTTTAACCC CGGATGGGAT TTTGGTACTC CCCTTTATGG GGAGTCCCAC CGGCCGTACT CCTATAGCAA TTCCGGTTGA TCCTGCCGGA GGCCACTGCT ATGGGGGTCC GACTAAG
16SBHB_8_2 GGTAGCCGT AGGGGAACCT ACGGCTCGAT CACCTCCTAT CGCCGGAAAC CCATCCGGGG GGTTTAACCC CGGATGGGAT TTTGGTACTC CCCTTTATGG GGAGTCCCAC CGGCCGTACT CCTATAGCAA TTCCGGTTGA TCCTGCCGGA GGCCACTGCT ATGGGGGTCC GACTAAGCCA T
16SBHB_5_4 GTAGCCGT AGGGGAACCT ACGGCTCGAT CACCTCCTAT CGCCGGAAAC CCATCCGGGG GGTTTAACCC CGGATGGGAT TTTGGTACTC CCCTTTATGG GGAGTCCCAC CGGCCGTACT CCTATAGCAA TTCCGGTTGA TCCTGCCGGA GGCCACTGCT ATGGGGGTCC GACTAAGCCA TG
16SBHB_5_4 GGTAGCCGT AGGGGAACCT ACGGCTCGAT CACCTCCTAT CGCCGGAAAC CCATCCGGGG GGTTTAACCC CGGATGGGAT TTTGGTACTC CCCTTTATGG GGAGTCCCAC CGGCCGTACT CCTATAGCAA TTCCGGTTGA TCCTGCCGGA GGCCACTGCT ATGGGGGTCC GACTAAGCC
16SBHB_4_7 A AGGTAGCCGT AGGGGAACCT ACGGCTCGAT CACCTCCTAT CGCCGGAAAC CCATCCGGGG GGTTTAACCC CGGATGGGAT TTTGGTACTC CCCTTTATGG GGAGTCCCAC CGGCCGTACT CCTATAGCAA TTCCGGTTGA TCCTGCCGGA GGCCACTGCT ATGGGGGTCC GACTAAG
16SBHB_41_ CGT AGGGGAACCT ACGGCTCGAT CACCTCCTAT CGCCGGAAAC CCATCCGGGG GGTTTAACCC CGGATGGGAT TTTGGTACTC CCCTTTATGG GGAGTCCCAC CGGCCGTACT CCTATAGCAA TTCCGGTTGA TCCTGCCGGA GGCCACTGCT ATGGGGGTCC GACTAAGCCA TGCG
16SBHB_6_4 CGT AGGGGAACCT ACGGCTCGAT CACCTCCTAT CGCCGGAAAC CCATCCGGGG GGTTTAACCC CGGATGGGAT TTTGGTACTC CCCTTTATGG GGAGTCCCAC CGGCCGTACT CCTATAGCAA TTCCGGTTGA TCCTGCCGGA GGCCACTGCT ATGGGGGTCC GACTAAGCCA TGCGAG
16SBHB_13_ CCGT AGGGGAACCT ACGGCTCGAT CACCTCCTAT CGCCGGAAAC CCATCCGGGG GGTTTAACCC CGGATGGGAT TTTGGTACTC CCCTTTATGG GGAGTCCCAC CGGCCGTACT CCTATAGCAA TTCCGGTTGA TCCTGCCGGA GGCCACTGCT ATGGGGGTCC GACTAAGCCA TGCG
16SBHB_5_5 CCGT AGGGGAACCT ACGGCTCGAT CACCTCCTAT CGCCGGAAAC CCATCCGGGG GGTTTAACCC CGGATGGGAT TTTGGTACTC CCCTTTATGG GGAGTCCCAC CGGCCGTACT CCTATAGCAA TTCCGGTTGA TCCTGCCGGA GGCCACTGCT ATGGGGGTCC GACTAAGCCA TGC
16SBHB_16_ CGT AGGGGAACCT ACGGCTCGAT CACCTCCTAT CGCCGGAAAC CCATCCGGGG GGTTTAACCC CGGATGGGAT TTTGGTACTC CCCTTTATGG GGAGTCCCAC CGGCCGTACT CCTATAGCAA TTCCGGTTGA TCCTGCCGGA GGCCACTGCT ATGGGGGTCC GACTAAGCCA TGC
16SBHB_27_ CGT AGGGGAACCT ACGGCTCGAT CACCTCCTAT CGCCGGAAAC CCATCCGGGG GGTTTAACCC CGGATGGGAT TTTGGTACTC CCCTTTATGG GGAGTCCCAC CGGCCGTACT CCTATAGCAA TTCCGGTTGA TCCTGCCGGA GGCCACTGCT ATGGGGGTCC GACTAAGCCA TG
16SBHB_4_5 GT AGGGGAACCT ACGGCTCGAT CACCTCCTAT CGCCGGAAAC CCATCCGGGG GGTTTAACCC CGGATGGGAT TTTGGTACTC CCCTTTATGG GGAGTCCCAC CGGCCGTACT CCTATAGCAA TTCCGGTTGA TCCTGCCGGA GGCCACTGCT ATGGGGGTCC GACTAAGCCA TGC
16SBHB_5_5 CCGT AGGGGAACCT ACGGCTCGAT CACCTCCTAT CGCCGGAAAC CCATCCGGGG GGTTTAACCC CGGATGGGAT TTTGGTACTC CCCTTTATGG GGAGTCCCAC CGGCCGTACT CCTATAGCAA TTCCGGTTGA TCCTGCCGGA GGCCACTGCT ATGGGGGTCC GACTAAGCCA TG
16SBHB_26_ CGT AGGGGAACCT ACGGCTCGAT CACCTCCTAT CGCCGGAAAC CCATCCGGGG GGTTTAACCC CGGATGGGAT TTTGGTACTC CCCTTTATGG GGAGTCCCAC CGGCCGTACT CCTATAGCAA TTCCGGTTGA TCCTGCCGGA GGCCACTGCT ATGGGGGTCC GACTAAGCCA T
16SBHB_5_5 AGGGGAACCT ACGGCTCGAT CACCTCCTAT CGCCGGAAAC CCATCCGGGG GGTTTAACCC CGGATGGGAT TTTGGTACTC CCCTTTATGG GGAGTCCCAC CGGCCGTACT CCTATAGCAA TTCCGGTTGA TCCTGCCGGA GGCCACTGCT ATGGGGGTCC GACTAAGCCA TGCG
16SBHB_8_2 GGGGAACCT ACGGCTCGAT CACCTCCTAT CGCCGGAAAC CCATCCGGGG GGTTTAACCC CGGATGGGAT TTTGGTACTC CCCTTTATGG GGAGTCCCAC CGGCCGTACT CCTATAGCAA TTCCGGTTGA TCCTGCCGGA GGCCACTGCT ATGGGGGTCC GACTAAGCCA TGCGAG
16SBHB_4_7 CCT ACGGCTCGAT CACCTCCTAT CGCCGGAAAC CCATCCGGGG GGTTTAACCC CGGATGGGAT TTTGGTACTC CCCTTTATGG GGAGTCCCAC CGGCCGTACT CCTATAGCAA TTCCGGTTGA TCCTGCCGGA GGCCACTGCT ATGGGGGTCC GACTAAGCCA TGCGAGTCAT GGG
16SBHB_27_ GGGGAACCT ACGGCTCGAT CACCTCCTAT CGCCGGAAAC CCATCCGGGG GGTTTAACCC CGGATGGGAT TTTGGTACTC CCCTTTATGG GGAGTCCCAC CGGCCGTACT CCTATAGCAA TTCCGGTTGA TCCTGCCGGA GGCCACTGCT ATGGGGGTCC GACTAAGCCA TGCG
16SBHB_11_ GGGGAACCT ACGGCTCGAT CACCTCCTAT CGCCGGAAAC CCATCCGGGG GGTTTAACCC CGGATGGGAT TTTGGTACTC CCCTTTATGG GGAGTCCCAC CGGCCGTACT CCTATAGCAA TTCCGGTTGA TCCTGCCGGA GGCCACTGCT ATGGGGGTCC GACTAAGCCA TGC
16SBHB_9_2 GGGAACCT ACGGCTCGAT CACCTCCTAT CGCCGGAAAC CCATCCGGGG GGTTTAACCC CGGATGGGAT TTTGGTACTC CCCTTTATGG GGAGTCCCAC CGGCCGTACT CCTATAGCAA TTCCGGTTGA TCCTGCCGGA GGCCACTGCT ATGGGGGTCC GACTAAGCCA TGCG
16SBHB_6_3 GGGAACCT ACGGCTCGAT CACCTCCTAT CGCCGGAAAC CCATCCGGGG GGTTTAACCC CGGATGGGAT TTTGGTACTC CCCTTTATGG GGAGTCCCAC CGGCCGTACT CCTATAGCAA TTCCGGTTGA TCCTGCCGGA GGCCACTGCT ATGGGGGTCC GACTAAGCCA TGCGAG
16SBHB_3_8 GGAACCT ACGGCTCGAT CACCTCCTAT CGCCGGAAAC CCATCCGGGG GGTTTAACCC CGGATGGGAT TTTGGTACTC CCCTTTATGG GGAGTCCCAC CGGCCGTACT CCTATAGCAA TTCCGGTTGA TCCTGCCGGA GGCCACTGCT ATGGGGGTCC GACTAAGCCA TGCGAG
16SBHB_17_ GGGGAACCT ACGGCTCGAT CACCTCCTAT CGCCGGAAAC CCATCCGGGG GGTTTAACCC CGGATGGGAT TTTGGTACTC CCCTTTATGG GGAGTCCCAC CGGCCGTACT CCTATAGCAA TTCCGGTTGA TCCTGCCGGA GGCCACTGCT ATGGGGGTCC GACTAAGCCA TG
16SBHB_4_7 AGGGGAACCT ACGGCTCGAT CACCTCCTAT CGCCGGAAAC CCATCCGGGG GGTTTAACCC CGGATGGGAT TTTGGTACTC CCCTTTATGG GGAGTCCCAC CGGCCGTACT CCTATAGCAA TTCCGGTTGA TCCTGCCGGA GGCCACTGCT ATGGGGGTCC GACTAAGCCA TG
16SBHB_14_ GGAACCT ACGGCTCGAT CACCTCCTAT CGCCGGAAAC CCATCCGGGG GGTTTAACCC CGGATGGGAT TTTGGTACTC CCCTTTATGG GGAGTCCCAC CGGCCGTACT CCTATAGCAA TTCCGGTTGA TCCTGCCGGA GGCCACTGCT ATGGGGGTCC GACTAAGCCA TGCG
16SBHB_7_3 GGGAACCT ACGGCTCGAT CACCTCCTAT CGCCGGAAAC CCATCCGGGG GGTTTAACCC CGGATGGGAT TTTGGTACTC CCCTTTATGG GGAGTCCCAC CGGCCGTACT CCTATAGCAA TTCCGGTTGA TCCTGCCGGA GGCCACTGCT ATGGGGGTCC GACTAAGCCA TGC
16SBHB_31_ GGGGAACCT ACGGCTCGAT CACCTCCTAT CGCCGGAAAC CCATCCGGGG GGTTTAACCC CGGATGGGAT TTTGGTACTC CCCTTTATGG GGAGTCCCAC CGGCCGTACT CCTATAGCAA TTCCGGTTGA TCCTGCCGGA GGCCACTGCT ATGGGGGTCC GACTAAGCC
16SBHB_21_ GGAACCT ACGGCTCGAT CACCTCCTAT CGCCGGAAAC CCATCCGGGG GGTTTAACCC CGGATGGGAT TTTGGTACTC CCCTTTATGG GGAGTCCCAC CGGCCGTACT CCTATAGCAA TTCCGGTTGA TCCTGCCGGA GGCCACTGCT ATGGGGGTCC GACTAAGCCA TG
16SBHB_11_ GGGAACCT ACGGCTCGAT CACCTCCTAT CGCCGGAAAC CCATCCGGGG GGTTTAACCC CGGATGGGAT TTTGGTACTC CCCTTTATGG GGAGTCCCAC CGGCCGTACT CCTATAGCAA TTCCGGTTGA TCCTGCCGGA GGCCACTGCT ATGGGGGTCC GACTAAGCCA TG
16SBHB_16_ GGGGAACCT ACGGCTCGAT CACCTCCTAT CGCCGGAAAC CCATCCGGGG GGTTTAACCC CGGATGGGAT TTTGGTACTC CCCTTTATGG GGAGTCCCAC CGGCCGTACT CCTATAGCAA TTCCGGTTGA TCCTGCCGGA GGCCACTGCT ATGGGGGTCC GACTAAGCCA
16SBHB_15_ GGGGAACCT ACGGCTCGAT CACCTCCTAT CGCCGGAAAC CCATCCGGGG GGTTTAACCC CGGATGGGAT TTTGGTACTC CCCTTTATGG GGAGTCCCAC CGGCCGTACT CCTATAGCAA TTCCGGTTGA TCCTGCCGGA GGCCACTGCT ATGGGGGTCC GACTAAGCCA T
16SBHB_8_2 GGGAACCT ACGGCTCGAT CACCTCCTAT CGCCGGAAAC CCATCCGGGG GGTTTAACCC CGGATGGGAT TTTGGTACTC CCCTTTATGG GGAGTCCCAC CGGCCGTACT CCTATAGCAA TTCCGGTTGA TCCTGCCGGA GGCCACTGCT ATGGGGGTCC GACTAAGCCA T
16SBHB_5_4 GGGAACCT ACGGCTCGAT CACCTCCTAT CGCCGGAAAC CCATCCGGGG GGTTTAACCC CGGATGGGAT TTTGGTACTC CCCTTTATGG GGAGTCCCAC CGGCCGTACT CCTATAGCAA TTCCGGTTGA TCCTGCCGGA GGCCACTGCT ATGGGGGTCC GACTAAGCCA G
16SBHB_14_ GGAACCT ACGGCTCGAT CACCTCCTAT CGCCGGAAAC CCATCCGGGG GGTTTAACCC CGGATGGGAT TTTGGTACTC CCCTTTATGG GGAGTCCCAC CGGCCGTACT CCTATAGCAA TTCCGGTTGA TCCTGCCGGA GGCCACTGCT ATGGGGGTCC GACTAAGCCA T
16SBHB_4_6 GGGAACCT ACGGCTCGAT CACCTCCTAT CGCCGGAAAC CCATCCGGGG GGTTTAACCC CGGATGGGAT TTTGGTACTC CCCTTTATGG GGAGTCCCAC CGGCCGTACT CCTATAGCAA TTCCGGTTGA TCCTGCCGGA GGCCACTGCT ATGGGGGTCC GACTAAGCCG
16SBHB_3_8 GGGGAACCT ACGGCTCGAT CACCTCCTAT CGCCGGAAAC CCATCCGGGG GGTTTAACCC CGGATGGGAT TTTGGTACTC CCCTTTATGG GGAGTCCCAC CGGCCGTACT CCTATAGCAA TTCCGGTTGA TCCTGCCGGA GGCCACTGCT ATGGGGGTCC GACTAAGCCG
16SBHB_14_ CCT ACGGCTCGAT CACCTCCTAT CGCCGGAAAC CCATCCGGGG GGTTTAACCC CGGATGGGAT TTTGGTACTC CCCTTTATGG GGAGTCCCAC CGGCCGTACT CCTATAGCAA TTCCGGTTGA TCCTGCCGGA GGCCACTGCT ATGGGGGTCC GACTAAGCCA TGCGAG
16SBHB_7_3 CT ACGGCTCGAT CACCTCCTAT CGCCGGAAAC CCATCCGGGG GGTTTAACCC CGGATGGGAT TTTGGTACTC CCCTTTATGG GGAGTCCCAC CGGCCGTACT CCTATAGCAA TTCCGGTTGA TCCTGCCGGA GGCCACTGCT ATGGGGGTCC GACTAAGCCA TGCGAG
16SBHB_4_6 GAACCT ACGGCTCGAT CACCTCCTAT CGCCGGAAAC CCATCCGGGG GGTTTAACCC CGGATGGGAT TTTGGTACTC CCCTTTATGG GGAGTCCCAC CGGCCGTACT CCTATAGCAA TTCCGGTTGA TCCTGCCGGA GGCCACTGCT ATGGGGGTCC GACTAAGCCA TGC
16SBHB_52_ CCT ACGGCTCGAT CACCTCCTAT CGCCGGAAAC CCATCCGGGG GGTTTAACCC CGGATGGGAT TTTGGTACTC CCCTTTATGG GGAGTCCCAC CGGCCGTACT CCTATAGCAA TTCCGGTTGA TCCTGCCGGA GGCCACTGCT ATGGGGGTCC GACTAAGCCA TGCG
16SBHB_19_ CT ACGGCTCGAT CACCTCCTAT CGCCGGAAAC CCATCCGGGG GGTTTAACCC CGGATGGGAT TTTGGTACTC CCCTTTATGG GGAGTCCCAC CGGCCGTACT CCTATAGCAA TTCCGGTTGA TCCTGCCGGA GGCCACTGCT ATGGGGGTCC GACTAAGCCA TGCG
16SBHB_17_ CCT ACGGCTCGAT CACCTCCTAT CGCCGGAAAC CCATCCGGGG GGTTTAACCC CGGATGGGAT TTTGGTACTC CCCTTTATGG GGAGTCCCAC CGGCCGTACT CCTATAGCAA TTCCGGTTGA TCCTGCCGGA GGCCACTGCT ATGGGGGTCC GACTAAGCCA TGC
16SBHB_5_5 CCT ACGGCTCGAT CACCTCCTAT CGCCGGAAAC CCATCCGGGG GGTTTAACCC CGGATGGGAT TTTGGTACTC CCCTTTATGG GGAGTCCCAC CGGCCGTACT CCTATAGCAA TTCCGGTTGA TCCTGCCGGA GGCCACTGCT ATGGGGGTCC GACTAAGCCA TGCT
16SBHB_17_ GGAACCT ACGGCTCGAT CACCTCCTAT CGCCGGAAAC CCATCCGGGG GGTTTAACCC CGGATGGGAT TTTGGTACTC CCCTTTATGG GGAGTCCCAC CGGCCGTACT CCTATAGCAA TTCCGGTTGA TCCTGCCGGA GGCCACTGCT ATGGGGGTCC GACTAAGCC
16SBHB_14_ GGAACCT ACGGCTCGAT CACCTCCTAT CGCCGGAAAC CCATCCGGGG GGTTTAACCC CGGATGGGAT TTTGGTACTC CCCTTTATGG GGAGTCCCAC CGGCCGTACT CCTATAGCAA TTCCGGTTGA TCCTGCCGGA GGCCACTGCT ATGGGGGTCC GACTAAGCCA
16SBHB_5_4 GGAACCT ACGGCTCGAT CACCTCCTAT CGCCGGAAAC CCATCCGGGG GGTTTAACCC CGGATGGGAT TTTGGTACTC CCCTTTATGG GGAGTCCCAC CGGCCGTACT CCTATAGCAA TTCCGGTTGA TCCTGCCGGA GGCCACTGCT ATGGGGGTCC GACTAAGCCG
16SBHB_3_9 GAACCT ACGGCTCGAT CACCTCCTAT CGCCGGAAAC CCATCCGGGG GGTTTAACCC CGGATGGGAT TTTGGTACTC CCCTTTATGG GGAGTCCCAC CGGCCGTACT CCTATAGCAA TTCCGGTTGA TCCTGCCGGA GGCCACTGCT ATGGGGGTCC GACTAAGCCA
16SBHB_97_ GGGGAACCT ACGGCTCGAT CACCTCCTAT CGCCGGAAAC CCATCCGGGG GGTTTAACCC CGGATGGGAT TTTGGTACTC CCCTTTATGG GGAGTCCCAC CGGCCGTACT CCTATAGCAA TTCCGGTTGA TCCTGCCGGA GGCCACTGCT ATGGGGGTCC GACTAAG
16SBHB_51_ GGGGAACCT ACGGCTCGAT CACCTCCTAT CGCCGGAAAC CCATCCGGGG GGTTTAACCC CGGATGGGAT TTTGGTACTC CCCTTTATGG GGAGTCCCAC CGGCCGTACT CCTATAGCAA TTCCGGTTGA TCCTGCCGGA GGCCACTGCT ATGGGGGTCC GACTAAGC
16SBHB_28_ AGGGGAACCT ACGGCTCGAT CACCTCCTAT CGCCGGAAAC CCATCCGGGG GGTTTAACCC CGGATGGGAT TTTGGTACTC CCCTTTATGG GGAGTCCCAC CGGCCGTACT CCTATAGCAA TTCCGGTTGA TCCTGCCGGA GGCCACTGCT ATGGGGGTCC GACTAAG
16SBHB_17_ GGGAACCT ACGGCTCGAT CACCTCCTAT CGCCGGAAAC CCATCCGGGG GGTTTAACCC CGGATGGGAT TTTGGTACTC CCCTTTATGG GGAGTCCCAC CGGCCGTACT CCTATAGCAA TTCCGGTTGA TCCTGCCGGA GGCCACTGCT ATGGGGGTCC GACTAAGC
16SBHB_15_ GGGAACCT ACGGCTCGAT CACCTCCTAT CGCCGGAAAC CCATCCGGGG GGTTTAACCC CGGATGGGAT TTTGGTACTC CCCTTTATGG GGAGTCCCAC CGGCCGTACT CCTATAGCAA TTCCGGTTGA TCCTGCCGGA GGCCACTGCT ATGGGGGTCC GACTAAGCC
16SBHB_3_8 GGGGAACCT ACGGCTCGAT CACCTCCTAT CGCCGGAAAC CCATCCGGGG GGTTTAACCC CGGATGGGAT TTTGGTACTC CCCTTTATGG GGAGTCCCAC CGGCCGTACT CCTATAGCAA TTCCGGTTGA TCCTGCCGGA GGCCACTGCT ATGGGGGTCC GACTAAGCC
16SBHB_36_ GGGAACCT ACGGCTCGAT CACCTCCTAT CGCCGGAAAC CCATCCGGGG GGTTTAACCC CGGATGGGAT TTTGGTACTC CCCTTTATGG GGAGTCCCAC CGGCCGTACT CCTATAGCAA TTCCGGTTGA TCCTGCCGGA GGCCACTGCT ATGGGGGTCC GACTAAG
16SBHB_22_ GGAACCT ACGGCTCGAT CACCTCCTAT CGCCGGAAAC CCATCCGGGG GGTTTAACCC CGGATGGGAT TTTGGTACTC CCCTTTATGG GGAGTCCCAC CGGCCGTACT CCTATAGCAA TTCCGGTTGA TCCTGCCGGA GGCCACTGCT ATGGGGGTCC GACTAAGC
16SBHB_26_ CCT ACGGCTCGAT CACCTCCTAT CGCCGGAAAC CCATCCGGGG GGTTTAACCC CGGATGGGAT TTTGGTACTC CCCTTTATGG GGAGTCCCAC CGGCCGTACT CCTATAGCAA TTCCGGTTGA TCCTGCCGGA GGCCACTGCT ATGGGGGTCC GACTAAGCCA TG
16SBHB_4_7 ACCT ACGGCTCGAT CACCTCCTAT CGCCGGAAAC CCATCCGGGG GGTTTAACCC CGGATGGGAT TTTGGTACTC CCCTTTATGG GGAGTCCCAC CGGCCGTACT CCTATAGCAA TTCCGGTTGA TCCTGCCGGA GGCCACTGCT ATGGGGGTCC GACTAAGCCA TG
16SBHB_18_ CT ACGGCTCGAT CACCTCCTAT CGCCGGAAAC CCATCCGGGG GGTTTAACCC CGGATGGGAT TTTGGTACTC CCCTTTATGG GGAGTCCCAC CGGCCGTACT CCTATAGCAA TTCCGGTTGA TCCTGCCGGA GGCCACTGCT ATGGGGGTCC GACTAAGCCA TG
16SBHB_8_2 CT ACGGCTCGAT CACCTCCTAT CGCCGGAAAC CCATCCGGGG GGTTTAACCC CGGATGGGAT TTTGGTACTC CCCTTTATGG GGAGTCCCAC CGGCCGTACT CCTATAGCAA TTCCGGTTGA TCCTGCCGGA GGCCACTGCT ATGGGGGTCC GACTAAGCCA TGC
16SBHB_3_9 CT ACGGCTCGAT CACCTCCTAT CGCCGGAAAC CCATCCGGGG GGTTTAACCC CGGATGGGAT TTTGGTACTC CCCTTTATGG GGAGTCCCAC CGGCCGTACT CCTATAGCAA TTCCGGTTGA TCCTGCCGGA GGCCACTGCT ATGGGGGTCC GACTAAGCCA CGCG
16SBHB_4_6 GAACCT ACGGCTCGAT CACCTCCTAT CGCCGGAAAC CCATCCGGGG GGTTTAACCC CGGATGGGAT TTTGGTACTC CCCTTTATGG GGAGTCCCAC CGGCCGTACT CCTATAGCAA TTCCGGTTGA TCCTGCCGGA GGCCACTGCT ATGGGGGTCC GACTAAGCC
16SBHB_3_9 GAACCT ACGGCTCGAT CACCTCCTAT CGCCGGAAAC CCATCCGGGG GGTTTAACCC CGGATGGGAT TTTGGTACTC CCCTTTATGG GGAGTCCCAC CGGCCGTACT CCTATAGCAA TTCCGGTTGA TCCTGCCGGA GGCCACTGCT ATGGGGGTCC GACTAAGC
16SBHB_64_ GGAACCT ACGGCTCGAT CACCTCCTAT CGCCGGAAAC CCATCCGGGG GGTTTAACCC CGGATGGGAT TTTGGTACTC CCCTTTATGG GGAGTCCCAC CGGCCGTACT CCTATAGCAA TTCCGGTTGA TCCTGCCGGA GGCCACTGCT ATGGGGGTCC GACTAAG
16SBHB_11_ GAACCT ACGGCTCGAT CACCTCCTAT CGCCGGAAAC CCATCCGGGG GGTTTAACCC CGGATGGGAT TTTGGTACTC CCCTTTATGG GGAGTCCCAC CGGCCGTACT CCTATAGCAA TTCCGGTTGA TCCTGCCGGA GGCCACTGCT ATGGGGGTCC GACTAAG
16SBHB_31_ CCT ACGGCTCGAT CACCTCCTAT CGCCGGAAAC CCATCCGGGG GGTTTAACCC CGGATGGGAT TTTGGTACTC CCCTTTATGG GGAGTCCCAC CGGCCGTACT CCTATAGCAA TTCCGGTTGA TCCTGCCGGA GGCCACTGCT ATGGGGGTCC GACTAAGCCA T
16SBHB_15_ CT ACGGCTCGAT CACCTCCTAT CGCCGGAAAC CCATCCGGGG GGTTTAACCC CGGATGGGAT TTTGGTACTC CCCTTTATGG GGAGTCCCAC CGGCCGTACT CCTATAGCAA TTCCGGTTGA TCCTGCCGGA GGCCACTGCT ATGGGGGTCC GACTAAGCCA T
16SBHB_9_2 CCT ACGGCTCGAT CACCTCCTAT CGCCGGAAAC CCATCCGGGG GGTTTAACCC CGGATGGGAT TTTGGTACTC CCCTTTATGG GGAGTCCCAC CGGCCGTACT CCTATAGCAA TTCCGGTTGA TCCTGCCGGA GGCCACTGCT ATGGGGGTCC GACTAAGCCA
16SBHB_5_5 CCT ACGGCTCGAT CACCTCCTAT CGCCGGAAAC CCATCCGGGG GGTTTAACCC CGGATGGGAT TTTGGTACTC CCCTTTATGG GGAGTCCCAC CGGCCGTACT CCTATAGCAA TTCCGGTTGA TCCTGCCGGA GGCCACTGCT ATGGGGGTCC GACTAAGCCA G
16SBHB_19_ CGGCTCGAT CACCTCCTAT CGCCGGAAAC CCATCCGGGG GGTTTAACCC CGGATGGGAT TTTGGTACTC CCCTTTATGG GGAGTCCCAC CGGCCGTACT CCTATAGCAA TTCCGGTTGA TCCTGCCGGA GGCCACTGCT ATGGGGGTCC GACTAAGCCA TGCG
16SBHB_5_5 CGGCTCGAT CACCTCCTAT CGCCGGAAAC CCATCCGGGG GGTTTAACCC CGGATGGGAT TTTGGTACTC CCCTTTATGG GGAGTCCCAC CGGCCGTACT CCTATAGCAA TTCCGGTTGA TCCTGCCGGA GGCCACTGCT ATGGGGGTCC GACTAAGCCA TGC
16SBHB_4_6 GGCTCGAT CACCTCCTAT CGCCGGAAAC CCATCCGGGG GGTTTAACCC CGGATGGGAT TTTGGTACTC CCCTTTATGG GGAGTCCCAC CGGCCGTACT CCTATAGCAA TTCCGGTTGA TCCTGCCGGA GGCCACTGCT ATGGGGGTCC GACTAAGCCA TGCG
16SBHB_13_ T ACGGCTCGAT CACCTCCTAT CGCCGGAAAC CCATCCGGGG GGTTTAACCC CGGATGGGAT TTTGGTACTC CCCTTTATGG GGAGTCCCAC CGGCCGTACT CCTATAGCAA TTCCGGTTGA TCCTGCCGGA GGCCACTGCT ATGGGGGTCC GACTAAGCCA TG
16SBHB_5_4 T ACGGCTCGAT CACCTCCTAT CGCCGGAAAC CCATCCGGGG GGTTTAACCC CGGATGGGAT TTTGGTACTC CCCTTTATGG GGAGTCCCAC CGGCCGTACT CCTATAGCAA TTCCGGTTGA TCCTGCCGGA GGCCACTGCT ATGGGGGTCC GACTAAGCCA T
16SBHB_9_2 CACCTCCTAT CGCCGGAAAC CCATCCGGGG GGTTTAACCC CGGATGGGAT TTTGGTACTC CCCTTTATGG GGAGTCCCAC CGGCCGTACT CCTATAGCAA TTCCGGTTGA TCCTGCCGGA GGCCACTGCT ATGGGGGTCC GACTAAGCCA TGCGAGTCAT GGG
16SBHB_5_5 CACCTCCTAT CGCCGGAAAC CCATCCGGGG GGTTTAACCC CGGATGGGAT TTTGGTACTC CCCTTTATGG GGAGTCCCAC CGGCCGTACT CCTATAGCAA TTCCGGTTGA TCCTGCCGGA GGCCACTGCT ATGGGGGTCC GACTAAGCCA TGCGAGTCAT GGGG
16SBHB_3_8 GGGGAACCT ACGGCTCGAT CACCTCCTAT CGCCGGAAAC CCATCCGGGG GGTTTAACCC CGGATGGGAT TTTGGTACTC CCCTTTATGG GGAGTCCCAC CGGCCGTACT CCTATAGCAA TTCCGGTTGA TCCTGCCGGA GGCCACTGCT ATGGGGGTCC GAC
16SBHB_3_9 CGT AGGGGAACCT ACGGCTCGAT CACCTCCTAT CGCCGGAAAC CCATCCGGGG GGTTTAACCC CGGATGGGAT TTTGGTACTC CCCTTTATGG GGAGTCCCAC CGGCCGTACT CCTATAGCAA TTCCGGTTGA TCCTGCCGGA GGCCACTGCT ATGGGGGTCC GAC
16SBHB_5_5 ACCT ACGGCTCGAT CACCTCCTAT CGCCGGAAAC CCATCCGGGG GGTTTAACCC CGGATGGGAT TTTGGTACTC CCCTTTATGG GGAGTCCCAC CGGCCGTACT CCTATAGCAA TTCCGGTTGA TCCTGCCGGA GGCCACTGCT ATGGGGGTCC GACTAAG
16SBHB_4_7 AACCT ACGGCTCGAT CACCTCCTAT CGCCGGAAAC CCATCCGGGG GGTTTAACCC CGGATGGGAT TTTGGTACTC CCCTTTATGG GGAGTCCCAC CGGCCGTACT CCTATAGCAA TTCCGGTTGA TCCTGCCGGA GGCCACTGCT ATGGGGGTCC GACTAAG
16SBHB_14_ CGGCTCGAT CACCTCCTAT CGCCGGAAAC CCATCCGGGG GGTTTAACCC CGGATGGGAT TTTGGTACTC CCCTTTATGG GGAGTCCCAC CGGCCGTACT CCTATAGCAA TTCCGGTTGA TCCTGCCGGA GGCCACTGCT ATGGGGGTCC GACTAAGCCA TG
16SBHB_9_2 CGGCTCGAT CACCTCCTAT CGCCGGAAAC CCATCCGGGG GGTTTAACCC CGGATGGGAT TTTGGTACTC CCCTTTATGG GGAGTCCCAC CGGCCGTACT CCTATAGCAA TTCCGGTTGA TCCTGCCGGA GGCCACTGCT ATGGGGGTCC GACTAAGCCA T
16SBHB_5_5 CGAT CACCTCCTAT CGCCGGAAAC CCATCCGGGG GGTTTAACCC CGGATGGGAT TTTGGTACTC CCCTTTATGG GGAGTCCCAC CGGCCGTACT CCTATAGCAA TTCCGGTTGA TCCTGCCGGA GGCCACTGCT ATGGGGGTCC GACTAAGCCA TGCGAG
16SBHB_52_ CT ACGGCTCGAT CACCTCCTAT CGCCGGAAAC CCATCCGGGG GGTTTAACCC CGGATGGGAT TTTGGTACTC CCCTTTATGG GGAGTCCCAC CGGCCGTACT CCTATAGCAA TTCCGGTTGA TCCTGCCGGA GGCCACTGCT ATGGGGGTCC GACTAAG
16SBHB_16_ CT ACGGCTCGAT CACCTCCTAT CGCCGGAAAC CCATCCGGGG GGTTTAACCC CGGATGGGAT TTTGGTACTC CCCTTTATGG GGAGTCCCAC CGGCCGTACT CCTATAGCAA TTCCGGTTGA TCCTGCCGGA GGCCACTGCT ATGGGGGTCC GACTAAGC
16SBHB_4_5 T ACGGCTCGAT CACCTCCTAT CGCCGGAAAC CCATCCGGGG GGTTTAACCC CGGATGGGAT TTTGGTACTC CCCTTTATGG GGAGTCCCAC CGGCCGTACT CCTATAGCAA TTCCGGTTGA TCCTGCCGGA GGCCACTGCT ATGGGGGTCC GACTAAGCC
16SBHB_11_ T ACGGCTCGAT CACCTCCTAT CGCCGGAAAC CCATCCGGGG GGTTTAACCC CGGATGGGAT TTTGGTACTC CCCTTTATGG GGAGTCCCAC CGGCCGTACT CCTATAGCAA TTCCGGTTGA TCCTGCCGGA GGCCACTGCT ATGGGGGTCC GACTAAGC
16SBHB_7_3 CTCCTAT CGCCGGAAAC CCATCCGGGG GGTTTAACCC CGGATGGGAT TTTGGTACTC CCCTTTATGG GGAGTCCCAC CGGCCGTACT CCTATAGCAA TTCCGGTTGA TCCTGCCGGA GGCCACTGCT ATGGGGGTCC GACTAAGCCA TGCGAGTCAT GGG
16SBHB_4_7 CCTCCTAT CGCCGGAAAC CCATCCGGGG GGTTTAACCC CGGATGGGAT TTTGGTACTC CCCTTTATGG GGAGTCCCAC CGGCCGTACT CCTATAGCAA TTCCGGTTGA TCCTGCCGGA GGCCACTGCT ATGGGGGTCC GACTAAGCCA TGCGAGTCAT GGG
16SBHB_3_7 TTCGTAACA AGGTAGCCGT AGGGGAACCT ACGGCTCGAT CACCTCCTAT CGCCGGAAAC CCATCCGGGG GGTTTAACCC CGGATGGGAT TTTGGTACTC CCCTTTATGG GGAGTCCCAC CGGCCGTACT CCTATAGCAA TTCCGGTTGA TCCTGCCGGA GGCCACTGCT ATGGGGG
16SBHB_20_ CGGCTCGAT CACCTCCTAT CGCCGGAAAC CCATCCGGGG GGTTTAACCC CGGATGGGAT TTTGGTACTC CCCTTTATGG GGAGTCCCAC CGGCCGTACT CCTATAGCAA TTCCGGTTGA TCCTGCCGGA GGCCACTGCT ATGGGGGTCC GACTAAGCC
```

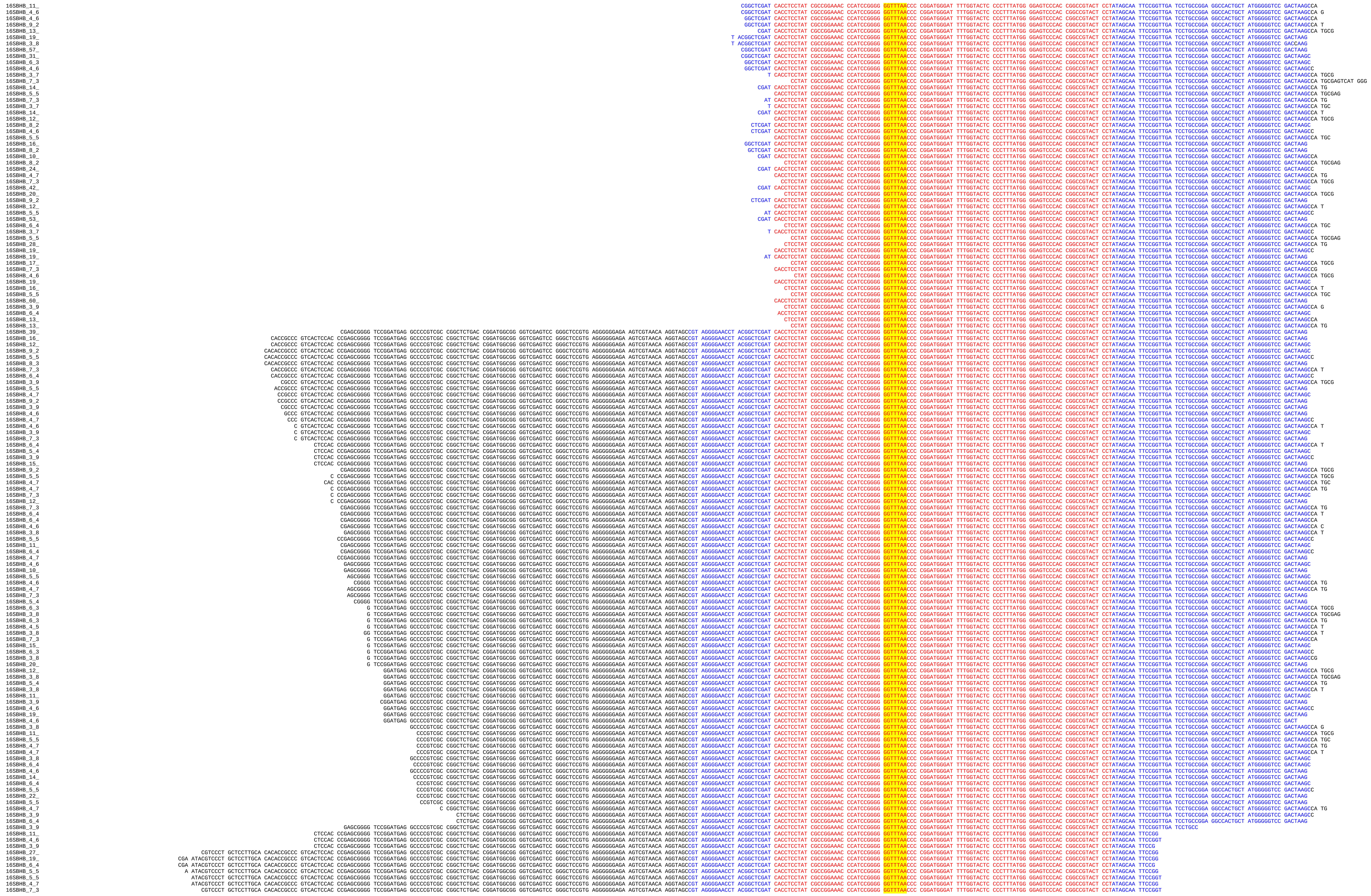

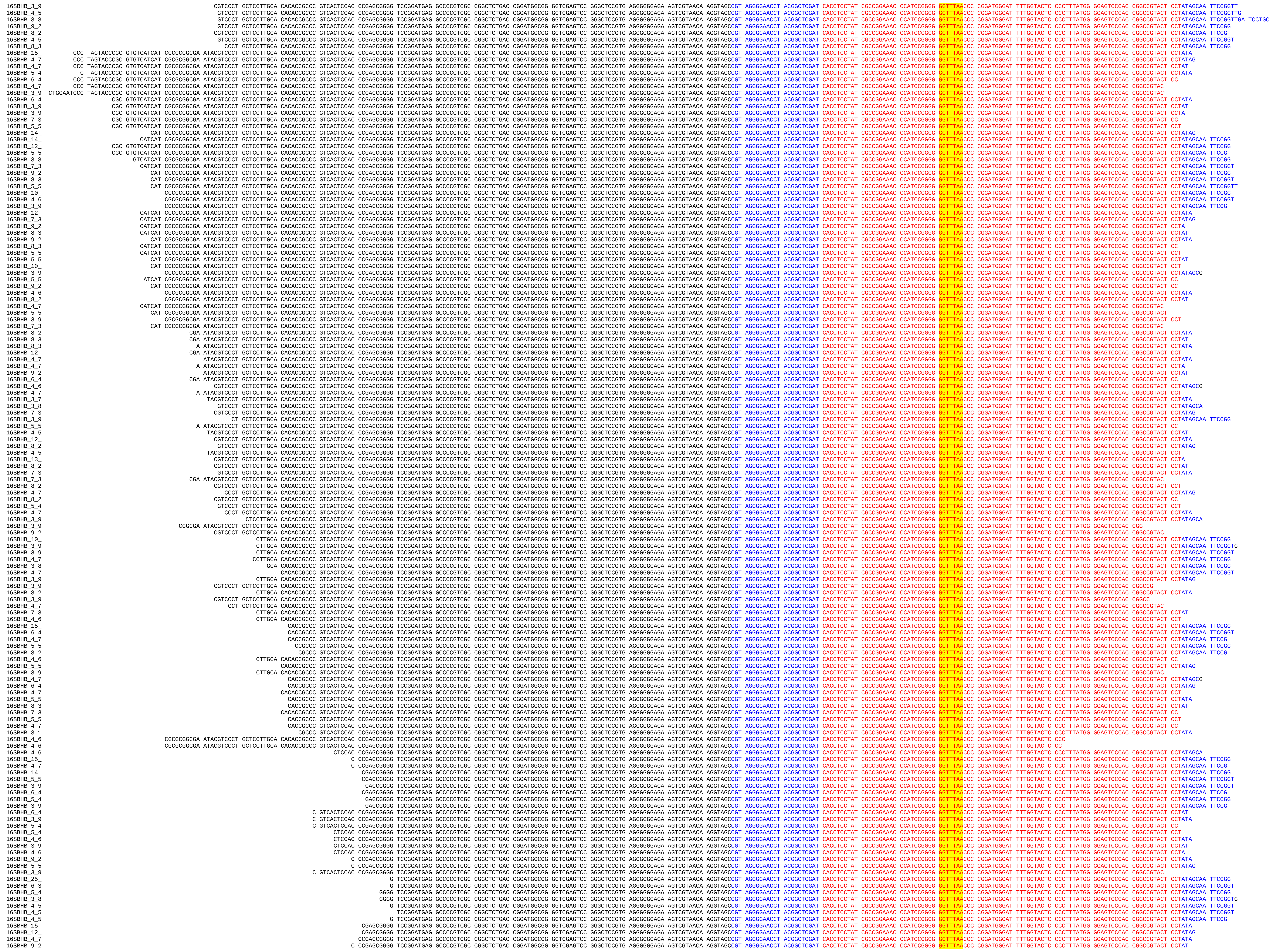

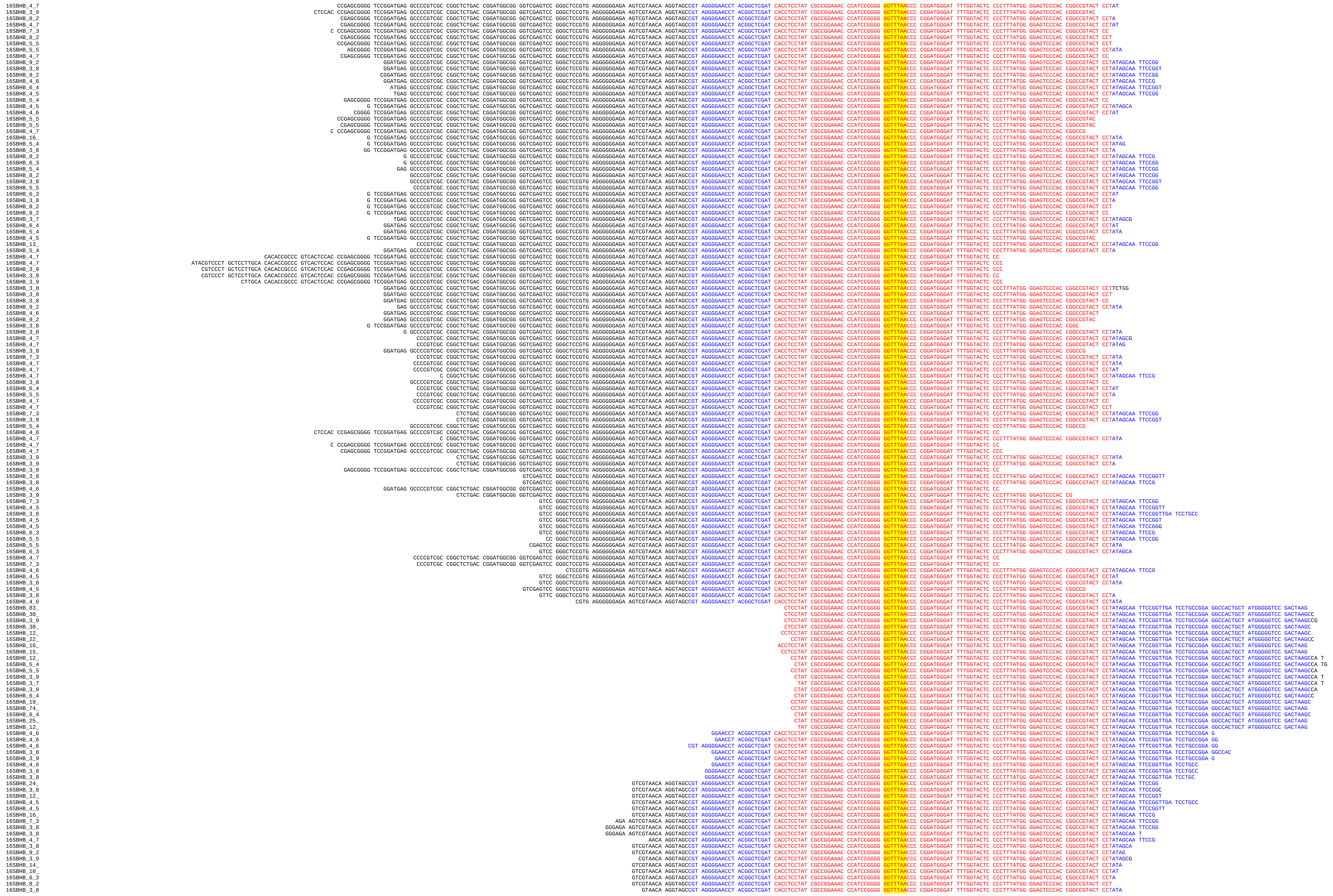

```
16SBHB_3_8  TAACA AGGTAGCCGT AGGGGAACCT ACGGCTCGAT CACCTCCTAT CGCCGGAAAC CCATCCGGGG GGTTTAACCC CGGATGGGAT TTTGGTACTC CCCTTTATGG GGAGTCCCAC CGGCCGTACT CCTATA
16SBHB_8_2  GTCGTAACA AGGTAGCCGT AGGGGAACCT ACGGCTCGAT CACCTCCTAT CGCCGGAAAC CCATCCGGGG GGTTTAACCC CGGATGGGAT TTTGGTACTC CCCTTTATGG GGAGTCCCAC CGGCCGTACT CC
16SBHB_5_4  CGTG AGGGGGGAGA AGTCGTAACA AGGTAGCCGT AGGGGAACCT ACGGCTCGAT CACCTCCTAT CGCCGGAAAC CCATCCGGGG GGTTTAACCC CGGATGGGAT TTTGGTACTC CCCTTTATGG GGAGTCCCAC CGGCCGTAC
16SBHB_3_9  CTCCGTG AGGGGGGAGA AGTCGTAACA AGGTAGCCGT AGGGGAACCT ACGGCTCGAT CACCTCCTAT CGCCGGAAAC CCATCCGGGG GGTTTAACCC CGGATGGGAT TTTGGTACTC CCCTTTATGG GGAGTCCCAC CGGCCGTAC
16SBHB_3_8  GGGATA AGTCGTAACA AGGTAGCCGT AGGGGAACCT ACGGCTCGAT CACCTCCTAT CGCCGGAAAC CCATCCGGGG GGTTTAACCC CGGATGGGAT TTTGGTACTC CCCTTTATGG GGAGTCCCAC CGGCCGTACT CCA
16SBHB_12_  CGGCTCGAT CACCTCCTAT CGCCGGAAAC CCATCCGGGG GGTTTAACCC CGGATGGGAT TTTGGTACTC CCCTTTATGG GGAGTCCCAC CGGCCGTACT CCTATAGCAA TTCCGGTTGA TCCTGCCGGA GGCCACTGCT ATGGGG
16SBHB_4_6  CTCGAT CACCTCCTAT CGCCGGAAAC CCATCCGGGG GGTTTAACCC CGGATGGGAT TTTGGTACTC CCCTTTATGG GGAGTCCCAC CGGCCGTACT CCTATAGCAA TTCCGGTTGA TCCTGCCGGA GGCCACTGCT ATGAG
16SBHB_4_6  CGAT CACCTCCTAT CGCCGGAAAC CCATCCGGGG GGTTTAACCC CGGATGGGAT TTTGGTACTC CCCTTTATGG GGAGTCCCAC CGGCCGTACT CCTATAGCAA TTCCGGTTGA TCCTGCCGGA GGCCACTGCT ATGGGG
16SBHB_3_9  CGGCTCGAT CACCTCCTAT CGCCGGAAAC CCATCCGGGG GGTTTAACCC CGGATGGGAT TTTGGTACTC CCCTTTATGG GGAGTCCCAC CGGCCGTACT CCTATAGCAA TTCCGGTTGA TCCTGCCGGA GGCCACTGC
16SBHB_4_7  CACCTCCTAT CGCCGGAAAC CCATCCGGGG GGTTTAACCC CGGATGGGAT TTTGGTACTC CCCTTTATGG GGAGTCCCAC CGGCCGTACT CCTATAGCAA TTCCGGTTGA TCCTGCCGGA GGCCACTGCT ATGGGG
16SBHB_5_4  CTCCTAT CGCCGGAAAC CCATCCGGGG GGTTTAACCC CGGATGGGAT TTTGGTACTC CCCTTTATGG GGAGTCCCAC CGGCCGTACT CCTATAGCAA TTCCGGTTGA TCCTGCCGGA GGCCACTGCT ATGGGG
16SBHB_44_  CGT AGGGGAACCT ACGGCTCGAT CACCTCCTAT CGCCGGAAAC CCATCCGGGG GGTTTAACCC CGGATGGGAT TTTGGTACTC CCCTTTATGG GGAGTCCCAC CGGCCGTACT CCTATAGCAA TTCCGG
16SBHB_18_  CGT AGGGGAACCT ACGGCTCGAT CACCTCCTAT CGCCGGAAAC CCATCCGGGG GGTTTAACCC CGGATGGGAT TTTGGTACTC CCCTTTATGG GGAGTCCCAC CGGCCGTACT CCTATAGCAA TTCCGGT
16SBHB_13_  CGT AGGGGAACCT ACGGCTCGAT CACCTCCTAT CGCCGGAAAC CCATCCGGGG GGTTTAACCC CGGATGGGAT TTTGGTACTC CCCTTTATGG GGAGTCCCAC CGGCCGTACT CCTATAGCAA TTCCGGTT
16SBHB_6_4  CGT AGGGGAACCT ACGGCTCGAT CACCTCCTAT CGCCGGAAAC CCATCCGGGG GGTTTAACCC CGGATGGGAT TTTGGTACTC CCCTTTATGG GGAGTCCCAC CGGCCGTACT CCTATAGCAA TTCCGGTTGA T
16SBHB_3_8  GT AGGGGAACCT ACGGCTCGAT CACCTCCTAT CGCCGGAAAC CCATCCGGGG GGTTTAACCC CGGATGGGAT TTTGGTACTC CCCTTTATGG GGAGTCCCAC CGGCCGTACT CCTATAGCAA TTCCGGT
16SBHB_11_  GGTAGCCGT AGGGGAACCT ACGGCTCGAT CACCTCCTAT CGCCGGAAAC CCATCCGGGG GGTTTAACCC CGGATGGGAT TTTGGTACTC CCCTTTATGG GGAGTCCCAC CGGCCGTACT CCTATAGCAA TTCCGG
16SBHB_4_5  GTAGCCGT AGGGGAACCT ACGGCTCGAT CACCTCCTAT CGCCGGAAAC CCATCCGGGG GGTTTAACCC CGGATGGGAT TTTGGTACTC CCCTTTATGG GGAGTCCCAC CGGCCGTACT CCTATAGCAA TTCCGG
16SBHB_3_7  TAGCCGT AGGGGAACCT ACGGCTCGAT CACCTCCTAT CGCCGGAAAC CCATCCGGGG GGTTTAACCC CGGATGGGAT TTTGGTACTC CCCTTTATGG GGAGTCCCAC CGGCCGTACT CCTATAGCAA TTCCGGT
16SBHB_4_5  GTAGCCGT AGGGGAACCT ACGGCTCGAT CACCTCCTAT CGCCGGAAAC CCATCCGGGG GGTTTAACCC CGGATGGGAT TTTGGTACTC CCCTTTATGG GGAGTCCCAC CGGCCGTACT CCTATAGCAA TTCCG
16SBHB_7_3  CCGT AGGGGAACCT ACGGCTCGAT CACCTCCTAT CGCCGGAAAC CCATCCGGGG GGTTTAACCC CGGATGGGAT TTTGGTACTC CCCTTTATGG GGAGTCCCAC CGGCCGTACT CCTATAGCAA TTCCGG
16SBHB_26_  CGT AGGGGAACCT ACGGCTCGAT CACCTCCTAT CGCCGGAAAC CCATCCGGGG GGTTTAACCC CGGATGGGAT TTTGGTACTC CCCTTTATGG GGAGTCCCAC CGGCCGTACT CCTATAGCAA TTCCG
16SBHB_30_  GGGGAACCT ACGGCTCGAT CACCTCCTAT CGCCGGAAAC CCATCCGGGG GGTTTAACCC CGGATGGGAT TTTGGTACTC CCCTTTATGG GGAGTCCCAC CGGCCGTACT CCTATAGCAA TTCCGG
16SBHB_4_7  AGGGGAACCT ACGGCTCGAT CACCTCCTAT CGCCGGAAAC CCATCCGGGG GGTTTAACCC CGGATGGGAT TTTGGTACTC CCCTTTATGG GGAGTCCCAC CGGCCGTACT CCTATAGCAA TTCCGG
16SBHB_5_5  AGGGGAACCT ACGGCTCGAT CACCTCCTAT CGCCGGAAAC CCATCCGGGG GGTTTAACCC CGGATGGGAT TTTGGTACTC CCCTTTATGG GGAGTCCCAC CGGCCGTACT CCTATAGCAA TTCCG
16SBHB_16_  GGGGAACCT ACGGCTCGAT CACCTCCTAT CGCCGGAAAC CCATCCGGGG GGTTTAACCC CGGATGGGAT TTTGGTACTC CCCTTTATGG GGAGTCCCAC CGGCCGTACT CCTATAGCAA TTCCGGT
16SBHB_5_4  GGGGAACCT ACGGCTCGAT CACCTCCTAT CGCCGGAAAC CCATCCGGGG GGTTTAACCC CGGATGGGAT TTTGGTACTC CCCTTTATGG GGAGTCCCAC CGGCCGTACT CCTATAGCAA TTCCGGTT
16SBHB_3_8  GGGGAACCT ACGGCTCGAT CACCTCCTAT CGCCGGAAAC CCATCCGGGG GGTTTAACCC CGGATGGGAT TTTGGTACTC CCCTTTATGG GGAGTCCCAC CGGCCGTACT CCTATAGCAA TTCCGGTTG
16SBHB_4_6  GGGGAACCT ACGGCTCGAT CACCTCCTAT CGCCGGAAAC CCATCCGGGG GGTTTAACCC CGGATGGGAT TTTGGTACTC CCCTTTATGG GGAGTCCCAC CGGCCGTACT CCTATAGCAA TTCCGGAT
16SBHB_6_3  GGGAACCT ACGGCTCGAT CACCTCCTAT CGCCGGAAAC CCATCCGGGG GGTTTAACCC CGGATGGGAT TTTGGTACTC CCCTTTATGG GGAGTCCCAC CGGCCGTACT CCTATAGCAA TTCCGG
16SBHB_14_  GGGGAACCT ACGGCTCGAT CACCTCCTAT CGCCGGAAAC CCATCCGGGG GGTTTAACCC CGGATGGGAT TTTGGTACTC CCCTTTATGG GGAGTCCCAC CGGCCGTACT CCTATAGCAA TTCCG
16SBHB_4_6  GGAACCT ACGGCTCGAT CACCTCCTAT CGCCGGAAAC CCATCCGGGG GGTTTAACCC CGGATGGGAT TTTGGTACTC CCCTTTATGG GGAGTCCCAC CGGCCGTACT CCTATAGCAA TTCCGGT
16SBHB_3_8  GGAACCT ACGGCTCGAT CACCTCCTAT CGCCGGAAAC CCATCCGGGG GGTTTAACCC CGGATGGGAT TTTGGTACTC CCCTTTATGG GGAGTCCCAC CGGCCGTACT CCTATAGCAA TTCCGGTTG
16SBHB_4_6  CGGCTCGAT CACCTCCTAT CGCCGGAAAC CCATCCGGGG GGTTTAACCC CGGATGGGAT TTTGGTACTC CCCTTTATGG GGAGTCCCAC CGGCCGTACT CCTATAGCAA TTCCGGTTGA TCCTGCCGGA G
16SBHB_3_9  CT ACGGCTCGAT CACCTCCTAT CGCCGGAAAC CCATCCGGGG GGTTTAACCC CGGATGGGAT TTTGGTACTC CCCTTTATGG GGAGTCCCAC CGGCCGTACT CCTATAGCAA TTCCGGTTGA TCCTGCC
16SBHB_21_  GGAACCT ACGGCTCGAT CACCTCCTAT CGCCGGAAAC CCATCCGGGG GGTTTAACCC CGGATGGGAT TTTGGTACTC CCCTTTATGG GGAGTCCCAC CGGCCGTACT CCTATAGCAA TTCCGG
16SBHB_4_6  GAACCT ACGGCTCGAT CACCTCCTAT CGCCGGAAAC CCATCCGGGG GGTTTAACCC CGGATGGGAT TTTGGTACTC CCCTTTATGG GGAGTCCCAC CGGCCGTACT CCTATAGCAA TTCCGG
16SBHB_9_2  GGAACCT ACGGCTCGAT CACCTCCTAT CGCCGGAAAC CCATCCGGGG GGTTTAACCC CGGATGGGAT TTTGGTACTC CCCTTTATGG GGAGTCCCAC CGGCCGTACT CCTATAGCAA TTCCG
16SBHB_5_4  GGGAACCT ACGGCTCGAT CACCTCCTAT CGCCGGAAAC CCATCCGGGG GGTTTAACCC CGGATGGGAT TTTGGTACTC CCCTTTATGG GGAGTCCCAC CGGCCGTACT CCTATAGCAA TTCCG
16SBHB_27_  CCT ACGGCTCGAT CACCTCCTAT CGCCGGAAAC CCATCCGGGG GGTTTAACCC CGGATGGGAT TTTGGTACTC CCCTTTATGG GGAGTCCCAC CGGCCGTACT CCTATAGCAA TTCCGG
16SBHB_12_  CCT ACGGCTCGAT CACCTCCTAT CGCCGGAAAC CCATCCGGGG GGTTTAACCC CGGATGGGAT TTTGGTACTC CCCTTTATGG GGAGTCCCAC CGGCCGTACT CCTATAGCAA TTCCGGT
16SBHB_6_4  CCT ACGGCTCGAT CACCTCCTAT CGCCGGAAAC CCATCCGGGG GGTTTAACCC CGGATGGGAT TTTGGTACTC CCCTTTATGG GGAGTCCCAC CGGCCGTACT CCTATAGCAA TTCCGGTT
16SBHB_12_  CCT ACGGCTCGAT CACCTCCTAT CGCCGGAAAC CCATCCGGGG GGTTTAACCC CGGATGGGAT TTTGGTACTC CCCTTTATGG GGAGTCCCAC CGGCCGTACT CCTATAGCAA TTCCG
16SBHB_3_8  T ACGGCTCGAT CACCTCCTAT CGCCGGAAAC CCATCCGGGG GGTTTAACCC CGGATGGGAT TTTGGTACTC CCCTTTATGG GGAGTCCCAC CGGCCGTACT CCTATAGCAA TTCCGGT
16SBHB_6_4  CT ACGGCTCGAT CACCTCCTAT CGCCGGAAAC CCATCCGGGG GGTTTAACCC CGGATGGGAT TTTGGTACTC CCCTTTATGG GGAGTCCCAC CGGCCGTACT CCTATAGCAA TTCCGG
16SBHB_12_  GTCGTAACA AGGTAGCCGT AGGGGAACCT ACGGCTCGAT CACCTCCTAT CGCCGGAAAC CCATCCGGGG GGTTTAACCC CGGATGGGAT TTTGGTACTC CCCTTTATGG GGAGTCCCAC CGGCCGTAC
16SBHB_3_8  GTCGTAACA AGGTAGCCGT AGGGGAACCT ACGGCTCGAT CACCTCCTAT CGCCGGAAAC CCATCCGGGG GGTTTAACCC CGGATGGGAT TTTGGTACTC CCCTTTATGG GGAGTCCCAC CGGCCGTACT
16SBHB_3_8  GTCGTAACA AGGTAGCCGT AGGGGAACCT ACGGCTCGAT CACCTCCTAT CGCCGGAAAC CCATCCGGGG GGTTTAACCC CGGATGGGAT TTTGGTACTC CCCTTTATGG GGAGTCCCAC CGGCCGTA
16SBHB_3_8  GTCGTAACA AGGTAGCCGT AGGGGAACCT ACGGCTCGAT CACCTCCTAT CGCCGGAAAC CCATCCGGGG GGTTTAACCC CGGATGGGAT TTTGGTACTC CCCTTTATGG GGAGTCCCAC CGGCC
16SBHB_3_8  GGGAGA AGTCGTAACA AGGTAGCCGT AGGGGAACCT ACGGCTCGAT CACCTCCTAT CGCCGGAAAC CCATCCGGGG GGTTTAACCC CGGATGGGAT TTTGGTACTC CCCTTTATGG GGAGTCCCAC CGGC
16SBHB_8_2  GGTAGCCGT AGGGGAACCT ACGGCTCGAT CACCTCCTAT CGCCGGAAAC CCATCCGGGG GGTTTAACCC CGGATGGGAT TTTGGTACTC CCCTTTATGG GGAGTCCCAC CGGCCGTACT CCTATA
16SBHB_4_7  CA AGGTAGCCGT AGGGGAACCT ACGGCTCGAT CACCTCCTAT CGCCGGAAAC CCATCCGGGG GGTTTAACCC CGGATGGGAT TTTGGTACTC CCCTTTATGG GGAGTCCCAC CGGCCGTACT CCTATA
16SBHB_4_7  AACA AGGTAGCCGT AGGGGAACCT ACGGCTCGAT CACCTCCTAT CGCCGGAAAC CCATCCGGGG GGTTTAACCC CGGATGGGAT TTTGGTACTC CCCTTTATGG GGAGTCCCAC CGGCCGTACT CCTAT
16SBHB_4_6  GGTAGCCGT AGGGGAACCT ACGGCTCGAT CACCTCCTAT CGCCGGAAAC CCATCCGGGG GGTTTAACCC CGGATGGGAT TTTGGTACTC CCCTTTATGG GGAGTCCCAC CGGCCGTACT CCTAT
16SBHB_4_7  CA AGGTAGCCGT AGGGGAACCT ACGGCTCGAT CACCTCCTAT CGCCGGAAAC CCATCCGGGG GGTTTAACCC CGGATGGGAT TTTGGTACTC CCCTTTATGG GGAGTCCCAC CGGCCGTACT CC
16SBHB_4_6  GGTAGCCGT AGGGGAACCT ACGGCTCGAT CACCTCCTAT CGCCGGAAAC CCATCCGGGG GGTTTAACCC CGGATGGGAT TTTGGTACTC CCCTTTATGG GGAGTCCCAC CGGCCGTACT CCTA
16SBHB_3_8  GTAGCCGT AGGGGAACCT ACGGCTCGAT CACCTCCTAT CGCCGGAAAC CCATCCGGGG GGTTTAACCC CGGATGGGAT TTTGGTACTC CCCTTTATGG GGAGTCCCAC CGGCCGTACT CC
16SBHB_6_4  CTCCTAT CGCCGGAAAC CCATCCGGGG GGTTTAACCC CGGATGGGAT TTTGGTACTC CCCTTTATGG GGAGTCCCAC CGGCCGTACT CCTATAGCAA TTCCGGTTGA TCCTGCCGGA GGC
16SBHB_6_4  CGAT CACCTCCTAT CGCCGGAAAC CCATCCGGGG GGTTTAACCC CGGATGGGAT TTTGGTACTC CCCTTTATGG GGAGTCCCAC CGGCCGTACT CCTATAGCAA TTCCGGTTGA TCCTGCCGGA GGCCAC
16SBHB_5_4  CTCCTAT CGCCGGAAAC CCATCCGGGG GGTTTAACCC CGGATGGGAT TTTGGTACTC CCCTTTATGG GGAGTCCCAC CGGCCGTACT CCTATAGCAA TTCCGGTTGA TCCTGCCGGA GGCCAC
16SBHB_4_7  CACCTCCTAT CGCCGGAAAC CCATCCGGGG GGTTTAACCC CGGATGGGAT TTTGGTACTC CCCTTTATGG GGAGTCCCAC CGGCCGTACT CCTATAGCAA TTCCGGTTGA TCCTGCCGGA GG
16SBHB_4_6  CTCCTAT CGCCGGAAAC CCATCCGGGG GGTTTAACCC CGGATGGGAT TTTGGTACTC CCCTTTATGG GGAGTCCCAC CGGCCGTACT CCTATAGCAA TTCCGGTTGA TCCTGCCGGA GG
16SBHB_3_9  CTCCTAT CGCCGGAAAC CCATCCGGGG GGTTTAACCC CGGATGGGAT TTTGGTACTC CCCTTTATGG GGAGTCCCAC CGGCCGTACT CCTATAGCAA TTCCGGTTGA TCCTGCCGGA G
16SBHB_21_  GGGGAACCT ACGGCTCGAT CACCTCCTAT CGCCGGAAAC CCATCCGGGG GGTTTAACCC CGGATGGGAT TTTGGTACTC CCCTTTATGG GGAGTCCCAC CGGCCGTACT CCTATA
16SBHB_6_3  GGGGAACCT ACGGCTCGAT CACCTCCTAT CGCCGGAAAC CCATCCGGGG GGTTTAACCC CGGATGGGAT TTTGGTACTC CCCTTTATGG GGAGTCCCAC CGGCCGTACT CCTATAG
16SBHB_5_4  GGGGAACCT ACGGCTCGAT CACCTCCTAT CGCCGGAAAC CCATCCGGGG GGTTTAACCC CGGATGGGAT TTTGGTACTC CCCTTTATGG GGAGTCCCAC CGGCCGTACT CCTATAGCG
16SBHB_4_6  GGGAACCT ACGGCTCGAT CACCTCCTAT CGCCGGAAAC CCATCCGGGG GGTTTAACCC CGGATGGGAT TTTGGTACTC CCCTTTATGG GGAGTCCCAC CGGCCGTACT CCTATAG
16SBHB_4_7  CCT ACGGCTCGAT CACCTCCTAT CGCCGGAAAC CCATCCGGGG GGTTTAACCC CGGATGGGAT TTTGGTACTC CCCTTTATGG GGAGTCCCAC CGGCCGTACT CCTATAGCAA TT
16SBHB_16_  CGGCTCGAT CACCTCCTAT CGCCGGAAAC CCATCCGGGG GGTTTAACCC CGGATGGGAT TTTGGTACTC CCCTTTATGG GGAGTCCCAC CGGCCGTACT CCTATAGCAA TTCCGG
16SBHB_7_3  CGGCTCGAT CACCTCCTAT CGCCGGAAAC CCATCCGGGG GGTTTAACCC CGGATGGGAT TTTGGTACTC CCCTTTATGG GGAGTCCCAC CGGCCGTACT CCTATAGCAA TTCCGGTT
16SBHB_3_9  CGGCTCGAT CACCTCCTAT CGCCGGAAAC CCATCCGGGG GGTTTAACCC CGGATGGGAT TTTGGTACTC CCCTTTATGG GGAGTCCCAC CGGCCGTACT CCTATAGCAA TTCCGGTC
16SBHB_3_9  CGGCTCGAT CACCTCCTAT CGCCGGAAAC CCATCCGGGG GGTTTAACCC CGGATGGGAT TTTGGTACTC CCCTTTATGG GGAGTCCCAC CGGCCGTACT CCTATAGCAA TTCCGGTG
16SBHB_11_  CGGCTCGAT CACCTCCTAT CGCCGGAAAC CCATCCGGGG GGTTTAACCC CGGATGGGAT TTTGGTACTC CCCTTTATGG GGAGTCCCAC CGGCCGTACT CCTATAGCAA TTCCG
16SBHB_20_  CGT AGGGGAACCT ACGGCTCGAT CACCTCCTAT CGCCGGAAAC CCATCCGGGG GGTTTAACCC CGGATGGGAT TTTGGTACTC CCCTTTATGG GGAGTCCCAC CGGCCGTACT CCTAT
16SBHB_18_  CGT AGGGGAACCT ACGGCTCGAT CACCTCCTAT CGCCGGAAAC CCATCCGGGG GGTTTAACCC CGGATGGGAT TTTGGTACTC CCCTTTATGG GGAGTCCCAC CGGCCGTACT CCTATA
16SBHB_3_9  CGT AGGGGAACCT ACGGCTCGAT CACCTCCTAT CGCCGGAAAC CCATCCGGGG GGTTTAACCC CGGATGGGAT TTTGGTACTC CCCTTTATGG GGAGTCCCAC CGGCCGTACT CCTATAG
16SBHB_14_  CGT AGGGGAACCT ACGGCTCGAT CACCTCCTAT CGCCGGAAAC CCATCCGGGG GGTTTAACCC CGGATGGGAT TTTGGTACTC CCCTTTATGG GGAGTCCCAC CGGCCGTACT CCTA
16SBHB_16_  CGT AGGGGAACCT ACGGCTCGAT CACCTCCTAT CGCCGGAAAC CCATCCGGGG GGTTTAACCC CGGATGGGAT TTTGGTACTC CCCTTTATGG GGAGTCCCAC CGGCCGTACT CCT
16SBHB_6_4  AGGGGAACCT ACGGCTCGAT CACCTCCTAT CGCCGGAAAC CCATCCGGGG GGTTTAACCC CGGATGGGAT TTTGGTACTC CCCTTTATGG GGAGTCCCAC CGGCCGTACT CCTAT
16SBHB_11_  CGT AGGGGAACCT ACGGCTCGAT CACCTCCTAT CGCCGGAAAC CCATCCGGGG GGTTTAACCC CGGATGGGAT TTTGGTACTC CCCTTTATGG GGAGTCCCAC CGGCCGTACT CC
16SBHB_15_  GGGGAACCT ACGGCTCGAT CACCTCCTAT CGCCGGAAAC CCATCCGGGG GGTTTAACCC CGGATGGGAT TTTGGTACTC CCCTTTATGG GGAGTCCCAC CGGCCGTACT CCTAT
16SBHB_8_2  GGAACCT ACGGCTCGAT CACCTCCTAT CGCCGGAAAC CCATCCGGGG GGTTTAACCC CGGATGGGAT TTTGGTACTC CCCTTTATGG GGAGTCCCAC CGGCCGTACT CCTATAG
16SBHB_5_4  GGAACCT ACGGCTCGAT CACCTCCTAT CGCCGGAAAC CCATCCGGGG GGTTTAACCC CGGATGGGAT TTTGGTACTC CCCTTTATGG GGAGTCCCAC CGGCCGTACT CCTATA
16SBHB_5_4  GGGAACCT ACGGCTCGAT CACCTCCTAT CGCCGGAAAC CCATCCGGGG GGTTTAACCC CGGATGGGAT TTTGGTACTC CCCTTTATGG GGAGTCCCAC CGGCCGTACT CCTATA
16SBHB_7_3  GGGAACCT ACGGCTCGAT CACCTCCTAT CGCCGGAAAC CCATCCGGGG GGTTTAACCC CGGATGGGAT TTTGGTACTC CCCTTTATGG GGAGTCCCAC CGGCCGTACT CCTAT
16SBHB_4_6  GGAACCT ACGGCTCGAT CACCTCCTAT CGCCGGAAAC CCATCCGGGG GGTTTAACCC CGGATGGGAT TTTGGTACTC CCCTTTATGG GGAGTCCCAC CGGCCGTACT CCTAT
16SBHB_8_3  CCT ACGGCTCGAT CACCTCCTAT CGCCGGAAAC CCATCCGGGG GGTTTAACCC CGGATGGGAT TTTGGTACTC CCCTTTATGG GGAGTCCCAC CGGCCGTACT CCTATAGCA
16SBHB_3_9  CT ACGGCTCGAT CACCTCCTAT CGCCGGAAAC CCATCCGGGG GGTTTAACCC CGGATGGGAT TTTGGTACTC CCCTTTATGG GGAGTCCCAC CGGCCGTACT CCTATAGCA
16SBHB_4_7  CCT ACGGCTCGAT CACCTCCTAT CGCCGGAAAC CCATCCGGGG GGTTTAACCC CGGATGGGAT TTTGGTACTC CCCTTTATGG GGAGTCCCAC CGGCCGTACT CCTATAGCG
16SBHB_3_7  TGAGCC TTTGGAGCTC TGACCCGGAG AAAGGTGTGA AGACAGGT-- ---------- ---------- -------GGG TTCCTCCTAT CGCCGGAAAC CCATCCGGGG GGTTTAACCC CGGATGGGAT TTTGGTACTC CCCTTTATGG GGAGTCCCAC CGGCCGTACT CCTATAGCAA TTCCGGTC
16SBHB_9_2  GGGGAACCT ACGGCTCGAT CACCTCCTAT CGCCGGAAAC CCATCCGGGG GGTTTAACCC CGGATGGGAT TTTGGTACTC CCCTTTATGG GGAGTCCCAC CGGCCGTACT CC
16SBHB_8_2  GGGGAACCT ACGGCTCGAT CACCTCCTAT CGCCGGAAAC CCATCCGGGG GGTTTAACCC CGGATGGGAT TTTGGTACTC CCCTTTATGG GGAGTCCCAC CGGCCGTACT CCT
16SBHB_5_5  AGGGGAACCT ACGGCTCGAT CACCTCCTAT CGCCGGAAAC CCATCCGGGG GGTTTAACCC CGGATGGGAT TTTGGTACTC CCCTTTATGG GGAGTCCCAC CGGCCGTACT CC
16SBHB_3_8  GGGGAACCT ACGGCTCGAT CACCTCCTAT CGCCGGAAAC CCATCCGGGG GGTTTAACCC CGGATGGGAT TTTGGTACTC CCCTTTATGG GGAGTCCCAC CGGCCGTACT CCC
16SBHB_9_2  CGT AGGGGAACCT ACGGCTCGAT CACCTCCTAT CGCCGGAAAC CCATCCGGGG GGTTTAACCC CGGATGGGAT TTTGGTACTC CCCTTTATGG GGAGTCCCAC CGGCCGTAC
16SBHB_4_6  CGT AGGGGAACCT ACGGCTCGAT CACCTCCTAT CGCCGGAAAC CCATCCGGGG GGTTTAACCC CGGATGGGAT TTTGGTACTC CCCTTTATGG GGAGTCCCAC CGGCCGTACT
16SBHB_9_2  GGAACCT ACGGCTCGAT CACCTCCTAT CGCCGGAAAC CCATCCGGGG GGTTTAACCC CGGATGGGAT TTTGGTACTC CCCTTTATGG GGAGTCCCAC CGGCCGTACT CCT
16SBHB_7_3  GGAACCT ACGGCTCGAT CACCTCCTAT CGCCGGAAAC CCATCCGGGG GGTTTAACCC CGGATGGGAT TTTGGTACTC CCCTTTATGG GGAGTCCCAC CGGCCGTACT CCTA
16SBHB_3_9  CGGCTCGAT CACCTCCTAT CGCCGGAAAC CCATCCGGGG GGTTTAACCC CGGATGGGAT TTTGGTACTC CCCTTTATGG GGAGTCCCAC CGGCCGTACT CCTATAGCAA TT
16SBHB_3_8  GGTAGCCGT AGGGGAACCT ACGGCTCGAT CACCTCCTAT CGCCGGAAAC CCATCCGGGG GGTTTAACCC CGGATGGGAT TTTGGTACTC CCCTTTATGG GGAGTCCCAC CGGCCG
16SBHB_25_  CCT ACGGCTCGAT CACCTCCTAT CGCCGGAAAC CCATCCGGGG GGTTTAACCC CGGATGGGAT TTTGGTACTC CCCTTTATGG GGAGTCCCAC CGGCCGTACT CCTATA
16SBHB_12_  CCT ACGGCTCGAT CACCTCCTAT CGCCGGAAAC CCATCCGGGG GGTTTAACCC CGGATGGGAT TTTGGTACTC CCCTTTATGG GGAGTCCCAC CGGCCGTACT CCTATAG
16SBHB_4_6  CT ACGGCTCGAT CACCTCCTAT CGCCGGAAAC CCATCCGGGG GGTTTAACCC CGGATGGGAT TTTGGTACTC CCCTTTATGG GGAGTCCCAC CGGCCGTACT CCTATAGCG
16SBHB_3_9  CT ACGGCTCGAT CACCTCCTAT CGCCGGAAAC CCATCCGGGG GGTTTAACCC CGGATGGGAT TTTGGTACTC CCCTTTATGG GGAGTCCCAC CGGCCGTACT CCTATAG
16SBHB_19_  CGAT CACCTCCTAT CGCCGGAAAC CCATCCGGGG GGTTTAACCC CGGATGGGAT TTTGGTACTC CCCTTTATGG GGAGTCCCAC CGGCCGTACT CCTATAGCAA TTCCGG
16SBHB_6_4  CGAT CACCTCCTAT CGCCGGAAAC CCATCCGGGG GGTTTAACCC CGGATGGGAT TTTGGTACTC CCCTTTATGG GGAGTCCCAC CGGCCGTACT CCTATAGCAA TTCCGGT
16SBHB_6_4  CGAT CACCTCCTAT CGCCGGAAAC CCATCCGGGG GGTTTAACCC CGGATGGGAT TTTGGTACTC CCCTTTATGG GGAGTCCCAC CGGCCGTACT CCTATAGCAA TTCCG
16SBHB_3_9  GAACCT ACGGCTCGAT CACCTCCTAT CGCCGGAAAC CCATCCGGGG GGTTTAACCC CGGATGGGAT TTTGGTACTC CCCTTTATGG GGAGTCCCAC CGGCCGTACT CCT
16SBHB_16_  CCT ACGGCTCGAT CACCTCCTAT CGCCGGAAAC CCATCCGGGG GGTTTAACCC CGGATGGGAT TTTGGTACTC CCCTTTATGG GGAGTCCCAC CGGCCGTACT CCTAT
16SBHB_7_3  CT ACGGCTCGAT CACCTCCTAT CGCCGGAAAC CCATCCGGGG GGTTTAACCC CGGATGGGAT TTTGGTACTC CCCTTTATGG GGAGTCCCAC CGGCCGTACT CCTATA
16SBHB_4_6  GGGGAACCT ACGGCTCGAT CACCTCCTAT CGCCGGAAAC CCATCCGGGG GGTTTAACCC CGGATGGGAT TTTGGTACTC CCCTTTATGG GGAGTCCCAC CGGCCGTAC
16SBHB_6_4  CGGCTCGAT CACCTCCTAT CGCCGGAAAC CCATCCGGGG GGTTTAACCC CGGATGGGAT TTTGGTACTC CCCTTTATGG GGAGTCCCAC CGGCCGTACT CCTATAG
16SBHB_3_9  CGGCTCGAT CACCTCCTAT CGCCGGAAAC CCATCCGGGG GGTTTAACCC CGGATGGGAT TTTGGTACTC CCCTTTATGG GGAGTCCCAC CGGCCGTACT CCTATAGCG
16SBHB_5_4  CGT AGGGGAACCT ACGGCTCGAT CACCTCCTAT CGCCGGAAAC CCATCCGGGG GGTTTAACCC CGGATGGGAT TTTGGTACTC CCCTTTATGG GGAGTCCCAC CGGCCG
16SBHB_8_3  CCT ACGGCTCGAT CACCTCCTAT CGCCGGAAAC CCATCCGGGG GGTTTAACCC CGGATGGGAT TTTGGTACTC CCCTTTATGG GGAGTCCCAC CGGCCGTACT CC
16SBHB_7_3  CCT ACGGCTCGAT CACCTCCTAT CGCCGGAAAC CCATCCGGGG GGTTTAACCC CGGATGGGAT TTTGGTACTC CCCTTTATGG GGAGTCCCAC CGGCCGTACT CCT
16SBHB_4_7  CCT ACGGCTCGAT TACCTCCTAT CGCCGGAAAC CCATCCGGGG GGTTTAACCC CGGATGGGAT TTTGGTACTC CCCTTTATGG GGAGTCCCAC CGGCCGTACT CCT
16SBHB_5_4  CT ACGGCTCGAT CACCTCCTAT CGCCGGAAAC CCATCCGGGG GGTTTAACCC CGGATGGGAT TTTGGTACTC CCCTTTATGG GGAGTCCCAC CGGCCGTACT CCTA
16SBHB_3_8  GTCC GGGCTCCGTG AGGGGGGAGA AGTCGTAACA AGGTAGCCGT AGGGGAACCT ACGGCTCGAT CACCTCCTAT CGCCGGAAAC CCATCCGGGG GGTTTAACCC CGGATGGGAT TTTGGTACTC CC
16SBHB_10_  CACCTCCTAT CGCCGGAAAC CCATCCGGGG GGTTTAACCC CGGATGGGAT TTTGGTACTC CCCTTTATGG GGAGTCCCAC CGGCCGTACT CCTATAGCAA TTCCGG
16SBHB_4_7  CACCTCCTAT CGCCGGAAAC CCATCCGGGG GGTTTAACCC CGGATGGGAT TTTGGTACTC CCCTTTATGG GGAGTCCCAC CGGCCGTACT CCTATAGCAA TTCCG
16SBHB_7_3  CTCCTAT CGCCGGAAAC CCATCCGGGG GGTTTAACCC CGGATGGGAT TTTGGTACTC CCCTTTATGG GGAGTCCCAC CGGCCGTACT CCTATAGCAA TTCCGGT
16SBHB_3_9  CTCCTAT CGCCGGAAAC CCATCCGGGG GGTTTAACCC CGGATGGGAT TTTGGTACTC CCCTTTATGG GGAGTCCCAC CGGCCGTACT CCTATAGCAA TTCCGGTT
16SBHB_10_  CGGCTCGAT CACCTCCTAT CGCCGGAAAC CCATCCGGGG GGTTTAACCC CGGATGGGAT TTTGGTACTC CCCTTTATGG GGAGTCCCAC CGGCCGTACT CCTATA
16SBHB_4_6  CGGCTCGAT CACCTCCTAT CGCCGGAAAC CCATCCGGGG GGTTTAACCC CGGATGGGAT TTTGGTACTC CCCTTTATGG GGAGTCCCAC CGGCCGTACT CCTAT
16SBHB_3_8  GTCGTAACA AGGTAGCCGT AGGGGAACCT ACGGCTCGAT CACCTCCTAT CGCCGGAAAC CCATCCGGGG GGTTTAACCC CGGATGGGAT TTTGGTACTC CCCTTTATGG GGAG
16SBHB_4_6  CGAT CACCTCCTAT CGCCGGAAAC CCATCCGGGG GGTTTAACCC CGGATGGGAT TTTGGTACTC CCCTTTATGG GGAGTCCCAC CGGCCGTACT CCTATAGCA
16SBHB_4_7  CCT ACGGCTCGAT CACCTCCTAT CGCCGGAAAC CCATCCGGGG GGTTTAACCC CGGATGGGAT TTTGGTACTC CCCTTTATGG GGAGTCCCAC CGGCCGTACT
16SBHB_5_5  CGGCTCGAT CACCTCCTAT CGCCGGAAAC CCATCCGGGG GGTTTAACCC CGGATGGGAT TTTGGTACTC CCCTTTATGG GGAGTCCCAC CGGCCGTACT CCTA
16SBHB_3_9  CGGCTCGAT CACCTCCTAT CGCCGGAAAC CCATCCGGGG GGTTTAACCC CGGATGGGAT TTTGGTACTC CCCTTTATGG GGAGTCCCAC CGGCCGTACT CCGAT
16SBHB_4_7  CCT ACGGCTCGAT CACCTCCTAT CGCCGGAAAC CCATCCGGGG GGTTTAACCC CGGATGGGAT TTTGGTACTC CCCTTTATGG GGAGTCCCAC CGGCCGTAC
16SBHB_10_  CTCCTAT CGCCGGAAAC CCATCCGGGG GGTTTAACCC CGGATGGGAT TTTGGTACTC CCCTTTATGG GGAGTCCCAC CGGCCGTACT CCTATAGCAA TTCCGG
16SBHB_3_9  CTCCTAT CGCCGGAAAC CCATCCGGGG GGTTTAACCC CGGATGGGAT TTTGGTACTC CCCTTTATGG GGAGTCCCAC CGGCCGTACT CCTATAGCAA TTCCGGC
16SBHB_3_9  CTCGAT CACCTCCTAT CGCCGGAAAC CCATCCGGGG GGTTTAACCC CGGATGGGAT TTTGGTACTC CCCTTTATGG GGAGTCCCAC CGGCCGTACT CCTATA
16SBHB_5_5  CCTAT CGCCGGAAAC CCATCCGGGG GGTTTAACCC CGGATGGGAT TTTGGTACTC CCCTTTATGG GGAGTCCCAC CGGCCGTACT CCTATAGCAA TTCCGG
16SBHB_3_9  CTAT CGCCGGAAAC CCATCCGGGG GGTTTAACCC CGGATGGGAT TTTGGTACTC CCCTTTATGG GGAGTCCCAC CGGCCGTACT CCTATAGCAA TTCCGGT
16SBHB_5_5  CGAT CACCTCCTAT CGCCGGAAAC CCATCCGGGG GGTTTAACCC CGGATGGGAT TTTGGTACTC CCCTTTATGG GGAGTCCCAC CGGCCGTACT CCTATA
16SBHB_4_6  CGGCTCGAT CACCTCCTAT CGCCGGAAAC CCATCCGGGG GGTTTAACCC CGGATGGGAT TTTGGTACTC CCCTTTATGG GGAGTCCCAC CGGCCGTACT CC
16SBHB_6_4  CGAT CACCTCCTAT CGCCGGAAAC CCATCCGGGG GGTTTAACCC CGGATGGGAT TTTGGTACTC CCCTTTATGG GGAGTCCCAC CGGCCGTACT CCTAT
16SBHB_4_6  CGAT CACCTCCTAT CGCCGGAAAC CCATCCGGGG GGTTTAACCC CGGATGGGAT TTTGGTACTC CCCTTTATGG GGAGTCCCAC CGGCCGTACT CCTA
16SBHB_3_9  CTCGAT CACCTCCTAT CGCCGGAAAC CCATCCGGGG GGTTTAACCC CGGATGGGAT TTTGGTACTC CCCTTTATGG GGAGTCCCAC CGGCCGTACT CCT
16SBHB_6_4  CACCTCCTAT CGCCGGAAAC CCATCCGGGG GGTTTAACCC CGGATGGGAT TTTGGTACTC CCCTTTATGG GGAGTCCCAC CGGCCGTACT CCTATAGCG
```

```
16SBHB_5_5                                                                             CGAT CACCTCCTAT CGCCGGAAAC CCATCCGGGG GGTTTAACCC CGGATGGGAT TTTGGTACTC CCCTTTATGG GGAGTCCCAC CGGCCGTACT CCT
16SBHB_4_6                                                                             CGAT CACCTCCTAT CGCCGGAAAC CCATCCGGGG GGTTTAACCC CGGATGGGAT TTTGGTACTC CCCTTTATGG GGAGTCCCAC CGGCCGTACT CC
16SBHB_4_6                                                      CGT AGGGGAACCT ACGGCTCGAT CACCTCCTAT CGCCGGAAAC CCATCCGGGG GGTTTAACCC CGGATGGGAT TTTGGTACTC CCCTTTATGG GGAG
16SBHB_4_6                                                                                       CTCCTAT CGCCGGAAAC CCATCCGGGG GGTTTAACCC CGGATGGGAT TTTGGTACTC CCCTTTATGG GGAGTCCCAC CGGCCGTACT CCTATA
16SBHB_3_8                                           GTCGTAACA AGGTAGCCGT AGGGGAACCT ACGGCTCGAT CACCTCCTAT CGCCGGAAAC CCATCCGGGG GGTTTAACCC CGGATGGGAT TTTGGTACTC CCC
16SBHB_7_3                                                                                       CTCCTAT CGCCGGAAAC CCATCCGGGG GGTTTAACCC CGGATGGGAT TTTGGTACTC CCCTTTATGG GGAGTCCCAC CGGCCGTACT CCTAT
16SBHB_4_7                                                                                         CCTAT CGCCGGAAAC CCATCCGGGG GGTTTAACCC CGGATGGGAT TTTGGTACTC CCCTTTATGG GGAGTCCCAC CGGCCGTACT CCTAT
16SBHB_4_7                                                                                         CCTAT CGCCGGAAAC CCATCCGGGG GGTTTAACCC CGGATGGGAT TTTGGTACTC CCCTTTATGG GGAGTCCCAC CGGCCGTACT CCTA
 Consensus ...... .......... .......... .......... .......... .......... .......... .......... .......... .......... .......... .......... .......cgt aggggaacct acggctcgat CACCTCCTAT CGCCGGAAAC CCATCCGGGG GGTTTAACCC CGGATGGGAT TTTGGTACTC CCCTTTATGG GGAGTCCCAC CGGCCGTACT CCTatagcaa ttccggttga tcctgccgga ggccactgct atgggggtcc gactaagc.. .......... .......... .......... .......... .......... ..........
```
