## Supplementary Figure S2b for "Permuted 23S rRNA is integrated in 50S ribosome particles in *Thermococcus barophilus*"

```
1                                                                                                                                                                                                                          397
23SBHB_440      CGCAGAAGA CGGGTTTGAT GGGGCGGGGA TGTAAGCGGG AAGGGAAACC GACCCGTTTA GTCTGCCGCT CCCAACAGCC CGAGTTTCTT GCCCCTGAAA GAGGGGCTGG GCATTTGATA TGCCCAGCCC CTTGAGTAAG GGGCAAGAAA CCTAAGCCGC CTGGTGGATG GCTCGGCTCG GGGCGCTGAC GAAGGG
23SBHB_229      CGCAGAAGA CGGGTTTGAT GGGGCGGGGA TGTAAGCGGG AAGGGAAACC GACCCGTTTA GTCTGCCGCT CCCAACAGCC CGAGTTTCTT GCCCCTGAAA GAGGGGCTGG GCATTTGATA TGCCCAGCCC CTTGAGTAAG GGGCAAGAAA CCTAAGCCGC CTGGTGGATG GCTCGGCTCG GGGCGCTGAC GAAGGGC
23SBHB_421      CGCAGAAGA CGGGTTTGAT GGGGCGGGGA TGTAAGCGGG AAGGGAAACC GACCCGTTTA GTCTGCCGCT CCCAACAGCC CGAGTTTCTT GCCCCTGAAA GAGGGGCTGG GCATTTGATA TGCCCAGCCC CTTGAGTAAG GGGCAAGAAA CCTAAGCCGC CTGGTGGATG GCTCGGCTCG GGGCGCTGAC GAAGGGCC
23SBHB_185     CCGCAGAAGA CGGGTTTGAT GGGGCGGGGA TGTAAGCGGG AAGGGAAACC GACCCGTTTA GTCTGCCGCT CCCAACAGCC CGAGTTTCTT GCCCCTGAAA GAGGGGCTGG GCATTTGATA TGCCCAGCCC CTTGAGTAAG GGGCAAGAAA CCTAAGCCGC CTGGTGGATG GCTCGGCTCG GGGCGCTGAC GAAGGGCC
23SBHB_198     CCGCAGAAGA CGGGTTTGAT GGGGCGGGGA TGTAAGCGGG AAGGGAAACC GACCCGTTTA GTCTGCCGCT CCCAACAGCC CGAGTTTCTT GCCCCTGAAA GAGGGGCTGG GCATTTGATA TGCCCAGCCC CTTGAGTAAG GGGCAAGAAA CCTAAGCCGC CTGGTGGATG GCTCGGCTCG GGGCGCTGAC GAAGGG
23SBHB_558     CCGCAGAAGA CGGGTTTGAT GGGGCGGGGA TGTAAGCGGG AAGGGAAACC GACCCGTTTA GTCTGCCGCT CCCAACAGCC CGAGTTTCTT GCCCCTGAAA GAGGGGCTGG GCATTTGATA TGCCCAGCCC CTTGAGTAAG GGGCAAGAAA CCTAAGCCGC CTGGTGGATG GCTCGGCTCG GGGCGCTGAC GAAGGGG
23SBHB_492   C CCGCAGAAGA CGGGTTTGAT GGGGCGGGGA TGTAAGCGGG AAGGGAAACC GACCCGTTTA GTCTGCCGCT CCCAACAGCC CGAGTTTCTT GCCCCTGAAA GAGGGGCTGG GCATTTGATA TGCCCAGCCC CTTGAGTAAG GGGCAAGAAA CCTAAGCCGC CTGGTGGATG GCTCGGCTCG GGGCGCTGAC GAAGGGG
23SBHB_176  GC CCGCAGAAGA CGGGTTTGAT GGGGCGGGGA TGTAAGCGGG AAGGGAAACC GACCCGTTTA GTCTGCCGCT CCCAACAGCC CGAGTTTCTT GCCCCTGAAA GAGGGGCTGG GCATTTGATA TGCCCAGCCC CTTGAGTAAG GGGCAAGAAA CCTAAGCCGC CTGGTGGATG GCTCGGCTCG GGGCGCTGAC GAAGGG
23SBHB_114   C CCGCAGAAGA CGGGTTTGAT GGGGCGGGGA TGTAAGCGGG AAGGGAAACC GACCCGTTTA GTCTGCCGCT CCCAACAGCC CGAGTTTCTT GCCCCTGAAA GAGGGGCTGG GCATTTGATA TGCCCAGCCC CTTGAGTAAG GGGCAAGAAA CCTAAGCCGC CTGGTGGATG GCTCGGCTCG GGGCGCTGAC GAAGG
23SBHB_161 CGC CCGCAGAAGA CGGGTTTGAT GGGGCGGGGA TGTAAGCGGG AAGGGAAACC GACCCGTTTA GTCTGCCGCT CCCAACAGCC CGAGTTTCTT GCCCCTGAAA GAGGGGCTGG GCATTTGATA TGCCCAGCCC CTTGAGTAAG GGGCAAGAAA CCTAAGCCGC CTGGTGGATG GCTCGGCTCG GGGCGCTGAC GAAG
23SBHB_137  GC CCGCAGAAGA CGGGTTTGAT GGGGCGGGGA TGTAAGCGGG AAGGGAAACC GACCCGTTTA GTCTGCCGCT CCCAACAGCC CGAGTTTCTT GCCCCTGAAA GAGGGGCTGG GCATTTGATA TGCCCAGCCC CTTGAGTAAG GGGCAAGAAA CCTAAGCCGC CTGGTGGATG GCTCGGCTCG GGGCGCTGAC GAAGG
23SBHB_129 CGC CCGCAGAAGA CGGGTTTGAT GGGGCGGGGA TGTAAGCGGG AAGGGAAACC GACCCGTTTA GTCTGCCGCT CCCAACAGCC CGAGTTTCTT GCCCCTGAAA GAGGGGCTGG GCATTTGATA TGCCCAGCCC CTTGAGTAAG GGGCAAGAAA CCTAAGCCGC CTGGTGGATG GCTCGGCTCG GGGCGCTGAC GAAGG
23SBHB_161   C CCGCAGAAGA CGGGTTTGAT GGGGCGGGGA TGTAAGCGGG AAGGGAAACC GACCCGTTTA GTCTGCCGCT CCCAACAGCC CGAGTTTCTT GCCCCTGAAA GAGGGGCTGG GCATTTGATA TGCCCAGCCC CTTGAGTAAG GGGCAAGAAA CCTAAGCCGC CTGGTGGATG GCTCGGCTCG GGGCGCTGAC GAAGAG
23SBHB_128     CCGCAGAAGA CGGGTTTGAT GGGGCGGGGA TGTAAGCGGG AAGGGAAACC GACCCGTTTA GTCTGCCGCT CCCAACAGCC CGAGTTTCTT GCCCCTGAAA GAGGGGCTGG GCATTTGATA TGCCCAGCCC CTTGAGTAAG GGGCAAGAAA CCTAAGCCGC CTGGTGGATG GCTCGGCTCG GGGCGCTGAC GAAGG
23SBHB_122   C CCGCAGAAGA CGGGTTTGAT GGGGCGGGGA TGTAAGCGGG AAGGGAAACC GACCCGTTTA GTCTGCCGCT CCCAACAGCC CGAGTTTCTT GCCCCTGAAA GAGGGGCTGG GCATTTGATA TGCCCAGCCC CTTGAGTAAG GGGCAAGAAA CCTAAGCCGC CTGGTGGATG GCTCGGCTCG GGGCGCTGAC GAAG
23SBHB_125      CGCAGAAGA CGGGTTTGAT GGGGCGGGGA TGTAAGCGGG AAGGGAAACC GACCCGTTTA GTCTGCCGCT CCCAACAGCC CGAGTTTCTT GCCCCTGAAA GAGGGGCTGG GCATTTGATA TGCCCAGCCC CTTGAGTAAG GGGCAAGAAA CCTAAGCCGC CTGGTGGATG GCTCGGCTCG GGGCGCTGAC GAAGGGG
23SBHB_176     CCGCAGAAGA CGGGTTTGAT GGGGCGGGGA TGTAAGCGGG AAGGGAAACC GACCCGTTTA GTCTGCCGCT CCCAACAGCC CGAGTTTCTT GCCCCTGAAA GAGGGGCTGG GCATTTGATA TGCCCAGCCC CTTGAGTAAG GGGCAAGAAA CCTAAGCCGC CTGGTGGATG GCTCGGCTCG GGGCGCTGAC GAAGAG
23SBHB_170      CGCAGAAGA CGGGTTTGAT GGGGCGGGGA TGTAAGCGGG AAGGGAAACC GACCCGTTTA GTCTGCCGCT CCCAACAGCC CGAGTTTCTT GCCCCTGAAA GAGGGGCTGG GCATTTGATA TGCCCAGCCC CTTGAGTAAG GGGCAAGAAA CCTAAGCCGC CTGGTGGATG GCTCGGCTCG GGGCGCTGAC GAAGGGCA
23SBHB_143      CGCAGAAGA CGGGTTTGAT GGGGCGGGGA TGTAAGCGGG AAGGGAAACC GACCCGTTTA GTCTGCCGCT CCCAACAGCC CGAGTTTCTT GCCCCTGAAA GAGGGGCTGG GCATTTGATA TGCCCAGCCC CTTGAGTAAG GGGCAAGAAA CCTAAGCCGC CTGGTGGATG GCTCGGCTCG GGGCGCTGAC GAAGGGCT
23SBHB_118  GC CCGCAGAAGA CGGGTTTGAT GGGGCGGGGA TGTAAGCGGG AAGGGAAACC GACCCGTTTA GTCTGCCGCT CCCAACAGCC CGAGTTTCTT GCCCCTGAAA GAGGGGCTGG GCATTTGATA TGCCCAGCCC CTTGAGTAAG GGGCAAGAAA CCTAAGCCGC CTGGTGGATG GCTCGGCTCG GGGCGCTGAC GAAG
23SBHB_280      CGCAGAAGA CGGGTTTGAT GGGGCGGGGA TGTAAGCGGG AAGGGAAACC GACCCGTTTA GTCTGCCGCT CCCAACAGCC CGAGTTTCTT GCCCCTGAAA GAGGGGCTGG GCATTTGATA TGCCCAGCCC CTTGAGTAAG GGGCAAGAAA CCTAAGCCGC CTGGTGGATG GCTCGGCTCG GGGCGCTGAC GAAGG
23SBHB_185   C CCGCAGAAGA CGGGTTTGAT GGGGCGGGGA TGTAAGCGGG AAGGGAAACC GACCCGTTTA GTCTGCCGCT CCCAACAGCC CGAGTTTCTT GCCCCTGAAA GAGGGGCTGG GCATTTGATA TGCCCAGCCC CTTGAGTAAG GGGCAAGAAA CCTAAGCCGC CTGGTGGATG GCTCGGCTCG GGGCGCTGAC GAAGGG
23SBHB_985   C CCGCAGAAGA CGGGTTTGAT GGGGCGGGGA TGTAAGCGGG AAGGGAAACC GACCCGTTTA GTCTGCCGCT CCCAACAGCC CGAGTTTCTT GCCCCTGAAA GAGGGGCTGG GCATTTGATA TGCCCAGCCC CTTGAGTAAG GGGCAAGAAA CCTAAGCCGC CTGGTGGATG GCTCGGCTCG GGGCGCTGAC GAAGGGC
23SBHB_493   C CCGCAGAAGA CGGGTTTGAT GGGGCGGGGA TGTAAGCGGG AAGGGAAACC GACCCGTTTA GTCTGCCGCT CCCAACAGCC CGAGTTTCTT GCCCCTGAAA GAGGGGCTGG GCATTTGATA TGCCCAGCCC CTTGAGTAAG GGGCAAGAAA CCTAAGCCGC CTGGTGGATG GCTCGGCTCG GGGCGCTGAC GAAGGGCG
23SBHB_558     CCGCAGAAGA CGGGTTTGAT GGGGCGGGGA TGTAAGCGGG AAGGGAAACC GACCCGTTTA GTCTGCCGCT CCCAACAGCC CGAGTTTCTT GCCCCTGAAA GAGGGGCTGG GCATTTGATA TGCCCAGCCC CTTGAGTAAG GGGCAAGAAA CCTAAGCCGC CTGGTGGATG GCTCGGCTCG GGGCGCTGAC GAAGGGCG
23SBHB_475      CGCAGAAGA CGGGTTTGAT GGGGCGGGGA TGTAAGCGGG AAGGGAAACC GACCCGTTTA GTCTGCCGCT CCCAACAGCC CGAGTTTCTT GCCCCTGAAA GAGGGGCTGG GCATTTGATA TGCCCAGCCC CTTGAGTAAG GGGCAAGAAA CCTAAGCCGC CTGGTGGATG GCTCGGCTCG GGGCGCTGAC GAAGGGCGTG
23SBHB_209     CCGCAGAAGA CGGGTTTGAT GGGGCGGGGA TGTAAGCGGG AAGGGAAACC GACCCGTTTA GTCTGCCGCT CCCAACAGCC CGAGTTTCTT GCCCCTGAAA GAGGGGCTGG GCATTTGATA TGCCCAGCCC CTTGAGTAAG GGGCAAGAAA CCTAAGCCGC CTGGTGGATG GCTCGGCTCG GGGCGCTGAC GAAGGGCGTG
23SBHB_131     CCGCAGAAGA CGGGTTTGAT GGGGCGGGGA TGTAAGCGGG AAGGGAAACC GACCCGTTTA GTCTGCCGCT CCCAACAGCC CGAGTTTCTT GCCCCTGAAA GAGGGGCTGG GCATTTGATA TGCCCAGCCC CTTGAGTAAG GGGCAAGAAA CCTAAGCCGC CTGGTGGATG GCTCGGCTCG GGGCGCTGAC GAAGGGCGT
23SBHB_141   C CCGCAGAAGA CGGGTTTGAT GGGGCGGGGA TGTAAGCGGG AAGGGAAACC GACCCGTTTA GTCTGCCGCT CCCAACAGCC CGAGTTTCTT GCCCCTGAAA GAGGGGCTGG GCATTTGATA TGCCCAGCCC CTTGAGTAAG GGGCAAGAAA CCTAAGCCGC CTGGTGGATG GCTCGGCTCG GGGCGCTGAC GAAGGGCC
23SBHB_243      CGCAGAAGA CGGGTTTGAT GGGGCGGGGA TGTAAGCGGG AAGGGAAACC GACCCGTTTA GTCTGCCGCT CCCAACAGCC CGAGTTTCTT GCCCCTGAAA GAGGGGCTGG GCATTTGATA TGCCCAGCCC CTTGAGTAAG GGGCAAGAAA CCTAAGCCGC CTGGTGGATG GCTCGGCTCG GGGCGCTGAC GAAGGGCGT
23SBHB_205      CGCAGAAGA CGGGTTTGAT GGGGCGGGGA TGTAAGCGGG AAGGGAAACC GACCCGTTTA GTCTGCCGCT CCCAACAGCC CGAGTTTCTT GCCCCTGAAA GAGGGGCTGG GCATTTGATA TGCCCAGCCC CTTGAGTAAG GGGCAAGAAA CCTAAGCCGC CTGGTGGATG GCTCGGCTCG GGGCGCTGAC GAAGGGCGTA G
23SBHB_180 CGC CCGCAGAAGA CGGGTTTGAT GGGGCGGGGA TGTAAGCGGG AAGGGAAACC GACCCGTTTA GTCTGCCGCT CCCAACAGCC CGAGTTTCTT GCCCCTGAAA GAGGGGCTGG GCATTTGATA TGCCCAGCCC CTTGAGTAAG GGGCAAGAAA CCTAAGCCGC CTGGTGGATG GCTCGGCTCG GGGCGCTGAC GAAGGG
23SBHB_123  GC CCGCAGAAGA CGGGTTTGAT GGGGCGGGGA TGTAAGCGGG AAGGGAAACC GACCCGTTTA GTCTGCCGCT CCCAACAGCC CGAGTTTCTT GCCCCTGAAA GAGGGGCTGG GCATTTGATA TGCCCAGCCC CTTGAGTAAG GGGCAAGAAA CCTAAGCCGC CTGGTGGATG GCTCGGCTCG GGGCGCTGAC GAAGGGC
23SBHB_115 CGC CCGCAGAAGA CGGGTTTGAT GGGGCGGGGA TGTAAGCGGG AAGGGAAACC GACCCGTTTA GTCTGCCGCT CCCAACAGCC CGAGTTTCTT GCCCCTGAAA GAGGGGCTGG GCATTTGATA TGCCCAGCCC CTTGAGTAAG GGGCAAGAAA CCTAAGCCGC CTGGTGGATG GCTCGGCTCG GGGCGCTGAC GAAGGGC
23SBHB_128      CGCAGAAGA CGGGTTTGAT GGGGCGGGGA TGTAAGCGGG AAGGGAAACC GACCCGTTTA GTCTGCCGCT CCCAACAGCC CGAGTTTCTT GCCCCTGAAA GAGGGGCTGG GCATTTGATA TGCCCAGCCC CTTGAGTAAG GGGCAAGAAA CCTAAGCCGC CTGGTGGATG GCTCGGCTCG GGGCGCTGAC GAAGGGCG
23SBHB_110     CCGCAGAAGA CGGGTTTGAT GGGGCGGGGA TGTAAGCGGG AAGGGAAACC GACCCGTTTA GTCTGCCGCT CCCAACAGCC CGAGTTTCTT GCCCCTGAAA GAGGGGCTGG GCATTTGATA TGCCCAGCCC CTTGAGTAAG GGGCAAGAAA CCTAAGCCGC CTGGTGGATG GCTCGGCTCG GGGCGCTGAC GAAGGGC
23SBHB_843      CGCAGAAGA CGGGTTTGAT GGGGCGGGGA TGTAAGCGGG AAGGGAAACC GACCCGTTTA GTCTGCCGCT CCCAACAGCC CGAGTTTCTT GCCCCTGAAA GAGGGGCTGG GCATTTGATA TGCCCAGCCC CTTGAGTAAG GGGCAAGAAA CCTAAGCCGC CTGGTGGATG GCTCGGCTCG GGGCGCTGAC GAAGGGCGTG G
23SBHB_378     CCGCAGAAGA CGGGTTTGAT GGGGCGGGGA TGTAAGCGGG AAGGGAAACC GACCCGTTTA GTCTGCCGCT CCCAACAGCC CGAGTTTCTT GCCCCTGAAA GAGGGGCTGG GCATTTGATA TGCCCAGCCC CTTGAGTAAG GGGCAAGAAA CCTAAGCCGC CTGGTGGATG GCTCGGCTCG GGGCGCTGAC GAAGGGCGTG G
23SBHB_347   C CCGCAGAAGA CGGGTTTGAT GGGGCGGGGA TGTAAGCGGG AAGGGAAACC GACCCGTTTA GTCTGCCGCT CCCAACAGCC CGAGTTTCTT GCCCCTGAAA GAGGGGCTGG GCATTTGATA TGCCCAGCCC CTTGAGTAAG GGGCAAGAAA CCTAAGCCGC CTGGTGGATG GCTCGGCTCG GGGCGCTGAC GAAGGGCGTG G
23SBHB_755   C CCGCAGAAGA CGGGTTTGAT GGGGCGGGGA TGTAAGCGGG AAGGGAAACC GACCCGTTTA GTCTGCCGCT CCCAACAGCC CGAGTTTCTT GCCCCTGAAA GAGGGGCTGG GCATTTGATA TGCCCAGCCC CTTGAGTAAG GGGCAAGAAA CCTAAGCCGC CTGGTGGATG GCTCGGCTCG GGGCGCTGAC GAAGGGCGTG GC
23SBHB_873     CCGCAGAAGA CGGGTTTGAT GGGGCGGGGA TGTAAGCGGG AAGGGAAACC GACCCGTTTA GTCTGCCGCT CCCAACAGCC CGAGTTTCTT GCCCCTGAAA GAGGGGCTGG GCATTTGATA TGCCCAGCCC CTTGAGTAAG GGGCAAGAAA CCTAAGCCGC CTGGTGGATG GCTCGGCTCG GGGCGCTGAC GAAGGGCGTG GC
23SBHB_186      CGCAGAAGA CGGGTTTGAT GGGGCGGGGA TGTAAGCGGG AAGGGAAACC GACCCGTTTA GTCTGCCGCT CCCAACAGCC CGAGTTTCTT GCCCCTGAAA GAGGGGCTGG GCATTTGATA TGCCCAGCCC CTTGAGTAAG GGGCAAGAAA CCTAAGCCGC CTGGTGGATG GCTCGGCTCG GGGCGCTGAC GAAGGGCGTG GC
23SBHB_185   C CCGCAGAAGA CGGGTTTGAT GGGGCGGGGA TGTAAGCGGG AAGGGAAACC GACCCGTTTA GTCTGCCGCT CCCAACAGCC CGAGTTTCTT GCCCCTGAAA GAGGGGCTGG GCATTTGATA TGCCCAGCCC CTTGAGTAAG GGGCAAGAAA CCTAAGCCGC CTGGTGGATG GCTCGGCTCG GGGCGCTGAC GAAGGGCGTG
23SBHB_123      CGCAGAAGA CGGGTTTGAT GGGGCGGGGA TGTAAGCGGG AAGGGAAACC GACCCGTTTA GTCTGCCGCT CCCAACAGCC CGAGTTTCTT GCCCCTGAAA GAGGGGCTGG GCATTTGATA TGCCCAGCCC CTTGAGTAAG GGGCAAGAAA CCTAAGCCGC CTGGTGGATG GCTCGGCTCG GGGCGCTGAC GAAGGGCGTG GCA
23SBHB_231      CGCAGAAGA CGGGTTTGAT GGGGCGGGGA TGTAAGCGGG AAGGGAAACC GACCCGTTTA GTCTGCCGCT CCCAACAGCC CGAGTTTCTT GCCCCTGAAA GAGGGGCTGG GCATTTGATA TGCCCAGCCC CTTGAGTAAG GGGCAAGAAA CCTAAGCCGC CTGGTGGATG GCTCGGCTCG GGGCGCTGAC GAAGGGCGTG GCG
23SBHB_283      CGCAGAAGA CGGGTTTGAT GGGGCGGGGA TGTAAGCGGG AAGGGAAACC GACCCGTTTA GTCTGCCGCT CCCAACAGCC CGAGTTTCTT GCCCCTGAAA GAGGGGCTGG GCATTTGATA TGCCCAGCCC CTTGAGTAAG GGGCAAGAAA CCTAAGCCGC CTGGTGGATG GCTCGGCTCG GGGCGCTGAC GAAGGGCGTG GCAAGCTGCG
23SBHB_124     CCGCAGAAGA CGGGTTTGAT GGGGCGGGGA TGTAAGCGGG AAGGGAAACC GACCCGTTTA GTCTGCCGCT CCCAACAGCC CGAGTTTCTT GCCCCTGAAA GAGGGGCTGG GCATTTGATA TGCCCAGCCC CTTGAGTAAG GGGCAAGAAA CCTAAGCCGC CTGGTGGATG GCTCGGCTCG GGGCGCTGAC GAAGGGCGTG GCAAGCTGCG
23SBHB_112   C CCGCAGAAGA CGGGTTTGAT GGGGCGGGGA TGTAAGCGGG AAGGGAAACC GACCCGTTTA GTCTGCCGCT CCCAACAGCC CGAGTTTCTT GCCCCTGAAA GAGGGGCTGG GCATTTGATA TGCCCAGCCC CTTGAGTAAG GGGCAAGAAA CCTAAGCCGC CTGGTGGATG GCTCGGCTCG GGGCGCTGAC GAAGGGCGTG GCAAGCTGCG
23SBHB_199   C CCGCAGAAGA CGGGTTTGAT GGGGCGGGGA TGTAAGCGGG AAGGGAAACC GACCCGTTTA GTCTGCCGCT CCCAACAGCC CGAGTTTCTT GCCCCTGAAA GAGGGGCTGG GCATTTGATA TGCCCAGCCC CTTGAGTAAG GGGCAAGAAA CCTAAGCCGC CTGGTGGATG GCTCGGCTCG GGGCGCTGAC GAAGGGCGTG GCAAGCTGC
23SBHB_344   C CCGCAGAAGA CGGGTTTGAT GGGGCGGGGA TGTAAGCGGG AAGGGAAACC GACCCGTTTA GTCTGCCGCT CCCAACAGCC CGAGTTTCTT GCCCCTGAAA GAGGGGCTGG GCATTTGATA TGCCCAGCCC CTTGAGTAAG GGGCAAGAAA CCTAAGCCGC CTGGTGGATG GCTCGGCTCG GGGCGCTGAC GAAGGGCGTG GCAAGCTG
23SBHB_265     CCGCAGAAGA CGGGTTTGAT GGGGCGGGGA TGTAAGCGGG AAGGGAAACC GACCCGTTTA GTCTGCCGCT CCCAACAGCC CGAGTTTCTT GCCCCTGAAA GAGGGGCTGG GCATTTGATA TGCCCAGCCC CTTGAGTAAG GGGCAAGAAA CCTAAGCCGC CTGGTGGATG GCTCGGCTCG GGGCGCTGAC GAAGGGCGTG GCAAGCTGC
23SBHB_152      CGCAGAAGA CGGGTTTGAT GGGGCGGGGA TGTAAGCGGG AAGGGAAACC GACCCGTTTA GTCTGCCGCT CCCAACAGCC CGAGTTTCTT GCCCCTGAAA GAGGGGCTGG GCATTTGATA TGCCCAGCCC CTTGAGTAAG GGGCAAGAAA CCTAAGCCGC CTGGTGGATG GCTCGGCTCG GGGCGCTGAC GAAGGGCGTG GCAAGCTGCG A
23SBHB_117      CGCAGAAGA CGGGTTTGAT GGGGCGGGGA TGTAAGCGGG AAGGGAAACC GACCCGTTTA GTCTGCCGCT CCCAACAGCC CGAGTTTCTT GCCCCTGAAA GAGGGGCTGG GCATTTGATA TGCCCAGCCC CTTGAGTAAG GGGCAAGAAA CCTAAGCCGC CTGGTGGATG GCTCGGCTCG GGGCGCTGAC GAAGGGCGTG GCAAGCTGCG G
23SBHB_228      CGCAGAAGA CGGGTTTGAT GGGGCGGGGA TGTAAGCGGG AAGGGAAACC GACCCGTTTA GTCTGCCGCT CCCAACAGCC CGAGTTTCTT GCCCCTGAAA GAGGGGCTGG GCATTTGATA TGCCCAGCCC CTTGAGTAAG GGGCAAGAAA CCTAAGCCGC CTGGTGGATG GCTCGGCTCG GGGCGCTGAC GAAGGGCGTG GCAAGCTGCG ATAAGCCCCG GTGAG
23SBHB_923     CCGCAGAAGA CGGGTTTGAT GGGGCGGGGA TGTAAGCGGG AAGGGAAACC GACCCGTTTA GTCTGCCGCT CCCAACAGCC CGAGTTTCTT GCCCCTGAAA GAGGGGCTGG GCATTTGATA TGCCCAGCCC CTTGAGTAAG GGGCAAGAAA CCTAAGCCGC CTGGTGGATG GCTCGGCTCG GGGCGCTGAC GAAGGGCGTG GCAAGCTGCG ATAAGCCCCG GTGAG
23SBHB_544      CGCAGAAGA CGGGTTTGAT GGGGCGGGGA TGTAAGCGGG AAGGGAAACC GACCCGTTTA GTCTGCCGCT CCCAACAGCC CGAGTTTCTT GCCCCTGAAA GAGGGGCTGG GCATTTGATA TGCCCAGCCC CTTGAGTAAG GGGCAAGAAA CCTAAGCCGC CTGGTGGATG GCTCGGCTCG GGGCGCTGAC GAAGGGCGTG GCAAGCTGCG ATAAGCCCCG GTGAGG
23SBHB_150      CGCAGAAGA CGGGTTTGAT GGGGCGGGGA TGTAAGCGGG AAGGGAAACC GACCCGTTTA GTCTGCCGCT CCCAACAGCC CGAGTTTCTT GCCCCTGAAA GAGGGGCTGG GCATTTGATA TGCCCAGCCC CTTGAGTAAG GGGCAAGAAA CCTAAGCCGC CTGGTGGATG GCTCGGCTCG GGGCGCTGAC GAAGGGCGTG GCAAGCTGCG ATAAGCCCCG GTGAGGCGCA GGCAGCCGTC GAGCCGGGGA TTCCCGAATG GGAC
23SBHB_710     CCGCAGAAGA CGGGTTTGAT GGGGCGGGGA TGTAAGCGGG AAGGGAAACC GACCCGTTTA GTCTGCCGCT CCCAACAGCC CGAGTTTCTT GCCCCTGAAA GAGGGGCTGG GCATTTGATA TGCCCAGCCC CTTGAGTAAG GGGCAAGAAA CCTAAGCCGC CTGGTGGATG GCTCGGCTCG GGGCGCTGAC GAAGGGCGTG GCAAGCTGCG ATAAGCCCCG GTGAGGCGCA GGCAGCCGTC GAGCCGGGGA TTCCCGAATG GGAC
23SBHB_606   C CCGCAGAAGA CGGGTTTGAT GGGGCGGGGA TGTAAGCGGG AAGGGAAACC GACCCGTTTA GTCTGCCGCT CCCAACAGCC CGAGTTTCTT GCCCCTGAAA GAGGGGCTGG GCATTTGATA TGCCCAGCCC CTTGAGTAAG GGGCAAGAAA CCTAAGCCGC CTGGTGGATG GCTCGGCTCG GGGCGCTGAC GAAGGGCGTG GCAAGCTGCG ATAAGCCCCG GTGAGGCGCA GGCAGCCGTC GAGCCGGGGA TTCCCGAATG GGAC
23SBHB_173      CGCAGAAGA CGGGTTTGAT GGGGCGGGGA TGTAAGCGGG AAGGGAAACC GACCCGTTTA GTCTGCCGCT CCCAACAGCC CGAGTTTCTT GCCCCTGAAA GAGGGGCTGG GCATTTGATA TGCCCAGCCC CTTGAGTAAG GGGCAAGAAA CCTAAGCCGC CTGGTGGATG GCTCGGCTCG GGGCGCTGAC GAAGGGCGTG GCAAGCTGCG ATAAGCCCCG GTGAGGCGCA GGCAGCCGTC GAGCCGGGGA TTCCCGAATG GGA
23SBHB_159     CCGCAGAAGA CGGGTTTGAT GGGGCGGGGA TGTAAGCGGG AAGGGAAACC GACCCGTTTA GTCTGCCGCT CCCAACAGCC CGAGTTTCTT GCCCCTGAAA GAGGGGCTGG GCATTTGATA TGCCCAGCCC CTTGAGTAAG GGGCAAGAAA CCTAAGCCGC CTGGTGGATG GCTCGGCTCG GGGCGCTGAC GAAGGGCGTG GCAAGCTGCG ATAAGCCCCG GTGAGGCGCA GGCAGCCGTC GAGCCGGGGA TTCCCGAATG GG
23SBHB_152       GCAGAAGA CGGGTTTGAT GGGGCGGGGA TGTAAGCGGG AAGGGAAACC GACCCGTTTA GTCTGCCGCT CCCAACAGCC CGAGTTTCTT GCCCCTGAAA GAGGGGCTGG GCATTTGATA TGCCCAGCCC CTTGAGTAAG GGGCAAGAAA CCTAAGCCGC CTGGTGGATG GCTCGGCTCG GGGCGCTGAC GAAGGGCGTG GCAAGCTGCG ATAAGCCCCG GTGAGGCGCA GGCAGCCGTC GAGCCGGGGA TTCCCGAATG GGAC
23SBHB_133   C CCGCAGAAGA CGGGTTTGAT GGGGCGGGGA TGTAAGCGGG AAGGGAAACC GACCCGTTTA GTCTGCCGCT CCCAACAGCC CGAGTTTCTT GCCCCTGAAA GAGGGGCTGG GCATTTGATA TGCCCAGCCC CTTGAGTAAG GGGCAAGAAA CCTAAGCCGC CTGGTGGATG GCTCGGCTCG GGGCGCTGAC GAAGGGCGTG GCAAGCTGCG ATAAGCCCCG GTGAGGCGCA GGCAGCCGTC GAGCCGGGGA TTCCCGAATG GG
23SBHB_657        CAGAAGA CGGGTTTGAT GGGGCGGGGA TGTAAGCGGG AAGGGAAACC GACCCGTTTA GTCTGCCGCT CCCAACAGCC CGAGTTTCTT GCCCCTGAAA GAGGGGCTGG GCATTTGATA TGCCCAGCCC CTTGAGTAAG GGGCAAGAAA CCTAAGCCGC CTGGTGGATG GCTCGGCTCG GGGCGCTGAC GAAGGGCGTG GCAAGCTGCG ATAAGCCCCG GTGAGGCGCA GGCAGCCGTC GAGCCGGGGA TTCCCGAATG GGAC
23SBHB_303      CGCAGAAGA CGGGTTTGAT GGGGCGGGGA TGTAAGCGGG AAGGGAAACC GACCCGTTTA GTCTGCCGCT CCCAACAGCC CGAGTTTCTT GCCCCTGAAA GAGGGGCTGG GCATTTGATA TGCCCAGCCC CTTGAGTAAG GGGCAAGAAA CCTAAGCCGC CTGGTGGATG GCTCGGCTCG GGGCGCTGAC GAAGGGCGTG GCAAGCTGCG ATAAGCCCCG GTGAGGCGCA GGCAGCCGTC GAGCCGGGGA TTCCCGAATG GG
23SBHB_225         AGAAGA CGGGTTTGAT GGGGCGGGGA TGTAAGCGGG AAGGGAAACC GACCCGTTTA GTCTGCCGCT CCCAACAGCC CGAGTTTCTT GCCCCTGAAA GAGGGGCTGG GCATTTGATA TGCCCAGCCC CTTGAGTAAG GGGCAAGAAA CCTAAGCCGC CTGGTGGATG GCTCGGCTCG GGGCGCTGAC GAAGGGCGTG GCAAGCTGCG ATAAGCCCCG GTGAGGCGCA GGCAGCCGTC GAGCCGGGGA TTCCCGAATG GGAC
23SBHB_126      CGCAGAAGA CGGGTTTGAT GGGGCGGGGA TGTAAGCGGG AAGGGAAACC GACCCGTTTA GTCTGCCGCT CCCAACAGCC CGAGTTTCTT GCCCCTGAAA GAGGGGCTGG GCATTTGATA TGCCCAGCCC CTTGAGTAAG GGGCAAGAAA CCTAAGCCGC CTGGTGGATG GCTCGGCTCG GGGCGCTGAC GAAGGGCGTG GCAAGCTGCG ATAAGCCCCG GTGAGGCGCA GGCAGCCGTC GAGCCGGGGA TTCCCGAATG G
23SBHB_119        CAGAAGA CGGGTTTGAT GGGGCGGGGA TGTAAGCGGG AAGGGAAACC GACCCGTTTA GTCTGCCGCT CCCAACAGCC CGAGTTTCTT GCCCCTGAAA GAGGGGCTGG GCATTTGATA TGCCCAGCCC CTTGAGTAAG GGGCAAGAAA CCTAAGCCGC CTGGTGGATG GCTCGGCTCG GGGCGCTGAC GAAGGGCGTG GCAAGCTGCG ATAAGCCCCG GTGAGGCGCA GGCAGCCGTC GAGCCGGGGA TTCCCGAATG GG
23SBHB_127     CCGCAGAAGA CGGGTTTGAT GGGGCGGGGA TGTAAGCGGG AAGGGAAACC GACCCGTTTA GTCTGCCGCT CCCAACAGCC CGAGTTTCTT GCCCCTGAAA GAGGGGCTGG GCATTTGATA TGCCCAGCCC CTTGAGTAAG GGGCAAGAAA CCTAAGCCGC CTGGTGGATG GCTCGGCTCG GGGCGCTGAC GAAGGGCGTG GCAAGCTGCG ATAAGCCCCG GTGAGGCGCA GGCAGCCGTC GAGCCGGGGA TTCCCGAAT
23SBHB_237      CGCAGAAGA CGGGTTTGAT GGGGCGGGGA TGTAAGCGGG AAGGGAAACC GACCCGTTTA GTCTGCCGCT CCCAACAGCC CGAGTTTCTT GCCCCTGAAA GAGGGGCTGG GCATTTGATA TGCCCAGCCC CTTGAGTAAG GGGCAAGAAA CCTAAGCCGC CTGGTGGATG GCTCGGCTCG GGGCGCTGAC GAAGGGCGTG GCAAGCTGCG ATAAGCCCCG GTGAGGCGCA GGCAGCCGTC GAGCCGGGGA TTCCCGAAT
23SBHB_129            AGA CGGGTTTGAT GGGGCGGGGA TGTAAGCGGG AAGGGAAACC GACCCGTTTA GTCTGCCGCT CCCAACAGCC CGAGTTTCTT GCCCCTGAAA GAGGGGCTGG GCATTTGATA TGCCCAGCCC CTTGAGTAAG GGGCAAGAAA CCTAAGCCGC CTGGTGGATG GCTCGGCTCG GGGCGCTGAC GAAGGGCGTG GCAAGCTGCG ATAAGCCCCG GTGAGGCGCA GGCAGCCGTC GAGCCGGGGA TTCCCGAATG GGAC
23SBHB_319             GA CGGGTTTGAT GGGGCGGGGA TGTAAGCGGG AAGGGAAACC GACCCGTTTA GTCTGCCGCT CCCAACAGCC CGAGTTTCTT GCCCCTGAAA GAGGGGCTGG GCATTTGATA TGCCCAGCCC CTTGAGTAAG GGGCAAGAAA CCTAAGCCGC CTGGTGGATG GCTCGGCTCG GGGCGCTGAC GAAGGGCGTG GCAAGCTGCG ATAAGCCCCG GTGAGGCGCA GGCAGCCGTC GAGCCGGGGA TTCCCGAATG GGAC
23SBHB_479                CGGGTTTGAT GGGGCGGGGA TGTAAGCGGG AAGGGAAACC GACCCGTTTA GTCTGCCGCT CCCAACAGCC CGAGTTTCTT GCCCCTGAAA GAGGGGCTGG GCATTTGATA TGCCCAGCCC CTTGAGTAAG GGGCAAGAAA CCTAAGCCGC CTGGTGGATG GCTCGGCTCG GGGCGCTGAC GAAGGGCGTG GCAAGCTGCG ATAAGCCCCG GTGAGGCGCA GGCAGCCGTC GAGCCGGGGA TTCCCGAATG GGAC
23SBHB_132                   GTTTGAT GGGGCGGGGA TGTAAGCGGG AAGGGAAACC GACCCGTTTA GTCTGCCGCT CCCAACAGCC CGAGTTTCTT GCCCCTGAAA GAGGGGCTGG GCATTTGATA TGCCCAGCCC CTTGAGTAAG GGGCAAGAAA CCTAAGCCGC CTGGTGGATG GCTCGGCTCG GGGCGCTGAC GAAGGGCGTG GCAAGCTGCG ATAAGCCCCG GTGAGGCGCA GGCAGCCGTC GAGCCGGGGA TTCCCGAATG GGAC
23SBHB_368                               CGGGGA TGTAAGCGGG AAGGGAAACC GACCCGTTTA GTCTGCCGCT CCCAACAGCC CGAGTTTCTT GCCCCTGAAA GAGGGGCTGG GCATTTGATA TGCCCAGCCC CTTGAGTAAG GGGCAAGAAA CCTAAGCCGC CTGGTGGATG GCTCGGCTCG GGGCGCTGAC GAAGGGCGTG GCAAGCTGCG ATAAGCCCCG GTGAGGCGCA GGCAGCCGTC GAGCCGGGGA TTCCCGAATG GGAC
23SBHB_115      CGCAGAAGA CGGGTTTGAT GGGGCGGGGA TGTAAGCGGG AAGGGAAACC GACCCGTTTA GTCTGCCGCT CCCAACAGCC CGAGTTTCTT GCCCCTGAAA GAGGGGCTGG GCATTTGATA TGCCCAGCCC CTTGAGTAAG GGGCAAGAAA CCTAAGCCGC CTGGTGGATG GCTCGGCTCG GGGCGCTGAC GAAGGGCGTG GCAAGCTGCG ATAAGCCCCG GTGAGGCGCA GGCAGCCGTC GAG
23SBHB_231      CGCAGAAGA CGGGTTTGAT GGGGCGGGGA TGTAAGCGGG AAGGGAAACC GACCCGTTTA GTCTGCCGCT CCCAACAGCC CGAGTTTCTT GCCCCTGAAA GAGGGGCTGG GCATTTGATA TGCCCAGCCC CTTGAGTAAG GGGCAAGAAA CCTAAGCCGC CTGGTGGATG GCTCGGCTCG GGGCGCTGAC GAAGGGCGTG GCAAGCTGCG ATAAGCCCCG GTGAGGCGCA GGCAGCCG
23SBHB_127     CCGCAGAAGA CGGGTTTGAT GGGGCGGGGA TGTAAGCGGG AAGGGAAACC GACCCGTTTA GTCTGCCGCT CCCAACAGCC CGAGTTTCTT GCCCCTGAAA GAGGGGCTGG GCATTTGATA TGCCCAGCCC CTTGAGTAAG GGGCAAGAAA CCTAAGCCGC CTGGTGGATG GCTCGGCTCG GGGCGCTGAC GAAGGGCGTG GCAAGCTGCG ATAAGCCCCG GTGAGGCGCA GGCAGCCG
23SBHB_137      CGCAGAAGA CGGGTTTGAT GGGGCGGGGA TGTAAGCGGG AAGGGAAACC GACCCGTTTA GTCTGCCGCT CCCAACAGCC CGAGTTTCTT GCCCCTGAAA GAGGGGCTGG GCATTTGATA TGCCCAGCCC CTTGAGTAAG GGGCAAGAAA CCTAAGCCGC CTGGTGGATG GCTCGGCTCG GGGCGCTGAC GAAGGGCGTG GCAAGCTGCG ATAAGCCCCG GTGAGGCGCA GGCAGC
23SBHB_416      CGCAGAAGA CGGGTTTGAT GGGGCGGGGA TGTAAGCGGG AAGGGAAACC GACCCGTTTA GTCTGCCGCT CCCAACAGCC CGAGTTTCTT GCCCCTGAAA GAGGGGCTGG GCATTTGATA TGCCCAGCCC CTTGAGTAAG GGGCAAGAAA CCTAAGCCGC CTGGTGGATG GCTCGGCTCG GGGCGCTGAC GAAGGGCGTG GCAAGCTGCG ATAAGCCCCG GTGAGGCGCA G
23SBHB_122      CGCAGAAGA CGGGTTTGAT GGGGCGGGGA TGTAAGCGGG AAGGGAAACC GACCCGTTTA GTCTGCCGCT CCCAACAGCC CGAGTTTCTT GCCCCTGAAA GAGGGGCTGG GCATTTGATA TGCCCAGCCC CTTGAGTAAG GGGCAAGAAA CCTAAGCCGC CTGGTGGATG GCTCGGCTCG GGGCGCTGAC GAAGGGCGTG GCAAGCTGCG ATAAGCCCCG GTGAGGCGCA GG
23SBHB_200     CCGCAGAAGA CGGGTTTGAT GGGGCGGGGA TGTAAGCGGG AAGGGAAACC GACCCGTTTA GTCTGCCGCT CCCAACAGCC CGAGTTTCTT GCCCCTGAAA GAGGGGCTGG GCATTTGATA TGCCCAGCCC CTTGAGTAAG GGGCAAGAAA CCTAAGCCGC CTGGTGGATG GCTCGGCTCG GGGCGCTGAC GAAGGGCGTG GCAAGCTGCG ATAAGCCCCG GTGAGGCGCA G
23SBHB_191   C CCGCAGAAGA CGGGTTTGAT GGGGCGGGGA TGTAAGCGGG AAGGGAAACC GACCCGTTTA GTCTGCCGCT CCCAACAGCC CGAGTTTCTT GCCCCTGAAA GAGGGGCTGG GCATTTGATA TGCCCAGCCC CTTGAGTAAG GGGCAAGAAA CCTAAGCCGC CTGGTGGATG GCTCGGCTCG GGGCGCTGAC GAAGGGCGTG GCAAGCTGCG ATAAGCCCCG GTGAGGCGCA G
23SBHB_183                CGGGTTTGAT GGGGCGGGGA TGTAAGCGGG AAGGGAAACC GACCCGTTTA GTCTGCCGCT CCCAACAGCC CGAGTTTCTT GCCCCTGAAA GAGGGGCTGG GCATTTGATA TGCCCAGCCC CTTGAGTAAG GGGCAAGAAA CCTAAGCCGC CTGGTGGATG GCTCGGCTCG GGGCGCTGAC GAAGGGCGTG GCAAGCTGCG ATAAGCCCCG GTGAGGCGCA GGCAGCCG
23SBHB_163        CAGAAGA CGGGTTTGAT GGGGCGGGGA TGTAAGCGGG AAGGGAAACC GACCCGTTTA GTCTGCCGCT CCCAACAGCC CGAGTTTCTT GCCCCTGAAA GAGGGGCTGG GCATTTGATA TGCCCAGCCC CTTGAGTAAG GGGCAAGAAA CCTAAGCCGC CTGGTGGATG GCTCGGCTCG GGGCGCTGAC GAAGGGCGTG GCAAGCTGCG ATAAGCCCCG GTGAGGCGCA G
23SBHB_976   C CCGCAGAAGA CGGGTTTGAT GGGGCGGGGA TGTAAGCGGG AAGGGAAACC GACCCGTTTA GTCTGCCGCT CCCAACAGCC CGAGTTTCTT GCCCCTGAAA GAGGGGCTGG GCATTTGATA TGCCCAGCCC CTTGAGTAAG GGGCAAGAAA CCTAAGCCGC CTGGTGGATG GCTCGGCTCG GGGCGCTGAC GAAGGGCGTG GCAAGCTGCG ATAAGCCCCG GTGAG
23SBHB_340      CGCAGAAGA CGGGTTTGAT GGGGCGGGGA TGTAAGCGGG AAGGGAAACC GACCCGTTTA GTCTGCCGCT CCCAACAGCC CGAGTTTCTT GCCCCTGAAA GAGGGGCTGG GCATTTGATA TGCCCAGCCC CTTGAGTAAG GGGCAAGAAA CCTAAGCCGC CTGGTGGATG GCTCGGCTCG GGGCGCTGAC GAAGGGCGTG GCAAGCTGCG ATAAGCCCCG GTGAGGCG
23SBHB_178     CCGCAGAAGA CGGGTTTGAT GGGGCGGGGA TGTAAGCGGG AAGGGAAACC GACCCGTTTA GTCTGCCGCT CCCAACAGCC CGAGTTTCTT GCCCCTGAAA GAGGGGCTGG GCATTTGATA TGCCCAGCCC CTTGAGTAAG GGGCAAGAAA CCTAAGCCGC CTGGTGGATG GCTCGGCTCG GGGCGCTGAC GAAGGGCGTG GCAAGCTGCG ATAAGCCCCG GTGAGGCG
23SBHB_168   C CCGCAGAAGA CGGGTTTGAT GGGGCGGGGA TGTAAGCGGG AAGGGAAACC GACCCGTTTA GTCTGCCGCT CCCAACAGCC CGAGTTTCTT GCCCCTGAAA GAGGGGCTGG GCATTTGATA TGCCCAGCCC CTTGAGTAAG GGGCAAGAAA CCTAAGCCGC CTGGTGGATG GCTCGGCTCG GGGCGCTGAC GAAGGGCGTG GCAAGCTGCG ATAAGCCCCG GTGAGGCG
23SBHB_155   C CCGCAGAAGA CGGGTTTGAT GGGGCGGGGA TGTAAGCGGG AAGGGAAACC GACCCGTTTA GTCTGCCGCT CCCAACAGCC CGAGTTTCTT GCCCCTGAAA GAGGGGCTGG GCATTTGATA TGCCCAGCCC CTTGAGTAAG GGGCAAGAAA CCTAAGCCGC CTGGTGGATG GCTCGGCTCG GGGCGCTGAC GAAGGGCGTG GCAAGCTGCG ATAAGCCCCG GTGAGGC
23SBHB_246   C CCGCAGAAGA CGGGTTTGAT GGGGCGGGGA TGTAAGCGGG AAGGGAAACC GACCCGTTTA GTCTGCCGCT CCCAACAGCC CGAGTTTCTT GCCCCTGAAA GAGGGGCTGG GCATTTGATA TGCCCAGCCC CTTGAGTAAG GGGCAAGAAA CCTAAGCCGC CTGGTGGATG GCTCGGCTCG GGGCGCTGAC GAAGGGCGTG GCAAGCTGCG ATAAGCCCCG GTGAGG
23SBHB_219     CCGCAGAAGA CGGGTTTGAT GGGGCGGGGA TGTAAGCGGG AAGGGAAACC GACCCGTTTA GTCTGCCGCT CCCAACAGCC CGAGTTTCTT GCCCCTGAAA GAGGGGCTGG GCATTTGATA TGCCCAGCCC CTTGAGTAAG GGGCAAGAAA CCTAAGCCGC CTGGTGGATG GCTCGGCTCG GGGCGCTGAC GAAGGGCGTG GCAAGCTGCG ATAAGCCCCG GTGAGGC
23SBHB_418      CGCAGAAGA CGGGTTTGAT GGGGCGGGGA TGTAAGCGGG AAGGGAAACC GACCCGTTTA GTCTGCCGCT CCCAACAGCC CGAGTTTCTT GCCCCTGAAA GAGGGGCTGG GCATTTGATA TGCCCAGCCC CTTGAGTAAG GGGCAAGAAA CCTAAGCCGC CTGGTGGATG GCTCGGCTCG GGGCGCTGAC GAAGGGCGTG GCAAGCTGCG ATAAGCCCCG GTGAGGC
23SBHB_255     CCGCAGAAGA CGGGTTTGAT GGGGCGGGGA TGTAAGCGGG AAGGGAAACC GACCCGTTTA GTCTGCCGCT CCCAACAGCC CGAGTTTCTT GCCCCTGAAA GAGGGGCTGG GCATTTGATA TGCCCAGCCC CTTGAGTAAG GGGCAAGAAA CCTAAGCCGC CTGGTGGATG GCTCGGCTCG GGGCGCTGAC GAAGGGCGTG GCAAGCTGCG ATAAGCCCCG GTGAGG
23SBHB_172        CAGAAGA CGGGTTTGAT GGGGCGGGGA TGTAAGCGGG AAGGGAAACC GACCCGTTTA GTCTGCCGCT CCCAACAGCC CGAGTTTCTT GCCCCTGAAA GAGGGGCTGG GCATTTGATA TGCCCAGCCC CTTGAGTAAG GGGCAAGAAA CCTAAGCCGC CTGGTGGATG GCTCGGCTCG GGGCGCTGAC GAAGGGCGTG GCAAGCTGCG ATAAGCCCCG GTGAGGC
23SBHB_122        CAGAAGA CGGGTTTGAT GGGGCGGGGA TGTAAGCGGG AAGGGAAACC GACCCGTTTA GTCTGCCGCT CCCAACAGCC CGAGTTTCTT GCCCCTGAAA GAGGGGCTGG GCATTTGATA TGCCCAGCCC CTTGAGTAAG GGGCAAGAAA CCTAAGCCGC CTGGTGGATG GCTCGGCTCG GGGCGCTGAC GAAGGGCGTG GCAAGCTGCG ATAAGCCCCG GTGAGGCG
23SBHB_116             GA CGGGTTTGAT GGGGCGGGGA TGTAAGCGGG AAGGGAAACC GACCCGTTTA GTCTGCCGCT CCCAACAGCC CGAGTTTCTT GCCCCTGAAA GAGGGGCTGG GCATTTGATA TGCCCAGCCC CTTGAGTAAG GGGCAAGAAA CCTAAGCCGC CTGGTGGATG GCTCGGCTCG GGGCGCTGAC GAAGGGCGTG GCAAGCTGCG ATAAGCCCCG GTGAGGCGCA G
23SBHB_983        CAGAAGA CGGGTTTGAT GGGGCGGGGA TGTAAGCGGG AAGGGAAACC GACCCGTTTA GTCTGCCGCT CCCAACAGCC CGAGTTTCTT GCCCCTGAAA GAGGGGCTGG GCATTTGATA TGCCCAGCCC CTTGAGTAAG GGGCAAGAAA CCTAAGCCGC CTGGTGGATG GCTCGGCTCG GGGCGCTGAC GAAGGGCGTG GCAAGCTGCG ATAAGCCCCG GTGAG
23SBHB_236       GCAGAAGA CGGGTTTGAT GGGGCGGGGA TGTAAGCGGG AAGGGAAACC GACCCGTTTA GTCTGCCGCT CCCAACAGCC CGAGTTTCTT GCCCCTGAAA GAGGGGCTGG GCATTTGATA TGCCCAGCCC CTTGAGTAAG GGGCAAGAAA CCTAAGCCGC CTGGTGGATG GCTCGGCTCG GGGCGCTGAC GAAGGGCGTG GCAAGCTGCG ATAAGCCCCG GTGAG
23SBHB_232        CAGAAGA CGGGTTTGAT GGGGCGGGGA TGTAAGCGGG AAGGGAAACC GACCCGTTTA GTCTGCCGCT CCCAACAGCC CGAGTTTCTT GCCCCTGAAA GAGGGGCTGG GCATTTGATA TGCCCAGCCC CTTGAGTAAG GGGCAAGAAA CCTAAGCCGC CTGGTGGATG GCTCGGCTCG GGGCGCTGAC GAAGGGCGTG GCAAGCTGCG ATAAGCCCCG GTGAGG
23SBHB_353         AGAAGA CGGGTTTGAT GGGGCGGGGA TGTAAGCGGG AAGGGAAACC GACCCGTTTA GTCTGCCGCT CCCAACAGCC CGAGTTTCTT GCCCCTGAAA GAGGGGCTGG GCATTTGATA TGCCCAGCCC CTTGAGTAAG GGGCAAGAAA CCTAAGCCGC CTGGTGGATG GCTCGGCTCG GGGCGCTGAC GAAGGGCGTG GCAAGCTGCG ATAAGCCCCG GTGAG
23SBHB_216      CGCAGAAGA CGGGTTTGAT GGGGCGGGGA TGTAAGCGGG AAGGGAAACC GACCCGTTTA GTCTGCCGCT CCCAACAGCC CGAGTTTCTT GCCCCTGAAA GAGGGGCTGG GCATTTGATA TGCCCAGCCC CTTGAGTAAG GGGCAAGAAA CCTAAGCCGC CTGGTGGATG GCTCGGCTCG GGGCGCTGAC GAAGGGCGTG GCAAGCTGCG ATAAGCCCCG GTG
23SBHB_213                CGGGTTTGAT GGGGCGGGGA TGTAAGCGGG AAGGGAAACC GACCCGTTTA GTCTGCCGCT CCCAACAGCC CGAGTTTCTT GCCCCTGAAA GAGGGGCTGG GCATTTGATA TGCCCAGCCC CTTGAGTAAG GGGCAAGAAA CCTAAGCCGC CTGGTGGATG GCTCGGCTCG GGGCGCTGAC GAAGGGCGTG GCAAGCTGCG ATAAGCCCCG GTGAGGCGCA G
23SBHB_106      CGCAGAAGA CGGGTTTGAT GGGGCGGGGA TGTAAGCGGG AAGGGAAACC GACCCGTTTA GTCTGCCGCT CCCAACAGCC CGAGTTTCTT GCCCCTGAAA GAGGGGCTGG GCATTTGATA TGCCCAGCCC CTTGAGTAAG GGGCAAGAAA CCTAAGCCGC CTGGTGGATG GCTCGGCTCG GGGCGCTGAC GAAGGGCGTG GCAAGCTGCG ATAAGCCCCG G
23SBHB_515     CCGCAGAAGA CGGGTTTGAT GGGGCGGGGA TGTAAGCGGG AAGGGAAACC GACCCGTTTA GTCTGCCGCT CCCAACAGCC CGAGTTTCTT GCCCCTGAAA GAGGGGCTGG GCATTTGATA TGCCCAGCCC CTTGAGTAAG GGGCAAGAAA CCTAAGCCGC CTGGTGGATG GCTCGGCTCG GGGCGCTGAC GAAGGGCGTG GCAAGCTGCG ATAAGCCCCG G
23SBHB_437   C CCGCAGAAGA CGGGTTTGAT GGGGCGGGGA TGTAAGCGGG AAGGGAAACC GACCCGTTTA GTCTGCCGCT CCCAACAGCC CGAGTTTCTT GCCCCTGAAA GAGGGGCTGG GCATTTGATA TGCCCAGCCC CTTGAGTAAG GGGCAAGAAA CCTAAGCCGC CTGGTGGATG GCTCGGCTCG GGGCGCTGAC GAAGGGCGTG GCAAGCTGCG ATAAGCCCCG G
23SBHB_258   C CCGCAGAAGA CGGGTTTGAT GGGGCGGGGA TGTAAGCGGG AAGGGAAACC GACCCGTTTA GTCTGCCGCT CCCAACAGCC CGAGTTTCTT GCCCCTGAAA GAGGGGCTGG GCATTTGATA TGCCCAGCCC CTTGAGTAAG GGGCAAGAAA CCTAAGCCGC CTGGTGGATG GCTCGGCTCG GGGCGCTGAC GAAGGGCGTG GCAAGCTGCG ATAAGCCCCG
23SBHB_290     CCGCAGAAGA CGGGTTTGAT GGGGCGGGGA TGTAAGCGGG AAGGGAAACC GACCCGTTTA GTCTGCCGCT CCCAACAGCC CGAGTTTCTT GCCCCTGAAA GAGGGGCTGG GCATTTGATA TGCCCAGCCC CTTGAGTAAG GGGCAAGAAA CCTAAGCCGC CTGGTGGATG GCTCGGCTCG GGGCGCTGAC GAAGGGCGTG GCAAGCTGCG ATAAGCCCCG
23SBHB_651      CGCAGAAGA CGGGTTTGAT GGGGCGGGGA TGTAAGCGGG AAGGGAAACC GACCCGTTTA GTCTGCCGCT CCCAACAGCC CGAGTTTCTT GCCCCTGAAA GAGGGGCTGG GCATTTGATA TGCCCAGCCC CTTGAGTAAG GGGCAAGAAA CCTAAGCCGC CTGGTGGATG GCTCGGCTCG GGGCGCTGAC GAAGGGCGTG GCAAGCTGCG ATAAGCCCCG
23SBHB_173      CGCAGAAGA CGGGTTTGAT GGGGCGGGGA TGTAAGCGGG AAGGGAAACC GACCCGTTTA GTCTGCCGCT CCCAACAGCC CGAGTTTCTT GCCCCTGAAA GAGGGGCTGG GCATTTGATA TGCCCAGCCC CTTGAGTAAG GGGCAAGAAA CCTAAGCCGC CTGGTGGATG GCTCGGCTCG GGGCGCTGAC GAAGGGCGTG GCAAGCTGCG ATAAGCCCC
23SBHB_121       GCAGAAGA CGGGTTTGAT GGGGCGGGGA TGTAAGCGGG AAGGGAAACC GACCCGTTTA GTCTGCCGCT CCCAACAGCC CGAGTTTCTT GCCCCTGAAA GAGGGGCTGG GCATTTGATA TGCCCAGCCC CTTGAGTAAG GGGCAAGAAA CCTAAGCCGC CTGGTGGATG GCTCGGCTCG GGGCGCTGAC GAAGGGCGTG GCAAGCTGCG ATAAGCCCCG G
23SBHB_148            AGA CGGGTTTGAT GGGGCGGGGA TGTAAGCGGG AAGGGAAACC GACCCGTTTA GTCTGCCGCT CCCAACAGCC CGAGTTTCTT GCCCCTGAAA GAGGGGCTGG GCATTTGATA TGCCCAGCCC CTTGAGTAAG GGGCAAGAAA CCTAAGCCGC CTGGTGGATG GCTCGGCTCG GGGCGCTGAC GAAGGGCGTG GCAAGCTGCG ATAAGCCCCG GTGAG
23SBHB_492             GA CGGGTTTGAT GGGGCGGGGA TGTAAGCGGG AAGGGAAACC GACCCGTTTA GTCTGCCGCT CCCAACAGCC CGAGTTTCTT GCCCCTGAAA GAGGGGCTGG GCATTTGATA TGCCCAGCCC CTTGAGTAAG GGGCAAGAAA CCTAAGCCGC CTGGTGGATG GCTCGGCTCG GGGCGCTGAC GAAGGGCGTG GCAAGCTGCG ATAAGCCCCG GTGAG
23SBHB_346        CAGAAGA CGGGTTTGAT GGGGCGGGGA TGTAAGCGGG AAGGGAAACC GACCCGTTTA GTCTGCCGCT CCCAACAGCC CGAGTTTCTT GCCCCTGAAA GAGGGGCTGG GCATTTGATA TGCCCAGCCC CTTGAGTAAG GGGCAAGAAA CCTAAGCCGC CTGGTGGATG GCTCGGCTCG GGGCGCTGAC GAAGGGCGTG GCAAGCTGCG ATAAGCCCCG G
23SBHB_255        CAGAAGA CGGGTTTGAT GGGGCGGGGA TGTAAGCGGG AAGGGAAACC GACCCGTTTA GTCTGCCGCT CCCAACAGCC CGAGTTTCTT GCCCCTGAAA GAGGGGCTGG GCATTTGATA TGCCCAGCCC CTTGAGTAAG GGGCAAGAAA CCTAAGCCGC CTGGTGGATG GCTCGGCTCG GGGCGCTGAC GAAGGGCGTG GCAAGCTGCG ATAAGCCCCG
23SBHB_156         AGAAGA CGGGTTTGAT GGGGCGGGGA TGTAAGCGGG AAGGGAAACC GACCCGTTTA GTCTGCCGCT CCCAACAGCC CGAGTTTCTT GCCCCTGAAA GAGGGGCTGG GCATTTGATA TGCCCAGCCC CTTGAGTAAG GGGCAAGAAA CCTAAGCCGC CTGGTGGATG GCTCGGCTCG GGGCGCTGAC GAAGGGCGTG GCAAGCTGCG ATAAGCCCCG G
23SBHB_166                CGGGTTTGAT GGGGCGGGGA TGTAAGCGGG AAGGGAAACC GACCCGTTTA GTCTGCCGCT CCCAACAGCC CGAGTTTCTT GCCCCTGAAA GAGGGGCTGG GCATTTGATA TGCCCAGCCC CTTGAGTAAG GGGCAAGAAA CCTAAGCCGC CTGGTGGATG GCTCGGCTCG GGGCGCTGAC GAAGGGCGTG GCAAGCTGCG ATAAGCCCCG GTGAGGC
23SBHB_137                CGGGTTTGAT GGGGCGGGGA TGTAAGCGGG AAGGGAAACC GACCCGTTTA GTCTGCCGCT CCCAACAGCC CGAGTTTCTT GCCCCTGAAA GAGGGGCTGG GCATTTGATA TGCCCAGCCC CTTGAGTAAG GGGCAAGAAA CCTAAGCCGC CTGGTGGATG GCTCGGCTCG GGGCGCTGAC GAAGGGCGTG GCAAGCTGCG ATAAGCCCCG GTGAGGCG
23SBHB_213                CGGGTTTGAT GGGGCGGGGA TGTAAGCGGG AAGGGAAACC GACCCGTTTA GTCTGCCGCT CCCAACAGCC CGAGTTTCTT GCCCCTGAAA GAGGGGCTGG GCATTTGATA TGCCCAGCCC CTTGAGTAAG GGGCAAGAAA CCTAAGCCGC CTGGTGGATG GCTCGGCTCG GGGCGCTGAC GAAGGGCGTG GCAAGCTGCG ATAAGCCCCG GTGAGG
23SBHB_178              A CGGGTTTGAT GGGGCGGGGA TGTAAGCGGG AAGGGAAACC GACCCGTTTA GTCTGCCGCT CCCAACAGCC CGAGTTTCTT GCCCCTGAAA GAGGGGCTGG GCATTTGATA TGCCCAGCCC CTTGAGTAAG GGGCAAGAAA CCTAAGCCGC CTGGTGGATG GCTCGGCTCG GGGCGCTGAC GAAGGGCGTG GCAAGCTGCG ATAAGCCCCG GTGAG
23SBHB_843                CGGGTTTGAT GGGGCGGGGA TGTAAGCGGG AAGGGAAACC GACCCGTTTA GTCTGCCGCT CCCAACAGCC CGAGTTTCTT GCCCCTGAAA GAGGGGCTGG GCATTTGATA TGCCCAGCCC CTTGAGTAAG GGGCAAGAAA CCTAAGCCGC CTGGTGGATG GCTCGGCTCG GGGCGCTGAC GAAGGGCGTG GCAAGCTGCG ATAAGCCCCG GTGAG
23SBHB_160      CGCAGAAGA CGGGTTTGAT GGGGCGGGGA TGTAAGCGGG AAGGGAAACC GACCCGTTTA GTCTGCCGCT CCCAACAGCC CGAGTTTCTT GCCCCTGAAA GAGGGGCTGG GCATTTGATA TGCCCAGCCC CTTGAGTAAG GGGCAAGAAA CCTAAGCCGC CTGGTGGATG GCTCGGCTCG GGGCGCTGAC GAAGGGCGTG GCAAGCTGCG ATAAGCC
23SBHB_199             GA CGGGTTTGAT GGGGCGGGGA TGTAAGCGGG AAGGGAAACC GACCCGTTTA GTCTGCCGCT CCCAACAGCC CGAGTTTCTT GCCCCTGAAA GAGGGGCTGG GCATTTGATA TGCCCAGCCC CTTGAGTAAG GGGCAAGAAA CCTAAGCCGC CTGGTGGATG GCTCGGCTCG GGGCGCTGAC GAAGGGCGTG GCAAGCTGCG ATAAGCCCCG G
23SBHB_134             GA CGGGTTTGAT GGGGCGGGGA TGTAAGCGGG AAGGGAAACC GACCCGTTTA GTCTGCCGCT CCCAACAGCC CGAGTTTCTT GCCCCTGAAA GAGGGGCTGG GCATTTGATA TGCCCAGCCC CTTGAGTAAG GGGCAAGAAA CCTAAGCCGC CTGGTGGATG GCTCGGCTCG GGGCGCTGAC GAAGGGCGTG GCAAGCTGCG ATAAGCCCCG
23SBHB_305                   GTTTGAT GGGGCGGGGA TGTAAGCGGG AAGGGAAACC GACCCGTTTA GTCTGCCGCT CCCAACAGCC CGAGTTTCTT GCCCCTGAAA GAGGGGCTGG GCATTTGATA TGCCCAGCCC CTTGAGTAAG GGGCAAGAAA CCTAAGCCGC CTGGTGGATG GCTCGGCTCG GGGCGCTGAC GAAGGGCGTG GCAAGCTGCG ATAAGCCCCG GTGAG
23SBHB_130      CGCAGAAGA CGGGTTTGAT GGGGCGGGGA TGTAAGCGGG AAGGGAAACC GACCCGTTTA GTCTGCCGCT CCCAACAGCC CGAGTTTCTT GCCCCTGAAA GAGGGGCTGG GCATTTGATA TGCCCAGCCC CTTGAGTAAG GGGCAAGAAA CCTAAGCCGC CTGGTGGATG GCTCGGCTCG GGGCGCTGAC GAAGGGCGTG GCAAGCTGCG AT
23SBHB_612     CCGCAGAAGA CGGGTTTGAT GGGGCGGGGA TGTAAGCGGG AAGGGAAACC GACCCGTTTA GTCTGCCGCT CCCAACAGCC CGAGTTTCTT GCCCCTGAAA GAGGGGCTGG GCATTTGATA TGCCCAGCCC CTTGAGTAAG GGGCAAGAAA CCTAAGCCGC CTGGTGGATG GCTCGGCTCG GGGCGCTGAC GAAGGGCGTG GCAAGCTGCG AT
23SBHB_476   C CCGCAGAAGA CGGGTTTGAT GGGGCGGGGA TGTAAGCGGG AAGGGAAACC GACCCGTTTA GTCTGCCGCT CCCAACAGCC CGAGTTTCTT GCCCCTGAAA GAGGGGCTGG GCATTTGATA TGCCCAGCCC CTTGAGTAAG GGGCAAGAAA CCTAAGCCGC CTGGTGGATG GCTCGGCTCG GGGCGCTGAC GAAGGGCGTG GCAAGCTGCG AT
23SBHB_336                CGGGTTTGAT GGGGCGGGGA TGTAAGCGGG AAGGGAAACC GACCCGTTTA GTCTGCCGCT CCCAACAGCC CGAGTTTCTT GCCCCTGAAA GAGGGGCTGG GCATTTGATA TGCCCAGCCC CTTGAGTAAG GGGCAAGAAA CCTAAGCCGC CTGGTGGATG GCTCGGCTCG GGGCGCTGAC GAAGGGCGTG GCAAGCTGCG ATAAGCCCCG G
23SBHB_233                CGGGTTTGAT GGGGCGGGGA TGTAAGCGGG AAGGGAAACC GACCCGTTTA GTCTGCCGCT CCCAACAGCC CGAGTTTCTT GCCCCTGAAA GAGGGGCTGG GCATTTGATA TGCCCAGCCC CTTGAGTAAG GGGCAAGAAA CCTAAGCCGC CTGGTGGATG GCTCGGCTCG GGGCGCTGAC GAAGGGCGTG GCAAGCTGCG ATAAGCCCCG
23SBHB_512        CAGAAGA CGGGTTTGAT GGGGCGGGGA TGTAAGCGGG AAGGGAAACC GACCCGTTTA GTCTGCCGCT CCCAACAGCC CGAGTTTCTT GCCCCTGAAA GAGGGGCTGG GCATTTGATA TGCCCAGCCC CTTGAGTAAG GGGCAAGAAA CCTAAGCCGC CTGGTGGATG GCTCGGCTCG GGGCGCTGAC GAAGGGCGTG GCAAGCTGCG AT
23SBHB_134       GCAGAAGA CGGGTTTGAT GGGGCGGGGA TGTAAGCGGG AAGGGAAACC GACCCGTTTA GTCTGCCGCT CCCAACAGCC CGAGTTTCTT GCCCCTGAAA GAGGGGCTGG GCATTTGATA TGCCCAGCCC CTTGAGTAAG GGGCAAGAAA CCTAAGCCGC CTGGTGGATG GCTCGGCTCG GGGCGCTGAC GAAGGGCGTG GCAAGCTGCG AT
23SBHB_187         AGAAGA CGGGTTTGAT GGGGCGGGGA TGTAAGCGGG AAGGGAAACC GACCCGTTTA GTCTGCCGCT CCCAACAGCC CGAGTTTCTT GCCCCTGAAA GAGGGGCTGG GCATTTGATA TGCCCAGCCC CTTGAGTAAG GGGCAAGAAA CCTAAGCCGC CTGGTGGATG GCTCGGCTCG GGGCGCTGAC GAAGGGCGTG GCAAGCTGCG AT
23SBHB_105        CAGAAGA CGGGTTTGAT GGGGCGGGGA TGTAAGCGGG AAGGGAAACC GACCCGTTTA GTCTGCCGCT CCCAACAGCC CGAGTTTCTT GCCCCTGAAA GAGGGGCTGG GCATTTGATA TGCCCAGCCC CTTGAGTAAG GGGCAAGAAA CCTAAGCCGC CTGGTGGATG GCTCGGCTCG GGGCGCTGAC GAAGGGCGTG GCAAGCTGCG
23SBHB_313       GCAGAAGA CGGGTTTGAT GGGGCGGGGA TGTAAGCGGG AAGGGAAACC GACCCGTTTA GTCTGCCGCT CCCAACAGCC CGAGTTTCTT GCCCCTGAAA GAGGGGCTGG GCATTTGATA TGCCCAGCCC CTTGAGTAAG GGGCAAGAAA CCTAAGCCGC CTGGTGGATG GCTCGGCTCG GGGCGCTGAC GAAGGGCGTG GCAAGCTGCG
23SBHB_854      CGCAGAAGA CGGGTTTGAT GGGGCGGGGA TGTAAGCGGG AAGGGAAACC GACCCGTTTA GTCTGCCGCT CCCAACAGCC CGAGTTTCTT GCCCCTGAAA GAGGGGCTGG GCATTTGATA TGCCCAGCCC CTTGAGTAAG GGGCAAGAAA CCTAAGCCGC CTGGTGGATG GCTCGGCTCG GGGCGCTGAC GAAGGGCGTG GCAAGCTG
23SBHB_460      CGCAGAAGA CGGGTTTGAT GGGGCGGGGA TGTAAGCGGG AAGGGAAACC GACCCGTTTA GTCTGCCGCT CCCAACAGCC CGAGTTTCTT GCCCCTGAAA GAGGGGCTGG GCATTTGATA TGCCCAGCCC CTTGAGTAAG GGGCAAGAAA CCTAAGCCGC CTGGTGGATG GCTCGGCTCG GGGCGCTGAC GAAGGGCGTG GCAAGCTGC
23SBHB_196      CGCAGAAGA CGGGTTTGAT GGGGCGGGGA TGTAAGCGGG AAGGGAAACC GACCCGTTTA GTCTGCCGCT CCCAACAGCC CGAGTTTCTT GCCCCTGAAA GAGGGGCTGG GCATTTGATA TGCCCAGCCC CTTGAGTAAG GGGCAAGAAA CCTAAGCCGC CTGGTGGATG GCTCGGCTCG GGGCGCTGAC GAAGGGCGTG GCAAGCTGCT
23SBHB_325     CCGCAGAAGA CGGGTTTGAT GGGGCGGGGA TGTAAGCGGG AAGGGAAACC GACCCGTTTA GTCTGCCGCT CCCAACAGCC CGAGTTTCTT GCCCCTGAAA GAGGGGCTGG GCATTTGATA TGCCCAGCCC CTTGAGTAAG GGGCAAGAAA CCTAAGCCGC CTGGTGGATG GCTCGGCTCG GGGCGCTGAC GAAGGGCGTG GCAAGCTG
23SBHB_447         AGAAGA CGGGTTTGAT GGGGCGGGGA TGTAAGCGGG AAGGGAAACC GACCCGTTTA GTCTGCCGCT CCCAACAGCC CGAGTTTCTT GCCCCTGAAA GAGGGGCTGG GCATTTGATA TGCCCAGCCC CTTGAGTAAG GGGCAAGAAA CCTAAGCCGC CTGGTGGATG GCTCGGCTCG GGGCGCTGAC GAAGGGCGTG GCAAGCTGCG
```

```
23SBHB_331       CAGAAGA CGGGTTTGAT GGGGCGGGGA TGTAAGCGGG AAGGGAAACC GACCCGTTTA GTCTGCCGCT CCCAACAGCC CGAGTTTCTT GCCCCTGAAA GAGGGGCTGG GCATTTGATA TGCCCAGCCC CTTGAGTAAG GGGCAAGAAA CCTAAGCCGC CTGGTGGATG GCTCGGCTCG GGGCGCTGAC GAAGGGCGTG GCAAGCTG
23SBHB_159       CAGAAGA CGGGTTTGAT GGGGCGGGGA TGTAAGCGGG AAGGGAAACC GACCCGTTTA GTCTGCCGCT CCCAACAGCC CGAGTTTCTT GCCCCTGAAA GAGGGGCTGG GCATTTGATA TGCCCAGCCC CTTGAGTAAG GGGCAAGAAA CCTAAGCCGC CTGGTGGATG GCTCGGCTCG GGGCGCTGAC GAAGGGCGTG GCAAGCTGC
23SBHB_237     CGCAGAAGA CGGGTTTGAT GGGGCGGGGA TGTAAGCGGG AAGGGAAACC GACCCGTTTA GTCTGCCGCT CCCAACAGCC CGAGTTTCTT GCCCCTGAAA GAGGGGCTGG GCATTTGATA TGCCCAGCCC CTTGAGTAAG GGGCAAGAAA CCTAAGCCGC CTGGTGGATG GCTCGGCTCG GGGCGCTGAC GAAGGGCGTG GCAAGC
23SBHB_171     CGCAGAAGA CGGGTTTGAT GGGGCGGGGA TGTAAGCGGG AAGGGAAACC GACCCGTTTA GTCTGCCGCT CCCAACAGCC CGAGTTTCTT GCCCCTGAAA GAGGGGCTGG GCATTTGATA TGCCCAGCCC CTTGAGTAAG GGGCAAGAAA CCTAAGCCGC CTGGTGGATG GCTCGGCTCG GGGCGCTGAC GAAGGGCGTG GCAAGCT
23SBHB_724     CGCAGAAGA CGGGTTTGAT GGGGCGGGGA TGTAAGCGGG AAGGGAAACC GACCCGTTTA GTCTGCCGCT CCCAACAGCC CGAGTTTCTT GCCCCTGAAA GAGGGGCTGG GCATTTGATA TGCCCAGCCC CTTGAGTAAG GGGCAAGAAA CCTAAGCCGC CTGGTGGATG GCTCGGCTCG GGGCGCTGAC GAAGGGCGTG GCAAG
23SBHB_362    CCGCAGAAGA CGGGTTTGAT GGGGCGGGGA TGTAAGCGGG AAGGGAAACC GACCCGTTTA GTCTGCCGCT CCCAACAGCC CGAGTTTCTT GCCCCTGAAA GAGGGGCTGG GCATTTGATA TGCCCAGCCC CTTGAGTAAG GGGCAAGAAA CCTAAGCCGC CTGGTGGATG GCTCGGCTCG GGGCGCTGAC GAAGGGCGTG GCAAG
23SBHB_323  C CCGCAGAAGA CGGGTTTGAT GGGGCGGGGA TGTAAGCGGG AAGGGAAACC GACCCGTTTA GTCTGCCGCT CCCAACAGCC CGAGTTTCTT GCCCCTGAAA GAGGGGCTGG GCATTTGATA TGCCCAGCCC CTTGAGTAAG GGGCAAGAAA CCTAAGCCGC CTGGTGGATG GCTCGGCTCG GGGCGCTGAC GAAGGGCGTG GCAAG
23SBHB_242            GA CGGGTTTGAT GGGGCGGGGA TGTAAGCGGG AAGGGAAACC GACCCGTTTA GTCTGCCGCT CCCAACAGCC CGAGTTTCTT GCCCCTGAAA GAGGGGCTGG GCATTTGATA TGCCCAGCCC CTTGAGTAAG GGGCAAGAAA CCTAAGCCGC CTGGTGGATG GCTCGGCTCG GGGCGCTGAC GAAGGGCGTG GCAAGCTGCG AT
23SBHB_130        AGAAGA CGGGTTTGAT GGGGCGGGGA TGTAAGCGGG AAGGGAAACC GACCCGTTTA GTCTGCCGCT CCCAACAGCC CGAGTTTCTT GCCCCTGAAA GAGGGGCTGG GCATTTGATA TGCCCAGCCC CTTGAGTAAG GGGCAAGAAA CCTAAGCCGC CTGGTGGATG GCTCGGCTCG GGGCGCTGAC GAAGGGCGTG GCAAGCTG
23SBHB_172           AGA CGGGTTTGAT GGGGCGGGGA TGTAAGCGGG AAGGGAAACC GACCCGTTTA GTCTGCCGCT CCCAACAGCC CGAGTTTCTT GCCCCTGAAA GAGGGGCTGG GCATTTGATA TGCCCAGCCC CTTGAGTAAG GGGCAAGAAA CCTAAGCCGC CTGGTGGATG GCTCGGCTCG GGGCGCTGAC GAAGGGCGTG GCAAGCTGCG
23SBHB_547            GA CGGGTTTGAT GGGGCGGGGA TGTAAGCGGG AAGGGAAACC GACCCGTTTA GTCTGCCGCT CCCAACAGCC CGAGTTTCTT GCCCCTGAAA GAGGGGCTGG GCATTTGATA TGCCCAGCCC CTTGAGTAAG GGGCAAGAAA CCTAAGCCGC CTGGTGGATG GCTCGGCTCG GGGCGCTGAC GAAGGGCGTG GCAAGCTGCG
23SBHB_360               CGGGTTTGAT GGGGCGGGGA TGTAAGCGGG AAGGGAAACC GACCCGTTTA GTCTGCCGCT CCCAACAGCC CGAGTTTCTT GCCCCTGAAA GAGGGGCTGG GCATTTGATA TGCCCAGCCC CTTGAGTAAG GGGCAAGAAA CCTAAGCCGC CTGGTGGATG GCTCGGCTCG GGGCGCTGAC GAAGGGCGTG GCAAGCTGCG AT
23SBHB_303       CAGAAGA CGGGTTTGAT GGGGCGGGGA TGTAAGCGGG AAGGGAAACC GACCCGTTTA GTCTGCCGCT CCCAACAGCC CGAGTTTCTT GCCCCTGAAA GAGGGGCTGG GCATTTGATA TGCCCAGCCC CTTGAGTAAG GGGCAAGAAA CCTAAGCCGC CTGGTGGATG GCTCGGCTCG GGGCGCTGAC GAAGGGCGTG GCAAG
23SBHB_798               CGGGTTTGAT GGGGCGGGGA TGTAAGCGGG AAGGGAAACC GACCCGTTTA GTCTGCCGCT CCCAACAGCC CGAGTTTCTT GCCCCTGAAA GAGGGGCTGG GCATTTGATA TGCCCAGCCC CTTGAGTAAG GGGCAAGAAA CCTAAGCCGC CTGGTGGATG GCTCGGCTCG GGGCGCTGAC GAAGGGCGTG GCAAGCTGCG
23SBHB_257             A CGGGTTTGAT GGGGCGGGGA TGTAAGCGGG AAGGGAAACC GACCCGTTTA GTCTGCCGCT CCCAACAGCC CGAGTTTCTT GCCCCTGAAA GAGGGGCTGG GCATTTGATA TGCCCAGCCC CTTGAGTAAG GGGCAAGAAA CCTAAGCCGC CTGGTGGATG GCTCGGCTCG GGGCGCTGAC GAAGGGCGTG GCAAGCTGCG
23SBHB_171            GA CGGGTTTGAT GGGGCGGGGA TGTAAGCGGG AAGGGAAACC GACCCGTTTA GTCTGCCGCT CCCAACAGCC CGAGTTTCTT GCCCCTGAAA GAGGGGCTGG GCATTTGATA TGCCCAGCCC CTTGAGTAAG GGGCAAGAAA CCTAAGCCGC CTGGTGGATG GCTCGGCTCG GGGCGCTGAC GAAGGGCGTG GCAAGCTG
23SBHB_918      GCAGAAGA CGGGTTTGAT GGGGCGGGGA TGTAAGCGGG AAGGGAAACC GACCCGTTTA GTCTGCCGCT CCCAACAGCC CGAGTTTCTT GCCCCTGAAA GAGGGGCTGG GCATTTGATA TGCCCAGCCC CTTGAGTAAG GGGCAAGAAA CCTAAGCCGC CTGGTGGATG GCTCGGCTCG GGGCGCTGAC GAAGGGCGTG G
23SBHB_219      GCAGAAGA CGGGTTTGAT GGGGCGGGGA TGTAAGCGGG AAGGGAAACC GACCCGTTTA GTCTGCCGCT CCCAACAGCC CGAGTTTCTT GCCCCTGAAA GAGGGGCTGG GCATTTGATA TGCCCAGCCC CTTGAGTAAG GGGCAAGAAA CCTAAGCCGC CTGGTGGATG GCTCGGCTCG GGGCGCTGAC GAAGGGCGTG GC
23SBHB_676       CAGAAGA CGGGTTTGAT GGGGCGGGGA TGTAAGCGGG AAGGGAAACC GACCCGTTTA GTCTGCCGCT CCCAACAGCC CGAGTTTCTT GCCCCTGAAA GAGGGGCTGG GCATTTGATA TGCCCAGCCC CTTGAGTAAG GGGCAAGAAA CCTAAGCCGC CTGGTGGATG GCTCGGCTCG GGGCGCTGAC GAAGGGCGTG GC
23SBHB_149               CGGGTTTGAT GGGGCGGGGA TGTAAGCGGG AAGGGAAACC GACCCGTTTA GTCTGCCGCT CCCAACAGCC CGAGTTTCTT GCCCCTGAAA GAGGGGCTGG GCATTTGATA TGCCCAGCCC CTTGAGTAAG GGGCAAGAAA CCTAAGCCGC CTGGTGGATG GCTCGGCTCG GGGCGCTGAC GAAGGGCGTG GCAAGCTGC
23SBHB_280        AGAAGA CGGGTTTGAT GGGGCGGGGA TGTAAGCGGG AAGGGAAACC GACCCGTTTA GTCTGCCGCT CCCAACAGCC CGAGTTTCTT GCCCCTGAAA GAGGGGCTGG GCATTTGATA TGCCCAGCCC CTTGAGTAAG GGGCAAGAAA CCTAAGCCGC CTGGTGGATG GCTCGGCTCG GGGCGCTGAC GAAGGGCGTG GC
23SBHB_315       CAGAAGA CGGGTTTGAT GGGGCGGGGA TGTAAGCGGG AAGGGAAACC GACCCGTTTA GTCTGCCGCT CCCAACAGCC CGAGTTTCTT GCCCCTGAAA GAGGGGCTGG GCATTTGATA TGCCCAGCCC CTTGAGTAAG GGGCAAGAAA CCTAAGCCGC CTGGTGGATG GCTCGGCTCG GGGCGCTGAC GAAGGGCGTG G
23SBHB_188       CAGAAGA CGGGTTTGAT GGGGCGGGGA TGTAAGCGGG AAGGGAAACC GACCCGTTTA GTCTGCCGCT CCCAACAGCC CGAGTTTCTT GCCCCTGAAA GAGGGGCTGG GCATTTGATA TGCCCAGCCC CTTGAGTAAG GGGCAAGAAA CCTAAGCCGC CTGGTGGATG GCTCGGCTCG GGGCGCTGAC GAAGGGCGTG
23SBHB_134        AGAAGA CGGGTTTGAT GGGGCGGGGA TGTAAGCGGG AAGGGAAACC GACCCGTTTA GTCTGCCGCT CCCAACAGCC CGAGTTTCTT GCCCCTGAAA GAGGGGCTGG GCATTTGATA TGCCCAGCCC CTTGAGTAAG GGGCAAGAAA CCTAAGCCGC CTGGTGGATG GCTCGGCTCG GGGCGCTGAC GAAGGGCGTG G
23SBHB_525      GCAGAAGA CGGGTTTGAT GGGGCGGGGA TGTAAGCGGG AAGGGAAACC GACCCGTTTA GTCTGCCGCT CCCAACAGCC CGAGTTTCTT GCCCCTGAAA GAGGGGCTGG GCATTTGATA TGCCCAGCCC CTTGAGTAAG GGGCAAGAAA CCTAAGCCGC CTGGTGGATG GCTCGGCTCG GGGCGCTGAC GAAGGGCGTG
23SBHB_295               CGGGTTTGAT GGGGCGGGGA TGTAAGCGGG AAGGGAAACC GACCCGTTTA GTCTGCCGCT CCCAACAGCC CGAGTTTCTT GCCCCTGAAA GAGGGGCTGG GCATTTGATA TGCCCAGCCC CTTGAGTAAG GGGCAAGAAA CCTAAGCCGC CTGGTGGATG GCTCGGCTCG GGGCGCTGAC GAAGGGCGTG GCAAGCTG
23SBHB_797        AGAAGA CGGGTTTGAT GGGGCGGGGA TGTAAGCGGG AAGGGAAACC GACCCGTTTA GTCTGCCGCT CCCAACAGCC CGAGTTTCTT GCCCCTGAAA GAGGGGCTGG GCATTTGATA TGCCCAGCCC CTTGAGTAAG GGGCAAGAAA CCTAAGCCGC CTGGTGGATG GCTCGGCTCG GGGCGCTGAC GAAGGGCGTG
23SBHB_313                  GTTTGAT GGGGCGGGGA TGTAAGCGGG AAGGGAAACC GACCCGTTTA GTCTGCCGCT CCCAACAGCC CGAGTTTCTT GCCCCTGAAA GAGGGGCTGG GCATTTGATA TGCCCAGCCC CTTGAGTAAG GGGCAAGAAA CCTAAGCCGC CTGGTGGATG GCTCGGCTCG GGGCGCTGAC GAAGGGCGTG GCAAGCTGCG
23SBHB_137                  GTTTGAT GGGGCGGGGA TGTAAGCGGG AAGGGAAACC GACCCGTTTA GTCTGCCGCT CCCAACAGCC CGAGTTTCTT GCCCCTGAAA GAGGGGCTGG GCATTTGATA TGCCCAGCCC CTTGAGTAAG GGGCAAGAAA CCTAAGCCGC CTGGTGGATG GCTCGGCTCG GGGCGCTGAC GAAGGGCGTG GCAAGCTGCG AT
23SBHB_218         GAAGA CGGGTTTGAT GGGGCGGGGA TGTAAGCGGG AAGGGAAACC GACCCGTTTA GTCTGCCGCT CCCAACAGCC CGAGTTTCTT GCCCCTGAAA GAGGGGCTGG GCATTTGATA TGCCCAGCCC CTTGAGTAAG GGGCAAGAAA CCTAAGCCGC CTGGTGGATG GCTCGGCTCG GGGCGCTGAC GAAGGGCGTG G
23SBHB_154            GA CGGGTTTGAT GGGGCGGGGA TGTAAGCGGG AAGGGAAACC GACCCGTTTA GTCTGCCGCT CCCAACAGCC CGAGTTTCTT GCCCCTGAAA GAGGGGCTGG GCATTTGATA TGCCCAGCCC CTTGAGTAAG GGGCAAGAAA CCTAAGCCGC CTGGTGGATG GCTCGGCTCG GGGCGCTGAC GAAGGGCGTG GCAAG
23SBHB_501       CAGAAGA CGGGTTTGAT GGGGCGGGGA TGTAAGCGGG AAGGGAAACC GACCCGTTTA GTCTGCCGCT CCCAACAGCC CGAGTTTCTT GCCCCTGAAA GAGGGGCTGG GCATTTGATA TGCCCAGCCC CTTGAGTAAG GGGCAAGAAA CCTAAGCCGC CTGGTGGATG GCTCGGCTCG GGGCGCTGAC GAAGGGCG
23SBHB_128      GCAGAAGA CGGGTTTGAT GGGGCGGGGA TGTAAGCGGG AAGGGAAACC GACCCGTTTA GTCTGCCGCT CCCAACAGCC CGAGTTTCTT GCCCCTGAAA GAGGGGCTGG GCATTTGATA TGCCCAGCCC CTTGAGTAAG GGGCAAGAAA CCTAAGCCGC CTGGTGGATG GCTCGGCTCG GGGCGCTGAC GAAGGGCG
23SBHB_279      GCAGAAGA CGGGTTTGAT GGGGCGGGGA TGTAAGCGGG AAGGGAAACC GACCCGTTTA GTCTGCCGCT CCCAACAGCC CGAGTTTCTT GCCCCTGAAA GAGGGGCTGG GCATTTGATA TGCCCAGCCC CTTGAGTAAG GGGCAAGAAA CCTAAGCCGC CTGGTGGATG GCTCGGCTCG GGGCGCTGAC GAAGGGC
23SBHB_121       CAGAAGA CGGGTTTGAT GGGGCGGGGA TGTAAGCGGG AAGGGAAACC GACCCGTTTA GTCTGCCGCT CCCAACAGCC CGAGTTTCTT GCCCCTGAAA GAGGGGCTGG GCATTTGATA TGCCCAGCCC CTTGAGTAAG GGGCAAGAAA CCTAAGCCGC CTGGTGGATG GCTCGGCTCG GGGCGCTGAC GAAGGGCGT
23SBHB_122         GAAGA CGGGTTTGAT GGGGCGGGGA TGTAAGCGGG AAGGGAAACC GACCCGTTTA GTCTGCCGCT CCCAACAGCC CGAGTTTCTT GCCCCTGAAA GAGGGGCTGG GCATTTGATA TGCCCAGCCC CTTGAGTAAG GGGCAAGAAA CCTAAGCCGC CTGGTGGATG GCTCGGCTCG GGGCGCTGAC GAAGGGCGTG
23SBHB_331          AAGA CGGGTTTGAT GGGGCGGGGA TGTAAGCGGG AAGGGAAACC GACCCGTTTA GTCTGCCGCT CCCAACAGCC CGAGTTTCTT GCCCCTGAAA GAGGGGCTGG GCATTTGATA TGCCCAGCCC CTTGAGTAAG GGGCAAGAAA CCTAAGCCGC CTGGTGGATG GCTCGGCTCG GGGCGCTGAC GAAGGGCGTG G
23SBHB_170          AAGA CGGGTTTGAT GGGGCGGGGA TGTAAGCGGG AAGGGAAACC GACCCGTTTA GTCTGCCGCT CCCAACAGCC CGAGTTTCTT GCCCCTGAAA GAGGGGCTGG GCATTTGATA TGCCCAGCCC CTTGAGTAAG GGGCAAGAAA CCTAAGCCGC CTGGTGGATG GCTCGGCTCG GGGCGCTGAC GAAGGGCGTG
23SBHB_205        AGAAGA CGGGTTTGAT GGGGCGGGGA TGTAAGCGGG AAGGGAAACC GACCCGTTTA GTCTGCCGCT CCCAACAGCC CGAGTTTCTT GCCCCTGAAA GAGGGGCTGG GCATTTGATA TGCCCAGCCC CTTGAGTAAG GGGCAAGAAA CCTAAGCCGC CTGGTGGATG GCTCGGCTCG GGGCGCTGAC GAAGGGCG
23SBHB_525           AGA CGGGTTTGAT GGGGCGGGGA TGTAAGCGGG AAGGGAAACC GACCCGTTTA GTCTGCCGCT CCCAACAGCC CGAGTTTCTT GCCCCTGAAA GAGGGGCTGG GCATTTGATA TGCCCAGCCC CTTGAGTAAG GGGCAAGAAA CCTAAGCCGC CTGGTGGATG GCTCGGCTCG GGGCGCTGAC GAAGGGCGTG G
23SBHB_116           AGA CGGGTTTGAT GGGGCGGGGA TGTAAGCGGG AAGGGAAACC GACCCGTTTA GTCTGCCGCT CCCAACAGCC CGAGTTTCTT GCCCCTGAAA GAGGGGCTGG GCATTTGATA TGCCCAGCCC CTTGAGTAAG GGGCAAGAAA CCTAAGCCGC CTGGTGGATG GCTCGGCTCG GGGCGCTGAC GAAGGGCGTG GC
23SBHB_354            GA CGGGTTTGAT GGGGCGGGGA TGTAAGCGGG AAGGGAAACC GACCCGTTTA GTCTGCCGCT CCCAACAGCC CGAGTTTCTT GCCCCTGAAA GAGGGGCTGG GCATTTGATA TGCCCAGCCC CTTGAGTAAG GGGCAAGAAA CCTAAGCCGC CTGGTGGATG GCTCGGCTCG GGGCGCTGAC GAAGGGCGTG GC
23SBHB_256     CGCAGAAGA CGGGTTTGAT GGGGCGGGGA TGTAAGCGGG AAGGGAAACC GACCCGTTTA GTCTGCCGCT CCCAACAGCC CGAGTTTCTT GCCCCTGAAA GAGGGGCTGG GCATTTGATA TGCCCAGCCC CTTGAGTAAG GGGCAAGAAA CCTAAGCCGC CTGGTGGATG GCTCGGCTCG GGGCGCTGAC GAAGGAC
23SBHB_121     CGCAGAAGA CGGGTTTGAT GGGGCGGGGA TGTAAGCGGG AAGGGAAACC GACCCGTTTA GTCTGCCGCT CCCAACAGCC CGAGTTTCTT GCCCCTGAAA GAGGGGCTGG GCATTTGATA TGCCCAGCCC CTTGAGTAAG GGGCAAGAAA CCTAAGCCGC CTGGTGGATG GCTCGGCTCG GGGCGCTGAC GAAGGC
23SBHB_311     CGCAGAAGA CGGGTTTGAT GGGGCGGGGA TGTAAGCGGG AAGGGAAACC GACCCGTTTA GTCTGCCGCT CCCAACAGCC CGAGTTTCTT GCCCCTGAAA GAGGGGCTGG GCATTTGATA TGCCCAGCCC CTTGAGTAAG GGGCAAGAAA CCTAAGCCGC CTGGTGGATG GCTCGGCTCG GGGCGCTGAC GAAG
23SBHB_131    CCGCAGAAGA CGGGTTTGAT GGGGCGGGGA TGTAAGCGGG AAGGGAAACC GACCCGTTTA GTCTGCCGCT CCCAACAGCC CGAGTTTCTT GCCCCTGAAA GAGGGGCTGG GCATTTGATA TGCCCAGCCC CTTGAGTAAG GGGCAAGAAA CCTAAGCCGC CTGGTGGATG GCTCGGCTCG GGGCGCTGAC GAAG
23SBHB_617    CCGCAGAAGA CGGGTTTGAT GGGGCGGGGA TGTAAGCGGG AAGGGAAACC GACCCGTTTA GTCTGCCGCT CCCAACAGCC CGAGTTTCTT GCCCCTGAAA GAGGGGCTGG GCATTTGATA TGCCCAGCCC CTTGAGTAAG GGGCAAGAAA CCTAAGCCGC CTGGTGGATG GCTCGGCTCG GGGCGCTGAC GAA
23SBHB_468     CGCAGAAGA CGGGTTTGAT GGGGCGGGGA TGTAAGCGGG AAGGGAAACC GACCCGTTTA GTCTGCCGCT CCCAACAGCC CGAGTTTCTT GCCCCTGAAA GAGGGGCTGG GCATTTGATA TGCCCAGCCC CTTGAGTAAG GGGCAAGAAA CCTAAGCCGC CTGGTGGATG GCTCGGCTCG GGGCGCTGAC GAAGAG
23SBHB_427  C CCGCAGAAGA CGGGTTTGAT GGGGCGGGGA TGTAAGCGGG AAGGGAAACC GACCCGTTTA GTCTGCCGCT CCCAACAGCC CGAGTTTCTT GCCCCTGAAA GAGGGGCTGG GCATTTGATA TGCCCAGCCC CTTGAGTAAG GGGCAAGAAA CCTAAGCCGC CTGGTGGATG GCTCGGCTCG GGGCGCTGAC GAA
23SBHB_178  C CCGCAGAAGA CGGGTTTGAT GGGGCGGGGA TGTAAGCGGG AAGGGAAACC GACCCGTTTA GTCTGCCGCT CCCAACAGCC CGAGTTTCTT GCCCCTGAAA GAGGGGCTGG GCATTTGATA TGCCCAGCCC CTTGAGTAAG GGGCAAGAAA CCTAAGCCGC CTGGTGGATG GCTCGGCTCG GGGCGCTGAC AAAG
23SBHB_176    CCGCAGAAGA CGGGTTTGAT GGGGCGGGGA TGTAAGCGGG AAGGGAAACC GACCCGTTTA GTCTGCCGCT CCCAACAGCC CGAGTTTCTT GCCCCTGAAA GAGGGGCTGG GCATTTGATA TGCCCAGCCC CTTGAGTAAG GGGCAAGAAA CCTAAGCCGC CTGGTGGATG GCTCGGCTCG GGGCGCTGAC AAAG
23SBHB_167       CAGAAGA CGGGTTTGAT GGGGCGGGGA TGTAAGCGGG AAGGGAAACC GACCCGTTTA GTCTGCCGCT CCCAACAGCC CGAGTTTCTT GCCCCTGAAA GAGGGGCTGG GCATTTGATA TGCCCAGCCC CTTGAGTAAG GGGCAAGAAA CCTAAGCCGC CTGGTGGATG GCTCGGCTCG GGGCGCTGAC GAAGGG
23SBHB_887       CAGAAGA CGGGTTTGAT GGGGCGGGGA TGTAAGCGGG AAGGGAAACC GACCCGTTTA GTCTGCCGCT CCCAACAGCC CGAGTTTCTT GCCCCTGAAA GAGGGGCTGG GCATTTGATA TGCCCAGCCC CTTGAGTAAG GGGCAAGAAA CCTAAGCCGC CTGGTGGATG GCTCGGCTCG GGGCGCTGAC GAAGGGC
23SBHB_165       CAGAAGA CGGGTTTGAT GGGGCGGGGA TGTAAGCGGG AAGGGAAACC GACCCGTTTA GTCTGCCGCT CCCAACAGCC CGAGTTTCTT GCCCCTGAAA GAGGGGCTGG GCATTTGATA TGCCCAGCCC CTTGAGTAAG GGGCAAGAAA CCTAAGCCGC CTGGTGGATG GCTCGGCTCG GGGCGCTGAC GAAGGGCC
23SBHB_436      GCAGAAGA CGGGTTTGAT GGGGCGGGGA TGTAAGCGGG AAGGGAAACC GACCCGTTTA GTCTGCCGCT CCCAACAGCC CGAGTTTCTT GCCCCTGAAA GAGGGGCTGG GCATTTGATA TGCCCAGCCC CTTGAGTAAG GGGCAAGAAA CCTAAGCCGC CTGGTGGATG GCTCGGCTCG GGGCGCTGAC GAAGGG
23SBHB_126      GCAGAAGA CGGGTTTGAT GGGGCGGGGA TGTAAGCGGG AAGGGAAACC GACCCGTTTA GTCTGCCGCT CCCAACAGCC CGAGTTTCTT GCCCCTGAAA GAGGGGCTGG GCATTTGATA TGCCCAGCCC CTTGAGTAAG GGGCAAGAAA CCTAAGCCGC CTGGTGGATG GCTCGGCTCG GGGCGCTGAC GAAGGGG
23SBHB_503       CAGAAGA CGGGTTTGAT GGGGCGGGGA TGTAAGCGGG AAGGGAAACC GACCCGTTTA GTCTGCCGCT CCCAACAGCC CGAGTTTCTT GCCCCTGAAA GAGGGGCTGG GCATTTGATA TGCCCAGCCC CTTGAGTAAG GGGCAAGAAA CCTAAGCCGC CTGGTGGATG GCTCGGCTCG GGGCGCTGAC GAAGGGG
23SBHB_348        AGAAGA CGGGTTTGAT GGGGCGGGGA TGTAAGCGGG AAGGGAAACC GACCCGTTTA GTCTGCCGCT CCCAACAGCC CGAGTTTCTT GCCCCTGAAA GAGGGGCTGG GCATTTGATA TGCCCAGCCC CTTGAGTAAG GGGCAAGAAA CCTAAGCCGC CTGGTGGATG GCTCGGCTCG GGGCGCTGAC GAAGGGC
23SBHB_293      GCAGAAGA CGGGTTTGAT GGGGCGGGGA TGTAAGCGGG AAGGGAAACC GACCCGTTTA GTCTGCCGCT CCCAACAGCC CGAGTTTCTT GCCCCTGAAA GAGGGGCTGG GCATTTGATA TGCCCAGCCC CTTGAGTAAG GGGCAAGAAA CCTAAGCCGC CTGGTGGATG GCTCGGCTCG GGGCGCTGAC GAAGG
23SBHB_249     CGCAGAAGA CGGGTTTGAT GGGGCGGGGA TGTAAGCGGG AAGGGAAACC GACCCGTTTA GTCTGCCGCT CCCAACAGCC CGAGTTTCTT GCCCCTGAAA GAGGGGCTGG GCATTTGATA TGCCCAGCCC CTTGAGTAAG GGGCAAGAAA CCTAAGCCGC CTGGTGGATG GCTCGGCTCG GGGCGCTGAC AAAGGG
23SBHB_229     CGCAGAAGA CGGGTTTGAT GGGGCGGGGA TGTAAGCGGG AAGGGAAACC GACCCGTTTA GTCTGCCGCT CCCAACAGCC CGAGTTTCTT GCCCCTGAAA GAGGGGCTGG GCATTTGATA TGCCCAGCCC CTTGAGTAAG GGGCAAGAAA CCTAAGCCGC CTGGTGGATG GCTCGGCTCG GGGCGCTGAC AAAGG
23SBHB_149     NGCAGAAGA CGGGTTTGAT GGGGCGGGGA TGTAAGCGGG AAGGGAAACC GACCCGTTTA GTCTGCCGCT CCCAACAGCC CGAGTTTCTT GCCCCTGAAA GAGGGGCTGG GCATTTGATA TGCCCAGCCC CTTGAGTAAG GGGCAAGAAA CCTAAGCCGC CTGGTGGATG GCTCGGCTCG GGGCGCTGAC GAAGGN
23SBHB_151     CGCAGAAGA CGGGTTTGAT GGGGCGGGGA TGTAAGCGGG CAGGGAAACC GACCCGTTTA GTCTGCCGCT CCCAACAGCC CGAGTTTCTT GCCCCTGAAA GAGGGGCTGG GCATTTGATA TGCCCAGCCC CTTGAGTAAG GGGCAAGAAA CCTAAGCCGC CTGGTGGATG GCTCGGCTCG GGGCGCTGAC GAAGG
23SBHB_124     CGCAGAAGA CGGGTTTGAT GGGGCGGGGA TGTAAGCGGG CAGGGAAACC GACCCGTTTA GTCTGCCGCT CCCAACAGCC CGAGTTTCTT GCCCCTGAAA GAGGGGCTGG GCATTTGATA TGCCCAGCCC CTTGAGTAAG GGGCAAGAAA CCTAAGCCGC CTGGTGGATG GCTCGGCTCG GGGCGCTGAC GAAGGG
23SBHB_202        AGAAGA CGGGTTTGAT GGGGCGGGGA TGTAAGCGGG AAGGGAAACC GACCCGTTTA GTCTGCCGCT CCCAACAGCC CGAGTTTCTT GCCCCTGAAA GAGGGGCTGG GCATTTGATA TGCCCAGCCC CTTGAGTAAG GGGCAAGAAA CCTAAGCCGC CTGGTGGATG GCTCGGCTCG GGGCGCTGAC GAAGGGG
23SBHB_149            GA CGGGTTTGAT GGGGCGGGGA TGTAAGCGGG AAGGGAAACC GACCCGTTTA GTCTGCCGCT CCCAACAGCC CGAGTTTCTT GCCCCTGAAA GAGGGGCTGG GCATTTGATA TGCCCAGCCC CTTGAGTAAG GGGCAAGAAA CCTAAGCCGC CTGGTGGATG GCTCGGCTCG GGGCGCTGAC GAAGGGCGTG G
23SBHB_333         GAAGA CGGGTTTGAT GGGGCGGGGA TGTAAGCGGG AAGGGAAACC GACCCGTTTA GTCTGCCGCT CCCAACAGCC CGAGTTTCTT GCCCCTGAAA GAGGGGCTGG GCATTTGATA TGCCCAGCCC CTTGAGTAAG GGGCAAGAAA CCTAAGCCGC CTGGTGGATG GCTCGGCTCG GGGCGCTGAC GAAGGGCG
23SBHB_315           AGA CGGGTTTGAT GGGGCGGGGA TGTAAGCGGG AAGGGAAACC GACCCGTTTA GTCTGCCGCT CCCAACAGCC CGAGTTTCTT GCCCCTGAAA GAGGGGCTGG GCATTTGATA TGCCCAGCCC CTTGAGTAAG GGGCAAGAAA CCTAAGCCGC CTGGTGGATG GCTCGGCTCG GGGCGCTGAC GAAGGGCGTG
23SBHB_247               CGGGTTTGAT GGGGCGGGGA TGTAAGCGGG AAGGGAAACC GACCCGTTTA GTCTGCCGCT CCCAACAGCC CGAGTTTCTT GCCCCTGAAA GAGGGGCTGG GCATTTGATA TGCCCAGCCC CTTGAGTAAG GGGCAAGAAA CCTAAGCCGC CTGGTGGATG GCTCGGCTCG GGGCGCTGAC GAAGGGCGTG GCAAG
23SBHB_688        AGAAGA CGGGTTTGAT GGGGCGGGGA TGTAAGCGGG AAGGGAAACC GACCCGTTTA GTCTGCCGCT CCCAACAGCC CGAGTTTCTT GCCCCTGAAA GAGGGGCTGG GCATTTGATA TGCCCAGCCC CTTGAGTAAG GGGCAAGAAA CCTAAGCCGC CTGGTGGATG GCTCGGCTCG GGGCGCTGAC GAAGGG
23SBHB_119     CGCAGAAGA CGGGTTTGAT GGGGCGGGGA TGTAAGCGGG AAGGGAAACC GACCCGTTTA GTCTGCCGCT CCCAACAGCC CGAGTTTCTT GCCCCTGAAA GAGGGGCTGG GCATTTGATA TGCCCAGCCC CTTGAGTAAG GGGCAAGAAA CCTAAGCCGC CTGGTGGATG GCTCGGCTCG GGGCGCTGAC GAA
23SBHB_222     CGCAGAAGA CGGGTTTGAT GGGGCGGGGA TGTAAGCGGG AAGGGAAACC GACCCGTTTA GTCTGCCGCT CCCAACAGCC CGAGTTTCTT GCCCCTGAAA GAGGGGCTGG GCATTTGATA TGCCCAGCCC CTTGAGTAAG GGGCAAGAAA CCTAAGCCGC CTGGTGGATG GCTCGGCTCG GGGCGCTGAC GAAAG
23SBHB_874            GA CGGGTTTGAT GGGGCGGGGA TGTAAGCGGG AAGGGAAACC GACCCGTTTA GTCTGCCGCT CCCAACAGCC CGAGTTTCTT GCCCCTGAAA GAGGGGCTGG GCATTTGATA TGCCCAGCCC CTTGAGTAAG GGGCAAGAAA CCTAAGCCGC CTGGTGGATG GCTCGGCTCG GGGCGCTGAC GAAGGGCGTG
23SBHB_638             A CGGGTTTGAT GGGGCGGGGA TGTAAGCGGG AAGGGAAACC GACCCGTTTA GTCTGCCGCT CCCAACAGCC CGAGTTTCTT GCCCCTGAAA GAGGGGCTGG GCATTTGATA TGCCCAGCCC CTTGAGTAAG GGGCAAGAAA CCTAAGCCGC CTGGTGGATG GCTCGGCTCG GGGCGCTGAC GAAGGGCGTG G
23SBHB_143             A CGGGTTTGAT GGGGCGGGGA TGTAAGCGGG AAGGGAAACC GACCCGTTTA GTCTGCCGCT CCCAACAGCC CGAGTTTCTT GCCCCTGAAA GAGGGGCTGG GCATTTGATA TGCCCAGCCC CTTGAGTAAG GGGCAAGAAA CCTAAGCCGC CTGGTGGATG GCTCGGCTCG GGGCGCTGAC GAAGGGCGTG GC
23SBHB_576               CGGGTTTGAT GGGGCGGGGA TGTAAGCGGG AAGGGAAACC GACCCGTTTA GTCTGCCGCT CCCAACAGCC CGAGTTTCTT GCCCCTGAAA GAGGGGCTGG GCATTTGATA TGCCCAGCCC CTTGAGTAAG GGGCAAGAAA CCTAAGCCGC CTGGTGGATG GCTCGGCTCG GGGCGCTGAC GAAGGGCGTG GC
23SBHB_569         GAAGA CGGGTTTGAT GGGGCGGGGA TGTAAGCGGG AAGGGAAACC GACCCGTTTA GTCTGCCGCT CCCAACAGCC CGAGTTTCTT GCCCCTGAAA GAGGGGCTGG GCATTTGATA TGCCCAGCCC CTTGAGTAAG GGGCAAGAAA CCTAAGCCGC CTGGTGGATG GCTCGGCTCG GGGCGCTGAC GAAGGGC
23SBHB_536          AAGA CGGGTTTGAT GGGGCGGGGA TGTAAGCGGG AAGGGAAACC GACCCGTTTA GTCTGCCGCT CCCAACAGCC CGAGTTTCTT GCCCCTGAAA GAGGGGCTGG GCATTTGATA TGCCCAGCCC CTTGAGTAAG GGGCAAGAAA CCTAAGCCGC CTGGTGGATG GCTCGGCTCG GGGCGCTGAC GAAGGGCG
23SBHB_120       CAGAAGA CGGGTTTGAT GGGGCGGGGA TGTAAGCGGG AAGGGAAACC GACCCGTTTA GTCTGCCGCT CCCAACAGCC CGAGTTTCTT GCCCCTGAAA GAGGGGCTGG GCATTTGATA TGCCCAGCCC CTTGAGTAAG GGGCAAGAAA CCTAAGCCGC CTGGTGGATG GCTCGGCTCG GGGCGCTGAC GAAG
23SBHB_106       CAGAAGA CGGGTTTGAT GGGGCGGGGA TGTAAGCGGG AAGGGAAACC GACCCGTTTA GTCTGCCGCT CCCAACAGCC CGAGTTTCTT GCCCCTGAAA GAGGGGCTGG GCATTTGATA TGCCCAGCCC CTTGAGTAAG GGGCAAGAAA CCTAAGCCGC CTGGTGGATG GCTCGGCTCG GGGCGCTGAC GAAGG
23SBHB_175       CAGAAGA CGGGTTTGAT GGGGCGGGGA TGTAAGCGGG AAGGGAAACC GACCCGTTTA GTCTGCCGCT CCCAACAGCC CGAGTTTCTT GCCCCTGAAA GAGGGGCTGG GCATTTGATA TGCCCAGCCC CTTGAGTAAG GGGCAAGAAA CCTAAGCCGC CTGGTGGATG GCTCGGCTCG GGGCGCTGAC GAAGAG
23SBHB_445        AGAAGA CGGGTTTGAT GGGGCGGGGA TGTAAGCGGG AAGGGAAACC GACCCGTTTA GTCTGCCGCT CCCAACAGCC CGAGTTTCTT GCCCCTGAAA GAGGGGCTGG GCATTTGATA TGCCCAGCCC CTTGAGTAAG GGGCAAGAAA CCTAAGCCGC CTGGTGGATG GCTCGGCTCG GGGCGCTGAC GAAGG
23SBHB_300      GCAGAAGA CGGGTTTGAT GGGGCGGGGA TGTAAGCGGG AAGGGAAACC GACCCGTTTA GTCTGCCGCT CCCAACAGCC CGAGTTTCTT GCCCCTGAAA GAGGGGCTGG GCATTTGATA TGCCCAGCCC CTTGAGTAAG GGGCAAGAAA CCTAAGCCGC CTGGTGGATG GCTCGGCTCG GGGCGCTGAC GAAG
23SBHB_233               CGGGTTTGAT GGGGCGGGGA TGTAAGCGGG AAGGGAAACC GACCCGTTTA GTCTGCCGCT CCCAACAGCC CGAGTTTCTT GCCCCTGAAA GAGGGGCTGG GCATTTGATA TGCCCAGCCC CTTGAGTAAG GGGCAAGAAA CCTAAGCCGC CTGGTGGATG GCTCGGCTCG GGGCGCTGAC GAAGGGCGTG G
23SBHB_100         GAAGA CGGGTTTGAT GGGGCGGGGA TGTAAGCGGG AAGGGAAACC GACCCGTTTA GTCTGCCGCT CCCAACAGCC CGAGTTTCTT GCCCCTGAAA GAGGGGCTGG GCATTTGATA TGCCCAGCCC CTTGAGTAAG GGGCAAGAAA CCTAAGCCGC CTGGTGGATG GCTCGGCTCG GGGCGCTGAC GAAGGG
23SBHB_821           AGA CGGGTTTGAT GGGGCGGGGA TGTAAGCGGG AAGGGAAACC GACCCGTTTA GTCTGCCGCT CCCAACAGCC CGAGTTTCTT GCCCCTGAAA GAGGGGCTGG GCATTTGATA TGCCCAGCCC CTTGAGTAAG GGGCAAGAAA CCTAAGCCGC CTGGTGGATG GCTCGGCTCG GGGCGCTGAC GAAGGGCG
23SBHB_811          AAGA CGGGTTTGAT GGGGCGGGGA TGTAAGCGGG AAGGGAAACC GACCCGTTTA GTCTGCCGCT CCCAACAGCC CGAGTTTCTT GCCCCTGAAA GAGGGGCTGG GCATTTGATA TGCCCAGCCC CTTGAGTAAG GGGCAAGAAA CCTAAGCCGC CTGGTGGATG GCTCGGCTCG GGGCGCTGAC GAAGGGC
23SBHB_353             A CGGGTTTGAT GGGGCGGGGA TGTAAGCGGG AAGGGAAACC GACCCGTTTA GTCTGCCGCT CCCAACAGCC CGAGTTTCTT GCCCCTGAAA GAGGGGCTGG GCATTTGATA TGCCCAGCCC CTTGAGTAAG GGGCAAGAAA CCTAAGCCGC CTGGTGGATG GCTCGGCTCG GGGCGCTGAC GAAGGGCGTG
23SBHB_492        AGAAGA CGGGTTTGAT GGGGCGGGGA TGTAAGCGGG AAGGGAAACC GACCCGTTTA GTCTGCCGCT CCCAACAGCC CGAGTTTCTT GCCCCTGAAA GAGGGGCTGG GCATTTGATA TGCCCAGCCC CTTGAGTAAG GGGCAAGAAA CCTAAGCCGC CTGGTGGATG GCTCGGCTCG GGGCGCTGAC GAAG
23SBHB_183                GGGTTTGAT GGGGCGGGGA TGTAAGCGGG AAGGGAAACC GACCCGTTTA GTCTGCCGCT CCCAACAGCC CGAGTTTCTT GCCCCTGAAA GAGGGGCTGG GCATTTGATA TGCCCAGCCC CTTGAGTAAG GGGCAAGAAA CCTAAGCCGC CTGGTGGATG GCTCGGCTCG GGGCGCTGAC GAAGGGCGTG G
23SBHB_262            GA CGGGTTTGAT GGGGCGGGGA TGTAAGCGGG AAGGGAAACC GACCCGTTTA GTCTGCCGCT CCCAACAGCC CGAGTTTCTT GCCCCTGAAA GAGGGGCTGG GCATTTGATA TGCCCAGCCC CTTGAGTAAG GGGCAAGAAA CCTAAGCCGC CTGGTGGATG GCTCGGCTCG GGGCGCTGAC GAAGGGCG
23SBHB_152          AAGA CGGGTTTGAT GGGGCGGGGA TGTAAGCGGG AAGGGAAACC GACCCGTTTA GTCTGCCGCT CCCAACAGCC CGAGTTTCTT GCCCCTGAAA GAGGGGCTGG GCATTTGATA TGCCCAGCCC CTTGAGTAAG GGGCAAGAAA CCTAAGCCGC CTGGTGGATG GCTCGGCTCG GGGCGCTGAC GAAGGG
23SBHB_132               CGGGTTTGAT GGGGCGGGGA TGTAAGCGGG AAGGGAAACC GACCCGTTTA GTCTGCCGCT CCCAACAGCC CGAGTTTCTT GCCCCTGAAA GAGGGGCTGG GCATTTGATA TGCCCAGCCC CTTGAGTAAG GGGCAAGAAA CCTAAGCCGC CTGGTGGATG GCTCGGCTCG GGGCGCTGAC GAAGGGCGTG
23SBHB_131     CGCAGAAGA CGGGTTTGAT GGGGCGGGGA TGTAAGCGGG AAGGGAAACC GACCCGTTTA GTCTGCCGCT CCCAACAGCC CGAGTTTCTT GCCCCTGAAA GAGGGGCTGG GCATTTGATA TGCCCAGCCC CTTGAGTAAG GGGCAAGAAA CCTAAGCCGC CTGGTGGATG GCTCGGCTCG GGGCGCTGAC G
23SBHB_568    CCGCAGAAGA CGGGTTTGAT GGGGCGGGGA TGTAAGCGGG AAGGGAAACC GACCCGTTTA GTCTGCCGCT CCCAACAGCC CGAGTTTCTT GCCCCTGAAA GAGGGGCTGG GCATTTGATA TGCCCAGCCC CTTGAGTAAG GGGCAAGAAA CCTAAGCCGC CTGGTGGATG GCTCGGCTCG GGGCGCTGAC G
23SBHB_534  C CCGCAGAAGA CGGGTTTGAT GGGGCGGGGA TGTAAGCGGG AAGGGAAACC GACCCGTTTA GTCTGCCGCT CCCAACAGCC CGAGTTTCTT GCCCCTGAAA GAGGGGCTGG GCATTTGATA TGCCCAGCCC CTTGAGTAAG GGGCAAGAAA CCTAAGCCGC CTGGTGGATG GCTCGGCTCG GGGCGCTGAC G
23SBHB_271  C CCGCAGAAGA CGGGTTTGAT GGGGCGGGGA TGTAAGCGGG AAGGGAAACC GACCCGTTTA GTCTGCCGCT CCCAACAGCC CGAGTTTCTT GCCCCTGAAA GAGGGGCTGG GCATTTGATA TGCCCAGCCC CTTGAGTAAG GGGCAAGAAA CCTAAGCCGC CTGGTGGATG GCTCGGCTCG GGGCGCTGAC
23SBHB_294    CCGCAGAAGA CGGGTTTGAT GGGGCGGGGA TGTAAGCGGG AAGGGAAACC GACCCGTTTA GTCTGCCGCT CCCAACAGCC CGAGTTTCTT GCCCCTGAAA GAGGGGCTGG GCATTTGATA TGCCCAGCCC CTTGAGTAAG GGGCAAGAAA CCTAAGCCGC CTGGTGGATG GCTCGGCTCG GGGCGCTGAC
23SBHB_145     CGCAGAAGA CGGGTTTGAT GGGGCGGGGA TGTAAGCGGG AAGGGAAACC GACCCGTTTA GTCTGCCGCT CCCAACAGCC CGAGTTTCTT GCCCCTGAAA GAGGGGCTGG GCATTTGATA TGCCCAGCCC CTTGAGTAAG GGGCAAGAAA CCTAAGCCGC CTGGTGGATG GCTCGGCTCG GGGCGCTGAC GA
23SBHB_129           AGA CGGGTTTGAT GGGGCGGGGA TGTAAGCGGG AAGGGAAACC GACCCGTTTA GTCTGCCGCT CCCAACAGCC CGAGTTTCTT GCCCCTGAAA GAGGGGCTGG GCATTTGATA TGCCCAGCCC CTTGAGTAAG GGGCAAGAAA CCTAAGCCGC CTGGTGGATG GCTCGGCTCG GGGCGCTGAC GAAGGGC
23SBHB_627         GAAGA CGGGTTTGAT GGGGCGGGGA TGTAAGCGGG AAGGGAAACC GACCCGTTTA GTCTGCCGCT CCCAACAGCC CGAGTTTCTT GCCCCTGAAA GAGGGGCTGG GCATTTGATA TGCCCAGCCC CTTGAGTAAG GGGCAAGAAA CCTAAGCCGC CTGGTGGATG GCTCGGCTCG GGGCGCTGAC GAAGG
23SBHB_408     CGCAGAAGA CGGGTTTGAT GGGGCGGGGA TGTAAGCGGG AAGGGAAACC GACCCGTTTA GTCTGCCGCT CCCAACAGCC CGAGTTTCTT GCCCCTGAAA GAGGGGCTGG GCATTTGATA TGCCCAGCCC CTTGAGTAAG GGGCAAGAAA CCTAAGCCGC CTGGTGGATG GCTCGGCTCG GGGCGCTGAC AAAG
23SBHB_169       CAGAAGA CGGGTTTGAT GGGGCGGGGA TGTAAGCGGG AAGGGAAACC GACCCGTTTA GTCTGCCGCT CCCAACAGCC CGAGTTTCTT GCCCCTGAAA GAGGGGCTGG GCATTTGATA TGCCCAGCCC CTTGAGTAAG GGGCAAGAAA CCTAAGCCGC CTGGTGGATG GCTCGGCTCG GGGCGCTGAC AAAG
23SBHB_162      GCAGAAGA CGGGTTTGAT GGGGCGGGGA TGTAAGCGGG AAGGGAAACC GACCCGTTTA GTCTGCCGCT CCCAACAGCC CGAGTTTCTT GCCCCTGAAA GAGGGGCTGG GCATTTGATA TGCCCAGCCC CTTGAGTAAG GGGCAAGAAA CCTAAGCCGC CTGGTGGATG GCTCGGCTCG GGGCGCTGAC GAA
23SBHB_421            GA CGGGTTTGAT GGGGCGGGGA TGTAAGCGGG AAGGGAAACC GACCCGTTTA GTCTGCCGCT CCCAACAGCC CGAGTTTCTT GCCCCTGAAA GAGGGGCTGG GCATTTGATA TGCCCAGCCC CTTGAGTAAG GGGCAAGAAA CCTAAGCCGC CTGGTGGATG GCTCGGCTCG GGGCGCTGAC GAAGGGC
23SBHB_266           AGA CGGGTTTGAT GGGGCGGGGA TGTAAGCGGG AAGGGAAACC GACCCGTTTA GTCTGCCGCT CCCAACAGCC CGAGTTTCTT GCCCCTGAAA GAGGGGCTGG GCATTTGATA TGCCCAGCCC CTTGAGTAAG GGGCAAGAAA CCTAAGCCGC CTGGTGGATG GCTCGGCTCG GGGCGCTGAC GAAGGG
23SBHB_101             A CGGGTTTGAT GGGGCGGGGA TGTAAGCGGG AAGGGAAACC GACCCGTTTA GTCTGCCGCT CCCAACAGCC CGAGTTTCTT GCCCCTGAAA GAGGGGCTGG GCATTTGATA TGCCCAGCCC CTTGAGTAAG GGGCAAGAAA CCTAAGCCGC CTGGTGGATG GCTCGGCTCG GGGCGCTGAC GAAGGGCG
23SBHB_958          AAGA CGGGTTTGAT GGGGCGGGGA TGTAAGCGGG AAGGGAAACC GACCCGTTTA GTCTGCCGCT CCCAACAGCC CGAGTTTCTT GCCCCTGAAA GAGGGGCTGG GCATTTGATA TGCCCAGCCC CTTGAGTAAG GGGCAAGAAA CCTAAGCCGC CTGGTGGATG GCTCGGCTCG GGGCGCTGAC GAAGG
23SBHB_717         GAAGA CGGGTTTGAT GGGGCGGGGA TGTAAGCGGG AAGGGAAACC GACCCGTTTA GTCTGCCGCT CCCAACAGCC CGAGTTTCTT GCCCCTGAAA GAGGGGCTGG GCATTTGATA TGCCCAGCCC CTTGAGTAAG GGGCAAGAAA CCTAAGCCGC CTGGTGGATG GCTCGGCTCG GGGCGCTGAC GAAG
23SBHB_639                              CGGGGA TGTAAGCGGG AAGGGAAACC GACCCGTTTA GTCTGCCGCT CCCAACAGCC CGAGTTTCTT GCCCCTGAAA GAGGGGCTGG GCATTTGATA TGCCCAGCCC CTTGAGTAAG GGGCAAGAAA CCTAAGCCGC CTGGTGGATG GCTCGGCTCG GGGCGCTGAC GAAGGGCGTG GCAAGCTGCG ATAAGCCCCG GTGAG
23SBHB_173                              CGGGGA TGTAAGCGGG AAGGGAAACC GACCCGTTTA GTCTGCCGCT CCCAACAGCC CGAGTTTCTT GCCCCTGAAA GAGGGGCTGG GCATTTGATA TGCCCAGCCC CTTGAGTAAG GGGCAAGAAA CCTAAGCCGC CTGGTGGATG GCTCGGCTCG GGGCGCTGAC GAAGGGCGTG GCAAGCTGCG ATAAGCCCCG GTGAGGCGCA G
23SBHB_155                              CGGGGA TGTAAGCGGG AAGGGAAACC GACCCGTTTA GTCTGCCGCT CCCAACAGCC CGAGTTTCTT GCCCCTGAAA GAGGGGCTGG GCATTTGATA TGCCCAGCCC CTTGAGTAAG GGGCAAGAAA CCTAAGCCGC CTGGTGGATG GCTCGGCTCG GGGCGCTGAC GAAGGGCGTG GCAAGCTGCG ATAAGCCCCG GTGAGG
23SBHB_127                                GGGA TGTAAGCGGG AAGGGAAACC GACCCGTTTA GTCTGCCGCT CCCAACAGCC CGAGTTTCTT GCCCCTGAAA GAGGGGCTGG GCATTTGATA TGCCCAGCCC CTTGAGTAAG GGGCAAGAAA CCTAAGCCGC CTGGTGGATG GCTCGGCTCG GGGCGCTGAC GAAGGGCGTG GCAAGCTGCG ATAAGCCCCG GTGAG
23SBHB_311                              CGGGGA TGTAAGCGGG AAGGGAAACC GACCCGTTTA GTCTGCCGCT CCCAACAGCC CGAGTTTCTT GCCCCTGAAA GAGGGGCTGG GCATTTGATA TGCCCAGCCC CTTGAGTAAG GGGCAAGAAA CCTAAGCCGC CTGGTGGATG GCTCGGCTCG GGGCGCTGAC GAAGGGCGTG GCAAGCTGCG ATAAGCCCCG G
23SBHB_203                              CGGGGA TGTAAGCGGG AAGGGAAACC GACCCGTTTA GTCTGCCGCT CCCAACAGCC CGAGTTTCTT GCCCCTGAAA GAGGGGCTGG GCATTTGATA TGCCCAGCCC CTTGAGTAAG GGGCAAGAAA CCTAAGCCGC CTGGTGGATG GCTCGGCTCG GGGCGCTGAC GAAGGGCGTG GCAAGCTGCG ATAAGCCCCG
23SBHB_357       CAGAAGA CGGGTTTGAT GGGGCGGGGA TGTAAGCGGG AAGGGAAACC GACCCGTTTA GTCTGCCGCT CCCAACAGCC CGAGTTTCTT GCCCCTGAAA GAGGGGCTGG GCATTTGATA TGCCCAGCCC CTTGAGTAAG GGGCAAGAAA CCTAAGCCGC CTGGTGGATG GCTCGGCTCG GGGCGCTGAC GAA
23SBHB_176        AGAAGA CGGGTTTGAT GGGGCGGGGA TGTAAGCGGG AAGGGAAACC GACCCGTTTA GTCTGCCGCT CCCAACAGCC CGAGTTTCTT GCCCCTGAAA GAGGGGCTGG GCATTTGATA TGCCCAGCCC CTTGAGTAAG GGGCAAGAAA CCTAAGCCGC CTGGTGGATG GCTCGGCTCG GGGCGCTGAC GAA
23SBHB_119                 GGTTTGAT GGGGCGGGGA TGTAAGCGGG AAGGGAAACC GACCCGTTTA GTCTGCCGCT CCCAACAGCC CGAGTTTCTT GCCCCTGAAA GAGGGGCTGG GCATTTGATA TGCCCAGCCC CTTGAGTAAG GGGCAAGAAA CCTAAGCCGC CTGGTGGATG GCTCGGCTCG GGGCGCTGAC GAAGGGCGTG G
23SBHB_733            GA CGGGTTTGAT GGGGCGGGGA TGTAAGCGGG AAGGGAAACC GACCCGTTTA GTCTGCCGCT CCCAACAGCC CGAGTTTCTT GCCCCTGAAA GAGGGGCTGG GCATTTGATA TGCCCAGCCC CTTGAGTAAG GGGCAAGAAA CCTAAGCCGC CTGGTGGATG GCTCGGCTCG GGGCGCTGAC GAAGGG
23SBHB_242            GA CGGGTTTGAT GGGGCGGGGA TGTAAGCGGG AAGGGAAACC GACCCGTTTA GTCTGCCGCT CCCAACAGCC CGAGTTTCTT GCCCCTGAAA GAGGGGCTGG GCATTTGATA TGCCCAGCCC CTTGAGTAAG GGGCAAGAAA CCTAAGCCGC CTGGTGGATG GCTCGGCTCG GGGCGCTGAC GAAGGGG
23SBHB_384               CGGGTTTGAT GGGGCGGGGA TGTAAGCGGG AAGGGAAACC GACCCGTTTA GTCTGCCGCT CCCAACAGCC CGAGTTTCTT GCCCCTGAAA GAGGGGCTGG GCATTTGATA TGCCCAGCCC CTTGAGTAAG GGGCAAGAAA CCTAAGCCGC CTGGTGGATG GCTCGGCTCG GGGCGCTGAC GAAGGGCG
23SBHB_179           AGA CGGGTTTGAT GGGGCGGGGA TGTAAGCGGG AAGGGAAACC GACCCGTTTA GTCTGCCGCT CCCAACAGCC CGAGTTTCTT GCCCCTGAAA GAGGGGCTGG GCATTTGATA TGCCCAGCCC CTTGAGTAAG GGGCAAGAAA CCTAAGCCGC CTGGTGGATG GCTCGGCTCG GGGCGCTGAC GAAGG
23SBHB_162             A CGGGTTTGAT GGGGCGGGGA TGTAAGCGGG AAGGGAAACC GACCCGTTTA GTCTGCCGCT CCCAACAGCC CGAGTTTCTT GCCCCTGAAA GAGGGGCTGG GCATTTGATA TGCCCAGCCC CTTGAGTAAG GGGCAAGAAA CCTAAGCCGC CTGGTGGATG GCTCGGCTCG GGGCGCTGAC GAAGGGC
23SBHB_125          AAGA CGGGTTTGAT GGGGCGGGGA TGTAAGCGGG AAGGGAAACC GACCCGTTTA GTCTGCCGCT CCCAACAGCC CGAGTTTCTT GCCCCTGAAA GAGGGGCTGG GCATTTGATA TGCCCAGCCC CTTGAGTAAG GGGCAAGAAA CCTAAGCCGC CTGGTGGATG GCTCGGCTCG GGGCGCTGAC GAAG
23SBHB_745     CGCAGAAGA CGGGTTTGAT GGGGCGGGGA TGTAAGCGGG AAGGGAAACC GACCCGTTTA GTCTGCCGCT CCCAACAGCC CGAGTTTCTT GCCCCTGAAA GAGGGGCTGG GCATTTGATA TGCCCAGCCC CTTGAGTAAG GGGCAAGAAA CCTAAGCCGC CTGGTGGATG GCTCGGCTCG GGGCGCTGAC
23SBHB_135      GCAGAAGA CGGGTTTGAT GGGGCGGGGA TGTAAGCGGG AAGGGAAACC GACCCGTTTA GTCTGCCGCT CCCAACAGCC CGAGTTTCTT GCCCCTGAAA GAGGGGCTGG GCATTTGATA TGCCCAGCCC CTTGAGTAAG GGGCAAGAAA CCTAAGCCGC CTGGTGGATG GCTCGGCTCG GGGCGCTGAC G
23SBHB_599       CAGAAGA CGGGTTTGAT GGGGCGGGGA TGTAAGCGGG AAGGGAAACC GACCCGTTTA GTCTGCCGCT CCCAACAGCC CGAGTTTCTT GCCCCTGAAA GAGGGGCTGG GCATTTGATA TGCCCAGCCC CTTGAGTAAG GGGCAAGAAA CCTAAGCCGC CTGGTGGATG GCTCGGCTCG GGGCGCTGAC G
23SBHB_606                  GTTTGAT GGGGCGGGGA TGTAAGCGGG AAGGGAAACC GACCCGTTTA GTCTGCCGCT CCCAACAGCC CGAGTTTCTT GCCCCTGAAA GAGGGGCTGG GCATTTGATA TGCCCAGCCC CTTGAGTAAG GGGCAAGAAA CCTAAGCCGC CTGGTGGATG GCTCGGCTCG GGGCGCTGAC GAAGGGCGTG G
23SBHB_135                  GTTTGAT GGGGCGGGGA TGTAAGCGGG AAGGGAAACC GACCCGTTTA GTCTGCCGCT CCCAACAGCC CGAGTTTCTT GCCCCTGAAA GAGGGGCTGG GCATTTGATA TGCCCAGCCC CTTGAGTAAG GGGCAAGAAA CCTAAGCCGC CTGGTGGATG GCTCGGCTCG GGGCGCTGAC GAAGGGCGTG GC
23SBHB_624               CGGGTTTGAT GGGGCGGGGA TGTAAGCGGG AAGGGAAACC GACCCGTTTA GTCTGCCGCT CCCAACAGCC CGAGTTTCTT GCCCCTGAAA GAGGGGCTGG GCATTTGATA TGCCCAGCCC CTTGAGTAAG GGGCAAGAAA CCTAAGCCGC CTGGTGGATG GCTCGGCTCG GGGCGCTGAC GAAGGGC
23SBHB_117               CGGGTTTGAT GGGGCGGGGA TGTAAGCGGG AAGGGAAACC GACCCGTTTA GTCTGCCGCT CCCAACAGCC CGAGTTTCTT GCCCCTGAAA GAGGGGCTGG GCATTTGATA TGCCCAGCCC CTTGAGTAAG GGGCAAGAAA CCTAAGCCGC CTGGTGGATG GCTCGGCTCG GGGCGCTGAC GAAGGGCC
23SBHB_488            GA CGGGTTTGAT GGGGCGGGGA TGTAAGCGGG AAGGGAAACC GACCCGTTTA GTCTGCCGCT CCCAACAGCC CGAGTTTCTT GCCCCTGAAA GAGGGGCTGG GCATTTGATA TGCCCAGCCC CTTGAGTAAG GGGCAAGAAA CCTAAGCCGC CTGGTGGATG GCTCGGCTCG GGGCGCTGAC GAAGG
23SBHB_291             A CGGGTTTGAT GGGGCGGGGA TGTAAGCGGG AAGGGAAACC GACCCGTTTA GTCTGCCGCT CCCAACAGCC CGAGTTTCTT GCCCCTGAAA GAGGGGCTGG GCATTTGATA TGCCCAGCCC CTTGAGTAAG GGGCAAGAAA CCTAAGCCGC CTGGTGGATG GCTCGGCTCG GGGCGCTGAC GAAGGG
23SBHB_195           AGA CGGGTTTGAT GGGGCGGGGA TGTAAGCGGG AAGGGAAACC GACCCGTTTA GTCTGCCGCT CCCAACAGCC CGAGTTTCTT GCCCCTGAAA GAGGGGCTGG GCATTTGATA TGCCCAGCCC CTTGAGTAAG GGGCAAGAAA CCTAAGCCGC CTGGTGGATG GCTCGGCTCG GGGCGCTGAC GAAG
23SBHB_216        AGAAGA CGGGTTTGAT GGGGCGGGGA TGTAAGCGGG AAGGGAAACC GACCCGTTTA GTCTGCCGCT CCCAACAGCC CGAGTTTCTT GCCCCTGAAA GAGGGGCTGG GCATTTGATA TGCCCAGCCC CTTGAGTAAG GGGCAAGAAA CCTAAGCCGC CTGGTGGATG GCTCGGCTCG GGGCGCTGAC G
23SBHB_358                  GTTTGAT GGGGCGGGGA TGTAAGCGGG AAGGGAAACC GACCCGTTTA GTCTGCCGCT CCCAACAGCC CGAGTTTCTT GCCCCTGAAA GAGGGGCTGG GCATTTGATA TGCCCAGCCC CTTGAGTAAG GGGCAAGAAA CCTAAGCCGC CTGGTGGATG GCTCGGCTCG GGGCGCTGAC GAAGGGCGTG
23SBHB_310                GGGTTTGAT GGGGCGGGGA TGTAAGCGGG AAGGGAAACC GACCCGTTTA GTCTGCCGCT CCCAACAGCC CGAGTTTCTT GCCCCTGAAA GAGGGGCTGG GCATTTGATA TGCCCAGCCC CTTGAGTAAG GGGCAAGAAA CCTAAGCCGC CTGGTGGATG GCTCGGCTCG GGGCGCTGAC GAAGGGCG
23SBHB_110               CGGGTTTGAT GGGGCGGGGA TGTAAGCGGG AAGGGAAACC GACCCGTTTA GTCTGCCGCT CCCAACAGCC CGAGTTTCTT GCCCCTGAAA GAGGGGCTGG GCATTTGATA TGCCCAGCCC CTTGAGTAAG GGGCAAGAAA CCTAAGCCGC CTGGTGGATG GCTCGGCTCG GGGCGCTGAC GAAGGG
23SBHB_322               CGGGTTTGAT GGGGCGGGGA TGTAAGCGGG AAGGGAAACC GACCCGTTTA GTCTGCCGCT CCCAACAGCC CGAGTTTCTT GCCCCTGAAA GAGGGGCTGG GCATTTGATA TGCCCAGCCC CTTGAGTAAG GGGCAAGAAA CCTAAGCCGC CTGGTGGATG GCTCGGCTCG GGGCGCTGAC GAAGGGG
23SBHB_561            GA CGGGTTTGAT GGGGCGGGGA TGTAAGCGGG AAGGGAAACC GACCCGTTTA GTCTGCCGCT CCCAACAGCC CGAGTTTCTT GCCCCTGAAA GAGGGGCTGG GCATTTGATA TGCCCAGCCC CTTGAGTAAG GGGCAAGAAA CCTAAGCCGC CTGGTGGATG GCTCGGCTCG GGGCGCTGAC GAAG
23SBHB_203             A CGGGTTTGAT GGGGCGGGGA TGTAAGCGGG AAGGGAAACC GACCCGTTTA GTCTGCCGCT CCCAACAGCC CGAGTTTCTT GCCCCTGAAA GAGGGGCTGG GCATTTGATA TGCCCAGCCC CTTGAGTAAG GGGCAAGAAA CCTAAGCCGC CTGGTGGATG GCTCGGCTCG GGGCGCTGAC GAAGG
23SBHB_515                GGGTTTGAT GGGGCGGGGA TGTAAGCGGG AAGGGAAACC GACCCGTTTA GTCTGCCGCT CCCAACAGCC CGAGTTTCTT GCCCCTGAAA GAGGGGCTGG GCATTTGATA TGCCCAGCCC CTTGAGTAAG GGGCAAGAAA CCTAAGCCGC CTGGTGGATG GCTCGGCTCG GGGCGCTGAC GAAGGGC
```

```
23SBHB_318 CAGAAGA CGGGTTTGAT GGGGCGGGGA TGTAAGCGGG AAGGGAAACC GACCCGTTTA GTCTGCCGCT CCCAACAGCC CGAGTTTCTT GCCCCTGAAA GAGGGGCTGG GCATTTGATA TGCCCAGCCC CTTGAGTAAG GGGCAAGAAA CCTAAGCCGC CTGGTGGATG GCTCGGCTCG GGGCGCTGAC
23SBHB_125 AGAAGA CGGGTTTGAT GGGGCGGGGA TGTAAGCGGG AAGGGAAACC GACCCGTTTA GTCTGCCGCT CCCAACAGCC CGAGTTTCTT GCCCCTGAAA GAGGGGCTGG GCATTTGATA TGCCCAGCCC CTTGAGTAAG GGGCAAGAAA CCTAAGCCGC CTGGTGGATG GCTCGGCTCG GGGCGCTGAC
23SBHB_266 GGTTTGAT GGGGCGGGGA TGTAAGCGGG AAGGGAAACC GACCCGTTTA GTCTGCCGCT CCCAACAGCC CGAGTTTCTT GCCCCTGAAA GAGGGGCTGG GCATTTGATA TGCCCAGCCC CTTGAGTAAG GGGCAAGAAA CCTAAGCCGC CTGGTGGATG GCTCGGCTCG GGGCGCTGAC GAAGGGCG
23SBHB_746 CGGGTTTGAT GGGGCGGGGA TGTAAGCGGG AAGGGAAACC GACCCGTTTA GTCTGCCGCT CCCAACAGCC CGAGTTTCTT GCCCCTGAAA GAGGGGCTGG GCATTTGATA TGCCCAGCCC CTTGAGTAAG GGGCAAGAAA CCTAAGCCGC CTGGTGGATG GCTCGGCTCG GGGCGCTGAC GAAGG
23SBHB_225 A CGGGTTTGAT GGGGCGGGGA TGTAAGCGGG AAGGGAAACC GACCCGTTTA GTCTGCCGCT CCCAACAGCC CGAGTTTCTT GCCCCTGAAA GAGGGGCTGG GCATTTGATA TGCCCAGCCC CTTGAGTAAG GGGCAAGAAA CCTAAGCCGC CTGGTGGATG GCTCGGCTCG GGGCGCTGAC GAAG
23SBHB_102 GTTTGAT GGGGCGGGGA TGTAAGCGGG AAGGGAAACC GACCCGTTTA GTCTGCCGCT CCCAACAGCC CGAGTTTCTT GCCCCTGAAA GAGGGGCTGG GCATTTGATA TGCCCAGCCC CTTGAGTAAG GGGCAAGAAA CCTAAGCCGC CTGGTGGATG GCTCGGCTCG GGGCGCTGAC GAAGGGCG
23SBHB_964 GGGTTTGAT GGGGCGGGGA TGTAAGCGGG AAGGGAAACC GACCCGTTTA GTCTGCCGCT CCCAACAGCC CGAGTTTCTT GCCCCTGAAA GAGGGGCTGG GCATTTGATA TGCCCAGCCC CTTGAGTAAG GGGCAAGAAA CCTAAGCCGC CTGGTGGATG GCTCGGCTCG GGGCGCTGAC GAAGGG
23SBHB_350 GGTTTGAT GGGGCGGGGA TGTAAGCGGG AAGGGAAACC GACCCGTTTA GTCTGCCGCT CCCAACAGCC CGAGTTTCTT GCCCCTGAAA GAGGGGCTGG GCATTTGATA TGCCCAGCCC CTTGAGTAAG GGGCAAGAAA CCTAAGCCGC CTGGTGGATG GCTCGGCTCG GGGCGCTGAC GAAGGGC
23SBHB_165 CGGGTTTGAT GGGGCGGGGA TGTAAGCGGG AAGGGAAACC GACCCGTTTA GTCTGCCGCT CCCAACAGCC CGAGTTTCTT GCCCCTGAAA GAGGGGCTGG GCATTTGATA TGCCCAGCCC CTTGAGTAAG GGGCAAGAAA CCTAAGCCGC CTGGTGGATG GCTCGGCTCG GGGCGCTGAC GAAGAG
23SBHB_277 GA CGGGTTTGAT GGGGCGGGGA TGTAAGCGGG AAGGGAAACC GACCCGTTTA GTCTGCCGCT CCCAACAGCC CGAGTTTCTT GCCCCTGAAA GAGGGGCTGG GCATTTGATA TGCCCAGCCC CTTGAGTAAG GGGCAAGAAA CCTAAGCCGC CTGGTGGATG GCTCGGCTCG GGGCGCTGAC GAA
23SBHB_861 CGGGTTTGAT GGGGCGGGGA TGTAAGCGGG AAGGGAAACC GACCCGTTTA GTCTGCCGCT CCCAACAGCC CGAGTTTCTT GCCCCTGAAA GAGGGGCTGG GCATTTGATA TGCCCAGCCC CTTGAGTAAG GGGCAAGAAA CCTAAGCCGC CTGGTGGATG GCTCGGCTCG GGGCGCTGAC GAAG
23SBHB_117 A CGGGTTTGAT GGGGCGGGGA TGTAAGCGGG AAGGGAAACC GACCCGTTTA GTCTGCCGCT CCCAACAGCC CGAGTTTCTT GCCCCTGAAA GAGGGGCTGG GCATTTGATA TGCCCAGCCC CTTGAGTAAG GGGCAAGAAA CCTAAGCCGC CTGGTGGATG GCTCGGCTCG GGGCGCTGAC GAA
23SBHB_154 GTTTGAT GGGGCGGGGA TGTAAGCGGG AAGGGAAACC GACCCGTTTA GTCTGCCGCT CCCAACAGCC CGAGTTTCTT GCCCCTGAAA GAGGGGCTGG GCATTTGATA TGCCCAGCCC CTTGAGTAAG GGGCAAGAAA CCTAAGCCGC CTGGTGGATG GCTCGGCTCG GGGCGCTGAC GAAGGGC
23SBHB_804 CGGGGA TGTAAGCGGG AAGGGAAACC GACCCGTTTA GTCTGCCGCT CCCAACAGCC CGAGTTTCTT GCCCCTGAAA GAGGGGCTGG GCATTTGATA TGCCCAGCCC CTTGAGTAAG GGGCAAGAAA CCTAAGCCGC CTGGTGGATG GCTCGGCTCG GGGCGCTGAC GAAGGGCGTG GCAAGCTGCG
23SBHB_351 CGGGGA TGTAAGCGGG AAGGGAAACC GACCCGTTTA GTCTGCCGCT CCCAACAGCC CGAGTTTCTT GCCCCTGAAA GAGGGGCTGG GCATTTGATA TGCCCAGCCC CTTGAGTAAG GGGCAAGAAA CCTAAGCCGC CTGGTGGATG GCTCGGCTCG GGGCGCTGAC GAAGGGCGTG GCAAGCTGCG AT
23SBHB_162 GGCGGGGA TGTAAGCGGG AAGGGAAACC GACCCGTTTA GTCTGCCGCT CCCAACAGCC CGAGTTTCTT GCCCCTGAAA GAGGGGCTGG GCATTTGATA TGCCCAGCCC CTTGAGTAAG GGGCAAGAAA CCTAAGCCGC CTGGTGGATG GCTCGGCTCG GGGCGCTGAC GAAGGGCGTG GCAAGCTGCG
23SBHB_136 GGGGCGGGGA TGTAAGCGGG AAGGGAAACC GACCCGTTTA GTCTGCCGCT CCCAACAGCC CGAGTTTCTT GCCCCTGAAA GAGGGGCTGG GCATTTGATA TGCCCAGCCC CTTGAGTAAG GGGCAAGAAA CCTAAGCCGC CTGGTGGATG GCTCGGCTCG GGGCGCTGAC GAAGGGCGTG GCAAGCTGCG
23SBHB_153 CGGGGA TGTAAGCGGG AAGGGAAACC GACCCGTTTA GTCTGCCGCT CCCAACAGCC CGAGTTTCTT GCCCCTGAAA GAGGGGCTGG GCATTTGATA TGCCCAGCCC CTTGAGTAAG GGGCAAGAAA CCTAAGCCGC CTGGTGGATG GCTCGGCTCG GGGCGCTGAC GAAGGGCGTG GCAAGCTGC
23SBHB_159 GGGA TGTAAGCGGG AAGGGAAACC GACCCGTTTA GTCTGCCGCT CCCAACAGCC CGAGTTTCTT GCCCCTGAAA GAGGGGCTGG GCATTTGATA TGCCCAGCCC CTTGAGTAAG GGGCAAGAAA CCTAAGCCGC CTGGTGGATG GCTCGGCTCG GGGCGCTGAC GAAGGGCGTG GCAAGCTGCG
23SBHB_240 CGGGGA TGTAAGCGGG AAGGGAAACC GACCCGTTTA GTCTGCCGCT CCCAACAGCC CGAGTTTCTT GCCCCTGAAA GAGGGGCTGG GCATTTGATA TGCCCAGCCC CTTGAGTAAG GGGCAAGAAA CCTAAGCCGC CTGGTGGATG GCTCGGCTCG GGGCGCTGAC GAAGGGCGTG GCAAGCTG
23SBHB_728 GGTTTGAT GGGGCGGGGA TGTAAGCGGG AAGGGAAACC GACCCGTTTA GTCTGCCGCT CCCAACAGCC CGAGTTTCTT GCCCCTGAAA GAGGGGCTGG GCATTTGATA TGCCCAGCCC CTTGAGTAAG GGGCAAGAAA CCTAAGCCGC CTGGTGGATG GCTCGGCTCG GGGCGCTGAC GAAGGG
23SBHB_658 GGGTTTGAT GGGGCGGGGA TGTAAGCGGG AAGGGAAACC GACCCGTTTA GTCTGCCGCT CCCAACAGCC CGAGTTTCTT GCCCCTGAAA GAGGGGCTGG GCATTTGATA TGCCCAGCCC CTTGAGTAAG GGGCAAGAAA CCTAAGCCGC CTGGTGGATG GCTCGGCTCG GGGCGCTGAC GAAGG
23SBHB_306 GTTTGAT GGGGCGGGGA TGTAAGCGGG AAGGGAAACC GACCCGTTTA GTCTGCCGCT CCCAACAGCC CGAGTTTCTT GCCCCTGAAA GAGGGGCTGG GCATTTGATA TGCCCAGCCC CTTGAGTAAG GGGCAAGAAA CCTAAGCCGC CTGGTGGATG GCTCGGCTCG GGGCGCTGAC GAAGGG
23SBHB_772 GGGTTTGAT GGGGCGGGGA TGTAAGCGGG AAGGGAAACC GACCCGTTTA GTCTGCCGCT CCCAACAGCC CGAGTTTCTT GCCCCTGAAA GAGGGGCTGG GCATTTGATA TGCCCAGCCC CTTGAGTAAG GGGCAAGAAA CCTAAGCCGC CTGGTGGATG GCTCGGCTCG GGGCGCTGAC GAAG
23SBHB_488 GGTTTGAT GGGGCGGGGA TGTAAGCGGG AAGGGAAACC GACCCGTTTA GTCTGCCGCT CCCAACAGCC CGAGTTTCTT GCCCCTGAAA GAGGGGCTGG GCATTTGATA TGCCCAGCCC CTTGAGTAAG GGGCAAGAAA CCTAAGCCGC CTGGTGGATG GCTCGGCTCG GGGCGCTGAC GAAGG
23SBHB_418 CGGGTTTGAT GGGGCGGGGA TGTAAGCGGG AAGGGAAACC GACCCGTTTA GTCTGCCGCT CCCAACAGCC CGAGTTTCTT GCCCCTGAAA GAGGGGCTGG GCATTTGATA TGCCCAGCCC CTTGAGTAAG GGGCAAGAAA CCTAAGCCGC CTGGTGGATG GCTCGGCTCG GGGCGCTGAC GAA
23SBHB_122 CGGGTTTGAT GGGGCGGGGA TGTAAGCGGG AAGGGAAACC GACCCGTTTA GTCTGCCGCT CCCAACAGCC CGAGTTTCTT GCCCCTGAAA GAGGGGCTGG GCATTTGATA TGCCCAGCCC CTTGAGTAAG GGGCAAGAAA CCTAAGCCGC CTGGTGGATG GCTCGGCTCG GGGCGCTGAC AAAG
23SBHB_342 GA CGGGTTTGAT GGGGCGGGGA TGTAAGCGGG AAGGGAAACC GACCCGTTTA GTCTGCCGCT CCCAACAGCC CGAGTTTCTT GCCCCTGAAA GAGGGGCTGG GCATTTGATA TGCCCAGCCC CTTGAGTAAG GGGCAAGAAA CCTAAGCCGC CTGGTGGATG GCTCGGCTCG GGGCGCTGAC G
23SBHB_195 GTTTGAT GGGGCGGGGA TGTAAGCGGG AAGGGAAACC GACCCGTTTA GTCTGCCGCT CCCAACAGCC CGAGTTTCTT GCCCCTGAAA GAGGGGCTGG GCATTTGATA TGCCCAGCCC CTTGAGTAAG GGGCAAGAAA CCTAAGCCGC CTGGTGGATG GCTCGGCTCG GGGCGCTGAC GAAGG
23SBHB_115 A CGGGTTTGAT GGGGCGGGGA TGTAAGCGGG AAGGGAAACC GACCCGTTTA GTCTGCCGCT CCCAACAGCC CGAGTTTCTT GCCCCTGAAA GAGGGGCTGG GCATTTGATA TGCCCAGCCC CTTGAGTAAG GGGCAAGAAA CCTAAGCCGC CTGGTGGATG GCTCGGCTCG GGGCGCTGAC G
23SBHB_581 GGTTTGAT GGGGCGGGGA TGTAAGCGGG AAGGGAAACC GACCCGTTTA GTCTGCCGCT CCCAACAGCC CGAGTTTCTT GCCCCTGAAA GAGGGGCTGG GCATTTGATA TGCCCAGCCC CTTGAGTAAG GGGCAAGAAA CCTAAGCCGC CTGGTGGATG GCTCGGCTCG GGGCGCTGAC GAAG
23SBHB_251 GA CGGGTTTGAT GGGGCGGGGA TGTAAGCGGG AAGGGAAACC GACCCGTTTA GTCTGCCGCT CCCAACAGCC CGAGTTTCTT GCCCCTGAAA GAGGGGCTGG GCATTTGATA TGCCCAGCCC CTTGAGTAAG GGGCAAGAAA CCTAAGCCGC CTGGTGGATG GCTCGGCTCG GGGCGCTGAC
23SBHB_249 GTTTGAT GGGGCGGGGA TGTAAGCGGG AAGGGAAACC GACCCGTTTA GTCTGCCGCT CCCAACAGCC CGAGTTTCTT GCCCCTGAAA GAGGGGCTGG GCATTTGATA TGCCCAGCCC CTTGAGTAAG GGGCAAGAAA CCTAAGCCGC CTGGTGGATG GCTCGGCTCG GGGCGCTGAC GAAG
23SBHB_460 CGGGTTTGAT GGGGCGGGGA TGTAAGCGGG AAGGGAAACC GACCCGTTTA GTCTGCCGCT CCCAACAGCC CGAGTTTCTT GCCCCTGAAA GAGGGGCTGG GCATTTGATA TGCCCAGCCC CTTGAGTAAG GGGCAAGAAA CCTAAGCCGC CTGGTGGATG GCTCGGCTCG GGGCGCTGAC G
23SBHB_124 TGAT GGGGCGGGGA TGTAAGCGGG AAGGGAAACC GACCCGTTTA GTCTGCCGCT CCCAACAGCC CGAGTTTCTT GCCCCTGAAA GAGGGGCTGG GCATTTGATA TGCCCAGCCC CTTGAGTAAG GGGCAAGAAA CCTAAGCCGC CTGGTGGATG GCTCGGCTCG GGGCGCTGAC GAAGGG
23SBHB_133 GAT GGGGCGGGGA TGTAAGCGGG AAGGGAAACC GACCCGTTTA GTCTGCCGCT CCCAACAGCC CGAGTTTCTT GCCCCTGAAA GAGGGGCTGG GCATTTGATA TGCCCAGCCC CTTGAGTAAG GGGCAAGAAA CCTAAGCCGC CTGGTGGATG GCTCGGCTCG GGGCGCTGAC GAAGGG
23SBHB_266 CGCAGAAGA CGGGTTTGAT GGGGCGGGGA TGTAAGCGGG AAGGGAAACC GACCCGTTTA GTCTGCCGCT CCCAACAGCC CGAGTTTCTT GCCCCTGAAA GAGGGGCTGG GCATTTGATA TGCCCAGCCC CTTGAGTAAG GGGCAAGAAA CCTAAGCCGC CTGGTGGATG GCTCGGCTCG GG
23SBHB_115 CCGCAGAAGA CGGGTTTGAT GGGGCGGGGA TGTAAGCGGG AAGGGAAACC GACCCGTTTA GTCTGCCGCT CCCAACAGCC CGAGTTTCTT GCCCCTGAAA GAGGGGCTGG GCATTTGATA TGCCCAGCCC CTTGAGTAAG GGGCAAGAAA CCTAAGCCGC CTGGTGGATG GCTCGGCTCG GG
23SBHB_124 CCGCAGAAGA CGGGTTTGAT GGGGCGGGGA TGTAAGCGGG AAGGGAAACC GACCCGTTTA GTCTGCCGCT CCCAACAGCC CGAGTTTCTT GCCCCTGAAA GAGGGGCTGG GCATTTGATA TGCCCAGCCC CTTGAGTAAG GGGCAAGAAA CCTAAGCCGC CTGGTGGATG GCTCGGCTCG GGG
23SBHB_101 C CCGCAGAAGA CGGGTTTGAT GGGGCGGGGA TGTAAGCGGG AAGGGAAACC GACCCGTTTA GTCTGCCGCT CCCAACAGCC CGAGTTTCTT GCCCCTGAAA GAGGGGCTGG GCATTTGATA TGCCCAGCCC CTTGAGTAAG GGGCAAGAAA CCTAAGCCGC CTGGTGGATG GCTCGGCTCG GG
23SBHB_136 CTCCCTGGCG CAGGGCCTGG GTGACCGGCC CATGCCAGGG ACGAGGGCGC CCGCAGAAGA CGGGTTTGAT GGGGCGGGGA TGTAAGCGGG AAGGGAAACC GACCCGTTTA GTCTGCCGCT CCCAACAGCC CGAGTTTCTT GCCCCTGAAA GAGGGGCTGG GCATTTGATA TGCCCAGCCC CTTGAGTAAG GGGCAAGAAA CCTAAGCCGC CTGGTGGATG GCTCGGCTCG GG
23SBHB_175 C CCGCAGAAGA CGGGTTTGAT GGGGCGGGGA TGTAAGCGGG AAGGGAAACC GACCCGTTTA GTCTGCCGCT CCCAACAGCC CGAGTTTCTT GCCCCTGAAA GAGGGGCTGG GCATTTGATA TGCCCAGCCC CTTGAGTAAG GGGCAAGAAA CCTAAGCCGC CTGGTGGATG GCTCGGCTCG G
23SBHB_182 CCGCAGAAGA CGGGTTTGAT GGGGCGGGGA TGTAAGCGGG AAGGGAAACC GACCCGTTTA GTCTGCCGCT CCCAACAGCC CGAGTTTCTT GCCCCTGAAA GAGGGGCTGG GCATTTGATA TGCCCAGCCC CTTGAGTAAG GGGCAAGAAA CCTAAGCCGC CTGGTGGATG GCTCGGCTCG G
23SBHB_253 CGCAGAAGA CGGGTTTGAT GGGGCGGGGA TGTAAGCGGG AAGGGAAACC GACCCGTTTA GTCTGCCGCT CCCAACAGCC CGAGTTTCTT GCCCCTGAAA GAGGGGCTGG GCATTTGATA TGCCCAGCCC CTTGAGTAAG GGGCAAGAAA CCTAAGCCGC CTGGTGGATG GCTCGGCTCG GGG
23SBHB_157 CCGCAGAAGA CGGGTTTGAT GGGGCGGGGA TGTAAGCGGG AAGGGAAACC GACCCGTTTA GTCTGCCGCT CCCAACAGCC CGAGTTTCTT GCCCCTGAAA GAGGGGCTGG GCATTTGATA TGCCCAGCCC CTTGAGTAAG GGGCAAGAAA CCTAAGCCGC CTGGTGGATG GCTCGGCTCG
23SBHB_390 CGCAGAAGA CGGGTTTGAT GGGGCGGGGA TGTAAGCGGG AAGGGAAACC GACCCGTTTA GTCTGCCGCT CCCAACAGCC CGAGTTTCTT GCCCCTGAAA GAGGGGCTGG GCATTTGATA TGCCCAGCCC CTTGAGTAAG GGGCAAGAAA CCTAAGCCGC CTGGTGGATG GCTCGGCTCG G
23SBHB_280 GCAGAAGA CGGGTTTGAT GGGGCGGGGA TGTAAGCGGG AAGGGAAACC GACCCGTTTA GTCTGCCGCT CCCAACAGCC CGAGTTTCTT GCCCCTGAAA GAGGGGCTGG GCATTTGATA TGCCCAGCCC CTTGAGTAAG GGGCAAGAAA CCTAAGCCGC CTGGTGGATG GCTCGGCTCG GG
23SBHB_328 CGGGTTTGAT GGGGCGGGGA TGTAAGCGGG AAGGGAAACC GACCCGTTTA GTCTGCCGCT CCCAACAGCC CGAGTTTCTT GCCCCTGAAA GAGGGGCTGG GCATTTGATA TGCCCAGCCC CTTGAGTAAG GGGCAAGAAA CCTAAGCCGC CTGGTGGATG GCTCGGCTCG GGGCGCTGAC
23SBHB_115 TGAT GGGGCGGGGA TGTAAGCGGG AAGGGAAACC GACCCGTTTA GTCTGCCGCT CCCAACAGCC CGAGTTTCTT GCCCCTGAAA GAGGGGCTGG GCATTTGATA TGCCCAGCCC CTTGAGTAAG GGGCAAGAAA CCTAAGCCGC CTGGTGGATG GCTCGGCTCG GGGCGCTGAC GAAG
23SBHB_196 CGGGGA TGTAAGCGGG AAGGGAAACC GACCCGTTTA GTCTGCCGCT CCCAACAGCC CGAGTTTCTT GCCCCTGAAA GAGGGGCTGG GCATTTGATA TGCCCAGCCC CTTGAGTAAG GGGCAAGAAA CCTAAGCCGC CTGGTGGATG GCTCGGCTCG GGGCGCTGAC GAAGGGCGTG GCAAG
23SBHB_118 GGGGCGGGGA TGTAAGCGGG AAGGGAAACC GACCCGTTTA GTCTGCCGCT CCCAACAGCC CGAGTTTCTT GCCCCTGAAA GAGGGGCTGG GCATTTGATA TGCCCAGCCC CTTGAGTAAG GGGCAAGAAA CCTAAGCCGC CTGGTGGATG GCTCGGCTCG GGGCGCTGAC GAAGGGCG
23SBHB_989 CAGAAGA CGGGTTTGAT GGGGCGGGGA TGTAAGCGGG AAGGGAAACC GACCCGTTTA GTCTGCCGCT CCCAACAGCC CGAGTTTCTT GCCCCTGAAA GAGGGGCTGG GCATTTGATA TGCCCAGCCC CTTGAGTAAG GGGCAAGAAA CCTAAGCCGC CTGGTGGATG GCTCGGCTCG GG
23SBHB_423 AGAAGA CGGGTTTGAT GGGGCGGGGA TGTAAGCGGG AAGGGAAACC GACCCGTTTA GTCTGCCGCT CCCAACAGCC CGAGTTTCTT GCCCCTGAAA GAGGGGCTGG GCATTTGATA TGCCCAGCCC CTTGAGTAAG GGGCAAGAAA CCTAAGCCGC CTGGTGGATG GCTCGGCTCG GG
23SBHB_152 CAGAAGA CGGGTTTGAT GGGGCGGGGA TGTAAGCGGG AAGGGAAACC GACCCGTTTA GTCTGCCGCT CCCAACAGCC CGAGTTTCTT GCCCCTGAAA GAGGGGCTGG GCATTTGATA TGCCCAGCCC CTTGAGTAAG GGGCAAGAAA CCTAAGCCGC CTGGTGGATG GCTCGGCTCG G
23SBHB_166 T GGGGCGGGGA TGTAAGCGGG AAGGGAAACC GACCCGTTTA GTCTGCCGCT CCCAACAGCC CGAGTTTCTT GCCCCTGAAA GAGGGGCTGG GCATTTGATA TGCCCAGCCC CTTGAGTAAG GGGCAAGAAA CCTAAGCCGC CTGGTGGATG GCTCGGCTCG GGGCGCTGAC GAAGGG
23SBHB_116 GTTTGAT GGGGCGGGGA TGTAAGCGGG AAGGGAAACC GACCCGTTTA GTCTGCCGCT CCCAACAGCC CGAGTTTCTT GCCCCTGAAA GAGGGGCTGG GCATTTGATA TGCCCAGCCC CTTGAGTAAG GGGCAAGAAA CCTAAGCCGC CTGGTGGATG GCTCGGCTCG GGGCGCTGAC G
23SBHB_131 GAT GGGGCGGGGA TGTAAGCGGG AAGGGAAACC GACCCGTTTA GTCTGCCGCT CCCAACAGCC CGAGTTTCTT GCCCCTGAAA GAGGGGCTGG GCATTTGATA TGCCCAGCCC CTTGAGTAAG GGGCAAGAAA CCTAAGCCGC CTGGTGGATG GCTCGGCTCG GGGCGCTGAC GAAG
23SBHB_356 CGCAGAAGA CGGGTTTGAT GGGGCGGGGA TGTAAGCGGG AAGGGAAACC GACCCGTTTA GTCTGCCGCT CCCAACAGCC CGAGTTTCTT GCCCCTGAAA GAGGGGCTGG GCATTTGATA TGCCCAGCCC CTTGAGTAAG GGGCAAGAAA CCTAAGCCGC CTGGTGGATG GCTCGGCTCG
23SBHB_222 GGGGCGGGGA TGTAAGCGGG AAGGGAAACC GACCCGTTTA GTCTGCCGCT CCCAACAGCC CGAGTTTCTT GCCCCTGAAA GAGGGGCTGG GCATTTGATA TGCCCAGCCC CTTGAGTAAG GGGCAAGAAA CCTAAGCCGC CTGGTGGATG GCTCGGCTCG GGGCGCTGAC GAAGGGC
23SBHB_184 CGGGGA TGTAAGCGGG AAGGGAAACC GACCCGTTTA GTCTGCCGCT CCCAACAGCC CGAGTTTCTT GCCCCTGAAA GAGGGGCTGG GCATTTGATA TGCCCAGCCC CTTGAGTAAG GGGCAAGAAA CCTAAGCCGC CTGGTGGATG GCTCGGCTCG GGGCGCTGAC GAAGGGCGTG G
23SBHB_430 CGGGGA TGTAAGCGGG AAGGGAAACC GACCCGTTTA GTCTGCCGCT CCCAACAGCC CGAGTTTCTT GCCCCTGAAA GAGGGGCTGG GCATTTGATA TGCCCAGCCC CTTGAGTAAG GGGCAAGAAA CCTAAGCCGC CTGGTGGATG GCTCGGCTCG GGGCGCTGAC GAAGGGCGTG GC
23SBHB_120 GTTTGAT GGGGCGGGGA TGTAAGCGGG AAGGGAAACC GACCCGTTTA GTCTGCCGCT CCCAACAGCC CGAGTTTCTT GCCCCTGAAA GAGGGGCTGG GCATTTGATA TGCCCAGCCC CTTGAGTAAG GGGCAAGAAA CCTAAGCCGC CTGGTGGATG GCTCGGCTCG GGGCGCTGAC
23SBHB_133 T GGGGCGGGGA TGTAAGCGGG AAGGGAAACC GACCCGTTTA GTCTGCCGCT CCCAACAGCC CGAGTTTCTT GCCCCTGAAA GAGGGGCTGG GCATTTGATA TGCCCAGCCC CTTGAGTAAG GGGCAAGAAA CCTAAGCCGC CTGGTGGATG GCTCGGCTCG GGGCGCTGAC GAAGG
23SBHB_418 GGGGCGGGGA TGTAAGCGGG AAGGGAAACC GACCCGTTTA GTCTGCCGCT CCCAACAGCC CGAGTTTCTT GCCCCTGAAA GAGGGGCTGG GCATTTGATA TGCCCAGCCC CTTGAGTAAG GGGCAAGAAA CCTAAGCCGC CTGGTGGATG GCTCGGCTCG GGGCGCTGAC GAAGGG
23SBHB_219 GGGCGGGGA TGTAAGCGGG AAGGGAAACC GACCCGTTTA GTCTGCCGCT CCCAACAGCC CGAGTTTCTT GCCCCTGAAA GAGGGGCTGG GCATTTGATA TGCCCAGCCC CTTGAGTAAG GGGCAAGAAA CCTAAGCCGC CTGGTGGATG GCTCGGCTCG GGGCGCTGAC GAAGGG
23SBHB_193 GGCGGGGA TGTAAGCGGG AAGGGAAACC GACCCGTTTA GTCTGCCGCT CCCAACAGCC CGAGTTTCTT GCCCCTGAAA GAGGGGCTGG GCATTTGATA TGCCCAGCCC CTTGAGTAAG GGGCAAGAAA CCTAAGCCGC CTGGTGGATG GCTCGGCTCG GGGCGCTGAC GAAGGGCG
23SBHB_110 CGGGGA TGTAAGCGGG AAGGGAAACC GACCCGTTTA GTCTGCCGCT CCCAACAGCC CGAGTTTCTT GCCCCTGAAA GAGGGGCTGG GCATTTGATA TGCCCAGCCC CTTGAGTAAG GGGCAAGAAA CCTAAGCCGC CTGGTGGATG GCTCGGCTCG GGGCGCTGAC GAAGGGCGTG
23SBHB_139 AAGA CGGGTTTGAT GGGGCGGGGA TGTAAGCGGG AAGGGAAACC GACCCGTTTA GTCTGCCGCT CCCAACAGCC CGAGTTTCTT GCCCCTGAAA GAGGGGCTGG GCATTTGATA TGCCCAGCCC CTTGAGTAAG GGGCAAGAAA CCTAAGCCGC CTGGTGGATG GCTCGGCTCG GG
23SBHB_161 T GGGGCGGGGA TGTAAGCGGG AAGGGAAACC GACCCGTTTA GTCTGCCGCT CCCAACAGCC CGAGTTTCTT GCCCCTGAAA GAGGGGCTGG GCATTTGATA TGCCCAGCCC CTTGAGTAAG GGGCAAGAAA CCTAAGCCGC CTGGTGGATG GCTCGGCTCG GGGCGCTGAC GAAG
23SBHB_116 GGGCGGGGA TGTAAGCGGG AAGGGAAACC GACCCGTTTA GTCTGCCGCT CCCAACAGCC CGAGTTTCTT GCCCCTGAAA GAGGGGCTGG GCATTTGATA TGCCCAGCCC CTTGAGTAAG GGGCAAGAAA CCTAAGCCGC CTGGTGGATG GCTCGGCTCG GGGCGCTGAC GAAGGGC
23SBHB_282 GGGGCGGGGA TGTAAGCGGG AAGGGAAACC GACCCGTTTA GTCTGCCGCT CCCAACAGCC CGAGTTTCTT GCCCCTGAAA GAGGGGCTGG GCATTTGATA TGCCCAGCCC CTTGAGTAAG GGGCAAGAAA CCTAAGCCGC CTGGTGGATG GCTCGGCTCG GGGCGCTGAC GAAGG
23SBHB_141 GTAAGCGGG AAGGGAAACC GACCCGTTTA GTCTGCCGCT CCCAACAGCC CGAGTTTCTT GCCCCTGAAA GAGGGGCTGG GCATTTGATA TGCCCAGCCC CTTGAGTAAG GGGCAAGAAA CCTAAGCCGC CTGGTGGATG GCTCGGCTCG GGGCGCTGAC GAAGGGCGTG GCAAGCTGCG
23SBHB_138 GCGGGGA TGTAAGCGGG AAGGGAAACC GACCCGTTTA GTCTGCCGCT CCCAACAGCC CGAGTTTCTT GCCCCTGAAA GAGGGGCTGG GCATTTGATA TGCCCAGCCC CTTGAGTAAG GGGCAAGAAA CCTAAGCCGC CTGGTGGATG GCTCGGCTCG GGGCGCTGAC GAAGGGCG
23SBHB_149 GGGCGGGGA TGTAAGCGGG AAGGGAAACC GACCCGTTTA GTCTGCCGCT CCCAACAGCC CGAGTTTCTT GCCCCTGAAA GAGGGGCTGG GCATTTGATA TGCCCAGCCC CTTGAGTAAG GGGCAAGAAA CCTAAGCCGC CTGGTGGATG GCTCGGCTCG GGGCGCTGAC GAAGG
23SBHB_275 AGA CGGGTTTGAT GGGGCGGGGA TGTAAGCGGG AAGGGAAACC GACCCGTTTA GTCTGCCGCT CCCAACAGCC CGAGTTTCTT GCCCCTGAAA GAGGGGCTGG GCATTTGATA TGCCCAGCCC CTTGAGTAAG GGGCAAGAAA CCTAAGCCGC CTGGTGGATG GCTCGGCTCG GG
23SBHB_494 GGCGGGGA TGTAAGCGGG AAGGGAAACC GACCCGTTTA GTCTGCCGCT CCCAACAGCC CGAGTTTCTT GCCCCTGAAA GAGGGGCTGG GCATTTGATA TGCCCAGCCC CTTGAGTAAG GGGCAAGAAA CCTAAGCCGC CTGGTGGATG GCTCGGCTCG GGGCGCTGAC GAAGGG
23SBHB_224 CGCAGAAGA CGGGTTTGAT GGGGCGGGGA TGTAAGCGGG AAGGGAAACC GACCCGTTTA GTCTGCCGCT CCCAACAGCC CGAGTTTCTT GCCCCTGAAA GAGGGGCTGG GCATTTGATA TGCCCAGCCC CTTGAGTAAG GGGCAAGAAA CCTAAGCCGC CTGGTGGATG GCTCGGCT
23SBHB_166 CGCAGAAGA CGGGTTTGAT GGGGCGGGGA TGTAAGCGGG AAGGGAAACC GACCCGTTTA GTCTGCCGCT CCCAACAGCC CGAGTTTCTT GCCCCTGAAA GAGGGGCTGG GCATTTGATA TGCCCAGCCC CTTGAGTAAG GGGCAAGAAA CCTAAGCCGC CTGGTGGATG GCTCGGC
23SBHB_117 CCGCAGAAGA CGGGTTTGAT GGGGCGGGGA TGTAAGCGGG AAGGGAAACC GACCCGTTTA GTCTGCCGCT CCCAACAGCC CGAGTTTCTT GCCCCTGAAA GAGGGGCTGG GCATTTGATA TGCCCAGCCC CTTGAGTAAG GGGCAAGAAA CCTAAGCCGC CTGGTGGATG GCTCGGC
23SBHB_207 CGCAGAAGA CGGGTTTGAT GGGGCGGGGA TGTAAGCGGG AAGGGAAACC GACCCGTTTA GTCTGCCGCT CCCAACAGCC CGAGTTTCTT GCCCCTGAAA GAGGGGCTGG GCATTTGATA TGCCCAGCCC CTTGAGTAAG GGGCAAGAAA CCTAAGCCGC CTGGTGGATG GCTCGG
23SBHB_344 GGGGCGGGGA TGTAAGCGGG AAGGGAAACC GACCCGTTTA GTCTGCCGCT CCCAACAGCC CGAGTTTCTT GCCCCTGAAA GAGGGGCTGG GCATTTGATA TGCCCAGCCC CTTGAGTAAG GGGCAAGAAA CCTAAGCCGC CTGGTGGATG GCTCGGCTCG GGGCGCTGAC GAAG
23SBHB_312 GGCGGGGA TGTAAGCGGG AAGGGAAACC GACCCGTTTA GTCTGCCGCT CCCAACAGCC CGAGTTTCTT GCCCCTGAAA GAGGGGCTGG GCATTTGATA TGCCCAGCCC CTTGAGTAAG GGGCAAGAAA CCTAAGCCGC CTGGTGGATG GCTCGGCTCG GGGCGCTGAC GAAGGGC
23SBHB_285 GGGA TGTAAGCGGG AAGGGAAACC GACCCGTTTA GTCTGCCGCT CCCAACAGCC CGAGTTTCTT GCCCCTGAAA GAGGGGCTGG GCATTTGATA TGCCCAGCCC CTTGAGTAAG GGGCAAGAAA CCTAAGCCGC CTGGTGGATG GCTCGGCTCG GGGCGCTGAC GAAGGGCGTG G
23SBHB_202 GGGGA TGTAAGCGGG AAGGGAAACC GACCCGTTTA GTCTGCCGCT CCCAACAGCC CGAGTTTCTT GCCCCTGAAA GAGGGGCTGG GCATTTGATA TGCCCAGCCC CTTGAGTAAG GGGCAAGAAA CCTAAGCCGC CTGGTGGATG GCTCGGCTCG GGGCGCTGAC GAAGGGCGTG G
23SBHB_123 GGGGA TGTAAGCGGG AAGGGAAACC GACCCGTTTA GTCTGCCGCT CCCAACAGCC CGAGTTTCTT GCCCCTGAAA GAGGGGCTGG GCATTTGATA TGCCCAGCCC CTTGAGTAAG GGGCAAGAAA CCTAAGCCGC CTGGTGGATG GCTCGGCTCG GGGCGCTGAC GAAGGGCGTG
23SBHB_268 CGGGGA TGTAAGCGGG AAGGGAAACC GACCCGTTTA GTCTGCCGCT CCCAACAGCC CGAGTTTCTT GCCCCTGAAA GAGGGGCTGG GCATTTGATA TGCCCAGCCC CTTGAGTAAG GGGCAAGAAA CCTAAGCCGC CTGGTGGATG GCTCGGCTCG GGGCGCTGAC GAAGGGCG
23SBHB_860 GA CGGGTTTGAT GGGGCGGGGA TGTAAGCGGG AAGGGAAACC GACCCGTTTA GTCTGCCGCT CCCAACAGCC CGAGTTTCTT GCCCCTGAAA GAGGGGCTGG GCATTTGATA TGCCCAGCCC CTTGAGTAAG GGGCAAGAAA CCTAAGCCGC CTGGTGGATG GCTCGGCTCG GG
23SBHB_163 GGGA TGTAAGCGGG AAGGGAAACC GACCCGTTTA GTCTGCCGCT CCCAACAGCC CGAGTTTCTT GCCCCTGAAA GAGGGGCTGG GCATTTGATA TGCCCAGCCC CTTGAGTAAG GGGCAAGAAA CCTAAGCCGC CTGGTGGATG GCTCGGCTCG GGGCGCTGAC GAAGGGCGTG
23SBHB_432 GCGGGGA TGTAAGCGGG AAGGGAAACC GACCCGTTTA GTCTGCCGCT CCCAACAGCC CGAGTTTCTT GCCCCTGAAA GAGGGGCTGG GCATTTGATA TGCCCAGCCC CTTGAGTAAG GGGCAAGAAA CCTAAGCCGC CTGGTGGATG GCTCGGCTCG GGGCGCTGAC GAAGGG
23SBHB_378 GGCGGGGA TGTAAGCGGG AAGGGAAACC GACCCGTTTA GTCTGCCGCT CCCAACAGCC CGAGTTTCTT GCCCCTGAAA GAGGGGCTGG GCATTTGATA TGCCCAGCCC CTTGAGTAAG GGGCAAGAAA CCTAAGCCGC CTGGTGGATG GCTCGGCTCG GGGCGCTGAC GAAGG
23SBHB_159 GGGCGGGGA TGTAAGCGGG AAGGGAAACC GACCCGTTTA GTCTGCCGCT CCCAACAGCC CGAGTTTCTT GCCCCTGAAA GAGGGGCTGG GCATTTGATA TGCCCAGCCC CTTGAGTAAG GGGCAAGAAA CCTAAGCCGC CTGGTGGATG GCTCGGCTCG GGGCGCTGAC GAAG
23SBHB_281 GCGGGGA TGTAAGCGGG AAGGGAAACC GACCCGTTTA GTCTGCCGCT CCCAACAGCC CGAGTTTCTT GCCCCTGAAA GAGGGGCTGG GCATTTGATA TGCCCAGCCC CTTGAGTAAG GGGCAAGAAA CCTAAGCCGC CTGGTGGATG GCTCGGCTCG GGGCGCTGAC GAAGGGC
23SBHB_920 CGGGGA TGTAAGCGGG AAGGGAAACC GACCCGTTTA GTCTGCCGCT CCCAACAGCC CGAGTTTCTT GCCCCTGAAA GAGGGGCTGG GCATTTGATA TGCCCAGCCC CTTGAGTAAG GGGCAAGAAA CCTAAGCCGC CTGGTGGATG GCTCGGCTCG GGGCGCTGAC GAAGGG
23SBHB_498 CGGGGA TGTAAGCGGG AAGGGAAACC GACCCGTTTA GTCTGCCGCT CCCAACAGCC CGAGTTTCTT GCCCCTGAAA GAGGGGCTGG GCATTTGATA TGCCCAGCCC CTTGAGTAAG GGGCAAGAAA CCTAAGCCGC CTGGTGGATG GCTCGGCTCG GGGCGCTGAC GAAGGGC
23SBHB_255 GGGGA TGTAAGCGGG AAGGGAAACC GACCCGTTTA GTCTGCCGCT CCCAACAGCC CGAGTTTCTT GCCCCTGAAA GAGGGGCTGG GCATTTGATA TGCCCAGCCC CTTGAGTAAG GGGCAAGAAA CCTAAGCCGC CTGGTGGATG GCTCGGCTCG GGGCGCTGAC GAAGGGCG
23SBHB_146 CGCAGAAGA CGGGTTTGAT GGGGCGGGGA TGTAAGCGGG AAGGGAAACC GACCCGTTTA GTCTGCCGCT CCCAACAGCC CGAGTTTCTT GCCCCTGAAA GAGGGGCTGG GCATTTGATA TGCCCAGCCC CTTGAGTAAG GGGCAAGAAA CCTAAGCCGC CTGGTGGATG GCTCG
23SBHB_290 CGGGTTTGAT GGGGCGGGGA TGTAAGCGGG AAGGGAAACC GACCCGTTTA GTCTGCCGCT CCCAACAGCC CGAGTTTCTT GCCCCTGAAA GAGGGGCTGG GCATTTGATA TGCCCAGCCC CTTGAGTAAG GGGCAAGAAA CCTAAGCCGC CTGGTGGATG GCTCGGCTCG GGG
23SBHB_378 GGCGGGGA TGTAAGCGGG AAGGGAAACC GACCCGTTTA GTCTGCCGCT CCCAACAGCC CGAGTTTCTT GCCCCTGAAA GAGGGGCTGG GCATTTGATA TGCCCAGCCC CTTGAGTAAG GGGCAAGAAA CCTAAGCCGC CTGGTGGATG GCTCGGCTCG GGGCGCTGAC GAAG
23SBHB_376 A CGGGTTTGAT GGGGCGGGGA TGTAAGCGGG AAGGGAAACC GACCCGTTTA GTCTGCCGCT CCCAACAGCC CGAGTTTCTT GCCCCTGAAA GAGGGGCTGG GCATTTGATA TGCCCAGCCC CTTGAGTAAG GGGCAAGAAA CCTAAGCCGC CTGGTGGATG GCTCGGCTCG GG
23SBHB_131 GA CGGGTTTGAT GGGGCGGGGA TGTAAGCGGG AAGGGAAACC GACCCGTTTA GTCTGCCGCT CCCAACAGCC CGAGTTTCTT GCCCCTGAAA GAGGGGCTGG GCATTTGATA TGCCCAGCCC CTTGAGTAAG GGGCAAGAAA CCTAAGCCGC CTGGTGGATG GCTCGGCTCG G
23SBHB_224 CGGGGA TGTAAGCGGG AAGGGAAACC GACCCGTTTA GTCTGCCGCT CCCAACAGCC CGAGTTTCTT GCCCCTGAAA GAGGGGCTGG GCATTTGATA TGCCCAGCCC CTTGAGTAAG GGGCAAGAAA CCTAAGCCGC CTGGTGGATG GCTCGGCTCG GGGCGCTGAC GAAGGGG
23SBHB_314 GCGGGGA TGTAAGCGGG AAGGGAAACC GACCCGTTTA GTCTGCCGCT CCCAACAGCC CGAGTTTCTT GCCCCTGAAA GAGGGGCTGG GCATTTGATA TGCCCAGCCC CTTGAGTAAG GGGCAAGAAA CCTAAGCCGC CTGGTGGATG GCTCGGCTCG GGGCGCTGAC GAAGG
23SBHB_598 CGGGGA TGTAAGCGGG AAGGGAAACC GACCCGTTTA GTCTGCCGCT CCCAACAGCC CGAGTTTCTT GCCCCTGAAA GAGGGGCTGG GCATTTGATA TGCCCAGCCC CTTGAGTAAG GGGCAAGAAA CCTAAGCCGC CTGGTGGATG GCTCGGCTCG GGGCGCTGAC GAAGG
23SBHB_235 CGGGTTTGAT GGGGCGGGGA TGTAAGCGGG AAGGGAAACC GACCCGTTTA GTCTGCCGCT CCCAACAGCC CGAGTTTCTT GCCCCTGAAA GAGGGGCTGG GCATTTGATA TGCCCAGCCC CTTGAGTAAG GGGCAAGAAA CCTAAGCCGC CTGGTGGATG GCTCGGCTCG GG
23SBHB_865 GGGGA TGTAAGCGGG AAGGGAAACC GACCCGTTTA GTCTGCCGCT CCCAACAGCC CGAGTTTCTT GCCCCTGAAA GAGGGGCTGG GCATTTGATA TGCCCAGCCC CTTGAGTAAG GGGCAAGAAA CCTAAGCCGC CTGGTGGATG GCTCGGCTCG GGGCGCTGAC GAAGGG
23SBHB_505 GGGGA TGTAAGCGGG AAGGGAAACC GACCCGTTTA GTCTGCCGCT CCCAACAGCC CGAGTTTCTT GCCCCTGAAA GAGGGGCTGG GCATTTGATA TGCCCAGCCC CTTGAGTAAG GGGCAAGAAA CCTAAGCCGC CTGGTGGATG GCTCGGCTCG GGGCGCTGAC GAAGGGC
23SBHB_364 GGGA TGTAAGCGGG AAGGGAAACC GACCCGTTTA GTCTGCCGCT CCCAACAGCC CGAGTTTCTT GCCCCTGAAA GAGGGGCTGG GCATTTGATA TGCCCAGCCC CTTGAGTAAG GGGCAAGAAA CCTAAGCCGC CTGGTGGATG GCTCGGCTCG GGGCGCTGAC GAAGGGCG
23SBHB_329 GCGGGGA TGTAAGCGGG AAGGGAAACC GACCCGTTTA GTCTGCCGCT CCCAACAGCC CGAGTTTCTT GCCCCTGAAA GAGGGGCTGG GCATTTGATA TGCCCAGCCC CTTGAGTAAG GGGCAAGAAA CCTAAGCCGC CTGGTGGATG GCTCGGCTCG GGGCGCTGAC GAAG
23SBHB_181 GGGTTTGAT GGGGCGGGGA TGTAAGCGGG AAGGGAAACC GACCCGTTTA GTCTGCCGCT CCCAACAGCC CGAGTTTCTT GCCCCTGAAA GAGGGGCTGG GCATTTGATA TGCCCAGCCC CTTGAGTAAG GGGCAAGAAA CCTAAGCCGC CTGGTGGATG GCTCGGCTCG GG
23SBHB_693 CGGGGA TGTAAGCGGG AAGGGAAACC GACCCGTTTA GTCTGCCGCT CCCAACAGCC CGAGTTTCTT GCCCCTGAAA GAGGGGCTGG GCATTTGATA TGCCCAGCCC CTTGAGTAAG GGGCAAGAAA CCTAAGCCGC CTGGTGGATG GCTCGGCTCG GGGCGCTGAC GAAG
23SBHB_106 GGGA TGTAAGCGGG AAGGGAAACC GACCCGTTTA GTCTGCCGCT CCCAACAGCC CGAGTTTCTT GCCCCTGAAA GAGGGGCTGG GCATTTGATA TGCCCAGCCC CTTGAGTAAG GGGCAAGAAA CCTAAGCCGC CTGGTGGATG GCTCGGCTCG GGGCGCTGAC GAAGGG
23SBHB_657 GGGGA TGTAAGCGGG AAGGGAAACC GACCCGTTTA GTCTGCCGCT CCCAACAGCC CGAGTTTCTT GCCCCTGAAA GAGGGGCTGG GCATTTGATA TGCCCAGCCC CTTGAGTAAG GGGCAAGAAA CCTAAGCCGC CTGGTGGATG GCTCGGCTCG GGGCGCTGAC GAAGG
23SBHB_639 GGGA TGTAAGCGGG AAGGGAAACC GACCCGTTTA GTCTGCCGCT CCCAACAGCC CGAGTTTCTT GCCCCTGAAA GAGGGGCTGG GCATTTGATA TGCCCAGCCC CTTGAGTAAG GGGCAAGAAA CCTAAGCCGC CTGGTGGATG GCTCGGCTCG GGGCGCTGAC GAAGGGC
23SBHB_172 GGA TGTAAGCGGG AAGGGAAACC GACCCGTTTA GTCTGCCGCT CCCAACAGCC CGAGTTTCTT GCCCCTGAAA GAGGGGCTGG GCATTTGATA TGCCCAGCCC CTTGAGTAAG GGGCAAGAAA CCTAAGCCGC CTGGTGGATG GCTCGGCTCG GGGCGCTGAC GAAGGGCG
23SBHB_335 CGGGTTTGAT GGGGCGGGGA TGTAAGCGGG AAGGGAAACC GACCCGTTTA GTCTGCCGCT CCCAACAGCC CGAGTTTCTT GCCCCTGAAA GAGGGGCTGG GCATTTGATA TGCCCAGCCC CTTGAGTAAG GGGCAAGAAA CCTAAGCCGC CTGGTGGATG GCTCGGCTCG G
23SBHB_253 CGGGGA TGTAAGCGGG AAGGGAAACC GACCCGTTTA GTCTGCCGCT CCCAACAGCC CGAGTTTCTT GCCCCTGAAA GAGGGGCTGG GCATTTGATA TGCCCAGCCC CTTGAGTAAG GGGCAAGAAA CCTAAGCCGC CTGGTGGATG GCTCGGCTCG GGGCGCTGAC GAA
23SBHB_269 GGA TGTAAGCGGG AAGGGAAACC GACCCGTTTA GTCTGCCGCT CCCAACAGCC CGAGTTTCTT GCCCCTGAAA GAGGGGCTGG GCATTTGATA TGCCCAGCCC CTTGAGTAAG GGGCAAGAAA CCTAAGCCGC CTGGTGGATG GCTCGGCTCG GGGCGCTGAC GAAGGGC
23SBHB_135 GTTTGAT GGGGCGGGGA TGTAAGCGGG AAGGGAAACC GACCCGTTTA GTCTGCCGCT CCCAACAGCC CGAGTTTCTT GCCCCTGAAA GAGGGGCTGG GCATTTGATA TGCCCAGCCC CTTGAGTAAG GGGCAAGAAA CCTAAGCCGC CTGGTGGATG GCTCGGCTCG GGG
23SBHB_815 GGGA TGTAAGCGGG AAGGGAAACC GACCCGTTTA GTCTGCCGCT CCCAACAGCC CGAGTTTCTT GCCCCTGAAA GAGGGGCTGG GCATTTGATA TGCCCAGCCC CTTGAGTAAG GGGCAAGAAA CCTAAGCCGC CTGGTGGATG GCTCGGCTCG GGGCGCTGAC GAAGG
23SBHB_653 GGGGA TGTAAGCGGG AAGGGAAACC GACCCGTTTA GTCTGCCGCT CCCAACAGCC CGAGTTTCTT GCCCCTGAAA GAGGGGCTGG GCATTTGATA TGCCCAGCCC CTTGAGTAAG GGGCAAGAAA CCTAAGCCGC CTGGTGGATG GCTCGGCTCG GGGCGCTGAC GAAG
23SBHB_564 CGGG AAGGGAAACC GACCCGTTTA GTCTGCCGCT CCCAACAGCC CGAGTTTCTT GCCCCTGAAA GAGGGGCTGG GCATTTGATA TGCCCAGCCC CTTGAGTAAG GGGCAAGAAA CCTAAGCCGC CTGGTGGATG GCTCGGCTCG GGGCGCTGAC GAAGGGCGTG GCAAGCTGCG
23SBHB_289 CGGG AAGGGAAACC GACCCGTTTA GTCTGCCGCT CCCAACAGCC CGAGTTTCTT GCCCCTGAAA GAGGGGCTGG GCATTTGATA TGCCCAGCCC CTTGAGTAAG GGGCAAGAAA CCTAAGCCGC CTGGTGGATG GCTCGGCTCG GGGCGCTGAC GAAGGGCGTG GCAAGCTGCG AT
23SBHB_129 CGGG AAGGGAAACC GACCCGTTTA GTCTGCCGCT CCCAACAGCC CGAGTTTCTT GCCCCTGAAA GAGGGGCTGG GCATTTGATA TGCCCAGCCC CTTGAGTAAG GGGCAAGAAA CCTAAGCCGC CTGGTGGATG GCTCGGCTCG GGGCGCTGAC GAAGGGCGTG GCAAGCTG
23SBHB_146 GGTTTGAT GGGGCGGGGA TGTAAGCGGG AAGGGAAACC GACCCGTTTA GTCTGCCGCT CCCAACAGCC CGAGTTTCTT GCCCCTGAAA GAGGGGCTGG GCATTTGATA TGCCCAGCCC CTTGAGTAAG GGGCAAGAAA CCTAAGCCGC CTGGTGGATG GCTCGGCTCG GG
23SBHB_509 CGCAGAAGA CGGGTTTGAT GGGGCGGGGA TGTAAGCGGG AAGGGAAACC GACCCGTTTA GTCTGCCGCT CCCAACAGCC CGAGTTTCTT GCCCCTGAAA GAGGGGCTGG GCATTTGATA TGCCCAGCCC CTTGAGTAAG GGGCAAGAAA CCTAAGCCGC CTGGTGGATG G
23SBHB_426 CGCAGAAGA CGGGTTTGAT GGGGCGGGGA TGTAAGCGGG AAGGGAAACC GACCCGTTTA GTCTGCCGCT CCCAACAGCC CGAGTTTCTT GCCCCTGAAA GAGGGGCTGG GCATTTGATA TGCCCAGCCC CTTGAGTAAG GGGCAAGAAA CCTAAGCCGC CTGGTGGATG GC
23SBHB_211 C CCGCAGAAGA CGGGTTTGAT GGGGCGGGGA TGTAAGCGGG AAGGGAAACC GACCCGTTTA GTCTGCCGCT CCCAACAGCC CGAGTTTCTT GCCCCTGAAA GAGGGGCTGG GCATTTGATA TGCCCAGCCC CTTGAGTAAG GGGCAAGAAA CCTAAGCCGC CTGGTGGATG G
23SBHB_198 CCGCAGAAGA CGGGTTTGAT GGGGCGGGGA TGTAAGCGGG AAGGGAAACC GACCCGTTTA GTCTGCCGCT CCCAACAGCC CGAGTTTCTT GCCCCTGAAA GAGGGGCTGG GCATTTGATA TGCCCAGCCC CTTGAGTAAG GGGCAAGAAA CCTAAGCCGC CTGGTGGATG GC
23SBHB_155 C CCGCAGAAGA CGGGTTTGAT GGGGCGGGGA TGTAAGCGGG AAGGGAAACC GACCCGTTTA GTCTGCCGCT CCCAACAGCC CGAGTTTCTT GCCCCTGAAA GAGGGGCTGG GCATTTGATA TGCCCAGCCC CTTGAGTAAG GGGCAAGAAA CCTAAGCCGC CTGGTGGATG GC
23SBHB_404 C CCGCAGAAGA CGGGTTTGAT GGGGCGGGGA TGTAAGCGGG AAGGGAAACC GACCCGTTTA GTCTGCCGCT CCCAACAGCC CGAGTTTCTT GCCCCTGAAA GAGGGGCTGG GCATTTGATA TGCCCAGCCC CTTGAGTAAG GGGCAAGAAA CCTAAGCCGC CTGGTGGATG
23SBHB_235 CCGCAGAAGA CGGGTTTGAT GGGGCGGGGA TGTAAGCGGG AAGGGAAACC GACCCGTTTA GTCTGCCGCT CCCAACAGCC CGAGTTTCTT GCCCCTGAAA GAGGGGCTGG GCATTTGATA TGCCCAGCCC CTTGAGTAAG GGGCAAGAAA CCTAAGCCGC CTGGTGGATG G
23SBHB_450 CCGCAGAAGA CGGGTTTGAT GGGGCGGGGA TGTAAGCGGG AAGGGAAACC GACCCGTTTA GTCTGCCGCT CCCAACAGCC CGAGTTTCTT GCCCCTGAAA GAGGGGCTGG GCATTTGATA TGCCCAGCCC CTTGAGTAAG GGGCAAGAAA CCTAAGCCGC CTGGTGGATG
23SBHB_418 C CCGCAGAAGA CGGGTTTGAT GGGGCGGGGA TGTAAGCGGG AAGGGAAACC GACCCGTTTA GTCTGCCGCT CCCAACAGCC CGAGTTTCTT GCCCCTGAAA GAGGGGCTGG GCATTTGATA TGCCCAGCCC CTTGAGTAAG GGGCAAGAAA CCTAAGCCGC CTGGTGGAT
23SBHB_224 CGGGTTTGAT GGGGCGGGGA TGTAAGCGGG AAGGGAAACC GACCCGTTTA GTCTGCCGCT CCCAACAGCC CGAGTTTCTT GCCCCTGAAA GAGGGGCTGG GCATTTGATA TGCCCAGCCC CTTGAGTAAG GGGCAAGAAA CCTAAGCCGC CTGGTGGATG GCTCGGCTCG
23SBHB_126 GTTTGAT GGGGCGGGGA TGTAAGCGGG AAGGGAAACC GACCCGTTTA GTCTGCCGCT CCCAACAGCC CGAGTTTCTT GCCCCTGAAA GAGGGGCTGG GCATTTGATA TGCCCAGCCC CTTGAGTAAG GGGCAAGAAA CCTAAGCCGC CTGGTGGATG GCTCGGCTCG GG
23SBHB_887 GGGA TGTAAGCGGG AAGGGAAACC GACCCGTTTA GTCTGCCGCT CCCAACAGCC CGAGTTTCTT GCCCCTGAAA GAGGGGCTGG GCATTTGATA TGCCCAGCCC CTTGAGTAAG GGGCAAGAAA CCTAAGCCGC CTGGTGGATG GCTCGGCTCG GGGCGCTGAC GAAG
23SBHB_466 GGA TGTAAGCGGG AAGGGAAACC GACCCGTTTA GTCTGCCGCT CCCAACAGCC CGAGTTTCTT GCCCCTGAAA GAGGGGCTGG GCATTTGATA TGCCCAGCCC CTTGAGTAAG GGGCAAGAAA CCTAAGCCGC CTGGTGGATG GCTCGGCTCG GGGCGCTGAC GAAGGG
23SBHB_369 GGA TGTAAGCGGG AAGGGAAACC GACCCGTTTA GTCTGCCGCT CCCAACAGCC CGAGTTTCTT GCCCCTGAAA GAGGGGCTGG GCATTTGATA TGCCCAGCCC CTTGAGTAAG GGGCAAGAAA CCTAAGCCGC CTGGTGGATG GCTCGGCTCG GGGCGCTGAC GAAGG
23SBHB_204 CAGAAGA CGGGTTTGAT GGGGCGGGGA TGTAAGCGGG AAGGGAAACC GACCCGTTTA GTCTGCCGCT CCCAACAGCC CGAGTTTCTT GCCCCTGAAA GAGGGGCTGG GCATTTGATA TGCCCAGCCC CTTGAGTAAG GGGCAAGAAA CCTAAGCCGC CTGGTGGATG G
23SBHB_196 CAGAAGA CGGGTTTGAT GGGGCGGGGA TGTAAGCGGG AAGGGAAACC GACCCGTTTA GTCTGCCGCT CCCAACAGCC CGAGTTTCTT GCCCCTGAAA GAGGGGCTGG GCATTTGATA TGCCCAGCCC CTTGAGTAAG GGGCAAGAAA CCTAAGCCGC CTGGTGGATG GC
23SBHB_170 GTTTGAT GGGGCGGGGA TGTAAGCGGG AAGGGAAACC GACCCGTTTA GTCTGCCGCT CCCAACAGCC CGAGTTTCTT GCCCCTGAAA GAGGGGCTGG GCATTTGATA TGCCCAGCCC CTTGAGTAAG GGGCAAGAAA CCTAAGCCGC CTGGTGGATG GCTCGGCTCG G
23SBHB_108 CGCAGAAGA CGGGTTTGAT GGGGCGGGGA TGTAAGCGGG AAGGGAAACC GACCCGTTTA GTCTGCCGCT CCCAACAGCC CGAGTTTCTT GCCCCTGAAA GAGGGGCTGG GCATTTGATA TGCCCAGCCC CTTGAGTAAG GGGCAAGAAA CCTAAGCCGC CTGGTGGAT
```

```
23SBHB_101  CGCAGAAGA CGGGTTTGAT GGGGCGGGGA TGTAAGCGGG AAGGGAAACC GACCCGTTTA GTCTGCCGCT CCCAACAGCC CGAGTTTCTT GCCCCTGAAA GAGGGGCTGG GCATTTGATA TGCCCAGCCC CTTGAGTAAG GGGCAAGAAA CCTAAGCCGC CTGGTGGATG
23SBHB_450  CCGCAGAAGA CGGGTTTGAT GGGGCGGGGA TGTAAGCGGG AAGGGAAACC GACCCGTTTA GTCTGCCGCT CCCAACAGCC CGAGTTTCTT GCCCCTGAAA GAGGGGCTGG GCATTTGATA TGCCCAGCCC CTTGAGTAAG GGGCAAGAAA CCTAAGCCGC CTGGTGGAT
23SBHB_123  GA TGTAAGCGGG AAGGGAAACC GACCCGTTTA GTCTGCCGCT CCCAACAGCC CGAGTTTCTT GCCCCTGAAA GAGGGGCTGG GCATTTGATA TGCCCAGCCC CTTGAGTAAG GGGCAAGAAA CCTAAGCCGC CTGGTGGATG GCTCGGCTCG GGGCGCTGAC GAAGG
23SBHB_190  GTAAGCGGG AAGGGAAACC GACCCGTTTA GTCTGCCGCT CCCAACAGCC CGAGTTTCTT GCCCCTGAAA GAGGGGCTGG GCATTTGATA TGCCCAGCCC CTTGAGTAAG GGGCAAGAAA CCTAAGCCGC CTGGTGGATG GCTCGGCTCG GGGCGCTGAC GAAGGGCGTG G
23SBHB_117  GTAAGCGGG AAGGGAAACC GACCCGTTTA GTCTGCCGCT CCCAACAGCC CGAGTTTCTT GCCCCTGAAA GAGGGGCTGG GCATTTGATA TGCCCAGCCC CTTGAGTAAG GGGCAAGAAA CCTAAGCCGC CTGGTGGATG GCTCGGCTCG GGGCGCTGAC GAAGGGCGTG
23SBHB_462  GGA TGTAAGCGGG AAGGGAAACC GACCCGTTTA GTCTGCCGCT CCCAACAGCC CGAGTTTCTT GCCCCTGAAA GAGGGGCTGG GCATTTGATA TGCCCAGCCC CTTGAGTAAG GGGCAAGAAA CCTAAGCCGC CTGGTGGATG GCTCGGCTCG GGGCGCTGAC GAAG
23SBHB_328  CGGGGA TGTAAGCGGG AAGGGAAACC GACCCGTTTA GTCTGCCGCT CCCAACAGCC CGAGTTTCTT GCCCCTGAAA GAGGGGCTGG GCATTTGATA TGCCCAGCCC CTTGAGTAAG GGGCAAGAAA CCTAAGCCGC CTGGTGGATG GCTCGGCTCG GGGCGCTGAC G
23SBHB_165  GA TGTAAGCGGG AAGGGAAACC GACCCGTTTA GTCTGCCGCT CCCAACAGCC CGAGTTTCTT GCCCCTGAAA GAGGGGCTGG GCATTTGATA TGCCCAGCCC CTTGAGTAAG GGGCAAGAAA CCTAAGCCGC CTGGTGGATG GCTCGGCTCG GGGCGCTGAC GAAGGG
23SBHB_127  CGGGTTTGAT GGGGCGGGGA TGTAAGCGGG AAGGGAAACC GACCCGTTTA GTCTGCCGCT CCCAACAGCC CGAGTTTCTT GCCCCTGAAA GAGGGGCTGG GCATTTGATA TGCCCAGCCC CTTGAGTAAG GGGCAAGAAA CCTAAGCCGC CTGGTGGATG GCTCGGCT
23SBHB_170  A TGTAAGCGGG AAGGGAAACC GACCCGTTTA GTCTGCCGCT CCCAACAGCC CGAGTTTCTT GCCCCTGAAA GAGGGGCTGG GCATTTGATA TGCCCAGCCC CTTGAGTAAG GGGCAAGAAA CCTAAGCCGC CTGGTGGATG GCTCGGCTCG GGGCGCTGAC GAAGG
23SBHB_221  CGGGGA TGTAAGCGGG AAGGGAAACC GACCCGTTTA GTCTGCCGCT CCCAACAGCC CGAGTTTCTT GCCCCTGAAA GAGGGGCTGG GCATTTGATA TGCCCAGCCC CTTGAGTAAG GGGCAAGAAA CCTAAGCCGC CTGGTGGATG GCTCGGCTCG GGGCGCTGAC
23SBHB_188  GA TGTAAGCGGG AAGGGAAACC GACCCGTTTA GTCTGCCGCT CCCAACAGCC CGAGTTTCTT GCCCCTGAAA GAGGGGCTGG GCATTTGATA TGCCCAGCCC CTTGAGTAAG GGGCAAGAAA CCTAAGCCGC CTGGTGGATG GCTCGGCTCG GGGCGCTGAC GAAG
23SBHB_226  A TGTAAGCGGG AAGGGAAACC GACCCGTTTA GTCTGCCGCT CCCAACAGCC CGAGTTTCTT GCCCCTGAAA GAGGGGCTGG GCATTTGATA TGCCCAGCCC CTTGAGTAAG GGGCAAGAAA CCTAAGCCGC CTGGTGGATG GCTCGGCTCG GGGCGCTGAC GAAGGG
23SBHB_115  TGTAAGCGGG AAGGGAAACC GACCCGTTTA GTCTGCCGCT CCCAACAGCC CGAGTTTCTT GCCCCTGAAA GAGGGGCTGG GCATTTGATA TGCCCAGCCC CTTGAGTAAG GGGCAAGAAA CCTAAGCCGC CTGGTGGATG GCTCGGCTCG GGGCGCTGAC GAAGGG
23SBHB_274  GAT GGGGCGGGGA TGTAAGCGGG AAGGGAAACC GACCCGTTTA GTCTGCCGCT CCCAACAGCC CGAGTTTCTT GCCCCTGAAA GAGGGGCTGG GCATTTGATA TGCCCAGCCC CTTGAGTAAG GGGCAAGAAA CCTAAGCCGC CTGGTGGATG GCTCGGCTCG GG
23SBHB_258  A TGTAAGCGGG AAGGGAAACC GACCCGTTTA GTCTGCCGCT CCCAACAGCC CGAGTTTCTT GCCCCTGAAA GAGGGGCTGG GCATTTGATA TGCCCAGCCC CTTGAGTAAG GGGCAAGAAA CCTAAGCCGC CTGGTGGATG GCTCGGCTCG GGGCGCTGAC GAAG
23SBHB_424  CAGAAGA CGGGTTTGAT GGGGCGGGGA TGTAAGCGGG AAGGGAAACC GACCCGTTTA GTCTGCCGCT CCCAACAGCC CGAGTTTCTT GCCCCTGAAA GAGGGGCTGG GCATTTGATA TGCCCAGCCC CTTGAGTAAG GGGCAAGAAA CCTAAGCCGC CTGGTGGAT
23SBHB_126  GCAGAAGA CGGGTTTGAT GGGGCGGGGA TGTAAGCGGG AAGGGAAACC GACCCGTTTA GTCTGCCGCT CCCAACAGCC CGAGTTTCTT GCCCCTGAAA GAGGGGCTGG GCATTTGATA TGCCCAGCCC CTTGAGTAAG GGGCAAGAAA CCTAAGCCGC CTGGTGGAT
23SBHB_394  CAGAAGA CGGGTTTGAT GGGGCGGGGA TGTAAGCGGG AAGGGAAACC GACCCGTTTA GTCTGCCGCT CCCAACAGCC CGAGTTTCTT GCCCCTGAAA GAGGGGCTGG GCATTTGATA TGCCCAGCCC CTTGAGTAAG GGGCAAGAAA CCTAAGCCGC CTGGTGGATG
23SBHB_161  AGAAGA CGGGTTTGAT GGGGCGGGGA TGTAAGCGGG AAGGGAAACC GACCCGTTTA GTCTGCCGCT CCCAACAGCC CGAGTTTCTT GCCCCTGAAA GAGGGGCTGG GCATTTGATA TGCCCAGCCC CTTGAGTAAG GGGCAAGAAA CCTAAGCCGC CTGGTGGATG
23SBHB_419  GTAAGCGGG AAGGGAAACC GACCCGTTTA GTCTGCCGCT CCCAACAGCC CGAGTTTCTT GCCCCTGAAA GAGGGGCTGG GCATTTGATA TGCCCAGCCC CTTGAGTAAG GGGCAAGAAA CCTAAGCCGC CTGGTGGATG GCTCGGCTCG GGGCGCTGAC GAAGGGC
23SBHB_292  GTAAGCGGG AAGGGAAACC GACCCGTTTA GTCTGCCGCT CCCAACAGCC CGAGTTTCTT GCCCCTGAAA GAGGGGCTGG GCATTTGATA TGCCCAGCCC CTTGAGTAAG GGGCAAGAAA CCTAAGCCGC CTGGTGGATG GCTCGGCTCG GGGCGCTGAC GAAGGGCG
23SBHB_404  CGCAGAAGA CGGGTTTGAT GGGGCGGGGA TGTAAGCGGG AAGGGAAACC GACCCGTTTA GTCTGCCGCT CCCAACAGCC CGAGTTTCTT GCCCCTGAAA GAGGGGCTGG GCATTTGATA TGCCCAGCCC CTTGAGTAAG GGGCAAGAAA CCTAAGCCGC CTGGTG
23SBHB_176  CCGCAGAAGA CGGGTTTGAT GGGGCGGGGA TGTAAGCGGG AAGGGAAACC GACCCGTTTA GTCTGCCGCT CCCAACAGCC CGAGTTTCTT GCCCCTGAAA GAGGGGCTGG GCATTTGATA TGCCCAGCCC CTTGAGTAAG GGGCAAGAAA CCTAAGCCGC CTGGTG
23SBHB_159  C CCGCAGAAGA CGGGTTTGAT GGGGCGGGGA TGTAAGCGGG AAGGGAAACC GACCCGTTTA GTCTGCCGCT CCCAACAGCC CGAGTTTCTT GCCCCTGAAA GAGGGGCTGG GCATTTGATA TGCCCAGCCC CTTGAGTAAG GGGCAAGAAA CCTAAGCCGC CTGGTG
23SBHB_124  C CCGCAGAAGA CGGGTTTGAT GGGGCGGGGA TGTAAGCGGG AAGGGAAACC GACCCGTTTA GTCTGCCGCT CCCAACAGCC CGAGTTTCTT GCCCCTGAAA GAGGGGCTGG GCATTTGATA TGCCCAGCCC CTTGAGTAAG GGGCAAGAAA CCTAAGCCGC CTGGTGG
23SBHB_141  CCGCAGAAGA CGGGTTTGAT GGGGCGGGGA TGTAAGCGGG AAGGGAAACC GACCCGTTTA GTCTGCCGCT CCCAACAGCC CGAGTTTCTT GCCCCTGAAA GAGGGGCTGG GCATTTGATA TGCCCAGCCC CTTGAGTAAG GGGCAAGAAA CCTAAGCCGC CTGGTGG
23SBHB_353  CGCAGAAGA CGGGTTTGAT GGGGCGGGGA TGTAAGCGGG AAGGGAAACC GACCCGTTTA GTCTGCCGCT CCCAACAGCC CGAGTTTCTT GCCCCTGAAA GAGGGGCTGG GCATTTGATA TGCCCAGCCC CTTGAGTAAG GGGCAAGAAA CCTAAGCCGC CTGGTGG
23SBHB_824  GTAAGCGGG AAGGGAAACC GACCCGTTTA GTCTGCCGCT CCCAACAGCC CGAGTTTCTT GCCCCTGAAA GAGGGGCTGG GCATTTGATA TGCCCAGCCC CTTGAGTAAG GGGCAAGAAA CCTAAGCCGC CTGGTGGATG GCTCGGCTCG GGGCGCTGAC GAAGGG
23SBHB_675  GTAAGCGGG AAGGGAAACC GACCCGTTTA GTCTGCCGCT CCCAACAGCC CGAGTTTCTT GCCCCTGAAA GAGGGGCTGG GCATTTGATA TGCCCAGCCC CTTGAGTAAG GGGCAAGAAA CCTAAGCCGC CTGGTGGATG GCTCGGCTCG GGGCGCTGAC GAAGG
23SBHB_153  AGAAGA CGGGTTTGAT GGGGCGGGGA TGTAAGCGGG AAGGGAAACC GACCCGTTTA GTCTGCCGCT CCCAACAGCC CGAGTTTCTT GCCCCTGAAA GAGGGGCTGG GCATTTGATA TGCCCAGCCC CTTGAGTAAG GGGCAAGAAA CCTAAGCCGC CTGGTGGAT
23SBHB_116  GGGA TGTAAGCGGG AAGGGAAACC GACCCGTTTA GTCTGCCGCT CCCAACAGCC CGAGTTTCTT GCCCCTGAAA GAGGGGCTGG GCATTTGATA TGCCCAGCCC CTTGAGTAAG GGGCAAGAAA CCTAAGCCGC CTGGTGGATG GCTCGGCTCG GGGCGCTGAC
23SBHB_222  AAGCGGG AAGGGAAACC GACCCGTTTA GTCTGCCGCT CCCAACAGCC CGAGTTTCTT GCCCCTGAAA GAGGGGCTGG GCATTTGATA TGCCCAGCCC CTTGAGTAAG GGGCAAGAAA CCTAAGCCGC CTGGTGGATG GCTCGGCTCG GGGCGCTGAC GAAGGGC
23SBHB_130  AAGCGGG AAGGGAAACC GACCCGTTTA GTCTGCCGCT CCCAACAGCC CGAGTTTCTT GCCCCTGAAA GAGGGGCTGG GCATTTGATA TGCCCAGCCC CTTGAGTAAG GGGCAAGAAA CCTAAGCCGC CTGGTGGATG GCTCGGCTCG GGGCGCTGAC GAAGGGCG
23SBHB_132  TAAGCGGG AAGGGAAACC GACCCGTTTA GTCTGCCGCT CCCAACAGCC CGAGTTTCTT GCCCCTGAAA GAGGGGCTGG GCATTTGATA TGCCCAGCCC CTTGAGTAAG GGGCAAGAAA CCTAAGCCGC CTGGTGGATG GCTCGGCTCG GGGCGCTGAC GAAGG
23SBHB_737  GTAAGCGGG AAGGGAAACC GACCCGTTTA GTCTGCCGCT CCCAACAGCC CGAGTTTCTT GCCCCTGAAA GAGGGGCTGG GCATTTGATA TGCCCAGCCC CTTGAGTAAG GGGCAAGAAA CCTAAGCCGC CTGGTGGATG GCTCGGCTCG GGGCGCTGAC GAAG
23SBHB_133  TAAGCGGG AAGGGAAACC GACCCGTTTA GTCTGCCGCT CCCAACAGCC CGAGTTTCTT GCCCCTGAAA GAGGGGCTGG GCATTTGATA TGCCCAGCCC CTTGAGTAAG GGGCAAGAAA CCTAAGCCGC CTGGTGGATG GCTCGGCTCG GGGCGCTGAC GAAGGG
23SBHB_300  T GGGGCGGGGA TGTAAGCGGG AAGGGAAACC GACCCGTTTA GTCTGCCGCT CCCAACAGCC CGAGTTTCTT GCCCCTGAAA GAGGGGCTGG GCATTTGATA TGCCCAGCCC CTTGAGTAAG GGGCAAGAAA CCTAAGCCGC CTGGTGGATG GCTCGGCTCG GG
23SBHB_153  TAAGCGGG AAGGGAAACC GACCCGTTTA GTCTGCCGCT CCCAACAGCC CGAGTTTCTT GCCCCTGAAA GAGGGGCTGG GCATTTGATA TGCCCAGCCC CTTGAGTAAG GGGCAAGAAA CCTAAGCCGC CTGGTGGATG GCTCGGCTCG GGGCGCTGAC GAAG
23SBHB_156  CAGAAGA CGGGTTTGAT GGGGCGGGGA TGTAAGCGGG AAGGGAAACC GACCCGTTTA GTCTGCCGCT CCCAACAGCC CGAGTTTCTT GCCCCTGAAA GAGGGGCTGG GCATTTGATA TGCCCAGCCC CTTGAGTAAG GGGCAAGAAA CCTAAGCCGC CTGGTG
23SBHB_132  CAGAAGA CGGGTTTGAT GGGGCGGGGA TGTAAGCGGG AAGGGAAACC GACCCGTTTA GTCTGCCGCT CCCAACAGCC CGAGTTTCTT GCCCCTGAAA GAGGGGCTGG GCATTTGATA TGCCCAGCCC CTTGAGTAAG GGGCAAGAAA CCTAAGCCGC CTGGTGG
23SBHB_416  GCAGAAGA CGGGTTTGAT GGGGCGGGGA TGTAAGCGGG AAGGGAAACC GACCCGTTTA GTCTGCCGCT CCCAACAGCC CGAGTTTCTT GCCCCTGAAA GAGGGGCTGG GCATTTGATA TGCCCAGCCC CTTGAGTAAG GGGCAAGAAA CCTAAGCCGC CTGGTG
23SBHB_745  GGGGCGGGGA TGTAAGCGGG AAGGGAAACC GACCCGTTTA GTCTGCCGCT CCCAACAGCC CGAGTTTCTT GCCCCTGAAA GAGGGGCTGG GCATTTGATA TGCCCAGCCC CTTGAGTAAG GGGCAAGAAA CCTAAGCCGC CTGGTGGATG GCTCGGCTCG GG
23SBHB_628  CGGG AAGGGAAACC GACCCGTTTA GTCTGCCGCT CCCAACAGCC CGAGTTTCTT GCCCCTGAAA GAGGGGCTGG GCATTTGATA TGCCCAGCCC CTTGAGTAAG GGGCAAGAAA CCTAAGCCGC CTGGTGGATG GCTCGGCTCG GGGCGCTGAC GAAGGGCGTG G
23SBHB_138  CGGG AAGGGAAACC GACCCGTTTA GTCTGCCGCT CCCAACAGCC CGAGTTTCTT GCCCCTGAAA GAGGGGCTGG GCATTTGATA TGCCCAGCCC CTTGAGTAAG GGGCAAGAAA CCTAAGCCGC CTGGTGGATG GCTCGGCTCG GGGCGCTGAC GAAGGGCGTG GC
23SBHB_115  GGG AAGGGAAACC GACCCGTTTA GTCTGCCGCT CCCAACAGCC CGAGTTTCTT GCCCCTGAAA GAGGGGCTGG GCATTTGATA TGCCCAGCCC CTTGAGTAAG GGGCAAGAAA CCTAAGCCGC CTGGTGGATG GCTCGGCTCG GGGCGCTGAC GAAGGGCGTG G
23SBHB_386  CGGG AAGGGAAACC GACCCGTTTA GTCTGCCGCT CCCAACAGCC CGAGTTTCTT GCCCCTGAAA GAGGGGCTGG GCATTTGATA TGCCCAGCCC CTTGAGTAAG GGGCAAGAAA CCTAAGCCGC CTGGTGGATG GCTCGGCTCG GGGCGCTGAC GAAGGGCGTG
23SBHB_606  GGGAAACC GACCCGTTTA GTCTGCCGCT CCCAACAGCC CGAGTTTCTT GCCCCTGAAA GAGGGGCTGG GCATTTGATA TGCCCAGCCC CTTGAGTAAG GGGCAAGAAA CCTAAGCCGC CTGGTGGATG GCTCGGCTCG GGGCGCTGAC GAAGGGCGTG GCAAGCTGCG AT
23SBHB_143  GGAAACC GACCCGTTTA GTCTGCCGCT CCCAACAGCC CGAGTTTCTT GCCCCTGAAA GAGGGGCTGG GCATTTGATA TGCCCAGCCC CTTGAGTAAG GGGCAAGAAA CCTAAGCCGC CTGGTGGATG GCTCGGCTCG GGGCGCTGAC GAAGGGCGTG GCAAGCTGCG AT
23SBHB_450  GGGAAACC GACCCGTTTA GTCTGCCGCT CCCAACAGCC CGAGTTTCTT GCCCCTGAAA GAGGGGCTGG GCATTTGATA TGCCCAGCCC CTTGAGTAAG GGGCAAGAAA CCTAAGCCGC CTGGTGGATG GCTCGGCTCG GGGCGCTGAC GAAGGGCGTG GCAAGCTGCG
23SBHB_279  AAGCGGG AAGGGAAACC GACCCGTTTA GTCTGCCGCT CCCAACAGCC CGAGTTTCTT GCCCCTGAAA GAGGGGCTGG GCATTTGATA TGCCCAGCCC CTTGAGTAAG GGGCAAGAAA CCTAAGCCGC CTGGTGGATG GCTCGGCTCG GGGCGCTGAC GAAGG
23SBHB_344  AAGCGGG AAGGGAAACC GACCCGTTTA GTCTGCCGCT CCCAACAGCC CGAGTTTCTT GCCCCTGAAA GAGGGGCTGG GCATTTGATA TGCCCAGCCC CTTGAGTAAG GGGCAAGAAA CCTAAGCCGC CTGGTGGATG GCTCGGCTCG GGGCGCTGAC GAAGGG
23SBHB_239  GA CGGGTTTGAT GGGGCGGGGA TGTAAGCGGG AAGGGAAACC GACCCGTTTA GTCTGCCGCT CCCAACAGCC CGAGTTTCTT GCCCCTGAAA GAGGGGCTGG GCATTTGATA TGCCCAGCCC CTTGAGTAAG GGGCAAGAAA CCTAAGCCGC CTGGTGGATG
23SBHB_144  AGCGGG AAGGGAAACC GACCCGTTTA GTCTGCCGCT CCCAACAGCC CGAGTTTCTT GCCCCTGAAA GAGGGGCTGG GCATTTGATA TGCCCAGCCC CTTGAGTAAG GGGCAAGAAA CCTAAGCCGC CTGGTGGATG GCTCGGCTCG GGGCGCTGAC GAAGG
23SBHB_190  CGGGTTTGAT GGGGCGGGGA TGTAAGCGGG AAGGGAAACC GACCCGTTTA GTCTGCCGCT CCCAACAGCC CGAGTTTCTT GCCCCTGAAA GAGGGGCTGG GCATTTGATA TGCCCAGCCC CTTGAGTAAG GGGCAAGAAA CCTAAGCCGC CTGGTGGATG GC
23SBHB_320  AAGCGGG AAGGGAAACC GACCCGTTTA GTCTGCCGCT CCCAACAGCC CGAGTTTCTT GCCCCTGAAA GAGGGGCTGG GCATTTGATA TGCCCAGCCC CTTGAGTAAG GGGCAAGAAA CCTAAGCCGC CTGGTGGATG GCTCGGCTCG GGGCGCTGAC GAAG
23SBHB_114  CGCAGAAGA CGGGTTTGAT GGGGCGGGGA TGTAAGCGGG AAGGGAAACC GACCCGTTTA GTCTGCCGCT CCCAACAGCC CGAGTTTCTT GCCCCTGAAA GAGGGGCTGG GCATTTGATA TGCCCAGCCC CTTGAGTAAG GGGCAAGAAA CCTAAGCCGC CTG
23SBHB_192  CGCAGAAGA CGGGTTTGAT GGGGCGGGGA TGTAAGCGGG AAGGGAAACC GACCCGTTTA GTCTGCCGCT CCCAACAGCC CGAGTTTCTT GCCCCTGAAA GAGGGGCTGG GCATTTGATA TGCCCAGCCC CTTGAGTAAG GGGCAAGAAA CCTAAGCCGC CTGG
23SBHB_530  CCGCAGAAGA CGGGTTTGAT GGGGCGGGGA TGTAAGCGGG AAGGGAAACC GACCCGTTTA GTCTGCCGCT CCCAACAGCC CGAGTTTCTT GCCCCTGAAA GAGGGGCTGG GCATTTGATA TGCCCAGCCC CTTGAGTAAG GGGCAAGAAA CCTAAGCCGC CTG
23SBHB_444  C CCGCAGAAGA CGGGTTTGAT GGGGCGGGGA TGTAAGCGGG AAGGGAAACC GACCCGTTTA GTCTGCCGCT CCCAACAGCC CGAGTTTCTT GCCCCTGAAA GAGGGGCTGG GCATTTGATA TGCCCAGCCC CTTGAGTAAG GGGCAAGAAA CCTAAGCCGC CTG
23SBHB_646  AGAAGA CGGGTTTGAT GGGGCGGGGA TGTAAGCGGG AAGGGAAACC GACCCGTTTA GTCTGCCGCT CCCAACAGCC CGAGTTTCTT GCCCCTGAAA GAGGGGCTGG GCATTTGATA TGCCCAGCCC CTTGAGTAAG GGGCAAGAAA CCTAAGCCGC CTGGTG
23SBHB_162  AGCGGG AAGGGAAACC GACCCGTTTA GTCTGCCGCT CCCAACAGCC CGAGTTTCTT GCCCCTGAAA GAGGGGCTGG GCATTTGATA TGCCCAGCCC CTTGAGTAAG GGGCAAGAAA CCTAAGCCGC CTGGTGGATG GCTCGGCTCG GGGCGCTGAC GAAGGG
23SBHB_371  GGGCGGGGA TGTAAGCGGG AAGGGAAACC GACCCGTTTA GTCTGCCGCT CCCAACAGCC CGAGTTTCTT GCCCCTGAAA GAGGGGCTGG GCATTTGATA TGCCCAGCCC CTTGAGTAAG GGGCAAGAAA CCTAAGCCGC CTGGTGGATG GCTCGGCTCG GG
23SBHB_182  CGGGTTTGAT GGGGCGGGGA TGTAAGCGGG AAGGGAAACC GACCCGTTTA GTCTGCCGCT CCCAACAGCC CGAGTTTCTT GCCCCTGAAA GAGGGGCTGG GCATTTGATA TGCCCAGCCC CTTGAGTAAG GGGCAAGAAA CCTAAGCCGC CTGGTGGATG G
23SBHB_120  GCGGG AAGGGAAACC GACCCGTTTA GTCTGCCGCT CCCAACAGCC CGAGTTTCTT GCCCCTGAAA GAGGGGCTGG GCATTTGATA TGCCCAGCCC CTTGAGTAAG GGGCAAGAAA CCTAAGCCGC CTGGTGGATG GCTCGGCTCG GGGCGCTGAC GAAGG
23SBHB_147  CGGG AAGGGAAACC GACCCGTTTA GTCTGCCGCT CCCAACAGCC CGAGTTTCTT GCCCCTGAAA GAGGGGCTGG GCATTTGATA TGCCCAGCCC CTTGAGTAAG GGGCAAGAAA CCTAAGCCGC CTGGTGGATG GCTCGGCTCG GGGCGCTGAC GAAGGGC
23SBHB_102  CGGG AAGGGAAACC GACCCGTTTA GTCTGCCGCT CCCAACAGCC CGAGTTTCTT GCCCCTGAAA GAGGGGCTGG GCATTTGATA TGCCCAGCCC CTTGAGTAAG GGGCAAGAAA CCTAAGCCGC CTGGTGGATG GCTCGGCTCG GGGCGCTGAC GAAGGGCG
23SBHB_165  GGG AAGGGAAACC GACCCGTTTA GTCTGCCGCT CCCAACAGCC CGAGTTTCTT GCCCCTGAAA GAGGGGCTGG GCATTTGATA TGCCCAGCCC CTTGAGTAAG GGGCAAGAAA CCTAAGCCGC CTGGTGGATG GCTCGGCTCG GGGCGCTGAC GAAGGGCG
23SBHB_115  GCGGG AAGGGAAACC GACCCGTTTA GTCTGCCGCT CCCAACAGCC CGAGTTTCTT GCCCCTGAAA GAGGGGCTGG GCATTTGATA TGCCCAGCCC CTTGAGTAAG GGGCAAGAAA CCTAAGCCGC CTGGTGGATG GCTCGGCTCG GGGCGCTGAC GAAGGGC
23SBHB_149  AGCGGG AAGGGAAACC GACCCGTTTA GTCTGCCGCT CCCAACAGCC CGAGTTTCTT GCCCCTGAAA GAGGGGCTGG GCATTTGATA TGCCCAGCCC CTTGAGTAAG GGGCAAGAAA CCTAAGCCGC CTGGTGGATG GCTCGGCTCG GGGCGCTGAC GAAG
23SBHB_132  GAAGA CGGGTTTGAT GGGGCGGGGA TGTAAGCGGG AAGGGAAACC GACCCGTTTA GTCTGCCGCT CCCAACAGCC CGAGTTTCTT GCCCCTGAAA GAGGGGCTGG GCATTTGATA TGCCCAGCCC CTTGAGTAAG GGGCAAGAAA CCTAAGCCGC CTGGTG
23SBHB_733  GGCGGGGA TGTAAGCGGG AAGGGAAACC GACCCGTTTA GTCTGCCGCT CCCAACAGCC CGAGTTTCTT GCCCCTGAAA GAGGGGCTGG GCATTTGATA TGCCCAGCCC CTTGAGTAAG GGGCAAGAAA CCTAAGCCGC CTGGTGGATG GCTCGGCTCG GG
23SBHB_209  GA CGGGTTTGAT GGGGCGGGGA TGTAAGCGGG AAGGGAAACC GACCCGTTTA GTCTGCCGCT CCCAACAGCC CGAGTTTCTT GCCCCTGAAA GAGGGGCTGG GCATTTGATA TGCCCAGCCC CTTGAGTAAG GGGCAAGAAA CCTAAGCCGC CTGGTGGAT
23SBHB_148  GCGGG AAGGGAAACC GACCCGTTTA GTCTGCCGCT CCCAACAGCC CGAGTTTCTT GCCCCTGAAA GAGGGGCTGG GCATTTGATA TGCCCAGCCC CTTGAGTAAG GGGCAAGAAA CCTAAGCCGC CTGGTGGATG GCTCGGCTCG GGGCGCTGAC GAAGGG
23SBHB_343  GCGGGGA TGTAAGCGGG AAGGGAAACC GACCCGTTTA GTCTGCCGCT CCCAACAGCC CGAGTTTCTT GCCCCTGAAA GAGGGGCTGG GCATTTGATA TGCCCAGCCC CTTGAGTAAG GGGCAAGAAA CCTAAGCCGC CTGGTGGATG GCTCGGCTCG GG
23SBHB_178  AAGA CGGGTTTGAT GGGGCGGGGA TGTAAGCGGG AAGGGAAACC GACCCGTTTA GTCTGCCGCT CCCAACAGCC CGAGTTTCTT GCCCCTGAAA GAGGGGCTGG GCATTTGATA TGCCCAGCCC CTTGAGTAAG GGGCAAGAAA CCTAAGCCGC CTGGTG
23SBHB_294  CGGG AAGGGAAACC GACCCGTTTA GTCTGCCGCT CCCAACAGCC CGAGTTTCTT GCCCCTGAAA GAGGGGCTGG GCATTTGATA TGCCCAGCCC CTTGAGTAAG GGGCAAGAAA CCTAAGCCGC CTGGTGGATG GCTCGGCTCG GGGCGCTGAC GAAGGG
23SBHB_211  CGGG AAGGGAAACC GACCCGTTTA GTCTGCCGCT CCCAACAGCC CGAGTTTCTT GCCCCTGAAA GAGGGGCTGG GCATTTGATA TGCCCAGCCC CTTGAGTAAG GGGCAAGAAA CCTAAGCCGC CTGGTGGATG GCTCGGCTCG GGGCGCTGAC GAAGG
23SBHB_380  CGGGTTTGAT GGGGCGGGGA TGTAAGCGGG AAGGGAAACC GACCCGTTTA GTCTGCCGCT CCCAACAGCC CGAGTTTCTT GCCCCTGAAA GAGGGGCTGG GCATTTGATA TGCCCAGCCC CTTGAGTAAG GGGCAAGAAA CCTAAGCCGC CTGGTGGATG
23SBHB_505  CAGAAGA CGGGTTTGAT GGGGCGGGGA TGTAAGCGGG AAGGGAAACC GACCCGTTTA GTCTGCCGCT CCCAACAGCC CGAGTTTCTT GCCCCTGAAA GAGGGGCTGG GCATTTGATA TGCCCAGCCC CTTGAGTAAG GGGCAAGAAA CCTAAGCCGC CTG
23SBHB_202  CGCAGAAGA CGGGTTTGAT GGGGCGGGGA TGTAAGCGGG AAGGGAAACC GACCCGTTTA GTCTGCCGCT CCCAACAGCC CGAGTTTCTT GCCCCTGAAA GAGGGGCTGG GCATTTGATA TGCCCAGCCC CTTGAGTAAG GGGCAAGAAA CCTAAGCCGC CT
23SBHB_117  GCAGAAGA CGGGTTTGAT GGGGCGGGGA TGTAAGCGGG AAGGGAAACC GACCCGTTTA GTCTGCCGCT CCCAACAGCC CGAGTTTCTT GCCCCTGAAA GAGGGGCTGG GCATTTGATA TGCCCAGCCC CTTGAGTAAG GGGCAAGAAA CCTAAGCCGC CTG
23SBHB_165  CGGGGA TGTAAGCGGG AAGGGAAACC GACCCGTTTA GTCTGCCGCT CCCAACAGCC CGAGTTTCTT GCCCCTGAAA GAGGGGCTGG GCATTTGATA TGCCCAGCCC CTTGAGTAAG GGGCAAGAAA CCTAAGCCGC CTGGTGGATG GCTCGGCTCG GGG
23SBHB_251  CGGG AAGGGAAACC GACCCGTTTA GTCTGCCGCT CCCAACAGCC CGAGTTTCTT GCCCCTGAAA GAGGGGCTGG GCATTTGATA TGCCCAGCCC CTTGAGTAAG GGGCAAGAAA CCTAAGCCGC CTGGTGGATG GCTCGGCTCG GGGCGCTGAC GAAG
23SBHB_353  CGGGTTTGAT GGGGCGGGGA TGTAAGCGGG AAGGGAAACC GACCCGTTTA GTCTGCCGCT CCCAACAGCC CGAGTTTCTT GCCCCTGAAA GAGGGGCTGG GCATTTGATA TGCCCAGCCC CTTGAGTAAG GGGCAAGAAA CCTAAGCCGC CTGGTGGAT
23SBHB_187  CGGGGA TGTAAGCGGG AAGGGAAACC GACCCGTTTA GTCTGCCGCT CCCAACAGCC CGAGTTTCTT GCCCCTGAAA GAGGGGCTGG GCATTTGATA TGCCCAGCCC CTTGAGTAAG GGGCAAGAAA CCTAAGCCGC CTGGTGGATG GCTCGGCTCG GG
23SBHB_336  GGG AAGGGAAACC GACCCGTTTA GTCTGCCGCT CCCAACAGCC CGAGTTTCTT GCCCCTGAAA GAGGGGCTGG GCATTTGATA TGCCCAGCCC CTTGAGTAAG GGGCAAGAAA CCTAAGCCGC CTGGTGGATG GCTCGGCTCG GGGCGCTGAC GAAGG
23SBHB_267  CGCAGAAGA CGGGTTTGAT GGGGCGGGGA TGTAAGCGGG AAGGGAAACC GACCCGTTTA GTCTGCCGCT CCCAACAGCC CGAGTTTCTT GCCCCTGAAA GAGGGGCTGG GCATTTGATA TGCCCAGCCC CTTGAGTAAG GGGCAAGAAA CCTAAGCCGC C
23SBHB_559  GGG AAGGGAAACC GACCCGTTTA GTCTGCCGCT CCCAACAGCC CGAGTTTCTT GCCCCTGAAA GAGGGGCTGG GCATTTGATA TGCCCAGCCC CTTGAGTAAG GGGCAAGAAA CCTAAGCCGC CTGGTGGATG GCTCGGCTCG GGGCGCTGAC GAAGGG
23SBHB_255  GGG AAGGGAAACC GACCCGTTTA GTCTGCCGCT CCCAACAGCC CGAGTTTCTT GCCCCTGAAA GAGGGGCTGG GCATTTGATA TGCCCAGCCC CTTGAGTAAG GGGCAAGAAA CCTAAGCCGC CTGGTGGATG GCTCGGCTCG GGGCGCTGAC GAAGGGC
23SBHB_154  GG AAGGGAAACC GACCCGTTTA GTCTGCCGCT CCCAACAGCC CGAGTTTCTT GCCCCTGAAA GAGGGGCTGG GCATTTGATA TGCCCAGCCC CTTGAGTAAG GGGCAAGAAA CCTAAGCCGC CTGGTGGATG GCTCGGCTCG GGGCGCTGAC GAAGGGC
23SBHB_137  GG AAGGGAAACC GACCCGTTTA GTCTGCCGCT CCCAACAGCC CGAGTTTCTT GCCCCTGAAA GAGGGGCTGG GCATTTGATA TGCCCAGCCC CTTGAGTAAG GGGCAAGAAA CCTAAGCCGC CTGGTGGATG GCTCGGCTCG GGGCGCTGAC GAAGGGCG
23SBHB_172  AGAAGA CGGGTTTGAT GGGGCGGGGA TGTAAGCGGG AAGGGAAACC GACCCGTTTA GTCTGCCGCT CCCAACAGCC CGAGTTTCTT GCCCCTGAAA GAGGGGCTGG GCATTTGATA TGCCCAGCCC CTTGAGTAAG GGGCAAGAAA CCTAAGCCGC CTG
23SBHB_356  AGA CGGGTTTGAT GGGGCGGGGA TGTAAGCGGG AAGGGAAACC GACCCGTTTA GTCTGCCGCT CCCAACAGCC CGAGTTTCTT GCCCCTGAAA GAGGGGCTGG GCATTTGATA TGCCCAGCCC CTTGAGTAAG GGGCAAGAAA CCTAAGCCGC CTGGTG
23SBHB_226  GG AAGGGAAACC GACCCGTTTA GTCTGCCGCT CCCAACAGCC CGAGTTTCTT GCCCCTGAAA GAGGGGCTGG GCATTTGATA TGCCCAGCCC CTTGAGTAAG GGGCAAGAAA CCTAAGCCGC CTGGTGGATG GCTCGGCTCG GGGCGCTGAC GAAGG
23SBHB_365  GGG AAGGGAAACC GACCCGTTTA GTCTGCCGCT CCCAACAGCC CGAGTTTCTT GCCCCTGAAA GAGGGGCTGG GCATTTGATA TGCCCAGCCC CTTGAGTAAG GGGCAAGAAA CCTAAGCCGC CTGGTGGATG GCTCGGCTCG GGGCGCTGAC GAAG
23SBHB_319  CC CGAGTTTCTT GCCCCTGAAA GAGGGGCTGG GCATTTGATA TGCCCAGCCC CTTGAGTAAG GGGCAAGAAA CCTAAGCCGC CTGGTGGATG GCTCGGCTCG GGGCGCTGAC GAAGGGCGTG GCAAGCTGCG ATAAGCCCCG GTGAGGCGCA GGCAGCCGTC GAGCCGGGGA TTCCGAATGG GGACCTCCCG CGGCTTTTGC CGCACTCCCG TCAGGGAGGG GGAACGCGGG GAATTGAAAC ATCTTAGTAC CCGCAGGAAA AGAAAG
23SBHB_129  CAACAGCC CGAGTTTCTT GCCCCTGAAA GAGGGGCTGG GCATTTGATA TGCCCAGCCC CTTGAGTAAG GGGCAAGAAA CCTAAGCCGC CTGGTGGATG GCTCGGCTCG GGGCGCTGAC GAAGGGCGTG GCAAGCTGCG ATAAGCCCCG GTGAGGCGCA GGCAGCCGTC GAGCCGGGGA TTCCGAATGG GGACCTCCCG CGGCTTTTGC CGCACTCCCG TCAGGGAGGG GGAACGCGGG GAATTGAAAC ATCTTAGTAC CCGCAGGAAA AGAAAG
23SBHB_128  CCCAACAGCC CGAGTTTCTT GCCCCTGAAA GAGGGGCTGG GCATTTGATA TGCCCAGCCC CTTGAGTAAG GGGCAAGAAA CCTAAGCCGC CTGGTGGATG GCTCGGCTCG GGGCGCTGAC GAAGGGCGTG GCAAGCTGCG ATAAGCCCCG GTGAGGCGCA GGCAGCCGTC GAGCCGGGGA TTCCGAATGG GGACCTCCCG CGGCTTTTGC CGCACTCCCG TCAGGGAGGG GGAACGCGGG GAATTGAAAC ATCTTAGTAC CCGCAGGAAA AGAAAG
23SBHB_922  CT CCCAACAGCC CGAGTTTCTT GCCCCTGAAA GAGGGGCTGG GCATTTGATA TGCCCAGCCC CTTGAGTAAG GGGCAAGAAA CCTAAGCCGC CTGGTGGATG GCTCGGCTCG GGGCGCTGAC GAAGGGCGTG GCAAGCTGCG ATAAGCCCCG GTGAGGCGCA GGCAGCCGTC GAGCCGGGGA TTCCGAATGG GGACCTCCCG CGGCTTTTGC CGCACTCCCG TCAGGGAGGG GGAACGCGGG GAATTGAAAC ATCTTAGTAC CCGCAGGAAA AGAAAG
23SBHB_567  CTGCCGCT CCCAACAGCC CGAGTTTCTT GCCCCTGAAA GAGGGGCTGG GCATTTGATA TGCCCAGCCC CTTGAGTAAG GGGCAAGAAA CCTAAGCCGC CTGGTGGATG GCTCGGCTCG GGGCGCTGAC GAAGGGCGTG GCAAGCTGCG ATAAGCCCCG GTGAGGCGCA GGCAGCCGTC GAGCCGGGGA TTCCGAATGG GGACCTCCCG CGGCTTTTGC CGCACTCCCG TCAGGGAGGG GGAACGCGGG GAATTGAAAC ATCTTAGTAC CCGCAGGAAA AGAAAG
23SBHB_295  GTCTGCCGCT CCCAACAGCC CGAGTTTCTT GCCCCTGAAA GAGGGGCTGG GCATTTGATA TGCCCAGCCC CTTGAGTAAG GGGCAAGAAA CCTAAGCCGC CTGGTGGATG GCTCGGCTCG GGGCGCTGAC GAAGGGCGTG GCAAGCTGCG ATAAGCCCCG GTGAGGCGCA GGCAGCCGTC GAGCCGGGGA TTCCGAATGG GGACCTCCCG CGGCTTTTGC CGCACTCCCG TCAGGGAGGG GGAACGCGGG GAATTGAAAC ATCTTAGTAC CCGCAGGAAA AGAAAG
23SBHB_173  TCTGCCGCT CCCAACAGCC CGAGTTTCTT GCCCCTGAAA GAGGGGCTGG GCATTTGATA TGCCCAGCCC CTTGAGTAAG GGGCAAGAAA CCTAAGCCGC CTGGTGGATG GCTCGGCTCG GGGCGCTGAC GAAGGGCGTG GCAAGCTGCG ATAAGCCCCG GTGAGGCGCA GGCAGCCGTC GAGCCGGGGA TTCCGAATGG GGACCTCCCG CGGCTTTTGC CGCACTCCCG TCAGGGAGGG GGAACGCGGG GAATTGAAAC ATCTTAGTAC CCGCAGGAAA AGAAAG
23SBHB_140  TGCCGCT CCCAACAGCC CGAGTTTCTT GCCCCTGAAA GAGGGGCTGG GCATTTGATA TGCCCAGCCC CTTGAGTAAG GGGCAAGAAA CCTAAGCCGC CTGGTGGATG GCTCGGCTCG GGGCGCTGAC GAAGGGCGTG GCAAGCTGCG ATAAGCCCCG GTGAGGCGCA GGCAGCCGTC GAGCCGGGGA TTCCGAATGG GGACCTCCCG CGGCTTTTGC CGCACTCCCG TCAGGGAGGG GGAACGCGGG GAATTGAAAC ATCTTAGTAC CCGCAGGAAA AGAAAG
23SBHB_122  GCCGCT CCCAACAGCC CGAGTTTCTT GCCCCTGAAA GAGGGGCTGG GCATTTGATA TGCCCAGCCC CTTGAGTAAG GGGCAAGAAA CCTAAGCCGC CTGGTGGATG GCTCGGCTCG GGGCGCTGAC GAAGGGCGTG GCAAGCTGCG ATAAGCCCCG GTGAGGCGCA GGCAGCCGTC GAGCCGGGGA TTCCGAATGG GGACCTCCCG CGGCTTTTGC CGCACTCCCG TCAGGGAGGG GGAACGCGGG GAATTGAAAC ATCTTAGTAC CCGCAGGAAA AGAAAG
23SBHB_317  CGCT CCCAACAGCC CGAGTTTCTT GCCCCTGAAA GAGGGGCTGG GCATTTGATA TGCCCAGCCC CTTGAGTAAG GGGCAAGAAA CCTAAGCCGC CTGGTGGATG GCTCGGCTCG GGGCGCTGAC GAAGGGCGTG GCAAGCTGCG ATAAGCCCCG GTGAGGCGCA GGCAGCCGTC GAGCCGGGGA TTCCGAATGG GGACCTCCCG CGGCTTTTGC CGCACTCCCG TCAGGGAGGG GGAACGCGGG GAATTGAAAC ATCTTAGTAC CCGCAGGAAA AGAAAG
23SBHB_157  CCCAACAGCC CGAGTTTCTT GCCCCTGAAA GAGGGGCTGG GCATTTGATA TGCCCAGCCC CTTGAGTAAG GGGCAAGAAA CCTAAGCCGC CTGGTGGATG GCTCGGCTCG GGGCGCTGAC GAAGGGCGTG GCAAGCTGCG ATAAGCCCCG GTGAGGCGCA GGCAGCCGTC GAGCCGGGGA TTCCGAATGG GGACCTCCCG CGGCTTTTGC CGCACTCCCG TCAGGGAGGG GGAACGCGGG GAATTGAAAC ATCTTAGTAC CCGCAGGAAA AGAAAGC
23SBHB_523  CCAACAGCC CGAGTTTCTT GCCCCTGAAA GAGGGGCTGG GCATTTGATA TGCCCAGCCC CTTGAGTAAG GGGCAAGAAA CCTAAGCCGC CTGGTGGATG GCTCGGCTCG GGGCGCTGAC GAAGGGCGTG GCAAGCTGCG ATAAGCCCCG GTGAGGCGCA GGCAGCCGTC GAGCCGGGGA TTCCGAATGG GGACCTCCCG CGGCTTTTGC CGCACTCCCG TCAGGGAGGG GGAACGCGGG GAATTGAAAC ATCTTAGTAC CCGCAGGAAA AGAAAG
23SBHB_157  CAACAGCC CGAGTTTCTT GCCCCTGAAA GAGGGGCTGG GCATTTGATA TGCCCAGCCC CTTGAGTAAG GGGCAAGAAA CCTAAGCCGC CTGGTGGATG GCTCGGCTCG GGGCGCTGAC GAAGGGCGTG GCAAGCTGCG ATAAGCCCCG GTGAGGCGCA GGCAGCCGTC GAGCCGGGGA TTCCGAATGG GGACCTCCCG CGGCTTTTGC CGCACTCCCG TCAGGGAGGG GGAACGCGGG GAATTGAAAC ATCTTAGTAC CCGCAGGAAA AGAAAGC
23SBHB_989  AACAGCC CGAGTTTCTT GCCCCTGAAA GAGGGGCTGG GCATTTGATA TGCCCAGCCC CTTGAGTAAG GGGCAAGAAA CCTAAGCCGC CTGGTGGATG GCTCGGCTCG GGGCGCTGAC GAAGGGCGTG GCAAGCTGCG ATAAGCCCCG GTGAGGCGCA GGCAGCCGTC GAGCCGGGGA TTCCGAATGG GGACCTCCCG CGGCTTTTGC CGCACTCCCG TCAGGGAGGG GGAACGCGGG GAATTGAAAC ATCTTAGTAC CCGCAGGAAA AGAAAG
23SBHB_472  CAGCC CGAGTTTCTT GCCCCTGAAA GAGGGGCTGG GCATTTGATA TGCCCAGCCC CTTGAGTAAG GGGCAAGAAA CCTAAGCCGC CTGGTGGATG GCTCGGCTCG GGGCGCTGAC GAAGGGCGTG GCAAGCTGCG ATAAGCCCCG GTGAGGCGCA GGCAGCCGTC GAGCCGGGGA TTCCGAATGG GGACCTCCCG CGGCTTTTGC CGCACTCCCG TCAGGGAGGG GGAACGCGGG GAATTGAAAC ATCTTAGTAC CCGCAGGAAA AGAAAG
23SBHB_200  AGCC CGAGTTTCTT GCCCCTGAAA GAGGGGCTGG GCATTTGATA TGCCCAGCCC CTTGAGTAAG GGGCAAGAAA CCTAAGCCGC CTGGTGGATG GCTCGGCTCG GGGCGCTGAC GAAGGGCGTG GCAAGCTGCG ATAAGCCCCG GTGAGGCGCA GGCAGCCGTC GAGCCGGGGA TTCCGAATGG GGACCTCCCG CGGCTTTTGC CGCACTCCCG TCAGGGAGGG GGAACGCGGG GAATTGAAAC ATCTTAGTAC CCGCAGGAAA AGAAAG
23SBHB_116  GCC CGAGTTTCTT GCCCCTGAAA GAGGGGCTGG GCATTTGATA TGCCCAGCCC CTTGAGTAAG GGGCAAGAAA CCTAAGCCGC CTGGTGGATG GCTCGGCTCG GGGCGCTGAC GAAGGGCGTG GCAAGCTGCG ATAAGCCCCG GTGAGGCGCA GGCAGCCGTC GAGCCGGGGA TTCCGAATGG GGACCTCCCG CGGCTTTTGC CGCACTCCCG TCAGGGAGGG GGAACGCGGG GAATTGAAAC ATCTTAGTAC CCGCAGGAAA AGAAAG
23SBHB_144  GCC CGAGTTTCTT GCCCCTGAAA GAGGGGCTGG GCATTTGATA TGCCCAGCCC CTTGAGTAAG GGGCAAGAAA CCTAAGCCGC CTGGTGGATG GCTCGGCTCG GGGCGCTGAC GAAGGGCGTG GCAAGCTGCG ATAAGCCCCG GTGAGGCGCA GGCAGCCGTC GAGCCGGGGA TTCCGAATGG GGACCTCCCG CGGCTTTTGC CGCACTCCCG TCAGGGAGGG GGAACGCGGG GAATTGAAAC ATCTTAGTAC CCGCAGGAAA AGAAAGC
23SBHB_344  CC CGAGTTTCTT GCCCCTGAAA GAGGGGCTGG GCATTTGATA TGCCCAGCCC CTTGAGTAAG GGGCAAGAAA CCTAAGCCGC CTGGTGGATG GCTCGGCTCG GGGCGCTGAC GAAGGGCGTG GCAAGCTGCG ATAAGCCCCG GTGAGGCGCA GGCAGCCGTC GAGCCGGGGA TTCCGAATGG GGACCTCCCG CGGCTTTTGC CGCACTCCCG TCAGGGAGGG GGAACGCGGG GAATTGAAAC ATCTTAGTAC CCGCAGGAAA AGAAAGC
23SBHB_202  C CGAGTTTCTT GCCCCTGAAA GAGGGGCTGG GCATTTGATA TGCCCAGCCC CTTGAGTAAG GGGCAAGAAA CCTAAGCCGC CTGGTGGATG GCTCGGCTCG GGGCGCTGAC GAAGGGCGTG GCAAGCTGCG ATAAGCCCCG GTGAGGCGCA GGCAGCCGTC GAGCCGGGGA TTCCGAATGG GGACCTCCCG CGGCTTTTGC CGCACTCCCG TCAGGGAGGG GGAACGCGGG GAATTGAAAC ATCTTAGTAC CCGCAGGAAA AGAAAG
23SBHB_208  C CGAGTTTCTT GCCCCTGAAA GAGGGGCTGG GCATTTGATA TGCCCAGCCC CTTGAGTAAG GGGCAAGAAA CCTAAGCCGC CTGGTGGATG GCTCGGCTCG GGGCGCTGAC GAAGGGCGTG GCAAGCTGCG ATAAGCCCCG GTGAGGCGCA GGCAGCCGTC GAGCCGGGGA TTCCGAATGG GGACCTCCCG CGGCTTTTGC CGCACTCCCG TCAGGGAGGG GGAACGCGGG GAATTGAAAC ATCTTAGTAC CCGCAGGAAA AGAAAGC
23SBHB_159  CGAGTTTCTT GCCCCTGAAA GAGGGGCTGG GCATTTGATA TGCCCAGCCC CTTGAGTAAG GGGCAAGAAA CCTAAGCCGC CTGGTGGATG GCTCGGCTCG GGGCGCTGAC GAAGGGCGTG GCAAGCTGCG ATAAGCCCCG GTGAGGCGCA GGCAGCCGTC GAGCCGGGGA TTCCGAATGG GGACCTCCCG CGGCTTTTGC CGCACTCCCG TCAGGGAGGG GGAACGCGGG GAATTGAAAC ATCTTAGTAC CCGCAGGAAA AGAAAG
23SBHB_195  CGAGTTTCTT GCCCCTGAAA GAGGGGCTGG GCATTTGATA TGCCCAGCCC CTTGAGTAAG GGGCAAGAAA CCTAAGCCGC CTGGTGGATG GCTCGGCTCG GGGCGCTGAC GAAGGGCGTG GCAAGCTGCG ATAAGCCCCG GTGAGGCGCA GGCAGCCGTC GAGCCGGGGA TTCCGAATGG GGACCTCCCG CGGCTTTTGC CGCACTCCCG TCAGGGAGGG GGAACGCGGG GAATTGAAAC ATCTTAGTAC CCGCAGGAAA AGAAAGC
23SBHB_130  GTTTCTT GCCCCTGAAA GAGGGGCTGG GCATTTGATA TGCCCAGCCC CTTGAGTAAG GGGCAAGAAA CCTAAGCCGC CTGGTGGATG GCTCGGCTCG GGGCGCTGAC GAAGGGCGTG GCAAGCTGCG ATAAGCCCCG GTGAGGCGCA GGCAGCCGTC GAGCCGGGGA TTCCGAATGG GGACCTCCCG CGGCTTTTGC CGCACTCCCG TCAGGGAGGG GGAACGCGGG GAATTGAAAC ATCTTAGTAC CCGCAGGAAA AGAAAG
23SBHB_150  GTTTCTT GCCCCTGAAA GAGGGGCTGG GCATTTGATA TGCCCAGCCC CTTGAGTAAG GGGCAAGAAA CCTAAGCCGC CTGGTGGATG GCTCGGCTCG GGGCGCTGAC GAAGGGCGTG GCAAGCTGCG ATAAGCCCCG GTGAGGCGCA GGCAGCCGTC GAGCCGGGGA TTCCGAATGG GGACCTCCCG CGGCTTTTGC CGCACTCCCG TCAGGGAGGG GGAACGCGGG GAATTGAAAC ATCTTAGTAC CCGCAGGAAA AGAAAGC
23SBHB_134  CTT GCCCCTGAAA GAGGGGCTGG GCATTTGATA TGCCCAGCCC CTTGAGTAAG GGGCAAGAAA CCTAAGCCGC CTGGTGGATG GCTCGGCTCG GGGCGCTGAC GAAGGGCGTG GCAAGCTGCG ATAAGCCCCG GTGAGGCGCA GGCAGCCGTC GAGCCGGGGA TTCCGAATGG GGACCTCCCG CGGCTTTTGC CGCACTCCCG TCAGGGAGGG GGAACGCGGG GAATTGAAAC ATCTTAGTAC CCGCAGGAAA AGAAAG
23SBHB_155  CTT GCCCCTGAAA GAGGGGCTGG GCATTTGATA TGCCCAGCCC CTTGAGTAAG GGGCAAGAAA CCTAAGCCGC CTGGTGGATG GCTCGGCTCG GGGCGCTGAC GAAGGGCGTG GCAAGCTGCG ATAAGCCCCG GTGAGGCGCA GGCAGCCGTC GAGCCGGGGA TTCCGAATGG GGACCTCCCG CGGCTTTTGC CGCACTCCCG TCAGGGAGGG GGAACGCGGG GAATTGAAAC ATCTTAGTAC CCGCAGGAAA AGAAAGC
23SBHB_121  T GCCCCTGAAA GAGGGGCTGG GCATTTGATA TGCCCAGCCC CTTGAGTAAG GGGCAAGAAA CCTAAGCCGC CTGGTGGATG GCTCGGCTCG GGGCGCTGAC GAAGGGCGTG GCAAGCTGCG ATAAGCCCCG GTGAGGCGCA GGCAGCCGTC GAGCCGGGGA TTCCGAATGG GGACCTCCCG CGGCTTTTGC CGCACTCCCG TCAGGGAGGG GGAACGCGGG GAATTGAAAC ATCTTAGTAC CCGCAGGAAA AGAAAG
23SBHB_287  CCCTGAAA GAGGGGCTGG GCATTTGATA TGCCCAGCCC CTTGAGTAAG GGGCAAGAAA CCTAAGCCGC CTGGTGGATG GCTCGGCTCG GGGCGCTGAC GAAGGGCGTG GCAAGCTGCG ATAAGCCCCG GTGAGGCGCA GGCAGCCGTC GAGCCGGGGA TTCCGAATGG GGACCTCCCG CGGCTTTTGC CGCACTCCCG TCAGGGAGGG GGAACGCGGG GAATTGAAAC ATCTTAGTAC CCGCAGGAAA AGAAAG
23SBHB_334  CCTGAAA GAGGGGCTGG GCATTTGATA TGCCCAGCCC CTTGAGTAAG GGGCAAGAAA CCTAAGCCGC CTGGTGGATG GCTCGGCTCG GGGCGCTGAC GAAGGGCGTG GCAAGCTGCG ATAAGCCCCG GTGAGGCGCA GGCAGCCGTC GAGCCGGGGA TTCCGAATGG GGACCTCCCG CGGCTTTTGC CGCACTCCCG TCAGGGAGGG GGAACGCGGG GAATTGAAAC ATCTTAGTAC CCGCAGGAAA AGAAAG
23SBHB_147  CCCAACAGCC CGAGTTTCTT GCCCCTGAAA GAGGGGCTGG GCATTTGATA TGCCCAGCCC CTTGAGTAAG GGGCAAGAAA CCTAAGCCGC CTGGTGGATG GCTCGGCTCG GGGCGCTGAC GAAGGGCGTG GCAAGCTGCG ATAAGCCCCG GTGAGGCGCA GGCAGCCGTC GAGCCGGGGA TTCCGAATGG GGACCTCCCG CGGCTTTTGC CGCACTCCCG TCAGGGAGGG GGAACGCGGG
23SBHB_133  CCCAACAGCC CGAGTTTCTT GCCCCTGAAA GAGGGGCTGG GCATTTGATA TGCCCAGCCC CTTGAGTAAG GGGCAAGAAA CCTAAGCCGC CTGGTGGATG GCTCGGCTCG GGGCGCTGAC GAAGGGCGTG GCAAGCTGCG ATAAGCCCCG GTGAGGCGCA GGCAGCCGTC GAGCCGGGGA TTCCGAATGG GGACCTCCCG CGGCTTTTGC CGCACTCCCG TCAGGGAGGG GGAACGCGGG G
23SBHB_131  CAACAGCC CGAGTTTCTT GCCCCTGAAA GAGGGGCTGG GCATTTGATA TGCCCAGCCC CTTGAGTAAG GGGCAAGAAA CCTAAGCCGC CTGGTGGATG GCTCGGCTCG GGGCGCTGAC GAAGGGCGTG GCAAGCTGCG ATAAGCCCCG GTGAGGCGCA GGCAGCCGTC GAGCCGGGGA TTCCGAATGG GGACCTCCCG CGGCTTTTGC CGCACTCCCG TCAGGGAGGG GGAACGCGGG G
23SBHB_120  CAACAGCC CGAGTTTCTT GCCCCTGAAA GAGGGGCTGG GCATTTGATA TGCCCAGCCC CTTGAGTAAG GGGCAAGAAA CCTAAGCCGC CTGGTGGATG GCTCGGCTCG GGGCGCTGAC GAAGGGCGTG GCAAGCTGCG ATAAGCCCCG GTGAGGCGCA GGCAGCCGTC GAGCCGGGGA TTCCGAATGG GGACCTCCCG CGGCTTTTGC CGCACTCCCG TCAGGGAGGG GGAACGCGGG
23SBHB_400  CC CGAGTTTCTT GCCCCTGAAA GAGGGGCTGG GCATTTGATA TGCCCAGCCC CTTGAGTAAG GGGCAAGAAA CCTAAGCCGC CTGGTGGATG GCTCGGCTCG GGGCGCTGAC GAAGGGCGTG GCAAGCTGCG ATAAGCCCCG GTGAGGCGCA GGCAGCCGTC GAGCCGGGGA TTCCGAATGG GGACCTCCCG CGGCTTTTGC CGCACTCCCG TCAGGGAGGG GGAACGCGGG G
23SBHB_129  GCC CGAGTTTCTT GCCCCTGAAA GAGGGGCTGG GCATTTGATA TGCCCAGCCC CTTGAGTAAG GGGCAAGAAA CCTAAGCCGC CTGGTGGATG GCTCGGCTCG GGGCGCTGAC GAAGGGCGTG GCAAGCTGCG ATAAGCCCCG GTGAGGCGCA GGCAGCCGTC GAGCCGGGGA TTCCGAATGG GGACCTCCCG CGGCTTTTGC CGCACTCCCG TCAGGGAGGG GGAACGCGGG G
23SBHB_366  CC CGAGTTTCTT GCCCCTGAAA GAGGGGCTGG GCATTTGATA TGCCCAGCCC CTTGAGTAAG GGGCAAGAAA CCTAAGCCGC CTGGTGGATG GCTCGGCTCG GGGCGCTGAC GAAGGGCGTG GCAAGCTGCG ATAAGCCCCG GTGAGGCGCA GGCAGCCGTC GAGCCGGGGA TTCCGAATGG GGACCTCCCG CGGCTTTTGC CGCACTCCCG TCAGGGAGGG GGAACGCGGG
23SBHB_260  C CGAGTTTCTT GCCCCTGAAA GAGGGGCTGG GCATTTGATA TGCCCAGCCC CTTGAGTAAG GGGCAAGAAA CCTAAGCCGC CTGGTGGATG GCTCGGCTCG GGGCGCTGAC GAAGGGCGTG GCAAGCTGCG ATAAGCCCCG GTGAGGCGCA GGCAGCCGTC GAGCCGGGGA TTCCGAATGG GGACCTCCCG CGGCTTTTGC CGCACTCCCG TCAGGGAGGG GGAACGCGGG G
23SBHB_137  GCC CGAGTTTCTT GCCCCTGAAA GAGGGGCTGG GCATTTGATA TGCCCAGCCC CTTGAGTAAG GGGCAAGAAA CCTAAGCCGC CTGGTGGATG GCTCGGCTCG GGGCGCTGAC GAAGGGCGTG GCAAGCTGCG ATAAGCCCCG GTGAGGCGCA GGCAGCCGTC GAGCCGGGGA TTCCGAATGG GGACCTCCCG CGGCTTTTGC CGCACTCCCG TCAGGGAGGG GGAACGCGGG
23SBHB_284  C CGAGTTTCTT GCCCCTGAAA GAGGGGCTGG GCATTTGATA TGCCCAGCCC CTTGAGTAAG GGGCAAGAAA CCTAAGCCGC CTGGTGGATG GCTCGGCTCG GGGCGCTGAC GAAGGGCGTG GCAAGCTGCG ATAAGCCCCG GTGAGGCGCA GGCAGCCGTC GAGCCGGGGA TTCCGAATGG GGACCTCCCG CGGCTTTTGC CGCACTCCCG TCAGGGAGGG GGAACGCGGG
23SBHB_256  CGAGTTTCTT GCCCCTGAAA GAGGGGCTGG GCATTTGATA TGCCCAGCCC CTTGAGTAAG GGGCAAGAAA CCTAAGCCGC CTGGTGGATG GCTCGGCTCG GGGCGCTGAC GAAGGGCGTG GCAAGCTGCG ATAAGCCCCG GTGAGGCGCA GGCAGCCGTC GAGCCGGGGA TTCCGAATGG GGACCTCCCG CGGCTTTTGC CGCACTCCCG TCAGGGAGGG GGAACGCGGG G
23SBHB_274  CGAGTTTCTT GCCCCTGAAA GAGGGGCTGG GCATTTGATA TGCCCAGCCC CTTGAGTAAG GGGCAAGAAA CCTAAGCCGC CTGGTGGATG GCTCGGCTCG GGGCGCTGAC GAAGGGCGTG GCAAGCTGCG ATAAGCCCCG GTGAGGCGCA GGCAGCCGTC GAGCCGGGGA TTCCGAATGG GGACCTCCCG CGGCTTTTGC CGCACTCCCG TCAGGGAGGG GGAACGCGGG
23SBHB_181  GTTTCTT GCCCCTGAAA GAGGGGCTGG GCATTTGATA TGCCCAGCCC CTTGAGTAAG GGGCAAGAAA CCTAAGCCGC CTGGTGGATG GCTCGGCTCG GGGCGCTGAC GAAGGGCGTG GCAAGCTGCG ATAAGCCCCG GTGAGGCGCA GGCAGCCGTC GAGCCGGGGA TTCCGAATGG GGACCTCCCG CGGCTTTTGC CGCACTCCCG TCAGGGAGGG GGAACGCGGG G
23SBHB_194  GTTTCTT GCCCCTGAAA GAGGGGCTGG GCATTTGATA TGCCCAGCCC CTTGAGTAAG GGGCAAGAAA CCTAAGCCGC CTGGTGGATG GCTCGGCTCG GGGCGCTGAC GAAGGGCGTG GCAAGCTGCG ATAAGCCCCG GTGAGGCGCA GGCAGCCGTC GAGCCGGGGA TTCCGAATGG GGACCTCCCG CGGCTTTTGC CGCACTCCCG TCAGGGAGGG GGAACGCGGG
23SBHB_126  CGGGGA TGTAAGCGGG AAGGGAAACC GACCCGTTTA GTCTGCCGCT CCCAACAGCC CGAGTTTCTT GCCCCTGAAA GAGGGGCTGG GCATTTGATA TGCCCAGCCC CTTGAGTAAG GGGCAAGAAA CCTAAGCCGC CTGGTGGATG GCTCGGCTCG GGGCGCTGAC GAAGGGCGTG GCAAGCTGCG ATAAGCCCCG GTGAGGCGCA GGCAGCCGTC GAGCCGGGGA TTCCGAATGG GGACCTCCCG CGGCTTTTGC CGCACTCCCG TCAGGGAGGG G
23SBHB_262  CGGGGA TGTAAGCGGG AAGGGAAACC GACCCGTTTA GTCTGCCGCT CCCAACAGCC CGAGTTTCTT GCCCCTGAAA GAGGGGCTGG GCATTTGATA TGCCCAGCCC CTTGAGTAAG GGGCAAGAAA CCTAAGCCGC CTGGTGGATG GCTCGGCTCG GGGCGCTGAC GAAGGGCGTG GCAAGCTGCG ATAAGCCCCG GTGAGGCGCA GGCAGCCGTC GAGCCGGGGA TTCCGAATGG GGACCTCCCG CGGCTTTTGC CGCACTCCCG TCAGGGAGGG GG
```

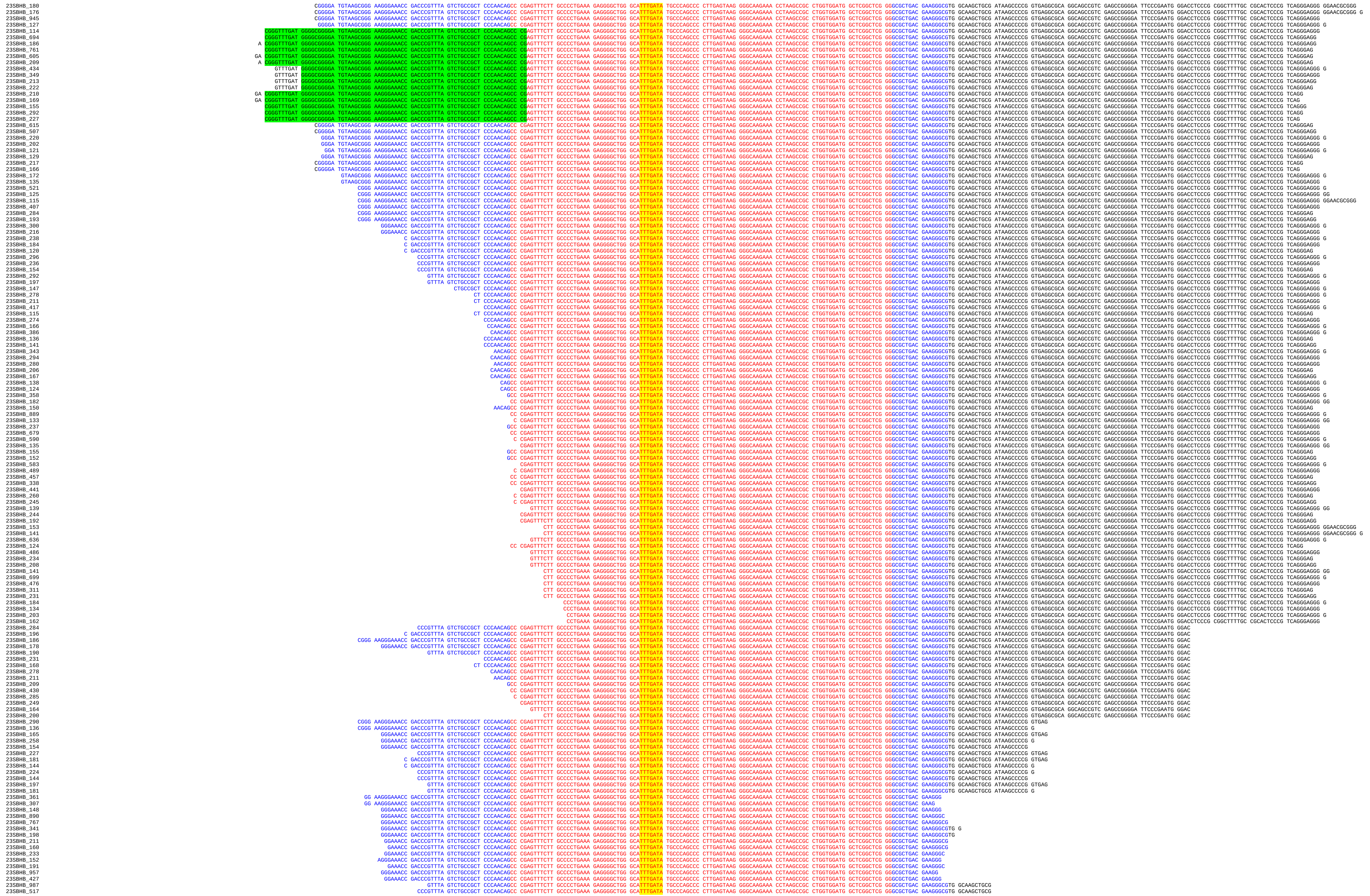

```
23SBHB_313 C GACCCGTTTA GTCTGCCGCT CCCAACAGCC CGAGTTTCTT GCCCCTGAAA GAGGGGCTGG GCATTTGATA TGCCCAGCCC CTTGAGTAAG GGGCAAGAAA CCTAAGCCGC CTGGTGGATG GCTCGGCTCG GGGCGCTGAC GAAGGGCGTG GCAAGCTGCG
23SBHB_185 C GACCCGTTTA GTCTGCCGCT CCCAACAGCC CGAGTTTCTT GCCCCTGAAA GAGGGGCTGG GCATTTGATA TGCCCAGCCC CTTGAGTAAG GGGCAAGAAA CCTAAGCCGC CTGGTGGATG GCTCGGCTCG GGGCGCTGAC GAAGGGCGTG GCAAGCTGCG AT
23SBHB_258 CCCGTTTA GTCTGCCGCT CCCAACAGCC CGAGTTTCTT GCCCCTGAAA GAGGGGCTGG GCATTTGATA TGCCCAGCCC CTTGAGTAAG GGGCAAGAAA CCTAAGCCGC CTGGTGGATG GCTCGGCTCG GGGCGCTGAC GAAGGGCGTG GCAAGCTGCG AT
23SBHB_249 CGTTTA GTCTGCCGCT CCCAACAGCC CGAGTTTCTT GCCCCTGAAA GAGGGGCTGG GCATTTGATA TGCCCAGCCC CTTGAGTAAG GGGCAAGAAA CCTAAGCCGC CTGGTGGATG GCTCGGCTCG GGGCGCTGAC GAAGGGCGTG GCAAGCTGCG
23SBHB_318 GTTTA GTCTGCCGCT CCCAACAGCC CGAGTTTCTT GCCCCTGAAA GAGGGGCTGG GCATTTGATA TGCCCAGCCC CTTGAGTAAG GGGCAAGAAA CCTAAGCCGC CTGGTGGATG GCTCGGCTCG GGGCGCTGAC GAAGGGCGTG GCAAGCTGCG AT
23SBHB_148 CCCGTTTA GTCTGCCGCT CCCAACAGCC CGAGTTTCTT GCCCCTGAAA GAGGGGCTGG GCATTTGATA TGCCCAGCCC CTTGAGTAAG GGGCAAGAAA CCTAAGCCGC CTGGTGGATG GCTCGGCTCG GGGCGCTGAC GAAGGGCGTG GCAAGCTG
23SBHB_155 GTTTA GTCTGCCGCT CCCAACAGCC CGAGTTTCTT GCCCCTGAAA GAGGGGCTGG GCATTTGATA TGCCCAGCCC CTTGAGTAAG GGGCAAGAAA CCTAAGCCGC CTGGTGGATG GCTCGGCTCG GGGCGCTGAC GAAGGGCGTG GCAAGCTGC
23SBHB_345 GTTTA GTCTGCCGCT CCCAACAGCC CGAGTTTCTT GCCCCTGAAA GAGGGGCTGG GCATTTGATA TGCCCAGCCC CTTGAGTAAG GGGCAAGAAA CCTAAGCCGC CTGGTGGATG GCTCGGCTCG GGGCGCTGAC GAAGGGCGTG GCAAGCTG
23SBHB_191 CCCGTTTA GTCTGCCGCT CCCAACAGCC CGAGTTTCTT GCCCCTGAAA GAGGGGCTGG GCATTTGATA TGCCCAGCCC CTTGAGTAAG GGGCAAGAAA CCTAAGCCGC CTGGTGGATG GCTCGGCTCG GGGCGCTGAC GAAGGGCGTG GCAAG
23SBHB_261 CGGG AAGGGAAACC GACCCGTTTA GTCTGCCGCT CCCAACAGCC CGAGTTTCTT GCCCCTGAAA GAGGGGCTGG GCATTTGATA TGCCCAGCCC CTTGAGTAAG GGGCAAGAAA CCTAAGCCGC CTGGTGGATG GCTCGGCTCG GGGCGCTGAC
23SBHB_155 CGGG AAGGGAAACC GACCCGTTTA GTCTGCCGCT CCCAACAGCC CGAGTTTCTT GCCCCTGAAA GAGGGGCTGG GCATTTGATA TGCCCAGCCC CTTGAGTAAG GGGCAAGAAA CCTAAGCCGC CTGGTGGATG GCTCGGCTCG GGGCGCTGAC G
23SBHB_106 GGGAAACC GACCCGTTTA GTCTGCCGCT CCCAACAGCC CGAGTTTCTT GCCCCTGAAA GAGGGGCTGG GCATTTGATA TGCCCAGCCC CTTGAGTAAG GGGCAAGAAA CCTAAGCCGC CTGGTGGATG GCTCGGCTCG GGGCGCTGAC GAAG
23SBHB_295 GGAAACC GACCCGTTTA GTCTGCCGCT CCCAACAGCC CGAGTTTCTT GCCCCTGAAA GAGGGGCTGG GCATTTGATA TGCCCAGCCC CTTGAGTAAG GGGCAAGAAA CCTAAGCCGC CTGGTGGATG GCTCGGCTCG GGGCGCTGAC GAAG
23SBHB_275 GGAAACC GACCCGTTTA GTCTGCCGCT CCCAACAGCC CGAGTTTCTT GCCCCTGAAA GAGGGGCTGG GCATTTGATA TGCCCAGCCC CTTGAGTAAG GGGCAAGAAA CCTAAGCCGC CTGGTGGATG GCTCGGCTCG GGGCGCTGAC GAAGG
23SBHB_188 GAAACC GACCCGTTTA GTCTGCCGCT CCCAACAGCC CGAGTTTCTT GCCCCTGAAA GAGGGGCTGG GCATTTGATA TGCCCAGCCC CTTGAGTAAG GGGCAAGAAA CCTAAGCCGC CTGGTGGATG GCTCGGCTCG GGGCGCTGAC GAAGG
23SBHB_301 GAAACC GACCCGTTTA GTCTGCCGCT CCCAACAGCC CGAGTTTCTT GCCCCTGAAA GAGGGGCTGG GCATTTGATA TGCCCAGCCC CTTGAGTAAG GGGCAAGAAA CCTAAGCCGC CTGGTGGATG GCTCGGCTCG GGGCGCTGAC GAAGGG
23SBHB_209 GAAACC GACCCGTTTA GTCTGCCGCT CCCAACAGCC CGAGTTTCTT GCCCCTGAAA GAGGGGCTGG GCATTTGATA TGCCCAGCCC CTTGAGTAAG GGGCAAGAAA CCTAAGCCGC CTGGTGGATG GCTCGGCTCG GGGCGCTGAC GAAG
23SBHB_167 AAACC GACCCGTTTA GTCTGCCGCT CCCAACAGCC CGAGTTTCTT GCCCCTGAAA GAGGGGCTGG GCATTTGATA TGCCCAGCCC CTTGAGTAAG GGGCAAGAAA CCTAAGCCGC CTGGTGGATG GCTCGGCTCG GGGCGCTGAC GAAGGG
23SBHB_512 GGGA TGTAAGCGGG AAGGGAAACC GACCCGTTTA GTCTGCCGCT CCCAACAGCC CGAGTTTCTT GCCCCTGAAA GAGGGGCTGG GCATTTGATA TGCCCAGCCC CTTGAGTAAG GGGCAAGAAA CCTAAGCCGC CTGGTGGATG GCTCGGCTCG GG
23SBHB_219 GGA TGTAAGCGGG AAGGGAAACC GACCCGTTTA GTCTGCCGCT CCCAACAGCC CGAGTTTCTT GCCCCTGAAA GAGGGGCTGG GCATTTGATA TGCCCAGCCC CTTGAGTAAG GGGCAAGAAA CCTAAGCCGC CTGGTGGATG GCTCGGCTCG GG
23SBHB_247 CGGGGA TGTAAGCGGG AAGGGAAACC GACCCGTTTA GTCTGCCGCT CCCAACAGCC CGAGTTTCTT GCCCCTGAAA GAGGGGCTGG GCATTTGATA TGCCCAGCCC CTTGAGTAAG GGGCAAGAAA CCTAAGCCGC CTGGTGGATG GCTCGGCTCG G
23SBHB_124 GGGGA TGTAAGCGGG AAGGGAAACC GACCCGTTTA GTCTGCCGCT CCCAACAGCC CGAGTTTCTT GCCCCTGAAA GAGGGGCTGG GCATTTGATA TGCCCAGCCC CTTGAGTAAG GGGCAAGAAA CCTAAGCCGC CTGGTGGATG GCTCGGCTCG GG
23SBHB_172 CGGGGA TGTAAGCGGG AAGGGAAACC GACCCGTTTA GTCTGCCGCT CCCAACAGCC CGAGTTTCTT GCCCCTGAAA GAGGGGCTGG GCATTTGATA TGCCCAGCCC CTTGAGTAAG GGGCAAGAAA CCTAAGCCGC CTGGTGGATG GCTCGGCTCG
23SBHB_338 GTAAGCGGG AAGGGAAACC GACCCGTTTA GTCTGCCGCT CCCAACAGCC CGAGTTTCTT GCCCCTGAAA GAGGGGCTGG GCATTTGATA TGCCCAGCCC CTTGAGTAAG GGGCAAGAAA CCTAAGCCGC CTGGTGGATG GCTCGGCTCG GG
23SBHB_512 C GACCCGTTTA GTCTGCCGCT CCCAACAGCC CGAGTTTCTT GCCCCTGAAA GAGGGGCTGG GCATTTGATA TGCCCAGCCC CTTGAGTAAG GGGCAAGAAA CCTAAGCCGC CTGGTGGATG GCTCGGCTCG GGGCGCTGAC GAAGGGCG
23SBHB_205 CC GACCCGTTTA GTCTGCCGCT CCCAACAGCC CGAGTTTCTT GCCCCTGAAA GAGGGGCTGG GCATTTGATA TGCCCAGCCC CTTGAGTAAG GGGCAAGAAA CCTAAGCCGC CTGGTGGATG GCTCGGCTCG GGGCGCTGAC GAAGGGC
23SBHB_200 AACC GACCCGTTTA GTCTGCCGCT CCCAACAGCC CGAGTTTCTT GCCCCTGAAA GAGGGGCTGG GCATTTGATA TGCCCAGCCC CTTGAGTAAG GGGCAAGAAA CCTAAGCCGC CTGGTGGATG GCTCGGCTCG GGGCGCTGAC GAAGGGCG
23SBHB_162 AACC GACCCGTTTA GTCTGCCGCT CCCAACAGCC CGAGTTTCTT GCCCCTGAAA GAGGGGCTGG GCATTTGATA TGCCCAGCCC CTTGAGTAAG GGGCAAGAAA CCTAAGCCGC CTGGTGGATG GCTCGGCTCG GGGCGCTGAC GAAGGGC
23SBHB_183 CC GACCCGTTTA GTCTGCCGCT CCCAACAGCC CGAGTTTCTT GCCCCTGAAA GAGGGGCTGG GCATTTGATA TGCCCAGCCC CTTGAGTAAG GGGCAAGAAA CCTAAGCCGC CTGGTGGATG GCTCGGCTCG GGGCGCTGAC GAAGGGCG
23SBHB_322 AACC GACCCGTTTA GTCTGCCGCT CCCAACAGCC CGAGTTTCTT GCCCCTGAAA GAGGGGCTGG GCATTTGATA TGCCCAGCCC CTTGAGTAAG GGGCAAGAAA CCTAAGCCGC CTGGTGGATG GCTCGGCTCG GGGCGCTGAC GAAGGG
23SBHB_201 AACC GACCCGTTTA GTCTGCCGCT CCCAACAGCC CGAGTTTCTT GCCCCTGAAA GAGGGGCTGG GCATTTGATA TGCCCAGCCC CTTGAGTAAG GGGCAAGAAA CCTAAGCCGC CTGGTGGATG GCTCGGCTCG GGGCGCTGAC GAAGG
23SBHB_335 CC GACCCGTTTA GTCTGCCGCT CCCAACAGCC CGAGTTTCTT GCCCCTGAAA GAGGGGCTGG GCATTTGATA TGCCCAGCCC CTTGAGTAAG GGGCAAGAAA CCTAAGCCGC CTGGTGGATG GCTCGGCTCG GGGCGCTGAC GAAGGG
23SBHB_450 C GACCCGTTTA GTCTGCCGCT CCCAACAGCC CGAGTTTCTT GCCCCTGAAA GAGGGGCTGG GCATTTGATA TGCCCAGCCC CTTGAGTAAG GGGCAAGAAA CCTAAGCCGC CTGGTGGATG GCTCGGCTCG GGGCGCTGAC GAAGGGC
23SBHB_649 CCCGTTTA GTCTGCCGCT CCCAACAGCC CGAGTTTCTT GCCCCTGAAA GAGGGGCTGG GCATTTGATA TGCCCAGCCC CTTGAGTAAG GGGCAAGAAA CCTAAGCCGC CTGGTGGATG GCTCGGCTCG GGGCGCTGAC GAAGGGCGTG G
23SBHB_389 C GACCCGTTTA GTCTGCCGCT CCCAACAGCC CGAGTTTCTT GCCCCTGAAA GAGGGGCTGG GCATTTGATA TGCCCAGCCC CTTGAGTAAG GGGCAAGAAA CCTAAGCCGC CTGGTGGATG GCTCGGCTCG GGGCGCTGAC GAAGGGCGTG G
23SBHB_233 C GACCCGTTTA GTCTGCCGCT CCCAACAGCC CGAGTTTCTT GCCCCTGAAA GAGGGGCTGG GCATTTGATA TGCCCAGCCC CTTGAGTAAG GGGCAAGAAA CCTAAGCCGC CTGGTGGATG GCTCGGCTCG GGGCGCTGAC GAAGGGCGTG
23SBHB_165 GACCCGTTTA GTCTGCCGCT CCCAACAGCC CGAGTTTCTT GCCCCTGAAA GAGGGGCTGG GCATTTGATA TGCCCAGCCC CTTGAGTAAG GGGCAAGAAA CCTAAGCCGC CTGGTGGATG GCTCGGCTCG GGGCGCTGAC GAAGGGCGTG G
23SBHB_135 CCCGTTTA GTCTGCCGCT CCCAACAGCC CGAGTTTCTT GCCCCTGAAA GAGGGGCTGG GCATTTGATA TGCCCAGCCC CTTGAGTAAG GGGCAAGAAA CCTAAGCCGC CTGGTGGATG GCTCGGCTCG GGGCGCTGAC GAAGGGCGTG GC
23SBHB_351 CCCGTTTA GTCTGCCGCT CCCAACAGCC CGAGTTTCTT GCCCCTGAAA GAGGGGCTGG GCATTTGATA TGCCCAGCCC CTTGAGTAAG GGGCAAGAAA CCTAAGCCGC CTGGTGGATG GCTCGGCTCG GGGCGCTGAC GAAGGGCGTG
23SBHB_165 GACCCGTTTA GTCTGCCGCT CCCAACAGCC CGAGTTTCTT GCCCCTGAAA GAGGGGCTGG GCATTTGATA TGCCCAGCCC CTTGAGTAAG GGGCAAGAAA CCTAAGCCGC CTGGTGGATG GCTCGGCTCG GGGCGCTGAC GAAGGGCG
23SBHB_214 AACC GACCCGTTTA GTCTGCCGCT CCCAACAGCC CGAGTTTCTT GCCCCTGAAA GAGGGGCTGG GCATTTGATA TGCCCAGCCC CTTGAGTAAG GGGCAAGAAA CCTAAGCCGC CTGGTGGATG GCTCGGCTCG GGGCGCTGAC GAAG
23SBHB_117 AAACC GACCCGTTTA GTCTGCCGCT CCCAACAGCC CGAGTTTCTT GCCCCTGAAA GAGGGGCTGG GCATTTGATA TGCCCAGCCC CTTGAGTAAG GGGCAAGAAA CCTAAGCCGC CTGGTGGATG GCTCGGCTCG GGGCGCTGAC GAAG
23SBHB_386 GTTTA GTCTGCCGCT CCCAACAGCC CGAGTTTCTT GCCCCTGAAA GAGGGGCTGG GCATTTGATA TGCCCAGCCC CTTGAGTAAG GGGCAAGAAA CCTAAGCCGC CTGGTGGATG GCTCGGCTCG GGGCGCTGAC GAAGGGCGTG GCAAG
23SBHB_139 GTTTA GTCTGCCGCT CCCAACAGCC CGAGTTTCTT GCCCCTGAAA GAGGGGCTGG GCATTTGATA TGCCCAGCCC CTTGAGTAAG GGGCAAGAAA CCTAAGCCGC CTGGTGGATG GCTCGGCTCG GGGCGCTGAC GAAGGGCGTG GCAAGC
23SBHB_168 GGGAAACC GACCCGTTTA GTCTGCCGCT CCCAACAGCC CGAGTTTCTT GCCCCTGAAA GAGGGGCTGG GCATTTGATA TGCCCAGCCC CTTGAGTAAG GGGCAAGAAA CCTAAGCCGC CTGGTGGATG GCTCGGCTCG GGGCGCTGAC
23SBHB_172 CGGGTTTGAT GGGGCGGGGA TGTAAGCGGG AAGGGAAACC GACCCGTTTA GTCTGCCGCT CCCAACAGCC CGAGTTTCTT GCCCCTGAAA GAGGGGCTGG GCATTTGATA TGCCCAGCCC CTTGAGTAAG GGGCAAGAAA CCTAAGCCGC CTGGTG
23SBHB_150 CGGGTTTGAT GGGGCGGGGA TGTAAGCGGG AAGGGAAACC GACCCGTTTA GTCTGCCGCT CCCAACAGCC CGAGTTTCTT GCCCCTGAAA GAGGGGCTGG GCATTTGATA TGCCCAGCCC CTTGAGTAAG GGGCAAGAAA CCTAAGCCGC CTGGTGG
23SBHB_977 GA CGGGTTTGAT GGGGCGGGGA TGTAAGCGGG AAGGGAAACC GACCCGTTTA GTCTGCCGCT CCCAACAGCC CGAGTTTCTT GCCCCTGAAA GAGGGGCTGG GCATTTGATA TGCCCAGCCC CTTGAGTAAG GGGCAAGAAA CCTAAGCCGC CTGGTG
23SBHB_405 A CGGGTTTGAT GGGGCGGGGA TGTAAGCGGG AAGGGAAACC GACCCGTTTA GTCTGCCGCT CCCAACAGCC CGAGTTTCTT GCCCCTGAAA GAGGGGCTGG GCATTTGATA TGCCCAGCCC CTTGAGTAAG GGGCAAGAAA CCTAAGCCGC CTGGTG
23SBHB_172 GGGTTTGAT GGGGCGGGGA TGTAAGCGGG AAGGGAAACC GACCCGTTTA GTCTGCCGCT CCCAACAGCC CGAGTTTCTT GCCCCTGAAA GAGGGGCTGG GCATTTGATA TGCCCAGCCC CTTGAGTAAG GGGCAAGAAA CCTAAGCCGC CTGGTG
23SBHB_381 GA CGGGTTTGAT GGGGCGGGGA TGTAAGCGGG AAGGGAAACC GACCCGTTTA GTCTGCCGCT CCCAACAGCC CGAGTTTCTT GCCCCTGAAA GAGGGGCTGG GCATTTGATA TGCCCAGCCC CTTGAGTAAG GGGCAAGAAA CCTAAGCCGC CTG
23SBHB_119 A CGGGTTTGAT GGGGCGGGGA TGTAAGCGGG AAGGGAAACC GACCCGTTTA GTCTGCCGCT CCCAACAGCC CGAGTTTCTT GCCCCTGAAA GAGGGGCTGG GCATTTGATA TGCCCAGCCC CTTGAGTAAG GGGCAAGAAA CCTAAGCCGC CTG
23SBHB_142 GTTTGAT GGGGCGGGGA TGTAAGCGGG AAGGGAAACC GACCCGTTTA GTCTGCCGCT CCCAACAGCC CGAGTTTCTT GCCCCTGAAA GAGGGGCTGG GCATTTGATA TGCCCAGCCC CTTGAGTAAG GGGCAAGAAA CCTAAGCCGC CTGGTGGAT
23SBHB_121 GTTTGAT GGGGCGGGGA TGTAAGCGGG AAGGGAAACC GACCCGTTTA GTCTGCCGCT CCCAACAGCC CGAGTTTCTT GCCCCTGAAA GAGGGGCTGG GCATTTGATA TGCCCAGCCC CTTGAGTAAG GGGCAAGAAA CCTAAGCCGC CTGGTGGATG
23SBHB_639 GTTTGAT GGGGCGGGGA TGTAAGCGGG AAGGGAAACC GACCCGTTTA GTCTGCCGCT CCCAACAGCC CGAGTTTCTT GCCCCTGAAA GAGGGGCTGG GCATTTGATA TGCCCAGCCC CTTGAGTAAG GGGCAAGAAA CCTAAGCCGC CTGGTG
23SBHB_776 CGGGTTTGAT GGGGCGGGGA TGTAAGCGGG AAGGGAAACC GACCCGTTTA GTCTGCCGCT CCCAACAGCC CGAGTTTCTT GCCCCTGAAA GAGGGGCTGG GCATTTGATA TGCCCAGCCC CTTGAGTAAG GGGCAAGAAA CCTAAGCCGC CTG
23SBHB_137 GA CGGGTTTGAT GGGGCGGGGA TGTAAGCGGG AAGGGAAACC GACCCGTTTA GTCTGCCGCT CCCAACAGCC CGAGTTTCTT GCCCCTGAAA GAGGGGCTGG GCATTTGATA TGCCCAGCCC CTTGAGTAAG GGGCAAGAAA CCTAAGCCGC C
23SBHB_134 CGGGTTTGAT GGGGCGGGGA TGTAAGCGGG AAGGGAAACC GACCCGTTTA GTCTGCCGCT CCCAACAGCC CGAGTTTCTT GCCCCTGAAA GAGGGGCTGG GCATTTGATA TGCCCAGCCC CTTGAGTAAG GGGCAAGAAA CCTAAGCCGC CT
23SBHB_302 GTTTGAT GGGGCGGGGA TGTAAGCGGG AAGGGAAACC GACCCGTTTA GTCTGCCGCT CCCAACAGCC CGAGTTTCTT GCCCCTGAAA GAGGGGCTGG GCATTTGATA TGCCCAGCCC CTTGAGTAAG GGGCAAGAAA CCTAAGCCGC CTG
23SBHB_300 CGGGTTTGAT GGGGCGGGGA TGTAAGCGGG AAGGGAAACC GACCCGTTTA GTCTGCCGCT CCCAACAGCC CGAGTTTCTT GCCCCTGAAA GAGGGGCTGG GCATTTGATA TGCCCAGCCC CTTGAGTAAG GGGCAAGAAA CCTAAGCCGC C
23SBHB_236 GGGGCGGGGA TGTAAGCGGG AAGGGAAACC GACCCGTTTA GTCTGCCGCT CCCAACAGCC CGAGTTTCTT GCCCCTGAAA GAGGGGCTGG GCATTTGATA TGCCCAGCCC CTTGAGTAAG GGGCAAGAAA CCTAAGCCGC CTGGTG
23SBHB_861 C GACCCGTTTA GTCTGCCGCT CCCAACAGCC CGAGTTTCTT GCCCCTGAAA GAGGGGCTGG GCATTTGATA TGCCCAGCCC CTTGAGTAAG GGGCAAGAAA CCTAAGCCGC CTGGTGGATG GCTCGGCTCG GGGCGCTGAC GAAGGG
23SBHB_262 GACCCGTTTA GTCTGCCGCT CCCAACAGCC CGAGTTTCTT GCCCCTGAAA GAGGGGCTGG GCATTTGATA TGCCCAGCCC CTTGAGTAAG GGGCAAGAAA CCTAAGCCGC CTGGTGGATG GCTCGGCTCG GGGCGCTGAC GAAGGG
23SBHB_464 C GACCCGTTTA GTCTGCCGCT CCCAACAGCC CGAGTTTCTT GCCCCTGAAA GAGGGGCTGG GCATTTGATA TGCCCAGCCC CTTGAGTAAG GGGCAAGAAA CCTAAGCCGC CTGGTGGATG GCTCGGCTCG GGGCGCTGAC GAAGG
23SBHB_223 CC GACCCGTTTA GTCTGCCGCT CCCAACAGCC CGAGTTTCTT GCCCCTGAAA GAGGGGCTGG GCATTTGATA TGCCCAGCCC CTTGAGTAAG GGGCAAGAAA CCTAAGCCGC CTGGTGGATG GCTCGGCTCG GGGCGCTGAC GAAGG
23SBHB_267 CC GACCCGTTTA GTCTGCCGCT CCCAACAGCC CGAGTTTCTT GCCCCTGAAA GAGGGGCTGG GCATTTGATA TGCCCAGCCC CTTGAGTAAG GGGCAAGAAA CCTAAGCCGC CTGGTGGATG GCTCGGCTCG GGGCGCTGAC GAAG
23SBHB_550 C GACCCGTTTA GTCTGCCGCT CCCAACAGCC CGAGTTTCTT GCCCCTGAAA GAGGGGCTGG GCATTTGATA TGCCCAGCCC CTTGAGTAAG GGGCAAGAAA CCTAAGCCGC CTGGTGGATG GCTCGGCTCG GGGCGCTGAC GAAG
23SBHB_145 GACCCGTTTA GTCTGCCGCT CCCAACAGCC CGAGTTTCTT GCCCCTGAAA GAGGGGCTGG GCATTTGATA TGCCCAGCCC CTTGAGTAAG GGGCAAGAAA CCTAAGCCGC CTGGTGGATG GCTCGGCTCG GGGCGCTGAC GAAGG
23SBHB_420 CTGCCGCT CCCAACAGCC CGAGTTTCTT GCCCCTGAAA GAGGGGCTGG GCATTTGATA TGCCCAGCCC CTTGAGTAAG GGGCAAGAAA CCTAAGCCGC CTGGTGGATG GCTCGGCTCG GGGCGCTGAC GAAGGGCGTG GCAAGCTGCG
23SBHB_293 GTCTGCCGCT CCCAACAGCC CGAGTTTCTT GCCCCTGAAA GAGGGGCTGG GCATTTGATA TGCCCAGCCC CTTGAGTAAG GGGCAAGAAA CCTAAGCCGC CTGGTGGATG GCTCGGCTCG GGGCGCTGAC GAAGGGCGTG GCAAGCTGCG
23SBHB_188 CTGCCGCT CCCAACAGCC CGAGTTTCTT GCCCCTGAAA GAGGGGCTGG GCATTTGATA TGCCCAGCCC CTTGAGTAAG GGGCAAGAAA CCTAAGCCGC CTGGTGGATG GCTCGGCTCG GGGCGCTGAC GAAGGGCGTG GCAAGCTGCG AT
23SBHB_115 GTCTGCCGCT CCCAACAGCC CGAGTTTCTT GCCCCTGAAA GAGGGGCTGG GCATTTGATA TGCCCAGCCC CTTGAGTAAG GGGCAAGAAA CCTAAGCCGC CTGGTGGATG GCTCGGCTCG GGGCGCTGAC GAAGGGCGTG GCAAGCTGCG AT
23SBHB_141 TCTGCCGCT CCCAACAGCC CGAGTTTCTT GCCCCTGAAA GAGGGGCTGG GCATTTGATA TGCCCAGCCC CTTGAGTAAG GGGCAAGAAA CCTAAGCCGC CTGGTGGATG GCTCGGCTCG GGGCGCTGAC GAAGGGCGTG GCAAGCTGCG
23SBHB_117 TGCCGCT CCCAACAGCC CGAGTTTCTT GCCCCTGAAA GAGGGGCTGG GCATTTGATA TGCCCAGCCC CTTGAGTAAG GGGCAAGAAA CCTAAGCCGC CTGGTGGATG GCTCGGCTCG GGGCGCTGAC GAAGGGCGTG GCAAGCTGCG
23SBHB_154 CTGCCGCT CCCAACAGCC CGAGTTTCTT GCCCCTGAAA GAGGGGCTGG GCATTTGATA TGCCCAGCCC CTTGAGTAAG GGGCAAGAAA CCTAAGCCGC CTGGTGGATG GCTCGGCTCG GGGCGCTGAC GAAGGGCGTG GCAAGCTG
23SBHB_152 GTCTGCCGCT CCCAACAGCC CGAGTTTCTT GCCCCTGAAA GAGGGGCTGG GCATTTGATA TGCCCAGCCC CTTGAGTAAG GGGCAAGAAA CCTAAGCCGC CTGGTGGATG GCTCGGCTCG GGGCGCTGAC GAAGGGCGTG GCAAG
23SBHB_879 CGGG AAGGGAAACC GACCCGTTTA GTCTGCCGCT CCCAACAGCC CGAGTTTCTT GCCCCTGAAA GAGGGGCTGG GCATTTGATA TGCCCAGCCC CTTGAGTAAG GGGCAAGAAA CCTAAGCCGC CTGGTGGATG GCTCGGCTCG GG
23SBHB_127 AAGCGGG AAGGGAAACC GACCCGTTTA GTCTGCCGCT CCCAACAGCC CGAGTTTCTT GCCCCTGAAA GAGGGGCTGG GCATTTGATA TGCCCAGCCC CTTGAGTAAG GGGCAAGAAA CCTAAGCCGC CTGGTGGATG GCTCGGCTCG GG
23SBHB_132 GGG AAGGGAAACC GACCCGTTTA GTCTGCCGCT CCCAACAGCC CGAGTTTCTT GCCCCTGAAA GAGGGGCTGG GCATTTGATA TGCCCAGCCC CTTGAGTAAG GGGCAAGAAA CCTAAGCCGC CTGGTGGATG GCTCGGCTCG GG
23SBHB_845 CCCGTTTA GTCTGCCGCT CCCAACAGCC CGAGTTTCTT GCCCCTGAAA GAGGGGCTGG GCATTTGATA TGCCCAGCCC CTTGAGTAAG GGGCAAGAAA CCTAAGCCGC CTGGTGGATG GCTCGGCTCG GGGCGCTGAC GAAGGG
23SBHB_570 CCCGTTTA GTCTGCCGCT CCCAACAGCC CGAGTTTCTT GCCCCTGAAA GAGGGGCTGG GCATTTGATA TGCCCAGCCC CTTGAGTAAG GGGCAAGAAA CCTAAGCCGC CTGGTGGATG GCTCGGCTCG GGGCGCTGAC GAAGG
23SBHB_460 CCCGTTTA GTCTGCCGCT CCCAACAGCC CGAGTTTCTT GCCCCTGAAA GAGGGGCTGG GCATTTGATA TGCCCAGCCC CTTGAGTAAG GGGCAAGAAA CCTAAGCCGC CTGGTGGATG GCTCGGCTCG GGGCGCTGAC GAAGGGCG
23SBHB_450 CCCGTTTA GTCTGCCGCT CCCAACAGCC CGAGTTTCTT GCCCCTGAAA GAGGGGCTGG GCATTTGATA TGCCCAGCCC CTTGAGTAAG GGGCAAGAAA CCTAAGCCGC CTGGTGGATG GCTCGGCTCG GGGCGCTGAC GAAGGGC
23SBHB_151 GACCCGTTTA GTCTGCCGCT CCCAACAGCC CGAGTTTCTT GCCCCTGAAA GAGGGGCTGG GCATTTGATA TGCCCAGCCC CTTGAGTAAG GGGCAAGAAA CCTAAGCCGC CTGGTGGATG GCTCGGCTCG GGGCGCTGAC GAAGGGC
23SBHB_385 GTTTA GTCTGCCGCT CCCAACAGCC CGAGTTTCTT GCCCCTGAAA GAGGGGCTGG GCATTTGATA TGCCCAGCCC CTTGAGTAAG GGGCAAGAAA CCTAAGCCGC CTGGTGGATG GCTCGGCTCG GGGCGCTGAC GAAGGGCGTG G
23SBHB_148 CGTTTA GTCTGCCGCT CCCAACAGCC CGAGTTTCTT GCCCCTGAAA GAGGGGCTGG GCATTTGATA TGCCCAGCCC CTTGAGTAAG GGGCAAGAAA CCTAAGCCGC CTGGTGGATG GCTCGGCTCG GGGCGCTGAC GAAGGGCGTG G
23SBHB_183 GTTTA GTCTGCCGCT CCCAACAGCC CGAGTTTCTT GCCCCTGAAA GAGGGGCTGG GCATTTGATA TGCCCAGCCC CTTGAGTAAG GGGCAAGAAA CCTAAGCCGC CTGGTGGATG GCTCGGCTCG GGGCGCTGAC GAAGGGCGTG
23SBHB_119 CGTTTA GTCTGCCGCT CCCAACAGCC CGAGTTTCTT GCCCCTGAAA GAGGGGCTGG GCATTTGATA TGCCCAGCCC CTTGAGTAAG GGGCAAGAAA CCTAAGCCGC CTGGTGGATG GCTCGGCTCG GGGCGCTGAC GAAGGGCG
23SBHB_331 GTTTA GTCTGCCGCT CCCAACAGCC CGAGTTTCTT GCCCCTGAAA GAGGGGCTGG GCATTTGATA TGCCCAGCCC CTTGAGTAAG GGGCAAGAAA CCTAAGCCGC CTGGTGGATG GCTCGGCTCG GGGCGCTGAC GAAGGGCG
23SBHB_145 CGTTTA GTCTGCCGCT CCCAACAGCC CGAGTTTCTT GCCCCTGAAA GAGGGGCTGG GCATTTGATA TGCCCAGCCC CTTGAGTAAG GGGCAAGAAA CCTAAGCCGC CTGGTGGATG GCTCGGCTCG GGGCGCTGAC GAAGGGC
23SBHB_511 CC CGAGTTTCTT GCCCCTGAAA GAGGGGCTGG GCATTTGATA TGCCCAGCCC CTTGAGTAAG GGGCAAGAAA CCTAAGCCGC CTGGTGGATG GCTCGGCTCG GGGCGCTGAC GAAGGGCGTG GCAAGCTGCG ATAAGCCCCG G
23SBHB_479 CC CGAGTTTCTT GCCCCTGAAA GAGGGGCTGG GCATTTGATA TGCCCAGCCC CTTGAGTAAG GGGCAAGAAA CCTAAGCCGC CTGGTGGATG GCTCGGCTCG GGGCGCTGAC GAAGGGCGTG GCAAGCTGCG ATAAGCCCCG GTGAG
23SBHB_175 GCC CGAGTTTCTT GCCCCTGAAA GAGGGGCTGG GCATTTGATA TGCCCAGCCC CTTGAGTAAG GGGCAAGAAA CCTAAGCCGC CTGGTGGATG GCTCGGCTCG GGGCGCTGAC GAAGGGCGTG GCAAGCTGCG ATAAGCCCCG GTGAG
23SBHB_149 CC CGAGTTTCTT GCCCCTGAAA GAGGGGCTGG GCATTTGATA TGCCCAGCCC CTTGAGTAAG GGGCAAGAAA CCTAAGCCGC CTGGTGGATG GCTCGGCTCG GGGCGCTGAC GAAGGGCGTG GCAAGCTGCG ATAAGCCCCG GTGAGGCGCA G
23SBHB_304 C CGAGTTTCTT GCCCCTGAAA GAGGGGCTGG GCATTTGATA TGCCCAGCCC CTTGAGTAAG GGGCAAGAAA CCTAAGCCGC CTGGTGGATG GCTCGGCTCG GGGCGCTGAC GAAGGGCGTG GCAAGCTGCG ATAAGCCCCG GTGAG
23SBHB_255 CCCAACAGCC CGAGTTTCTT GCCCCTGAAA GAGGGGCTGG GCATTTGATA TGCCCAGCCC CTTGAGTAAG GGGCAAGAAA CCTAAGCCGC CTGGTGGATG GCTCGGCTCG GGGCGCTGAC GAAGGGCGTG GCAAGCTGCG ATAAGCCCCG G
23SBHB_203 CT CCCAACAGCC CGAGTTTCTT GCCCCTGAAA GAGGGGCTGG GCATTTGATA TGCCCAGCCC CTTGAGTAAG GGGCAAGAAA CCTAAGCCGC CTGGTGGATG GCTCGGCTCG GGGCGCTGAC GAAGGGCGTG GCAAGCTGCG ATAAGCCCCG G
23SBHB_137 CTGCCGCT CCCAACAGCC CGAGTTTCTT GCCCCTGAAA GAGGGGCTGG GCATTTGATA TGCCCAGCCC CTTGAGTAAG GGGCAAGAAA CCTAAGCCGC CTGGTGGATG GCTCGGCTCG GGGCGCTGAC GAAGGGCGTG GCAAGCTGCG ATAAGCCCCG G
23SBHB_153 CT CCCAACAGCC CGAGTTTCTT GCCCCTGAAA GAGGGGCTGG GCATTTGATA TGCCCAGCCC CTTGAGTAAG GGGCAAGAAA CCTAAGCCGC CTGGTGGATG GCTCGGCTCG GGGCGCTGAC GAAGGGCGTG GCAAGCTGCG ATAAGCCCCG
23SBHB_163 CCCAACAGCC CGAGTTTCTT GCCCCTGAAA GAGGGGCTGG GCATTTGATA TGCCCAGCCC CTTGAGTAAG GGGCAAGAAA CCTAAGCCGC CTGGTGGATG GCTCGGCTCG GGGCGCTGAC GAAGGGCGTG GCAAGCTGCG ATAAGCCCCG
23SBHB_216 CCCAACAGCC CGAGTTTCTT GCCCCTGAAA GAGGGGCTGG GCATTTGATA TGCCCAGCCC CTTGAGTAAG GGGCAAGAAA CCTAAGCCGC CTGGTGGATG GCTCGGCTCG GGGCGCTGAC GAAGGGCGTG GCAAGCTGCG ATAAGCCCCG GTGAG
23SBHB_180 CT CCCAACAGCC CGAGTTTCTT GCCCCTGAAA GAGGGGCTGG GCATTTGATA TGCCCAGCCC CTTGAGTAAG GGGCAAGAAA CCTAAGCCGC CTGGTGGATG GCTCGGCTCG GGGCGCTGAC GAAGGGCGTG GCAAGCTGCG ATAAGCCCCG GTGAG
23SBHB_214 CAACAGCC CGAGTTTCTT GCCCCTGAAA GAGGGGCTGG GCATTTGATA TGCCCAGCCC CTTGAGTAAG GGGCAAGAAA CCTAAGCCGC CTGGTGGATG GCTCGGCTCG GGGCGCTGAC GAAGGGCGTG GCAAGCTGCG ATAAGCCCCG GTGAG
23SBHB_184 AACAGCC CGAGTTTCTT GCCCCTGAAA GAGGGGCTGG GCATTTGATA TGCCCAGCCC CTTGAGTAAG GGGCAAGAAA CCTAAGCCGC CTGGTGGATG GCTCGGCTCG GGGCGCTGAC GAAGGGCGTG GCAAGCTGCG ATAAGCCCCG GTGAG
23SBHB_225 CAACAGCC CGAGTTTCTT GCCCCTGAAA GAGGGGCTGG GCATTTGATA TGCCCAGCCC CTTGAGTAAG GGGCAAGAAA CCTAAGCCGC CTGGTGGATG GCTCGGCTCG GGGCGCTGAC GAAGGGCGTG GCAAGCTGCG ATAAGCCCCG G
23SBHB_168 CAACAGCC CGAGTTTCTT GCCCCTGAAA GAGGGGCTGG GCATTTGATA TGCCCAGCCC CTTGAGTAAG GGGCAAGAAA CCTAAGCCGC CTGGTGGATG GCTCGGCTCG GGGCGCTGAC GAAGGGCGTG GCAAGCTGCG ATAAGCCCCG
23SBHB_204 AACAGCC CGAGTTTCTT GCCCCTGAAA GAGGGGCTGG GCATTTGATA TGCCCAGCCC CTTGAGTAAG GGGCAAGAAA CCTAAGCCGC CTGGTGGATG GCTCGGCTCG GGGCGCTGAC GAAGGGCGTG GCAAGCTGCG ATAAGCCCCG G
23SBHB_121 AACAGCC CGAGTTTCTT GCCCCTGAAA GAGGGGCTGG GCATTTGATA TGCCCAGCCC CTTGAGTAAG GGGCAAGAAA CCTAAGCCGC CTGGTGGATG GCTCGGCTCG GGGCGCTGAC GAAGGGCGTG GCAAGCTGCG ATAAGCCCCG
23SBHB_203 GCC CGAGTTTCTT GCCCCTGAAA GAGGGGCTGG GCATTTGATA TGCCCAGCCC CTTGAGTAAG GGGCAAGAAA CCTAAGCCGC CTGGTGGATG GCTCGGCTCG GGGCGCTGAC GAAGGGCGTG GCAAGCTGCG ATAAGCCCCG G
23SBHB_115 CAACAGCC CGAGTTTCTT GCCCCTGAAA GAGGGGCTGG GCATTTGATA TGCCCAGCCC CTTGAGTAAG GGGCAAGAAA CCTAAGCCGC CTGGTGGATG GCTCGGCTCG GGGCGCTGAC GAAGGGCGTG GCAAGCTGCG ATAAGCC
23SBHB_273 CGAGTTTCTT GCCCCTGAAA GAGGGGCTGG GCATTTGATA TGCCCAGCCC CTTGAGTAAG GGGCAAGAAA CCTAAGCCGC CTGGTGGATG GCTCGGCTCG GGGCGCTGAC GAAGGGCGTG GCAAGCTGCG ATAAGCCCCG GTGAG
23SBHB_124 GCC CGAGTTTCTT GCCCCTGAAA GAGGGGCTGG GCATTTGATA TGCCCAGCCC CTTGAGTAAG GGGCAAGAAA CCTAAGCCGC CTGGTGGATG GCTCGGCTCG GGGCGCTGAC GAAGGGCGTG GCAAGCTGCG ATAAGCCCCG
23SBHB_328 C CGAGTTTCTT GCCCCTGAAA GAGGGGCTGG GCATTTGATA TGCCCAGCCC CTTGAGTAAG GGGCAAGAAA CCTAAGCCGC CTGGTGGATG GCTCGGCTCG GGGCGCTGAC GAAGGGCGTG GCAAGCTGCG ATAAGCCCCG G
23SBHB_363 CC CGAGTTTCTT GCCCCTGAAA GAGGGGCTGG GCATTTGATA TGCCCAGCCC CTTGAGTAAG GGGCAAGAAA CCTAAGCCGC CTGGTGGATG GCTCGGCTCG GGGCGCTGAC GAAGGGCGTG GCAAGCTGCG ATAAGCCCCG
23SBHB_205 CGAGTTTCTT GCCCCTGAAA GAGGGGCTGG GCATTTGATA TGCCCAGCCC CTTGAGTAAG GGGCAAGAAA CCTAAGCCGC CTGGTGGATG GCTCGGCTCG GGGCGCTGAC GAAGGGCGTG GCAAGCTGCG ATAAGCCCCG G
23SBHB_217 C CGAGTTTCTT GCCCCTGAAA GAGGGGCTGG GCATTTGATA TGCCCAGCCC CTTGAGTAAG GGGCAAGAAA CCTAAGCCGC CTGGTGGATG GCTCGGCTCG GGGCGCTGAC GAAGGGCGTG GCAAGCTGCG ATAAGCCCCG
23SBHB_202 CCCAACAGCC CGAGTTTCTT GCCCCTGAAA GAGGGGCTGG GCATTTGATA TGCCCAGCCC CTTGAGTAAG GGGCAAGAAA CCTAAGCCGC CTGGTGGATG GCTCGGCTCG GGGCGCTGAC GAAGGGCGTG GCAAGCTGCG AT
23SBHB_156 CT CCCAACAGCC CGAGTTTCTT GCCCCTGAAA GAGGGGCTGG GCATTTGATA TGCCCAGCCC CTTGAGTAAG GGGCAAGAAA CCTAAGCCGC CTGGTGGATG GCTCGGCTCG GGGCGCTGAC GAAGGGCGTG GCAAGCTGCG AT
23SBHB_124 CC CGAGTTTCTT GCCCCTGAAA GAGGGGCTGG GCATTTGATA TGCCCAGCCC CTTGAGTAAG GGGCAAGAAA CCTAAGCCGC CTGGTGGATG GCTCGGCTCG GGGCGCTGAC GAAGGGCGTG GCAAGCTGCG ATAAGCCCC
23SBHB_647 GTTTA GTCTGCCGCT CCCAACAGCC CGAGTTTCTT GCCCCTGAAA GAGGGGCTGG GCATTTGATA TGCCCAGCCC CTTGAGTAAG GGGCAAGAAA CCTAAGCCGC CTGGTGGATG GCTCGGCTCG GGGCGCTGAC GAAGGG
23SBHB_257 CGTTTA GTCTGCCGCT CCCAACAGCC CGAGTTTCTT GCCCCTGAAA GAGGGGCTGG GCATTTGATA TGCCCAGCCC CTTGAGTAAG GGGCAAGAAA CCTAAGCCGC CTGGTGGATG GCTCGGCTCG GGGCGCTGAC GAAGGG
23SBHB_177 CGTTTA GTCTGCCGCT CCCAACAGCC CGAGTTTCTT GCCCCTGAAA GAGGGGCTGG GCATTTGATA TGCCCAGCCC CTTGAGTAAG GGGCAAGAAA CCTAAGCCGC CTGGTGGATG GCTCGGCTCG GGGCGCTGAC GAAGG
23SBHB_350 GTTTA GTCTGCCGCT CCCAACAGCC CGAGTTTCTT GCCCCTGAAA GAGGGGCTGG GCATTTGATA TGCCCAGCCC CTTGAGTAAG GGGCAAGAAA CCTAAGCCGC CTGGTGGATG GCTCGGCTCG GGGCGCTGAC GAAGGGC
23SBHB_191 CGTTTA GTCTGCCGCT CCCAACAGCC CGAGTTTCTT GCCCCTGAAA GAGGGGCTGG GCATTTGATA TGCCCAGCCC CTTGAGTAAG GGGCAAGAAA CCTAAGCCGC CTGGTGGATG GCTCGGCTCG GGGCGCTGAC GAAG
23SBHB_449 GTTTA GTCTGCCGCT CCCAACAGCC CGAGTTTCTT GCCCCTGAAA GAGGGGCTGG GCATTTGATA TGCCCAGCCC CTTGAGTAAG GGGCAAGAAA CCTAAGCCGC CTGGTGGATG GCTCGGCTCG GGGCGCTGAC GAAGG
23SBHB_593 CCCGTTTA GTCTGCCGCT CCCAACAGCC CGAGTTTCTT GCCCCTGAAA GAGGGGCTGG GCATTTGATA TGCCCAGCCC CTTGAGTAAG GGGCAAGAAA CCTAAGCCGC CTGGTGGATG GCTCGGCTCG GGGCGCTGAC GAAG
23SBHB_166 GACCCGTTTA GTCTGCCGCT CCCAACAGCC CGAGTTTCTT GCCCCTGAAA GAGGGGCTGG GCATTTGATA TGCCCAGCCC CTTGAGTAAG GGGCAAGAAA CCTAAGCCGC CTGGTGGATG GCTCGGCTCG GGGCGCTGAC GAAG
23SBHB_456 GTTTA GTCTGCCGCT CCCAACAGCC CGAGTTTCTT GCCCCTGAAA GAGGGGCTGG GCATTTGATA TGCCCAGCCC CTTGAGTAAG GGGCAAGAAA CCTAAGCCGC CTGGTGGATG GCTCGGCTCG GGGCGCTGAC GAAG
23SBHB_218 CTGCCGCT CCCAACAGCC CGAGTTTCTT GCCCCTGAAA GAGGGGCTGG GCATTTGATA TGCCCAGCCC CTTGAGTAAG GGGCAAGAAA CCTAAGCCGC CTGGTGGATG GCTCGGCTCG GGGCGCTGAC GAAGGGCGTG GCAAG
23SBHB_200 CTGCCGCT CCCAACAGCC CGAGTTTCTT GCCCCTGAAA GAGGGGCTGG GCATTTGATA TGCCCAGCCC CTTGAGTAAG GGGCAAGAAA CCTAAGCCGC CTGGTGGATG GCTCGGCTCG GGGCGCTGAC GAAGGGCGTG G
23SBHB_406 GGGAAACC GACCCGTTTA GTCTGCCGCT CCCAACAGCC CGAGTTTCTT GCCCCTGAAA GAGGGGCTGG GCATTTGATA TGCCCAGCCC CTTGAGTAAG GGGCAAGAAA CCTAAGCCGC CTGGTGGATG GCTCGGCTCG GG
23SBHB_340 CAACAGCC CGAGTTTCTT GCCCCTGAAA GAGGGGCTGG GCATTTGATA TGCCCAGCCC CTTGAGTAAG GGGCAAGAAA CCTAAGCCGC CTGGTGGATG GCTCGGCTCG GGGCGCTGAC GAAGGGCGTG GCAAGCTGCG
23SBHB_287 CAACAGCC CGAGTTTCTT GCCCCTGAAA GAGGGGCTGG GCATTTGATA TGCCCAGCCC CTTGAGTAAG GGGCAAGAAA CCTAAGCCGC CTGGTGGATG GCTCGGCTCG GGGCGCTGAC GAAGGGCGTG GCAAGCTGCG AT
23SBHB_200 AACAGCC CGAGTTTCTT GCCCCTGAAA GAGGGGCTGG GCATTTGATA TGCCCAGCCC CTTGAGTAAG GGGCAAGAAA CCTAAGCCGC CTGGTGGATG GCTCGGCTCG GGGCGCTGAC GAAGGGCGTG GCAAGCTGCG AT
23SBHB_329 CT CCCAACAGCC CGAGTTTCTT GCCCCTGAAA GAGGGGCTGG GCATTTGATA TGCCCAGCCC CTTGAGTAAG GGGCAAGAAA CCTAAGCCGC CTGGTGGATG GCTCGGCTCG GGGCGCTGAC GAAGGGCGTG GCAAGCTGCG
23SBHB_137 CGCT CCCAACAGCC CGAGTTTCTT GCCCCTGAAA GAGGGGCTGG GCATTTGATA TGCCCAGCCC CTTGAGTAAG GGGCAAGAAA CCTAAGCCGC CTGGTGGATG GCTCGGCTCG GGGCGCTGAC GAAGGGCGTG GCAAGCTGCG
23SBHB_305 CCCAACAGCC CGAGTTTCTT GCCCCTGAAA GAGGGGCTGG GCATTTGATA TGCCCAGCCC CTTGAGTAAG GGGCAAGAAA CCTAAGCCGC CTGGTGGATG GCTCGGCTCG GGGCGCTGAC GAAGGGCGTG GCAAGCTGCG
23SBHB_125 CCAACAGCC CGAGTTTCTT GCCCCTGAAA GAGGGGCTGG GCATTTGATA TGCCCAGCCC CTTGAGTAAG GGGCAAGAAA CCTAAGCCGC CTGGTGGATG GCTCGGCTCG GGGCGCTGAC GAAGGGCGTG GCAAGCTGCG
23SBHB_292 AACAGCC CGAGTTTCTT GCCCCTGAAA GAGGGGCTGG GCATTTGATA TGCCCAGCCC CTTGAGTAAG GGGCAAGAAA CCTAAGCCGC CTGGTGGATG GCTCGGCTCG GGGCGCTGAC GAAGGGCGTG GCAAGCTGCG
23SBHB_145 CAGCC CGAGTTTCTT GCCCCTGAAA GAGGGGCTGG GCATTTGATA TGCCCAGCCC CTTGAGTAAG GGGCAAGAAA CCTAAGCCGC CTGGTGGATG GCTCGGCTCG GGGCGCTGAC GAAGGGCGTG GCAAGCTGCG
23SBHB_326 CTGCCGCT CCCAACAGCC CGAGTTTCTT GCCCCTGAAA GAGGGGCTGG GCATTTGATA TGCCCAGCCC CTTGAGTAAG GGGCAAGAAA CCTAAGCCGC CTGGTGGATG GCTCGGCTCG GGGCGCTGAC GAAGGG
23SBHB_222 GTCTGCCGCT CCCAACAGCC CGAGTTTCTT GCCCCTGAAA GAGGGGCTGG GCATTTGATA TGCCCAGCCC CTTGAGTAAG GGGCAAGAAA CCTAAGCCGC CTGGTGGATG GCTCGGCTCG GGGCGCTGAC GAAGGG
```

```
23SBHB_116 GTCTGCCGCT CCCAACAGCC CGAGTTTCTT GCCCCTGAAA GAGGGGCTGG GCATTTGATA TGCCCAGCCC CTTGAGTAAG GGGCAAGAAA CCTAAGCCGC CTGGTGGATG GCTCGGCTCG GGGCGCTGAC GAAGGGC
23SBHB_177 CTGCCGCT CCCAACAGCC CGAGTTTCTT GCCCCTGAAA GAGGGGCTGG GCATTTGATA TGCCCAGCCC CTTGAGTAAG GGGCAAGAAA CCTAAGCCGC CTGGTGGATG GCTCGGCTCG GGGCGCTGAC GAAGGGCG
23SBHB_183 CTGCCGCT CCCAACAGCC CGAGTTTCTT GCCCCTGAAA GAGGGGCTGG GCATTTGATA TGCCCAGCCC CTTGAGTAAG GGGCAAGAAA CCTAAGCCGC CTGGTGGATG GCTCGGCTCG GGGCGCTGAC GAAGGGC
23SBHB_134 GTCTGCCGCT CCCAACAGCC CGAGTTTCTT GCCCCTGAAA GAGGGGCTGG GCATTTGATA TGCCCAGCCC CTTGAGTAAG GGGCAAGAAA CCTAAGCCGC CTGGTGGATG GCTCGGCTCG GGGCGCTGAC GAAGG
23SBHB_131 GTCTGCCGCT CCCAACAGCC CGAGTTTCTT GCCCCTGAAA GAGGGGCTGG GCATTTGATA TGCCCAGCCC CTTGAGTAAG GGGCAAGAAA CCTAAGCCGC CTGGTGGATG GCTCGGCTCG GGGCGCTGAC GAAG
23SBHB_107 CGGGGA TGTAAGCGGG AAGGGAAACC GACCCGTTTA GTCTGCCGCT CCCAACAGCC CGAGTTTCTT GCCCCTGAAA GAGGGGCTGG GCATTTGATA TGCCCAGCCC CTTGAGTAAG GGGCAAGAAA CCTAAGCCGC CTGGTG
23SBHB_212 CGGGGA TGTAAGCGGG AAGGGAAACC GACCCGTTTA GTCTGCCGCT CCCAACAGCC CGAGTTTCTT GCCCCTGAAA GAGGGGCTGG GCATTTGATA TGCCCAGCCC CTTGAGTAAG GGGCAAGAAA CCTAAGCCGC CTGGTGGATG
23SBHB_123 CGGGGA TGTAAGCGGG AAGGGAAACC GACCCGTTTA GTCTGCCGCT CCCAACAGCC CGAGTTTCTT GCCCCTGAAA GAGGGGCTGG GCATTTGATA TGCCCAGCCC CTTGAGTAAG GGGCAAGAAA CCTAAGCCGC CTGGTGGATG G
23SBHB_207 CGGGGA TGTAAGCGGG AAGGGAAACC GACCCGTTTA GTCTGCCGCT CCCAACAGCC CGAGTTTCTT GCCCCTGAAA GAGGGGCTGG GCATTTGATA TGCCCAGCCC CTTGAGTAAG GGGCAAGAAA CCTAAGCCGC CTGGTGGAT
23SBHB_199 GGCGGGGA TGTAAGCGGG AAGGGAAACC GACCCGTTTA GTCTGCCGCT CCCAACAGCC CGAGTTTCTT GCCCCTGAAA GAGGGGCTGG GCATTTGATA TGCCCAGCCC CTTGAGTAAG GGGCAAGAAA CCTAAGCCGC CTGGTG
23SBHB_121 GCGGGGA TGTAAGCGGG AAGGGAAACC GACCCGTTTA GTCTGCCGCT CCCAACAGCC CGAGTTTCTT GCCCCTGAAA GAGGGGCTGG GCATTTGATA TGCCCAGCCC CTTGAGTAAG GGGCAAGAAA CCTAAGCCGC CTGGTG
23SBHB_266 GGGA TGTAAGCGGG AAGGGAAACC GACCCGTTTA GTCTGCCGCT CCCAACAGCC CGAGTTTCTT GCCCCTGAAA GAGGGGCTGG GCATTTGATA TGCCCAGCCC CTTGAGTAAG GGGCAAGAAA CCTAAGCCGC CTGGTG
23SBHB_400 CGGGGA TGTAAGCGGG AAGGGAAACC GACCCGTTTA GTCTGCCGCT CCCAACAGCC CGAGTTTCTT GCCCCTGAAA GAGGGGCTGG GCATTTGATA TGCCCAGCCC CTTGAGTAAG GGGCAAGAAA CCTAAGCCGC CTG
23SBHB_140 GGA TGTAAGCGGG AAGGGAAACC GACCCGTTTA GTCTGCCGCT CCCAACAGCC CGAGTTTCTT GCCCCTGAAA GAGGGGCTGG GCATTTGATA TGCCCAGCCC CTTGAGTAAG GGGCAAGAAA CCTAAGCCGC CTGGTG
23SBHB_157 GGGA TGTAAGCGGG AAGGGAAACC GACCCGTTTA GTCTGCCGCT CCCAACAGCC CGAGTTTCTT GCCCCTGAAA GAGGGGCTGG GCATTTGATA TGCCCAGCCC CTTGAGTAAG GGGCAAGAAA CCTAAGCCGC CTG
23SBHB_184 GTAAGCGGG AAGGGAAACC GACCCGTTTA GTCTGCCGCT CCCAACAGCC CGAGTTTCTT GCCCCTGAAA GAGGGGCTGG GCATTTGATA TGCCCAGCCC CTTGAGTAAG GGGCAAGAAA CCTAAGCCGC CTGGTG
23SBHB_222 CTGCCGCT CCCAACAGCC CGAGTTTCTT GCCCCTGAAA GAGGGGCTGG GCATTTGATA TGCCCAGCCC CTTGAGTAAG GGGCAAGAAA CCTAAGCCGC CTGGTGGATG GCTCGGCTCG GGGCGCTGAC GAAG
23SBHB_190 CTGCCGCT CCCAACAGCC CGAGTTTCTT GCCCCTGAAA GAGGGGCTGG GCATTTGATA TGCCCAGCCC CTTGAGTAAG GGGCAAGAAA CCTAAGCCGC CTGGTGGATG GCTCGGCTCG GGGCGCTGAC GAAGG
23SBHB_206 C GACCCGTTTA GTCTGCCGCT CCCAACAGCC CGAGTTTCTT GCCCCTGAAA GAGGGGCTGG GCATTTGATA TGCCCAGCCC CTTGAGTAAG GGGCAAGAAA CCTAAGCCGC CTGGTGGATG GCTCGGCTCG GG
23SBHB_446 CT CCCAACAGCC CGAGTTTCTT GCCCCTGAAA GAGGGGCTGG GCATTTGATA TGCCCAGCCC CTTGAGTAAG GGGCAAGAAA CCTAAGCCGC CTGGTGGATG GCTCGGCTCG GGGCGCTGAC GAAGGGCG
23SBHB_289 CT CCCAACAGCC CGAGTTTCTT GCCCCTGAAA GAGGGGCTGG GCATTTGATA TGCCCAGCCC CTTGAGTAAG GGGCAAGAAA CCTAAGCCGC CTGGTGGATG GCTCGGCTCG GGGCGCTGAC GAAGGGCGTG G
23SBHB_159 CT CCCAACAGCC CGAGTTTCTT GCCCCTGAAA GAGGGGCTGG GCATTTGATA TGCCCAGCCC CTTGAGTAAG GGGCAAGAAA CCTAAGCCGC CTGGTGGATG GCTCGGCTCG GGGCGCTGAC GAAGGGCGTG
23SBHB_273 CCCAACAGCC CGAGTTTCTT GCCCCTGAAA GAGGGGCTGG GCATTTGATA TGCCCAGCCC CTTGAGTAAG GGGCAAGAAA CCTAAGCCGC CTGGTGGATG GCTCGGCTCG GGGCGCTGAC GAAGGGCGTG G
23SBHB_199 CCCAACAGCC CGAGTTTCTT GCCCCTGAAA GAGGGGCTGG GCATTTGATA TGCCCAGCCC CTTGAGTAAG GGGCAAGAAA CCTAAGCCGC CTGGTGGATG GCTCGGCTCG GGGCGCTGAC GAAGGGCGTG
23SBHB_149 CCAACAGCC CGAGTTTCTT GCCCCTGAAA GAGGGGCTGG GCATTTGATA TGCCCAGCCC CTTGAGTAAG GGGCAAGAAA CCTAAGCCGC CTGGTGGATG GCTCGGCTCG GGGCGCTGAC GAAGGGCGTG G
23SBHB_305 CAACAGCC CGAGTTTCTT GCCCCTGAAA GAGGGGCTGG GCATTTGATA TGCCCAGCCC CTTGAGTAAG GGGCAAGAAA CCTAAGCCGC CTGGTGGATG GCTCGGCTCG GGGCGCTGAC GAAGGGCGTG G
23SBHB_417 CT CCCAACAGCC CGAGTTTCTT GCCCCTGAAA GAGGGGCTGG GCATTTGATA TGCCCAGCCC CTTGAGTAAG GGGCAAGAAA CCTAAGCCGC CTGGTGGATG GCTCGGCTCG GGGCGCTGAC GAAGGGC
23SBHB_225 CGCT CCCAACAGCC CGAGTTTCTT GCCCCTGAAA GAGGGGCTGG GCATTTGATA TGCCCAGCCC CTTGAGTAAG GGGCAAGAAA CCTAAGCCGC CTGGTGGATG GCTCGGCTCG GGGCGCTGAC GAAGG
23SBHB_130 CGCT CCCAACAGCC CGAGTTTCTT GCCCCTGAAA GAGGGGCTGG GCATTTGATA TGCCCAGCCC CTTGAGTAAG GGGCAAGAAA CCTAAGCCGC CTGGTGGATG GCTCGGCTCG GGGCGCTGAC GAAGGGCG
23SBHB_125 CGCT CCCAACAGCC CGAGTTTCTT GCCCCTGAAA GAGGGGCTGG GCATTTGATA TGCCCAGCCC CTTGAGTAAG GGGCAAGAAA CCTAAGCCGC CTGGTGGATG GCTCGGCTCG GGGCGCTGAC GAAGGGC
23SBHB_147 CGCT CCCAACAGCC CGAGTTTCTT GCCCCTGAAA GAGGGGCTGG GCATTTGATA TGCCCAGCCC CTTGAGTAAG GGGCAAGAAA CCTAAGCCGC CTGGTGGATG GCTCGGCTCG GGGCGCTGAC GAAGG
23SBHB_306 CCCGTTTA GTCTGCCGCT CCCAACAGCC CGAGTTTCTT GCCCCTGAAA GAGGGGCTGG GCATTTGATA TGCCCAGCCC CTTGAGTAAG GGGCAAGAAA CCTAAGCCGC CTGGTGGATG GCTCGGCTCG GG
23SBHB_845 CC CGAGTTTCTT GCCCCTGAAA GAGGGGCTGG GCATTTGATA TGCCCAGCCC CTTGAGTAAG GGGCAAGAAA CCTAAGCCGC CTGGTGGATG GCTCGGCTCG GGGCGCTGAC GAAGGGCGTG GCAAGCTGCG
23SBHB_730 CC CGAGTTTCTT GCCCCTGAAA GAGGGGCTGG GCATTTGATA TGCCCAGCCC CTTGAGTAAG GGGCAAGAAA CCTAAGCCGC CTGGTGGATG GCTCGGCTCG GGGCGCTGAC GAAGGGCGTG GCAAGCTGCG AT
23SBHB_247 GCC CGAGTTTCTT GCCCCTGAAA GAGGGGCTGG GCATTTGATA TGCCCAGCCC CTTGAGTAAG GGGCAAGAAA CCTAAGCCGC CTGGTGGATG GCTCGGCTCG GGGCGCTGAC GAAGGGCGTG GCAAGCTGCG AT
23SBHB_428 C CGAGTTTCTT GCCCCTGAAA GAGGGGCTGG GCATTTGATA TGCCCAGCCC CTTGAGTAAG GGGCAAGAAA CCTAAGCCGC CTGGTGGATG GCTCGGCTCG GGGCGCTGAC GAAGGGCGTG GCAAGCTGCG AT
23SBHB_309 CGAGTTTCTT GCCCCTGAAA GAGGGGCTGG GCATTTGATA TGCCCAGCCC CTTGAGTAAG GGGCAAGAAA CCTAAGCCGC CTGGTGGATG GCTCGGCTCG GGGCGCTGAC GAAGGGCGTG GCAAGCTGCG AT
23SBHB_367 GCC CGAGTTTCTT GCCCCTGAAA GAGGGGCTGG GCATTTGATA TGCCCAGCCC CTTGAGTAAG GGGCAAGAAA CCTAAGCCGC CTGGTGGATG GCTCGGCTCG GGGCGCTGAC GAAGGGCGTG GCAAGCTGCG
23SBHB_514 C CGAGTTTCTT GCCCCTGAAA GAGGGGCTGG GCATTTGATA TGCCCAGCCC CTTGAGTAAG GGGCAAGAAA CCTAAGCCGC CTGGTGGATG GCTCGGCTCG GGGCGCTGAC GAAGGGCGTG GCAAGCTGCG
23SBHB_190 CTT GCCCCTGAAA GAGGGGCTGG GCATTTGATA TGCCCAGCCC CTTGAGTAAG GGGCAAGAAA CCTAAGCCGC CTGGTGGATG GCTCGGCTCG GGGCGCTGAC GAAGGGCGTG GCAAGCTGCG ATAAGCCCCG GTGAG
23SBHB_168 GTTTCTT GCCCCTGAAA GAGGGGCTGG GCATTTGATA TGCCCAGCCC CTTGAGTAAG GGGCAAGAAA CCTAAGCCGC CTGGTGGATG GCTCGGCTCG GGGCGCTGAC GAAGGGCGTG GCAAGCTGCG ATAAGCCCCG GTGAG
23SBHB_124 GTTTCTT GCCCCTGAAA GAGGGGCTGG GCATTTGATA TGCCCAGCCC CTTGAGTAAG GGGCAAGAAA CCTAAGCCGC CTGGTGGATG GCTCGGCTCG GGGCGCTGAC GAAGGGCGTG GCAAGCTGCG ATAAGCCCCG G
23SBHB_188 CC CGAGTTTCTT GCCCCTGAAA GAGGGGCTGG GCATTTGATA TGCCCAGCCC CTTGAGTAAG GGGCAAGAAA CCTAAGCCGC CTGGTGGATG GCTCGGCTCG GGGCGCTGAC GAAGGGCGTG GCAAGCTGCG ATAAGCC
23SBHB_175 CGAGTTTCTT GCCCCTGAAA GAGGGGCTGG GCATTTGATA TGCCCAGCCC CTTGAGTAAG GGGCAAGAAA CCTAAGCCGC CTGGTGGATG GCTCGGCTCG GGGCGCTGAC GAAGGGCGTG GCAAGCTGCG ATAAGCCCCG
23SBHB_124 CGAGTTTCTT GCCCCTGAAA GAGGGGCTGG GCATTTGATA TGCCCAGCCC CTTGAGTAAG GGGCAAGAAA CCTAAGCCGC CTGGTGGATG GCTCGGCTCG GGGCGCTGAC GAAGGGCGTG GCAAGCTGCG ATAAGCC
23SBHB_187 GTTTCTT GCCCCTGAAA GAGGGGCTGG GCATTTGATA TGCCCAGCCC CTTGAGTAAG GGGCAAGAAA CCTAAGCCGC CTGGTGGATG GCTCGGCTCG GGGCGCTGAC GAAGGGCGTG GCAAGCTGCG AT
23SBHB_144 CC CGAGTTTCTT GCCCCTGAAA GAGGGGCTGG GCATTTGATA TGCCCAGCCC CTTGAGTAAG GGGCAAGAAA CCTAAGCCGC CTGGTGGATG GCTCGGCTCG GGGCGCTGAC GAAGGGCGTG GCAAGCTG
23SBHB_138 C CGAGTTTCTT GCCCCTGAAA GAGGGGCTGG GCATTTGATA TGCCCAGCCC CTTGAGTAAG GGGCAAGAAA CCTAAGCCGC CTGGTGGATG GCTCGGCTCG GGGCGCTGAC GAAGGGCGTG GCAAGCTG
23SBHB_407 CGAGTTTCTT GCCCCTGAAA GAGGGGCTGG GCATTTGATA TGCCCAGCCC CTTGAGTAAG GGGCAAGAAA CCTAAGCCGC CTGGTGGATG GCTCGGCTCG GGGCGCTGAC GAAGGGCGTG GCAAGCTGCG
23SBHB_171 CGCT CCCAACAGCC CGAGTTTCTT GCCCCTGAAA GAGGGGCTGG GCATTTGATA TGCCCAGCCC CTTGAGTAAG GGGCAAGAAA CCTAAGCCGC CTGGTGGATG GCTCGGCTCG GGGCGCTGAC GAAG
23SBHB_722 CT CCCAACAGCC CGAGTTTCTT GCCCCTGAAA GAGGGGCTGG GCATTTGATA TGCCCAGCCC CTTGAGTAAG GGGCAAGAAA CCTAAGCCGC CTGGTGGATG GCTCGGCTCG GGGCGCTGAC GAAGGG
23SBHB_443 CT CCCAACAGCC CGAGTTTCTT GCCCCTGAAA GAGGGGCTGG GCATTTGATA TGCCCAGCCC CTTGAGTAAG GGGCAAGAAA CCTAAGCCGC CTGGTGGATG GCTCGGCTCG GGGCGCTGAC GAAGG
23SBHB_934 CCCAACAGCC CGAGTTTCTT GCCCCTGAAA GAGGGGCTGG GCATTTGATA TGCCCAGCCC CTTGAGTAAG GGGCAAGAAA CCTAAGCCGC CTGGTGGATG GCTCGGCTCG GGGCGCTGAC GAAGGG
23SBHB_562 CCCAACAGCC CGAGTTTCTT GCCCCTGAAA GAGGGGCTGG GCATTTGATA TGCCCAGCCC CTTGAGTAAG GGGCAAGAAA CCTAAGCCGC CTGGTGGATG GCTCGGCTCG GGGCGCTGAC GAAGGGC
23SBHB_555 CCCAACAGCC CGAGTTTCTT GCCCCTGAAA GAGGGGCTGG GCATTTGATA TGCCCAGCCC CTTGAGTAAG GGGCAAGAAA CCTAAGCCGC CTGGTGGATG GCTCGGCTCG GGGCGCTGAC GAAGGGCG
23SBHB_236 CCAACAGCC CGAGTTTCTT GCCCCTGAAA GAGGGGCTGG GCATTTGATA TGCCCAGCCC CTTGAGTAAG GGGCAAGAAA CCTAAGCCGC CTGGTGGATG GCTCGGCTCG GGGCGCTGAC GAAGGGCG
23SBHB_222 CCAACAGCC CGAGTTTCTT GCCCCTGAAA GAGGGGCTGG GCATTTGATA TGCCCAGCCC CTTGAGTAAG GGGCAAGAAA CCTAAGCCGC CTGGTGGATG GCTCGGCTCG GGGCGCTGAC GAAGGGC
23SBHB_558 CAACAGCC CGAGTTTCTT GCCCCTGAAA GAGGGGCTGG GCATTTGATA TGCCCAGCCC CTTGAGTAAG GGGCAAGAAA CCTAAGCCGC CTGGTGGATG GCTCGGCTCG GGGCGCTGAC GAAGGGCG
23SBHB_245 AACAGCC CGAGTTTCTT GCCCCTGAAA GAGGGGCTGG GCATTTGATA TGCCCAGCCC CTTGAGTAAG GGGCAAGAAA CCTAAGCCGC CTGGTGGATG GCTCGGCTCG GGGCGCTGAC GAAGGGCGTG G
23SBHB_174 CAACAGCC CGAGTTTCTT GCCCCTGAAA GAGGGGCTGG GCATTTGATA TGCCCAGCCC CTTGAGTAAG GGGCAAGAAA CCTAAGCCGC CTGGTGGATG GCTCGGCTCG GGGCGCTGAC GAAGGGCGTG
23SBHB_143 AACAGCC CGAGTTTCTT GCCCCTGAAA GAGGGGCTGG GCATTTGATA TGCCCAGCCC CTTGAGTAAG GGGCAAGAAA CCTAAGCCGC CTGGTGGATG GCTCGGCTCG GGGCGCTGAC GAAGGGCGTG
23SBHB_236 GTTTA GTCTGCCGCT CCCAACAGCC CGAGTTTCTT GCCCCTGAAA GAGGGGCTGG GCATTTGATA TGCCCAGCCC CTTGAGTAAG GGGCAAGAAA CCTAAGCCGC CTGGTGGATG GCTCGGCTCG GG
23SBHB_576 CCCAACAGCC CGAGTTTCTT GCCCCTGAAA GAGGGGCTGG GCATTTGATA TGCCCAGCCC CTTGAGTAAG GGGCAAGAAA CCTAAGCCGC CTGGTGGATG GCTCGGCTCG GGGCGCTGAC GAAGG
23SBHB_522 CT CCCAACAGCC CGAGTTTCTT GCCCCTGAAA GAGGGGCTGG GCATTTGATA TGCCCAGCCC CTTGAGTAAG GGGCAAGAAA CCTAAGCCGC CTGGTGGATG GCTCGGCTCG GGGCGCTGAC GAAG
23SBHB_102 CAACAGCC CGAGTTTCTT GCCCCTGAAA GAGGGGCTGG GCATTTGATA TGCCCAGCCC CTTGAGTAAG GGGCAAGAAA CCTAAGCCGC CTGGTGGATG GCTCGGCTCG GGGCGCTGAC GAAGGG
23SBHB_588 CAACAGCC CGAGTTTCTT GCCCCTGAAA GAGGGGCTGG GCATTTGATA TGCCCAGCCC CTTGAGTAAG GGGCAAGAAA CCTAAGCCGC CTGGTGGATG GCTCGGCTCG GGGCGCTGAC GAAGGGC
23SBHB_420 CCAACAGCC CGAGTTTCTT GCCCCTGAAA GAGGGGCTGG GCATTTGATA TGCCCAGCCC CTTGAGTAAG GGGCAAGAAA CCTAAGCCGC CTGGTGGATG GCTCGGCTCG GGGCGCTGAC GAAGGG
23SBHB_532 AACAGCC CGAGTTTCTT GCCCCTGAAA GAGGGGCTGG GCATTTGATA TGCCCAGCCC CTTGAGTAAG GGGCAAGAAA CCTAAGCCGC CTGGTGGATG GCTCGGCTCG GGGCGCTGAC GAAGGGC
23SBHB_459 AACAGCC CGAGTTTCTT GCCCCTGAAA GAGGGGCTGG GCATTTGATA TGCCCAGCCC CTTGAGTAAG GGGCAAGAAA CCTAAGCCGC CTGGTGGATG GCTCGGCTCG GGGCGCTGAC GAAGGGCG
23SBHB_263 CCAACAGCC CGAGTTTCTT GCCCCTGAAA GAGGGGCTGG GCATTTGATA TGCCCAGCCC CTTGAGTAAG GGGCAAGAAA CCTAAGCCGC CTGGTGGATG GCTCGGCTCG GGGCGCTGAC GAAGG
23SBHB_619 CCCAACAGCC CGAGTTTCTT GCCCCTGAAA GAGGGGCTGG GCATTTGATA TGCCCAGCCC CTTGAGTAAG GGGCAAGAAA CCTAAGCCGC CTGGTGGATG GCTCGGCTCG GGGCGCTGAC GAAG
23SBHB_201 CC CGAGTTTCTT GCCCCTGAAA GAGGGGCTGG GCATTTGATA TGCCCAGCCC CTTGAGTAAG GGGCAAGAAA CCTAAGCCGC CTGGTGGATG GCTCGGCTCG GGGCGCTGAC GAAGGGCGTG GCAAG
23SBHB_136 C CGAGTTTCTT GCCCCTGAAA GAGGGGCTGG GCATTTGATA TGCCCAGCCC CTTGAGTAAG GGGCAAGAAA CCTAAGCCGC CTGGTGGATG GCTCGGCTCG GGGCGCTGAC GAAGGGCGTG GCAAG
23SBHB_126 CGAGTTTCTT GCCCCTGAAA GAGGGGCTGG GCATTTGATA TGCCCAGCCC CTTGAGTAAG GGGCAAGAAA CCTAAGCCGC CTGGTGGATG GCTCGGCTCG GGGCGCTGAC GAAGGGCGTG GCAAG
23SBHB_119 CC CGAGTTTCTT GCCCCTGAAA GAGGGGCTGG GCATTTGATA TGCCCAGCCC CTTGAGTAAG GGGCAAGAAA CCTAAGCCGC CTGGTGGATG GCTCGGCTCG GGGCGCTGAC GAAGGGCGTG GC
23SBHB_633 CAACAGCC CGAGTTTCTT GCCCCTGAAA GAGGGGCTGG GCATTTGATA TGCCCAGCCC CTTGAGTAAG GGGCAAGAAA CCTAAGCCGC CTGGTGGATG GCTCGGCTCG GGGCGCTGAC GAAGG
23SBHB_269 CCAACAGCC CGAGTTTCTT GCCCCTGAAA GAGGGGCTGG GCATTTGATA TGCCCAGCCC CTTGAGTAAG GGGCAAGAAA CCTAAGCCGC CTGGTGGATG GCTCGGCTCG GGGCGCTGAC GAAG
23SBHB_725 CAACAGCC CGAGTTTCTT GCCCCTGAAA GAGGGGCTGG GCATTTGATA TGCCCAGCCC CTTGAGTAAG GGGCAAGAAA CCTAAGCCGC CTGGTGGATG GCTCGGCTCG GGGCGCTGAC GAAG
23SBHB_890 AACAGCC CGAGTTTCTT GCCCCTGAAA GAGGGGCTGG GCATTTGATA TGCCCAGCCC CTTGAGTAAG GGGCAAGAAA CCTAAGCCGC CTGGTGGATG GCTCGGCTCG GGGCGCTGAC GAAGGG
23SBHB_539 AACAGCC CGAGTTTCTT GCCCCTGAAA GAGGGGCTGG GCATTTGATA TGCCCAGCCC CTTGAGTAAG GGGCAAGAAA CCTAAGCCGC CTGGTGGATG GCTCGGCTCG GGGCGCTGAC GAAGG
23SBHB_453 CGGG AAGGGAAACC GACCCGTTTA GTCTGCCGCT CCCAACAGCC CGAGTTTCTT GCCCCTGAAA GAGGGGCTGG GCATTTGATA TGCCCAGCCC CTTGAGTAAG GGGCAAGAAA CCTAAGCCGC CTGGTG
23SBHB_161 CGGG AAGGGAAACC GACCCGTTTA GTCTGCCGCT CCCAACAGCC CGAGTTTCTT GCCCCTGAAA GAGGGGCTGG GCATTTGATA TGCCCAGCCC CTTGAGTAAG GGGCAAGAAA CCTAAGCCGC CTG
23SBHB_215 GGGAAACC GACCCGTTTA GTCTGCCGCT CCCAACAGCC CGAGTTTCTT GCCCCTGAAA GAGGGGCTGG GCATTTGATA TGCCCAGCCC CTTGAGTAAG GGGCAAGAAA CCTAAGCCGC CTGGTG
23SBHB_688 CC CGAGTTTCTT GCCCCTGAAA GAGGGGCTGG GCATTTGATA TGCCCAGCCC CTTGAGTAAG GGGCAAGAAA CCTAAGCCGC CTGGTGGATG GCTCGGCTCG GGGCGCTGAC GAAGGGCGTG G
23SBHB_235 GCC CGAGTTTCTT GCCCCTGAAA GAGGGGCTGG GCATTTGATA TGCCCAGCCC CTTGAGTAAG GGGCAAGAAA CCTAAGCCGC CTGGTGGATG GCTCGGCTCG GGGCGCTGAC GAAGGGCGTG G
23SBHB_122 CAGCC CGAGTTTCTT GCCCCTGAAA GAGGGGCTGG GCATTTGATA TGCCCAGCCC CTTGAGTAAG GGGCAAGAAA CCTAAGCCGC CTGGTGGATG GCTCGGCTCG GGGCGCTGAC GAAGGGCGTG G
23SBHB_130 GCC CGAGTTTCTT GCCCCTGAAA GAGGGGCTGG GCATTTGATA TGCCCAGCCC CTTGAGTAAG GGGCAAGAAA CCTAAGCCGC CTGGTGGATG GCTCGGCTCG GGGCGCTGAC GAAGGGCGTG
23SBHB_430 C CGAGTTTCTT GCCCCTGAAA GAGGGGCTGG GCATTTGATA TGCCCAGCCC CTTGAGTAAG GGGCAAGAAA CCTAAGCCGC CTGGTGGATG GCTCGGCTCG GGGCGCTGAC GAAGGGCGTG G
23SBHB_380 CC CGAGTTTCTT GCCCCTGAAA GAGGGGCTGG GCATTTGATA TGCCCAGCCC CTTGAGTAAG GGGCAAGAAA CCTAAGCCGC CTGGTGGATG GCTCGGCTCG GGGCGCTGAC GAAGGGCGTG
23SBHB_605 AACAGCC CGAGTTTCTT GCCCCTGAAA GAGGGGCTGG GCATTTGATA TGCCCAGCCC CTTGAGTAAG GGGCAAGAAA CCTAAGCCGC CTGGTGGATG GCTCGGCTCG GGGCGCTGAC GAAG
23SBHB_119 CC CGAGTTTCTT GCCCCTGAAA GAGGGGCTGG GCATTTGATA TGCCCAGCCC CTTGAGTAAG GGGCAAGAAA CCTAAGCCGC CTGGTGGATG GCTCGGCTCG GGGCGCTGAC GAAGGGCG
23SBHB_423 GCC CGAGTTTCTT GCCCCTGAAA GAGGGGCTGG GCATTTGATA TGCCCAGCCC CTTGAGTAAG GGGCAAGAAA CCTAAGCCGC CTGGTGGATG GCTCGGCTCG GGGCGCTGAC GAAGGGCG
23SBHB_229 CAGCC CGAGTTTCTT GCCCCTGAAA GAGGGGCTGG GCATTTGATA TGCCCAGCCC CTTGAGTAAG GGGCAAGAAA CCTAAGCCGC CTGGTGGATG GCTCGGCTCG GGGCGCTGAC GAAGGGCG
23SBHB_227 CAGCC CGAGTTTCTT GCCCCTGAAA GAGGGGCTGG GCATTTGATA TGCCCAGCCC CTTGAGTAAG GGGCAAGAAA CCTAAGCCGC CTGGTGGATG GCTCGGCTCG GGGCGCTGAC GAAGGGC
23SBHB_167 AGCC CGAGTTTCTT GCCCCTGAAA GAGGGGCTGG GCATTTGATA TGCCCAGCCC CTTGAGTAAG GGGCAAGAAA CCTAAGCCGC CTGGTGGATG GCTCGGCTCG GGGCGCTGAC GAAGGG
23SBHB_366 CAGCC CGAGTTTCTT GCCCCTGAAA GAGGGGCTGG GCATTTGATA TGCCCAGCCC CTTGAGTAAG GGGCAAGAAA CCTAAGCCGC CTGGTGGATG GCTCGGCTCG GGGCGCTGAC GAAGGG
23SBHB_558 GCC CGAGTTTCTT GCCCCTGAAA GAGGGGCTGG GCATTTGATA TGCCCAGCCC CTTGAGTAAG GGGCAAGAAA CCTAAGCCGC CTGGTGGATG GCTCGGCTCG GGGCGCTGAC GAAGGGC
23SBHB_293 CTT GCCCCTGAAA GAGGGGCTGG GCATTTGATA TGCCCAGCCC CTTGAGTAAG GGGCAAGAAA CCTAAGCCGC CTGGTGGATG GCTCGGCTCG GGGCGCTGAC GAAGGGCGTG GCAAGCTGCG
23SBHB_277 GTTTCTT GCCCCTGAAA GAGGGGCTGG GCATTTGATA TGCCCAGCCC CTTGAGTAAG GGGCAAGAAA CCTAAGCCGC CTGGTGGATG GCTCGGCTCG GGGCGCTGAC GAAGGGCGTG GCAAGCTGCG
23SBHB_170 CTT GCCCCTGAAA GAGGGGCTGG GCATTTGATA TGCCCAGCCC CTTGAGTAAG GGGCAAGAAA CCTAAGCCGC CTGGTGGATG GCTCGGCTCG GGGCGCTGAC GAAGGGCGTG GCAAGCTGCG AT
23SBHB_250 CAGCC CGAGTTTCTT GCCCCTGAAA GAGGGGCTGG GCATTTGATA TGCCCAGCCC CTTGAGTAAG GGGCAAGAAA CCTAAGCCGC CTGGTGGATG GCTCGGCTCG GGGCGCTGAC GAAGG
23SBHB_240 CAGCC CGAGTTTCTT GCCCCTGAAA GAGGGGCTGG GCATTTGATA TGCCCAGCCC CTTGAGTAAG GGGCAAGAAA CCTAAGCCGC CTGGTGGATG GCTCGGCTCG GGGCGCTGAC GAAG
23SBHB_668 C CGAGTTTCTT GCCCCTGAAA GAGGGGCTGG GCATTTGATA TGCCCAGCCC CTTGAGTAAG GGGCAAGAAA CCTAAGCCGC CTGGTGGATG GCTCGGCTCG GGGCGCTGAC GAAGGGCG
23SBHB_352 CGAGTTTCTT GCCCCTGAAA GAGGGGCTGG GCATTTGATA TGCCCAGCCC CTTGAGTAAG GGGCAAGAAA CCTAAGCCGC CTGGTGGATG GCTCGGCTCG GGGCGCTGAC GAAGGGCGTG G
23SBHB_240 C CGAGTTTCTT GCCCCTGAAA GAGGGGCTGG GCATTTGATA TGCCCAGCCC CTTGAGTAAG GGGCAAGAAA CCTAAGCCGC CTGGTGGATG GCTCGGCTCG GGGCGCTGAC GAAGGGCGTG
23SBHB_216 CGAGTTTCTT GCCCCTGAAA GAGGGGCTGG GCATTTGATA TGCCCAGCCC CTTGAGTAAG GGGCAAGAAA CCTAAGCCGC CTGGTGGATG GCTCGGCTCG GGGCGCTGAC GAAGGGCGTG
23SBHB_117 AGCC CGAGTTTCTT GCCCCTGAAA GAGGGGCTGG GCATTTGATA TGCCCAGCCC CTTGAGTAAG GGGCAAGAAA CCTAAGCCGC CTGGTGGATG GCTCGGCTCG GGGCGCTGAC GAAGG
23SBHB_232 CC CGAGTTTCTT GCCCCTGAAA GAGGGGCTGG GCATTTGATA TGCCCAGCCC CTTGAGTAAG GGGCAAGAAA CCTAAGCCGC CTGGTGGATG GCTCGGCTCG GGGCGCTGAC GAAGGG
23SBHB_757 GCC CGAGTTTCTT GCCCCTGAAA GAGGGGCTGG GCATTTGATA TGCCCAGCCC CTTGAGTAAG GGGCAAGAAA CCTAAGCCGC CTGGTGGATG GCTCGGCTCG GGGCGCTGAC GAAGGG
23SBHB_560 GCC CGAGTTTCTT GCCCCTGAAA GAGGGGCTGG GCATTTGATA TGCCCAGCCC CTTGAGTAAG GGGCAAGAAA CCTAAGCCGC CTGGTGGATG GCTCGGCTCG GGGCGCTGAC GAAGG
23SBHB_112 CC CGAGTTTCTT GCCCCTGAAA GAGGGGCTGG GCATTTGATA TGCCCAGCCC CTTGAGTAAG GGGCAAGAAA CCTAAGCCGC CTGGTGGATG GCTCGGCTCG GGGCGCTGAC GAAGGGC
23SBHB_120 AGCC CGAGTTTCTT GCCCCTGAAA GAGGGGCTGG GCATTTGATA TGCCCAGCCC CTTGAGTAAG GGGCAAGAAA CCTAAGCCGC CTGGTGGATG GCTCGGCTCG GGGCGCTGAC GAAG
23SBHB_151 CC CGAGTTTCTT GCCCCTGAAA GAGGGGCTGG GCATTTGATA TGCCCAGCCC CTTGAGTAAG GGGCAAGAAA CCTAAGCCGC CTGGTGGATG GCTCGGCTCG GGGCGCTGAC GAAGG
23SBHB_559 GCC CGAGTTTCTT GCCCCTGAAA GAGGGGCTGG GCATTTGATA TGCCCAGCCC CTTGAGTAAG GGGCAAGAAA CCTAAGCCGC CTGGTGGATG GCTCGGCTCG GGGCGCTGAC GAAG
23SBHB_255 GTTTCTT GCCCCTGAAA GAGGGGCTGG GCATTTGATA TGCCCAGCCC CTTGAGTAAG GGGCAAGAAA CCTAAGCCGC CTGGTGGATG GCTCGGCTCG GGGCGCTGAC GAAGGGCGTG G
23SBHB_132 GTTTCTT GCCCCTGAAA GAGGGGCTGG GCATTTGATA TGCCCAGCCC CTTGAGTAAG GGGCAAGAAA CCTAAGCCGC CTGGTGGATG GCTCGGCTCG GGGCGCTGAC GAAGGGCGTG
23SBHB_166 CC CGAGTTTCTT GCCCCTGAAA GAGGGGCTGG GCATTTGATA TGCCCAGCCC CTTGAGTAAG GGGCAAGAAA CCTAAGCCGC CTGGTGGATG GCTCGGCTCG GGGCGCTGAC GAAG
23SBHB_122 C CGAGTTTCTT GCCCCTGAAA GAGGGGCTGG GCATTTGATA TGCCCAGCCC CTTGAGTAAG GGGCAAGAAA CCTAAGCCGC CTGGTGGATG GCTCGGCTCG GGGCGCTGAC GAAGGG
23SBHB_706 C CGAGTTTCTT GCCCCTGAAA GAGGGGCTGG GCATTTGATA TGCCCAGCCC CTTGAGTAAG GGGCAAGAAA CCTAAGCCGC CTGGTGGATG GCTCGGCTCG GGGCGCTGAC GAAGGGC
23SBHB_577 CGAGTTTCTT GCCCCTGAAA GAGGGGCTGG GCATTTGATA TGCCCAGCCC CTTGAGTAAG GGGCAAGAAA CCTAAGCCGC CTGGTGGATG GCTCGGCTCG GGGCGCTGAC GAAGGGC
23SBHB_540 CGAGTTTCTT GCCCCTGAAA GAGGGGCTGG GCATTTGATA TGCCCAGCCC CTTGAGTAAG GGGCAAGAAA CCTAAGCCGC CTGGTGGATG GCTCGGCTCG GGGCGCTGAC GAAGGGCG
23SBHB_957 C CGAGTTTCTT GCCCCTGAAA GAGGGGCTGG GCATTTGATA TGCCCAGCCC CTTGAGTAAG GGGCAAGAAA CCTAAGCCGC CTGGTGGATG GCTCGGCTCG GGGCGCTGAC GAAGG
23SBHB_994 CGAGTTTCTT GCCCCTGAAA GAGGGGCTGG GCATTTGATA TGCCCAGCCC CTTGAGTAAG GGGCAAGAAA CCTAAGCCGC CTGGTGGATG GCTCGGCTCG GGGCGCTGAC GAAGGG
23SBHB_704 CGAGTTTCTT GCCCCTGAAA GAGGGGCTGG GCATTTGATA TGCCCAGCCC CTTGAGTAAG GGGCAAGAAA CCTAAGCCGC CTGGTGGATG GCTCGGCTCG GGGCGCTGAC GAAGG
23SBHB_107 C CGAGTTTCTT GCCCCTGAAA GAGGGGCTGG GCATTTGATA TGCCCAGCCC CTTGAGTAAG GGGCAAGAAA CCTAAGCCGC CTGGTGGATG GCTCGGCTCG GGGCGCTGAC GAAG
23SBHB_810 CGAGTTTCTT GCCCCTGAAA GAGGGGCTGG GCATTTGATA TGCCCAGCCC CTTGAGTAAG GGGCAAGAAA CCTAAGCCGC CTGGTGGATG GCTCGGCTCG GGGCGCTGAC GAAG
23SBHB_195 CT CCCAACAGCC CGAGTTTCTT GCCCCTGAAA GAGGGGCTGG GCATTTGATA TGCCCAGCCC CTTGAGTAAG GGGCAAGAAA CCTAAGCCGC CTGGTGGATG GCTCGGCTCG GG
23SBHB_199 CCCGTTTA GTCTGCCGCT CCCAACAGCC CGAGTTTCTT GCCCCTGAAA GAGGGGCTGG GCATTTGATA TGCCCAGCCC CTTGAGTAAG GGGCAAGAAA CCTAAGCCGC CTGGTG
23SBHB_135 C GACCCGTTTA GTCTGCCGCT CCCAACAGCC CGAGTTTCTT GCCCCTGAAA GAGGGGCTGG GCATTTGATA TGCCCAGCCC CTTGAGTAAG GGGCAAGAAA CCTAAGCCGC CTGGTG
23SBHB_647 GTTTCTT GCCCCTGAAA GAGGGGCTGG GCATTTGATA TGCCCAGCCC CTTGAGTAAG GGGCAAGAAA CCTAAGCCGC CTGGTGGATG GCTCGGCTCG GGGCGCTGAC GAAGGG
23SBHB_399 GTTTCTT GCCCCTGAAA GAGGGGCTGG GCATTTGATA TGCCCAGCCC CTTGAGTAAG GGGCAAGAAA CCTAAGCCGC CTGGTGGATG GCTCGGCTCG GGGCGCTGAC GAAGGGCG
23SBHB_364 GTTTCTT GCCCCTGAAA GAGGGGCTGG GCATTTGATA TGCCCAGCCC CTTGAGTAAG GGGCAAGAAA CCTAAGCCGC CTGGTGGATG GCTCGGCTCG GGGCGCTGAC GAAGGGC
23SBHB_174 CC CGAGTTTCTT GCCCCTGAAA GAGGGGCTGG GCATTTGATA TGCCCAGCCC CTTGAGTAAG GGGCAAGAAA CCTAAGCCGC CTGGTGGATG GCTCGGCTCG GGGCGCTGAC
23SBHB_144 CCCAACAGCC CGAGTTTCTT GCCCCTGAAA GAGGGGCTGG GCATTTGATA TGCCCAGCCC CTTGAGTAAG GGGCAAGAAA CCTAAGCCGC CTGGTGGATG GCTCGGCTCG GG
23SBHB_230 CTT GCCCCTGAAA GAGGGGCTGG GCATTTGATA TGCCCAGCCC CTTGAGTAAG GGGCAAGAAA CCTAAGCCGC CTGGTGGATG GCTCGGCTCG GGGCGCTGAC GAAGGGCGTG G
23SBHB_482 GTTTCTT GCCCCTGAAA GAGGGGCTGG GCATTTGATA TGCCCAGCCC CTTGAGTAAG GGGCAAGAAA CCTAAGCCGC CTGGTGGATG GCTCGGCTCG GGGCGCTGAC GAAGG
23SBHB_619 GTTTCTT GCCCCTGAAA GAGGGGCTGG GCATTTGATA TGCCCAGCCC CTTGAGTAAG GGGCAAGAAA CCTAAGCCGC CTGGTGGATG GCTCGGCTCG GGGCGCTGAC GAAG
23SBHB_165 CAACAGCC CGAGTTTCTT GCCCCTGAAA GAGGGGCTGG GCATTTGATA TGCCCAGCCC CTTGAGTAAG GGGCAAGAAA CCTAAGCCGC CTGGTGGATG GCTCGGCTCG GG
23SBHB_173 GTTTA GTCTGCCGCT CCCAACAGCC CGAGTTTCTT GCCCCTGAAA GAGGGGCTGG GCATTTGATA TGCCCAGCCC CTTGAGTAAG GGGCAAGAAA CCTAAGCCGC CTGGTG
23SBHB_187 AACAGCC CGAGTTTCTT GCCCCTGAAA GAGGGGCTGG GCATTTGATA TGCCCAGCCC CTTGAGTAAG GGGCAAGAAA CCTAAGCCGC CTGGTGGATG GCTCGGCTCG GG
23SBHB_590 CTT GCCCCTGAAA GAGGGGCTGG GCATTTGATA TGCCCAGCCC CTTGAGTAAG GGGCAAGAAA CCTAAGCCGC CTGGTGGATG GCTCGGCTCG GGGCGCTGAC GAAGGG
23SBHB_313 CTT GCCCCTGAAA GAGGGGCTGG GCATTTGATA TGCCCAGCCC CTTGAGTAAG GGGCAAGAAA CCTAAGCCGC CTGGTGGATG GCTCGGCTCG GGGCGCTGAC GAAGGGCG
```

```
23SBHB_291                                                        CTT GCCCCTGAAA GAGGGGCTGG GCATTTGATA TGCCCAGCCC CTTGAGTAAG GGGCAAGAAA CCTAAGCCGC CTGGTGGATG GCTCGGCTCG GGGCGCTGAC GAAGGGC
23SBHB_368                                                        CTT GCCCCTGAAA GAGGGGCTGG GCATTTGATA TGCCCAGCCC CTTGAGTAAG GGGCAAGAAA CCTAAGCCGC CTGGTGGATG GCTCGGCTCG GGGCGCTGAC GAAGG
23SBHB_494                                                        CTT GCCCCTGAAA GAGGGGCTGG GCATTTGATA TGCCCAGCCC CTTGAGTAAG GGGCAAGAAA CCTAAGCCGC CTGGTGGATG GCTCGGCTCG GGGCGCTGAC GAAG
23SBHB_134                                                GCC CGAGTTTCTT GCCCCTGAAA GAGGGGCTGG GCATTTGATA TGCCCAGCCC CTTGAGTAAG GGGCAAGAAA CCTAAGCCGC CTGGTGGATG GCTCGGCTCG GG
23SBHB_283                                                 CC CGAGTTTCTT GCCCCTGAAA GAGGGGCTGG GCATTTGATA TGCCCAGCCC CTTGAGTAAG GGGCAAGAAA CCTAAGCCGC CTGGTGGATG GCTCGGCTCG GG
23SBHB_234                                                  C CGAGTTTCTT GCCCCTGAAA GAGGGGCTGG GCATTTGATA TGCCCAGCCC CTTGAGTAAG GGGCAAGAAA CCTAAGCCGC CTGGTGGATG GCTCGGCTCG GG
23SBHB_133                                                    CGAGTTTCTT GCCCCTGAAA GAGGGGCTGG GCATTTGATA TGCCCAGCCC CTTGAGTAAG GGGCAAGAAA CCTAAGCCGC CTGGTGGATG GCTCGGCTCG GG
23SBHB_132                                                       GTTTCTT GCCCCTGAAA GAGGGGCTGG GCATTTGATA TGCCCAGCCC CTTGAGTAAG GGGCAAGAAA CCTAAGCCGC CTGGTGGATG GCTCGGCTCG GG
 Consensus .......... .......... .....gggga tgtaagcggg aagggaaacc gacccgttta gtctgccgct cccaacagCC CGAGTTTCTT GCCCCTGAAA GAGGGGCTGG GCATTTGATA TGCCCAGCCC CTTGAGTAAG GGGCAAGAAA CCTAAGCCGC CTGGTGGATG GCTCGGCTCG GGgcgctgac gaagggcg.. .......... .......... .......... .......... .......... .......... .......... .......... .......... ..........
```
