## Supplementary Figure S2c for "Permuted 23S rRNA is integrated in 50S ribosome particles in *Thermococcus barophilus*"

```
                                    _______________________________________23S : H99-101_____________________________________________|_23S : TH3p__|_Junction|_23S : TH5p_|____23S____________________________
                                                                                                                                      ..<< <<<<<<<<< ........>> >>>>>>>>>>

J23SpTT_6759                    CGCA GAAGACGGGT TTGATGGGGC GGGGATGTAA GCCGGGAAGG AAACCGACCC GTTTAGTCTG CCGCTCCCAA CAGCCCGAGT TTCTTGCCCC TTGAGTAAGG GGCAAGAAAC CTAAGCCGCC TGGTGGATGG CTCGGCTCGG GGCGCTGACG AAGGG
J23SpTT_3515                    CGCA GAAGACGGGT TTGATGGGGC GGGGATGTAA GCCGGGAAGG AAACCGACCC GTTTAGTCTG CCGCTCCCAA CAGCCCGAGT TTCTTGCCCC TTGAGTAAGG GGCAAGAAAC CTAAGCCGCC TGGTGGATGG CTCGGCTCGG GGCGCTGACG AAGGGC
J23SpTT_2810                  CCCGCA GAAGACGGGT TTGATGGGGC GGGGATGTAA GCCGGGAAGG AAACCGACCC GTTTAGTCTG CCGCTCCCAA CAGCCCGAGT TTCTTGCCCC TTGAGTAAGG GGCAAGAAAC CTAAGCCGCC TGGTGGATGG CTCGGCTCGG GGCGCTGACG AAGGG
J23SpTT_1482                  CCCGCA GAAGACGGGT TTGATGGGGC GGGGATGTAA GCCGGGAAGG AAACCGACCC GTTTAGTCTG CCGCTCCCAA CAGCCCGAGT TTCTTGCCCC TTGAGTAAGG GGCAAGAAAC CTAAGCCGCC TGGTGGATGG CTCGGCTCGG GGCGCTGACG AAGGGC
J23SpTT_835                   CCCGCA GAAGACGGGT TTGATGGGGC GGGGATGTAA GCCGGGAAGG AAACCGACCC GTTTAGTCTG CCGCTCCCAA CAGCCCGAGT TTCTTGCCCC TTGAGTAAGG GGCAAGAAAC CTAAGCCGCC TGGTGGATGG CTCGGCTCGG GGCGCTGACG AAGGGCG
J23SpTT_13                  CGCCCGCA GAAGACGGGT TTGATGGGGC GGGGATGTAA GCCGGGAAGG AAACCGACCC GTTTAGTCTG CCGCTCCCAA CAGCCCGAGT TTCTTGCCCC TTGAGTAAGG GGCAAGAAAC CTAAGCCGCC TGGTGGATGG CTCGGCTCGG GGCGCTGACG AAGGGCG
J23SpTT_10                   GCCCGCA GAAGACGGGT TTGATGGGGC GGGGATGTAA GCCGGGAAGG AAACCGACCC GTTTAGTCTG CCGCTCCCAA CAGCCCGAGT TTCTTGCCCC TTGAGTAAGG GGCAAGAAAC CTAAGCCGCC TGGTGGATGG CTCGGCTCGG GGCGCTGACG AAGGGCG
J23SpTT_12      GCCCTTCGAG GGCGCCCGCA GAAGACGGGT TTGATGGGGC GGGGATGTAA GCCGGGAAGG AAACCGACCC GTTTAGTCTG CCGCTCCCAA CAGCCCGAGT TTCTTGCCCC TTGAGTAAGG GGCAAGAAAC CTAAGCCGCC TGGTGGATGG CTCGGCTCGG GGCGCTGACG AAGGGC
J23SpTT_13                  CGCCCGCA GAAGACGGGT TTGATGGGGC GGGGATGTAA GCCGGGAAGG AAACCGACCC GTTTAGTCTG CCGCTCCCAA CAGCCCGAGT TTCTTGCCCC TTGAGTAAGG GGCAAGAAAC CTAAGCCGCC TGGTGGATGG CTCGGCTCGG GGCGCTGACG AAGGGC
J23SpTT_41                  CGCCCGCA GAAGACGGGT TTGATGGGGC GGGGATGTAA GCCGGGAAGG AAACCGACCC GTTTAGTCTG CCGCTCCCAA CAGCCCGAGT TTCTTGCCCC TTGAGTAAGG GGCAAGAAAC CTAAGCCGCC TGGTGGATGG CTCGGCTCGG GGCGCTGACG AAGGG
J23SpTT_23                    CCCGCA GAAGACGGGT TTGATGGGGC GGGGATGTAA GCCGGGAAGG AAACCGACCC GTTTAGTCTG CCGCTCCCAA CAGCCCGAGT TTCTTGCCCC TTGAGTAAGG GGCAAGAAAC CTAAGCCGCC TGGTGGATGG CTCGGCTCGG GGCGCTGACG AAGGGCGT
J23SpTT_21                   GCCCGCA GAAGACGGGT TTGATGGGGC GGGGATGTAA GCCGGGAAGG AAACCGACCC GTTTAGTCTG CCGCTCCCAA CAGCCCGAGT TTCTTGCCCC TTGAGTAAGG GGCAAGAAAC CTAAGCCGCC TGGTGGATGG CTCGGCTCGG GGCGCTGACG AAGGGC
J23SpTT_894                    CCGCA GAAGACGGGT TTGATGGGGC GGGGATGTAA GCCGGGAAGG AAACCGACCC GTTTAGTCTG CCGCTCCCAA CAGCCCGAGT TTCTTGCCCC TTGAGTAAGG GGCAAGAAAC CTAAGCCGCC TGGTGGATGG CTCGGCTCGG GGCGCTGACG AAGGGCG
J23SpTT_13                     CCGCA GAAGACGGGT TTGATGGGGC GGGGATGTAA GCCGGGAAGG AAACCGACCC GTTTAGTCTG CCGCTCCCAA CAGCCCGAGT TTCTTGCCCC TTGAGTAAGG GGCAAGAAAC CTAAGCCGCC TGGTGGATGG CTCGGCTCGG GGCGCTGACG AAGGGCGG
J23SpTT_11                    CCCGCA GAAGACGGGT TTGATGGGGC GGGGATGTAA GCCGGGAAGG AAACCGACCC GTTTAGTCTG CCGCTCCCAA CAGCCCGAGT TTCTTGCCCC TTGAGTAAGG GGCAAGAAAC CTAAGCCGCC TGGTGGATGG CTCGGCTCGG GGCGCTGACG AAGGGCA
J23SpTT_12                     CCGCA GAAGACGGGT TTGATGGGGC GGGGATGTAA GCCGGGAAGG AAACCGACCC GTTTAGTCTG CCGCTCCCAA CAGCCCGAGT TTCTTGCCCC TTGAGTAAGG GGCAAGAAAC CTAAGCCGCC TGGTGGATGG CTCGGCTCGG GGCGCTGACG AAGGGCGT
J23SpTT_34                   GCCCGCA GAAGACGGGT TTGATGGGGC GGGGATGTAA GCCGGGAAGG AAACCGACCC GTTTAGTCTG CCGCTCCCAA CAGCCCGAGT TTCTTGCCCC TTGAGTAAGG GGCAAGAAAC CTAAGCCGCC TGGTGGATGG CTCGGCTCGG GGCGCTGACG AAGGG
J23SpTT_23                    CCCGCA GAAGACGGGT TTGATGGGGC GGGGATGTAA GCCGGGAAGG AAACCGACCC GTTTAGTCTG CCGCTCCCAA CAGCCCGAGT TTCTTGCCCC TTGAGTAAGG GGCAAGAAAC CTAAGCCGCC TGGTGGATGG CTCGGCTCGG GGCGCTGACG AAGGGCC
J23SpTT_21                  CGCCCGCA GAAGACGGGT TTGATGGGGC GGGGATGTAA GCCGGGAAGG AAACCGACCC GTTTAGTCTG CCGCTCCCAA CAGCCCGAGT TTCTTGCCCC TTGAGTAAGG GGCAAGAAAC CTAAGCCGCC TGGTGGATGG CTCGGCTCGG GGCGCTGACG AAGG
J23SpTT_10                    CCCGCA GAAGACGGGT TTGATGGGGC GGGGATGTAA GCCGGGAAGG AAACCGACCC GTTTAGTCTG CCGCTCCCAA CAGCCCGAGT TTCTTGCCCC TTGAGTAAGG GGCAAGAAAC CTAAGCCGCC TGGTGGATGG CTCGGCTCGG GGCGCTGACG AAGGGCT
J23SpTT_1950                    CGCA GAAGACGGGT TTGATGGGGC GGGGATGTAA GCCGGGAAGG AAACCGACCC GTTTAGTCTG CCGCTCCCAA CAGCCCGAGT TTCTTGCCCC TTGAGTAAGG GGCAAGAAAC CTAAGCCGCC TGGTGGATGG CTCGGCTCGG GGCGCTGACG AAGGGCG
J23SpTT_1834                   CCGCA GAAGACGGGT TTGATGGGGC GGGGATGTAA GCCGGGAAGG AAACCGACCC GTTTAGTCTG CCGCTCCCAA CAGCCCGAGT TTCTTGCCCC TTGAGTAAGG GGCAAGAAAC CTAAGCCGCC TGGTGGATGG CTCGGCTCGG GGCGCTGACG AAGGGC
J23SpTT_1396                    CGCA GAAGACGGGT TTGATGGGGC GGGGATGTAA GCCGGGAAGG AAACCGACCC GTTTAGTCTG CCGCTCCCAA CAGCCCGAGT TTCTTGCCCC TTGAGTAAGG GGCAAGAAAC CTAAGCCGCC TGGTGGATGG CTCGGCTCGG GGCGCTGACG AAGGGCGTGG
J23SpTT_37                      CGCA GAAGACGGGT TTGATGGGGC GGGGATGTAA GCCGGGAAGG AAACCGACCC GTTTAGTCTG CCGCTCCCAA CAGCCCGAGT TTCTTGCCCC TTGAGTAAGG GGCAAGAAAC CTAAGCCGCC TGGTGGATGG CTCGGCTCGG GGCGCTGACG AAGGGCGTGG CG
J23SpTT_39                       GCA GAAGACGGGT TTGATGGGGC GGGGATGTAA GCCGGGAAGG AAACCGACCC GTTTAGTCTG CCGCTCCCAA CAGCCCGAGT TTCTTGCCCC TTGAGTAAGG GGCAAGAAAC CTAAGCCGCC TGGTGGATGG CTCGGCTCGG GGCGCTGACG AAGGGCGTGG C
J23SpTT_21                      CGCA GAAGACGGGT TTGATGGGGC GGGGATGTAA GCCGGGAAGG AAACCGACCC GTTTAGTCTG CCGCTCCCAA CAGCCCGAGT TTCTTGCCCC TTGAGTAAGG GGCAAGAAAC CTAAGCCGCC TGGTGGATGG CTCGGCTCGG GGCGCTGACG AAGGGCGTGA C
J23SpTT_11                     CCGCA GAAGACGGGT TTGATGGGGC GGGGATGTAA GCCGGGAAGG AAACCGACCC GTTTAGTCTG CCGCTCCCAA CAGCCCGAGT TTCTTGCCCC TTGAGTAAGG GGCAAGAAAC CTAAGCCGCC TGGTGGATGG CTCGGCTCGG GGCGCTGACG AAGGGCGTGA C
J23SpTT_10                      CGCA GAAGACGGGT TTGATGGGGC GGGGATGTAA GCCGGGAAGG AAACCGACCC GTTTAGTCTG CCGCTCCCAA CAGCCCGAGT TTCTTGCCCC TTGAGTAAGG GGCAAGAAAC CTAAGCCGCC TGGTGGATGG CTCGGCTCGG GGCGCTGACG AAGGGCGTGG G
J23SpTT_823                     CGCA GAAGACGGGT TTGATGGGGC GGGGATGTAA GCCGGGAAGG AAACCGACCC GTTTAGTCTG CCGCTCCCAA CAGCCCGAGT TTCTTGCCCC TTGAGTAAGG GGCAAGAAAC CTAAGCCGCC TGGTGGATGG CTCGGCTCGG GGCGCTGACG AAGGGCGTG
J23SpTT_357                    CCGCA GAAGACGGGT TTGATGGGGC GGGGATGTAA GCCGGGAAGG AAACCGACCC GTTTAGTCTG CCGCTCCCAA CAGCCCGAGT TTCTTGCCCC TTGAGTAAGG GGCAAGAAAC CTAAGCCGCC TGGTGGATGG CTCGGCTCGG GGCGCTGACG AAGGGCGTG
J23SpTT_147                      GCA GAAGACGGGT TTGATGGGGC GGGGATGTAA GCCGGGAAGG AAACCGACCC GTTTAGTCTG CCGCTCCCAA CAGCCCGAGT TTCTTGCCCC TTGAGTAAGG GGCAAGAAAC CTAAGCCGCC TGGTGGATGG CTCGGCTCGG GGCGCTGACG AAGGGCGTGG
J23SpTT_107                       CA GAAGACGGGT TTGATGGGGC GGGGATGTAA GCCGGGAAGG AAACCGACCC GTTTAGTCTG CCGCTCCCAA CAGCCCGAGT TTCTTGCCCC TTGAGTAAGG GGCAAGAAAC CTAAGCCGCC TGGTGGATGG CTCGGCTCGG GGCGCTGACG AAGGGCGTGG C
J23SpTT_11                    CCCGCA GAAGACGGGT TTGATGGGGC GGGGATGTAA GCCGGGAAGG AAACCGACCC GTTTAGTCTG CCGCTCCCAA CAGCCCGAGT TTCTTGCCCC TTGAGTAAGG GGCAAGAAAC CTAAGCCGCC TGGTGGATGG CTCGGCTCGG GGCGCTGACG AAGGGCATGG
J23SpTT_9                       CGCA GAAGACGGGT TTGATGGGGC GGGGATGTAA GCCGGGAAGG AAACCGACCC GTTTAGTCTG CCGCTCCCAA CAGCCCGAGT TTCTTGCCCC TTGAGTAAGG GGCAAGAAAC CTAAGCCGCC TGGTGGATGG CTCGGCTCGG GGCGCTGACG AAGGGCATGG
J23SpTT_36                      CGCA GAAGACGGGT TTGATGGGGC GGGGATGTAA GCCGGGAAGG AAACCGACCC GTTTAGTCTG CCGCTCCCAA CAGCCCGAGT TTCTTGCCCC TTGAGTAAGG GGCAAGAAAC CTAAGCCGCC TGGTGGATGG CTCGGCTCGG GGCGCTGACG AAGGGCGTAG
J23SpTT_16                    CCCGCA GAAGACGGGT TTGATGGGGC GGGGATGTAA GCCGGGAAGG AAACCGACCC GTTTAGTCTG CCGCTCCCAA CAGCCCGAGT TTCTTGCCCC TTGAGTAAGG GGCAAGAAAC CTAAGCCGCC TGGTGGATGG CTCGGCTCGG GGCGCTGACG AAGGGCGTAG
J23SpTT_11                     CCGCA GAAGACGGGT TTGATGGGGC GGGGATGTAA GCCGGGAAGG AAACCGACCC GTTTAGTCTG CCGCTCCCAA CAGCCCGAGT TTCTTGCCCC TTGAGTAAGG GGCAAGAAAC CTAAGCCGCC TGGTGGATGG CTCGGCTCGG GGCGCTGACG AAGGGCGTAG
J23SpTT_21                        CA GAAGACGGGT TTGATGGGGC GGGGATGTAA GCCGGGAAGG AAACCGACCC GTTTAGTCTG CCGCTCCCAA CAGCCCGAGT TTCTTGCCCC TTGAGTAAGG GGCAAGAAAC CTAAGCCGCC TGGTGGATGG CTCGGCTCGG GGCGCTGACG AAGGGCGTGG CG
J23SpTT_20                     CCGCA GAAGACGGGT TTGATGGGGC GGGGATGTAA GCCGGGAAGG AAACCGACCC GTTTAGTCTG CCGCTCCCAA CAGCCCGAGT TTCTTGCCCC TTGAGTAAGG GGCAAGAAAC CTAAGCCGCC TGGTGGATGG CTCGGCTCGG GGCGCTGACG AAGGGCGGGG
J23SpTT_13                      CGCA GAAGACGGGT TTGATGGGGC GGGGATGTAA GCCGGGAAGG AAACCGACCC GTTTAGTCTG CCGCTCCCAA CAGCCCGAGT TTCTTGCCCC TTGAGTAAGG GGCAAGAAAC CTAAGCCGCC TGGTGGATGG CTCGGCTCGG GGCGCTGACG AAGGGCGGGG
J23SpTT_11                    CCCGCA GAAGACGGGT TTGATGGGGC GGGGATGTAA GCCGGGAAGG AAACCGACCC GTTTAGTCTG CCGCTCCCAA CAGCCCGAGT TTCTTGCCCC TTGAGTAAGG GGCAAGAAAC CTAAGCCGCC TGGTGGATGG CTCGGCTCGG GGCGCTGACG AAGGGCGGG
J23SpTT_14                        CA GAAGACGGGT TTGATGGGGC GGGGATGTAA GCCGGGAAGG AAACCGACCC GTTTAGTCTG CCGCTCCCAA CAGCCCGAGT TTCTTGCCCC TTGAGTAAGG GGCAAGAAAC CTAAGCCGCC TGGTGGATGG CTCGGCTCGG GGCGCTGACG AAGGGCGTGG CA
J23SpTT_553                       CA GAAGACGGGT TTGATGGGGC GGGGATGTAA GCCGGGAAGG AAACCGACCC GTTTAGTCTG CCGCTCCCAA CAGCCCGAGT TTCTTGCCCC TTGAGTAAGG GGCAAGAAAC CTAAGCCGCC TGGTGGATGG CTCGGCTCGG GGCGCTGACG AAGGGCGTGG
J23SpTT_8                       NGCA GAAGACGGGT TTGATGGGGC GGGGATGTAA GCCGGGAAGG AAACCGACCC GTTTAGTCTG CCGCTCCCAA CAGCCCGAGT TTCTTGCCCC TTGAGTAAGG GGCAAGAAAC CTAAGCCGCC TGGTGGATGG CTCGGCTCGG GGCGCTGACG AAGGGCGTGN
J23SpTT_97                       GCA GAAGACGGGT TTGATGGGGC GGGGATGTAA GCCGGGAAGG AAACCGACCC GTTTAGTCTG CCGCTCCCAA CAGCCCGAGT TTCTTGCCCC TTGAGTAAGG GGCAAGAAAC CTAAGCCGCC TGGTGGATGG CTCGGCTCGG GGCGCTGACG AAGGGCGTG
J23SpTT_55                         A GAAGACGGGT TTGATGGGGC GGGGATGTAA GCCGGGAAGG AAACCGACCC GTTTAGTCTG CCGCTCCCAA CAGCCCGAGT TTCTTGCCCC TTGAGTAAGG GGCAAGAAAC CTAAGCCGCC TGGTGGATGG CTCGGCTCGG GGCGCTGACG AAGGGCGTGG C
J23SpTT_35                      CGCA GAAGACGGGT TTGATGGGGC GGGGATGTAA GCCGGGAAGG AAACCGACCC GTTTAGTCTG CCGCTCCCAA CAGCCCGAGT TTCTTGCCCC TTGAGTAAGG GGCAAGAAAC CTAAGCCGCC TGGTGGATGG CTCGGCTCGG GGCGCTGACG AAGGGCGT
J23SpTT_23                         A GAAGACGGGT TTGATGGGGC GGGGATGTAA GCCGGGAAGG AAACCGACCC GTTTAGTCTG CCGCTCCCAA CAGCCCGAGT TTCTTGCCCC TTGAGTAAGG GGCAAGAAAC CTAAGCCGCC TGGTGGATGG CTCGGCTCGG GGCGCTGACG AAGGGCGTGG CAAG
J23SpTT_21                      CGCA GAAGACGGGT TTGATGGGGC GGGGATGTAA GCCGGGAAGG AAACCGACCC GTTTAGTCTG CCGCTCCCAA CAGCCCGAGT TTCTTGCCCC TTGAGTAAGG GGCAAGAAAC CTAAGCCGCC TGGTGGATGG CTCGGCTCGG GGCGCTGACG AAGGGCGGG
J23SpTT_14                      CGCA GAAGACGGGT TTGATGGGGC GGGGATGTAA GCCGGGAAGG AAACCGACCC GTTTAGTCTG CCGCTCCCAA CAGCCCGAGT TTCTTGCCCC TTGAGTAAGG GGCAAGAAAC CTAAGCCGCC TGGTGGATGG CTCGGCTCGG GGCGCTGACG AAGGGCGG
J23SpTT_10                      CGCA GAAGACGGGT TTGATGGGGC GGGGATGTAA GCCGGGAAGG AAACCGACCC GTTTAGTCTG CCGCTCCCAA CAGCCCGAGT TTCTTGCCCC TTGAGTAAGG GGCAAGAAAC CTAAGCCGCC TGGTGGATGG CTCGGCTCGG GGCGCTGACG AAGGGCGC
J23SpTT_328                       CA GAAGACGGGT TTGATGGGGC GGGGATGTAA GCCGGGAAGG AAACCGACCC GTTTAGTCTG CCGCTCCCAA CAGCCCGAGT TTCTTGCCCC TTGAGTAAGG GGCAAGAAAC CTAAGCCGCC TGGTGGATGG CTCGGCTCGG GGCGCTGACG AAGGGCGTG
J23SpTT_204                      GCA GAAGACGGGT TTGATGGGGC GGGGATGTAA GCCGGGAAGG AAACCGACCC GTTTAGTCTG CCGCTCCCAA CAGCCCGAGT TTCTTGCCCC TTGAGTAAGG GGCAAGAAAC CTAAGCCGCC TGGTGGATGG CTCGGCTCGG GGCGCTGACG AAGGGCG
J23SpTT_204                        A GAAGACGGGT TTGATGGGGC GGGGATGTAA GCCGGGAAGG AAACCGACCC GTTTAGTCTG CCGCTCCCAA CAGCCCGAGT TTCTTGCCCC TTGAGTAAGG GGCAAGAAAC CTAAGCCGCC TGGTGGATGG CTCGGCTCGG GGCGCTGACG AAGGGCGTGG
J23SpTT_108                    CCGCA GAAGACGGGT TTGATGGGGC GGGGATGTAA GCCGGGAAGG AAACCGACCC GTTTAGTCTG CCGCTCCCAA CAGCCCGAGT TTCTTGCCCC TTGAGTAAGG GGCAAGAAAC CTAAGCCGCC TGGTGGATGG CTCGGCTCGG GGCGCTGACG AAGGGG
J23SpTT_83                    CCCGCA GAAGACGGGT TTGATGGGGC GGGGATGTAA GCCGGGAAGG AAACCGACCC GTTTAGTCTG CCGCTCCCAA CAGCCCGAGT TTCTTGCCCC TTGAGTAAGG GGCAAGAAAC CTAAGCCGCC TGGTGGATGG CTCGGCTCGG GGCGCTGACG AAGGGG
J23SpTT_9                     CCCGCA GAAGACGGGT TTGATGGGGC GGGGATGTAA GCCGGGAAGG AAACCGACCC GTTTAGTCTG CCGCTCCCAA CAGCCCGAGT TTCTTGCCCC TTGAGTAAGG GGCAAGAAAC CTAAGCCGCC TGGTGGATGG CTCGGCTCGG GGCGCTGACG AAGGGT
J23SpTT_14                      CGCA GAAGACGGGT TTGATGGGGC GGGGATGTAA GCCGGGAAGG AAACCGACCC GTTTAGTCTG CCGCTCCCAA CAGCCCGAGT TTCTTGCCCC TTGAGTAAGG GGCAAGAAAC CTAAGCCGCC TGGTGGATGG CTCGGCTCGG GGCGCTGACG AAGAGCG
J23SpTT_14                      CGCA GAAGACGGGT TTGATGGGGC GGGGATGTAA GCCGGGAAGG AAACCGACCC GTTTAGTCTG CCGCTCCCAA CAGCCCGAGT TTCTTGCCCC TTGAGTAAGG GGCAAGAAAC CTAAGCCGCC TGGTGGATGG CTCGGCTCGG GGCGCTGACG AAAGGCG
J23SpTT_50                      CGCA GAAGACGGGT TTGATGGGGC GGGGATGTAA GCCGGGAAGG AAACCGACCC GTTTAGTCTG CCGCTCCCAA CAGCCCGAGT TTCTTGCCCC TTGAGTAAGG GGCAAGAAAC CTAAGCCGCC TGGTGGATGG CTCGGCTCGG GGCGCTGACG AAGGGCC
J23SpTT_28                     CCGCA GAAGACGGGT TTGATGGGGC GGGGATGTAA GCCGGGAAGG AAACCGACCC GTTTAGTCTG CCGCTCCCAA CAGCCCGAGT TTCTTGCCCC TTGAGTAAGG GGCAAGAAAC CTAAGCCGCC TGGTGGATGG CTCGGCTCGG GGCGCTGACG AAGGGCC
J23SpTT_17                     CCGCA GAAGACGGGT TTGATGGGGC GGGGATGTAA GCCGGGAAGG AAACCGACCC GTTTAGTCTG CCGCTCCCAA CAGCCCGAGT TTCTTGCCCC TTGAGTAAGG GGCAAGAAAC CTAAGCCGCC TGGTGGATGG CTCGGCTCGG GGCGCTGACG AAGGGCT
J23SpTT_19                     CCGCA GAAGACGGGT TTGATGGGGC GGGGATGTAA GCCGGGAAGG AAACCGACCC GTTTAGTCTG CCGCTCCCAA CAGCCCGAGT TTCTTGCCCC TTGAGTAAGG GGCAAGAAAC CTAAGCCGCC TGGTGGATGG CTCGGCTCGG GGCGCTGACG AAGGGCA
J23SpTT_13                      CGCA GAAGACGGGT TTGATGGGGC GGGGATGTAA GCCGGGAAGG AAACCGACCC GTTTAGTCTG CCGCTCCCAA CAGCCCGAGT TTCTTGCCCC TTGAGTAAGG GGCAAGAAAC CTAAGCCGCC TGGTGGATGG CTCGGCTCGG GGCGCTGACG AAGGGAG
J23SpTT_11                        CA GAAGACGGGT TTGATGGGGC GGGGATGTAA GCCGGGAAGG AAACCGACCC GTTTAGTCTG CCGCTCCCAA CAGCCCGAGT TTCTTGCCCC TTGAGTAAGG GGCAAGAAAC CTAAGCCGCC TGGTGGATGG CTCGGCTCGG GGCGCTGACG AAGGGCGT
J23SpTT_27                      CGCA GAAGACGGGT TTGATGGGGC GGGGATGTAA GCCGGGAAGG AAACCGACCC GTTTAGTCTG CCGCTCCCAA CAGCCCGAGT TTCTTGCCCC TTGAGTAAGG GGCAAGAAAC CTAAGCCGCC TGGTGGATGG CTCGGCTCGG GGCGCTGACG AAGGGCA
J23SpTT_13                        CA GAAGACGGGT TTGATGGGGC GGGGATGTAA GCCGGGAAGG AAACCGACCC GTTTAGTCTG CCGCTCCCAA CAGCCCGAGT TTCTTGCCCC TTGAGTAAGG GGCAAGAAAC CTAAGCCGCC TGGTGGATGG CTCGGCTCGG GGCGCTGACG AAGGGCGGGG
J23SpTT_26                      CGCA GAAGACGGGT TTGATGGGGC GGGGATGTAA GCCGGGAAGG AAACCGACCC GTTTAGTCTG CCGCTCCCAA CAGCCCGAGT TTCTTGCCCC TTGAGTAAGG GGCAAGAAAC CTAAGCCGCC TGGTGGATGG CTCGGCTCGG GGCGCTGACG AAGGGCT
J23SpTT_21                    CCCGCA GAAGACGGGT TTGATGGGGC GGGGATGTAA GCCGGGAAGG AAACCGACCC GTTTAGTCTG CCGCTCCCAA CAGCCCGAGT TTCTTGCCCC TTGAGTAAGG GGCAAGAAAC CTAAGCCGCC TGGTGGATGG CTCGGCTCGG GGCGCTGACG AAGGAC
J23SpTT_14                     CCGCA GAAGACGGGT TTGATGGGGC GGGGATGTAA GCCGGGAAGG AAACCGACCC GTTTAGTCTG CCGCTCCCAA CAGCCCGAGT TTCTTGCCCC TTGAGTAAGG GGCAAGAAAC CTAAGCCGCC TGGTGGATGG CTCGGCTCGG GGCGCTGACG AAGGAC
J23SpTT_12                     CCGCA GAAGACGGGT TTGATGGGGC GGGGATGTAA GCCGGGAAGG AAACCGACCC GTTTAGTCTG CCGCTCCCAA CAGCCCGAGT TTCTTGCCCC TTGAGTAAGG GGCAAGAAAC CTAAGCCGCC TGGTGGATGG CTCGGCTCGG GGCGCTGACG AAGGACG
J23SpTT_11                      CGCA GAAGACGGGT TTGATGGGGC GGGGATGTAA GCCGGGAAGG AAACCGACCC GTTTAGTCTG CCGCTCCCAA CAGCCCGAGT TTCTTGCCCC TTGAGTAAGG GGCAAGAAAC CTAAGCCGCC TGGTGGATGG CTCGGCTCGG GGCGCTGACG AAGGACG
J23SpTT_19                        CA GAAGACGGGT TTGATGGGGC GGGGATGTAA GCCGGGAAGG AAACCGACCC GTTTAGTCTG CCGCTCCCAA CAGCCCGAGT TTCTTGCCCC TTGAGTAAGG GGCAAGAAAC CTAAGCCGCC TGGTGGATGG CTCGGCTCGG GGCGCTGACG AAGGGCGTAG
J23SpTT_11                      CGCA GAAGACGGGT TTGATGGGGC GGGGATGTAA GCCGGGAAGG AAACCGACCC GTTTAGTCTG CCGCTCCCAA CAGCCCGAGT TTCTTGCCCC TTGAGTAAGG GGCAAGAAAC CTAAGCCGCC TGGTGGATGG CTCGGCTCGG GGCGCTGACG AAGGGTG
J23SpTT_3141                   CCGCA GAAGACGGGT TTGATGGGGC GGGGATGTAA GCCGGGAAGG AAACCGACCC GTTTAGTCTG CCGCTCCCAA CAGCCCGAGT TTCTTGCCCC TTGAGTAAGG GGCAAGAAAC CTAAGCCGCC TGGTGGATGG CTCGGCTCGG GGCGCTGACG AAGGG
J23SpTT_8                     NCCGCA GAAGACGGGT TTGATGGGGC GGGGATGTAA GCCGGGAAGG AAACCGACCC GTTTAGTCTG CCGCTCCCAA CAGCCCGAGT TTCTTGCCCC TTGAGTAAGG GGCAAGAAAC CTAAGCCGCC TGGTGGATGG CTCGGCTCGG GGCGCTGACG AAGGGN
J23SpTT_13                    NCCGCA GAAGACGGGT TTGATGGGGC GGGGATGTAA GCCGGGAAGG AAACCGACCC GTTTAGTCTG CCGCTCCCAA CAGCCCGAGT TTCTTGCCCC TTGAGTAAGG GGCAAGAAAC CTAAGCCGCC TGGTGGATGG CTCGGCTCGG GGCGCTGACG AAGGN
J23SpTT_11                      CGCA GAAGACGGGT TTGATGGGGC GGGGATGTAA GCCGGGAAGG AAACCGACCC GTTTAGTCTG CCGCTCCCAA CAGCCCGAGT TTCTTGCCCC TTGAGTAAGG GGCAAGAAAC CTAAGCCGCC TGGTGGATGG CTCGGCTCGG GGCGCTGACG AAGAGC
J23SpTT_1796                  CCCGCA GAAGACGGGT TTGATGGGGC GGGGATGTAA GCCGGGAAGG AAACCGACCC GTTTAGTCTG CCGCTCCCAA CAGCCCGAGT TTCTTGCCCC TTGAGTAAGG GGCAAGAAAC CTAAGCCGCC TGGTGGATGG CTCGGCTCGG GGCGCTGACG AAGG
J23SpTT_29                   GCCCGCA GAAGACGGGT TTGATGGGGC GGGGATGTAA GCCGGGAAGG AAACCGACCC GTTTAGTCTG CCGCTCCCAA CAGCCCGAGT TTCTTGCCCC TTGAGTAAGG GGCAAGAAAC CTAAGCCGCC TGGTGGATGG CTCGGCTCGG GGCGCTGACG AAGG
J23SpTT_21                  CGCCCGCA GAAGACGGGT TTGATGGGGC GGGGATGTAA GCCGGGAAGG AAACCGACCC GTTTAGTCTG CCGCTCCCAA CAGCCCGAGT TTCTTGCCCC TTGAGTAAGG GGCAAGAAAC CTAAGCCGCC TGGTGGATGG CTCGGCTCGG GGCGCTGACG AAG
J23SpTT_17                   GCCCGCA GAAGACGGGT TTGATGGGGC GGGGATGTAA GCCGGGAAGG AAACCGACCC GTTTAGTCTG CCGCTCCCAA CAGCCCGAGT TTCTTGCCCC TTGAGTAAGG GGCAAGAAAC CTAAGCCGCC TGGTGGATGG CTCGGCTCGG GGCGCTGACG AAG
J23SpTT_693                    CCGCA GAAGACGGGT TTGATGGGGC GGGGATGTAA GCCGGGAAGG AAACCGACCC GTTTAGTCTG CCGCTCCCAA CAGCCCGAGT TTCTTGCCCC TTGAGTAAGG GGCAAGAAAC CTAAGCCGCC TGGTGGATGG CTCGGCTCGG GGCGCTGACG AAGGGCGTGG
J23SpTT_551                   CCCGCA GAAGACGGGT TTGATGGGGC GGGGATGTAA GCCGGGAAGG AAACCGACCC GTTTAGTCTG CCGCTCCCAA CAGCCCGAGT TTCTTGCCCC TTGAGTAAGG GGCAAGAAAC CTAAGCCGCC TGGTGGATGG CTCGGCTCGG GGCGCTGACG AAGGGCGTGG
J23SpTT_21                     CCGCA GAAGACGGGT TTGATGGGGC GGGGATGTAA GCCGGGAAGG AAACCGACCC GTTTAGTCTG CCGCTCCCAA CAGCCCGAGT TTCTTGCCCC TTGAGTAAGG GGCAAGAAAC CTAAGCCGCC TGGTGGATGG CTCGGCTCGG GGCGCTGACG AAGGGCGTGG CG
J23SpTT_19                    CCCGCA GAAGACGGGT TTGATGGGGC GGGGATGTAA GCCGGGAAGG AAACCGACCC GTTTAGTCTG CCGCTCCCAA CAGCCCGAGT TTCTTGCCCC TTGAGTAAGG GGCAAGAAAC CTAAGCCGCC TGGTGGATGG CTCGGCTCGG GGCGCTGACG AAGGGCGTGG CG
J23SpTT_12                   GCCCGCA GAAGACGGGT TTGATGGGGC GGGGATGTAA GCCGGGAAGG AAACCGACCC GTTTAGTCTG CCGCTCCCAA CAGCCCGAGT TTCTTGCCCC TTGAGTAAGG GGCAAGAAAC CTAAGCCGCC TGGTGGATGG CTCGGCTCGG GGCGCTGACG AAGGGCGTGG
J23SpTT_360                   CCCGCA GAAGACGGGT TTGATGGGGC GGGGATGTAA GCCGGGAAGG AAACCGACCC GTTTAGTCTG CCGCTCCCAA CAGCCCGAGT TTCTTGCCCC TTGAGTAAGG GGCAAGAAAC CTAAGCCGCC TGGTGGATGG CTCGGCTCGG GGCGCTGACG AAGGGCGTG
J23SpTT_333                     CGCA GAAGACGGGT TTGATGGGGC GGGGATGTAA GCCGGGAAGG AAACCGACCC GTTTAGTCTG CCGCTCCCAA CAGCCCGAGT TTCTTGCCCC TTGAGTAAGG GGCAAGAAAC CTAAGCCGCC TGGTGGATGG CTCGGCTCGG GGCGCTGACG AAGGGCGTGG C
J23SpTT_166                    CCGCA GAAGACGGGT TTGATGGGGC GGGGATGTAA GCCGGGAAGG AAACCGACCC GTTTAGTCTG CCGCTCCCAA CAGCCCGAGT TTCTTGCCCC TTGAGTAAGG GGCAAGAAAC CTAAGCCGCC TGGTGGATGG CTCGGCTCGG GGCGCTGACG AAGGGCGTGG C
J23SpTT_12                     CCGCA GAAGACGGGT TTGATGGGGC GGGGATGTAA GCCGGGAAGG AAACCGACCC GTTTAGTCTG CCGCTCCCAA CAGCCCGAGT TTCTTGCCCC TTGAGTAAGG GGCAAGAAAC CTAAGCCGCC TGGTGGATGG CTCGGCTCGG GGCGCTGACG AAGGGCGTGG CA
J23SpTT_122                     CGCA GAAGACGGGT TTGATGGGGC GGGGATGTAA GCCGGGAAGG AAACCGACCC GTTTAGTCTG CCGCTCCCAA CAGCCCGAGT TTCTTGCCCC TTGAGTAAGG GGCAAGAAAC CTAAGCCGCC TGGTGGATGG CTCGGCTCGG GGCGCTGACG AAGGGCGTGG CAAG
J23SpTT_69                     CCGCA GAAGACGGGT TTGATGGGGC GGGGATGTAA GCCGGGAAGG AAACCGACCC GTTTAGTCTG CCGCTCCCAA CAGCCCGAGT TTCTTGCCCC TTGAGTAAGG GGCAAGAAAC CTAAGCCGCC TGGTGGATGG CTCGGCTCGG GGCGCTGACG AAGGGCGTGG CAAG
J23SpTT_57                    CCCGCA GAAGACGGGT TTGATGGGGC GGGGATGTAA GCCGGGAAGG AAACCGACCC GTTTAGTCTG CCGCTCCCAA CAGCCCGAGT TTCTTGCCCC TTGAGTAAGG GGCAAGAAAC CTAAGCCGCC TGGTGGATGG CTCGGCTCGG GGCGCTGACG AAGGGCGTGG CAAG
J23SpTT_11                    CCCGCA GAAGACGGGT TTGATGGGGC GGGGATGTAA GCCGGGAAGG AAACCGACCC GTTTAGTCTG CCGCTCCCAA CAGCCCGAGT TTCTTGCCCC TTGAGTAAGG GGCAAGAAAC CTAAGCCGCC TGGTGGATGG CTCGGCTCGG GGCGCTGACG AAGGGCGTGG CAAGC
```

```
J23SpTT_19   CCGCA GAAGACGGGT TTGATGGGGC GGGGATGTAA GCCGGGAAGGG AAACCGACCC GTTTAGTCTG CCGCTCCCAA CAGCCCGAGT TTCTTGCCCC TTGAGTAAGG GGCAAGAAAC CTAAGCCGCC TGGTGGATGG CTCGGCTCGG GGCGCTGACG AAGGGCGTGG CAAGC
J23SpTT_42   CGCA GAAGACGGGT TTGATGGGGC GGGGATGTAA GCCGGGAAGGG AAACCGACCC GTTTAGTCTG CCGCTCCCAA CAGCCCGAGT TTCTTGCCCC TTGAGTAAGG GGCAAGAAAC CTAAGCCGCC TGGTGGATGG CTCGGCTCGG GGCGCTGACG AAGGGCGTGG CAAGC
J23SpTT_25   CGCA GAAGACGGGT TTGATGGGGC GGGGATGTAA GCCGGGAAGGG AAACCGACCC GTTTAGTCTG CCGCTCCCAA CAGCCCGAGT TTCTTGCCCC TTGAGTAAGG GGCAAGAAAC CTAAGCCGCC TGGTGGATGG CTCGGCTCGG GGCGCTGACG AAGGGCGTGG CAAGCT
J23SpTT_12   CCCGCA GAAGACGGGT TTGATGGGGC GGGGATGTAA GCCGGGAAGGG AAACCGACCC GTTTAGTCTG CCGCTCCCAA CAGCCCGAGT TTCTTGCCCC TTGAGTAAGG GGCAAGAAAC CTAAGCCGCC TGGTGGATGG CTCGGCTCGG GGCGCTGACG AAGGGCGTGG CA
J23SpTT_104  CCCGCA GAAGACGGGT TTGATGGGGC GGGGATGTAA GCCGGGAAGGG AAACCGACCC GTTTAGTCTG CCGCTCCCAA CAGCCCGAGT TTCTTGCCCC TTGAGTAAGG GGCAAGAAAC CTAAGCCGCC TGGTGGATGG CTCGGCTCGG GGCGCTGACG AAGGGCGTGG C
J23SpTT_53   A GAAGACGGGT TTGATGGGGC GGGGATGTAA GCCGGGAAGGG AAACCGACCC GTTTAGTCTG CCGCTCCCAA CAGCCCGAGT TTCTTGCCCC TTGAGTAAGG GGCAAGAAAC CTAAGCCGCC TGGTGGATGG CTCGGCTCGG GGCGCTGACG AAGGGCGTGG CAAGCTGCG
J23SpTT_9    CA GAAGACGGGT TTGATGGGGC GGGGATGTAA GCCGGGAAGGG AAACCGACCC GTTTAGTCTG CCGCTCCCAA CAGCCCGAGT TTCTTGCCCC TTGAGTAAGG GGCAAGAAAC CTAAGCCGCC TGGTGGATGG CTCGGCTCGG GGCGCTGACG AAGGGCGTGG CAAGCTGC
J23SpTT_42   CA GAAGACGGGT TTGATGGGGC GGGGATGTAA GCCGGGAAGGG AAACCGACCC GTTTAGTCTG CCGCTCCCAA CAGCCCGAGT TTCTTGCCCC TTGAGTAAGG GGCAAGAAAC CTAAGCCGCC TGGTGGATGG CTCGGCTCGG GGCGCTGACG AAGGGCGTGG CAAGCTG
J23SpTT_11   A GAAGACGGGT TTGATGGGGC GGGGATGTAA GCCGGGAAGGG AAACCGACCC GTTTAGTCTG CCGCTCCCAA CAGCCCGAGT TTCTTGCCCC TTGAGTAAGG GGCAAGAAAC CTAAGCCGCC TGGTGGATGG CTCGGCTCGG GGCGCTGACG AAGGGCGTGG CAAGCTGC
J23SpTT_11   CA GAAGACGGGT TTGATGGGGC GGGGATGTAA GCCGGGAAGGG AAACCGACCC GTTTAGTCTG CCGCTCCCAA CAGCCCGAGT TTCTTGCCCC TTGAGTAAGG GGCAAGAAAC CTAAGCCGCC TGGTGGATGG CTCGGCTCGG GGCGCTGACG AAGGGCGTGG CAAGCT
J23SpTT_13   A GAAGACGGGT TTGATGGGGC GGGGATGTAA GCCGGGAAGGG AAACCGACCC GTTTAGTCTG CCGCTCCCAA CAGCCCGAGT TTCTTGCCCC TTGAGTAAGG GGCAAGAAAC CTAAGCCGCC TGGTGGATGG CTCGGCTCGG GGCGCTGACG AAGGGCGTGG CAAGCTG
J23SpTT_10   CA GAAGACGGGT TTGATGGGGC GGGGATGTAA GCCGGGAAGGG AAACCGACCC GTTTAGTCTG CCGCTCCCAA CAGCCCGAGT TTCTTGCCCC TTGAGTAAGG GGCAAGAAAC CTAAGCCGCC TGGTGGATGG CTCGGCTCGG GGCGCTGACG AAGGGCGTGG CAAGC
J23SpTT_12   GCA GAAGACGGGT TTGATGGGGC GGGGATGTAA GCCGGGAAGGG AAACCGACCC GTTTAGTCTG CCGCTCCCAA CAGCCCGAGT TTCTTGCCCC TTGAGTAAGG GGCAAGAAAC CTAAGCCGCC TGGTGGATGG CTCGGCTCGG GGCGCTGACG AAGGGCGTGG CAAG
J23SpTT_47   CA GAAGACGGGT TTGATGGGGC GGGGATGTAA GCCGGGAAGGG AAACCGACCC GTTTAGTCTG CCGCTCCCAA CAGCCCGAGT TTCTTGCCCC TTGAGTAAGG GGCAAGAAAC CTAAGCCGCC TGGTGGATGG CTCGGCTCGG GGCGCTGACG AAGGGCGTGG CAAG
J23SpTT_27   CGCA GAAGACGGGT TTGATGGGGC GGGGATGTAA GCCGGGAAGGG AAACCGACCC GTTTAGTCTG CCGCTCCCAA CAGCCCGAGT TTCTTGCCCC TTGAGTAAGG GGCAAGAAAC CTAAGCCGCC TGGTGGATGG CTCGGCTCGG GGCGCTGACG AAGGGCGTGG CA
J23SpTT_604  CGCA GAAGACGGGT TTGATGGGGC GGGGATGTAA GCCGGGAAGGG AAACCGACCC GTTTAGTCTG CCGCTCCCAA CAGCCCGAGT TTCTTGCCCC TTGAGTAAGG GGCAAGAAAC CTAAGCCGCC TGGTGGATGG CTCGGCTCGG GGCGCTGACG AAGGGCGTGG CAAGCTGCGA TAAGCCCCGG TGAGGCGCAG GCAGCCGTCG AGCCGGGGAT TCCCGAATGG GACCTCCCGC GGCTTTTGCC GCACTCCCGT CAGGGAGGGG
J23SpTT_349  CCGCA GAAGACGGGT TTGATGGGGC GGGGATGTAA GCCGGGAAGGG AAACCGACCC GTTTAGTCTG CCGCTCCCAA CAGCCCGAGT TTCTTGCCCC TTGAGTAAGG GGCAAGAAAC CTAAGCCGCC TGGTGGATGG CTCGGCTCGG GGCGCTGACG AAGGGCGTGG CAAGCTGCGA TAAGCCCCGG TGAGGCGCAG GCAGCCGTCG AGCCGGGGAT TCCCGAATGG GACCTCCCGC GGCTTTTGCC GCACTCCCGT CAGGGAGGGG
J23SpTT_341  CCCGCA GAAGACGGGT TTGATGGGGC GGGGATGTAA GCCGGGAAGGG AAACCGACCC GTTTAGTCTG CCGCTCCCAA CAGCCCGAGT TTCTTGCCCC TTGAGTAAGG GGCAAGAAAC CTAAGCCGCC TGGTGGATGG CTCGGCTCGG GGCGCTGACG AAGGGCGTGG CAAGCTGCGA TAAGCCCCGG TGAGGCGCAG GCAGCCGTCG AGCCGGGGAT TCCCGAATGG GACCTCCCGC GGCTTTTGCC GCACTCCCGT CAGGGAGGGG
J23SpTT_90   CCCGCA GAAGACGGGT TTGATGGGGC GGGGATGTAA GCCGGGAAGGG AAACCGACCC GTTTAGTCTG CCGCTCCCAA CAGCCCGAGT TTCTTGCCCC TTGAGTAAGG GGCAAGAAAC CTAAGCCGCC TGGTGGATGG CTCGGCTCGG GGCGCTGACG AAGGGCGTGG CAAGCTGCGA TAAGCCCCGG TGAGGCGCAG GCAGCCGTCG AGCCGGGGAT TCCCGAATGG GACCTCCCGC GGCTTTTGCC GCACTCCCGT CAGGGAGGGG G
J23SpTT_57   CGCA GAAGACGGGT TTGATGGGGC GGGGATGTAA GCCGGGAAGGG AAACCGACCC GTTTAGTCTG CCGCTCCCAA CAGCCCGAGT TTCTTGCCCC TTGAGTAAGG GGCAAGAAAC CTAAGCCGCC TGGTGGATGG CTCGGCTCGG GGCGCTGACG AAGGGCGTGG CAAGCTGCGA TAAGCCCCGG TGAGGCGCAG GCAGCCGTCG AGCCGGGGAT TCCCGAATGG GACCTCCCGC GGCTTTTGCC GCACTCCCGT CAGGGAGGGG GAACGCGGG
J23SpTT_53   CGCA GAAGACGGGT TTGATGGGGC GGGGATGTAA GCCGGGAAGGG AAACCGACCC GTTTAGTCTG CCGCTCCCAA CAGCCCGAGT TTCTTGCCCC TTGAGTAAGG GGCAAGAAAC CTAAGCCGCC TGGTGGATGG CTCGGCTCGG GGCGCTGACG AAGGGCGTGG CAAGCTGCGA TAAGCCCCGG TGAGGCGCAG GCAGCCGTCG AGCCGGGGAT TCCCGAATGG GACCTCCCGC GGCTTTTGCC GCACTCCCGT CAGGGAGGGG GAACGCGGGG
J23SpTT_31   CCGCA GAAGACGGGT TTGATGGGGC GGGGATGTAA GCCGGGAAGGG AAACCGACCC GTTTAGTCTG CCGCTCCCAA CAGCCCGAGT TTCTTGCCCC TTGAGTAAGG GGCAAGAAAC CTAAGCCGCC TGGTGGATGG CTCGGCTCGG GGCGCTGACG AAGGGCGTGG CAAGCTGCGA TAAGCCCCGG TGAGGCGCAG GCAGCCGTCG AGCCGGGGAT TCCCGAATGG GACCTCCCGC GGCTTTTGCC GCACTCCCGT CAGGGAGGGG GAACGCGGGG
J23SpTT_26   CCCGCA GAAGACGGGT TTGATGGGGC GGGGATGTAA GCCGGGAAGGG AAACCGACCC GTTTAGTCTG CCGCTCCCAA CAGCCCGAGT TTCTTGCCCC TTGAGTAAGG GGCAAGAAAC CTAAGCCGCC TGGTGGATGG CTCGGCTCGG GGCGCTGACG AAGGGCGTGG CAAGCTGCGA TAAGCCCCGG TGAGGCGCAG GCAGCCGTCG AGCCGGGGAT TCCCGAATGG GACCTCCCGC GGCTTTTGCC GCACTCCCGT CAGGGAGGGG GAACGCGGGG
J23SpTT_16   CCCGCA GAAGACGGGT TTGATGGGGC GGGGATGTAA GCCGGGAAGGG AAACCGACCC GTTTAGTCTG CCGCTCCCAA CAGCCCGAGT TTCTTGCCCC TTGAGTAAGG GGCAAGAAAC CTAAGCCGCC TGGTGGATGG CTCGGCTCGG GGCGCTGACG AAGGGCGTGG CAAGCTGCGA TAAGCCCCGG TGAGGCGCAG GCAGCCGTCG AGCCGGGGAT TCCCGAATGG GACCTCCCGC GGCTTTTGCC GCACTCCCGT CAGGGAGGGG GAACGCGGG
J23SpTT_33   CCGCA GAAGACGGGT TTGATGGGGC GGGGATGTAA GCCGGGAAGGG AAACCGACCC GTTTAGTCTG CCGCTCCCAA CAGCCCGAGT TTCTTGCCCC TTGAGTAAGG GGCAAGAAAC CTAAGCCGCC TGGTGGATGG CTCGGCTCGG GGCGCTGACG AAGGGCGTGG CAAGCTGCGA TAAGCCCCGG TGAGGCGCAG GCAGCCGTCG AGCCGGGGAT TCCCGAATGG GACCTCCCGC GGCTTTTGCC GCACTCCCGT CAGGGAGGGG GAACGCGGG
J23SpTT_9    CCGCA GAAGACGGGT TTGATGGGGC GGGGATGTAA GCCGGGAAGGG AAACCGACCC GTTTAGTCTG CCGCTCCCAA CAGCCCGAGT TTCTTGCCCC TTGAGTAAGG GGCAAGAAAC CTAAGCCGCC TGGTGGATGG CTCGGCTCGG GGCGCTGACG AAGGGCGTGG CAAGCTGCGA TAAGCCCCGG TGAGGCGCAG GCAGCCGTCG AGCCGGGGAT TCCCGAATGG GACCTCCCGC GGCTTTTGCC GCACTCCCGT CAGGGAGGGG GAACGCGG
J23SpTT_21   CA GAAGACGGGT TTGATGGGGC GGGGATGTAA GCCGGGAAGGG AAACCGACCC GTTTAGTCTG CCGCTCCCAA CAGCCCGAGT TTCTTGCCCC TTGAGTAAGG GGCAAGAAAC CTAAGCCGCC TGGTGGATGG CTCGGCTCGG GGCGCTGACG AAGGGCGTGG CAAGCTGCGA TAAGCCCCGG TGAGGCGCAG GCAGCCGTCG AGCCGGGGAT TCCCGAATGG GACCTCCCGC GGCTTTTGCC GCACTCCCGT CAGGGAGGGG GAACGCGGGG
J23SpTT_9    CGCA GAAGACGGGT TTGATGGGGC GGGGATGTAA GCCGGGAAGGG AAACCGACCC GTTTAGTCTG CCGCTCCCAA CAGCCCGAGT TTCTTGCCCC TTGAGTAAGG GGCAAGAAAC CTAAGCCGCC TGGTGGATGG CTCGGCTCGG GGCGCTGACG AAGGGCGTGG CAAGCTGCGA TAAGCCCCGG TGAGGCGCAG GCAGCCGTCG AGCCGGGGAT TCCCGAATGG GACCTCCCGC GGCTTTTGCC GCACTCCCGT CAGGGAGGGG GAACGCGG
J23SpTT_26   CA GAAGACGGGT TTGATGGGGC GGGGATGTAA GCCGGGAAGGG AAACCGACCC GTTTAGTCTG CCGCTCCCAA CAGCCCGAGT TTCTTGCCCC TTGAGTAAGG GGCAAGAAAC CTAAGCCGCC TGGTGGATGG CTCGGCTCGG GGCGCTGACG AAGGGCGTGG CAAGCTGCGA TAAGCCCCGG TGAGGCGCAG GCAGCCGTCG AGCCGGGGAT TCCCGAATGG GACCTCCCGC GGCTTTTGCC GCACTCCCGT CAGGGAGGGG GAACGCGGG
J23SpTT_17   CCCGCA GAAGACGGGT TTGATGGGGC GGGGATGTAA GCCGGGAAGGG AAACCGACCC GTTTAGTCTG CCGCTCCCAA CAGCCCGAGT TTCTTGCCCC TTGAGTAAGG GGCAAGAAAC CTAAGCCGCC TGGTGGATGG CTCGGCTCGG GGCGCTGACG AAGGGCGTGG CAAGCTGCGA TAAGCCCCGG TGAGGCGCAG GCAGCCGTCG AGCCGGGGAT TCCCGAATGG GACCTCCCGC GGCTTTTGCC GCACTCCCGT CAGGGAGGGG GAAC
J23SpTT_19   CCGCA GAAGACGGGT TTGATGGGGC GGGGATGTAA GCCGGGAAGGG AAACCGACCC GTTTAGTCTG CCGCTCCCAA CAGCCCGAGT TTCTTGCCCC TTGAGTAAGG GGCAAGAAAC CTAAGCCGCC TGGTGGATGG CTCGGCTCGG GGCGCTGACG AAGGGCGTGG CAAGCTGCGA TAAGCCCCGG TGAGGCGCAG GCAGCCGTCG AGCCGGGGAT TCCCGAATGG GACCTCCCGC GGCTTTTGCC GCACTCCCGT CAGGGAGGGG GAAC
J23SpTT_31   CGCA GAAGACGGGT TTGATGGGGC GGGGATGTAA GCCGGGAAGGG AAACCGACCC GTTTAGTCTG CCGCTCCCAA CAGCCCGAGT TTCTTGCCCC TTGAGTAAGG GGCAAGAAAC CTAAGCCGCC TGGTGGATGG CTCGGCTCGG GGCGCTGACG AAGGGCGTGG CAAGCTGCGA TAAGCCCCGG TGAGGCGCAG GCAGCCGTCG AGCCGGGGAT TCCCGAATGG GACCTCCCGC GGCTTTTGCC GCACTCCCGT CAGGGAGGGG GAAC
J23SpTT_28   CCCGCA GAAGACGGGT TTGATGGGGC GGGGATGTAA GCCGGGAAGGG AAACCGACCC GTTTAGTCTG CCGCTCCCAA CAGCCCGAGT TTCTTGCCCC TTGAGTAAGG GGCAAGAAAC CTAAGCCGCC TGGTGGATGG CTCGGCTCGG GGCGCTGACG AAGGGCGTGG CAAGCTGCGA TAAGCCCCGG TGAGGCGCAG GCAGCCGTCG AGCCGGGGAT TCCCGAATGG GACCTCCCGC GGCTTTTGCC GCACTCCCGT CAGGGAGGGG GG
J23SpTT_10   CCGCA GAAGACGGGT TTGATGGGGC GGGGATGTAA GCCGGGAAGGG AAACCGACCC GTTTAGTCTG CCGCTCCCAA CAGCCCGAGT TTCTTGCCCC TTGAGTAAGG GGCAAGAAAC CTAAGCCGCC TGGTGGATGG CTCGGCTCGG GGCGCTGACG AAGGGCGTGG CAAGCTGCGA TAAGCCCCGG TGAGGCGCAG GCAGCCGTCG AGCCGGGGAT TCCCGAATGG GACCTCCCGC GGCTTTTGCC GCACTCCCGT CAGGGAGGGG GA
J23SpTT_80   CCGCA GAAGACGGGT TTGATGGGGC GGGGATGTAA GCCGGGAAGGG AAACCGACCC GTTTAGTCTG CCGCTCCCAA CAGCCCGAGT TTCTTGCCCC TTGAGTAAGG GGCAAGAAAC CTAAGCCGCC TGGTGGATGG CTCGGCTCGG GGCGCTGACG AAGGGCGTGG CAAGCTGCGA TAAGCCCCGG TGAGGCGCAG GCAGCCGTCG AGCCGGGGAT TCCCGAATGG GACCTCCCGC GGCTTTTGCC GCACTCCCGT CAGGGAGGGG G
J23SpTT_28   CCGCA GAAGACGGGT TTGATGGGGC GGGGATGTAA GCCGGGAAGGG AAACCGACCC GTTTAGTCTG CCGCTCCCAA CAGCCCGAGT TTCTTGCCCC TTGAGTAAGG GGCAAGAAAC CTAAGCCGCC TGGTGGATGG CTCGGCTCGG GGCGCTGACG AAGGGCGTGG CAAGCTGCGA TAAGCCCCGG TGAGGCGCAG GCAGCCGTCG AGCCGGGGAT TCCCGAATGG GACCTCCCGC GGCTTTTGCC GCACTCCCGT CAGGGAGGGG GG
J23SpTT_8    GCCCGCA GAAGACGGGT TTGATGGGGC GGGGATGTAA GCCGGGAAGGG AAACCGACCC GTTTAGTCTG CCGCTCCCAA CAGCCCGAGT TTCTTGCCCC TTGAGTAAGG GGCAAGAAAC CTAAGCCGCC TGGTGGATGG CTCGGCTCGG GGCGCTGACG AAGGGCGTGG CAAGCTGCGA TAAGCCCCGG TGAGGCGCAG GCAGCCGTCG AGCCGGGGAT TCCCGAATGG GACCTCCCGC GGCTTTTGCC GCACTCCCGT CAGGGAGGGG
J23SpTT_16   CA GAAGACGGGT TTGATGGGGC GGGGATGTAA GCCGGGAAGGG AAACCGACCC GTTTAGTCTG CCGCTCCCAA CAGCCCGAGT TTCTTGCCCC TTGAGTAAGG GGCAAGAAAC CTAAGCCGCC TGGTGGATGG CTCGGCTCGG GGCGCTGACG AAGGGCGTGG CAAGCTGCGA TAAGCCCCGG TGAGGCGCAG GCAGCCGTCG AGCCGGGGAT TCCCGAATGG GACCTCCCGC GGCTTTTGCC GCACTCCCGT CAGGGAGGGG GAAC
J23SpTT_13   CGCA GAAGACGGGT TTGATGGGGC GGGGATGTAA GCCGGGAAGGG AAACCGACCC GTTTAGTCTG CCGCTCCCAA CAGCCCGAGT TTCTTGCCCC TTGAGTAAGG GGCAAGAAAC CTAAGCCGCC TGGTGGATGG CTCGGCTCGG GGCGCTGACG AAGGGCGTGG CAAGCTGCGA TAAGCCCCGG TGAGGCGCAG GCAGCCGTCG AGCCGGGGAT TCCCGAATGG GACCTCCCGC GGCTTTTGCC GCACTCCCGT CAGGGAGGGG GAG
J23SpTT_12   CGCA GAAGACGGGT TTGATGGGGC GGGGATGTAA GCCGGGAAGGG AAACCGACCC GTTTAGTCTG CCGCTCCCAA CAGCCCGAGT TTCTTGCCCC TTGAGTAAGG GGCAAGAAAC CTAAGCCGCC TGGTGGATGG CTCGGCTCGG GGCGCTGACG AAGGGCGTGG CAAGCTGCGA TAAGCCCCGG TGAGGCGCAG GCAGCCGTCG AGCCGGGGAT TCCCGAATGG GACCTCCCGC GGCTTTTGCC GCACTCCCGT CAGGGAGGGG GA
J23SpTT_245  CCCGCA GAAGACGGGT TTGATGGGGC GGGGATGTAA GCCGGGAAGGG AAACCGACCC GTTTAGTCTG CCGCTCCCAA CAGCCCGAGT TTCTTGCCCC TTGAGTAAGG GGCAAGAAAC CTAAGCCGCC TGGTGGATGG CTCGGCTCGG GGCGCTGACG AAGGGCGTGG CAAGCTGCGA TAAGCCCCGG TGAGGCGCAG GCAGCCGTCG AGCCGGGGAT TCCCGAATGG GACCTCCCGC GGCTTTTGCC GCACTCCCGT CAGGGAGGG
J23SpTT_156  CGCA GAAGACGGGT TTGATGGGGC GGGGATGTAA GCCGGGAAGGG AAACCGACCC GTTTAGTCTG CCGCTCCCAA CAGCCCGAGT TTCTTGCCCC TTGAGTAAGG GGCAAGAAAC CTAAGCCGCC TGGTGGATGG CTCGGCTCGG GGCGCTGACG AAGGGCGTGG CAAGCTGCGA TAAGCCCCGG TGAGGCGCAG GCAGCCGTCG AGCCGGGGAT TCCCGAATGG GACCTCCCGC GGCTTTTGCC GCACTCCCGT CAGGGAGGGG G
J23SpTT_52   CGCA GAAGACGGGT TTGATGGGGC GGGGATGTAA GCCGGGAAGGG AAACCGACCC GTTTAGTCTG CCGCTCCCAA CAGCCCGAGT TTCTTGCCCC TTGAGTAAGG GGCAAGAAAC CTAAGCCGCC TGGTGGATGG CTCGGCTCGG GGCGCTGACG AAGGGCGTGG CAAGCTGCGA TAAGCCCCGG TGAGGCGCAG GCAGCCGTCG AGCCGGGGAT TCCCGAATGG GACCTCCCGC GGCTTTTGCC GCACTCCCGT CAGGGAGGGG GG
J23SpTT_15   GACGGGT TTGATGGGGC GGGGATGTAA GCCGGGAAGGG AAACCGACCC GTTTAGTCTG CCGCTCCCAA CAGCCCGAGT TTCTTGCCCC TTGAGTAAGG GGCAAGAAAC CTAAGCCGCC TGGTGGATGG CTCGGCTCGG GGCGCTGACG AAGGGCGTGG CAAGCTGCGA TAAGCCCCGG TGAGGCGCAG GCAGCCGTCG AGCCGGGGAT TCCCGAATGG GACCTCCCGC GGCTTTTGCC GCACTCCCGT CAGGGAGGGG GAACGCGGG
J23SpTT_10   GACGGGT TTGATGGGGC GGGGATGTAA GCCGGGAAGGG AAACCGACCC GTTTAGTCTG CCGCTCCCAA CAGCCCGAGT TTCTTGCCCC TTGAGTAAGG GGCAAGAAAC CTAAGCCGCC TGGTGGATGG CTCGGCTCGG GGCGCTGACG AAGGGCGTGG CAAGCTGCGA TAAGCCCCGG TGAGGCGCAG GCAGCCGTCG AGCCGGGGAT TCCCGAATGG GACCTCCCGC GGCTTTTGCC GCACTCCCGT CAGGGAGGGG GAACGCGGGG
J23SpTT_19   GCA GAAGACGGGT TTGATGGGGC GGGGATGTAA GCCGGGAAGGG AAACCGACCC GTTTAGTCTG CCGCTCCCAA CAGCCCGAGT TTCTTGCCCC TTGAGTAAGG GGCAAGAAAC CTAAGCCGCC TGGTGGATGG CTCGGCTCGG GGCGCTGACG AAGGGCGTGG CAAGCTGCGA TAAGCCCCGG TGAGGCGCAG GCAGCCGTCG AGCCGGGGAT TCCCGAATGG GACCTCCCGC GGCTTTTGCC GCACTCCCGT CAGGGAGGGG G
J23SpTT_9    CCCGCA GAAGACGGGT TTGATGGGGC GGGGATGTAA GCCGGGAAGGG AAACCGACCC GTTTAGTCTG CCGCTCCCAA CAGCCCGAGT TTCTTGCCCC TTGAGTAAGG GGCAAGAAAC CTAAGCCGCC TGGTGGATGG CTCGGCTCGG GGCGCTGACG AAGGGCGTGG CAAGCTGCGA TAAGCCCCGG TGAGGCGCAG GCAGCCGTCG AGCCGGGGAT TCCCGAATGG GACCTCCCGC GGCTTTTGCC GCACTCCCGT CAGGGAGGAG
J23SpTT_481  CGCA GAAGACGGGT TTGATGGGGC GGGGATGTAA GCCGGGAAGGG AAACCGACCC GTTTAGTCTG CCGCTCCCAA CAGCCCGAGT TTCTTGCCCC TTGAGTAAGG GGCAAGAAAC CTAAGCCGCC TGGTGGATGG CTCGGCTCGG GGCGCTGACG AAGGGCGTGG CAAGCTGCGA TAAGCCCCGG TGAGGCGCAG GCAGCCGTCG AGCCGGGGAT TCCCGAATGG GACCTCCCGC GGCTTTTGCC GCACTCCCGT CAGGGAGGG
J23SpTT_218  CCGCA GAAGACGGGT TTGATGGGGC GGGGATGTAA GCCGGGAAGGG AAACCGACCC GTTTAGTCTG CCGCTCCCAA CAGCCCGAGT TTCTTGCCCC TTGAGTAAGG GGCAAGAAAC CTAAGCCGCC TGGTGGATGG CTCGGCTCGG GGCGCTGACG AAGGGCGTGG CAAGCTGCGA TAAGCCCCGG TGAGGCGCAG GCAGCCGTCG AGCCGGGGAT TCCCGAATGG GACCTCCCGC GGCTTTTGCC GCACTCCCGT CAGGGAGGG
J23SpTT_111  CCCGCA GAAGACGGGT TTGATGGGGC GGGGATGTAA GCCGGGAAGGG AAACCGACCC GTTTAGTCTG CCGCTCCCAA CAGCCCGAGT TTCTTGCCCC TTGAGTAAGG GGCAAGAAAC CTAAGCCGCC TGGTGGATGG CTCGGCTCGG GGCGCTGACG AAGGGCGTGG CAAGCTGCGA TAAGCCCCGG TGAGGCGCAG GCAGCCGTCG AGCCGGGGAT TCCCGAATGG GACCTCCCGC GGCTTTTGCC GCACTCCCGT CAGGGAGG
J23SpTT_72   CA GAAGACGGGT TTGATGGGGC GGGGATGTAA GCCGGGAAGGG AAACCGACCC GTTTAGTCTG CCGCTCCCAA CAGCCCGAGT TTCTTGCCCC TTGAGTAAGG GGCAAGAAAC CTAAGCCGCC TGGTGGATGG CTCGGCTCGG GGCGCTGACG AAGGGCGTGG CAAGCTGCGA TAAGCCCCGG TGAGGCGCAG GCAGCCGTCG AGCCGGGGAT TCCCGAATGG GACCTCCCGC GGCTTTTGCC GCACTCCCGT CAGGGAGGGG G
J23SpTT_56   GCA GAAGACGGGT TTGATGGGGC GGGGATGTAA GCCGGGAAGGG AAACCGACCC GTTTAGTCTG CCGCTCCCAA CAGCCCGAGT TTCTTGCCCC TTGAGTAAGG GGCAAGAAAC CTAAGCCGCC TGGTGGATGG CTCGGCTCGG GGCGCTGACG AAGGGCGTGG CAAGCTGCGA TAAGCCCCGG TGAGGCGCAG GCAGCCGTCG AGCCGGGGAT TCCCGAATGG GACCTCCCGC GGCTTTTGCC GCACTCCCGT CAGGGAGGGG
J23SpTT_27   CA GAAGACGGGT TTGATGGGGC GGGGATGTAA GCCGGGAAGGG AAACCGACCC GTTTAGTCTG CCGCTCCCAA CAGCCCGAGT TTCTTGCCCC TTGAGTAAGG GGCAAGAAAC CTAAGCCGCC TGGTGGATGG CTCGGCTCGG GGCGCTGACG AAGGGCGTGG CAAGCTGCGA TAAGCCCCGG TGAGGCGCAG GCAGCCGTCG AGCCGGGGAT TCCCGAATGG GACCTCCCGC GGCTTTTGCC GCACTCCCGT CAGGGAGGGG GG
J23SpTT_10   CGGGT TTGATGGGGC GGGGATGTAA GCCGGGAAGGG AAACCGACCC GTTTAGTCTG CCGCTCCCAA CAGCCCGAGT TTCTTGCCCC TTGAGTAAGG GGCAAGAAAC CTAAGCCGCC TGGTGGATGG CTCGGCTCGG GGCGCTGACG AAGGGCGTGG CAAGCTGCGA TAAGCCCCGG TGAGGCGCAG GCAGCCGTCG AGCCGGGGAT TCCCGAATGG GACCTCCCGC GGCTTTTGCC GCACTCCCGT CAGGGAGGGG GAACGCGGG
J23SpTT_9    CCCGCA GAAGACGGGT TTGATGGGGC GGGGATGTAA GCCGGGAAGGG AAACCGACCC GTTTAGTCTG CCGCTCCCAA CAGCCCGAGT TTCTTGCCCC TTGAGTAAGG GGCAAGAAAC CTAAGCCGCC TGGTGGATGG CTCGGCTCGG GGCGCTGACG AAGGGCGTGG CAAGCTGCGA TAAGCCCCGG TGAGGCGCAG GCAGCCGTCG AGCCGGGGAT TCCCGAATGG GACCTCCCGC GGCTTTTGCC GCACTCCCGT CAGGGAGAG
J23SpTT_268  CGCA GAAGACGGGT TTGATGGGGC GGGGATGTAA GCCGGGAAGGG AAACCGACCC GTTTAGTCTG CCGCTCCCAA CAGCCCGAGT TTCTTGCCCC TTGAGTAAGG GGCAAGAAAC CTAAGCCGCC TGGTGGATGG CTCGGCTCGG GGCGCTGACG AAGGGCGTGG CAAGCTGCGA TAAGCCCCGG TGAGGCGCAG GCAGCCGTCG AGCCGGGGAT TCCCGAATGG GACCTCCCGC GGCTTTTGCC GCACTCCCGT CAGGGAGG
J23SpTT_260  CA GAAGACGGGT TTGATGGGGC GGGGATGTAA GCCGGGAAGGG AAACCGACCC GTTTAGTCTG CCGCTCCCAA CAGCCCGAGT TTCTTGCCCC TTGAGTAAGG GGCAAGAAAC CTAAGCCGCC TGGTGGATGG CTCGGCTCGG GGCGCTGACG AAGGGCGTGG CAAGCTGCGA TAAGCCCCGG TGAGGCGCAG GCAGCCGTCG AGCCGGGGAT TCCCGAATGG GACCTCCCGC GGCTTTTGCC GCACTCCCGT CAGGGAGGGG
J23SpTT_133  CCGCA GAAGACGGGT TTGATGGGGC GGGGATGTAA GCCGGGAAGGG AAACCGACCC GTTTAGTCTG CCGCTCCCAA CAGCCCGAGT TTCTTGCCCC TTGAGTAAGG GGCAAGAAAC CTAAGCCGCC TGGTGGATGG CTCGGCTCGG GGCGCTGACG AAGGGCGTGG CAAGCTGCGA TAAGCCCCGG TGAGGCGCAG GCAGCCGTCG AGCCGGGGAT TCCCGAATGG GACCTCCCGC GGCTTTTGCC GCACTCCCGT CAGGGAGG
J23SpTT_127  CCCGCA GAAGACGGGT TTGATGGGGC GGGGATGTAA GCCGGGAAGGG AAACCGACCC GTTTAGTCTG CCGCTCCCAA CAGCCCGAGT TTCTTGCCCC TTGAGTAAGG GGCAAGAAAC CTAAGCCGCC TGGTGGATGG CTCGGCTCGG GGCGCTGACG AAGGGCGTGG CAAGCTGCGA TAAGCCCCGG TGAGGCGCAG GCAGCCGTCG AGCCGGGGAT TCCCGAATGG GACCTCCCGC GGCTTTTGCC GCACTCCCGT CAGGGAG
J23SpTT_47   GCA GAAGACGGGT TTGATGGGGC GGGGATGTAA GCCGGGAAGGG AAACCGACCC GTTTAGTCTG CCGCTCCCAA CAGCCCGAGT TTCTTGCCCC TTGAGTAAGG GGCAAGAAAC CTAAGCCGCC TGGTGGATGG CTCGGCTCGG GGCGCTGACG AAGGGCGTGG CAAGCTGCGA TAAGCCCCGG TGAGGCGCAG GCAGCCGTCG AGCCGGGGAT TCCCGAATGG GACCTCCCGC GGCTTTTGCC GCACTCCCGT CAGGGAGGG
J23SpTT_31   A GAAGACGGGT TTGATGGGGC GGGGATGTAA GCCGGGAAGGG AAACCGACCC GTTTAGTCTG CCGCTCCCAA CAGCCCGAGT TTCTTGCCCC TTGAGTAAGG GGCAAGAAAC CTAAGCCGCC TGGTGGATGG CTCGGCTCGG GGCGCTGACG AAGGGCGTGG CAAGCTGCGA TAAGCCCCGG TGAGGCGCAG GCAGCCGTCG AGCCGGGGAT TCCCGAATGG GACCTCCCGC GGCTTTTGCC GCACTCCCGT CAGGGAGGGG G
J23SpTT_15   CGCA GAAGACGGGT TTGATGGGGC GGGGATGTAA GCCGGGAAGGG AAACCGACCC GTTTAGTCTG CCGCTCCCAA CAGCCCGAGT TTCTTGCCCC TTGAGTAAGG GGCAAGAAAC CTAAGCCGCC TGGTGGATGG CTCGGCTCGG GGCGCTGACG AAGGGCGTGG CAAGCTGCGA TAAGCCCCGG TGAGGCGCAG GCAGCCGTCG AGCCGGGGAT TCCCGAATGG GACCTCCCGC GGCTTTTGCC GCACTCCCGT CAGGGAGAG
J23SpTT_13   A GAAGACGGGT TTGATGGGGC GGGGATGTAA GCCGGGAAGGG AAACCGACCC GTTTAGTCTG CCGCTCCCAA CAGCCCGAGT TTCTTGCCCC TTGAGTAAGG GGCAAGAAAC CTAAGCCGCC TGGTGGATGG CTCGGCTCGG GGCGCTGACG AAGGGCGTGG CAAGCTGCGA TAAGCCCCGG TGAGGCGCAG GCAGCCGTCG AGCCGGGGAT TCCCGAATGG GACCTCCCGC GGCTTTTGCC GCACTCCCGT CAGGGAGGGG GG
J23SpTT_291  CGCA GAAGACGGGT TTGATGGGGC GGGGATGTAA GCCGGGAAGGG AAACCGACCC GTTTAGTCTG CCGCTCCCAA CAGCCCGAGT TTCTTGCCCC TTGAGTAAGG GGCAAGAAAC CTAAGCCGCC TGGTGGATGG CTCGGCTCGG GGCGCTGACG AAGGGCGTGG CAAGCTGCGA TAAGCCCCGG TGAGGCGCAG GCAGCCGTCG AGCCGGGGAT TCCCGAATGG GACCTCCCGC GGCTTTTGCC GCACTCCCGT CAGGGAG
J23SpTT_231  CA GAAGACGGGT TTGATGGGGC GGGGATGTAA GCCGGGAAGGG AAACCGACCC GTTTAGTCTG CCGCTCCCAA CAGCCCGAGT TTCTTGCCCC TTGAGTAAGG GGCAAGAAAC CTAAGCCGCC TGGTGGATGG CTCGGCTCGG GGCGCTGACG AAGGGCGTGG CAAGCTGCGA TAAGCCCCGG TGAGGCGCAG GCAGCCGTCG AGCCGGGGAT TCCCGAATGG GACCTCCCGC GGCTTTTGCC GCACTCCCGT CAGGGAGGG
J23SpTT_128  CCGCA GAAGACGGGT TTGATGGGGC GGGGATGTAA GCCGGGAAGGG AAACCGACCC GTTTAGTCTG CCGCTCCCAA CAGCCCGAGT TTCTTGCCCC TTGAGTAAGG GGCAAGAAAC CTAAGCCGCC TGGTGGATGG CTCGGCTCGG GGCGCTGACG AAGGGCGTGG CAAGCTGCGA TAAGCCCCGG TGAGGCGCAG GCAGCCGTCG AGCCGGGGAT TCCCGAATGG GACCTCCCGC GGCTTTTGCC GCACTCCCGT CAGGGAG
J23SpTT_116  A GAAGACGGGT TTGATGGGGC GGGGATGTAA GCCGGGAAGGG AAACCGACCC GTTTAGTCTG CCGCTCCCAA CAGCCCGAGT TTCTTGCCCC TTGAGTAAGG GGCAAGAAAC CTAAGCCGCC TGGTGGATGG CTCGGCTCGG GGCGCTGACG AAGGGCGTGG CAAGCTGCGA TAAGCCCCGG TGAGGCGCAG GCAGCCGTCG AGCCGGGGAT TCCCGAATGG GACCTCCCGC GGCTTTTGCC GCACTCCCGT CAGGGAGGGG
J23SpTT_24   GCA GAAGACGGGT TTGATGGGGC GGGGATGTAA GCCGGGAAGGG AAACCGACCC GTTTAGTCTG CCGCTCCCAA CAGCCCGAGT TTCTTGCCCC TTGAGTAAGG GGCAAGAAAC CTAAGCCGCC TGGTGGATGG CTCGGCTCGG GGCGCTGACG AAGGGCGTGG CAAGCTGCGA TAAGCCCCGG TGAGGCGCAG GCAGCCGTCG AGCCGGGGAT TCCCGAATGG GACCTCCCGC GGCTTTTGCC GCACTCCCGT CAGGGAGG
J23SpTT_105  CA GAAGACGGGT TTGATGGGGC GGGGATGTAA GCCGGGAAGGG AAACCGACCC GTTTAGTCTG CCGCTCCCAA CAGCCCGAGT TTCTTGCCCC TTGAGTAAGG GGCAAGAAAC CTAAGCCGCC TGGTGGATGG CTCGGCTCGG GGCGCTGACG AAGGGCGTGG CAAGCTGCGA TAAGCCCCGG TGAGGCGCAG GCAGCCGTCG AGCCGGGGAT TCCCGAATGG GACCTCCCGC GGCTTTTGCC GCACTCCCGT CAGGGAGG
J23SpTT_73   A GAAGACGGGT TTGATGGGGC GGGGATGTAA GCCGGGAAGGG AAACCGACCC GTTTAGTCTG CCGCTCCCAA CAGCCCGAGT TTCTTGCCCC TTGAGTAAGG GGCAAGAAAC CTAAGCCGCC TGGTGGATGG CTCGGCTCGG GGCGCTGACG AAGGGCGTGG CAAGCTGCGA TAAGCCCCGG TGAGGCGCAG GCAGCCGTCG AGCCGGGGAT TCCCGAATGG GACCTCCCGC GGCTTTTGCC GCACTCCCGT CAGGGAGGG
J23SpTT_37   GCA GAAGACGGGT TTGATGGGGC GGGGATGTAA GCCGGGAAGGG AAACCGACCC GTTTAGTCTG CCGCTCCCAA CAGCCCGAGT TTCTTGCCCC TTGAGTAAGG GGCAAGAAAC CTAAGCCGCC TGGTGGATGG CTCGGCTCGG GGCGCTGACG AAGGGCGTGG CAAGCTGCGA TAAGCCCCGG TGAGGCGCAG GCAGCCGTCG AGCCGGGGAT TCCCGAATGG GACCTCCCGC GGCTTTTGCC GCACTCCCGT CAGGGAG
J23SpTT_24   GAAGACGGGT TTGATGGGGC GGGGATGTAA GCCGGGAAGGG AAACCGACCC GTTTAGTCTG CCGCTCCCAA CAGCCCGAGT TTCTTGCCCC TTGAGTAAGG GGCAAGAAAC CTAAGCCGCC TGGTGGATGG CTCGGCTCGG GGCGCTGACG AAGGGCGTGG CAAGCTGCGA TAAGCCCCGG TGAGGCGCAG GCAGCCGTCG AGCCGGGGAT TCCCGAATGG GACCTCCCGC GGCTTTTGCC GCACTCCCGT CAGGGAGGGG
J23SpTT_9    CGCA GAAGACGGGT TTGATGGGGC GGGGATGTAA GCCGGGAAGGG AAACCGACCC GTTTAGTCTG CCGCTCCCAA CAGCCCGAGT TTCTTGCCCC TTGAGTAAGG GGCAAGAAAC CTAAGCCGCC TGGTGGATGG CTCGGCTCGG GGCGCTGACG AAGGGCGTGG CAAGCTGCGA TAAGCCCCGG TGAGGCGCAG GCAGCCGTCG AGCCGGGGAT TCCCGAATGG GACCTCCCGC GGCTTTTGCC GCACTCCCGT CAGGAAG
J23SpTT_18   CGCA GAAGACGGGT TTGATGGGGC GGGGATGTAA GCCGGGAAGGG AAACCGACCC GTTTAGTCTG CCGCTCCCAA CAGCCCGAGT TTCTTGCCCC TTGAGTAAGG GGCAAGAAAC CTAAGCCGCC TGGTGGATGG CTCGGCTCGG GGCGCTGACG AAGGGCGTGG CAAGCTGCGA TAAGCCCCGG TGAGGCGCAG GCAGCCGTCG AGCCGGGGAT TCCCGAATGG GACCTCCCGC GGCTTTTGCC GCACTCCCGT CAGGGA
J23SpTT_17   CCCGCA GAAGACGGGT TTGATGGGGC GGGGATGTAA GCCGGGAAGGG AAACCGACCC GTTTAGTCTG CCGCTCCCAA CAGCCCGAGT TTCTTGCCCC TTGAGTAAGG GGCAAGAAAC CTAAGCCGCC TGGTGGATGG CTCGGCTCGG GGCGCTGACG AAGGGCGTGG CAAGCTGCGA TAAGCCCCGG TGAGGCGCAG GCAGCCGTCG AGCCGGGGAT TCCCGAATGG GACCTCCCGC GGCTTTTGCC GCACTCCCGT CAGGG
J23SpTT_10   GT TTGATGGGGC GGGGATGTAA GCCGGGAAGGG AAACCGACCC GTTTAGTCTG CCGCTCCCAA CAGCCCGAGT TTCTTGCCCC TTGAGTAAGG GGCAAGAAAC CTAAGCCGCC TGGTGGATGG CTCGGCTCGG GGCGCTGACG AAGGGCGTGG CAAGCTGCGA TAAGCCCCGG TGAGGCGCAG GCAGCCGTCG AGCCGGGGAT TCCCGAATGG GACCTCCCGC GGCTTTTGCC GCACTCCCGT CAGGGAGGGG GAACGCGGG
J23SpTT_111  CA GAAGACGGGT TTGATGGGGC GGGGATGTAA GCCGGGAAGGG AAACCGACCC GTTTAGTCTG CCGCTCCCAA CAGCCCGAGT TTCTTGCCCC TTGAGTAAGG GGCAAGAAAC CTAAGCCGCC TGGTGGATGG CTCGGCTCGG GGCGCTGACG AAGGGCGTGG CAAGCTGCGA TAAGCCCCGG TGAGGCGCAG GCAGCCGTCG AGCCGGGGAT TCCCGAATGG GACCTCCCGC GGCTTTTGCC GCACTCCCGT CAGGGAG
J23SpTT_63   CCCGCA GAAGACGGGT TTGATGGGGC GGGGATGTAA GCCGGGAAGGG AAACCGACCC GTTTAGTCTG CCGCTCCCAA CAGCCCGAGT TTCTTGCCCC TTGAGTAAGG GGCAAGAAAC CTAAGCCGCC TGGTGGATGG CTCGGCTCGG GGCGCTGACG AAGGGCGTGG CAAGCTGCGA TAAGCCCCGG TGAGGCGCAG GCAGCCGTCG AGCCGGGGAT TCCCGAATGG GACCTCCCGC GGCTTTTGCC GCACTCCCGT CAGG
J23SpTT_42   CGCA GAAGACGGGT TTGATGGGGC GGGGATGTAA GCCGGGAAGGG AAACCGACCC GTTTAGTCTG CCGCTCCCAA CAGCCCGAGT TTCTTGCCCC TTGAGTAAGG GGCAAGAAAC CTAAGCCGCC TGGTGGATGG CTCGGCTCGG GGCGCTGACG AAGGGCGTGG CAAGCTGCGA TAAGCCCCGG TGAGGCGCAG GCAGCCGTCG AGCCGGGGAT TCCCGAATGG GACCTCCCGC GGCTTTTGCC GCACTCCCGT CAGGG
J23SpTT_29   A GAAGACGGGT TTGATGGGGC GGGGATGTAA GCCGGGAAGGG AAACCGACCC GTTTAGTCTG CCGCTCCCAA CAGCCCGAGT TTCTTGCCCC TTGAGTAAGG GGCAAGAAAC CTAAGCCGCC TGGTGGATGG CTCGGCTCGG GGCGCTGACG AAGGGCGTGG CAAGCTGCGA TAAGCCCCGG TGAGGCGCAG GCAGCCGTCG AGCCGGGGAT TCCCGAATGG GACCTCCCGC GGCTTTTGCC GCACTCCCGT CAGGGAGG
J23SpTT_22   AAGACGGGT TTGATGGGGC GGGGATGTAA GCCGGGAAGGG AAACCGACCC GTTTAGTCTG CCGCTCCCAA CAGCCCGAGT TTCTTGCCCC TTGAGTAAGG GGCAAGAAAC CTAAGCCGCC TGGTGGATGG CTCGGCTCGG GGCGCTGACG AAGGGCGTGG CAAGCTGCGA TAAGCCCCGG TGAGGCGCAG GCAGCCGTCG AGCCGGGGAT TCCCGAATGG GACCTCCCGC GGCTTTTGCC GCACTCCCGT CAGGGAGGGG
J23SpTT_12   CGGGT TTGATGGGGC GGGGATGTAA GCCGGGAAGGG AAACCGACCC GTTTAGTCTG CCGCTCCCAA CAGCCCGAGT TTCTTGCCCC TTGAGTAAGG GGCAAGAAAC CTAAGCCGCC TGGTGGATGG CTCGGCTCGG GGCGCTGACG AAGGGCGTGG CAAGCTGCGA TAAGCCCCGG TGAGGCGCAG GCAGCCGTCG AGCCGGGGAT TCCCGAATGG GACCTCCCGC GGCTTTTGCC GCACTCCCGT CAGGGAGGGG GAAC
J23SpTT_11   CCGCA GAAGACGGGT TTGATGGGGC GGGGATGTAA GCCGGGAAGGG AAACCGACCC GTTTAGTCTG CCGCTCCCAA CAGCCCGAGT TTCTTGCCCC TTGAGTAAGG GGCAAGAAAC CTAAGCCGCC TGGTGGATGG CTCGGCTCGG GGCGCTGACG AAGGGCGTGG CAAGCTGCGA TAAGCCCCGG TGAGGCGCAG GCAGCCGTCG AGCCGGGGAT TCCCGAATGG GACCTCCCGC GGCTTTTGCC GCACTCCCGT CAGGG
J23SpTT_9    GAAGACGGGT TTGATGGGGC GGGGATGTAA GCCGGGAAGGG AAACCGACCC GTTTAGTCTG CCGCTCCCAA CAGCCCGAGT TTCTTGCCCC TTGAGTAAGG GGCAAGAAAC CTAAGCCGCC TGGTGGATGG CTCGGCTCGG GGCGCTGACG AAGGGCGTGG CAAGCTGCGA TAAGCCCCGG TGAGGCGCAG GCAGCCGTCG AGCCGGGGAT TCCCGAATGG GACCTCCCGC GGCTTTTGCC GCACTCCCGT CAGGGAGGG
J23SpTT_114  CGCA GAAGACGGGT TTGATGGGGC GGGGATGTAA GCCGGGAAGGG AAACCGACCC GTTTAGTCTG CCGCTCCCAA CAGCCCGAGT TTCTTGCCCC TTGAGTAAGG GGCAAGAAAC CTAAGCCGCC TGGTGGATGG CTCGGCTCGG GGCGCTGACG AAGGGCGTGG CAAGCTGCGA TAAGCCCCGG TGAGGCGCAG GCAGCCGTCG AGCCGGGGAT TCCCGAATGG GACCTCCCGC GGCTTTTGCC GCACTCCCGT CAGG
J23SpTT_54   CCGCA GAAGACGGGT TTGATGGGGC GGGGATGTAA GCCGGGAAGGG AAACCGACCC GTTTAGTCTG CCGCTCCCAA CAGCCCGAGT TTCTTGCCCC TTGAGTAAGG GGCAAGAAAC CTAAGCCGCC TGGTGGATGG CTCGGCTCGG GGCGCTGACG AAGGGCGTGG CAAGCTGCGA TAAGCCCCGG TGAGGCGCAG GCAGCCGTCG AGCCGGGGAT TCCCGAATGG GACCTCCCGC GGCTTTTGCC GCACTCCCGT CAGG
J23SpTT_47   A GAAGACGGGT TTGATGGGGC GGGGATGTAA GCCGGGAAGGG AAACCGACCC GTTTAGTCTG CCGCTCCCAA CAGCCCGAGT TTCTTGCCCC TTGAGTAAGG GGCAAGAAAC CTAAGCCGCC TGGTGGATGG CTCGGCTCGG GGCGCTGACG AAGGGCGTGG CAAGCTGCGA TAAGCCCCGG TGAGGCGCAG GCAGCCGTCG AGCCGGGGAT TCCCGAATGG GACCTCCCGC GGCTTTTGCC GCACTCCCGT CAGGGAG
J23SpTT_40   AGACGGGT TTGATGGGGC GGGGATGTAA GCCGGGAAGGG AAACCGACCC GTTTAGTCTG CCGCTCCCAA CAGCCCGAGT TTCTTGCCCC TTGAGTAAGG GGCAAGAAAC CTAAGCCGCC TGGTGGATGG CTCGGCTCGG GGCGCTGACG AAGGGCGTGG CAAGCTGCGA TAAGCCCCGG TGAGGCGCAG GCAGCCGTCG AGCCGGGGAT TCCCGAATGG GACCTCCCGC GGCTTTTGCC GCACTCCCGT CAGGGAGGGG
J23SpTT_38   CCCGCA GAAGACGGGT TTGATGGGGC GGGGATGTAA GCCGGGAAGGG AAACCGACCC GTTTAGTCTG CCGCTCCCAA CAGCCCGAGT TTCTTGCCCC TTGAGTAAGG GGCAAGAAAC CTAAGCCGCC TGGTGGATGG CTCGGCTCGG GGCGCTGACG AAGGGCGTGG CAAGCTGCGA TAAGCCCCGG TGAGGCGCAG GCAGCCGTCG AGCCGGGGAT TCCCGAATGG GACCTCCCGC GGCTTTTGCC GCACTCCCGT CAG
J23SpTT_8    GCA GAAGACGGGT TTGATGGGGC GGGGATGTAA GCCGGGAAGGG AAACCGACCC GTTTAGTCTG CCGCTCCCAA CAGCCCGAGT TTCTTGCCCC TTGAGTAAGG GGCAAGAAAC CTAAGCCGCC TGGTGGATGG CTCGGCTCGG GGCGCTGACG AAGGGCGTGG CAAGCTGCGA TAAGCCCCGG TGAGGCGCAG GCAGCCGTCG AGCCGGGGAT TCCCGAATGG GACCTCCCGC GGCTTTTGCC GCACTCCCGT CAGGG
J23SpTT_21   AAGACGGGT TTGATGGGGC GGGGATGTAA GCCGGGAAGGG AAACCGACCC GTTTAGTCTG CCGCTCCCAA CAGCCCGAGT TTCTTGCCCC TTGAGTAAGG GGCAAGAAAC CTAAGCCGCC TGGTGGATGG CTCGGCTCGG GGCGCTGACG AAGGGCGTGG CAAGCTGCGA TAAGCCCCGG TGAGGCGCAG GCAGCCGTCG AGCCGGGGAT TCCCGAATGG GACCTCCCGC GGCTTTTGCC GCACTCCCGT CAGGGAGGG
J23SpTT_15   GACGGGT TTGATGGGGC GGGGATGTAA GCCGGGAAGGG AAACCGACCC GTTTAGTCTG CCGCTCCCAA CAGCCCGAGT TTCTTGCCCC TTGAGTAAGG GGCAAGAAAC CTAAGCCGCC TGGTGGATGG CTCGGCTCGG GGCGCTGACG AAGGGCGTGG CAAGCTGCGA TAAGCCCCGG TGAGGCGCAG GCAGCCGTCG AGCCGGGGAT TCCCGAATGG GACCTCCCGC GGCTTTTGCC GCACTCCCGT CAGGGAGGGG G
J23SpTT_14   GACGGGT TTGATGGGGC GGGGATGTAA GCCGGGAAGGG AAACCGACCC GTTTAGTCTG CCGCTCCCAA CAGCCCGAGT TTCTTGCCCC TTGAGTAAGG GGCAAGAAAC CTAAGCCGCC TGGTGGATGG CTCGGCTCGG GGCGCTGACG AAGGGCGTGG CAAGCTGCGA TAAGCCCCGG TGAGGCGCAG GCAGCCGTCG AGCCGGGGAT TCCCGAATGG GACCTCCCGC GGCTTTTGCC GCACTCCCGT CAGGGAGGGG GG
J23SpTT_10   GAAGACGGGT TTGATGGGGC GGGGATGTAA GCCGGGAAGGG AAACCGACCC GTTTAGTCTG CCGCTCCCAA CAGCCCGAGT TTCTTGCCCC TTGAGTAAGG GGCAAGAAAC CTAAGCCGCC TGGTGGATGG CTCGGCTCGG GGCGCTGACG AAGGGCGTGG CAAGCTGCGA TAAGCCCCGG TGAGGCGCAG GCAGCCGTCG AGCCGGGGAT TCCCGAATGG GACCTCCCGC GGCTTTTGCC GCACTCCCGT CAGGGAGG
J23SpTT_93   GACGGGT TTGATGGGGC GGGGATGTAA GCCGGGAAGGG AAACCGACCC GTTTAGTCTG CCGCTCCCAA CAGCCCGAGT TTCTTGCCCC TTGAGTAAGG GGCAAGAAAC CTAAGCCGCC TGGTGGATGG CTCGGCTCGG GGCGCTGACG AAGGGCGTGG CAAGCTGCGA TAAGCCCCGG TGAGGCGCAG GCAGCCGTCG AGCCGGGGAT TCCCGAATGG GACCTCCCGC GGCTTTTGCC GCACTCCCGT CAGGGAGGGG
J23SpTT_79   CGCA GAAGACGGGT TTGATGGGGC GGGGATGTAA GCCGGGAAGGG AAACCGACCC GTTTAGTCTG CCGCTCCCAA CAGCCCGAGT TTCTTGCCCC TTGAGTAAGG GGCAAGAAAC CTAAGCCGCC TGGTGGATGG CTCGGCTCGG GGCGCTGACG AAGGGCGTGG CAAGCTGCGA TAAGCCCCGG TGAGGCGCAG GCAGCCGTCG AGCCGGGGAT TCCCGAATGG GACCTCCCGC GGCTTTTGCC GCACTCCCGT CAG
J23SpTT_35   CCGCA GAAGACGGGT TTGATGGGGC GGGGATGTAA GCCGGGAAGGG AAACCGACCC GTTTAGTCTG CCGCTCCCAA CAGCCCGAGT TTCTTGCCCC TTGAGTAAGG GGCAAGAAAC CTAAGCCGCC TGGTGGATGG CTCGGCTCGG GGCGCTGACG AAGGGCGTGG CAAGCTGCGA TAAGCCCCGG TGAGGCGCAG GCAGCCGTCG AGCCGGGGAT TCCCGAATGG GACCTCCCGC GGCTTTTGCC GCACTCCCGT CAG
J23SpTT_30   AGACGGGT TTGATGGGGC GGGGATGTAA GCCGGGAAGGG AAACCGACCC GTTTAGTCTG CCGCTCCCAA CAGCCCGAGT TTCTTGCCCC TTGAGTAAGG GGCAAGAAAC CTAAGCCGCC TGGTGGATGG CTCGGCTCGG GGCGCTGACG AAGGGCGTGG CAAGCTGCGA TAAGCCCCGG TGAGGCGCAG GCAGCCGTCG AGCCGGGGAT TCCCGAATGG GACCTCCCGC GGCTTTTGCC GCACTCCCGT CAGGGAGGG
```

```
J23SpTT_25     CA GAAGACGGGT TTGATGGGGC GGGGATGTAA GCCGGGAAGG AAACCGACCC GTTTAGTCTG CCGCTCCCAA CAGCCCGAGT TTCTTGCCCC TTGAGTAAGG GGCAAGAAAC CTAAGCCGCC TGGTGGATGG CTCGGCTCGG GGCGCTGACG AAGGGCGTGG CAAGCTGCGA TAAGCCCCGG TGAGGCGCAG GCAGCCGTCG AGCCGGGGAT TCCCGAATGG GACCTCCCGC GGCTTTTGCC GCACTCCCGT CAGGG
J23SpTT_15     GCA GAAGACGGGT TTGATGGGGC GGGGATGTAA GCCGGGAAGG AAACCGACCC GTTTAGTCTG CCGCTCCCAA CAGCCCGAGT TTCTTGCCCC TTGAGTAAGG GGCAAGAAAC CTAAGCCGCC TGGTGGATGG CTCGGCTCGG GGCGCTGACG AAGGGCGTGG CAAGCTGCGA TAAGCCCCGG TGAGGCGCAG GCAGCCGTCG AGCCGGGGAT TCCCGAATGG GACCTCCCGC GGCTTTTGCC GCACTCCCGT CAGG
J23SpTT_14     AAGACGGGT TTGATGGGGC GGGGATGTAA GCCGGGAAGG AAACCGACCC GTTTAGTCTG CCGCTCCCAA CAGCCCGAGT TTCTTGCCCC TTGAGTAAGG GGCAAGAAAC CTAAGCCGCC TGGTGGATGG CTCGGCTCGG GGCGCTGACG AAGGGCGTGG CAAGCTGCGA TAAGCCCCGG TGAGGCGCAG GCAGCCGTCG AGCCGGGGAT TCCCGAATGG GACCTCCCGC GGCTTTTGCC GCACTCCCGT CAGGGAGG
J23SpTT_97     GACGGGT TTGATGGGGC GGGGATGTAA GCCGGGAAGG AAACCGACCC GTTTAGTCTG CCGCTCCCAA CAGCCCGAGT TTCTTGCCCC TTGAGTAAGG GGCAAGAAAC CTAAGCCGCC TGGTGGATGG CTCGGCTCGG GGCGCTGACG AAGGGCGTGG CAAGCTGCGA TAAGCCCCGG TGAGGCGCAG GCAGCCGTCG AGCCGGGGAT TCCCGAATGG GACCTCCCGC GGCTTTTGCC GCACTCCCGT CAGGGAGGG
J23SpTT_44     CA GAAGACGGGT TTGATGGGGC GGGGATGTAA GCCGGGAAGG AAACCGACCC GTTTAGTCTG CCGCTCCCAA CAGCCCGAGT TTCTTGCCCC TTGAGTAAGG GGCAAGAAAC CTAAGCCGCC TGGTGGATGG CTCGGCTCGG GGCGCTGACG AAGGGCGTGG CAAGCTGCGA TAAGCCCCGG TGAGGCGCAG GCAGCCGTCG AGCCGGGGAT TCCCGAATGG GACCTCCCGC GGCTTTTGCC GCACTCCCGT CAGG
J23SpTT_38     CGGGT TTGATGGGGC GGGGATGTAA GCCGGGAAGG AAACCGACCC GTTTAGTCTG CCGCTCCCAA CAGCCCGAGT TTCTTGCCCC TTGAGTAAGG GGCAAGAAAC CTAAGCCGCC TGGTGGATGG CTCGGCTCGG GGCGCTGACG AAGGGCGTGG CAAGCTGCGA TAAGCCCCGG TGAGGCGCAG GCAGCCGTCG AGCCGGGGAT TCCCGAATGG GACCTCCCGC GGCTTTTGCC GCACTCCCGT CAGGGAGGGG G
J23SpTT_8      GCA GAAGACGGGT TTGATGGGGC GGGGATGTAA GCCGGGAAGG AAACCGACCC GTTTAGTCTG CCGCTCCCAA CAGCCCGAGT TTCTTGCCCC TTGAGTAAGG GGCAAGAAAC CTAAGCCGCC TGGTGGATGG CTCGGCTCGG GGCGCTGACG AAGGGCGTGG CAAGCTGCGA TAAGCCCCGG TGAGGCGCAG GCAGCCGTCG AGCCGGGGAT TCCCGAATGG GACCTCCCGC GGCTTTTGCC GCACTCCCGT CAG
J23SpTT_30     ACGGGT TTGATGGGGC GGGGATGTAA GCCGGGAAGG AAACCGACCC GTTTAGTCTG CCGCTCCCAA CAGCCCGAGT TTCTTGCCCC TTGAGTAAGG GGCAAGAAAC CTAAGCCGCC TGGTGGATGG CTCGGCTCGG GGCGCTGACG AAGGGCGTGG CAAGCTGCGA TAAGCCCCGG TGAGGCGCAG GCAGCCGTCG AGCCGGGGAT TCCCGAATGG GACCTCCCGC GGCTTTTGCC GCACTCCCGT CAGGGAGGGG
J23SpTT_17     AAGACGGGT TTGATGGGGC GGGGATGTAA GCCGGGAAGG AAACCGACCC GTTTAGTCTG CCGCTCCCAA CAGCCCGAGT TTCTTGCCCC TTGAGTAAGG GGCAAGAAAC CTAAGCCGCC TGGTGGATGG CTCGGCTCGG GGCGCTGACG AAGGGCGTGG CAAGCTGCGA TAAGCCCCGG TGAGGCGCAG GCAGCCGTCG AGCCGGGGAT TCCCGAATGG GACCTCCCGC GGCTTTTGCC GCACTCCCGT CAGGGAG
J23SpTT_10     AGACGGGT TTGATGGGGC GGGGATGTAA GCCGGGAAGG AAACCGACCC GTTTAGTCTG CCGCTCCCAA CAGCCCGAGT TTCTTGCCCC TTGAGTAAGG GGCAAGAAAC CTAAGCCGCC TGGTGGATGG CTCGGCTCGG GGCGCTGACG AAGGGCGTGG CAAGCTGCGA TAAGCCCCGG TGAGGCGCAG GCAGCCGTCG AGCCGGGGAT TCCCGAATGG GACCTCCCGC GGCTTTTGCC GCACTCCCGT CAGGGAGG
J23SpTT_9      CGGGT TTGATGGGGC GGGGATGTAA GCCGGGAAGG AAACCGACCC GTTTAGTCTG CCGCTCCCAA CAGCCCGAGT TTCTTGCCCC TTGAGTAAGG GGCAAGAAAC CTAAGCCGCC TGGTGGATGG CTCGGCTCGG GGCGCTGACG AAGGGCGTGG CAAGCTGCGA TAAGCCCCGG TGAGGCGCAG GCAGCCGTCG AGCCGGGGAT TCCCGAATGG GACCTCCCGC GGCTTTTGCC GCACTCCCGT CAGGGAGGGG GG
J23SpTT_12     CGCA GAAGACGGGT TTGATGGGGC GGGGATGTAA GCCGGGAAGG AAACCGACCC GTTTAGTCTG CCGCTCCCAA CAGCCCGAGT TTCTTGCCCC TTGAGTAAGG GGCAAGAAAC CTAAGCCGCC TGGTGGATGG CTCGGCTCGG GGCGCTGACG AAGGGCGTGG CAAGCTGCGA TAAGCCCCGG TGAGGCGCAG GCAGCCGTCG AGCCGGGGAT TCCCGAATGG GACCTCCCGC GGCTTTTGCC GCACTCCCGT CA
J23SpTT_137    CGGGT TTGATGGGGC GGGGATGTAA GCCGGGAAGG AAACCGACCC GTTTAGTCTG CCGCTCCCAA CAGCCCGAGT TTCTTGCCCC TTGAGTAAGG GGCAAGAAAC CTAAGCCGCC TGGTGGATGG CTCGGCTCGG GGCGCTGACG AAGGGCGTGG CAAGCTGCGA TAAGCCCCGG TGAGGCGCAG GCAGCCGTCG AGCCGGGGAT TCCCGAATGG GACCTCCCGC GGCTTTTGCC GCACTCCCGT CAGGGAGGGG
J23SpTT_49     GACGGGT TTGATGGGGC GGGGATGTAA GCCGGGAAGG AAACCGACCC GTTTAGTCTG CCGCTCCCAA CAGCCCGAGT TTCTTGCCCC TTGAGTAAGG GGCAAGAAAC CTAAGCCGCC TGGTGGATGG CTCGGCTCGG GGCGCTGACG AAGGGCGTGG CAAGCTGCGA TAAGCCCCGG TGAGGCGCAG GCAGCCGTCG AGCCGGGGAT TCCCGAATGG GACCTCCCGC GGCTTTTGCC GCACTCCCGT CAGGGAGG
J23SpTT_47     ACGGGT TTGATGGGGC GGGGATGTAA GCCGGGAAGG AAACCGACCC GTTTAGTCTG CCGCTCCCAA CAGCCCGAGT TTCTTGCCCC TTGAGTAAGG GGCAAGAAAC CTAAGCCGCC TGGTGGATGG CTCGGCTCGG GGCGCTGACG AAGGGCGTGG CAAGCTGCGA TAAGCCCCGG TGAGGCGCAG GCAGCCGTCG AGCCGGGGAT TCCCGAATGG GACCTCCCGC GGCTTTTGCC GCACTCCCGT CAGGGAGGG
J23SpTT_25     A GAAGACGGGT TTGATGGGGC GGGGATGTAA GCCGGGAAGG AAACCGACCC GTTTAGTCTG CCGCTCCCAA CAGCCCGAGT TTCTTGCCCC TTGAGTAAGG GGCAAGAAAC CTAAGCCGCC TGGTGGATGG CTCGGCTCGG GGCGCTGACG AAGGGCGTGG CAAGCTGCGA TAAGCCCCGG TGAGGCGCAG GCAGCCGTCG AGCCGGGGAT TCCCGAATGG GACCTCCCGC GGCTTTTGCC GCACTCCCGT CAGG
J23SpTT_21     CA GAAGACGGGT TTGATGGGGC GGGGATGTAA GCCGGGAAGG AAACCGACCC GTTTAGTCTG CCGCTCCCAA CAGCCCGAGT TTCTTGCCCC TTGAGTAAGG GGCAAGAAAC CTAAGCCGCC TGGTGGATGG CTCGGCTCGG GGCGCTGACG AAGGGCGTGG CAAGCTGCGA TAAGCCCCGG TGAGGCGCAG GCAGCCGTCG AGCCGGGGAT TCCCGAATGG GACCTCCCGC GGCTTTTGCC GCACTCCCGT CAG
J23SpTT_21     AGACGGGT TTGATGGGGC GGGGATGTAA GCCGGGAAGG AAACCGACCC GTTTAGTCTG CCGCTCCCAA CAGCCCGAGT TTCTTGCCCC TTGAGTAAGG GGCAAGAAAC CTAAGCCGCC TGGTGGATGG CTCGGCTCGG GGCGCTGACG AAGGGCGTGG CAAGCTGCGA TAAGCCCCGG TGAGGCGCAG GCAGCCGTCG AGCCGGGGAT TCCCGAATGG GACCTCCCGC GGCTTTTGCC GCACTCCCGT CAGGGAG
J23SpTT_144    CGGGT TTGATGGGGC GGGGATGTAA GCCGGGAAGG AAACCGACCC GTTTAGTCTG CCGCTCCCAA CAGCCCGAGT TTCTTGCCCC TTGAGTAAGG GGCAAGAAAC CTAAGCCGCC TGGTGGATGG CTCGGCTCGG GGCGCTGACG AAGGGCGTGG CAAGCTGCGA TAAGCCCCGG TGAGGCGCAG GCAGCCGTCG AGCCGGGGAT TCCCGAATGG GACCTCCCGC GGCTTTTGCC GCACTCCCGT CAGGGAGGG
J23SpTT_71     GACGGGT TTGATGGGGC GGGGATGTAA GCCGGGAAGG AAACCGACCC GTTTAGTCTG CCGCTCCCAA CAGCCCGAGT TTCTTGCCCC TTGAGTAAGG GGCAAGAAAC CTAAGCCGCC TGGTGGATGG CTCGGCTCGG GGCGCTGACG AAGGGCGTGG CAAGCTGCGA TAAGCCCCGG TGAGGCGCAG GCAGCCGTCG AGCCGGGGAT TCCCGAATGG GACCTCCCGC GGCTTTTGCC GCACTCCCGT CAGGGAG
J23SpTT_15     ACGGGT TTGATGGGGC GGGGATGTAA GCCGGGAAGG AAACCGACCC GTTTAGTCTG CCGCTCCCAA CAGCCCGAGT TTCTTGCCCC TTGAGTAAGG GGCAAGAAAC CTAAGCCGCC TGGTGGATGG CTCGGCTCGG GGCGCTGACG AAGGGCGTGG CAAGCTGCGA TAAGCCCCGG TGAGGCGCAG GCAGCCGTCG AGCCGGGGAT TCCCGAATGG GACCTCCCGC GGCTTTTGCC GCACTCCCGT CAGGGAGG
J23SpTT_12     GGGT TTGATGGGGC GGGGATGTAA GCCGGGAAGG AAACCGACCC GTTTAGTCTG CCGCTCCCAA CAGCCCGAGT TTCTTGCCCC TTGAGTAAGG GGCAAGAAAC CTAAGCCGCC TGGTGGATGG CTCGGCTCGG GGCGCTGACG AAGGGCGTGG CAAGCTGCGA TAAGCCCCGG TGAGGCGCAG GCAGCCGTCG AGCCGGGGAT TCCCGAATGG GACCTCCCGC GGCTTTTGCC GCACTCCCGT CAGGGAGGGG
J23SpTT_63     CGGGT TTGATGGGGC GGGGATGTAA GCCGGGAAGG AAACCGACCC GTTTAGTCTG CCGCTCCCAA CAGCCCGAGT TTCTTGCCCC TTGAGTAAGG GGCAAGAAAC CTAAGCCGCC TGGTGGATGG CTCGGCTCGG GGCGCTGACG AAGGGCGTGG CAAGCTGCGA TAAGCCCCGG TGAGGCGCAG GCAGCCGTCG AGCCGGGGAT TCCCGAATGG GACCTCCCGC GGCTTTTGCC GCACTCCCGT CAGGGAGG
J23SpTT_35     ACGGGT TTGATGGGGC GGGGATGTAA GCCGGGAAGG AAACCGACCC GTTTAGTCTG CCGCTCCCAA CAGCCCGAGT TTCTTGCCCC TTGAGTAAGG GGCAAGAAAC CTAAGCCGCC TGGTGGATGG CTCGGCTCGG GGCGCTGACG AAGGGCGTGG CAAGCTGCGA TAAGCCCCGG TGAGGCGCAG GCAGCCGTCG AGCCGGGGAT TCCCGAATGG GACCTCCCGC GGCTTTTGCC GCACTCCCGT CAGGGAG
J23SpTT_16     GT TTGATGGGGC GGGGATGTAA GCCGGGAAGG AAACCGACCC GTTTAGTCTG CCGCTCCCAA CAGCCCGAGT TTCTTGCCCC TTGAGTAAGG GGCAAGAAAC CTAAGCCGCC TGGTGGATGG CTCGGCTCGG GGCGCTGACG AAGGGCGTGG CAAGCTGCGA TAAGCCCCGG TGAGGCGCAG GCAGCCGTCG AGCCGGGGAT TCCCGAATGG GACCTCCCGC GGCTTTTGCC GCACTCCCGT CAGGGAGGGG G
J23SpTT_13     GGGT TTGATGGGGC GGGGATGTAA GCCGGGAAGG AAACCGACCC GTTTAGTCTG CCGCTCCCAA CAGCCCGAGT TTCTTGCCCC TTGAGTAAGG GGCAAGAAAC CTAAGCCGCC TGGTGGATGG CTCGGCTCGG GGCGCTGACG AAGGGCGTGG CAAGCTGCGA TAAGCCCCGG TGAGGCGCAG GCAGCCGTCG AGCCGGGGAT TCCCGAATGG GACCTCCCGC GGCTTTTGCC GCACTCCCGT CAGGGAGGG
J23SpTT_11     GGT TTGATGGGGC GGGGATGTAA GCCGGGAAGG AAACCGACCC GTTTAGTCTG CCGCTCCCAA CAGCCCGAGT TTCTTGCCCC TTGAGTAAGG GGCAAGAAAC CTAAGCCGCC TGGTGGATGG CTCGGCTCGG GGCGCTGACG AAGGGCGTGG CAAGCTGCGA TAAGCCCCGG TGAGGCGCAG GCAGCCGTCG AGCCGGGGAT TCCCGAATGG GACCTCCCGC GGCTTTTGCC GCACTCCCGT CAGGGAGGGG
J23SpTT_70     CGGGT TTGATGGGGC GGGGATGTAA GCCGGGAAGG AAACCGACCC GTTTAGTCTG CCGCTCCCAA CAGCCCGAGT TTCTTGCCCC TTGAGTAAGG GGCAAGAAAC CTAAGCCGCC TGGTGGATGG CTCGGCTCGG GGCGCTGACG AAGGGCGTGG CAAGCTGCGA TAAGCCCCGG TGAGGCGCAG GCAGCCGTCG AGCCGGGGAT TCCCGAATGG GACCTCCCGC GGCTTTTGCC GCACTCCCGT CAGGGAG
J23SpTT_35     GT TTGATGGGGC GGGGATGTAA GCCGGGAAGG AAACCGACCC GTTTAGTCTG CCGCTCCCAA CAGCCCGAGT TTCTTGCCCC TTGAGTAAGG GGCAAGAAAC CTAAGCCGCC TGGTGGATGG CTCGGCTCGG GGCGCTGACG AAGGGCGTGG CAAGCTGCGA TAAGCCCCGG TGAGGCGCAG GCAGCCGTCG AGCCGGGGAT TCCCGAATGG GACCTCCCGC GGCTTTTGCC GCACTCCCGT CAGGGAGGGG
J23SpTT_39     GT TTGATGGGGC GGGGATGTAA GCCGGGAAGG AAACCGACCC GTTTAGTCTG CCGCTCCCAA CAGCCCGAGT TTCTTGCCCC TTGAGTAAGG GGCAAGAAAC CTAAGCCGCC TGGTGGATGG CTCGGCTCGG GGCGCTGACG AAGGGCGTGG CAAGCTGCGA TAAGCCCCGG TGAGGCGCAG GCAGCCGTCG AGCCGGGGAT TCCCGAATGG GACCTCCCGC GGCTTTTGCC GCACTCCCGT CAGGGAGGG
J23SpTT_19     GACGGGT TTGATGGGGC GGGGATGTAA GCCGGGAAGG AAACCGACCC GTTTAGTCTG CCGCTCCCAA CAGCCCGAGT TTCTTGCCCC TTGAGTAAGG GGCAAGAAAC CTAAGCCGCC TGGTGGATGG CTCGGCTCGG GGCGCTGACG AAGGGCGTGG CAAGCTGCGA TAAGCCCCGG TGAGGCGCAG GCAGCCGTCG AGCCGGGGAT TCCCGAATGG GACCTCCCGC GGCTTTTGCC GCACTCCCGT CAGG
J23SpTT_8      GGGT TTGATGGGGC GGGGATGTAA GCCGGGAAGG AAACCGACCC GTTTAGTCTG CCGCTCCCAA CAGCCCGAGT TTCTTGCCCC TTGAGTAAGG GGCAAGAAAC CTAAGCCGCC TGGTGGATGG CTCGGCTCGG GGCGCTGACG AAGGGCGTGG CAAGCTGCGA TAAGCCCCGG TGAGGCGCAG GCAGCCGTCG AGCCGGGGAT TCCCGAATGG GACCTCCCGC GGCTTTTGCC GCACTCCCGT CAGGGAG
J23SpTT_15     GT TTGATGGGGC GGGGATGTAA GCCGGGAAGG AAACCGACCC GTTTAGTCTG CCGCTCCCAA CAGCCCGAGT TTCTTGCCCC TTGAGTAAGG GGCAAGAAAC CTAAGCCGCC TGGTGGATGG CTCGGCTCGG GGCGCTGACG AAGGGCGTGG CAAGCTGCGA TAAGCCCCGG TGAGGCGCAG GCAGCCGTCG AGCCGGGGAT TCCCGAATGG GACCTCCCGC GGCTTTTGCC GCACTCCCGT CAGGGAGG
J23SpTT_13     GACGGGT TTGATGGGGC GGGGATGTAA GCCGGGAAGG AAACCGACCC GTTTAGTCTG CCGCTCCCAA CAGCCCGAGT TTCTTGCCCC TTGAGTAAGG GGCAAGAAAC CTAAGCCGCC TGGTGGATGG CTCGGCTCGG GGCGCTGACG AAGGGCGTGG CAAGCTGCGA TAAGCCCCGG TGAGGCGCAG GCAGCCGTCG AGCCGGGGAT TCCCGAATGG GACCTCCCGC GGCTTTTGCC GCACTCCCGT CAG
J23SpTT_13     CGGGT TTGATGGGGC GGGGATGTAA GCCGGGAAGG AAACCGACCC GTTTAGTCTG CCGCTCCCAA CAGCCCGAGT TTCTTGCCCC TTGAGTAAGG GGCAAGAAAC CTAAGCCGCC TGGTGGATGG CTCGGCTCGG GGCGCTGACG AAGGGCGTGG CAAGCTGCGA TAAGCCCCGG TGAGGCGCAG GCAGCCGTCG AGCCGGGGAT TCCCGAATGG GACCTCCCGC GGCTTTTGCC GCACTCCCGT CAGGG
J23SpTT_11     ACGGGT TTGATGGGGC GGGGATGTAA GCCGGGAAGG AAACCGACCC GTTTAGTCTG CCGCTCCCAA CAGCCCGAGT TTCTTGCCCC TTGAGTAAGG GGCAAGAAAC CTAAGCCGCC TGGTGGATGG CTCGGCTCGG GGCGCTGACG AAGGGCGTGG CAAGCTGCGA TAAGCCCCGG TGAGGCGCAG GCAGCCGTCG AGCCGGGGAT TCCCGAATGG GACCTCCCGC GGCTTTTGCC GCACTCCCGT CAGG
J23SpTT_31     CGGGT TTGATGGGGC GGGGATGTAA GCCGGGAAGG AAACCGACCC GTTTAGTCTG CCGCTCCCAA CAGCCCGAGT TTCTTGCCCC TTGAGTAAGG GGCAAGAAAC CTAAGCCGCC TGGTGGATGG CTCGGCTCGG GGCGCTGACG AAGGGCGTGG CAAGCTGCGA TAAGCCCCGG TGAGGCGCAG GCAGCCGTCG AGCCGGGGAT TCCCGAATGG GACCTCCCGC GGCTTTTGCC GCACTCCCGT CAGG
J23SpTT_20     GT TTGATGGGGC GGGGATGTAA GCCGGGAAGG AAACCGACCC GTTTAGTCTG CCGCTCCCAA CAGCCCGAGT TTCTTGCCCC TTGAGTAAGG GGCAAGAAAC CTAAGCCGCC TGGTGGATGG CTCGGCTCGG GGCGCTGACG AAGGGCGTGG CAAGCTGCGA TAAGCCCCGG TGAGGCGCAG GCAGCCGTCG AGCCGGGGAT TCCCGAATGG GACCTCCCGC GGCTTTTGCC GCACTCCCGT CAGGGAG
J23SpTT_13     C GGGGATGTAA GCCGGGAAGG AAACCGACCC GTTTAGTCTG CCGCTCCCAA CAGCCCGAGT TTCTTGCCCC TTGAGTAAGG GGCAAGAAAC CTAAGCCGCC TGGTGGATGG CTCGGCTCGG GGCGCTGACG AAGGGCGTGG CAAGCTGCGA TAAGCCCCGG TGAGGCGCAG GCAGCCGTCG AGCCGGGGAT TCCCGAATGG GACCTCCCGC GGCTTTTGCC GCACTCCCGT CAGGGAGGGG GAACGCGGGG
J23SpTT_12     C GGGGATGTAA GCCGGGAAGG AAACCGACCC GTTTAGTCTG CCGCTCCCAA CAGCCCGAGT TTCTTGCCCC TTGAGTAAGG GGCAAGAAAC CTAAGCCGCC TGGTGGATGG CTCGGCTCGG GGCGCTGACG AAGGGCGTGG CAAGCTGCGA TAAGCCCCGG TGAGGCGCAG GCAGCCGTCG AGCCGGGGAT TCCCGAATGG GACCTCCCGC GGCTTTTGCC GCACTCCCGT CAGGGAGGGG GAACGCGGG
J23SpTT_17     CGGGT TTGATGGGGC GGGGATGTAA GCCGGGAAGG AAACCGACCC GTTTAGTCTG CCGCTCCCAA CAGCCCGAGT TTCTTGCCCC TTGAGTAAGG GGCAAGAAAC CTAAGCCGCC TGGTGGATGG CTCGGCTCGG GGCGCTGACG AAGGGCGTGG CAAGCTGCGA TAAGCCCCGG TGAGGCGCAG GCAGCCGTCG AGCCGGGGAT TCCCGAATGG GACCTCCCGC GGCTTTTGCC GCACTCCCGT CAG
J23SpTT_11     TGGGGC GGGGATGTAA GCCGGGAAGG AAACCGACCC GTTTAGTCTG CCGCTCCCAA CAGCCCGAGT TTCTTGCCCC TTGAGTAAGG GGCAAGAAAC CTAAGCCGCC TGGTGGATGG CTCGGCTCGG GGCGCTGACG AAGGGCGTGG CAAGCTGCGA TAAGCCCCGG TGAGGCGCAG GCAGCCGTCG AGCCGGGGAT TCCCGAATGG GACCTCCCGC GGCTTTTGCC GCACTCCCGT CAGGGAGGGG
J23SpTT_8      GGGGC GGGGATGTAA GCCGGGAAGG AAACCGACCC GTTTAGTCTG CCGCTCCCAA CAGCCCGAGT TTCTTGCCCC TTGAGTAAGG GGCAAGAAAC CTAAGCCGCC TGGTGGATGG CTCGGCTCGG GGCGCTGACG AAGGGCGTGG CAAGCTGCGA TAAGCCCCGG TGAGGCGCAG GCAGCCGTCG AGCCGGGGAT TCCCGAATGG GACCTCCCGC GGCTTTTGCC GCACTCCCGT CAGGGAG
J23SpTT_24     C GGGGATGTAA GCCGGGAAGG AAACCGACCC GTTTAGTCTG CCGCTCCCAA CAGCCCGAGT TTCTTGCCCC TTGAGTAAGG GGCAAGAAAC CTAAGCCGCC TGGTGGATGG CTCGGCTCGG GGCGCTGACG AAGGGCGTGG CAAGCTGCGA TAAGCCCCGG TGAGGCGCAG GCAGCCGTCG AGCCGGGGAT TCCCGAATGG GACCTCCCGC GGCTTTTGCC GCACTCCCGT CAGGGAGGGG G
J23SpTT_12     C GGGGATGTAA GCCGGGAAGG AAACCGACCC GTTTAGTCTG CCGCTCCCAA CAGCCCGAGT TTCTTGCCCC TTGAGTAAGG GGCAAGAAAC CTAAGCCGCC TGGTGGATGG CTCGGCTCGG GGCGCTGACG AAGGGCGTGG CAAGCTGCGA TAAGCCCCGG TGAGGCGCAG GCAGCCGTCG AGCCGGGGAT TCCCGAATGG GACCTCCCGC GGCTTTTGCC GCACTCCCGT CAGGGAGGGG GG
J23SpTT_129    C GGGGATGTAA GCCGGGAAGG AAACCGACCC GTTTAGTCTG CCGCTCCCAA CAGCCCGAGT TTCTTGCCCC TTGAGTAAGG GGCAAGAAAC CTAAGCCGCC TGGTGGATGG CTCGGCTCGG GGCGCTGACG AAGGGCGTGG CAAGCTGCGA TAAGCCCCGG TGAGGCGCAG GCAGCCGTCG AGCCGGGGAT TCCCGAATGG GACCTCCCGC GGCTTTTGCC GCACTCCCGT CAGGGAGGGG
J23SpTT_91     C GGGGATGTAA GCCGGGAAGG AAACCGACCC GTTTAGTCTG CCGCTCCCAA CAGCCCGAGT TTCTTGCCCC TTGAGTAAGG GGCAAGAAAC CTAAGCCGCC TGGTGGATGG CTCGGCTCGG GGCGCTGACG AAGGGCGTGG CAAGCTGCGA TAAGCCCCGG TGAGGCGCAG GCAGCCGTCG AGCCGGGGAT TCCCGAATGG GACCTCCCGC GGCTTTTGCC GCACTCCCGT CAGGGAGGG
J23SpTT_19     GGGGATGTAA GCCGGGAAGG AAACCGACCC GTTTAGTCTG CCGCTCCCAA CAGCCCGAGT TTCTTGCCCC TTGAGTAAGG GGCAAGAAAC CTAAGCCGCC TGGTGGATGG CTCGGCTCGG GGCGCTGACG AAGGGCGTGG CAAGCTGCGA TAAGCCCCGG TGAGGCGCAG GCAGCCGTCG AGCCGGGGAT TCCCGAATGG GACCTCCCGC GGCTTTTGCC GCACTCCCGT CAGGGAGGGG
J23SpTT_63     C GGGGATGTAA GCCGGGAAGG AAACCGACCC GTTTAGTCTG CCGCTCCCAA CAGCCCGAGT TTCTTGCCCC TTGAGTAAGG GGCAAGAAAC CTAAGCCGCC TGGTGGATGG CTCGGCTCGG GGCGCTGACG AAGGGCGTGG CAAGCTGCGA TAAGCCCCGG TGAGGCGCAG GCAGCCGTCG AGCCGGGGAT TCCCGAATGG GACCTCCCGC GGCTTTTGCC GCACTCCCGT CAGGGAGG
J23SpTT_8      GGGGATGTAA GCCGGGAAGG AAACCGACCC GTTTAGTCTG CCGCTCCCAA CAGCCCGAGT TTCTTGCCCC TTGAGTAAGG GGCAAGAAAC CTAAGCCGCC TGGTGGATGG CTCGGCTCGG GGCGCTGACG AAGGGCGTGG CAAGCTGCGA TAAGCCCCGG TGAGGCGCAG GCAGCCGTCG AGCCGGGGAT TCCCGAATGG GACCTCCCGC GGCTTTTGCC GCACTCCCGT CAGGGAGGG
J23SpTT_79     C GGGGATGTAA GCCGGGAAGG AAACCGACCC GTTTAGTCTG CCGCTCCCAA CAGCCCGAGT TTCTTGCCCC TTGAGTAAGG GGCAAGAAAC CTAAGCCGCC TGGTGGATGG CTCGGCTCGG GGCGCTGACG AAGGGCGTGG CAAGCTGCGA TAAGCCCCGG TGAGGCGCAG GCAGCCGTCG AGCCGGGGAT TCCCGAATGG GACCTCCCGC GGCTTTTGCC GCACTCCCGT CAGGGAG
J23SpTT_10     GGGATGTAA GCCGGGAAGG AAACCGACCC GTTTAGTCTG CCGCTCCCAA CAGCCCGAGT TTCTTGCCCC TTGAGTAAGG GGCAAGAAAC CTAAGCCGCC TGGTGGATGG CTCGGCTCGG GGCGCTGACG AAGGGCGTGG CAAGCTGCGA TAAGCCCCGG TGAGGCGCAG GCAGCCGTCG AGCCGGGGAT TCCCGAATGG GACCTCCCGC GGCTTTTGCC GCACTCCCGT CAGGGAGGG
J23SpTT_12     GGATGTAA GCCGGGAAGG AAACCGACCC GTTTAGTCTG CCGCTCCCAA CAGCCCGAGT TTCTTGCCCC TTGAGTAAGG GGCAAGAAAC CTAAGCCGCC TGGTGGATGG CTCGGCTCGG GGCGCTGACG AAGGGCGTGG CAAGCTGCGA TAAGCCCCGG TGAGGCGCAG GCAGCCGTCG AGCCGGGGAT TCCCGAATGG GACCTCCCGC GGCTTTTGCC GCACTCCCGT CAGGGAGGGG
J23SpTT_9      GGGGATGTAA GCCGGGAAGG AAACCGACCC GTTTAGTCTG CCGCTCCCAA CAGCCCGAGT TTCTTGCCCC TTGAGTAAGG GGCAAGAAAC CTAAGCCGCC TGGTGGATGG CTCGGCTCGG GGCGCTGACG AAGGGCGTGG CAAGCTGCGA TAAGCCCCGG TGAGGCGCAG GCAGCCGTCG AGCCGGGGAT TCCCGAATGG GACCTCCCGC GGCTTTTGCC GCACTCCCGT CAGGGAG
J23SpTT_13     C GGGGATGTAA GCCGGGAAGG AAACCGACCC GTTTAGTCTG CCGCTCCCAA CAGCCCGAGT TTCTTGCCCC TTGAGTAAGG GGCAAGAAAC CTAAGCCGCC TGGTGGATGG CTCGGCTCGG GGCGCTGACG AAGGGCGTGG CAAGCTGCGA TAAGCCCCGG TGAGGCGCAG GCAGCCGTCG AGCCGGGGAT TCCCGAATGG GACCTCCCGC GGCTTTTGCC GCACTCCCGT CAGGG
J23SpTT_11     GGGATGTAA GCCGGGAAGG AAACCGACCC GTTTAGTCTG CCGCTCCCAA CAGCCCGAGT TTCTTGCCCC TTGAGTAAGG GGCAAGAAAC CTAAGCCGCC TGGTGGATGG CTCGGCTCGG GGCGCTGACG AAGGGCGTGG CAAGCTGCGA TAAGCCCCGG TGAGGCGCAG GCAGCCGTCG AGCCGGGGAT TCCCGAATGG GACCTCCCGC GGCTTTTGCC GCACTCCCGT CAGGGAG
J23SpTT_36     C GGGGATGTAA GCCGGGAAGG AAACCGACCC GTTTAGTCTG CCGCTCCCAA CAGCCCGAGT TTCTTGCCCC TTGAGTAAGG GGCAAGAAAC CTAAGCCGCC TGGTGGATGG CTCGGCTCGG GGCGCTGACG AAGGGCGTGG CAAGCTGCGA TAAGCCCCGG TGAGGCGCAG GCAGCCGTCG AGCCGGGGAT TCCCGAATGG GACCTCCCGC GGCTTTTGCC GCACTCCCGT CAGG
J23SpTT_22     C GGGGATGTAA GCCGGGAAGG AAACCGACCC GTTTAGTCTG CCGCTCCCAA CAGCCCGAGT TTCTTGCCCC TTGAGTAAGG GGCAAGAAAC CTAAGCCGCC TGGTGGATGG CTCGGCTCGG GGCGCTGACG AAGGGCGTGG CAAGCTGCGA TAAGCCCCGG TGAGGCGCAG GCAGCCGTCG AGCCGGGGAT TCCCGAATGG GACCTCCCGC GGCTTTTGCC GCACTCCCGT CAG
J23SpTT_10     GTAA GCCGGGAAGG AAACCGACCC GTTTAGTCTG CCGCTCCCAA CAGCCCGAGT TTCTTGCCCC TTGAGTAAGG GGCAAGAAAC CTAAGCCGCC TGGTGGATGG CTCGGCTCGG GGCGCTGACG AAGGGCGTGG CAAGCTGCGA TAAGCCCCGG TGAGGCGCAG GCAGCCGTCG AGCCGGGGAT TCCCGAATGG GACCTCCCGC GGCTTTTGCC GCACTCCCGT CAGGGAGGGG
J23SpTT_15     GTAA GCCGGGAAGG AAACCGACCC GTTTAGTCTG CCGCTCCCAA CAGCCCGAGT TTCTTGCCCC TTGAGTAAGG GGCAAGAAAC CTAAGCCGCC TGGTGGATGG CTCGGCTCGG GGCGCTGACG AAGGGCGTGG CAAGCTGCGA TAAGCCCCGG TGAGGCGCAG GCAGCCGTCG AGCCGGGGAT TCCCGAATGG GACCTCCCGC GGCTTTTGCC GCACTCCCGT CAGGGAGGG
J23SpTT_9      AA GCCGGGAAGG AAACCGACCC GTTTAGTCTG CCGCTCCCAA CAGCCCGAGT TTCTTGCCCC TTGAGTAAGG GGCAAGAAAC CTAAGCCGCC TGGTGGATGG CTCGGCTCGG GGCGCTGACG AAGGGCGTGG CAAGCTGCGA TAAGCCCCGG TGAGGCGCAG GCAGCCGTCG AGCCGGGGAT TCCCGAATGG GACCTCCCGC GGCTTTTGCC GCACTCCCGT CAGGGAGGGG
J23SpTT_8      GTAA GCCGGGAAGG AAACCGACCC GTTTAGTCTG CCGCTCCCAA CAGCCCGAGT TTCTTGCCCC TTGAGTAAGG GGCAAGAAAC CTAAGCCGCC TGGTGGATGG CTCGGCTCGG GGCGCTGACG AAGGGCGTGG CAAGCTGCGA TAAGCCCCGG TGAGGCGCAG GCAGCCGTCG AGCCGGGGAT TCCCGAATGG GACCTCCCGC GGCTTTTGCC GCACTCCCGT CAGGGAG
J23SpTT_9      CCGGGAAGG AAACCGACCC GTTTAGTCTG CCGCTCCCAA CAGCCCGAGT TTCTTGCCCC TTGAGTAAGG GGCAAGAAAC CTAAGCCGCC TGGTGGATGG CTCGGCTCGG GGCGCTGACG AAGGGCGTGG CAAGCTGCGA TAAGCCCCGG TGAGGCGCAG GCAGCCGTCG AGCCGGGGAT TCCCGAATGG GACCTCCCGC GGCTTTTGCC GCACTCCCGT CAGGGAGGGG G
J23SpTT_30     CCGGGAAGG AAACCGACCC GTTTAGTCTG CCGCTCCCAA CAGCCCGAGT TTCTTGCCCC TTGAGTAAGG GGCAAGAAAC CTAAGCCGCC TGGTGGATGG CTCGGCTCGG GGCGCTGACG AAGGGCGTGG CAAGCTGCGA TAAGCCCCGG TGAGGCGCAG GCAGCCGTCG AGCCGGGGAT TCCCGAATGG GACCTCCCGC GGCTTTTGCC GCACTCCCGT CAGGGAGGGG
J23SpTT_83     CCCGCA GAAGACGGGT TTGATGGGGC GGGGATGTAA GCCGGGAAGG AAACCGACCC GTTTAGTCTG CCGCTCCCAA CAGCCCGAGT TTCTTGCCCC TTGAGTAAGG GGCAAGAAAC CTAAGCCGCC TGGTGGATGG CTCGGCTCGG GGCGCTGACG AAGGGCGTGG CAAGCTGCGA TAAGCCCCGG TGAGGCGCAG GCAGCCGTCG AGCCGGGGAT TCCCGAATGG GAC
J23SpTT_36     CCGGGAAGG AAACCGACCC GTTTAGTCTG CCGCTCCCAA CAGCCCGAGT TTCTTGCCCC TTGAGTAAGG GGCAAGAAAC CTAAGCCGCC TGGTGGATGG CTCGGCTCGG GGCGCTGACG AAGGGCGTGG CAAGCTGCGA TAAGCCCCGG TGAGGCGCAG GCAGCCGTCG AGCCGGGGAT TCCCGAATGG GACCTCCCGC GGCTTTTGCC GCACTCCCGT CAGGGAGGG
J23SpTT_12     CGGGAAGG AAACCGACCC GTTTAGTCTG CCGCTCCCAA CAGCCCGAGT TTCTTGCCCC TTGAGTAAGG GGCAAGAAAC CTAAGCCGCC TGGTGGATGG CTCGGCTCGG GGCGCTGACG AAGGGCGTGG CAAGCTGCGA TAAGCCCCGG TGAGGCGCAG GCAGCCGTCG AGCCGGGGAT TCCCGAATGG GACCTCCCGC GGCTTTTGCC GCACTCCCGT CAGGGAGGGG
J23SpTT_180    CGCA GAAGACGGGT TTGATGGGGC GGGGATGTAA GCCGGGAAGG AAACCGACCC GTTTAGTCTG CCGCTCCCAA CAGCCCGAGT TTCTTGCCCC TTGAGTAAGG GGCAAGAAAC CTAAGCCGCC TGGTGGATGG CTCGGCTCGG GGCGCTGACG AAGGGCGTGG CAAGCTGCGA TAAGCCCCGG TGAGGCGCAG GCAGCCGTCG AGCCGGGGAT TCCCGAATGG GAC
J23SpTT_80     CCGCA GAAGACGGGT TTGATGGGGC GGGGATGTAA GCCGGGAAGG AAACCGACCC GTTTAGTCTG CCGCTCCCAA CAGCCCGAGT TTCTTGCCCC TTGAGTAAGG GGCAAGAAAC CTAAGCCGCC TGGTGGATGG CTCGGCTCGG GGCGCTGACG AAGGGCGTGG CAAGCTGCGA TAAGCCCCGG TGAGGCGCAG GCAGCCGTCG AGCCGGGGAT TCCCGAATGG GAC
J23SpTT_73     C GGGGATGTAA GCCGGGAAGG AAACCGACCC GTTTAGTCTG CCGCTCCCAA CAGCCCGAGT TTCTTGCCCC TTGAGTAAGG GGCAAGAAAC CTAAGCCGCC TGGTGGATGG CTCGGCTCGG GGCGCTGACG AAGGGCGTGG CAAGCTGCGA TAAGCCCCGG TGAGGCGCAG GCAGCCGTCG AGCCGGGGAT TCCCGAATGG GACCTCCCGC GGCTTTTGCC GCACTCCCGT CAGGGAGGGG GAACGCGGGG AATTGAAACA TCTTAGTACC CGCAGGAAAA GAAAG
J23SpTT_10     GGGGATGTAA GCCGGGAAGG AAACCGACCC GTTTAGTCTG CCGCTCCCAA CAGCCCGAGT TTCTTGCCCC TTGAGTAAGG GGCAAGAAAC CTAAGCCGCC TGGTGGATGG CTCGGCTCGG GGCGCTGACG AAGGGCGTGG CAAGCTGCGA TAAGCCCCGG TGAGGCGCAG GCAGCCGTCG AGCCGGGGAT TCCCGAATGG GACCTCCCGC GGCTTTTGCC GCACTCCCGT CAGGGAGGGG GAACGCGGGG AATTGAAACA TCTTAGTACC CGCAGGAAAA GAAAG
J23SpTT_8      GGGATGTAA GCCGGGAAGG AAACCGACCC GTTTAGTCTG CCGCTCCCAA CAGCCCGAGT TTCTTGCCCC TTGAGTAAGG GGCAAGAAAC CTAAGCCGCC TGGTGGATGG CTCGGCTCGG GGCGCTGACG AAGGGCGTGG CAAGCTGCGA TAAGCCCCGG TGAGGCGCAG GCAGCCGTCG AGCCGGGGAT TCCCGAATGG GACCTCCCGC GGCTTTTGCC GCACTCCCGT CAGGGAGGGG GAACGCGGGG AATTGAAACA TCTTAGTACC CGCAGGAAAA GAAAG
J23SpTT_23     CCGGGAAGG AAACCGACCC GTTTAGTCTG CCGCTCCCAA CAGCCCGAGT TTCTTGCCCC TTGAGTAAGG GGCAAGAAAC CTAAGCCGCC TGGTGGATGG CTCGGCTCGG GGCGCTGACG AAGGGCGTGG CAAGCTGCGA TAAGCCCCGG TGAGGCGCAG GCAGCCGTCG AGCCGGGGAT TCCCGAATGG GACCTCCCGC GGCTTTTGCC GCACTCCCGT CAGGGAGGGG GAACGCGGGG AATTGAAACA TCTTAGTACC CGCAGGAAAA GAAAG
J23SpTT_11     CGGGAAGG AAACCGACCC GTTTAGTCTG CCGCTCCCAA CAGCCCGAGT TTCTTGCCCC TTGAGTAAGG GGCAAGAAAC CTAAGCCGCC TGGTGGATGG CTCGGCTCGG GGCGCTGACG AAGGGCGTGG CAAGCTGCGA TAAGCCCCGG TGAGGCGCAG GCAGCCGTCG AGCCGGGGAT TCCCGAATGG GACCTCCCGC GGCTTTTGCC GCACTCCCGT CAGGGAGGGG GAACGCGGGG AATTGAAACA TCTTAGTACC CGCAGGAAAA GAAAG
J23SpTT_19     GGG AAACCGACCC GTTTAGTCTG CCGCTCCCAA CAGCCCGAGT TTCTTGCCCC TTGAGTAAGG GGCAAGAAAC CTAAGCCGCC TGGTGGATGG CTCGGCTCGG GGCGCTGACG AAGGGCGTGG CAAGCTGCGA TAAGCCCCGG TGAGGCGCAG GCAGCCGTCG AGCCGGGGAT TCCCGAATGG GACCTCCCGC GGCTTTTGCC GCACTCCCGT CAGGGAGGGG GAACGCGGGG AATTGAAACA TCTTAGTACC CGCAGGAAAA GAAAG
J23SpTT_11     CGACCC GTTTAGTCTG CCGCTCCCAA CAGCCCGAGT TTCTTGCCCC TTGAGTAAGG GGCAAGAAAC CTAAGCCGCC TGGTGGATGG CTCGGCTCGG GGCGCTGACG AAGGGCGTGG CAAGCTGCGA TAAGCCCCGG TGAGGCGCAG GCAGCCGTCG AGCCGGGGAT TCCCGAATGG GACCTCCCGC GGCTTTTGCC GCACTCCCGT CAGGGAGGGG GAACGCGGGG AATTGAAACA TCTTAGTACC CGCAGGAAAA GAAAG
J23SpTT_14     CCC GTTTAGTCTG CCGCTCCCAA CAGCCCGAGT TTCTTGCCCC TTGAGTAAGG GGCAAGAAAC CTAAGCCGCC TGGTGGATGG CTCGGCTCGG GGCGCTGACG AAGGGCGTGG CAAGCTGCGA TAAGCCCCGG TGAGGCGCAG GCAGCCGTCG AGCCGGGGAT TCCCGAATGG GACCTCCCGC GGCTTTTGCC GCACTCCCGT CAGGGAGGGG GAACGCGGGG AATTGAAACA TCTTAGTACC CGCAGGAAAA GAAAG
J23SpTT_8      GTTTAGTCTG CCGCTCCCAA CAGCCCGAGT TTCTTGCCCC TTGAGTAAGG GGCAAGAAAC CTAAGCCGCC TGGTGGATGG CTCGGCTCGG GGCGCTGACG AAGGGCGTGG CAAGCTGCGA TAAGCCCCGG TGAGGCGCAG GCAGCCGTCG AGCCGGGGAT TCCCGAATGG GACCTCCCGC GGCTTTTGCC GCACTCCCGT CAGGGAGGGG GAACGCGGGG AATTGAAACA TCTTAGTACC CGCAGGAAAA GAAAG
J23SpTT_17     CTCCCAA CAGCCCGAGT TTCTTGCCCC TTGAGTAAGG GGCAAGAAAC CTAAGCCGCC TGGTGGATGG CTCGGCTCGG GGCGCTGACG AAGGGCGTGG CAAGCTGCGA TAAGCCCCGG TGAGGCGCAG GCAGCCGTCG AGCCGGGGAT TCCCGAATGG GACCTCCCGC GGCTTTTGCC GCACTCCCGT CAGGGAGGGG GAACGCGGGG AATTGAAACA TCTTAGTACC CGCAGGAAAA GAAAG
J23SpTT_15     CCCAA CAGCCCGAGT TTCTTGCCCC TTGAGTAAGG GGCAAGAAAC CTAAGCCGCC TGGTGGATGG CTCGGCTCGG GGCGCTGACG AAGGGCGTGG CAAGCTGCGA TAAGCCCCGG TGAGGCGCAG GCAGCCGTCG AGCCGGGGAT TCCCGAATGG GACCTCCCGC GGCTTTTGCC GCACTCCCGT CAGGGAGGGG GAACGCGGGG AATTGAAACA TCTTAGTACC CGCAGGAAAA GAAAG
J23SpTT_10     CCAA CAGCCCGAGT TTCTTGCCCC TTGAGTAAGG GGCAAGAAAC CTAAGCCGCC TGGTGGATGG CTCGGCTCGG GGCGCTGACG AAGGGCGTGG CAAGCTGCGA TAAGCCCCGG TGAGGCGCAG GCAGCCGTCG AGCCGGGGAT TCCCGAATGG GACCTCCCGC GGCTTTTGCC GCACTCCCGT CAGGGAGGGG GAACGCGGGG AATTGAAACA TCTTAGTACC CGCAGGAAAA GAAAG
J23SpTT_17     CCGGGAAGG AAACCGACCC GTTTAGTCTG CCGCTCCCAA CAGCCCGAGT TTCTTGCCCC TTGAGTAAGG GGCAAGAAAC CTAAGCCGCC TGGTGGATGG CTCGGCTCGG GGCGCTGACG AAGGGCGTGG CAAGCTGCGA TAAGCCCCGG TGAGGCGCAG GCAGCCGTCG AGCCGGGGAT TCCCGAATGG GACCTCCCGC GGCTTTTGCC GCACTCCCGT CAGGGAGG
J23SpTT_18     CCGGGAAGG AAACCGACCC GTTTAGTCTG CCGCTCCCAA CAGCCCGAGT TTCTTGCCCC TTGAGTAAGG GGCAAGAAAC CTAAGCCGCC TGGTGGATGG CTCGGCTCGG GGCGCTGACG AAGGGCGTGG CAAGCTGCGA TAAGCCCCGG TGAGGCGCAG GCAGCCGTCG AGCCGGGGAT TCCCGAATGG GACCTCCCGC GGCTTTTGCC GCACTCCCGT CAGGGAG
J23SpTT_68     CA GAAGACGGGT TTGATGGGGC GGGGATGTAA GCCGGGAAGG AAACCGACCC GTTTAGTCTG CCGCTCCCAA CAGCCCGAGT TTCTTGCCCC TTGAGTAAGG GGCAAGAAAC CTAAGCCGCC TGGTGGATGG CTCGGCTCGG GGCGCTGACG AAGGGCGTGG CAAGCTGCGA TAAGCCCCGG TGAGGCGCAG GCAGCCGTCG AGCCGGGGAT TCCCGAATGG GAC
J23SpTT_17     GCA GAAGACGGGT TTGATGGGGC GGGGATGTAA GCCGGGAAGG AAACCGACCC GTTTAGTCTG CCGCTCCCAA CAGCCCGAGT TTCTTGCCCC TTGAGTAAGG GGCAAGAAAC CTAAGCCGCC TGGTGGATGG CTCGGCTCGG GGCGCTGACG AAGGGCGTGG CAAGCTGCGA TAAGCCCCGG TGAGGCGCAG GCAGCCGTCG AGCCGGGGAT TCCCGAATGG GAC
J23SpTT_46     CGCA GAAGACGGGT TTGATGGGGC GGGGATGTAA GCCGGGAAGG AAACCGACCC GTTTAGTCTG CCGCTCCCAA CAGCCCGAGT TTCTTGCCCC TTGAGTAAGG GGCAAGAAAC CTAAGCCGCC TGGTGGATGG CTCGGCTCGG GGCGCTGACG AAGGGCGTGG CAAGCTGCGA TAAGCCCCGG TGAGGCGCAG GCAGCCGTCG AGCCGGGGAT TCCCGAATGG G
J23SpTT_20     CCGCA GAAGACGGGT TTGATGGGGC GGGGATGTAA GCCGGGAAGG AAACCGACCC GTTTAGTCTG CCGCTCCCAA CAGCCCGAGT TTCTTGCCCC TTGAGTAAGG GGCAAGAAAC CTAAGCCGCC TGGTGGATGG CTCGGCTCGG GGCGCTGACG AAGGGCGTGG CAAGCTGCGA TAAGCCCCGG TGAGGCGCAG GCAGCCGTCG AGCCGGGGAT TCCCGAATGG GA
J23SpTT_16     CCGCA GAAGACGGGT TTGATGGGGC GGGGATGTAA GCCGGGAAGG AAACCGACCC GTTTAGTCTG CCGCTCCCAA CAGCCCGAGT TTCTTGCCCC TTGAGTAAGG GGCAAGAAAC CTAAGCCGCC TGGTGGATGG CTCGGCTCGG GGCGCTGACG AAGGGCGTGG CAAGCTGCGA TAAGCCCCGG TGAGGCGCAG GCAGCCGTCG AGCCGGGGAT TCCCGAATGG G
J23SpTT_14     CCCGCA GAAGACGGGT TTGATGGGGC GGGGATGTAA GCCGGGAAGG AAACCGACCC GTTTAGTCTG CCGCTCCCAA CAGCCCGAGT TTCTTGCCCC TTGAGTAAGG GGCAAGAAAC CTAAGCCGCC TGGTGGATGG CTCGGCTCGG GGCGCTGACG AAGGGCGTGG CAAGCTGCGA TAAGCCCCGG TGAGGCGCAG GCAGCCGTCG AGCCGGGGAT TCCCGAATGG G
J23SpTT_13     CGCA GAAGACGGGT TTGATGGGGC GGGGATGTAA GCCGGGAAGG AAACCGACCC GTTTAGTCTG CCGCTCCCAA CAGCCCGAGT TTCTTGCCCC TTGAGTAAGG GGCAAGAAAC CTAAGCCGCC TGGTGGATGG CTCGGCTCGG GGCGCTGACG AAGGGCGTGG CAAGCTGCGA TAAGCCCCGG TGAGGCGCAG GCAGCCGTCG AGCCGGGGAT TCCCGAATGG GA
J23SpTT_10     CCGCA GAAGACGGGT TTGATGGGGC GGGGATGTAA GCCGGGAAGG AAACCGACCC GTTTAGTCTG CCGCTCCCAA CAGCCCGAGT TTCTTGCCCC TTGAGTAAGG GGCAAGAAAC CTAAGCCGCC TGGTGGATGG CTCGGCTCGG GGCGCTGACG AAGGGCGTGG CAAGCTGCGA TAAGCCCCGG TGAGGCGCAG GCAGCCGTCG AGCCGGGGAT TCCCGAATGG
J23SpTT_10     CGCA GAAGACGGGT TTGATGGGGC GGGGATGTAA GCCGGGAAGG AAACCGACCC GTTTAGTCTG CCGCTCCCAA CAGCCCGAGT TTCTTGCCCC TTGAGTAAGG GGCAAGAAAC CTAAGCCGCC TGGTGGATGG CTCGGCTCGG GGCGCTGACG AAGGGCGTGG CAAGCTGCGA TAAGCCCCGG TGAGGCGCAG GCAGCCGTCG AGCCGGGGAT TCCCGAATGG
J23SpTT_29     A GAAGACGGGT TTGATGGGGC GGGGATGTAA GCCGGGAAGG AAACCGACCC GTTTAGTCTG CCGCTCCCAA CAGCCCGAGT TTCTTGCCCC TTGAGTAAGG GGCAAGAAAC CTAAGCCGCC TGGTGGATGG CTCGGCTCGG GGCGCTGACG AAGGGCGTGG CAAGCTGCGA TAAGCCCCGG TGAGGCGCAG GCAGCCGTCG AGCCGGGGAT TCCCGAATGG GAC
J23SpTT_21     CA GAAGACGGGT TTGATGGGGC GGGGATGTAA GCCGGGAAGG AAACCGACCC GTTTAGTCTG CCGCTCCCAA CAGCCCGAGT TTCTTGCCCC TTGAGTAAGG GGCAAGAAAC CTAAGCCGCC TGGTGGATGG CTCGGCTCGG GGCGCTGACG AAGGGCGTGG CAAGCTGCGA TAAGCCCCGG TGAGGCGCAG GCAGCCGTCG AGCCGGGGAT TCCCGAATGG G
J23SpTT_34     CGCA GAAGACGGGT TTGATGGGGC GGGGATGTAA GCCGGGAAGG AAACCGACCC GTTTAGTCTG CCGCTCCCAA CAGCCCGAGT TTCTTGCCCC TTGAGTAAGG GGCAAGAAAC CTAAGCCGCC TGGTGGATGG CTCGGCTCGG GGCGCTGACG AAGGGCGTGG CAAGCTGCGA TAAGCCCCGG TGAGGCGCAG GCAGCCGTCG AGCCGGGGAT TCCCGAAT
J23SpTT_16     CGCA GAAGACGGGT TTGATGGGGC GGGGATGTAA GCCGGGAAGG AAACCGACCC GTTTAGTCTG CCGCTCCCAA CAGCCCGAGT TTCTTGCCCC TTGAGTAAGG GGCAAGAAAC CTAAGCCGCC TGGTGGATGG CTCGGCTCGG GGCGCTGACG AAGGGCGTGG CAAGCTGCGA TAAGCCCCGG TGAGGCGCAG GCAGCCGTCG AGCCGGGGAT TCCCGAATG
J23SpTT_16     CCGCA GAAGACGGGT TTGATGGGGC GGGGATGTAA GCCGGGAAGG AAACCGACCC GTTTAGTCTG CCGCTCCCAA CAGCCCGAGT TTCTTGCCCC TTGAGTAAGG GGCAAGAAAC CTAAGCCGCC TGGTGGATGG CTCGGCTCGG GGCGCTGACG AAGGGCGTGG CAAGCTGCGA TAAGCCCCGG TGAGGCGCAG GCAGCCGTCG AGCCGGGGAT TCCCGAAT
J23SpTT_10     GCA GAAGACGGGT TTGATGGGGC GGGGATGTAA GCCGGGAAGG AAACCGACCC GTTTAGTCTG CCGCTCCCAA CAGCCCGAGT TTCTTGCCCC TTGAGTAAGG GGCAAGAAAC CTAAGCCGCC TGGTGGATGG CTCGGCTCGG GGCGCTGACG AAGGGCGTGG CAAGCTGCGA TAAGCCCCGG TGAGGCGCAG GCAGCCGTCG AGCCGGGGAT TCCCGAAT
```

```
J23SpTT_15 GGG AAACCGACCC GTTTAGTCTG CCGCTCCCAA CAGCCCGAGT TTCTTGCCCC TTGAGTAAGG GGCAAGAAAC CTAAGCCGCC TGGTGGATGG CTCGGCTCGG GGCGCTGACG AAGGGCGTGG CAAGCTGCGA TAAGCCCCGG TGAGGCGCAG GCAGCCGTCG AGCCGGGGAT TCCCGAATGG GACCTCCCGC GGCTTTTGCC GCACTCCCGT CAGGGAGGGG
J23SpTT_8 GG AAACCGACCC GTTTAGTCTG CCGCTCCCAA CAGCCCGAGT TTCTTGCCCC TTGAGTAAGG GGCAAGAAAC CTAAGCCGCC TGGTGGATGG CTCGGCTCGG GGCGCTGACG AAGGGCGTGG CAAGCTGCGA TAAGCCCCGG TGAGGCGCAG GCAGCCGTCG AGCCGGGGAT TCCCGAATGG GACCTCCCGC GGCTTTTGCC GCACTCCCGT CAGGGAGGGG
J23SpTT_12 GGG AAACCGACCC GTTTAGTCTG CCGCTCCCAA CAGCCCGAGT TTCTTGCCCC TTGAGTAAGG GGCAAGAAAC CTAAGCCGCC TGGTGGATGG CTCGGCTCGG GGCGCTGACG AAGGGCGTGG CAAGCTGCGA TAAGCCCCGG TGAGGCGCAG GCAGCCGTCG AGCCGGGGAT TCCCGAATGG GACCTCCCGC GGCTTTTGCC GCACTCCCGT CAGGGAGGG
J23SpTT_14 GGG AAACCGACCC GTTTAGTCTG CCGCTCCCAA CAGCCCGAGT TTCTTGCCCC TTGAGTAAGG GGCAAGAAAC CTAAGCCGCC TGGTGGATGG CTCGGCTCGG GGCGCTGACG AAGGGCGTGG CAAGCTGCGA TAAGCCCCGG TGAGGCGCAG GCAGCCGTCG AGCCGGGGAT TCCCGAATGG GACCTCCCGC GGCTTTTGCC GCACTCCCGT CAGGGAGG
J23SpTT_11 CCGGGAAGGG AAACCGACCC GTTTAGTCTG CCGCTCCCAA CAGCCCGAGT TTCTTGCCCC TTGAGTAAGG GGCAAGAAAC CTAAGCCGCC TGGTGGATGG CTCGGCTCGG GGCGCTGACG AAGGGCGTGG CAAGCTGCGA TAAGCCCCGG TGAGGCGCAG GCAGCCGTCG AGCCGGGGAT TCCCGAATGG GACCTCCCGC GGCTTTTGCC GCACTCCCGT CAGG
J23SpTT_9 CA GAAGACGGGT TTGATGGGGC GGGGATGTAA GCCGGGAAGGG AAACCGACCC GTTTAGTCTG CCGCTCCCAA CAGCCCGAGT TTCTTGCCCC TTGAGTAAGG GGCAAGAAAC CTAAGCCGCC TGGTGGATGG CTCGGCTCGG GGCGCTGACG AAGGGCGTGG CAAGCTGCGA TAAGCCCCGG TGAGGCGCAG GCAGCCGTCG AGCCGGGGAT TCCCGAAT
J23SpTT_43 CGGGT TTGATGGGGC GGGGATGTAA GCCGGGAAGGG AAACCGACCC GTTTAGTCTG CCGCTCCCAA CAGCCCGAGT TTCTTGCCCC TTGAGTAAGG GGCAAGAAAC CTAAGCCGCC TGGTGGATGG CTCGGCTCGG GGCGCTGACG AAGGGCGTGG CAAGCTGCGA TAAGCCCCGG TGAGGCGCAG GCAGCCGTCG AGCCGGGGAT TCCCGAATGG GAC
J23SpTT_31 GACGGGT TTGATGGGGC GGGGATGTAA GCCGGGAAGGG AAACCGACCC GTTTAGTCTG CCGCTCCCAA CAGCCCGAGT TTCTTGCCCC TTGAGTAAGG GGCAAGAAAC CTAAGCCGCC TGGTGGATGG CTCGGCTCGG GGCGCTGACG AAGGGCGTGG CAAGCTGCGA TAAGCCCCGG TGAGGCGCAG GCAGCCGTCG AGCCGGGGAT TCCCGAATGG GAC
J23SpTT_17 ACGGGT TTGATGGGGC GGGGATGTAA GCCGGGAAGGG AAACCGACCC GTTTAGTCTG CCGCTCCCAA CAGCCCGAGT TTCTTGCCCC TTGAGTAAGG GGCAAGAAAC CTAAGCCGCC TGGTGGATGG CTCGGCTCGG GGCGCTGACG AAGGGCGTGG CAAGCTGCGA TAAGCCCCGG TGAGGCGCAG GCAGCCGTCG AGCCGGGGAT TCCCGAATGG GAC
J23SpTT_12 CGGGT TTGATGGGGC GGGGATGTAA GCCGGGAAGGG AAACCGACCC GTTTAGTCTG CCGCTCCCAA CAGCCCGAGT TTCTTGCCCC TTGAGTAAGG GGCAAGAAAC CTAAGCCGCC TGGTGGATGG CTCGGCTCGG GGCGCTGACG AAGGGCGTGG CAAGCTGCGA TAAGCCCCGG TGAGGCGCAG GCAGCCGTCG AGCCGGGGAT TCCCGAATGG GA
J23SpTT_10 CGGGT TTGATGGGGC GGGGATGTAA GCCGGGAAGGG AAACCGACCC GTTTAGTCTG CCGCTCCCAA CAGCCCGAGT TTCTTGCCCC TTGAGTAAGG GGCAAGAAAC CTAAGCCGCC TGGTGGATGG CTCGGCTCGG GGCGCTGACG AAGGGCGTGG CAAGCTGCGA TAAGCCCCGG TGAGGCGCAG GCAGCCGTCG AGCCGGGGAT TCCCGAATGG G
J23SpTT_13 GT TTGATGGGGC GGGGATGTAA GCCGGGAAGGG AAACCGACCC GTTTAGTCTG CCGCTCCCAA CAGCCCGAGT TTCTTGCCCC TTGAGTAAGG GGCAAGAAAC CTAAGCCGCC TGGTGGATGG CTCGGCTCGG GGCGCTGACG AAGGGCGTGG CAAGCTGCGA TAAGCCCCGG TGAGGCGCAG GCAGCCGTCG AGCCGGGGAT TCCCGAATGG GAC
J23SpTT_9 GACGGGT TTGATGGGGC GGGGATGTAA GCCGGGAAGGG AAACCGACCC GTTTAGTCTG CCGCTCCCAA CAGCCCGAGT TTCTTGCCCC TTGAGTAAGG GGCAAGAAAC CTAAGCCGCC TGGTGGATGG CTCGGCTCGG GGCGCTGACG AAGGGCGTGG CAAGCTGCGA TAAGCCCCGG TGAGGCGCAG GCAGCCGTCG AGCCGGGGAT TCCCGAAT
J23SpTT_11 CGCA GAAGACGGGT TTGATGGGGC GGGGATGTAA GCCGGGAAGGG AAACCGACCC GTTTAGTCTG CCGCTCCCAA CAGCCCGAGT TTCTTGCCCC TTGAGTAAGG GGCAAGAAAC CTAAGCCGCC TGGTGGATGG CTCGGCTCGG GGCGCTGACG AAGGGCGTGG CAAGCTGCGA TAAGCCCCGG TGAGGCGCAG GCAGCCGTCG AGCCGGG
J23SpTT_9 CCGCA GAAGACGGGT TTGATGGGGC GGGGATGTAA GCCGGGAAGGG AAACCGACCC GTTTAGTCTG CCGCTCCCAA CAGCCCGAGT TTCTTGCCCC TTGAGTAAGG GGCAAGAAAC CTAAGCCGCC TGGTGGATGG CTCGGCTCGG GGCGCTGACG AAGGGCGTGG CAAGCTGCGA TAAGCCCCGG TGAGGCGCAG GCAGCCGTCG AGCCGGG
J23SpTT_33 C GGGGATGTAA GCCGGGAAGGG AAACCGACCC GTTTAGTCTG CCGCTCCCAA CAGCCCGAGT TTCTTGCCCC TTGAGTAAGG GGCAAGAAAC CTAAGCCGCC TGGTGGATGG CTCGGCTCGG GGCGCTGACG AAGGGCGTGG CAAGCTGCGA TAAGCCCCGG TGAGGCGCAG GCAGCCGTCG AGCCGGGGAT TCCCGAATGG GAC
J23SpTT_9 CTG CCGCTCCCAA CAGCCCGAGT TTCTTGCCCC TTGAGTAAGG GGCAAGAAAC CTAAGCCGCC TGGTGGATGG CTCGGCTCGG GGCGCTGACG AAGGGCGTGG CAAGCTGCGA TAAGCCCCGG TGAGGCGCAG GCAGCCGTCG AGCCGGGGAT TCCCGAATGG GACCTCCCGC GGCTTTTGCC GCACTCCCGT CAGGGAGGGG
J23SpTT_18 CGCA GAAGACGGGT TTGATGGGGC GGGGATGTAA GCCGGGAAGGG AAACCGACCC GTTTAGTCTG CCGCTCCCAA CAGCCCGAGT TTCTTGCCCC TTGAGTAAGG GGCAAGAAAC CTAAGCCGCC TGGTGGATGG CTCGGCTCGG GGCGCTGACG AAGGGCGTGG CAAGCTGCGA TAAGCCCCGG TGAGGCGCAG GCAGCCG
J23SpTT_12 CCCGCA GAAGACGGGT TTGATGGGGC GGGGATGTAA GCCGGGAAGGG AAACCGACCC GTTTAGTCTG CCGCTCCCAA CAGCCCGAGT TTCTTGCCCC TTGAGTAAGG GGCAAGAAAC CTAAGCCGCC TGGTGGATGG CTCGGCTCGG GGCGCTGACG AAGGGCGTGG CAAGCTGCGA TAAGCCCCGG TGAGGCGCAG GCAGCCG
J23SpTT_9 CCGCA GAAGACGGGT TTGATGGGGC GGGGATGTAA GCCGGGAAGGG AAACCGACCC GTTTAGTCTG CCGCTCCCAA CAGCCCGAGT TTCTTGCCCC TTGAGTAAGG GGCAAGAAAC CTAAGCCGCC TGGTGGATGG CTCGGCTCGG GGCGCTGACG AAGGGCGTGG CAAGCTGCGA TAAGCCCCGG TGAGGCGCAG GCAGCCG
J23SpTT_9 CGCA GAAGACGGGT TTGATGGGGC GGGGATGTAA GCCGGGAAGGG AAACCGACCC GTTTAGTCTG CCGCTCCCAA CAGCCCGAGT TTCTTGCCCC TTGAGTAAGG GGCAAGAAAC CTAAGCCGCC TGGTGGATGG CTCGGCTCGG GGCGCTGACG AAGGGCGTGG CAAGCTGCGA TAAGCCCCGG TGAGGCGCAG GCAGCC
J23SpTT_13 CCGCA GAAGACGGGT TTGATGGGGC GGGGATGTAA GCCGGGAAGGG AAACCGACCC GTTTAGTCTG CCGCTCCCAA CAGCCCGAGT TTCTTGCCCC TTGAGTAAGG GGCAAGAAAC CTAAGCCGCC TGGTGGATGG CTCGGCTCGG GGCGCTGACG AAGGGCGTGG CAAGCTGCGA TAAGCCCCGG TGAGGCGCAG GCAGC
J23SpTT_11 CGCA GAAGACGGGT TTGATGGGGC GGGGATGTAA GCCGGGAAGGG AAACCGACCC GTTTAGTCTG CCGCTCCCAA CAGCCCGAGT TTCTTGCCCC TTGAGTAAGG GGCAAGAAAC CTAAGCCGCC TGGTGGATGG CTCGGCTCGG GGCGCTGACG AAGGGCGTGG CAAGCTGCGA TAAGCCCCGG TGAGGCGCAG GCAGC
J23SpTT_11 CA GAAGACGGGT TTGATGGGGC GGGGATGTAA GCCGGGAAGGG AAACCGACCC GTTTAGTCTG CCGCTCCCAA CAGCCCGAGT TTCTTGCCCC TTGAGTAAGG GGCAAGAAAC CTAAGCCGCC TGGTGGATGG CTCGGCTCGG GGCGCTGACG AAGGGCGTGG CAAGCTGCGA TAAGCCCCGG TGAGGCGCAG GCAGCCG
J23SpTT_14 CGCA GAAGACGGGT TTGATGGGGC GGGGATGTAA GCCGGGAAGGG AAACCGACCC GTTTAGTCTG CCGCTCCCAA CAGCCCGAGT TTCTTGCCCC TTGAGTAAGG GGCAAGAAAC CTAAGCCGCC TGGTGGATGG CTCGGCTCGG GGCGCTGACG AAGGGCGTGG CAAGCTGCGA TAAGCCCCGG TGAGGCGCAG GCAG
J23SpTT_34 CGCA GAAGACGGGT TTGATGGGGC GGGGATGTAA GCCGGGAAGGG AAACCGACCC GTTTAGTCTG CCGCTCCCAA CAGCCCGAGT TTCTTGCCCC TTGAGTAAGG GGCAAGAAAC CTAAGCCGCC TGGTGGATGG CTCGGCTCGG GGCGCTGACG AAGGGCGTGG CAAGCTGCGA TAAGCCCCGG TGAGGCGCAG
J23SpTT_25 CCGCA GAAGACGGGT TTGATGGGGC GGGGATGTAA GCCGGGAAGGG AAACCGACCC GTTTAGTCTG CCGCTCCCAA CAGCCCGAGT TTCTTGCCCC TTGAGTAAGG GGCAAGAAAC CTAAGCCGCC TGGTGGATGG CTCGGCTCGG GGCGCTGACG AAGGGCGTGG CAAGCTGCGA TAAGCCCCGG TGAGGCGCAG
J23SpTT_18 CCCGCA GAAGACGGGT TTGATGGGGC GGGGATGTAA GCCGGGAAGGG AAACCGACCC GTTTAGTCTG CCGCTCCCAA CAGCCCGAGT TTCTTGCCCC TTGAGTAAGG GGCAAGAAAC CTAAGCCGCC TGGTGGATGG CTCGGCTCGG GGCGCTGACG AAGGGCGTGG CAAGCTGCGA TAAGCCCCGG TGAGGCGCAG
J23SpTT_12 CGCA GAAGACGGGT TTGATGGGGC GGGGATGTAA GCCGGGAAGGG AAACCGACCC GTTTAGTCTG CCGCTCCCAA CAGCCCGAGT TTCTTGCCCC TTGAGTAAGG GGCAAGAAAC CTAAGCCGCC TGGTGGATGG CTCGGCTCGG GGCGCTGACG AAGGGCGTGG CAAGCTGCGA TAAGCCCCGG TGAGGCGCAG G
J23SpTT_11 CGCA GAAGACGGGT TTGATGGGGC GGGGATGTAA GCCGGGAAGGG AAACCGACCC GTTTAGTCTG CCGCTCCCAA CAGCCCGAGT TTCTTGCCCC TTGAGTAAGG GGCAAGAAAC CTAAGCCGCC TGGTGGATGG CTCGGCTCGG GGCGCTGACG AAGGGCGTGG CAAGCTGCGA TAAGCCCCGG TGAGGCGCAG GC
J23SpTT_11 CA GAAGACGGGT TTGATGGGGC GGGGATGTAA GCCGGGAAGGG AAACCGACCC GTTTAGTCTG CCGCTCCCAA CAGCCCGAGT TTCTTGCCCC TTGAGTAAGG GGCAAGAAAC CTAAGCCGCC TGGTGGATGG CTCGGCTCGG GGCGCTGACG AAGGGCGTGG CAAGCTGCGA TAAGCCCCGG TGAGGCGCAG GCAG
J23SpTT_10 CCGGGAAGGG AAACCGACCC GTTTAGTCTG CCGCTCCCAA CAGCCCGAGT TTCTTGCCCC TTGAGTAAGG GGCAAGAAAC CTAAGCCGCC TGGTGGATGG CTCGGCTCGG GGCGCTGACG AAGGGCGTGG CAAGCTGCGA TAAGCCCCGG TGAGGCGCAG GCAGCCGTCG AGCCGGGGAT TCCCGAATGG GAC
J23SpTT_12 GACGGGT TTGATGGGGC GGGGATGTAA GCCGGGAAGGG AAACCGACCC GTTTAGTCTG CCGCTCCCAA CAGCCCGAGT TTCTTGCCCC TTGAGTAAGG GGCAAGAAAC CTAAGCCGCC TGGTGGATGG CTCGGCTCGG GGCGCTGACG AAGGGCGTGG CAAGCTGCGA TAAGCCCCGG TGAGGCGCAG GCAGCCG
J23SpTT_11 CGGGT TTGATGGGGC GGGGATGTAA GCCGGGAAGGG AAACCGACCC GTTTAGTCTG CCGCTCCCAA CAGCCCGAGT TTCTTGCCCC TTGAGTAAGG GGCAAGAAAC CTAAGCCGCC TGGTGGATGG CTCGGCTCGG GGCGCTGACG AAGGGCGTGG CAAGCTGCGA TAAGCCCCGG TGAGGCGCAG GCAGCCG
J23SpTT_135 CCCGCA GAAGACGGGT TTGATGGGGC GGGGATGTAA GCCGGGAAGGG AAACCGACCC GTTTAGTCTG CCGCTCCCAA CAGCCCGAGT TTCTTGCCCC TTGAGTAAGG GGCAAGAAAC CTAAGCCGCC TGGTGGATGG CTCGGCTCGG GGCGCTGACG AAGGGCGTGG CAAGCTGCGA TAAGCCCCGG TGAG
J23SpTT_57 CGCA GAAGACGGGT TTGATGGGGC GGGGATGTAA GCCGGGAAGGG AAACCGACCC GTTTAGTCTG CCGCTCCCAA CAGCCCGAGT TTCTTGCCCC TTGAGTAAGG GGCAAGAAAC CTAAGCCGCC TGGTGGATGG CTCGGCTCGG GGCGCTGACG AAGGGCGTGG CAAGCTGCGA TAAGCCCCGG TGAGGCG
J23SpTT_29 CCGCA GAAGACGGGT TTGATGGGGC GGGGATGTAA GCCGGGAAGGG AAACCGACCC GTTTAGTCTG CCGCTCCCAA CAGCCCGAGT TTCTTGCCCC TTGAGTAAGG GGCAAGAAAC CTAAGCCGCC TGGTGGATGG CTCGGCTCGG GGCGCTGACG AAGGGCGTGG CAAGCTGCGA TAAGCCCCGG TGAGGCG
J23SpTT_23 CCCGCA GAAGACGGGT TTGATGGGGC GGGGATGTAA GCCGGGAAGGG AAACCGACCC GTTTAGTCTG CCGCTCCCAA CAGCCCGAGT TTCTTGCCCC TTGAGTAAGG GGCAAGAAAC CTAAGCCGCC TGGTGGATGG CTCGGCTCGG GGCGCTGACG AAGGGCGTGG CAAGCTGCGA TAAGCCCCGG TGAGGCG
J23SpTT_17 CCCGCA GAAGACGGGT TTGATGGGGC GGGGATGTAA GCCGGGAAGGG AAACCGACCC GTTTAGTCTG CCGCTCCCAA CAGCCCGAGT TTCTTGCCCC TTGAGTAAGG GGCAAGAAAC CTAAGCCGCC TGGTGGATGG CTCGGCTCGG GGCGCTGACG AAGGGCGTGG CAAGCTGCGA TAAGCCCCGG TGAGGC
J23SpTT_33 CCCGCA GAAGACGGGT TTGATGGGGC GGGGATGTAA GCCGGGAAGGG AAACCGACCC GTTTAGTCTG CCGCTCCCAA CAGCCCGAGT TTCTTGCCCC TTGAGTAAGG GGCAAGAAAC CTAAGCCGCC TGGTGGATGG CTCGGCTCGG GGCGCTGACG AAGGGCGTGG CAAGCTGCGA TAAGCCCCGG TGAGG
J23SpTT_30 CCGCA GAAGACGGGT TTGATGGGGC GGGGATGTAA GCCGGGAAGGG AAACCGACCC GTTTAGTCTG CCGCTCCCAA CAGCCCGAGT TTCTTGCCCC TTGAGTAAGG GGCAAGAAAC CTAAGCCGCC TGGTGGATGG CTCGGCTCGG GGCGCTGACG AAGGGCGTGG CAAGCTGCGA TAAGCCCCGG TGAGGC
J23SpTT_63 CGCA GAAGACGGGT TTGATGGGGC GGGGATGTAA GCCGGGAAGGG AAACCGACCC GTTTAGTCTG CCGCTCCCAA CAGCCCGAGT TTCTTGCCCC TTGAGTAAGG GGCAAGAAAC CTAAGCCGCC TGGTGGATGG CTCGGCTCGG GGCGCTGACG AAGGGCGTGG CAAGCTGCGA TAAGCCCCGG TGAGGC
J23SpTT_30 CCGCA GAAGACGGGT TTGATGGGGC GGGGATGTAA GCCGGGAAGGG AAACCGACCC GTTTAGTCTG CCGCTCCCAA CAGCCCGAGT TTCTTGCCCC TTGAGTAAGG GGCAAGAAAC CTAAGCCGCC TGGTGGATGG CTCGGCTCGG GGCGCTGACG AAGGGCGTGG CAAGCTGCGA TAAGCCCCGG TGAGG
J23SpTT_113 CCGCA GAAGACGGGT TTGATGGGGC GGGGATGTAA GCCGGGAAGGG AAACCGACCC GTTTAGTCTG CCGCTCCCAA CAGCCCGAGT TTCTTGCCCC TTGAGTAAGG GGCAAGAAAC CTAAGCCGCC TGGTGGATGG CTCGGCTCGG GGCGCTGACG AAGGGCGTGG CAAGCTGCGA TAAGCCCCGG TGAG
J23SpTT_60 CGCA GAAGACGGGT TTGATGGGGC GGGGATGTAA GCCGGGAAGGG AAACCGACCC GTTTAGTCTG CCGCTCCCAA CAGCCCGAGT TTCTTGCCCC TTGAGTAAGG GGCAAGAAAC CTAAGCCGCC TGGTGGATGG CTCGGCTCGG GGCGCTGACG AAGGGCGTGG CAAGCTGCGA TAAGCCCCGG TGAGG
J23SpTT_19 CA GAAGACGGGT TTGATGGGGC GGGGATGTAA GCCGGGAAGGG AAACCGACCC GTTTAGTCTG CCGCTCCCAA CAGCCCGAGT TTCTTGCCCC TTGAGTAAGG GGCAAGAAAC CTAAGCCGCC TGGTGGATGG CTCGGCTCGG GGCGCTGACG AAGGGCGTGG CAAGCTGCGA TAAGCCCCGG TGAGGCGCAG
J23SpTT_9 A GAAGACGGGT TTGATGGGGC GGGGATGTAA GCCGGGAAGGG AAACCGACCC GTTTAGTCTG CCGCTCCCAA CAGCCCGAGT TTCTTGCCCC TTGAGTAAGG GGCAAGAAAC CTAAGCCGCC TGGTGGATGG CTCGGCTCGG GGCGCTGACG AAGGGCGTGG CAAGCTGCGA TAAGCCCCGG TGAGGCGCAG
J23SpTT_17 CA GAAGACGGGT TTGATGGGGC GGGGATGTAA GCCGGGAAGGG AAACCGACCC GTTTAGTCTG CCGCTCCCAA CAGCCCGAGT TTCTTGCCCC TTGAGTAAGG GGCAAGAAAC CTAAGCCGCC TGGTGGATGG CTCGGCTCGG GGCGCTGACG AAGGGCGTGG CAAGCTGCGA TAAGCCCCGG TGAGGCG
J23SpTT_336 CGCA GAAGACGGGT TTGATGGGGC GGGGATGTAA GCCGGGAAGGG AAACCGACCC GTTTAGTCTG CCGCTCCCAA CAGCCCGAGT TTCTTGCCCC TTGAGTAAGG GGCAAGAAAC CTAAGCCGCC TGGTGGATGG CTCGGCTCGG GGCGCTGACG AAGGGCGTGG CAAGCTGCGA TAAGCCCCGG TGAG
J23SpTT_40 GCA GAAGACGGGT TTGATGGGGC GGGGATGTAA GCCGGGAAGGG AAACCGACCC GTTTAGTCTG CCGCTCCCAA CAGCCCGAGT TTCTTGCCCC TTGAGTAAGG GGCAAGAAAC CTAAGCCGCC TGGTGGATGG CTCGGCTCGG GGCGCTGACG AAGGGCGTGG CAAGCTGCGA TAAGCCCCGG TGAG
J23SpTT_8 GCA GAAGACGGGT TTGATGGGGC GGGGATGTAA GCCGGGAAGGG AAACCGACCC GTTTAGTCTG CCGCTCCCAA CAGCCCGAGT TTCTTGCCCC TTGAGTAAGG GGCAAGAAAC CTAAGCCGCC TGGTGGATGG CTCGGCTCGG GGCGCTGACG AAGGGCGTGG CAAGCTGCGA TAAGCCCCGG TGAGG
J23SpTT_120 CA GAAGACGGGT TTGATGGGGC GGGGATGTAA GCCGGGAAGGG AAACCGACCC GTTTAGTCTG CCGCTCCCAA CAGCCCGAGT TTCTTGCCCC TTGAGTAAGG GGCAAGAAAC CTAAGCCGCC TGGTGGATGG CTCGGCTCGG GGCGCTGACG AAGGGCGTGG CAAGCTGCGA TAAGCCCCGG TGAG
J23SpTT_36 CGCA GAAGACGGGT TTGATGGGGC GGGGATGTAA GCCGGGAAGGG AAACCGACCC GTTTAGTCTG CCGCTCCCAA CAGCCCGAGT TTCTTGCCCC TTGAGTAAGG GGCAAGAAAC CTAAGCCGCC TGGTGGATGG CTCGGCTCGG GGCGCTGACG AAGGGCGTGG CAAGCTGCGA TAAGCCCCGG TG
J23SpTT_27 CA GAAGACGGGT TTGATGGGGC GGGGATGTAA GCCGGGAAGGG AAACCGACCC GTTTAGTCTG CCGCTCCCAA CAGCCCGAGT TTCTTGCCCC TTGAGTAAGG GGCAAGAAAC CTAAGCCGCC TGGTGGATGG CTCGGCTCGG GGCGCTGACG AAGGGCGTGG CAAGCTGCGA TAAGCCCCGG TGAGG
J23SpTT_17 CCGCA GAAGACGGGT TTGATGGGGC GGGGATGTAA GCCGGGAAGGG AAACCGACCC GTTTAGTCTG CCGCTCCCAA CAGCCCGAGT TTCTTGCCCC TTGAGTAAGG GGCAAGAAAC CTAAGCCGCC TGGTGGATGG CTCGGCTCGG GGCGCTGACG AAGGGCGTGG CAAGCTGCGA TAAGCCCCGG TG
J23SpTT_13 CCCGCA GAAGACGGGT TTGATGGGGC GGGGATGTAA GCCGGGAAGGG AAACCGACCC GTTTAGTCTG CCGCTCCCAA CAGCCCGAGT TTCTTGCCCC TTGAGTAAGG GGCAAGAAAC CTAAGCCGCC TGGTGGATGG CTCGGCTCGG GGCGCTGACG AAGGGCGTGG CAAGCTGCGA TAAGCCCCGG TG
J23SpTT_57 A GAAGACGGGT TTGATGGGGC GGGGATGTAA GCCGGGAAGGG AAACCGACCC GTTTAGTCTG CCGCTCCCAA CAGCCCGAGT TTCTTGCCCC TTGAGTAAGG GGCAAGAAAC CTAAGCCGCC TGGTGGATGG CTCGGCTCGG GGCGCTGACG AAGGGCGTGG CAAGCTGCGA TAAGCCCCGG TGAG
J23SpTT_37 CCCGCA GAAGACGGGT TTGATGGGGC GGGGATGTAA GCCGGGAAGGG AAACCGACCC GTTTAGTCTG CCGCTCCCAA CAGCCCGAGT TTCTTGCCCC TTGAGTAAGG GGCAAGAAAC CTAAGCCGCC TGGTGGATGG CTCGGCTCGG GGCGCTGACG AAGGGCGTGG CAAGCTGCGA TAAGCCCCGG
J23SpTT_18 CA GAAGACGGGT TTGATGGGGC GGGGATGTAA GCCGGGAAGGG AAACCGACCC GTTTAGTCTG CCGCTCCCAA CAGCCCGAGT TTCTTGCCCC TTGAGTAAGG GGCAAGAAAC CTAAGCCGCC TGGTGGATGG CTCGGCTCGG GGCGCTGACG AAGGGCGTGG CAAGCTGCGA TAAGCCCCGG TGAGGC
J23SpTT_9 A GAAGACGGGT TTGATGGGGC GGGGATGTAA GCCGGGAAGGG AAACCGACCC GTTTAGTCTG CCGCTCCCAA CAGCCCGAGT TTCTTGCCCC TTGAGTAAGG GGCAAGAAAC CTAAGCCGCC TGGTGGATGG CTCGGCTCGG GGCGCTGACG AAGGGCGTGG CAAGCTGCGA TAAGCCCCGG TGAGGCG
J23SpTT_9 A GAAGACGGGT TTGATGGGGC GGGGATGTAA GCCGGGAAGGG AAACCGACCC GTTTAGTCTG CCGCTCCCAA CAGCCCGAGT TTCTTGCCCC TTGAGTAAGG GGCAAGAAAC CTAAGCCGCC TGGTGGATGG CTCGGCTCGG GGCGCTGACG AAGGGCGTGG CAAGCTGCGA TAAGCCCCGG TGAGGC
J23SpTT_100 CGCA GAAGACGGGT TTGATGGGGC GGGGATGTAA GCCGGGAAGGG AAACCGACCC GTTTAGTCTG CCGCTCCCAA CAGCCCGAGT TTCTTGCCCC TTGAGTAAGG GGCAAGAAAC CTAAGCCGCC TGGTGGATGG CTCGGCTCGG GGCGCTGACG AAGGGCGTGG CAAGCTGCGA TAAGCCCCGG
J23SpTT_52 CCGCA GAAGACGGGT TTGATGGGGC GGGGATGTAA GCCGGGAAGGG AAACCGACCC GTTTAGTCTG CCGCTCCCAA CAGCCCGAGT TTCTTGCCCC TTGAGTAAGG GGCAAGAAAC CTAAGCCGCC TGGTGGATGG CTCGGCTCGG GGCGCTGACG AAGGGCGTGG CAAGCTGCGA TAAGCCCCGG
J23SpTT_32 CCCGCA GAAGACGGGT TTGATGGGGC GGGGATGTAA GCCGGGAAGGG AAACCGACCC GTTTAGTCTG CCGCTCCCAA CAGCCCGAGT TTCTTGCCCC TTGAGTAAGG GGCAAGAAAC CTAAGCCGCC TGGTGGATGG CTCGGCTCGG GGCGCTGACG AAGGGCGTGG CAAGCTGCGA TAAGCCCCG
J23SpTT_28 CCGCA GAAGACGGGT TTGATGGGGC GGGGATGTAA GCCGGGAAGGG AAACCGACCC GTTTAGTCTG CCGCTCCCAA CAGCCCGAGT TTCTTGCCCC TTGAGTAAGG GGCAAGAAAC CTAAGCCGCC TGGTGGATGG CTCGGCTCGG GGCGCTGACG AAGGGCGTGG CAAGCTGCGA TAAGCCCCG
J23SpTT_10 GCA GAAGACGGGT TTGATGGGGC GGGGATGTAA GCCGGGAAGGG AAACCGACCC GTTTAGTCTG CCGCTCCCAA CAGCCCGAGT TTCTTGCCCC TTGAGTAAGG GGCAAGAAAC CTAAGCCGCC TGGTGGATGG CTCGGCTCGG GGCGCTGACG AAGGGCGTGG CAAGCTGCGA TAAGCCCCGG
J23SpTT_68 CGCA GAAGACGGGT TTGATGGGGC GGGGATGTAA GCCGGGAAGGG AAACCGACCC GTTTAGTCTG CCGCTCCCAA CAGCCCGAGT TTCTTGCCCC TTGAGTAAGG GGCAAGAAAC CTAAGCCGCC TGGTGGATGG CTCGGCTCGG GGCGCTGACG AAGGGCGTGG CAAGCTGCGA TAAGCCCCG
J23SpTT_9 CCCGCA GAAGACGGGT TTGATGGGGC GGGGATGTAA GCCGGGAAGGG AAACCGACCC GTTTAGTCTG CCGCTCCCAA CAGCCCGAGT TTCTTGCCCC TTGAGTAAGG GGCAAGAAAC CTAAGCCGCC TGGTGGATGG CTCGGCTCGG GGCGCTGACG AAGGGCGTGG CAAGCTGCGA TAAGCCCC
J23SpTT_11 CGCA GAAGACGGGT TTGATGGGGC GGGGATGTAA GCCGGGAAGGG AAACCGACCC GTTTAGTCTG CCGCTCCCAA CAGCCCGAGT TTCTTGCCCC TTGAGTAAGG GGCAAGAAAC CTAAGCCGCC TGGTGGATGG CTCGGCTCGG GGCGCTGACG AAGGGCGTGG CAAGCTGCGA TAAGCCCC
J23SpTT_25 CA GAAGACGGGT TTGATGGGGC GGGGATGTAA GCCGGGAAGGG AAACCGACCC GTTTAGTCTG CCGCTCCCAA CAGCCCGAGT TTCTTGCCCC TTGAGTAAGG GGCAAGAAAC CTAAGCCGCC TGGTGGATGG CTCGGCTCGG GGCGCTGACG AAGGGCGTGG CAAGCTGCGA TAAGCCCCGG
J23SpTT_23 CA GAAGACGGGT TTGATGGGGC GGGGATGTAA GCCGGGAAGGG AAACCGACCC GTTTAGTCTG CCGCTCCCAA CAGCCCGAGT TTCTTGCCCC TTGAGTAAGG GGCAAGAAAC CTAAGCCGCC TGGTGGATGG CTCGGCTCGG GGCGCTGACG AAGGGCGTGG CAAGCTGCGA TAAGCCCCGG TG
J23SpTT_27 CA GAAGACGGGT TTGATGGGGC GGGGATGTAA GCCGGGAAGGG AAACCGACCC GTTTAGTCTG CCGCTCCCAA CAGCCCGAGT TTCTTGCCCC TTGAGTAAGG GGCAAGAAAC CTAAGCCGCC TGGTGGATGG CTCGGCTCGG GGCGCTGACG AAGGGCGTGG CAAGCTGCGA TAAGCCCCG
J23SpTT_16 A GAAGACGGGT TTGATGGGGC GGGGATGTAA GCCGGGAAGGG AAACCGACCC GTTTAGTCTG CCGCTCCCAA CAGCCCGAGT TTCTTGCCCC TTGAGTAAGG GGCAAGAAAC CTAAGCCGCC TGGTGGATGG CTCGGCTCGG GGCGCTGACG AAGGGCGTGG CAAGCTGCGA TAAGCCCCGG
J23SpTT_13 CCCGCA GAAGACGGGT TTGATGGGGC GGGGATGTAA GCCGGGAAGGG AAACCGACCC GTTTAGTCTG CCGCTCCCAA CAGCCCGAGT TTCTTGCCCC TTGAGTAAGG GGCAAGAAAC CTAAGCCGCC TGGTGGATGG CTCGGCTCGG GGCGCTGACG AAGGGCGTGG CAAGCTGCGA TAAGCC
J23SpTT_11 CGGGT TTGATGGGGC GGGGATGTAA GCCGGGAAGGG AAACCGACCC GTTTAGTCTG CCGCTCCCAA CAGCCCGAGT TTCTTGCCCC TTGAGTAAGG GGCAAGAAAC CTAAGCCGCC TGGTGGATGG CTCGGCTCGG GGCGCTGACG AAGGGCGTGG CAAGCTGCGA TAAGCCCCGG TGAGGCGCAG
J23SpTT_9 GACGGGT TTGATGGGGC GGGGATGTAA GCCGGGAAGGG AAACCGACCC GTTTAGTCTG CCGCTCCCAA CAGCCCGAGT TTCTTGCCCC TTGAGTAAGG GGCAAGAAAC CTAAGCCGCC TGGTGGATGG CTCGGCTCGG GGCGCTGACG AAGGGCGTGG CAAGCTGCGA TAAGCCCCGG TGAGGCGCAG
J23SpTT_10 GACGGGT TTGATGGGGC GGGGATGTAA GCCGGGAAGGG AAACCGACCC GTTTAGTCTG CCGCTCCCAA CAGCCCGAGT TTCTTGCCCC TTGAGTAAGG GGCAAGAAAC CTAAGCCGCC TGGTGGATGG CTCGGCTCGG GGCGCTGACG AAGGGCGTGG CAAGCTGCGA TAAGCCCCGG TGAGGCG
J23SpTT_10 CGGGT TTGATGGGGC GGGGATGTAA GCCGGGAAGGG AAACCGACCC GTTTAGTCTG CCGCTCCCAA CAGCCCGAGT TTCTTGCCCC TTGAGTAAGG GGCAAGAAAC CTAAGCCGCC TGGTGGATGG CTCGGCTCGG GGCGCTGACG AAGGGCGTGG CAAGCTGCGA TAAGCCCCGG TGAGGCG
J23SpTT_80 CGGGT TTGATGGGGC GGGGATGTAA GCCGGGAAGGG AAACCGACCC GTTTAGTCTG CCGCTCCCAA CAGCCCGAGT TTCTTGCCCC TTGAGTAAGG GGCAAGAAAC CTAAGCCGCC TGGTGGATGG CTCGGCTCGG GGCGCTGACG AAGGGCGTGG CAAGCTGCGA TAAGCCCCGG TGAG
J23SpTT_47 GACGGGT TTGATGGGGC GGGGATGTAA GCCGGGAAGGG AAACCGACCC GTTTAGTCTG CCGCTCCCAA CAGCCCGAGT TTCTTGCCCC TTGAGTAAGG GGCAAGAAAC CTAAGCCGCC TGGTGGATGG CTCGGCTCGG GGCGCTGACG AAGGGCGTGG CAAGCTGCGA TAAGCCCCGG TGAG
J23SpTT_21 AGACGGGT TTGATGGGGC GGGGATGTAA GCCGGGAAGGG AAACCGACCC GTTTAGTCTG CCGCTCCCAA CAGCCCGAGT TTCTTGCCCC TTGAGTAAGG GGCAAGAAAC CTAAGCCGCC TGGTGGATGG CTCGGCTCGG GGCGCTGACG AAGGGCGTGG CAAGCTGCGA TAAGCCCCGG TGAG
J23SpTT_19 AAGACGGGT TTGATGGGGC GGGGATGTAA GCCGGGAAGGG AAACCGACCC GTTTAGTCTG CCGCTCCCAA CAGCCCGAGT TTCTTGCCCC TTGAGTAAGG GGCAAGAAAC CTAAGCCGCC TGGTGGATGG CTCGGCTCGG GGCGCTGACG AAGGGCGTGG CAAGCTGCGA TAAGCCCCGG TGAG
J23SpTT_15 GACGGGT TTGATGGGGC GGGGATGTAA GCCGGGAAGGG AAACCGACCC GTTTAGTCTG CCGCTCCCAA CAGCCCGAGT TTCTTGCCCC TTGAGTAAGG GGCAAGAAAC CTAAGCCGCC TGGTGGATGG CTCGGCTCGG GGCGCTGACG AAGGGCGTGG CAAGCTGCGA TAAGCCCCGG TGAGG
J23SpTT_16 ACGGGT TTGATGGGGC GGGGATGTAA GCCGGGAAGGG AAACCGACCC GTTTAGTCTG CCGCTCCCAA CAGCCCGAGT TTCTTGCCCC TTGAGTAAGG GGCAAGAAAC CTAAGCCGCC TGGTGGATGG CTCGGCTCGG GGCGCTGACG AAGGGCGTGG CAAGCTGCGA TAAGCCCCGG TGAG
J23SpTT_15 CGGGT TTGATGGGGC GGGGATGTAA GCCGGGAAGGG AAACCGACCC GTTTAGTCTG CCGCTCCCAA CAGCCCGAGT TTCTTGCCCC TTGAGTAAGG GGCAAGAAAC CTAAGCCGCC TGGTGGATGG CTCGGCTCGG GGCGCTGACG AAGGGCGTGG CAAGCTGCGA TAAGCCCCGG TGAGG
J23SpTT_18 CCCAA CAGCCCGAGT TTCTTGCCCC TTGAGTAAGG GGCAAGAAAC CTAAGCCGCC TGGTGGATGG CTCGGCTCGG GGCGCTGACG AAGGGCGTGG CAAGCTGCGA TAAGCCCCGG TGAGGCGCAG GCAGCCGTCG AGCCGGGGAT TCCCGAATGG GACCTCCCGC GGCTTTTGCC GCACTCCCGT CAGGGAGGGG
J23SpTT_15 CTCCCAA CAGCCCGAGT TTCTTGCCCC TTGAGTAAGG GGCAAGAAAC CTAAGCCGCC TGGTGGATGG CTCGGCTCGG GGCGCTGACG AAGGGCGTGG CAAGCTGCGA TAAGCCCCGG TGAGGCGCAG GCAGCCGTCG AGCCGGGGAT TCCCGAATGG GACCTCCCGC GGCTTTTGCC GCACTCCCGT CAGGGAGGGG
J23SpTT_13 CTCCCAA CAGCCCGAGT TTCTTGCCCC TTGAGTAAGG GGCAAGAAAC CTAAGCCGCC TGGTGGATGG CTCGGCTCGG GGCGCTGACG AAGGGCGTGG CAAGCTGCGA TAAGCCCCGG TGAGGCGCAG GCAGCCGTCG AGCCGGGGAT TCCCGAATGG GACCTCCCGC GGCTTTTGCC GCACTCCCGT CAGGGAGGG
J23SpTT_14 CCCAA CAGCCCGAGT TTCTTGCCCC TTGAGTAAGG GGCAAGAAAC CTAAGCCGCC TGGTGGATGG CTCGGCTCGG GGCGCTGACG AAGGGCGTGG CAAGCTGCGA TAAGCCCCGG TGAGGCGCAG GCAGCCGTCG AGCCGGGGAT TCCCGAATGG GACCTCCCGC GGCTTTTGCC GCACTCCCGT CAGGGAGGG
J23SpTT_35 CGGGT TTGATGGGGC GGGGATGTAA GCCGGGAAGGG AAACCGACCC GTTTAGTCTG CCGCTCCCAA CAGCCCGAGT TTCTTGCCCC TTGAGTAAGG GGCAAGAAAC CTAAGCCGCC TGGTGGATGG CTCGGCTCGG GGCGCTGACG AAGGGCGTGG CAAGCTGCGA TAAGCCCCGG
J23SpTT_10 CGGGT TTGATGGGGC GGGGATGTAA GCCGGGAAGGG AAACCGACCC GTTTAGTCTG CCGCTCCCAA CAGCCCGAGT TTCTTGCCCC TTGAGTAAGG GGCAAGAAAC CTAAGCCGCC TGGTGGATGG CTCGGCTCGG GGCGCTGACG AAGGGCGTGG CAAGCTGCGA TAAGCCCCGG TG
J23SpTT_21 GACGGGT TTGATGGGGC GGGGATGTAA GCCGGGAAGGG AAACCGACCC GTTTAGTCTG CCGCTCCCAA CAGCCCGAGT TTCTTGCCCC TTGAGTAAGG GGCAAGAAAC CTAAGCCGCC TGGTGGATGG CTCGGCTCGG GGCGCTGACG AAGGGCGTGG CAAGCTGCGA TAAGCCCCGG
J23SpTT_9 ACGGGT TTGATGGGGC GGGGATGTAA GCCGGGAAGGG AAACCGACCC GTTTAGTCTG CCGCTCCCAA CAGCCCGAGT TTCTTGCCCC TTGAGTAAGG GGCAAGAAAC CTAAGCCGCC TGGTGGATGG CTCGGCTCGG GGCGCTGACG AAGGGCGTGG CAAGCTGCGA TAAGCCCCGG
J23SpTT_23 GT TTGATGGGGC GGGGATGTAA GCCGGGAAGGG AAACCGACCC GTTTAGTCTG CCGCTCCCAA CAGCCCGAGT TTCTTGCCCC TTGAGTAAGG GGCAAGAAAC CTAAGCCGCC TGGTGGATGG CTCGGCTCGG GGCGCTGACG AAGGGCGTGG CAAGCTGCGA TAAGCCCCGG TGAG
J23SpTT_266 CGCA GAAGACGGGT TTGATGGGGC GGGGATGTAA GCCGGGAAGGG AAACCGACCC GTTTAGTCTG CCGCTCCCAA CAGCCCGAGT TTCTTGCCCC TTGAGTAAGG GGCAAGAAAC CTAAGCCGCC TGGTGGATGG CTCGGCTCGG GGCGCTGACG AAGGGCGTGG CAAGCTGCG
J23SpTT_130 CGCA GAAGACGGGT TTGATGGGGC GGGGATGTAA GCCGGGAAGGG AAACCGACCC GTTTAGTCTG CCGCTCCCAA CAGCCCGAGT TTCTTGCCCC TTGAGTAAGG GGCAAGAAAC CTAAGCCGCC TGGTGGATGG CTCGGCTCGG GGCGCTGACG AAGGGCGTGG CAAGCTGCGA T
J23SpTT_53 CCCGCA GAAGACGGGT TTGATGGGGC GGGGATGTAA GCCGGGAAGGG AAACCGACCC GTTTAGTCTG CCGCTCCCAA CAGCCCGAGT TTCTTGCCCC TTGAGTAAGG GGCAAGAAAC CTAAGCCGCC TGGTGGATGG CTCGGCTCGG GGCGCTGACG AAGGGCGTGG CAAGCTGCGA T
J23SpTT_44 CCGCA GAAGACGGGT TTGATGGGGC GGGGATGTAA GCCGGGAAGGG AAACCGACCC GTTTAGTCTG CCGCTCCCAA CAGCCCGAGT TTCTTGCCCC TTGAGTAAGG GGCAAGAAAC CTAAGCCGCC TGGTGGATGG CTCGGCTCGG GGCGCTGACG AAGGGCGTGG CAAGCTGCGA T
```

```
J23SpTT_9      CCGCA GAAGACGGGT TTGATGGGGC GGGGATGTAA GCCGGGAAGGG AAACCGACCC GTTTAGTCTG CCGCTCCCAA CAGCCCGAGT TTCTTGCCCC TTGAGTAAGG GGCAAGAAAC CTAAGCCGCC TGGTGGATGG CTCGGCTCGG GGCGCTGACG AAGGGCGTGG CAAGCTGCGA G
J23SpTT_104    CCGCA GAAGACGGGT TTGATGGGGC GGGGATGTAA GCCGGGAAGGG AAACCGACCC GTTTAGTCTG CCGCTCCCAA CAGCCCGAGT TTCTTGCCCC TTGAGTAAGG GGCAAGAAAC CTAAGCCGCC TGGTGGATGG CTCGGCTCGG GGCGCTGACG AAGGGCGTGG CAAGCTGCG
J23SpTT_83    CCCGCA GAAGACGGGT TTGATGGGGC GGGGATGTAA GCCGGGAAGGG AAACCGACCC GTTTAGTCTG CCGCTCCCAA CAGCCCGAGT TTCTTGCCCC TTGAGTAAGG GGCAAGAAAC CTAAGCCGCC TGGTGGATGG CTCGGCTCGG GGCGCTGACG AAGGGCGTGG CAAGCTGCG
J23SpTT_10      CGCA GAAGACGGGT TTGATGGGGC GGGGATGTAA GCCGGGAAGGG AAACCGACCC GTTTAGTCTG CCGCTCCCAA CAGCCCGAGT TTCTTGCCCC TTGAGTAAGG GGCAAGAAAC CTAAGCCGCC TGGTGGATGG CTCGGCTCGG GGCGCTGACG AAGGGCGTGG CAAGCTGCGA C
J23SpTT_13       GCA GAAGACGGGT TTGATGGGGC GGGGATGTAA GCCGGGAAGGG AAACCGACCC GTTTAGTCTG CCGCTCCCAA CAGCCCGAGT TTCTTGCCCC TTGAGTAAGG GGCAAGAAAC CTAAGCCGCC TGGTGGATGG CTCGGCTCGG GGCGCTGACG AAGGGCGTGG CAAGCTGCGA T
J23SpTT_43        CA GAAGACGGGT TTGATGGGGC GGGGATGTAA GCCGGGAAGGG AAACCGACCC GTTTAGTCTG CCGCTCCCAA CAGCCCGAGT TTCTTGCCCC TTGAGTAAGG GGCAAGAAAC CTAAGCCGCC TGGTGGATGG CTCGGCTCGG GGCGCTGACG AAGGGCGTGG CAAGCTGCGA T
J23SpTT_25     CCGCA GAAGACGGGT TTGATGGGGC GGGGATGTAA GCCGGGAAGGG AAACCGACCC GTTTAGTCTG CCGCTCCCAA CAGCCCGAGT TTCTTGCCCC TTGAGTAAGG GGCAAGAAAC CTAAGCCGCC TGGTGGATGG CTCGGCTCGG GGCGCTGACG AAGGGCGTGG CAAGCTGC
J23SpTT_13    CCCGCA GAAGACGGGT TTGATGGGGC GGGGATGTAA GCCGGGAAGGG AAACCGACCC GTTTAGTCTG CCGCTCCCAA CAGCCCGAGT TTCTTGCCCC TTGAGTAAGG GGCAAGAAAC CTAAGCCGCC TGGTGGATGG CTCGGCTCGG GGCGCTGACG AAGGGCGTGG CAAGCTGC
J23SpTT_53      CGCA GAAGACGGGT TTGATGGGGC GGGGATGTAA GCCGGGAAGGG AAACCGACCC GTTTAGTCTG CCGCTCCCAA CAGCCCGAGT TTCTTGCCCC TTGAGTAAGG GGCAAGAAAC CTAAGCCGCC TGGTGGATGG CTCGGCTCGG GGCGCTGACG AAGGGCGTGG CAAGCTGC
J23SpTT_48     CCGCA GAAGACGGGT TTGATGGGGC GGGGATGTAA GCCGGGAAGGG AAACCGACCC GTTTAGTCTG CCGCTCCCAA CAGCCCGAGT TTCTTGCCCC TTGAGTAAGG GGCAAGAAAC CTAAGCCGCC TGGTGGATGG CTCGGCTCGG GGCGCTGACG AAGGGCGTGG CAAGCTG
J23SpTT_31       GCA GAAGACGGGT TTGATGGGGC GGGGATGTAA GCCGGGAAGGG AAACCGACCC GTTTAGTCTG CCGCTCCCAA CAGCCCGAGT TTCTTGCCCC TTGAGTAAGG GGCAAGAAAC CTAAGCCGCC TGGTGGATGG CTCGGCTCGG GGCGCTGACG AAGGGCGTGG CAAGCTGCG
J23SpTT_29      CGCA GAAGACGGGT TTGATGGGGC GGGGATGTAA GCCGGGAAGGG AAACCGACCC GTTTAGTCTG CCGCTCCCAA CAGCCCGAGT TTCTTGCCCC TTGAGTAAGG GGCAAGAAAC CTAAGCCGCC TGGTGGATGG CTCGGCTCGG GGCGCTGACG AAGGGCGTGG CAAGCTGCT
J23SpTT_24    CCCGCA GAAGACGGGT TTGATGGGGC GGGGATGTAA GCCGGGAAGGG AAACCGACCC GTTTAGTCTG CCGCTCCCAA CAGCCCGAGT TTCTTGCCCC TTGAGTAAGG GGCAAGAAAC CTAAGCCGCC TGGTGGATGG CTCGGCTCGG GGCGCTGACG AAGGGCGTGG CAAGCTG
J23SpTT_20         A GAAGACGGGT TTGATGGGGC GGGGATGTAA GCCGGGAAGGG AAACCGACCC GTTTAGTCTG CCGCTCCCAA CAGCCCGAGT TTCTTGCCCC TTGAGTAAGG GGCAAGAAAC CTAAGCCGCC TGGTGGATGG CTCGGCTCGG GGCGCTGACG AAGGGCGTGG CAAGCTGCGA T
J23SpTT_108     CGCA GAAGACGGGT TTGATGGGGC GGGGATGTAA GCCGGGAAGGG AAACCGACCC GTTTAGTCTG CCGCTCCCAA CAGCCCGAGT TTCTTGCCCC TTGAGTAAGG GGCAAGAAAC CTAAGCCGCC TGGTGGATGG CTCGGCTCGG GGCGCTGACG AAGGGCGTGG CAAGCTG
J23SpTT_98        CA GAAGACGGGT TTGATGGGGC GGGGATGTAA GCCGGGAAGGG AAACCGACCC GTTTAGTCTG CCGCTCCCAA CAGCCCGAGT TTCTTGCCCC TTGAGTAAGG GGCAAGAAAC CTAAGCCGCC TGGTGGATGG CTCGGCTCGG GGCGCTGACG AAGGGCGTGG CAAGCTGCG
J23SpTT_10    CCCGCA GAAGACGGGT TTGATGGGGC GGGGATGTAA GCCGGGAAGGG AAACCGACCC GTTTAGTCTG CCGCTCCCAA CAGCCCGAGT TTCTTGCCCC TTGAGTAAGG GGCAAGAAAC CTAAGCCGCC TGGTGGATGG CTCGGCTCGG GGCGCTGACG AAGGGCGTGG CAAGCT
J23SpTT_48                CGGGT TTGATGGGGC GGGGATGTAA GCCGGGAAGGG AAACCGACCC GTTTAGTCTG CCGCTCCCAA CAGCCCGAGT TTCTTGCCCC TTGAGTAAGG GGCAAGAAAC CTAAGCCGCC TGGTGGATGG CTCGGCTCGG GGCGCTGACG AAGGGCGTGG CAAGCTGCG
J23SpTT_38                CGGGT TTGATGGGGC GGGGATGTAA GCCGGGAAGGG AAACCGACCC GTTTAGTCTG CCGCTCCCAA CAGCCCGAGT TTCTTGCCCC TTGAGTAAGG GGCAAGAAAC CTAAGCCGCC TGGTGGATGG CTCGGCTCGG GGCGCTGACG AAGGGCGTGG CAAGCTGCGA T
J23SpTT_12              GACGGGT TTGATGGGGC GGGGATGTAA GCCGGGAAGGG AAACCGACCC GTTTAGTCTG CCGCTCCCAA CAGCCCGAGT TTCTTGCCCC TTGAGTAAGG GGCAAGAAAC CTAAGCCGCC TGGTGGATGG CTCGGCTCGG GGCGCTGACG AAGGGCGTGG CAAGCTGCGA T
J23SpTT_30              GACGGGT TTGATGGGGC GGGGATGTAA GCCGGGAAGGG AAACCGACCC GTTTAGTCTG CCGCTCCCAA CAGCCCGAGT TTCTTGCCCC TTGAGTAAGG GGCAAGAAAC CTAAGCCGCC TGGTGGATGG CTCGGCTCGG GGCGCTGACG AAGGGCGTGG CAAGCTGCG
J23SpTT_14             AGACGGGT TTGATGGGGC GGGGATGTAA GCCGGGAAGGG AAACCGACCC GTTTAGTCTG CCGCTCCCAA CAGCCCGAGT TTCTTGCCCC TTGAGTAAGG GGCAAGAAAC CTAAGCCGCC TGGTGGATGG CTCGGCTCGG GGCGCTGACG AAGGGCGTGG CAAGCTGCG
J23SpTT_11              GACGGGT TTGATGGGGC GGGGATGTAA GCCGGGAAGGG AAACCGACCC GTTTAGTCTG CCGCTCCCAA CAGCCCGAGT TTCTTGCCCC TTGAGTAAGG GGCAAGAAAC CTAAGCCGCC TGGTGGATGG CTCGGCTCGG GGCGCTGACG AAGGGCGTGG CAAGCTGC
J23SpTT_20               ACGGGT TTGATGGGGC GGGGATGTAA GCCGGGAAGGG AAACCGACCC GTTTAGTCTG CCGCTCCCAA CAGCCCGAGT TTCTTGCCCC TTGAGTAAGG GGCAAGAAAC CTAAGCCGCC TGGTGGATGG CTCGGCTCGG GGCGCTGACG AAGGGCGTGG CAAGCTGCG
J23SpTT_15              GACGGGT TTGATGGGGC GGGGATGTAA GCCGGGAAGGG AAACCGACCC GTTTAGTCTG CCGCTCCCAA CAGCCCGAGT TTCTTGCCCC TTGAGTAAGG GGCAAGAAAC CTAAGCCGCC TGGTGGATGG CTCGGCTCGG GGCGCTGACG AAGGGCGTGG CAAGCTG
J23SpTT_20                CGGGT TTGATGGGGC GGGGATGTAA GCCGGGAAGGG AAACCGACCC GTTTAGTCTG CCGCTCCCAA CAGCCCGAGT TTCTTGCCCC TTGAGTAAGG GGCAAGAAAC CTAAGCCGCC TGGTGGATGG CTCGGCTCGG GGCGCTGACG AAGGGCGTGG CAAGCTGC
J23SpTT_61                                 C GGGGATGTAA GCCGGGAAGGG AAACCGACCC GTTTAGTCTG CCGCTCCCAA CAGCCCGAGT TTCTTGCCCC TTGAGTAAGG GGCAAGAAAC CTAAGCCGCC TGGTGGATGG CTCGGCTCGG GGCGCTGACG AAGGGCGTGG CAAGCTGCGA TAAGCCCCGG TGAG
J23SpTT_19                                 C GGGGATGTAA GCCGGGAAGGG AAACCGACCC GTTTAGTCTG CCGCTCCCAA CAGCCCGAGT TTCTTGCCCC TTGAGTAAGG GGCAAGAAAC CTAAGCCGCC TGGTGGATGG CTCGGCTCGG GGCGCTGACG AAGGGCGTGG CAAGCTGCGA TAAGCCCCGG TGAGG
J23SpTT_16                                 C GGGGATGTAA GCCGGGAAGGG AAACCGACCC GTTTAGTCTG CCGCTCCCAA CAGCCCGAGT TTCTTGCCCC TTGAGTAAGG GGCAAGAAAC CTAAGCCGCC TGGTGGATGG CTCGGCTCGG GGCGCTGACG AAGGGCGTGG CAAGCTGCGA TAAGCCCCGG TGAGGC
J23SpTT_329        A GAAGACGGGT TTGATGGGGC GGGGATGTAA GCCGGGAAGGG AAACCGACCC GTTTAGTCTG CCGCTCCCAA CAGCCCGAGT TTCTTGCCCC TTGAGTAAGG GGCAAGAAAC CTAAGCCGCC TGGTGGATGG CTCGGCTCGG GGCGCTGACG AAGGGCG
J23SpTT_140        A GAAGACGGGT TTGATGGGGC GGGGATGTAA GCCGGGAAGGG AAACCGACCC GTTTAGTCTG CCGCTCCCAA CAGCCCGAGT TTCTTGCCCC TTGAGTAAGG GGCAAGAAAC CTAAGCCGCC TGGTGGATGG CTCGGCTCGG GGCGCTGACG AAGGGCGTG
J23SpTT_14           GAAGACGGGT TTGATGGGGC GGGGATGTAA GCCGGGAAGGG AAACCGACCC GTTTAGTCTG CCGCTCCCAA CAGCCCGAGT TTCTTGCCCC TTGAGTAAGG GGCAAGAAAC CTAAGCCGCC TGGTGGATGG CTCGGCTCGG GGCGCTGACG AAGGGCGTG
J23SpTT_24           GAAGACGGGT TTGATGGGGC GGGGATGTAA GCCGGGAAGGG AAACCGACCC GTTTAGTCTG CCGCTCCCAA CAGCCCGAGT TTCTTGCCCC TTGAGTAAGG GGCAAGAAAC CTAAGCCGCC TGGTGGATGG CTCGGCTCGG GGCGCTGACG AAGGGCGTGG
J23SpTT_37            AAGACGGGT TTGATGGGGC GGGGATGTAA GCCGGGAAGGG AAACCGACCC GTTTAGTCTG CCGCTCCCAA CAGCCCGAGT TTCTTGCCCC TTGAGTAAGG GGCAAGAAAC CTAAGCCGCC TGGTGGATGG CTCGGCTCGG GGCGCTGACG AAGGGCGTGG
J23SpTT_13            AAGACGGGT TTGATGGGGC GGGGATGTAA GCCGGGAAGGG AAACCGACCC GTTTAGTCTG CCGCTCCCAA CAGCCCGAGT TTCTTGCCCC TTGAGTAAGG GGCAAGAAAC CTAAGCCGCC TGGTGGATGG CTCGGCTCGG GGCGCTGACG AAGGGCGTGG C
J23SpTT_30            AAGACGGGT TTGATGGGGC GGGGATGTAA GCCGGGAAGGG AAACCGACCC GTTTAGTCTG CCGCTCCCAA CAGCCCGAGT TTCTTGCCCC TTGAGTAAGG GGCAAGAAAC CTAAGCCGCC TGGTGGATGG CTCGGCTCGG GGCGCTGACG AAGGGCGTG
J23SpTT_75             AGACGGGT TTGATGGGGC GGGGATGTAA GCCGGGAAGGG AAACCGACCC GTTTAGTCTG CCGCTCCCAA CAGCCCGAGT TTCTTGCCCC TTGAGTAAGG GGCAAGAAAC CTAAGCCGCC TGGTGGATGG CTCGGCTCGG GGCGCTGACG AAGGGCGTGG
J23SpTT_43              GACGGGT TTGATGGGGC GGGGATGTAA GCCGGGAAGGG AAACCGACCC GTTTAGTCTG CCGCTCCCAA CAGCCCGAGT TTCTTGCCCC TTGAGTAAGG GGCAAGAAAC CTAAGCCGCC TGGTGGATGG CTCGGCTCGG GGCGCTGACG AAGGGCGTGG C
J23SpTT_42             AGACGGGT TTGATGGGGC GGGGATGTAA GCCGGGAAGGG AAACCGACCC GTTTAGTCTG CCGCTCCCAA CAGCCCGAGT TTCTTGCCCC TTGAGTAAGG GGCAAGAAAC CTAAGCCGCC TGGTGGATGG CTCGGCTCGG GGCGCTGACG AAGGGCGTG
J23SpTT_29           GAAGACGGGT TTGATGGGGC GGGGATGTAA GCCGGGAAGGG AAACCGACCC GTTTAGTCTG CCGCTCCCAA CAGCCCGAGT TTCTTGCCCC TTGAGTAAGG GGCAAGAAAC CTAAGCCGCC TGGTGGATGG CTCGGCTCGG GGCGCTGACG AAGGGCG
J23SpTT_24                CGGGT TTGATGGGGC GGGGATGTAA GCCGGGAAGGG AAACCGACCC GTTTAGTCTG CCGCTCCCAA CAGCCCGAGT TTCTTGCCCC TTGAGTAAGG GGCAAGAAAC CTAAGCCGCC TGGTGGATGG CTCGGCTCGG GGCGCTGACG AAGGGCGTGG CAAG
J23SpTT_15              GACGGGT TTGATGGGGC GGGGATGTAA GCCGGGAAGGG AAACCGACCC GTTTAGTCTG CCGCTCCCAA CAGCCCGAGT TTCTTGCCCC TTGAGTAAGG GGCAAGAAAC CTAAGCCGCC TGGTGGATGG CTCGGCTCGG GGCGCTGACG AAGGGCGTGG CAAG
J23SpTT_15                CGGGT TTGATGGGGC GGGGATGTAA GCCGGGAAGGG AAACCGACCC GTTTAGTCTG CCGCTCCCAA CAGCCCGAGT TTCTTGCCCC TTGAGTAAGG GGCAAGAAAC CTAAGCCGCC TGGTGGATGG CTCGGCTCGG GGCGCTGACG AAGGGCGTGG CAAGC
J23SpTT_13                CGGGT TTGATGGGGC GGGGATGTAA GCCGGGAAGGG AAACCGACCC GTTTAGTCTG CCGCTCCCAA CAGCCCGAGT TTCTTGCCCC TTGAGTAAGG GGCAAGAAAC CTAAGCCGCC TGGTGGATGG CTCGGCTCGG GGCGCTGACG AAGGGCGTGG CAAGCTG
J23SpTT_16                   GT TTGATGGGGC GGGGATGTAA GCCGGGAAGGG AAACCGACCC GTTTAGTCTG CCGCTCCCAA CAGCCCGAGT TTCTTGCCCC TTGAGTAAGG GGCAAGAAAC CTAAGCCGCC TGGTGGATGG CTCGGCTCGG GGCGCTGACG AAGGGCGTGG CAAGCTGCG
J23SpTT_4662    CGCA GAAGACGGGT TTGATGGGGC GGGGATGTAA GCCGGGAAGGG AAACCGACCC GTTTAGTCTG CCGCTCCCAA CAGCCCGAGT TTCTTGCCCC TTGAGTAAGG GGCAAGAAAC CTAAGCCGCC TGGTGGATGG CTCGGCTCGG GGCGCTGACG AAG
J23SpTT_4341    CGCA GAAGACGGGT TTGATGGGGC GGGGATGTAA GCCGGGAAGGG AAACCGACCC GTTTAGTCTG CCGCTCCCAA CAGCCCGAGT TTCTTGCCCC TTGAGTAAGG GGCAAGAAAC CTAAGCCGCC TGGTGGATGG CTCGGCTCGG GGCGCTGACG AAGG
J23SpTT_1928   CCGCA GAAGACGGGT TTGATGGGGC GGGGATGTAA GCCGGGAAGGG AAACCGACCC GTTTAGTCTG CCGCTCCCAA CAGCCCGAGT TTCTTGCCCC TTGAGTAAGG GGCAAGAAAC CTAAGCCGCC TGGTGGATGG CTCGGCTCGG GGCGCTGACG AAGG
J23SpTT_2114   CCGCA GAAGACGGGT TTGATGGGGC GGGGATGTAA GCCGGGAAGGG AAACCGACCC GTTTAGTCTG CCGCTCCCAA CAGCCCGAGT TTCTTGCCCC TTGAGTAAGG GGCAAGAAAC CTAAGCCGCC TGGTGGATGG CTCGGCTCGG GGCGCTGACG AAG
J23SpTT_1910  CCCGCA GAAGACGGGT TTGATGGGGC GGGGATGTAA GCCGGGAAGGG AAACCGACCC GTTTAGTCTG CCGCTCCCAA CAGCCCGAGT TTCTTGCCCC TTGAGTAAGG GGCAAGAAAC CTAAGCCGCC TGGTGGATGG CTCGGCTCGG GGCGCTGACG AAG
J23SpTT_11    CCCGCA GAAGACGGGT TTGATGGGGC GGGGATGTAA GCCGGGAAGGG AAACCGACCC GTTTAGTCTG CCGCTCCCAA CAGCCCGAGT TTCTTGCCCC TTGAGTAAGG GGCAAGAAAC CTAAGCCGCC TGGTGGATGG CTCGGCTCGG GGCGCTGACG AAGN
J23SpTT_32     CCGCA GAAGACGGGT TTGATGGGGC GGGGATGTAA GCCGGGAAGGG AAACCGACCC GTTTAGTCTG CCGCTCCCAA CAGCCCGAGT TTCTTGCCCC TTGAGTAAGG GGCAAGAAAC CTAAGCCGCC TGGTGGATGG CTCGGCTCGG GGCGCTGACG AAGAG
J23SpTT_31    CCCGCA GAAGACGGGT TTGATGGGGC GGGGATGTAA GCCGGGAAGGG AAACCGACCC GTTTAGTCTG CCGCTCCCAA CAGCCCGAGT TTCTTGCCCC TTGAGTAAGG GGCAAGAAAC CTAAGCCGCC TGGTGGATGG CTCGGCTCGG GGCGCTGACG AAGAG
J23SpTT_17     CCCCA GAAGACGGGT TTGATGGGGC GGGGATGTAA GCCGGGAAGGG AAACCGACCC GTTTAGTCTG CCGCTCCCAA CAGCCCGAGT TTCTTGCCCC TTGAGTAAGG GGCAAGAAAC CTAAGCCGCC TGGTGGATGG CTCGGCTCGG GGCGCTGACG AAGGG
J23SpTT_10      CCCA GAAGACGGGT TTGATGGGGC GGGGATGTAA GCCGGGAAGGG AAACCGACCC GTTTAGTCTG CCGCTCCCAA CAGCCCGAGT TTCTTGCCCC TTGAGTAAGG GGCAAGAAAC CTAAGCCGCC TGGTGGATGG CTCGGCTCGG GGCGCTGACG AAGGGC
J23SpTT_13      CCCA GAAGACGGGT TTGATGGGGC GGGGATGTAA GCCGGGAAGGG AAACCGACCC GTTTAGTCTG CCGCTCCCAA CAGCCCGAGT TTCTTGCCCC TTGAGTAAGG GGCAAGAAAC CTAAGCCGCC TGGTGGATGG CTCGGCTCGG GGCGCTGACG AAGGG
J23SpTT_214     CGCA GAAGACGGGT TTGATGGGGC GGGGATGTAA GCCGGGAAGGG AAACCGACCC GTTTAGTCTG CCGCTCCCAA CAGCCCGAGT TTCTTGCCCC TTGAGTAAGG GGCAAGAAAC CTAAGCCGCC TGGTGGATGG CTCGGCTCGG GGCGCTGACG AAGGGG
J23SpTT_11      CGCA GAAGACGGGT TTGATGGGGC GGGGATGTAA GCCGGGAAGGG AAACCGACCC GTTTAGTCTG CCGCTCCCAA CAGCCCGAGT TTCTTGCCCC TTGAGTAAGG GGCAAGAAAC CTAAGCCGCC TGGTGGATGG CTCGGCTCGG GGCGCTGACG AAGGGAC
J23SpTT_20       GCA GAAGACGGGT TTGATGGGGC GGGGATGTAA GCCGGGAAGGG AAACCGACCC GTTTAGTCTG CCGCTCCCAA CAGCCCGAGT TTCTTGCCCC TTGAGTAAGG GGCAAGAAAC CTAAGCCGCC TGGTGGATGG CTCGGCTCGG GGCGCTGACG AAGGGG
J23SpTT_85      CGCA GAAGACGGGT TTGATGGGGC GGGGATGTAA GCCGGGAAGGG AAACCGACCC GTTTAGTCTG CCGCTCCCAA CAGCCCGAGT TTCTTGCCCC TTGAGTAAGG GGCAAGAAAC CTAAGCCGCC TGGTGGATGG CTCGGCTCGG GGCGCTGACG AAGAG
J23SpTT_34      CGCA GAAGACGGGT TTGATGGGGC GGGGATGTAA GCCGGGAAGGG AAACCGACCC GTTTAGTCTG CCGCTCCCAA CAGCCCGAGT TTCTTGCCCC TTGAGTAAGG GGCAAGAAAC CTAAGCCGCC TGGTGGATGG CTCGGCTCGG GGCGCTGACA AAGGG
J23SpTT_18     CCGCA GAAGACGGGT TTGATGGGGC GGGGATGTAA GCCGGGAAGGG AAACCGACCC GTTTAGTCTG CCGCTCCCAA CAGCCCGAGT TTCTTGCCCC TTGAGTAAGG GGCAAGAAAC CTAAGCCGCC TGGTGGATGG CTCGGCTCGG GGCGCTGACA AAGG
J23SpTT_15     CCGCA GAAGACGGGT TTGATGGGGC GGGGATGTAA GCCGGGAAGGG AAACCGACCC GTTTAGTCTG CCGCTCCCAA CAGCCCGAGT TTCTTGCCCC TTGAGTAAGG GGCAAGAAAC CTAAGCCGCC TGGTGGATGG CTCGGCTCGG GGCGCTGACA AAGGG
J23SpTT_14    CCCGCA GAAGACGGGT TTGATGGGGC GGGGATGTAA GCCGGGAAGGG AAACCGACCC GTTTAGTCTG CCGCTCCCAA CAGCCCGAGT TTCTTGCCCC TTGAGTAAGG GGCAAGAAAC CTAAGCCGCC TGGTGGATGG CTCGGCTCGG GGCGCTGACA AAGGG
J23SpTT_23    CCCGCA GAAGACGGGT TTGATGGGGC GGGGATGTAA GCCGGGAAGGG AAACCGACCC GTTTAGTCTG CCGCTCCCAA CAGCCCGAGT TTCTTGCCCC TTGAGTAAGG GGCAAGAAAC CTAAGCCGCC TGGTGGATGG CTCGGCTCGG GGCGCTGACA AAG
J23SpTT_8        GCA GAAGACGGGT TTGATGGGGC GGGGATGTAA GCCGGGAAGGG AAACCGACCC GTTTAGTCTG CCGCTCCCAA CAGCCCGAGT TTCTTGCCCC TTGAGTAAGG GGCAAGAAAC CTAAGCCGCC TGGTGGATGG CTCGGCTCGG GGCGCTGACA AAGGG
J23SpTT_1328      CA GAAGACGGGT TTGATGGGGC GGGGATGTAA GCCGGGAAGGG AAACCGACCC GTTTAGTCTG CCGCTCCCAA CAGCCCGAGT TTCTTGCCCC TTGAGTAAGG GGCAAGAAAC CTAAGCCGCC TGGTGGATGG CTCGGCTCGG GGCGCTGACG AAGGGC
J23SpTT_685       CA GAAGACGGGT TTGATGGGGC GGGGATGTAA GCCGGGAAGGG AAACCGACCC GTTTAGTCTG CCGCTCCCAA CAGCCCGAGT TTCTTGCCCC TTGAGTAAGG GGCAAGAAAC CTAAGCCGCC TGGTGGATGG CTCGGCTCGG GGCGCTGACG AAGGGCG
J23SpTT_10        CA GAAGACGGGT TTGATGGGGC GGGGATGTAA GCCGGGAAGGG AAACCGACCC GTTTAGTCTG CCGCTCCCAA CAGCCCGAGT TTCTTGCCCC TTGAGTAAGG GGCAAGAAAC CTAAGCCGCC TGGTGGATGG CTCGGCTCGG GGCGCTGACG AAGGGCGG
J23SpTT_426      GCA GAAGACGGGT TTGATGGGGC GGGGATGTAA GCCGGGAAGGG AAACCGACCC GTTTAGTCTG CCGCTCCCAA CAGCCCGAGT TTCTTGCCCC TTGAGTAAGG GGCAAGAAAC CTAAGCCGCC TGGTGGATGG CTCGGCTCGG GGCGCTGACG AAGGGC
J23SpTT_12      NGCA GAAGACGGGT TTGATGGGGC GGGGATGTAA GCCGGGAAGGG AAACCGACCC GTTTAGTCTG CCGCTCCCAA CAGCCCGAGT TTCTTGCCCC TTGAGTAAGG GGCAAGAAAC CTAAGCCGCC TGGTGGATGG CTCGGCTCGG GGCGCTGACG AAGGGCN
J23SpTT_15      NGCA GAAGACGGGT TTGATGGGGC GGGGATGTAA GCCGGGAAGGG AAACCGACCC GTTTAGTCTG CCGCTCCCAA CAGCCCGAGT TTCTTGCCCC TTGAGTAAGG GGCAAGAAAC CTAAGCCGCC TGGTGGATGG CTCGGCTCGG GGCGCTGACG AAGGGN
J23SpTT_9       GGCA GAAGACGGGT TTGATGGGGC GGGGATGTAA GCCGGGAAGGG AAACCGACCC GTTTAGTCTG CCGCTCCCAA CAGCCCGAGT TTCTTGCCCC TTGAGTAAGG GGCAAGAAAC CTAAGCCGCC TGGTGGATGG CTCGGCTCGG GGCGCTGACG AAGGGC
J23SpTT_646      GCA GAAGACGGGT TTGATGGGGC GGGGATGTAA GCCGGGAAGGG AAACCGACCC GTTTAGTCTG CCGCTCCCAA CAGCCCGAGT TTCTTGCCCC TTGAGTAAGG GGCAAGAAAC CTAAGCCGCC TGGTGGATGG CTCGGCTCGG GGCGCTGACG AAGGG
J23SpTT_9         CA GAAGACGGGT TTGATGGGGC GGGGATGTAA GCCGGGAAGGG AAACCGACCC GTTTAGTCTG CCGCTCCCAA CAGCCCGAGT TTCTTGCCCC TTGAGTAAGG GGCAAGAAAC CTAAGCCGCC TGGTGGATGG CTCGGCTCGG GGCGCTGACG AAGGGCT
J23SpTT_19      GGCA GAAGACGGGT TTGATGGGGC GGGGATGTAA GCCGGGAAGGG AAACCGACCC GTTTAGTCTG CCGCTCCCAA CAGCCCGAGT TTCTTGCCCC TTGAGTAAGG GGCAAGAAAC CTAAGCCGCC TGGTGGATGG CTCGGCTCGG GGCGCTGACG AAGGG
J23SpTT_36      CGCA GAAGACGGGT TTGATGGGGC GGGGATGTAA GCCGGGAAGGG AAACCGACCC GTTTAGTCTG CCGCTCCCAA CAGCCCGAGT TTCTTGCCCC TTGAGTAAGG GGCAAGAAAC CTAAGCCGCC TGGTGGATGG CTCGGCTCGG GGCGCTGACG AAGGAC
J23SpTT_22        CA GAAGACGGGT TTGATGGGGC GGGGATGTAA GCCGGGAAGGG AAACCGACCC GTTTAGTCTG CCGCTCCCAA CAGCCCGAGT TTCTTGCCCC TTGAGTAAGG GGCAAGAAAC CTAAGCCGCC TGGTGGATGG CTCGGCTCGG GGCGCTGACG AAGGGCC
J23SpTT_13      CGCA GAAGACGGGT TTGATGGGGC GGGGATGTAA GCCGGGAAGGG AAACCGACCC GTTTAGTCTG CCGCTCCCAA CAGCCCGAGT TTCTTGCCCC TTGAGTAAGG GGCAAGAAAC CTAAGCCGCC TGGTGGATGG CTCGGCTCGG GGCGCTGACG AGGG
J23SpTT_12      CGCA GAAGACGGGT TTGATGGGGC GGGGATGTAA GCCGGGAAGGG AAACCGACCC GTTTAGTCTG CCGCTCCCAA CAGCCCGAGT TTCTTGCCCC TTGAGTAAGG GGCAAGAAAC CTAAGCCGCC TGGTGGATGG CTCGGCTCGG GGCGCTGACG AGGGG
J23SpTT_12      CCCA GAAGACGGGT TTGATGGGGC GGGGATGTAA GCCGGGAAGGG AAACCGACCC GTTTAGTCTG CCGCTCCCAA CAGCCCGAGT TTCTTGCCCC TTGAGTAAGG GGCAAGAAAC CTAAGCCGCC TGGTGGATGG CTCGGCTCGG GGCGCTGACG AAGG
J23SpTT_10      CGCA GAAGACGGGT TTGATGGGGC GGGGATGTAA GCCGGGAAGGG AAACCGACCC GTTTAGTCTG CCGCTCCCAA CAGCCCGAGT TTCTTGCCCC TTGAGTAAGG GGCAAGAAAC CTAAGCCGCC TGGTGGATGG CTCGGCTCGG GGCGCTGACG AAGC
J23SpTT_48      CGCA GAAGACGGGT TTGATGGGGC GGGGATGTAA GCCGGGAAGGG AAACCGACCC GTTTAGTCTG CCGCTCCCAA CAGCCCGAGT TTCTTGCCCC TTGAGTAAGG GGCAAGAAAC CTAAGCCGCC TGGTGGATGG CTCGGCTCGG GGCGCTGACA AAGG
J23SpTT_11      CGCA GAAGGCGGGT TTGATGGGGC GGGGATGTAA GCCGGGAAGGG AAACCGACCC GTTTAGTCTG CCGCTCCCAA CAGCCCGAGT TTCTTGCCCC TTGAGTAAGG GGCAAGAAAC CTAAGCCGCC TGGTGGATGG CTCGGCTCGG GGCGCTGACG AAGGG
J23SpTT_33      CGCA GAAGACGGGT TTGATGGGGC GGGGATGTAA GCCGGGAAGGG AAACCGACCC GTTTAGTCTG CCGCTCCCAA CAGCCCGAGT TTCTTGCCCC TTGAGTAAGG GGCAAGAAAC CTAAGCCGCC TGGTGGATGG CTCGGCTCGG GGCGCTGACG AAAG
J23SpTT_10     CCGCA GAAGACGGGT TTGATGGGGC GGGGATGTAA GCCGGGAAGGG AAACCGACCC GTTTAGTCTG CCGCTCCCAA CAGCCCGAGT TTCTTGCCCC TTGAGTAAGG GGCAAGAAAC CTAAGCCGCC TGGTGGATGG CTCGGCTCGG GGCGCTGACG AAAG
J23SpTT_12      CGCA GAAGACGGGT TTGATGGGGC GGGGATGTAA GCCGGGAAGGG AAACCGACCC GTTTAGTCTG CCGCTCCCAA CAGCCCGAGT TTCTTGCCCC TTGAGTAAGG GGCAAGAAAC CTAAGCCGCC TGGTGGATGG CTCGGCTCGG GGCGCTGACG AAAGG
J23SpTT_9       CGCA GAAGACGGGT TTGATGGGGC GGGGATGTAA GCCGGGAAGGG AAACCGACCC GTTTAGTCTG CCGCTCCCAA CAGCCCGAGT TTCTTGCCCC TTGAGTAAGG GGCAAGAAAC CTAAGCCGCC TGGTGGATGG CTCGGCTCGG GGCGCTGCCG AAGGG
J23SpTT_26      NGCA GAAGACGGGT TTGATGGGGC GGGGATGTAA GCCGGGAAGGG AAACCGACCC GTTTAGTCTG CCGCTCCCAA CAGCCCGAGT TTCTTGCCCC TTGAGTAAGG GGCAAGAAAC CTAAGCCGCC TGGTGGATGG CTCGGCTCGG GGCGCTGACG AAGGN
J23SpTT_14     NCGCA GAAGACGGGT TTGATGGGGC GGGGATGTAA GCCGGGAAGGG AAACCGACCC GTTTAGTCTG CCGCTCCCAA CAGCCCGAGT TTCTTGCCCC TTGAGTAAGG GGCAAGAAAC CTAAGCCGCC TGGTGGATGG CTCGGCTCGG GGCGCTGACG AAGGN
J23SpTT_10      CGCA GAAGACGGGT TTGATGGGGC GGGGATGTAA GCCGGGAAGGG AAACCGACCC GTTTAGTCTG CCGCTCCCAA CAGCCCGAGT TTCTTGCCCC TTGAGTAAGG GGCAAGAAAC CTAAGCCGCC TGGTGGATGG CTCGGCTCGG GGCGCTGACG AAGGN
J23SpTT_13      CGCA GAAGACGGGT TTGATGGGGC GGGGATGTAA GCCGGGAAGGG AAACCGACCC GTTTAGTCTG CCGCTCCCAA CAGCCCGAGT TTCTTGCCCC TTGAGTAAGG GGCAAGAAAC CTAAGCCGCC TGGTGGATGG CTCGGCTCGG GGCGCTGACG AAGGC
J23SpTT_8        GCA GAAGACGGGT TTGATGGGGC GGGGATGTAA GCCGGGAAGGG AAACCGACCC GTTTAGTCTG CCGCTCCCAA CAGCCCGAGT TTCTTGCCCC TTGAGTAAGG GGCAAGAAAC CTAAGCCGCC TGGTGGATGG CTCGGCTCGG GGCGCTGACG AAGAG
J23SpTT_2583      CA GAAGACGGGT TTGATGGGGC GGGGATGTAA GCCGGGAAGGG AAACCGACCC GTTTAGTCTG CCGCTCCCAA CAGCCCGAGT TTCTTGCCCC TTGAGTAAGG GGCAAGAAAC CTAAGCCGCC TGGTGGATGG CTCGGCTCGG GGCGCTGACG AAGGG
J23SpTT_78        CA GAAGACGGGT TTGATGGGGC GGGGATGTAA GCCGGGAAGGG AAACCGACCC GTTTAGTCTG CCGCTCCCAA CAGCCCGAGT TTCTTGCCCC TTGAGTAAGG GGCAAGAAAC CTAAGCCGCC TGGTGGATGG CTCGGCTCGG GGCGCTGACG AAGGGG
J23SpTT_37         A GAAGACGGGT TTGATGGGGC GGGGATGTAA GCCGGGAAGGG AAACCGACCC GTTTAGTCTG CCGCTCCCAA CAGCCCGAGT TTCTTGCCCC TTGAGTAAGG GGCAAGAAAC CTAAGCCGCC TGGTGGATGG CTCGGCTCGG GGCGCTGACG AAGGGG
J23SpTT_522        A GAAGACGGGT TTGATGGGGC GGGGATGTAA GCCGGGAAGGG AAACCGACCC GTTTAGTCTG CCGCTCCCAA CAGCCCGAGT TTCTTGCCCC TTGAGTAAGG GGCAAGAAAC CTAAGCCGCC TGGTGGATGG CTCGGCTCGG GGCGCTGACG AAGGGC
J23SpTT_469      GCA GAAGACGGGT TTGATGGGGC GGGGATGTAA GCCGGGAAGGG AAACCGACCC GTTTAGTCTG CCGCTCCCAA CAGCCCGAGT TTCTTGCCCC TTGAGTAAGG GGCAAGAAAC CTAAGCCGCC TGGTGGATGG CTCGGCTCGG GGCGCTGACG AAGG
```

```
J23SpTT_451        GCA GAAGACGGGT TTGATGGGGC GGGGATGTAA GCCGGGAAGG AAACCGACCC GTTTAGTCTG CCGCTCCCAA CAGCCCGAGT TTCTTGCCCC TTGAGTAAGG GGCAAGAAAC CTAAGCCGCC TGGTGGATGG CTCGGCTCGG GGCGCTGACG AAG
J23SpTT_17        NGCA GAAGACGGGT TTGATGGGGC GGGGATGTAA GCCGGGAAGG AAACCGACCC GTTTAGTCTG CCGCTCCCAA CAGCCCGAGT TTCTTGCCCC TTGAGTAAGG GGCAAGAAAC CTAAGCCGCC TGGTGGATGG CTCGGCTCGG GGCGCTGACG AAGN
J23SpTT_8         NGCA GAAGACGGGT TTGATGGGGC GGGGATGTAA GCCGGGAAGG AAACCGACCC GTTTAGTCTG CCGCTCCCAA CAGCCCGAGT TTCTTGCCCC TTGAGTAAGG GGCAAGAAAC CTAAGCCGCC TGGTGGATGG CTCGGCTCGG GGCGCTGACG AAG
J23SpTT_128       CGCA GAAGACGGGT TTGATGGGGC GGGGATGTAA GCCGGGAAGG AAACCGACCC GTTTAGTCTG CCGCTCCCAA CAGCCCGAGT TTCTTGCCCC TTGAGTAAGG GGCAAGAAAC CTAAGCCGCC TGGTGGATGG CTCGGCTCGG GGCGCTGACG AA
J23SpTT_98       CCGCA GAAGACGGGT TTGATGGGGC GGGGATGTAA GCCGGGAAGG AAACCGACCC GTTTAGTCTG CCGCTCCCAA CAGCCCGAGT TTCTTGCCCC TTGAGTAAGG GGCAAGAAAC CTAAGCCGCC TGGTGGATGG CTCGGCTCGG GGCGCTGACG AA
J23SpTT_65      CCCGCA GAAGACGGGT TTGATGGGGC GGGGATGTAA GCCGGGAAGG AAACCGACCC GTTTAGTCTG CCGCTCCCAA CAGCCCGAGT TTCTTGCCCC TTGAGTAAGG GGCAAGAAAC CTAAGCCGCC TGGTGGATGG CTCGGCTCGG GGCGCTGACG AA
J23SpTT_13      NCCGCA GAAGACGGGT TTGATGGGGC GGGGATGTAA GCCGGGAAGG AAACCGACCC GTTTAGTCTG CCGCTCCCAA CAGCCCGAGT TTCTTGCCCC TTGAGTAAGG GGCAAGAAAC CTAAGCCGCC TGGTGGATGG CTCGGCTCGG GGCGCTGACG AAN
J23SpTT_21          CA GAAGACGGGT TTGATGGGGC GGGGATGTAA GCCGGGAAGG AAACCGACCC GTTTAGTCTG CCGCTCCCAA CAGCCCGAGT TTCTTGCCCC TTGAGTAAGG GGCAAGAAAC CTAAGCCGCC TGGTGGATGG CTCGGCTCGG GGCGCTGACG AAGAG
J23SpTT_66        CGCA GAAGACGGGT TTGATGGGGC GGGGATGTAA GCCGGGAAGG AAACCGACCC GTTTAGTCTG CCGCTCCCAA CAGCCCGAGT TTCTTGCCCC TTGAGTAAGG GGCAAGAAAC CTAAGCCGCC TGGTGGATGG CTCGGCTCGG GGCGCTGACA AAG
J23SpTT_25       CCGCA GAAGACGGGT TTGATGGGGC GGGGATGTAA GCCGGGAAGG AAACCGACCC GTTTAGTCTG CCGCTCCCAA CAGCCCGAGT TTCTTGCCCC TTGAGTAAGG GGCAAGAAAC CTAAGCCGCC TGGTGGATGG CTCGGCTCGG GGCGCTGACA AAG
J23SpTT_25          CA GAAGACGGGT TTGATGGGGC GGGGATGTAA GCCGGGAAGG AAACCGACCC GTTTAGTCTG CCGCTCCCAA CAGCCCGAGT TTCTTGCCCC TTGAGTAAGG GGCAAGAAAC CTAAGCCGCC TGGTGGATGG CTCGGCTCGG GGCGCTGACA AAGGG
J23SpTT_23          CA GAAGACGGGT TTGATGGGGC GGGGATGTAA GCCGGGAAGG AAACCGACCC GTTTAGTCTG CCGCTCCCAA CAGCCCGAGT TTCTTGCCCC TTGAGTAAGG GGCAAGAAAC CTAAGCCGCC TGGTGGATGG CTCGGCTCGG GGCGCTGACG AAGGAC
J23SpTT_13       NCGCA GAAGACGGGT TTGATGGGGC GGGGATGTAA GCCGGGAAGG AAACCGACCC GTTTAGTCTG CCGCTCCCAA CAGCCCGAGT TTCTTGCCCC TTGAGTAAGG GGCAAGAAAC CTAAGCCGCC TGGTGGATGG CTCGGCTCGG GGCGCTGACG AAN
J23SpTT_10        CCCA GAAGACGGGT TTGATGGGGC GGGGATGTAA GCCGGGAAGG AAACCGACCC GTTTAGTCTG CCGCTCCCAA CAGCCCGAGT TTCTTGCCCC TTGAGTAAGG GGCAAGAAAC CTAAGCCGCC TGGTGGATGG CTCGGCTCGG GGCGCTGACG AAG
J23SpTT_357             CGGGT TTGATGGGGC GGGGATGTAA GCCGGGAAGG AAACCGACCC GTTTAGTCTG CCGCTCCCAA CAGCCCGAGT TTCTTGCCCC TTGAGTAAGG GGCAAGAAAC CTAAGCCGCC TGGTGGATGG CTCGGCTCGG GGCGCTGACG AAGGGCGTGG
J23SpTT_237           GACGGGT TTGATGGGGC GGGGATGTAA GCCGGGAAGG AAACCGACCC GTTTAGTCTG CCGCTCCCAA CAGCCCGAGT TTCTTGCCCC TTGAGTAAGG GGCAAGAAAC CTAAGCCGCC TGGTGGATGG CTCGGCTCGG GGCGCTGACG AAGGGCGTGG
J23SpTT_115            ACGGGT TTGATGGGGC GGGGATGTAA GCCGGGAAGG AAACCGACCC GTTTAGTCTG CCGCTCCCAA CAGCCCGAGT TTCTTGCCCC TTGAGTAAGG GGCAAGAAAC CTAAGCCGCC TGGTGGATGG CTCGGCTCGG GGCGCTGACG AAGGGCGTGG
J23SpTT_114           GACGGGT TTGATGGGGC GGGGATGTAA GCCGGGAAGG AAACCGACCC GTTTAGTCTG CCGCTCCCAA CAGCCCGAGT TTCTTGCCCC TTGAGTAAGG GGCAAGAAAC CTAAGCCGCC TGGTGGATGG CTCGGCTCGG GGCGCTGACG AAGGGCGTG
J23SpTT_19             ACGGGT TTGATGGGGC GGGGATGTAA GCCGGGAAGG AAACCGACCC GTTTAGTCTG CCGCTCCCAA CAGCCCGAGT TTCTTGCCCC TTGAGTAAGG GGCAAGAAAC CTAAGCCGCC TGGTGGATGG CTCGGCTCGG GGCGCTGACG AAGGGCGTGG C
J23SpTT_76              CGGGT TTGATGGGGC GGGGATGTAA GCCGGGAAGG AAACCGACCC GTTTAGTCTG CCGCTCCCAA CAGCCCGAGT TTCTTGCCCC TTGAGTAAGG GGCAAGAAAC CTAAGCCGCC TGGTGGATGG CTCGGCTCGG GGCGCTGACG AAGGGCGTGG C
J23SpTT_15              CGGGT TTGATGGGGC GGGGATGTAA GCCGGGAAGG AAACCGACCC GTTTAGTCTG CCGCTCCCAA CAGCCCGAGT TTCTTGCCCC TTGAGTAAGG GGCAAGAAAC CTAAGCCGCC TGGTGGATGG CTCGGCTCGG GGCGCTGACG AAGGGCGTGG CG
J23SpTT_50             ACGGGT TTGATGGGGC GGGGATGTAA GCCGGGAAGG AAACCGACCC GTTTAGTCTG CCGCTCCCAA CAGCCCGAGT TTCTTGCCCC TTGAGTAAGG GGCAAGAAAC CTAAGCCGCC TGGTGGATGG CTCGGCTCGG GGCGCTGACG AAGGGCGTG
J23SpTT_206             CGGGT TTGATGGGGC GGGGATGTAA GCCGGGAAGG AAACCGACCC GTTTAGTCTG CCGCTCCCAA CAGCCCGAGT TTCTTGCCCC TTGAGTAAGG GGCAAGAAAC CTAAGCCGCC TGGTGGATGG CTCGGCTCGG GGCGCTGACG AAGGGCGTG
J23SpTT_8                GGGT TTGATGGGGC GGGGATGTAA GCCGGGAAGG AAACCGACCC GTTTAGTCTG CCGCTCCCAA CAGCCCGAGT TTCTTGCCCC TTGAGTAAGG GGCAAGAAAC CTAAGCCGCC TGGTGGATGG CTCGGCTCGG GGCGCTGACG AAGGGCGTGG C
J23SpTT_36               GGGT TTGATGGGGC GGGGATGTAA GCCGGGAAGG AAACCGACCC GTTTAGTCTG CCGCTCCCAA CAGCCCGAGT TTCTTGCCCC TTGAGTAAGG GGCAAGAAAC CTAAGCCGCC TGGTGGATGG CTCGGCTCGG GGCGCTGACG AAGGGCGTGG
J23SpTT_11            GACGGGT TTGATGGGGC GGGGATGTAA GCCGGGAAGG AAACCGACCC GTTTAGTCTG CCGCTCCCAA CAGCCCGAGT TTCTTGCCCC TTGAGTAAGG GGCAAGAAAC CTAAGCCGCC TGGTGGATGG CTCGGCTCGG GGCGCTGACG AAGGGCGT
J23SpTT_1782        CA GAAGACGGGT TTGATGGGGC GGGGATGTAA GCCGGGAAGG AAACCGACCC GTTTAGTCTG CCGCTCCCAA CAGCCCGAGT TTCTTGCCCC TTGAGTAAGG GGCAAGAAAC CTAAGCCGCC TGGTGGATGG CTCGGCTCGG GGCGCTGACG AAG
J23SpTT_1653        CA GAAGACGGGT TTGATGGGGC GGGGATGTAA GCCGGGAAGG AAACCGACCC GTTTAGTCTG CCGCTCCCAA CAGCCCGAGT TTCTTGCCCC TTGAGTAAGG GGCAAGAAAC CTAAGCCGCC TGGTGGATGG CTCGGCTCGG GGCGCTGACG AAGG
J23SpTT_1069         A GAAGACGGGT TTGATGGGGC GGGGATGTAA GCCGGGAAGG AAACCGACCC GTTTAGTCTG CCGCTCCCAA CAGCCCGAGT TTCTTGCCCC TTGAGTAAGG GGCAAGAAAC CTAAGCCGCC TGGTGGATGG CTCGGCTCGG GGCGCTGACG AAGGG
J23SpTT_10          CA GAAGACGGGT TTGATGGGGC GGGGATGTAA GCCGGGAAGG AAACCGACCC GTTTAGTCTG CCGCTCCCAA CAGCCCGAGT TTCTTGCCCC TTGAGTAAGG GGCAAGAAAC CTAAGCCGCC TGGTGGATGG CTCGGCTCGG GGCGCTGACA AAGG
J23SpTT_683          A GAAGACGGGT TTGATGGGGC GGGGATGTAA GCCGGGAAGG AAACCGACCC GTTTAGTCTG CCGCTCCCAA CAGCCCGAGT TTCTTGCCCC TTGAGTAAGG GGCAAGAAAC CTAAGCCGCC TGGTGGATGG CTCGGCTCGG GGCGCTGACG AAGG
J23SpTT_16          CA GAAGACGGGT TTGATGGGGC GGGGATGTAA GCCGGGAAGG AAACCGACCC GTTTAGTCTG CCGCTCCCAA CAGCCCGAGT TTCTTGCCCC TTGAGTAAGG GGCAAGAAAC CTAAGCCGCC TGGTGGATGG CTCGGCTCGG GGCGCTGACG AAAG
J23SpTT_9          GCA GAAGACGGGT TTGATGGGGC GGGGATGTAA GCCGGGAAGG AAACCGACCC GTTTAGTCTG CCGCTCCCAA CAGCCCGAGT TTCTTGCCCC TTGAGTAAGG GGCAAGAAAC CTAAGCCGCC TGGTGGATGG CTCGGCTCGG GGCGCTGACG AAAG
J23SpTT_12          NA GAAGACGGGT TTGATGGGGC GGGGATGTAA GCCGGGAAGG AAACCGACCC GTTTAGTCTG CCGCTCCCAA CAGCCCGAGT TTCTTGCCCC TTGAGTAAGG GGCAAGAAAC CTAAGCCGCC TGGTGGATGG CTCGGCTCGG GGCGCTGACG AAGGN
J23SpTT_729          A GAAGACGGGT TTGATGGGGC GGGGATGTAA GCCGGGAAGG AAACCGACCC GTTTAGTCTG CCGCTCCCAA CAGCCCGAGT TTCTTGCCCC TTGAGTAAGG GGCAAGAAAC CTAAGCCGCC TGGTGGATGG CTCGGCTCGG GGCGCTGACG AAG
J23SpTT_137            GAAGACGGGT TTGATGGGGC GGGGATGTAA GCCGGGAAGG AAACCGACCC GTTTAGTCTG CCGCTCCCAA CAGCCCGAGT TTCTTGCCCC TTGAGTAAGG GGCAAGAAAC CTAAGCCGCC TGGTGGATGG CTCGGCTCGG GGCGCTGACG AAGGG
J23SpTT_102            GAAGACGGGT TTGATGGGGC GGGGATGTAA GCCGGGAAGG AAACCGACCC GTTTAGTCTG CCGCTCCCAA CAGCCCGAGT TTCTTGCCCC TTGAGTAAGG GGCAAGAAAC CTAAGCCGCC TGGTGGATGG CTCGGCTCGG GGCGCTGACG AAGGGC
J23SpTT_100             AAGACGGGT TTGATGGGGC GGGGATGTAA GCCGGGAAGG AAACCGACCC GTTTAGTCTG CCGCTCCCAA CAGCCCGAGT TTCTTGCCCC TTGAGTAAGG GGCAAGAAAC CTAAGCCGCC TGGTGGATGG CTCGGCTCGG GGCGCTGACG AAGGGC
J23SpTT_65              AAGACGGGT TTGATGGGGC GGGGATGTAA GCCGGGAAGG AAACCGACCC GTTTAGTCTG CCGCTCCCAA CAGCCCGAGT TTCTTGCCCC TTGAGTAAGG GGCAAGAAAC CTAAGCCGCC TGGTGGATGG CTCGGCTCGG GGCGCTGACG AAGGGCG
J23SpTT_10          NA GAAGACGGGT TTGATGGGGC GGGGATGTAA GCCGGGAAGG AAACCGACCC GTTTAGTCTG CCGCTCCCAA CAGCCCGAGT TTCTTGCCCC TTGAGTAAGG GGCAAGAAAC CTAAGCCGCC TGGTGGATGG CTCGGCTCGG GGCGCTGACG AAGN
J23SpTT_90             GAAGACGGGT TTGATGGGGC GGGGATGTAA GCCGGGAAGG AAACCGACCC GTTTAGTCTG CCGCTCCCAA CAGCCCGAGT TTCTTGCCCC TTGAGTAAGG GGCAAGAAAC CTAAGCCGCC TGGTGGATGG CTCGGCTCGG GGCGCTGACG AAGG
J23SpTT_289           GACGGGT TTGATGGGGC GGGGATGTAA GCCGGGAAGG AAACCGACCC GTTTAGTCTG CCGCTCCCAA CAGCCCGAGT TTCTTGCCCC TTGAGTAAGG GGCAAGAAAC CTAAGCCGCC TGGTGGATGG CTCGGCTCGG GGCGCTGACG AAGGGCG
J23SpTT_97           AGACGGGT TTGATGGGGC GGGGATGTAA GCCGGGAAGG AAACCGACCC GTTTAGTCTG CCGCTCCCAA CAGCCCGAGT TTCTTGCCCC TTGAGTAAGG GGCAAGAAAC CTAAGCCGCC TGGTGGATGG CTCGGCTCGG GGCGCTGACG AAGGGCG
J23SpTT_199          AGACGGGT TTGATGGGGC GGGGATGTAA GCCGGGAAGG AAACCGACCC GTTTAGTCTG CCGCTCCCAA CAGCCCGAGT TTCTTGCCCC TTGAGTAAGG GGCAAGAAAC CTAAGCCGCC TGGTGGATGG CTCGGCTCGG GGCGCTGACG AAGGGC
J23SpTT_14           A GAAGACGGGT TTGATGGGGC GGGGATGTAA GCCGGGAAGG AAACCGACCC GTTTAGTCTG CCGCTCCCAA CAGCCCGAGT TTCTTGCCCC TTGAGTAAGG GGCAAGAAAC CTAAGCCGCC TGGTGGATGG CTCGGCTCGG GGCGCTGACG AA
J23SpTT_221             AAGACGGGT TTGATGGGGC GGGGATGTAA GCCGGGAAGG AAACCGACCC GTTTAGTCTG CCGCTCCCAA CAGCCCGAGT TTCTTGCCCC TTGAGTAAGG GGCAAGAAAC CTAAGCCGCC TGGTGGATGG CTCGGCTCGG GGCGCTGACG AAGGG
J23SpTT_9               AAGACGGGT TTGATGGGGC GGGGATGTAA GCCGGGAAGG AAACCGACCC GTTTAGTCTG CCGCTCCCAA CAGCCCGAGT TTCTTGCCCC TTGAGTAAGG GGCAAGAAAC CTAAGCCGCC TGGTGGATGG CTCGGCTCGG GGCGCTGACG AAGGGG
J23SpTT_16           AGACGGGT TTGATGGGGC GGGGATGTAA GCCGGGAAGG AAACCGACCC GTTTAGTCTG CCGCTCCCAA CAGCCCGAGT TTCTTGCCCC TTGAGTAAGG GGCAAGAAAC CTAAGCCGCC TGGTGGATGG CTCGGCTCGG GGCGCTGACG AAGGGG
J23SpTT_144             AAGACGGGT TTGATGGGGC GGGGATGTAA GCCGGGAAGG AAACCGACCC GTTTAGTCTG CCGCTCCCAA CAGCCCGAGT TTCTTGCCCC TTGAGTAAGG GGCAAGAAAC CTAAGCCGCC TGGTGGATGG CTCGGCTCGG GGCGCTGACG AAGG
J23SpTT_99             GAAGACGGGT TTGATGGGGC GGGGATGTAA GCCGGGAAGG AAACCGACCC GTTTAGTCTG CCGCTCCCAA CAGCCCGAGT TTCTTGCCCC TTGAGTAAGG GGCAAGAAAC CTAAGCCGCC TGGTGGATGG CTCGGCTCGG GGCGCTGACG AAG
J23SpTT_30          CA GAAGACGGGT TTGATGGGGC GGGGATGTAA GCCGGGAAGG AAACCGACCC GTTTAGTCTG CCGCTCCCAA CAGCCCGAGT TTCTTGCCCC TTGAGTAAGG GGCAAGAAAC CTAAGCCGCC TGGTGGATGG CTCGGCTCGG GGCGCTGACA AAG
J23SpTT_57          CA GAAGACGGGT TTGATGGGGC GGGGATGTAA GCCGGGAAGG AAACCGACCC GTTTAGTCTG CCGCTCCCAA CAGCCCGAGT TTCTTGCCCC TTGAGTAAGG GGCAAGAAAC CTAAGCCGCC TGGTGGATGG CTCGGCTCGG GGCGCTGACG AA
J23SpTT_8           NA GAAGACGGGT TTGATGGGGC GGGGATGTAA GCCGGGAAGG AAACCGACCC GTTTAGTCTG CCGCTCCCAA CAGCCCGAGT TTCTTGCCCC TTGAGTAAGG GGCAAGAAAC CTAAGCCGCC TGGTGGATGG CTCGGCTCGG GGCGCTGACG AAN
J23SpTT_197       CGCA GAAGACGGGT TTGATGGGGC GGGGATGTAA GCCGGGAAGG AAACCGACCC GTTTAGTCTG CCGCTCCCAA CAGCCCGAGT TTCTTGCCCC TTGAGTAAGG GGCAAGAAAC CTAAGCCGCC TGGTGGATGG CTCGGCTCGG GGCGCTGACG
J23SpTT_98       CCGCA GAAGACGGGT TTGATGGGGC GGGGATGTAA GCCGGGAAGG AAACCGACCC GTTTAGTCTG CCGCTCCCAA CAGCCCGAGT TTCTTGCCCC TTGAGTAAGG GGCAAGAAAC CTAAGCCGCC TGGTGGATGG CTCGGCTCGG GGCGCTGACG
J23SpTT_16        CGCA GAAGACGGGT TTGATGGGGC GGGGATGTAA GCCGGGAAGG AAACCGACCC GTTTAGTCTG CCGCTCCCAA CAGCCCGAGT TTCTTGCCCC TTGAGTAAGG GGCAAGAAAC CTAAGCCGCC TGGTGGATGG CTCGGCTCGG GGCGCTGACG AG
J23SpTT_9        CCGCA GAAGACGGGT TTGATGGGGC GGGGATGTAA GCCGGGAAGG AAACCGACCC GTTTAGTCTG CCGCTCCCAA CAGCCCGAGT TTCTTGCCCC TTGAGTAAGG GGCAAGAAAC CTAAGCCGCC TGGTGGATGG CTCGGCTCGG GGCGCTGACG A
J23SpTT_82      CCCGCA GAAGACGGGT TTGATGGGGC GGGGATGTAA GCCGGGAAGG AAACCGACCC GTTTAGTCTG CCGCTCCCAA CAGCCCGAGT TTCTTGCCCC TTGAGTAAGG GGCAAGAAAC CTAAGCCGCC TGGTGGATGG CTCGGCTCGG GGCGCTGACG
J23SpTT_29        CGCA GAAGACGGGT TTGATGGGGC GGGGATGTAA GCCGGGAAGG AAACCGACCC GTTTAGTCTG CCGCTCCCAA CAGCCCGAGT TTCTTGCCCC TTGAGTAAGG GGCAAGAAAC CTAAGCCGCC TGGTGGATGG CTCGGCTCGG GGCGCTGACG A
J23SpTT_52       CCGCA GAAGACGGGT TTGATGGGGC GGGGATGTAA GCCGGGAAGG AAACCGACCC GTTTAGTCTG CCGCTCCCAA CAGCCCGAGT TTCTTGCCCC TTGAGTAAGG GGCAAGAAAC CTAAGCCGCC TGGTGGATGG CTCGGCTCGG GGCGCTGAC
J23SpTT_40      CCCGCA GAAGACGGGT TTGATGGGGC GGGGATGTAA GCCGGGAAGG AAACCGACCC GTTTAGTCTG CCGCTCCCAA CAGCCCGAGT TTCTTGCCCC TTGAGTAAGG GGCAAGAAAC CTAAGCCGCC TGGTGGATGG CTCGGCTCGG GGCGCTGAC
J23SpTT_29         GCA GAAGACGGGT TTGATGGGGC GGGGATGTAA GCCGGGAAGG AAACCGACCC GTTTAGTCTG CCGCTCCCAA CAGCCCGAGT TTCTTGCCCC TTGAGTAAGG GGCAAGAAAC CTAAGCCGCC TGGTGGATGG CTCGGCTCGG GGCGCTGACG AA
J23SpTT_16        NGCA GAAGACGGGT TTGATGGGGC GGGGATGTAA GCCGGGAAGG AAACCGACCC GTTTAGTCTG CCGCTCCCAA CAGCCCGAGT TTCTTGCCCC TTGAGTAAGG GGCAAGAAAC CTAAGCCGCC TGGTGGATGG CTCGGCTCGG GGCGCTGACG AAN
J23SpTT_116       CGCA GAAGACGGGT TTGATGGGGC GGGGATGTAA GCCGGGAAGG AAACCGACCC GTTTAGTCTG CCGCTCCCAA CAGCCCGAGT TTCTTGCCCC TTGAGTAAGG GGCAAGAAAC CTAAGCCGCC TGGTGGATGG CTCGGCTCGG GGCGCTGAC
J23SpTT_19         GCA GAAGACGGGT TTGATGGGGC GGGGATGTAA GCCGGGAAGG AAACCGACCC GTTTAGTCTG CCGCTCCCAA CAGCCCGAGT TTCTTGCCCC TTGAGTAAGG GGCAAGAAAC CTAAGCCGCC TGGTGGATGG CTCGGCTCGG GGCGCTGACG
J23SpTT_13          CA GAAGACGGGT TTGATGGGGC GGGGATGTAA GCCGGGAAGG AAACCGACCC GTTTAGTCTG CCGCTCCCAA CAGCCCGAGT TTCTTGCCCC TTGAGTAAGG GGCAAGAAAC CTAAGCCGCC TGGTGGATGG CTCGGCTCGG GGCGCTGACG A
J23SpTT_70                 GT TTGATGGGGC GGGGATGTAA GCCGGGAAGG AAACCGACCC GTTTAGTCTG CCGCTCCCAA CAGCCCGAGT TTCTTGCCCC TTGAGTAAGG GGCAAGAAAC CTAAGCCGCC TGGTGGATGG CTCGGCTCGG GGCGCTGACG AAGGGCGTGG
J23SpTT_21                 GT TTGATGGGGC GGGGATGTAA GCCGGGAAGG AAACCGACCC GTTTAGTCTG CCGCTCCCAA CAGCCCGAGT TTCTTGCCCC TTGAGTAAGG GGCAAGAAAC CTAAGCCGCC TGGTGGATGG CTCGGCTCGG GGCGCTGACG AAGGGCGTGG C
J23SpTT_8                  GT TTGATGGGGC GGGGATGTAA GCCGGGAAGG AAACCGACCC GTTTAGTCTG CCGCTCCCAA CAGCCCGAGT TTCTTGCCCC TTGAGTAAGG GGCAAGAAAC CTAAGCCGCC TGGTGGATGG CTCGGCTCGG GGCGCTGACG AAGGGCGTGG CAAG
J23SpTT_23                GGT TTGATGGGGC GGGGATGTAA GCCGGGAAGG AAACCGACCC GTTTAGTCTG CCGCTCCCAA CAGCCCGAGT TTCTTGCCCC TTGAGTAAGG GGCAAGAAAC CTAAGCCGCC TGGTGGATGG CTCGGCTCGG GGCGCTGACG AAGGGCGTGG
J23SpTT_53                 GT TTGATGGGGC GGGGATGTAA GCCGGGAAGG AAACCGACCC GTTTAGTCTG CCGCTCCCAA CAGCCCGAGT TTCTTGCCCC TTGAGTAAGG GGCAAGAAAC CTAAGCCGCC TGGTGGATGG CTCGGCTCGG GGCGCTGACG AAGGGCGTG
J23SpTT_13                GGT TTGATGGGGC GGGGATGTAA GCCGGGAAGG AAACCGACCC GTTTAGTCTG CCGCTCCCAA CAGCCCGAGT TTCTTGCCCC TTGAGTAAGG GGCAAGAAAC CTAAGCCGCC TGGTGGATGG CTCGGCTCGG GGCGCTGACG AAGGGCGTG
J23SpTT_1361            CGGGT TTGATGGGGC GGGGATGTAA GCCGGGAAGG AAACCGACCC GTTTAGTCTG CCGCTCCCAA CAGCCCGAGT TTCTTGCCCC TTGAGTAAGG GGCAAGAAAC CTAAGCCGCC TGGTGGATGG CTCGGCTCGG GGCGCTGACG AAGGG
J23SpTT_886           GACGGGT TTGATGGGGC GGGGATGTAA GCCGGGAAGG AAACCGACCC GTTTAGTCTG CCGCTCCCAA CAGCCCGAGT TTCTTGCCCC TTGAGTAAGG GGCAAGAAAC CTAAGCCGCC TGGTGGATGG CTCGGCTCGG GGCGCTGACG AAGGG
J23SpTT_474           GACGGGT TTGATGGGGC GGGGATGTAA GCCGGGAAGG AAACCGACCC GTTTAGTCTG CCGCTCCCAA CAGCCCGAGT TTCTTGCCCC TTGAGTAAGG GGCAAGAAAC CTAAGCCGCC TGGTGGATGG CTCGGCTCGG GGCGCTGACG AAGGGC
J23SpTT_9             GACGGGT TTGATGGGGC GGGGATGTAA GCCGGGAAGG AAACCGACCC GTTTAGTCTG CCGCTCCCAA CAGCCCGAGT TTCTTGCCCC TTGAGTAAGG GGCAAGAAAC CTAAGCCGCC TGGTGGATGG CTCGGCTCGG GGCGCTGACG AAGGGCA
J23SpTT_361          AGACGGGT TTGATGGGGC GGGGATGTAA GCCGGGAAGG AAACCGACCC GTTTAGTCTG CCGCTCCCAA CAGCCCGAGT TTCTTGCCCC TTGAGTAAGG GGCAAGAAAC CTAAGCCGCC TGGTGGATGG CTCGGCTCGG GGCGCTGACG AAGGG
J23SpTT_247            ACGGGT TTGATGGGGC GGGGATGTAA GCCGGGAAGG AAACCGACCC GTTTAGTCTG CCGCTCCCAA CAGCCCGAGT TTCTTGCCCC TTGAGTAAGG GGCAAGAAAC CTAAGCCGCC TGGTGGATGG CTCGGCTCGG GGCGCTGACG AAGGGC
J23SpTT_120            ACGGGT TTGATGGGGC GGGGATGTAA GCCGGGAAGG AAACCGACCC GTTTAGTCTG CCGCTCCCAA CAGCCCGAGT TTCTTGCCCC TTGAGTAAGG GGCAAGAAAC CTAAGCCGCC TGGTGGATGG CTCGGCTCGG GGCGCTGACG AAGGGCG
J23SpTT_31            GACGGGT TTGATGGGGC GGGGATGTAA GCCGGGAAGG AAACCGACCC GTTTAGTCTG CCGCTCCCAA CAGCCCGAGT TTCTTGCCCC TTGAGTAAGG GGCAAGAAAC CTAAGCCGCC TGGTGGATGG CTCGGCTCGG GGCGCTGACG AAGGGG
J23SpTT_772             CGGGT TTGATGGGGC GGGGATGTAA GCCGGGAAGG AAACCGACCC GTTTAGTCTG CCGCTCCCAA CAGCCCGAGT TTCTTGCCCC TTGAGTAAGG GGCAAGAAAC CTAAGCCGCC TGGTGGATGG CTCGGCTCGG GGCGCTGACG AAGGGC
J23SpTT_598           GACGGGT TTGATGGGGC GGGGATGTAA GCCGGGAAGG AAACCGACCC GTTTAGTCTG CCGCTCCCAA CAGCCCGAGT TTCTTGCCCC TTGAGTAAGG GGCAAGAAAC CTAAGCCGCC TGGTGGATGG CTCGGCTCGG GGCGCTGACG AAGG
J23SpTT_464             CGGGT TTGATGGGGC GGGGATGTAA GCCGGGAAGG AAACCGACCC GTTTAGTCTG CCGCTCCCAA CAGCCCGAGT TTCTTGCCCC TTGAGTAAGG GGCAAGAAAC CTAAGCCGCC TGGTGGATGG CTCGGCTCGG GGCGCTGACG AAGGGCG
J23SpTT_9               CGGGT TTGATGGGGC GGGGATGTAA GCCGGGAAGG AAACCGACCC GTTTAGTCTG CCGCTCCCAA CAGCCCGAGT TTCTTGCCCC TTGAGTAAGG GGCAAGAAAC CTAAGCCGCC TGGTGGATGG CTCGGCTCGG GGCGCTGACG AAGGGCGT
J23SpTT_29               GGGT TTGATGGGGC GGGGATGTAA GCCGGGAAGG AAACCGACCC GTTTAGTCTG CCGCTCCCAA CAGCCCGAGT TTCTTGCCCC TTGAGTAAGG GGCAAGAAAC CTAAGCCGCC TGGTGGATGG CTCGGCTCGG GGCGCTGACG AAGGGCGTG
J23SpTT_353            ACGGGT TTGATGGGGC GGGGATGTAA GCCGGGAAGG AAACCGACCC GTTTAGTCTG CCGCTCCCAA CAGCCCGAGT TTCTTGCCCC TTGAGTAAGG GGCAAGAAAC CTAAGCCGCC TGGTGGATGG CTCGGCTCGG GGCGCTGACG AAGGG
J23SpTT_255          AGACGGGT TTGATGGGGC GGGGATGTAA GCCGGGAAGG AAACCGACCC GTTTAGTCTG CCGCTCCCAA CAGCCCGAGT TTCTTGCCCC TTGAGTAAGG GGCAAGAAAC CTAAGCCGCC TGGTGGATGG CTCGGCTCGG GGCGCTGACG AAGG
J23SpTT_11              CGGGT TTGATGGGGC GGGGATGTAA GCCGGGAAGG AAACCGACCC GTTTAGTCTG CCGCTCCCAA CAGCCCGAGT TTCTTGCCCC TTGAGTAAGG GGCAAGAAAC CTAAGCCGCC TGGTGGATGG CTCGGCTCGG GGCGCTGACG AAGGGCC
J23SpTT_14             ACGGGT TTGATGGGGC GGGGATGTAA GCCGGGAAGG AAACCGACCC GTTTAGTCTG CCGCTCCCAA CAGCCCGAGT TTCTTGCCCC TTGAGTAAGG GGCAAGAAAC CTAAGCCGCC TGGTGGATGG CTCGGCTCGG GGCGCTGACG AAGGGG
J23SpTT_16            GACGGGT TTGATGGGGC GGGGATGTAA GCCGGGAAGG AAACCGACCC GTTTAGTCTG CCGCTCCCAA CAGCCCGAGT TTCTTGCCCC TTGAGTAAGG GGCAAGAAAC CTAAGCCGCC TGGTGGATGG CTCGGCTCGG GGCGCTGACG AAGAG
J23SpTT_10              CGGGT TTGATGGGGC GGGGATGTAA GCCGGGAAGG AAACCGACCC GTTTAGTCTG CCGCTCCCAA CAGCCCGAGT TTCTTGCCCC TTGAGTAAGG GGCAAGAAAC CTAAGCCGCC TGGTGGATGG CTCGGCTCGG GGCGCTGACG AAGGGCA
J23SpTT_602           GACGGGT TTGATGGGGC GGGGATGTAA GCCGGGAAGG AAACCGACCC GTTTAGTCTG CCGCTCCCAA CAGCCCGAGT TTCTTGCCCC TTGAGTAAGG GGCAAGAAAC CTAAGCCGCC TGGTGGATGG CTCGGCTCGG GGCGCTGACG AAG
J23SpTT_263          AGACGGGT TTGATGGGGC GGGGATGTAA GCCGGGAAGG AAACCGACCC GTTTAGTCTG CCGCTCCCAA CAGCCCGAGT TTCTTGCCCC TTGAGTAAGG GGCAAGAAAC CTAAGCCGCC TGGTGGATGG CTCGGCTCGG GGCGCTGACG AAG
J23SpTT_177         AAGACGGGT TTGATGGGGC GGGGATGTAA GCCGGGAAGG AAACCGACCC GTTTAGTCTG CCGCTCCCAA CAGCCCGAGT TTCTTGCCCC TTGAGTAAGG GGCAAGAAAC CTAAGCCGCC TGGTGGATGG CTCGGCTCGG GGCGCTGACG AAG
J23SpTT_216            ACGGGT TTGATGGGGC GGGGATGTAA GCCGGGAAGG AAACCGACCC GTTTAGTCTG CCGCTCCCAA CAGCCCGAGT TTCTTGCCCC TTGAGTAAGG GGCAAGAAAC CTAAGCCGCC TGGTGGATGG CTCGGCTCGG GGCGCTGACG AAGG
J23SpTT_9             GACGGGT TTGATGGGGC GGGGATGTAA GCCGGGAAGG AAACCGACCC GTTTAGTCTG CCGCTCCCAA CAGCCCGAGT TTCTTGCCCC TTGAGTAAGG GGCAAGAAAC CTAAGCCGCC TGGTGGATGG CTCGGCTCGG GGCGCTGACA AAGG
J23SpTT_68               GGGT TTGATGGGGC GGGGATGTAA GCCGGGAAGG AAACCGACCC GTTTAGTCTG CCGCTCCCAA CAGCCCGAGT TTCTTGCCCC TTGAGTAAGG GGCAAGAAAC CTAAGCCGCC TGGTGGATGG CTCGGCTCGG GGCGCTGACG AAGGGC
J23SpTT_53              CGGGT TTGATGGGGC GGGGATGTAA GCCGGGAAGG AAACCGACCC GTTTAGTCTG CCGCTCCCAA CAGCCCGAGT TTCTTGCCCC TTGAGTAAGG GGCAAGAAAC CTAAGCCGCC TGGTGGATGG CTCGGCTCGG GGCGCTGACG AAGGGG
```

```
J23SpTT_51                   GGGT TTGATGGGGC GGGGATGTAA GCCGGGAAGGG AAACCGACCC GTTTAGTCTG CCGCTCCCAA CAGCCCGAGT TTCTTGCCCC TTGAGTAAGG GGCAAGAAAC CTAAGCCGCC TGGTGGATGG CTCGGCTCGG GGCGCTGACG AAGGGCG
J23SpTT_75          CA GAAGACGGGT TTGATGGGGC GGGGATGTAA GCCGGGAAGGG AAACCGACCC GTTTAGTCTG CCGCTCCCAA CAGCCCGAGT TTCTTGCCCC TTGAGTAAGG GGCAAGAAAC CTAAGCCGCC TGGTGGATGG CTCGGCTCGG GGCGCTGACG
J23SpTT_21         GCA GAAGACGGGT TTGATGGGGC GGGGATGTAA GCCGGGAAGGG AAACCGACCC GTTTAGTCTG CCGCTCCCAA CAGCCCGAGT TTCTTGCCCC TTGAGTAAGG GGCAAGAAAC CTAAGCCGCC TGGTGGATGG CTCGGCTCGG GGCGCTGAC
J23SpTT_46          CA GAAGACGGGT TTGATGGGGC GGGGATGTAA GCCGGGAAGGG AAACCGACCC GTTTAGTCTG CCGCTCCCAA CAGCCCGAGT TTCTTGCCCC TTGAGTAAGG GGCAAGAAAC CTAAGCCGCC TGGTGGATGG CTCGGCTCGG GGCGCTGAC
J23SpTT_34           A GAAGACGGGT TTGATGGGGC GGGGATGTAA GCCGGGAAGGG AAACCGACCC GTTTAGTCTG CCGCTCCCAA CAGCCCGAGT TTCTTGCCCC TTGAGTAAGG GGCAAGAAAC CTAAGCCGCC TGGTGGATGG CTCGGCTCGG GGCGCTGACG
J23SpTT_18           A GAAGACGGGT TTGATGGGGC GGGGATGTAA GCCGGGAAGGG AAACCGACCC GTTTAGTCTG CCGCTCCCAA CAGCCCGAGT TTCTTGCCCC TTGAGTAAGG GGCAAGAAAC CTAAGCCGCC TGGTGGATGG CTCGGCTCGG GGCGCTGAC
J23SpTT_17        CGCA GAAGACGGGT TTGATGGGGC GGGGATGTAA GCCGGGAAGGG AAACCGACCC GTTTAGTCTG CCGCTCCCAA CAGCCCGAGT TTCTTGCCCC TTGAGTAAGG GGCAAGAAAC CTAAGCCGCC TGGTGGATGG CTCGGCTCGG GGCGCTG
J23SpTT_11        CGCA GAAGACGGGT TTGATGGGGC GGGGATGTAA GCCGGGAAGGG AAACCGACCC GTTTAGTCTG CCGCTCCCAA CAGCCCGAGT TTCTTGCCCC TTGAGTAAGG GGCAAGAAAC CTAAGCCGCC TGGTGGATGG CTCGGCTCGG GGCGCT
J23SpTT_178                    GT TTGATGGGGC GGGGATGTAA GCCGGGAAGGG AAACCGACCC GTTTAGTCTG CCGCTCCCAA CAGCCCGAGT TTCTTGCCCC TTGAGTAAGG GGCAAGAAAC CTAAGCCGCC TGGTGGATGG CTCGGCTCGG GGCGCTGACG AAGGGC
J23SpTT_127                    GT TTGATGGGGC GGGGATGTAA GCCGGGAAGGG AAACCGACCC GTTTAGTCTG CCGCTCCCAA CAGCCCGAGT TTCTTGCCCC TTGAGTAAGG GGCAAGAAAC CTAAGCCGCC TGGTGGATGG CTCGGCTCGG GGCGCTGACG AAGGGCG
J23SpTT_25                    GGT TTGATGGGGC GGGGATGTAA GCCGGGAAGGG AAACCGACCC GTTTAGTCTG CCGCTCCCAA CAGCCCGAGT TTCTTGCCCC TTGAGTAAGG GGCAAGAAAC CTAAGCCGCC TGGTGGATGG CTCGGCTCGG GGCGCTGACG AAGGGCG
J23SpTT_59                    GGT TTGATGGGGC GGGGATGTAA GCCGGGAAGGG AAACCGACCC GTTTAGTCTG CCGCTCCCAA CAGCCCGAGT TTCTTGCCCC TTGAGTAAGG GGCAAGAAAC CTAAGCCGCC TGGTGGATGG CTCGGCTCGG GGCGCTGACG AAGGGC
J23SpTT_84                    GGT TTGATGGGGC GGGGATGTAA GCCGGGAAGGG AAACCGACCC GTTTAGTCTG CCGCTCCCAA CAGCCCGAGT TTCTTGCCCC TTGAGTAAGG GGCAAGAAAC CTAAGCCGCC TGGTGGATGG CTCGGCTCGG GGCGCTGACG AAGGG
J23SpTT_12          CA GAAGACGGGT TTGATGGGGC GGGGATGTAA GCCGGGAAGGG AAACCGACCC GTTTAGTCTG CCGCTCCCAA CAGCCCGAGT TTCTTGCCCC TTGAGTAAGG GGCAAGAAAC CTAAGCCGCC TGGTGGATGG CTCGGCTCGG GGCGCTG
J23SpTT_26                                                 CCGGGAAGGG AAACCGACCC GTTTAGTCTG CCGCTCCCAA CAGCCCGAGT TTCTTGCCCC TTGAGTAAGG GGCAAGAAAC CTAAGCCGCC TGGTGGATGG CTCGGCTCGG GGCGCTGACG AAGGGCGTGG CAAGCTGCGA TAAGCCCCGG TGAG
J23SpTT_20                                C GGGGATGTAA GCCGGGAAGGG AAACCGACCC GTTTAGTCTG CCGCTCCCAA CAGCCCGAGT TTCTTGCCCC TTGAGTAAGG GGCAAGAAAC CTAAGCCGCC TGGTGGATGG CTCGGCTCGG GGCGCTGACG AAGGGCGTGG CAAGCTGCGA TAAGCCCCGG
J23SpTT_12                                C GGGGATGTAA GCCGGGAAGGG AAACCGACCC GTTTAGTCTG CCGCTCCCAA CAGCCCGAGT TTCTTGCCCC TTGAGTAAGG GGCAAGAAAC CTAAGCCGCC TGGTGGATGG CTCGGCTCGG GGCGCTGACG AAGGGCGTGG CAAGCTGCGA TAAGCCCCG
J23SpTT_9                                         GTAA GCCGGGAAGGG AAACCGACCC GTTTAGTCTG CCGCTCCCAA CAGCCCGAGT TTCTTGCCCC TTGAGTAAGG GGCAAGAAAC CTAAGCCGCC TGGTGGATGG CTCGGCTCGG GGCGCTGACG AAGGGCGTGG CAAGCTGCGA TAAGCCCCGG TGAG
J23SpTT_1055                CGGGT TTGATGGGGC GGGGATGTAA GCCGGGAAGGG AAACCGACCC GTTTAGTCTG CCGCTCCCAA CAGCCCGAGT TTCTTGCCCC TTGAGTAAGG GGCAAGAAAC CTAAGCCGCC TGGTGGATGG CTCGGCTCGG GGCGCTGACG AAG
J23SpTT_941                 CGGGT TTGATGGGGC GGGGATGTAA GCCGGGAAGGG AAACCGACCC GTTTAGTCTG CCGCTCCCAA CAGCCCGAGT TTCTTGCCCC TTGAGTAAGG GGCAAGAAAC CTAAGCCGCC TGGTGGATGG CTCGGCTCGG GGCGCTGACG AAGG
J23SpTT_10                  CGGGT TTGATGGGGC GGGGATGTAA GCCGGGAAGGG AAACCGACCC GTTTAGTCTG CCGCTCCCAA CAGCCCGAGT TTCTTGCCCC TTGAGTAAGG GGCAAGAAAC CTAAGCCGCC TGGTGGATGG CTCGGCTCGG GGCGCTGACA AAGG
J23SpTT_24                  CGGGT TTGATGGGGC GGGGATGTAA GCCGGGAAGGG AAACCGACCC GTTTAGTCTG CCGCTCCCAA CAGCCCGAGT TTCTTGCCCC TTGAGTAAGG GGCAAGAAAC CTAAGCCGCC TGGTGGATGG CTCGGCTCGG GGCGCTGACG AAGAG
J23SpTT_263                ACGGGT TTGATGGGGC GGGGATGTAA GCCGGGAAGGG AAACCGACCC GTTTAGTCTG CCGCTCCCAA CAGCCCGAGT TTCTTGCCCC TTGAGTAAGG GGCAAGAAAC CTAAGCCGCC TGGTGGATGG CTCGGCTCGG GGCGCTGACG AAG
J23SpTT_130                  GGGT TTGATGGGGC GGGGATGTAA GCCGGGAAGGG AAACCGACCC GTTTAGTCTG CCGCTCCCAA CAGCCCGAGT TTCTTGCCCC TTGAGTAAGG GGCAAGAAAC CTAAGCCGCC TGGTGGATGG CTCGGCTCGG GGCGCTGACG AAGGG
J23SpTT_10                  CGGGT TTGATGGGGC GGGGATGTAA GCCGGGAAGGG AAACCGACCC GTTTAGTCTG CCGCTCCCAA CAGCCCGAGT TTCTTGCCCC TTGAGTAAGG GGCAAGAAAC CTAAGCCGCC TGGTGGATGG CTCGGCTCGG GGCGCTGACG AAAG
J23SpTT_73                   GGGT TTGATGGGGC GGGGATGTAA GCCGGGAAGGG AAACCGACCC GTTTAGTCTG CCGCTCCCAA CAGCCCGAGT TTCTTGCCCC TTGAGTAAGG GGCAAGAAAC CTAAGCCGCC TGGTGGATGG CTCGGCTCGG GGCGCTGACG AAGG
J23SpTT_32                GACGGGT TTGATGGGGC GGGGATGTAA GCCGGGAAGGG AAACCGACCC GTTTAGTCTG CCGCTCCCAA CAGCCCGAGT TTCTTGCCCC TTGAGTAAGG GGCAAGAAAC CTAAGCCGCC TGGTGGATGG CTCGGCTCGG GGCGCTGACG AA
J23SpTT_13                 ACGGGT TTGATGGGGC GGGGATGTAA GCCGGGAAGGG AAACCGACCC GTTTAGTCTG CCGCTCCCAA CAGCCCGAGT TTCTTGCCCC TTGAGTAAGG GGCAAGAAAC CTAAGCCGCC TGGTGGATGG CTCGGCTCGG GGCGCTGACG AA
J23SpTT_12                GACGGGT TTGATGGGGC GGGGATGTAA GCCGGGAAGGG AAACCGACCC GTTTAGTCTG CCGCTCCCAA CAGCCCGAGT TTCTTGCCCC TTGAGTAAGG GGCAAGAAAC CTAAGCCGCC TGGTGGATGG CTCGGCTCGG GGCGCTGACA AAG
J23SpTT_13                  CGGGT TTGATGGGGC GGGGATGTAA GCCGGGAAGGG AAACCGACCC GTTTAGTCTG CCGCTCCCAA CAGCCCGAGT TTCTTGCCCC TTGAGTAAGG GGCAAGAAAC CTAAGCCGCC TGGTGGATGG CTCGGCTCGG GGCGCTGACA AAG
J23SpTT_99                   GGGT TTGATGGGGC GGGGATGTAA GCCGGGAAGGG AAACCGACCC GTTTAGTCTG CCGCTCCCAA CAGCCCGAGT TTCTTGCCCC TTGAGTAAGG GGCAAGAAAC CTAAGCCGCC TGGTGGATGG CTCGGCTCGG GGCGCTGACG AAG
J23SpTT_43                  CGGGT TTGATGGGGC GGGGATGTAA GCCGGGAAGGG AAACCGACCC GTTTAGTCTG CCGCTCCCAA CAGCCCGAGT TTCTTGCCCC TTGAGTAAGG GGCAAGAAAC CTAAGCCGCC TGGTGGATGG CTCGGCTCGG GGCGCTGACG AA
J23SpTT_64                    GGT TTGATGGGGC GGGGATGTAA GCCGGGAAGGG AAACCGACCC GTTTAGTCTG CCGCTCCCAA CAGCCCGAGT TTCTTGCCCC TTGAGTAAGG GGCAAGAAAC CTAAGCCGCC TGGTGGATGG CTCGGCTCGG GGCGCTGACG AAGG
J23SpTT_59                    GGT TTGATGGGGC GGGGATGTAA GCCGGGAAGGG AAACCGACCC GTTTAGTCTG CCGCTCCCAA CAGCCCGAGT TTCTTGCCCC TTGAGTAAGG GGCAAGAAAC CTAAGCCGCC TGGTGGATGG CTCGGCTCGG GGCGCTGACG AAG
J23SpTT_30                GACGGGT TTGATGGGGC GGGGATGTAA GCCGGGAAGGG AAACCGACCC GTTTAGTCTG CCGCTCCCAA CAGCCCGAGT TTCTTGCCCC TTGAGTAAGG GGCAAGAAAC CTAAGCCGCC TGGTGGATGG CTCGGCTCGG GGCGCTGACG
J23SpTT_9                AGACGGGT TTGATGGGGC GGGGATGTAA GCCGGGAAGGG AAACCGACCC GTTTAGTCTG CCGCTCCCAA CAGCCCGAGT TTCTTGCCCC TTGAGTAAGG GGCAAGAAAC CTAAGCCGCC TGGTGGATGG CTCGGCTCGG GGCGCTGACG
J23SpTT_25                GACGGGT TTGATGGGGC GGGGATGTAA GCCGGGAAGGG AAACCGACCC GTTTAGTCTG CCGCTCCCAA CAGCCCGAGT TTCTTGCCCC TTGAGTAAGG GGCAAGAAAC CTAAGCCGCC TGGTGGATGG CTCGGCTCGG GGCGCTGAC
J23SpTT_48                                C GGGGATGTAA GCCGGGAAGGG AAACCGACCC GTTTAGTCTG CCGCTCCCAA CAGCCCGAGT TTCTTGCCCC TTGAGTAAGG GGCAAGAAAC CTAAGCCGCC TGGTGGATGG CTCGGCTCGG GGCGCTGACG AAGGGCGTGG CAAGCTGCG
J23SpTT_28                                C GGGGATGTAA GCCGGGAAGGG AAACCGACCC GTTTAGTCTG CCGCTCCCAA CAGCCCGAGT TTCTTGCCCC TTGAGTAAGG GGCAAGAAAC CTAAGCCGCC TGGTGGATGG CTCGGCTCGG GGCGCTGACG AAGGGCGTGG CAAGCTGCGA T
J23SpTT_10                                C GGGGATGTAA GCCGGGAAGGG AAACCGACCC GTTTAGTCTG CCGCTCCCAA CAGCCCGAGT TTCTTGCCCC TTGAGTAAGG GGCAAGAAAC CTAAGCCGCC TGGTGGATGG CTCGGCTCGG GGCGCTGACG AAGGGCGTGG CAAGCTGC
J23SpTT_16                                C GGGGATGTAA GCCGGGAAGGG AAACCGACCC GTTTAGTCTG CCGCTCCCAA CAGCCCGAGT TTCTTGCCCC TTGAGTAAGG GGCAAGAAAC CTAAGCCGCC TGGTGGATGG CTCGGCTCGG GGCGCTGACG AAGGGCGTGG CAAGCTG
J23SpTT_408                    GT TTGATGGGGC GGGGATGTAA GCCGGGAAGGG AAACCGACCC GTTTAGTCTG CCGCTCCCAA CAGCCCGAGT TTCTTGCCCC TTGAGTAAGG GGCAAGAAAC CTAAGCCGCC TGGTGGATGG CTCGGCTCGG GGCGCTGACG AAGGG
J23SpTT_10                     GT TTGATGGGGC GGGGATGTAA GCCGGGAAGGG AAACCGACCC GTTTAGTCTG CCGCTCCCAA CAGCCCGAGT TTCTTGCCCC TTGAGTAAGG GGCAAGAAAC CTAAGCCGCC TGGTGGATGG CTCGGCTCGG GGCGCTGACG AAGGGG
J23SpTT_201                    GT TTGATGGGGC GGGGATGTAA GCCGGGAAGGG AAACCGACCC GTTTAGTCTG CCGCTCCCAA CAGCCCGAGT TTCTTGCCCC TTGAGTAAGG GGCAAGAAAC CTAAGCCGCC TGGTGGATGG CTCGGCTCGG GGCGCTGACG AAGG
J23SpTT_295                    GT TTGATGGGGC GGGGATGTAA GCCGGGAAGGG AAACCGACCC GTTTAGTCTG CCGCTCCCAA CAGCCCGAGT TTCTTGCCCC TTGAGTAAGG GGCAAGAAAC CTAAGCCGCC TGGTGGATGG CTCGGCTCGG GGCGCTGACG AAG
J23SpTT_280       CGCA GAAGACGGGT TTGATGGGGC GGGGATGTAA GCCGGGAAGGG AAACCGACCC GTTTAGTCTG CCGCTCCCAA CAGCCCGAGT TTCTTGCCCC TTGAGTAAGG GGCAAGAAAC CTAAGCCGCC TGGTGGATGG CTCGGCTCGG G
J23SpTT_140      CCGCA GAAGACGGGT TTGATGGGGC GGGGATGTAA GCCGGGAAGGG AAACCGACCC GTTTAGTCTG CCGCTCCCAA CAGCCCGAGT TTCTTGCCCC TTGAGTAAGG GGCAAGAAAC CTAAGCCGCC TGGTGGATGG CTCGGCTCGG G
J23SpTT_109     CCCGCA GAAGACGGGT TTGATGGGGC GGGGATGTAA GCCGGGAAGGG AAACCGACCC GTTTAGTCTG CCGCTCCCAA CAGCCCGAGT TTCTTGCCCC TTGAGTAAGG GGCAAGAAAC CTAAGCCGCC TGGTGGATGG CTCGGCTCGG G
J23SpTT_12      CCCGCA GAAGACGGGT TTGATGGGGC GGGGATGTAA GCCGGGAAGGG AAACCGACCC GTTTAGTCTG CCGCTCCCAA CAGCCCGAGT TTCTTGCCCC TTGAGTAAGG GGCAAGAAAC CTAAGCCGCC TGGTGGATGG CTCGGCTCGG GG
J23SpTT_12       CCGCA GAAGACGGGT TTGATGGGGC GGGGATGTAA GCCGGGAAGGG AAACCGACCC GTTTAGTCTG CCGCTCCCAA CAGCCCGAGT TTCTTGCCCC TTGAGTAAGG GGCAAGAAAC CTAAGCCGCC TGGTGGATGG CTCGGCTCGG GG
J23SpTT_40        CGCA GAAGACGGGT TTGATGGGGC GGGGATGTAA GCCGGGAAGGG AAACCGACCC GTTTAGTCTG CCGCTCCCAA CAGCCCGAGT TTCTTGCCCC TTGAGTAAGG GGCAAGAAAC CTAAGCCGCC TGGTGGATGG CTCGGCTCGG GG
J23SpTT_9         CGCA GAAGACGGGT TTGATGGGGC GGGGATGTAA GCCGGGAAGGG AAACCGACCC GTTTAGTCTG CCGCTCCCAA CAGCCCGAGT TTCTTGCCCC TTGAGTAAGG GGCAAGAAAC CTAAGCCGCC TGGTGGATGG CTCGGCTCGG GGCG
J23SpTT_15      CCCGCA GAAGACGGGT TTGATGGGGC GGGGATGTAA GCCGGGAAGGG AAACCGACCC GTTTAGTCTG CCGCTCCCAA CAGCCCGAGT TTCTTGCCCC TTGAGTAAGG GGCAAGAAAC CTAAGCCGCC TGGTGGATGG CTCGGCTCGG
J23SpTT_16       CCGCA GAAGACGGGT TTGATGGGGC GGGGATGTAA GCCGGGAAGGG AAACCGACCC GTTTAGTCTG CCGCTCCCAA CAGCCCGAGT TTCTTGCCCC TTGAGTAAGG GGCAAGAAAC CTAAGCCGCC TGGTGGATGG CTCGGCTCGG
J23SpTT_15      CCCGCA GAAGACGGGT TTGATGGGGC GGGGATGTAA GCCGGGAAGGG AAACCGACCC GTTTAGTCTG CCGCTCCCAA CAGCCCGAGT TTCTTGCCCC TTGAGTAAGG GGCAAGAAAC CTAAGCCGCC TGGTGGATGG CTCGGCTCG
J23SpTT_10          CA GAAGACGGGT TTGATGGGGC GGGGATGTAA GCCGGGAAGGG AAACCGACCC GTTTAGTCTG CCGCTCCCAA CAGCCCGAGT TTCTTGCCCC TTGAGTAAGG GGCAAGAAAC CTAAGCCGCC TGGTGGATGG CTCGGCTCGG GG
J23SpTT_20       CCGCA GAAGACGGGT TTGATGGGGC GGGGATGTAA GCCGGGAAGGG AAACCGACCC GTTTAGTCTG CCGCTCCCAA CAGCCCGAGT TTCTTGCCCC TTGAGTAAGG GGCAAGAAAC CTAAGCCGCC TGGTGGATGG CTCGGCTCG
J23SpTT_35        CGCA GAAGACGGGT TTGATGGGGC GGGGATGTAA GCCGGGAAGGG AAACCGACCC GTTTAGTCTG CCGCTCCCAA CAGCCCGAGT TTCTTGCCCC TTGAGTAAGG GGCAAGAAAC CTAAGCCGCC TGGTGGATGG CTCGGCTCGG
J23SpTT_33         GCA GAAGACGGGT TTGATGGGGC GGGGATGTAA GCCGGGAAGGG AAACCGACCC GTTTAGTCTG CCGCTCCCAA CAGCCCGAGT TTCTTGCCCC TTGAGTAAGG GGCAAGAAAC CTAAGCCGCC TGGTGGATGG CTCGGCTCGG G
J23SpTT_47                  CGGGT TTGATGGGGC GGGGATGTAA GCCGGGAAGGG AAACCGACCC GTTTAGTCTG CCGCTCCCAA CAGCCCGAGT TTCTTGCCCC TTGAGTAAGG GGCAAGAAAC CTAAGCCGCC TGGTGGATGG CTCGGCTCGG GGCGCTGACG
J23SpTT_21                 ACGGGT TTGATGGGGC GGGGATGTAA GCCGGGAAGGG AAACCGACCC GTTTAGTCTG CCGCTCCCAA CAGCCCGAGT TTCTTGCCCC TTGAGTAAGG GGCAAGAAAC CTAAGCCGCC TGGTGGATGG CTCGGCTCGG GGCGCTGACG
J23SpTT_24                  CGGGT TTGATGGGGC GGGGATGTAA GCCGGGAAGGG AAACCGACCC GTTTAGTCTG CCGCTCCCAA CAGCCCGAGT TTCTTGCCCC TTGAGTAAGG GGCAAGAAAC CTAAGCCGCC TGGTGGATGG CTCGGCTCGG GGCGCTGAC
J23SpTT_21                              GGC GGGGATGTAA GCCGGGAAGGG AAACCGACCC GTTTAGTCTG CCGCTCCCAA CAGCCCGAGT TTCTTGCCCC TTGAGTAAGG GGCAAGAAAC CTAAGCCGCC TGGTGGATGG CTCGGCTCGG GGCGCTGACG AAGGGCGTGG
J23SpTT_11                            GGGGC GGGGATGTAA GCCGGGAAGGG AAACCGACCC GTTTAGTCTG CCGCTCCCAA CAGCCCGAGT TTCTTGCCCC TTGAGTAAGG GGCAAGAAAC CTAAGCCGCC TGGTGGATGG CTCGGCTCGG GGCGCTGACG AAGGGCGTGG
J23SpTT_9                             GGGGC GGGGATGTAA GCCGGGAAGGG AAACCGACCC GTTTAGTCTG CCGCTCCCAA CAGCCCGAGT TTCTTGCCCC TTGAGTAAGG GGCAAGAAAC CTAAGCCGCC TGGTGGATGG CTCGGCTCGG GGCGCTGACG AAGGGCGTG
J23SpTT_8                                GC GGGGATGTAA GCCGGGAAGGG AAACCGACCC GTTTAGTCTG CCGCTCCCAA CAGCCCGAGT TTCTTGCCCC TTGAGTAAGG GGCAAGAAAC CTAAGCCGCC TGGTGGATGG CTCGGCTCGG GGCGCTGACG AAGGGCGTGG C
J23SpTT_11                               GC GGGGATGTAA GCCGGGAAGGG AAACCGACCC GTTTAGTCTG CCGCTCCCAA CAGCCCGAGT TTCTTGCCCC TTGAGTAAGG GGCAAGAAAC CTAAGCCGCC TGGTGGATGG CTCGGCTCGG GGCGCTGACG AAGGGCGTGG
J23SpTT_19                         GATGGGGC GGGGATGTAA GCCGGGAAGGG AAACCGACCC GTTTAGTCTG CCGCTCCCAA CAGCCCGAGT TTCTTGCCCC TTGAGTAAGG GGCAAGAAAC CTAAGCCGCC TGGTGGATGG CTCGGCTCGG GGCGCTGACG AAGG
J23SpTT_12                        TGATGGGGC GGGGATGTAA GCCGGGAAGGG AAACCGACCC GTTTAGTCTG CCGCTCCCAA CAGCCCGAGT TTCTTGCCCC TTGAGTAAGG GGCAAGAAAC CTAAGCCGCC TGGTGGATGG CTCGGCTCGG GGCGCTGACG AAGGG
J23SpTT_9                         TGATGGGGC GGGGATGTAA GCCGGGAAGGG AAACCGACCC GTTTAGTCTG CCGCTCCCAA CAGCCCGAGT TTCTTGCCCC TTGAGTAAGG GGCAAGAAAC CTAAGCCGCC TGGTGGATGG CTCGGCTCGG GGCGCTGACG AAGG
J23SpTT_9                          GATGGGGC GGGGATGTAA GCCGGGAAGGG AAACCGACCC GTTTAGTCTG CCGCTCCCAA CAGCCCGAGT TTCTTGCCCC TTGAGTAAGG GGCAAGAAAC CTAAGCCGCC TGGTGGATGG CTCGGCTCGG GGCGCTGACG AAGGGC
J23SpTT_12                         GATGGGGC GGGGATGTAA GCCGGGAAGGG AAACCGACCC GTTTAGTCTG CCGCTCCCAA CAGCCCGAGT TTCTTGCCCC TTGAGTAAGG GGCAAGAAAC CTAAGCCGCC TGGTGGATGG CTCGGCTCGG GGCGCTGACG AAGGG
J23SpTT_12                        TGATGGGGC GGGGATGTAA GCCGGGAAGGG AAACCGACCC GTTTAGTCTG CCGCTCCCAA CAGCCCGAGT TTCTTGCCCC TTGAGTAAGG GGCAAGAAAC CTAAGCCGCC TGGTGGATGG CTCGGCTCGG GGCGCTGACG AAG
J23SpTT_25                            GGGGC GGGGATGTAA GCCGGGAAGGG AAACCGACCC GTTTAGTCTG CCGCTCCCAA CAGCCCGAGT TTCTTGCCCC TTGAGTAAGG GGCAAGAAAC CTAAGCCGCC TGGTGGATGG CTCGGCTCGG GGCGCTGACG AAGGGC
J23SpTT_17                            GGGGC GGGGATGTAA GCCGGGAAGGG AAACCGACCC GTTTAGTCTG CCGCTCCCAA CAGCCCGAGT TTCTTGCCCC TTGAGTAAGG GGCAAGAAAC CTAAGCCGCC TGGTGGATGG CTCGGCTCGG GGCGCTGACG AAGGGCG
J23SpTT_15                           TGGGGC GGGGATGTAA GCCGGGAAGGG AAACCGACCC GTTTAGTCTG CCGCTCCCAA CAGCCCGAGT TTCTTGCCCC TTGAGTAAGG GGCAAGAAAC CTAAGCCGCC TGGTGGATGG CTCGGCTCGG GGCGCTGACG AAGGGC
J23SpTT_113         CA GAAGACGGGT TTGATGGGGC GGGGATGTAA GCCGGGAAGGG AAACCGACCC GTTTAGTCTG CCGCTCCCAA CAGCCCGAGT TTCTTGCCCC TTGAGTAAGG GGCAAGAAAC CTAAGCCGCC TGGTGGATGG CTCGGCTCGG G
J23SpTT_38           A GAAGACGGGT TTGATGGGGC GGGGATGTAA GCCGGGAAGGG AAACCGACCC GTTTAGTCTG CCGCTCCCAA CAGCCCGAGT TTCTTGCCCC TTGAGTAAGG GGCAAGAAAC CTAAGCCGCC TGGTGGATGG CTCGGCTCGG G
J23SpTT_12          CA GAAGACGGGT TTGATGGGGC GGGGATGTAA GCCGGGAAGGG AAACCGACCC GTTTAGTCTG CCGCTCCCAA CAGCCCGAGT TTCTTGCCCC TTGAGTAAGG GGCAAGAAAC CTAAGCCGCC TGGTGGATGG CTCGGCTCGG
J23SpTT_18          CA GAAGACGGGT TTGATGGGGC GGGGATGTAA GCCGGGAAGGG AAACCGACCC GTTTAGTCTG CCGCTCCCAA CAGCCCGAGT TTCTTGCCCC TTGAGTAAGG GGCAAGAAAC CTAAGCCGCC TGGTGGATGG CTCGGCTCG
J23SpTT_10           A GAAGACGGGT TTGATGGGGC GGGGATGTAA GCCGGGAAGGG AAACCGACCC GTTTAGTCTG CCGCTCCCAA CAGCCCGAGT TTCTTGCCCC TTGAGTAAGG GGCAAGAAAC CTAAGCCGCC TGGTGGATGG CTCGGCTCGG
J23SpTT_54        CGCA GAAGACGGGT TTGATGGGGC GGGGATGTAA GCCGGGAAGGG AAACCGACCC GTTTAGTCTG CCGCTCCCAA CAGCCCGAGT TTCTTGCCCC TTGAGTAAGG GGCAAGAAAC CTAAGCCGCC TGGTGGATGG CTCGGCTCG
J23SpTT_15                     GT TTGATGGGGC GGGGATGTAA GCCGGGAAGGG AAACCGACCC GTTTAGTCTG CCGCTCCCAA CAGCCCGAGT TTCTTGCCCC TTGAGTAAGG GGCAAGAAAC CTAAGCCGCC TGGTGGATGG CTCGGCTCGG GGCGCTGACG
J23SpTT_52                            GGGGC GGGGATGTAA GCCGGGAAGGG AAACCGACCC GTTTAGTCTG CCGCTCCCAA CAGCCCGAGT TTCTTGCCCC TTGAGTAAGG GGCAAGAAAC CTAAGCCGCC TGGTGGATGG CTCGGCTCGG GGCGCTGACG AAGGG
J23SpTT_19                           TGGGGC GGGGATGTAA GCCGGGAAGGG AAACCGACCC GTTTAGTCTG CCGCTCCCAA CAGCCCGAGT TTCTTGCCCC TTGAGTAAGG GGCAAGAAAC CTAAGCCGCC TGGTGGATGG CTCGGCTCGG GGCGCTGACG AAGGG
J23SpTT_8                            TGGGGC GGGGATGTAA GCCGGGAAGGG AAACCGACCC GTTTAGTCTG CCGCTCCCAA CAGCCCGAGT TTCTTGCCCC TTGAGTAAGG GGCAAGAAAC CTAAGCCGCC TGGTGGATGG CTCGGCTCGG GGCGCTGACG AAGG
J23SpTT_206                               C GGGGATGTAA GCCGGGAAGGG AAACCGACCC GTTTAGTCTG CCGCTCCCAA CAGCCCGAGT TTCTTGCCCC TTGAGTAAGG GGCAAGAAAC CTAAGCCGCC TGGTGGATGG CTCGGCTCGG GGCGCTGACG AAGGGCGTGG
J23SpTT_54                                C GGGGATGTAA GCCGGGAAGGG AAACCGACCC GTTTAGTCTG CCGCTCCCAA CAGCCCGAGT TTCTTGCCCC TTGAGTAAGG GGCAAGAAAC CTAAGCCGCC TGGTGGATGG CTCGGCTCGG GGCGCTGACG AAGGGCGTGG C
J23SpTT_16                                C GGGGATGTAA GCCGGGAAGGG AAACCGACCC GTTTAGTCTG CCGCTCCCAA CAGCCCGAGT TTCTTGCCCC TTGAGTAAGG GGCAAGAAAC CTAAGCCGCC TGGTGGATGG CTCGGCTCGG GGCGCTGACG AAGGGCGTGG CAAG
J23SpTT_8                                   GGGGATGTAA GCCGGGAAGGG AAACCGACCC GTTTAGTCTG CCGCTCCCAA CAGCCCGAGT TTCTTGCCCC TTGAGTAAGG GGCAAGAAAC CTAAGCCGCC TGGTGGATGG CTCGGCTCGG GGCGCTGACG AAGGGCGTGG C
J23SpTT_142                               C GGGGATGTAA GCCGGGAAGGG AAACCGACCC GTTTAGTCTG CCGCTCCCAA CAGCCCGAGT TTCTTGCCCC TTGAGTAAGG GGCAAGAAAC CTAAGCCGCC TGGTGGATGG CTCGGCTCGG GGCGCTGACG AAGGGCGTG
J23SpTT_9                                    GGGATGTAA GCCGGGAAGGG AAACCGACCC GTTTAGTCTG CCGCTCCCAA CAGCCCGAGT TTCTTGCCCC TTGAGTAAGG GGCAAGAAAC CTAAGCCGCC TGGTGGATGG CTCGGCTCGG GGCGCTGACG AAGGGCGTGG C
J23SpTT_8                                GC GGGGATGTAA GCCGGGAAGGG AAACCGACCC GTTTAGTCTG CCGCTCCCAA CAGCCCGAGT TTCTTGCCCC TTGAGTAAGG GGCAAGAAAC CTAAGCCGCC TGGTGGATGG CTCGGCTCGG GGCGCTGACG AAGGGCGTG
J23SpTT_18                                  GGGGATGTAA GCCGGGAAGGG AAACCGACCC GTTTAGTCTG CCGCTCCCAA CAGCCCGAGT TTCTTGCCCC TTGAGTAAGG GGCAAGAAAC CTAAGCCGCC TGGTGGATGG CTCGGCTCGG GGCGCTGACG AAGGGCGTGG
J23SpTT_34                              GGC GGGGATGTAA GCCGGGAAGGG AAACCGACCC GTTTAGTCTG CCGCTCCCAA CAGCCCGAGT TTCTTGCCCC TTGAGTAAGG GGCAAGAAAC CTAAGCCGCC TGGTGGATGG CTCGGCTCGG GGCGCTGACG AAGGGC
J23SpTT_16                              GGC GGGGATGTAA GCCGGGAAGGG AAACCGACCC GTTTAGTCTG CCGCTCCCAA CAGCCCGAGT TTCTTGCCCC TTGAGTAAGG GGCAAGAAAC CTAAGCCGCC TGGTGGATGG CTCGGCTCGG GGCGCTGACG AAGGGCG
J23SpTT_16                               GC GGGGATGTAA GCCGGGAAGGG AAACCGACCC GTTTAGTCTG CCGCTCCCAA CAGCCCGAGT TTCTTGCCCC TTGAGTAAGG GGCAAGAAAC CTAAGCCGCC TGGTGGATGG CTCGGCTCGG GGCGCTGACG AAGGGCG
J23SpTT_20                                   GGGATGTAA GCCGGGAAGGG AAACCGACCC GTTTAGTCTG CCGCTCCCAA CAGCCCGAGT TTCTTGCCCC TTGAGTAAGG GGCAAGAAAC CTAAGCCGCC TGGTGGATGG CTCGGCTCGG GGCGCTGACG AAGGGCGTGG
```

```
J23SpTT_9                                          GGATGTAA GCCGGGAAGG AAACCGACCC GTTTAGTCTG CCGCTCCCAA CAGCCCGAGT TTCTTGCCCC TTGAGTAAGG GGCAAGAAAC CTAAGCCGCC TGGTGGATGG CTCGGCTCGG GGCGCTGACG AAGGGCGTGG
J23SpTT_16                                       GGGGATGTAA GCCGGGAAGG AAACCGACCC GTTTAGTCTG CCGCTCCCAA CAGCCCGAGT TTCTTGCCCC TTGAGTAAGG GGCAAGAAAC CTAAGCCGCC TGGTGGATGG CTCGGCTCGG GGCGCTGACG AAGGGCGTG
J23SpTT_8                                         GGGATGTAA GCCGGGAAGG AAACCGACCC GTTTAGTCTG CCGCTCCCAA CAGCCCGAGT TTCTTGCCCC TTGAGTAAGG GGCAAGAAAC CTAAGCCGCC TGGTGGATGG CTCGGCTCGG GGCGCTGACG AAGGGCGTG
J23SpTT_34            CGCA GAAGACGGGT TTGATGGGGC GGGGATGTAA GCCGGGAAGG AAACCGACCC GTTTAGTCTG CCGCTCCCAA CAGCCCGAGT TTCTTGCCCC TTGAGTAAGG GGCAAGAAAC CTAAGCCGCC TGGTGGATGG CTCGG
J23SpTT_31            CGCA GAAGACGGGT TTGATGGGGC GGGGATGTAA GCCGGGAAGG AAACCGACCC GTTTAGTCTG CCGCTCCCAA CAGCCCGAGT TTCTTGCCCC TTGAGTAAGG GGCAAGAAAC CTAAGCCGCC TGGTGGATGG CTCGGC
J23SpTT_22            CGCA GAAGACGGGT TTGATGGGGC GGGGATGTAA GCCGGGAAGG AAACCGACCC GTTTAGTCTG CCGCTCCCAA CAGCCCGAGT TTCTTGCCCC TTGAGTAAGG GGCAAGAAAC CTAAGCCGCC TGGTGGATGG CTCGGCT
J23SpTT_15           CCGCA GAAGACGGGT TTGATGGGGC GGGGATGTAA GCCGGGAAGG AAACCGACCC GTTTAGTCTG CCGCTCCCAA CAGCCCGAGT TTCTTGCCCC TTGAGTAAGG GGCAAGAAAC CTAAGCCGCC TGGTGGATGG CTCGGCT
J23SpTT_11           CCGCA GAAGACGGGT TTGATGGGGC GGGGATGTAA GCCGGGAAGG AAACCGACCC GTTTAGTCTG CCGCTCCCAA CAGCCCGAGT TTCTTGCCCC TTGAGTAAGG GGCAAGAAAC CTAAGCCGCC TGGTGGATGG CTCGGC
J23SpTT_13          CCCGCA GAAGACGGGT TTGATGGGGC GGGGATGTAA GCCGGGAAGG AAACCGACCC GTTTAGTCTG CCGCTCCCAA CAGCCCGAGT TTCTTGCCCC TTGAGTAAGG GGCAAGAAAC CTAAGCCGCC TGGTGGATGG CTCGG
J23SpTT_11           CCGCA GAAGACGGGT TTGATGGGGC GGGGATGTAA GCCGGGAAGG AAACCGACCC GTTTAGTCTG CCGCTCCCAA CAGCCCGAGT TTCTTGCCCC TTGAGTAAGG GGCAAGAAAC CTAAGCCGCC TGGTGGATGG CTCGG
J23SpTT_15            CGCA GAAGACGGGT TTGATGGGGC GGGGATGTAA GCCGGGAAGG AAACCGACCC GTTTAGTCTG CCGCTCCCAA CAGCCCGAGT TTCTTGCCCC TTGAGTAAGG GGCAAGAAAC CTAAGCCGCC TGGTGGATGG CTCG
J23SpTT_386                                    C GGGGATGTAA GCCGGGAAGG AAACCGACCC GTTTAGTCTG CCGCTCCCAA CAGCCCGAGT TTCTTGCCCC TTGAGTAAGG GGCAAGAAAC CTAAGCCGCC TGGTGGATGG CTCGGCTCGG GGCGCTGACG AAGGGCG
J23SpTT_13                                     C GGGGATGTAA GCCGGGAAGG AAACCGACCC GTTTAGTCTG CCGCTCCCAA CAGCCCGAGT TTCTTGCCCC TTGAGTAAGG GGCAAGAAAC CTAAGCCGCC TGGTGGATGG CTCGGCTCGG GGCGCTGACG AAGGGCGT
J23SpTT_48                                   GGC GGGGATGTAA GCCGGGAAGG AAACCGACCC GTTTAGTCTG CCGCTCCCAA CAGCCCGAGT TTCTTGCCCC TTGAGTAAGG GGCAAGAAAC CTAAGCCGCC TGGTGGATGG CTCGGCTCGG GGCGCTGACG AAGGG
J23SpTT_21                                  GGGC GGGGATGTAA GCCGGGAAGG AAACCGACCC GTTTAGTCTG CCGCTCCCAA CAGCCCGAGT TTCTTGCCCC TTGAGTAAGG GGCAAGAAAC CTAAGCCGCC TGGTGGATGG CTCGGCTCGG GGCGCTGACG AAGGG
J23SpTT_17                                 GGGGC GGGGATGTAA GCCGGGAAGG AAACCGACCC GTTTAGTCTG CCGCTCCCAA CAGCCCGAGT TTCTTGCCCC TTGAGTAAGG GGCAAGAAAC CTAAGCCGCC TGGTGGATGG CTCGGCTCGG GGCGCTGACG AAGG
J23SpTT_24                                  GGGC GGGGATGTAA GCCGGGAAGG AAACCGACCC GTTTAGTCTG CCGCTCCCAA CAGCCCGAGT TTCTTGCCCC TTGAGTAAGG GGCAAGAAAC CTAAGCCGCC TGGTGGATGG CTCGGCTCGG GGCGCTGACG AAGG
J23SpTT_30                                 GGGGC GGGGATGTAA GCCGGGAAGG AAACCGACCC GTTTAGTCTG CCGCTCCCAA CAGCCCGAGT TTCTTGCCCC TTGAGTAAGG GGCAAGAAAC CTAAGCCGCC TGGTGGATGG CTCGGCTCGG GGCGCTGACG AAG
J23SpTT_12                                TGGGGC GGGGATGTAA GCCGGGAAGG AAACCGACCC GTTTAGTCTG CCGCTCCCAA CAGCCCGAGT TTCTTGCCCC TTGAGTAAGG GGCAAGAAAC CTAAGCCGCC TGGTGGATGG CTCGGCTCGG GGCGCTGACG AAG
J23SpTT_638                                    C GGGGATGTAA GCCGGGAAGG AAACCGACCC GTTTAGTCTG CCGCTCCCAA CAGCCCGAGT TTCTTGCCCC TTGAGTAAGG GGCAAGAAAC CTAAGCCGCC TGGTGGATGG CTCGGCTCGG GGCGCTGACG AAGGGC
J23SpTT_38                                    GC GGGGATGTAA GCCGGGAAGG AAACCGACCC GTTTAGTCTG CCGCTCCCAA CAGCCCGAGT TTCTTGCCCC TTGAGTAAGG GGCAAGAAAC CTAAGCCGCC TGGTGGATGG CTCGGCTCGG GGCGCTGACG AAGGGC
J23SpTT_49                                    GC GGGGATGTAA GCCGGGAAGG AAACCGACCC GTTTAGTCTG CCGCTCCCAA CAGCCCGAGT TTCTTGCCCC TTGAGTAAGG GGCAAGAAAC CTAAGCCGCC TGGTGGATGG CTCGGCTCGG GGCGCTGACG AAGGG
J23SpTT_47                    GACGGGT TTGATGGGGC GGGGATGTAA GCCGGGAAGG AAACCGACCC GTTTAGTCTG CCGCTCCCAA CAGCCCGAGT TTCTTGCCCC TTGAGTAAGG GGCAAGAAAC CTAAGCCGCC TGGTGGATGG CTCGGCTCGG G
J23SpTT_20                   AGACGGGT TTGATGGGGC GGGGATGTAA GCCGGGAAGG AAACCGACCC GTTTAGTCTG CCGCTCCCAA CAGCCCGAGT TTCTTGCCCC TTGAGTAAGG GGCAAGAAAC CTAAGCCGCC TGGTGGATGG CTCGGCTCGG G
J23SpTT_11                  AAGACGGGT TTGATGGGGC GGGGATGTAA GCCGGGAAGG AAACCGACCC GTTTAGTCTG CCGCTCCCAA CAGCCCGAGT TTCTTGCCCC TTGAGTAAGG GGCAAGAAAC CTAAGCCGCC TGGTGGATGG CTCGGCTCGG G
J23SpTT_14                     ACGGGT TTGATGGGGC GGGGATGTAA GCCGGGAAGG AAACCGACCC GTTTAGTCTG CCGCTCCCAA CAGCCCGAGT TTCTTGCCCC TTGAGTAAGG GGCAAGAAAC CTAAGCCGCC TGGTGGATGG CTCGGCTCGG G
J23SpTT_33                                   GGC GGGGATGTAA GCCGGGAAGG AAACCGACCC GTTTAGTCTG CCGCTCCCAA CAGCCCGAGT TTCTTGCCCC TTGAGTAAGG GGCAAGAAAC CTAAGCCGCC TGGTGGATGG CTCGGCTCGG GGCGCTGACG AAG
J23SpTT_32                                   GGC GGGGATGTAA GCCGGGAAGG AAACCGACCC GTTTAGTCTG CCGCTCCCAA CAGCCCGAGT TTCTTGCCCC TTGAGTAAGG GGCAAGAAAC CTAAGCCGCC TGGTGGATGG CTCGGCTCGG GGCGCTGACG AAGG
J23SpTT_20                                    GC GGGGATGTAA GCCGGGAAGG AAACCGACCC GTTTAGTCTG CCGCTCCCAA CAGCCCGAGT TTCTTGCCCC TTGAGTAAGG GGCAAGAAAC CTAAGCCGCC TGGTGGATGG CTCGGCTCGG GGCGCTGACG AAGG
J23SpTT_13                                  GGGC GGGGATGTAA GCCGGGAAGG AAACCGACCC GTTTAGTCTG CCGCTCCCAA CAGCCCGAGT TTCTTGCCCC TTGAGTAAGG GGCAAGAAAC CTAAGCCGCC TGGTGGATGG CTCGGCTCGG GGCGCTGACG AAG
J23SpTT_76                                       GGGGATGTAA GCCGGGAAGG AAACCGACCC GTTTAGTCTG CCGCTCCCAA CAGCCCGAGT TTCTTGCCCC TTGAGTAAGG GGCAAGAAAC CTAAGCCGCC TGGTGGATGG CTCGGCTCGG GGCGCTGACG AAGGGC
J23SpTT_40                                       GGGGATGTAA GCCGGGAAGG AAACCGACCC GTTTAGTCTG CCGCTCCCAA CAGCCCGAGT TTCTTGCCCC TTGAGTAAGG GGCAAGAAAC CTAAGCCGCC TGGTGGATGG CTCGGCTCGG GGCGCTGACG AAGGGCG
J23SpTT_38                                        GGGATGTAA GCCGGGAAGG AAACCGACCC GTTTAGTCTG CCGCTCCCAA CAGCCCGAGT TTCTTGCCCC TTGAGTAAGG GGCAAGAAAC CTAAGCCGCC TGGTGGATGG CTCGGCTCGG GGCGCTGACG AAGGGCG
J23SpTT_1183                                   C GGGGATGTAA GCCGGGAAGG AAACCGACCC GTTTAGTCTG CCGCTCCCAA CAGCCCGAGT TTCTTGCCCC TTGAGTAAGG GGCAAGAAAC CTAAGCCGCC TGGTGGATGG CTCGGCTCGG GGCGCTGACG AAGGG
J23SpTT_34                                     C GGGGATGTAA GCCGGGAAGG AAACCGACCC GTTTAGTCTG CCGCTCCCAA CAGCCCGAGT TTCTTGCCCC TTGAGTAAGG GGCAAGAAAC CTAAGCCGCC TGGTGGATGG CTCGGCTCGG GGCGCTGACG AAGGGG
J23SpTT_700                                    C GGGGATGTAA GCCGGGAAGG AAACCGACCC GTTTAGTCTG CCGCTCCCAA CAGCCCGAGT TTCTTGCCCC TTGAGTAAGG GGCAAGAAAC CTAAGCCGCC TGGTGGATGG CTCGGCTCGG GGCGCTGACG AAGG
J23SpTT_9                                      C GGGGATGTAA GCCGGGAAGG AAACCGACCC GTTTAGTCTG CCGCTCCCAA CAGCCCGAGT TTCTTGCCCC TTGAGTAAGG GGCAAGAAAC CTAAGCCGCC TGGTGGATGG CTCGGCTCGG GGCGCTGACG AAGGAC
J23SpTT_10                                     C GGGGATGTAA GCCGGGAAGG AAACCGACCC GTTTAGTCTG CCGCTCCCAA CAGCCCGAGT TTCTTGCCCC TTGAGTAAGG GGCAAGAAAC CTAAGCCGCC TGGTGGATGG CTCGGCTCGG GGCGCTGACG AAGAG
J23SpTT_10                                     C GGGGATGTAA GCCGGGAAGG AAACCGACCC GTTTAGTCTG CCGCTCCCAA CAGCCCGAGT TTCTTGCCCC TTGAGTAAGG GGCAAGAAAC CTAAGCCGCC TGGTGGATGG CTCGGCTCGG GGCGCTGACG AAAGGC
J23SpTT_88                                       GGGGATGTAA GCCGGGAAGG AAACCGACCC GTTTAGTCTG CCGCTCCCAA CAGCCCGAGT TTCTTGCCCC TTGAGTAAGG GGCAAGAAAC CTAAGCCGCC TGGTGGATGG CTCGGCTCGG GGCGCTGACG AAGGG
J23SpTT_11              CA GAAGACGGGT TTGATGGGGC GGGGATGTAA GCCGGGAAGG AAACCGACCC GTTTAGTCTG CCGCTCCCAA CAGCCCGAGT TTCTTGCCCC TTGAGTAAGG GGCAAGAAAC CTAAGCCGCC TGGTGGATGG CTCGG
J23SpTT_793                                    C GGGGATGTAA GCCGGGAAGG AAACCGACCC GTTTAGTCTG CCGCTCCCAA CAGCCCGAGT TTCTTGCCCC TTGAGTAAGG GGCAAGAAAC CTAAGCCGCC TGGTGGATGG CTCGGCTCGG GGCGCTGACG AAG
J23SpTT_40                                    GC GGGGATGTAA GCCGGGAAGG AAACCGACCC GTTTAGTCTG CCGCTCCCAA CAGCCCGAGT TTCTTGCCCC TTGAGTAAGG GGCAAGAAAC CTAAGCCGCC TGGTGGATGG CTCGGCTCGG GGCGCTGACG AAG
J23SpTT_71                                        GGGATGTAA GCCGGGAAGG AAACCGACCC GTTTAGTCTG CCGCTCCCAA CAGCCCGAGT TTCTTGCCCC TTGAGTAAGG GGCAAGAAAC CTAAGCCGCC TGGTGGATGG CTCGGCTCGG GGCGCTGACG AAGGGC
J23SpTT_32                                         GGATGTAA GCCGGGAAGG AAACCGACCC GTTTAGTCTG CCGCTCCCAA CAGCCCGAGT TTCTTGCCCC TTGAGTAAGG GGCAAGAAAC CTAAGCCGCC TGGTGGATGG CTCGGCTCGG GGCGCTGACG AAGGGC
J23SpTT_10                                         GGATGTAA GCCGGGAAGG AAACCGACCC GTTTAGTCTG CCGCTCCCAA CAGCCCGAGT TTCTTGCCCC TTGAGTAAGG GGCAAGAAAC CTAAGCCGCC TGGTGGATGG CTCGGCTCGG GGCGCTGACG AAGGGCG
J23SpTT_65                      CGGGT TTGATGGGGC GGGGATGTAA GCCGGGAAGG AAACCGACCC GTTTAGTCTG CCGCTCCCAA CAGCCCGAGT TTCTTGCCCC TTGAGTAAGG GGCAAGAAAC CTAAGCCGCC TGGTGGATGG CTCGGCTCGG G
J23SpTT_15                      CGGGT TTGATGGGGC GGGGATGTAA GCCGGGAAGG AAACCGACCC GTTTAGTCTG CCGCTCCCAA CAGCCCGAGT TTCTTGCCCC TTGAGTAAGG GGCAAGAAAC CTAAGCCGCC TGGTGGATGG CTCGGCTCGG
J23SpTT_12                      CGGGT TTGATGGGGC GGGGATGTAA GCCGGGAAGG AAACCGACCC GTTTAGTCTG CCGCTCCCAA CAGCCCGAGT TTCTTGCCCC TTGAGTAAGG GGCAAGAAAC CTAAGCCGCC TGGTGGATGG CTCGGCTCG
J23SpTT_11                    GACGGGT TTGATGGGGC GGGGATGTAA GCCGGGAAGG AAACCGACCC GTTTAGTCTG CCGCTCCCAA CAGCCCGAGT TTCTTGCCCC TTGAGTAAGG GGCAAGAAAC CTAAGCCGCC TGGTGGATGG CTCGGCTCG
J23SpTT_15                                             GTAA GCCGGGAAGG AAACCGACCC GTTTAGTCTG CCGCTCCCAA CAGCCCGAGT TTCTTGCCCC TTGAGTAAGG GGCAAGAAAC CTAAGCCGCC TGGTGGATGG CTCGGCTCGG GGCGCTGACG AAGGGCGTGG
J23SpTT_15                                             GTAA GCCGGGAAGG AAACCGACCC GTTTAGTCTG CCGCTCCCAA CAGCCCGAGT TTCTTGCCCC TTGAGTAAGG GGCAAGAAAC CTAAGCCGCC TGGTGGATGG CTCGGCTCGG GGCGCTGACG AAGGGCGTG
J23SpTT_143                                       GGGATGTAA GCCGGGAAGG AAACCGACCC GTTTAGTCTG CCGCTCCCAA CAGCCCGAGT TTCTTGCCCC TTGAGTAAGG GGCAAGAAAC CTAAGCCGCC TGGTGGATGG CTCGGCTCGG GGCGCTGACG AAGGG
J23SpTT_84                                       GGGGATGTAA GCCGGGAAGG AAACCGACCC GTTTAGTCTG CCGCTCCCAA CAGCCCGAGT TTCTTGCCCC TTGAGTAAGG GGCAAGAAAC CTAAGCCGCC TGGTGGATGG CTCGGCTCGG GGCGCTGACG AAGG
J23SpTT_75                                       GGGGATGTAA GCCGGGAAGG AAACCGACCC GTTTAGTCTG CCGCTCCCAA CAGCCCGAGT TTCTTGCCCC TTGAGTAAGG GGCAAGAAAC CTAAGCCGCC TGGTGGATGG CTCGGCTCGG GGCGCTGACG AAG
J23SpTT_76                                        GGGATGTAA GCCGGGAAGG AAACCGACCC GTTTAGTCTG CCGCTCCCAA CAGCCCGAGT TTCTTGCCCC TTGAGTAAGG GGCAAGAAAC CTAAGCCGCC TGGTGGATGG CTCGGCTCGG GGCGCTGACG AAGG
J23SpTT_97            CGCA GAAGACGGGT TTGATGGGGC GGGGATGTAA GCCGGGAAGG AAACCGACCC GTTTAGTCTG CCGCTCCCAA CAGCCCGAGT TTCTTGCCCC TTGAGTAAGG GGCAAGAAAC CTAAGCCGCC TGGTGGATG
J23SpTT_60            CGCA GAAGACGGGT TTGATGGGGC GGGGATGTAA GCCGGGAAGG AAACCGACCC GTTTAGTCTG CCGCTCCCAA CAGCCCGAGT TTCTTGCCCC TTGAGTAAGG GGCAAGAAAC CTAAGCCGCC TGGTGGATGG
J23SpTT_59           CCGCA GAAGACGGGT TTGATGGGGC GGGGATGTAA GCCGGGAAGG AAACCGACCC GTTTAGTCTG CCGCTCCCAA CAGCCCGAGT TTCTTGCCCC TTGAGTAAGG GGCAAGAAAC CTAAGCCGCC TGGTGGATG
J23SpTT_44            CGCA GAAGACGGGT TTGATGGGGC GGGGATGTAA GCCGGGAAGG AAACCGACCC GTTTAGTCTG CCGCTCCCAA CAGCCCGAGT TTCTTGCCCC TTGAGTAAGG GGCAAGAAAC CTAAGCCGCC TGGTGGATGG C
J23SpTT_27          CCCGCA GAAGACGGGT TTGATGGGGC GGGGATGTAA GCCGGGAAGG AAACCGACCC GTTTAGTCTG CCGCTCCCAA CAGCCCGAGT TTCTTGCCCC TTGAGTAAGG GGCAAGAAAC CTAAGCCGCC TGGTGGATGG
J23SpTT_18          CCCGCA GAAGACGGGT TTGATGGGGC GGGGATGTAA GCCGGGAAGG AAACCGACCC GTTTAGTCTG CCGCTCCCAA CAGCCCGAGT TTCTTGCCCC TTGAGTAAGG GGCAAGAAAC CTAAGCCGCC TGGTGGATGG C
J23SpTT_19           CCGCA GAAGACGGGT TTGATGGGGC GGGGATGTAA GCCGGGAAGG AAACCGACCC GTTTAGTCTG CCGCTCCCAA CAGCCCGAGT TTCTTGCCCC TTGAGTAAGG GGCAAGAAAC CTAAGCCGCC TGGTGGATGG C
J23SpTT_20           CCGCA GAAGACGGGT TTGATGGGGC GGGGATGTAA GCCGGGAAGG AAACCGACCC GTTTAGTCTG CCGCTCCCAA CAGCCCGAGT TTCTTGCCCC TTGAGTAAGG GGCAAGAAAC CTAAGCCGCC TGGTGGATGG
J23SpTT_41          CCCGCA GAAGACGGGT TTGATGGGGC GGGGATGTAA GCCGGGAAGG AAACCGACCC GTTTAGTCTG CCGCTCCCAA CAGCCCGAGT TTCTTGCCCC TTGAGTAAGG GGCAAGAAAC CTAAGCCGCC TGGTGGATG
J23SpTT_40          CCCGCA GAAGACGGGT TTGATGGGGC GGGGATGTAA GCCGGGAAGG AAACCGACCC GTTTAGTCTG CCGCTCCCAA CAGCCCGAGT TTCTTGCCCC TTGAGTAAGG GGCAAGAAAC CTAAGCCGCC TGGTGGAT
J23SpTT_10             GCA GAAGACGGGT TTGATGGGGC GGGGATGTAA GCCGGGAAGG AAACCGACCC GTTTAGTCTG CCGCTCCCAA CAGCCCGAGT TTCTTGCCCC TTGAGTAAGG GGCAAGAAAC CTAAGCCGCC TGGTGGATGG C
J23SpTT_19              CA GAAGACGGGT TTGATGGGGC GGGGATGTAA GCCGGGAAGG AAACCGACCC GTTTAGTCTG CCGCTCCCAA CAGCCCGAGT TTCTTGCCCC TTGAGTAAGG GGCAAGAAAC CTAAGCCGCC TGGTGGATGG C
J23SpTT_48           CCGCA GAAGACGGGT TTGATGGGGC GGGGATGTAA GCCGGGAAGG AAACCGACCC GTTTAGTCTG CCGCTCCCAA CAGCCCGAGT TTCTTGCCCC TTGAGTAAGG GGCAAGAAAC CTAAGCCGCC TGGTGGAT
J23SpTT_10             GCA GAAGACGGGT TTGATGGGGC GGGGATGTAA GCCGGGAAGG AAACCGACCC GTTTAGTCTG CCGCTCCCAA CAGCCCGAGT TTCTTGCCCC TTGAGTAAGG GGCAAGAAAC CTAAGCCGCC TGGTGGATGG
J23SpTT_34                                     C GGGGATGTAA GCCGGGAAGG AAACCGACCC GTTTAGTCTG CCGCTCCCAA CAGCCCGAGT TTCTTGCCCC TTGAGTAAGG GGCAAGAAAC CTAAGCCGCC TGGTGGATGG CTCGGCTCGG GGCGCTGACG AA
J23SpTT_50                                         GGATGTAA GCCGGGAAGG AAACCGACCC GTTTAGTCTG CCGCTCCCAA CAGCCCGAGT TTCTTGCCCC TTGAGTAAGG GGCAAGAAAC CTAAGCCGCC TGGTGGATGG CTCGGCTCGG GGCGCTGACG AAGGG
J23SpTT_32                                         GGATGTAA GCCGGGAAGG AAACCGACCC GTTTAGTCTG CCGCTCCCAA CAGCCCGAGT TTCTTGCCCC TTGAGTAAGG GGCAAGAAAC CTAAGCCGCC TGGTGGATGG CTCGGCTCGG GGCGCTGACG AAGG
J23SpTT_12                                          GATGTAA GCCGGGAAGG AAACCGACCC GTTTAGTCTG CCGCTCCCAA CAGCCCGAGT TTCTTGCCCC TTGAGTAAGG GGCAAGAAAC CTAAGCCGCC TGGTGGATGG CTCGGCTCGG GGCGCTGACG AAGGG
J23SpTT_72                                        GGGATGTAA GCCGGGAAGG AAACCGACCC GTTTAGTCTG CCGCTCCCAA CAGCCCGAGT TTCTTGCCCC TTGAGTAAGG GGCAAGAAAC CTAAGCCGCC TGGTGGATGG CTCGGCTCGG GGCGCTGACG AAG
J23SpTT_27                                         GGATGTAA GCCGGGAAGG AAACCGACCC GTTTAGTCTG CCGCTCCCAA CAGCCCGAGT TTCTTGCCCC TTGAGTAAGG GGCAAGAAAC CTAAGCCGCC TGGTGGATGG CTCGGCTCGG GGCGCTGACG AAG
J23SpTT_17                         GT TTGATGGGGC GGGGATGTAA GCCGGGAAGG AAACCGACCC GTTTAGTCTG CCGCTCCCAA CAGCCCGAGT TTCTTGCCCC TTGAGTAAGG GGCAAGAAAC CTAAGCCGCC TGGTGGATGG CTCGGCTCGG G
J23SpTT_111           CGCA GAAGACGGGT TTGATGGGGC GGGGATGTAA GCCGGGAAGG AAACCGACCC GTTTAGTCTG CCGCTCCCAA CAGCCCGAGT TTCTTGCCCC TTGAGTAAGG GGCAAGAAAC CTAAGCCGCC TGGTGGAT
J23SpTT_34              CA GAAGACGGGT TTGATGGGGC GGGGATGTAA GCCGGGAAGG AAACCGACCC GTTTAGTCTG CCGCTCCCAA CAGCCCGAGT TTCTTGCCCC TTGAGTAAGG GGCAAGAAAC CTAAGCCGCC TGGTGGATG
J23SpTT_13             GCA GAAGACGGGT TTGATGGGGC GGGGATGTAA GCCGGGAAGG AAACCGACCC GTTTAGTCTG CCGCTCCCAA CAGCCCGAGT TTCTTGCCCC TTGAGTAAGG GGCAAGAAAC CTAAGCCGCC TGGTGGATG
J23SpTT_11             GCA GAAGACGGGT TTGATGGGGC GGGGATGTAA GCCGGGAAGG AAACCGACCC GTTTAGTCTG CCGCTCCCAA CAGCCCGAGT TTCTTGCCCC TTGAGTAAGG GGCAAGAAAC CTAAGCCGCC TGGTGGAT
J23SpTT_34              CA GAAGACGGGT TTGATGGGGC GGGGATGTAA GCCGGGAAGG AAACCGACCC GTTTAGTCTG CCGCTCCCAA CAGCCCGAGT TTCTTGCCCC TTGAGTAAGG GGCAAGAAAC CTAAGCCGCC TGGTGGAT
J23SpTT_9                A GAAGACGGGT TTGATGGGGC GGGGATGTAA GCCGGGAAGG AAACCGACCC GTTTAGTCTG CCGCTCCCAA CAGCCCGAGT TTCTTGCCCC TTGAGTAAGG GGCAAGAAAC CTAAGCCGCC TGGTGGATG
J23SpTT_20              CA GAAGACGGGT TTGATGGGGC GGGGATGTAA GCCGGGAAGG AAACCGACCC GTTTAGTCTG CCGCTCCCAA CAGCCCGAGT TTCTTGCCCC TTGAGTAAGG GGCAAGAAAC CTAAGCCGCC TGGTGGATGG
J23SpTT_13               A GAAGACGGGT TTGATGGGGC GGGGATGTAA GCCGGGAAGG AAACCGACCC GTTTAGTCTG CCGCTCCCAA CAGCCCGAGT TTCTTGCCCC TTGAGTAAGG GGCAAGAAAC CTAAGCCGCC TGGTGGATGG
J23SpTT_11               A GAAGACGGGT TTGATGGGGC GGGGATGTAA GCCGGGAAGG AAACCGACCC GTTTAGTCTG CCGCTCCCAA CAGCCCGAGT TTCTTGCCCC TTGAGTAAGG GGCAAGAAAC CTAAGCCGCC TGGTGGATGG C
J23SpTT_48                                             GTAA GCCGGGAAGG AAACCGACCC GTTTAGTCTG CCGCTCCCAA CAGCCCGAGT TTCTTGCCCC TTGAGTAAGG GGCAAGAAAC CTAAGCCGCC TGGTGGATGG CTCGGCTCGG GGCGCTGACG AAGGGC
J23SpTT_20                                             GTAA GCCGGGAAGG AAACCGACCC GTTTAGTCTG CCGCTCCCAA CAGCCCGAGT TTCTTGCCCC TTGAGTAAGG GGCAAGAAAC CTAAGCCGCC TGGTGGATGG CTCGGCTCGG GGCGCTGACG AAGGGCG
J23SpTT_9                       CGGGT TTGATGGGGC GGGGATGTAA GCCGGGAAGG AAACCGACCC GTTTAGTCTG CCGCTCCCAA CAGCCCGAGT TTCTTGCCCC TTGAGTAAGG GGCAAGAAAC CTAAGCCGCC TGGTGGATGG CTCGGCT
J23SpTT_38                                     C GGGGATGTAA GCCGGGAAGG AAACCGACCC GTTTAGTCTG CCGCTCCCAA CAGCCCGAGT TTCTTGCCCC TTGAGTAAGG GGCAAGAAAC CTAAGCCGCC TGGTGGATGG CTCGGCTCGG GGCGCTGACG
J23SpTT_26                                     C GGGGATGTAA GCCGGGAAGG AAACCGACCC GTTTAGTCTG CCGCTCCCAA CAGCCCGAGT TTCTTGCCCC TTGAGTAAGG GGCAAGAAAC CTAAGCCGCC TGGTGGATGG CTCGGCTCGG GGCGCTGAC
J23SpTT_28                                           ATGTAA GCCGGGAAGG AAACCGACCC GTTTAGTCTG CCGCTCCCAA CAGCCCGAGT TTCTTGCCCC TTGAGTAAGG GGCAAGAAAC CTAAGCCGCC TGGTGGATGG CTCGGCTCGG GGCGCTGACG AAGGG
J23SpTT_9                                           GATGTAA GCCGGGAAGG AAACCGACCC GTTTAGTCTG CCGCTCCCAA CAGCCCGAGT TTCTTGCCCC TTGAGTAAGG GGCAAGAAAC CTAAGCCGCC TGGTGGATGG CTCGGCTCGG GGCGCTGACG AAGG
J23SpTT_9                                           GATGTAA GCCGGGAAGG AAACCGACCC GTTTAGTCTG CCGCTCCCAA CAGCCCGAGT TTCTTGCCCC TTGAGTAAGG GGCAAGAAAC CTAAGCCGCC TGGTGGATGG CTCGGCTCGG GGCGCTGACG AAG
J23SpTT_21                                           ATGTAA GCCGGGAAGG AAACCGACCC GTTTAGTCTG CCGCTCCCAA CAGCCCGAGT TTCTTGCCCC TTGAGTAAGG GGCAAGAAAC CTAAGCCGCC TGGTGGATGG CTCGGCTCGG GGCGCTGACG AAGG
J23SpTT_12                                                 AA GCCGGGAAGG AAACCGACCC GTTTAGTCTG CCGCTCCCAA CAGCCCGAGT TTCTTGCCCC TTGAGTAAGG GGCAAGAAAC CTAAGCCGCC TGGTGGATGG CTCGGCTCGG GGCGCTGACG AAGGGCGTGG
J23SpTT_9                                                  AA GCCGGGAAGG AAACCGACCC GTTTAGTCTG CCGCTCCCAA CAGCCCGAGT TTCTTGCCCC TTGAGTAAGG GGCAAGAAAC CTAAGCCGCC TGGTGGATGG CTCGGCTCGG GGCGCTGACG AAGGGCGTG
J23SpTT_96                                             GTAA GCCGGGAAGG AAACCGACCC GTTTAGTCTG CCGCTCCCAA CAGCCCGAGT TTCTTGCCCC TTGAGTAAGG GGCAAGAAAC CTAAGCCGCC TGGTGGATGG CTCGGCTCGG GGCGCTGACG AAGGG
```

```
J23SpTT_8                                         TGTAA GCCGGGAAGGG AAACCGACCC GTTTAGTCTG CCGCTCCCAA CAGCCCGAGT TTCTTGCCCC TTGAGTAAGG GGCAAGAAAC CTAAGCCGCC TGGTGGATGG CTCGGCTCGG GGCGCTGACG AAGGG
J23SpTT_10                                                      GGG AAACCGACCC GTTTAGTCTG CCGCTCCCAA CAGCCCGAGT TTCTTGCCCC TTGAGTAAGG GGCAAGAAAC CTAAGCCGCC TGGTGGATGG CTCGGCTCGG GGCGCTGACG AAGGGCGTGG CAAGCTGCGA TAAGCCCCGG TGAG
J23SpTT_9                                                       GGG AAACCGACCC GTTTAGTCTG CCGCTCCCAA CAGCCCGAGT TTCTTGCCCC TTGAGTAAGG GGCAAGAAAC CTAAGCCGCC TGGTGGATGG CTCGGCTCGG GGCGCTGACG AAGGGCGTGG CAAGCTGCGA T
J23SpTT_8                                                       GGG AAACCGACCC GTTTAGTCTG CCGCTCCCAA CAGCCCGAGT TTCTTGCCCC TTGAGTAAGG GGCAAGAAAC CTAAGCCGCC TGGTGGATGG CTCGGCTCGG GGCGCTGACG AAGGGCGTGG CAAGCTGCG
J23SpTT_313       CGCA GAAGACGGGT TTGATGGGGC GGGGATGTAA GCCGGGAAGGG AAACCGACCC GTTTAGTCTG CCGCTCCCAA CAGCCCGAGT TTCTTGCCCC TTGAGTAAGG GGCAAGAAAC CTAAGCCGCC TGGTG
J23SpTT_180      CCGCA GAAGACGGGT TTGATGGGGC GGGGATGTAA GCCGGGAAGGG AAACCGACCC GTTTAGTCTG CCGCTCCCAA CAGCCCGAGT TTCTTGCCCC TTGAGTAAGG GGCAAGAAAC CTAAGCCGCC TGGTG
J23SpTT_150     CCCGCA GAAGACGGGT TTGATGGGGC GGGGATGTAA GCCGGGAAGGG AAACCGACCC GTTTAGTCTG CCGCTCCCAA CAGCCCGAGT TTCTTGCCCC TTGAGTAAGG GGCAAGAAAC CTAAGCCGCC TGGTG
J23SpTT_12      CCCGCA GAAGACGGGT TTGATGGGGC GGGGATGTAA GCCGGGAAGGG AAACCGACCC GTTTAGTCTG CCGCTCCCAA CAGCCCGAGT TTCTTGCCCC TTGAGTAAGG GGCAAGAAAC CTAAGCCGCC TGGTGG
J23SpTT_16       CCGCA GAAGACGGGT TTGATGGGGC GGGGATGTAA GCCGGGAAGGG AAACCGACCC GTTTAGTCTG CCGCTCCCAA CAGCCCGAGT TTCTTGCCCC TTGAGTAAGG GGCAAGAAAC CTAAGCCGCC TGGTGG
J23SpTT_38        CGCA GAAGACGGGT TTGATGGGGC GGGGATGTAA GCCGGGAAGGG AAACCGACCC GTTTAGTCTG CCGCTCCCAA CAGCCCGAGT TTCTTGCCCC TTGAGTAAGG GGCAAGAAAC CTAAGCCGCC TGGTGG
J23SpTT_9        CCGCA GAAGACGGGT TTGATGGGGC GGGGATGTAA GCCGGGAAGGG AAACCGACCC GTTTAGTCTG CCGCTCCCAA CAGCCCGAGT TTCTTGCCCC TTGAGTAAGG GGCAAGAAAC CTAAGCCGCC TGGT
J23SpTT_27      CCCGCA GAAGACGGGT TTGATGGGGC GGGGATGTAA GCCGGGAAGGG AAACCGACCC GTTTAGTCTG CCGCTCCCAA CAGCCCGAGT TTCTTGCCCC TTGAGTAAGG GGCAAGAAAC CTAAGCCGCC TG
J23SpTT_35         GCA GAAGACGGGT TTGATGGGGC GGGGATGTAA GCCGGGAAGGG AAACCGACCC GTTTAGTCTG CCGCTCCCAA CAGCCCGAGT TTCTTGCCCC TTGAGTAAGG GGCAAGAAAC CTAAGCCGCC TGGTG
J23SpTT_10          CA GAAGACGGGT TTGATGGGGC GGGGATGTAA GCCGGGAAGGG AAACCGACCC GTTTAGTCTG CCGCTCCCAA CAGCCCGAGT TTCTTGCCCC TTGAGTAAGG GGCAAGAAAC CTAAGCCGCC TGGTGG
J23SpTT_19                                           AA GCCGGGAAGGG AAACCGACCC GTTTAGTCTG CCGCTCCCAA CAGCCCGAGT TTCTTGCCCC TTGAGTAAGG GGCAAGAAAC CTAAGCCGCC TGGTGGATGG CTCGGCTCGG GGCGCTGACG AAGGGC
J23SpTT_9                                           TAA GCCGGGAAGGG AAACCGACCC GTTTAGTCTG CCGCTCCCAA CAGCCCGAGT TTCTTGCCCC TTGAGTAAGG GGCAAGAAAC CTAAGCCGCC TGGTGGATGG CTCGGCTCGG GGCGCTGACG AAGGGC
J23SpTT_15                                           AA GCCGGGAAGGG AAACCGACCC GTTTAGTCTG CCGCTCCCAA CAGCCCGAGT TTCTTGCCCC TTGAGTAAGG GGCAAGAAAC CTAAGCCGCC TGGTGGATGG CTCGGCTCGG GGCGCTGACG AAGGGCG
J23SpTT_59                                               CCGGGAAGGG AAACCGACCC GTTTAGTCTG CCGCTCCCAA CAGCCCGAGT TTCTTGCCCC TTGAGTAAGG GGCAAGAAAC CTAAGCCGCC TGGTGGATGG CTCGGCTCGG GGCGCTGACG AAGGGCGTGG
J23SpTT_11                                               CCGGGAAGGG AAACCGACCC GTTTAGTCTG CCGCTCCCAA CAGCCCGAGT TTCTTGCCCC TTGAGTAAGG GGCAAGAAAC CTAAGCCGCC TGGTGGATGG CTCGGCTCGG GGCGCTGACG AAGGGCGTGG C
J23SpTT_31                                               CCGGGAAGGG AAACCGACCC GTTTAGTCTG CCGCTCCCAA CAGCCCGAGT TTCTTGCCCC TTGAGTAAGG GGCAAGAAAC CTAAGCCGCC TGGTGGATGG CTCGGCTCGG GGCGCTGACG AAGGGCGTG
J23SpTT_19                                       ATGTAA GCCGGGAAGGG AAACCGACCC GTTTAGTCTG CCGCTCCCAA CAGCCCGAGT TTCTTGCCCC TTGAGTAAGG GGCAAGAAAC CTAAGCCGCC TGGTGGATGG CTCGGCTCGG GGCGCTGACG AAG
J23SpTT_54                                         GTAA GCCGGGAAGGG AAACCGACCC GTTTAGTCTG CCGCTCCCAA CAGCCCGAGT TTCTTGCCCC TTGAGTAAGG GGCAAGAAAC CTAAGCCGCC TGGTGGATGG CTCGGCTCGG GGCGCTGACG AAG
J23SpTT_43                                         GTAA GCCGGGAAGGG AAACCGACCC GTTTAGTCTG CCGCTCCCAA CAGCCCGAGT TTCTTGCCCC TTGAGTAAGG GGCAAGAAAC CTAAGCCGCC TGGTGGATGG CTCGGCTCGG GGCGCTGACG AAGG
J23SpTT_9                                           TAA GCCGGGAAGGG AAACCGACCC GTTTAGTCTG CCGCTCCCAA CAGCCCGAGT TTCTTGCCCC TTGAGTAAGG GGCAAGAAAC CTAAGCCGCC TGGTGGATGG CTCGGCTCGG GGCGCTGACG AAGG
J23SpTT_9            A GAAGACGGGT TTGATGGGGC GGGGATGTAA GCCGGGAAGGG AAACCGACCC GTTTAGTCTG CCGCTCCCAA CAGCCCGAGT TTCTTGCCCC TTGAGTAAGG GGCAAGAAAC CTAAGCCGCC TGGTGGAT
J23SpTT_92                                               CCGGGAAGGG AAACCGACCC GTTTAGTCTG CCGCTCCCAA CAGCCCGAGT TTCTTGCCCC TTGAGTAAGG GGCAAGAAAC CTAAGCCGCC TGGTGGATGG CTCGGCTCGG GGCGCTGACG AAGGGCG
J23SpTT_27                                           AA GCCGGGAAGGG AAACCGACCC GTTTAGTCTG CCGCTCCCAA CAGCCCGAGT TTCTTGCCCC TTGAGTAAGG GGCAAGAAAC CTAAGCCGCC TGGTGGATGG CTCGGCTCGG GGCGCTGACG AAGG
J23SpTT_25                                           AA GCCGGGAAGGG AAACCGACCC GTTTAGTCTG CCGCTCCCAA CAGCCCGAGT TTCTTGCCCC TTGAGTAAGG GGCAAGAAAC CTAAGCCGCC TGGTGGATGG CTCGGCTCGG GGCGCTGACG AAGGG
J23SpTT_17                                            A GCCGGGAAGGG AAACCGACCC GTTTAGTCTG CCGCTCCCAA CAGCCCGAGT TTCTTGCCCC TTGAGTAAGG GGCAAGAAAC CTAAGCCGCC TGGTGGATGG CTCGGCTCGG GGCGCTGACG AAGGG
J23SpTT_14                                              GCCGGGAAGGG AAACCGACCC GTTTAGTCTG CCGCTCCCAA CAGCCCGAGT TTCTTGCCCC TTGAGTAAGG GGCAAGAAAC CTAAGCCGCC TGGTGGATGG CTCGGCTCGG GGCGCTGACG AAGGG
J23SpTT_118         CA GAAGACGGGT TTGATGGGGC GGGGATGTAA GCCGGGAAGGG AAACCGACCC GTTTAGTCTG CCGCTCCCAA CAGCCCGAGT TTCTTGCCCC TTGAGTAAGG GGCAAGAAAC CTAAGCCGCC TGGTG
J23SpTT_49           A GAAGACGGGT TTGATGGGGC GGGGATGTAA GCCGGGAAGGG AAACCGACCC GTTTAGTCTG CCGCTCCCAA CAGCCCGAGT TTCTTGCCCC TTGAGTAAGG GGCAAGAAAC CTAAGCCGCC TGGTG
J23SpTT_16                  GACGGGT TTGATGGGGC GGGGATGTAA GCCGGGAAGGG AAACCGACCC GTTTAGTCTG CCGCTCCCAA CAGCCCGAGT TTCTTGCCCC TTGAGTAAGG GGCAAGAAAC CTAAGCCGCC TGGTGGATG
J23SpTT_10                  GACGGGT TTGATGGGGC GGGGATGTAA GCCGGGAAGGG AAACCGACCC GTTTAGTCTG CCGCTCCCAA CAGCCCGAGT TTCTTGCCCC TTGAGTAAGG GGCAAGAAAC CTAAGCCGCC TGGTGGATGG
J23SpTT_9                  AGACGGGT TTGATGGGGC GGGGATGTAA GCCGGGAAGGG AAACCGACCC GTTTAGTCTG CCGCTCCCAA CAGCCCGAGT TTCTTGCCCC TTGAGTAAGG GGCAAGAAAC CTAAGCCGCC TGGTGGATG
J23SpTT_9                    CGGGT TTGATGGGGC GGGGATGTAA GCCGGGAAGGG AAACCGACCC GTTTAGTCTG CCGCTCCCAA CAGCCCGAGT TTCTTGCCCC TTGAGTAAGG GGCAAGAAAC CTAAGCCGCC TGGTGGATGG C
J23SpTT_9                    CGGGT TTGATGGGGC GGGGATGTAA GCCGGGAAGGG AAACCGACCC GTTTAGTCTG CCGCTCCCAA CAGCCCGAGT TTCTTGCCCC TTGAGTAAGG GGCAAGAAAC CTAAGCCGCC TGGTGGATGG
J23SpTT_9                   ACGGGT TTGATGGGGC GGGGATGTAA GCCGGGAAGGG AAACCGACCC GTTTAGTCTG CCGCTCCCAA CAGCCCGAGT TTCTTGCCCC TTGAGTAAGG GGCAAGAAAC CTAAGCCGCC TGGTGGATG
J23SpTT_8                                                 CGGGAAGGG AAACCGACCC GTTTAGTCTG CCGCTCCCAA CAGCCCGAGT TTCTTGCCCC TTGAGTAAGG GGCAAGAAAC CTAAGCCGCC TGGTGGATGG CTCGGCTCGG GGCGCTGACG AAGGGCG
J23SpTT_134                                              CCGGGAAGGG AAACCGACCC GTTTAGTCTG CCGCTCCCAA CAGCCCGAGT TTCTTGCCCC TTGAGTAAGG GGCAAGAAAC CTAAGCCGCC TGGTGGATGG CTCGGCTCGG GGCGCTGACG AAGGGC
J23SpTT_18                                                CGGGAAGGG AAACCGACCC GTTTAGTCTG CCGCTCCCAA CAGCCCGAGT TTCTTGCCCC TTGAGTAAGG GGCAAGAAAC CTAAGCCGCC TGGTGGATGG CTCGGCTCGG GGCGCTGACG AAGGGC
J23SpTT_274                                              CCGGGAAGGG AAACCGACCC GTTTAGTCTG CCGCTCCCAA CAGCCCGAGT TTCTTGCCCC TTGAGTAAGG GGCAAGAAAC CTAAGCCGCC TGGTGGATGG CTCGGCTCGG GGCGCTGACG AAGGG
J23SpTT_11                                               CCGGGAAGGG AAACCGACCC GTTTAGTCTG CCGCTCCCAA CAGCCCGAGT TTCTTGCCCC TTGAGTAAGG GGCAAGAAAC CTAAGCCGCC TGGTGGATGG CTCGGCTCGG GGCGCTGACG AAGGGG
J23SpTT_23                                           AA GCCGGGAAGGG AAACCGACCC GTTTAGTCTG CCGCTCCCAA CAGCCCGAGT TTCTTGCCCC TTGAGTAAGG GGCAAGAAAC CTAAGCCGCC TGGTGGATGG CTCGGCTCGG GGCGCTGACG AAG
J23SpTT_13                                            A GCCGGGAAGGG AAACCGACCC GTTTAGTCTG CCGCTCCCAA CAGCCCGAGT TTCTTGCCCC TTGAGTAAGG GGCAAGAAAC CTAAGCCGCC TGGTGGATGG CTCGGCTCGG GGCGCTGACG AAG
J23SpTT_11                                            A GCCGGGAAGGG AAACCGACCC GTTTAGTCTG CCGCTCCCAA CAGCCCGAGT TTCTTGCCCC TTGAGTAAGG GGCAAGAAAC CTAAGCCGCC TGGTGGATGG CTCGGCTCGG GGCGCTGACG AAGG
J23SpTT_19                                                  GGAAGGG AAACCGACCC GTTTAGTCTG CCGCTCCCAA CAGCCCGAGT TTCTTGCCCC TTGAGTAAGG GGCAAGAAAC CTAAGCCGCC TGGTGGATGG CTCGGCTCGG GGCGCTGACG AAGGGCG
J23SpTT_91        CGCA GAAGACGGGT TTGATGGGGC GGGGATGTAA GCCGGGAAGGG AAACCGACCC GTTTAGTCTG CCGCTCCCAA CAGCCCGAGT TTCTTGCCCC TTGAGTAAGG GGCAAGAAAC CTAAGCCGCC TG
J23SpTT_37       CCGCA GAAGACGGGT TTGATGGGGC GGGGATGTAA GCCGGGAAGGG AAACCGACCC GTTTAGTCTG CCGCTCCCAA CAGCCCGAGT TTCTTGCCCC TTGAGTAAGG GGCAAGAAAC CTAAGCCGCC TG
J23SpTT_10        CGCA GAAGACGGGT TTGATGGGGC GGGGATGTAA GCCGGGAAGGG AAACCGACCC GTTTAGTCTG CCGCTCCCAA CAGCCCGAGT TTCTTGCCCC TTGAGTAAGG GGCAAGAAAC CTAAGCCGCC TGG
J23SpTT_8          GCA GAAGACGGGT TTGATGGGGC GGGGATGTAA GCCGGGAAGGG AAACCGACCC GTTTAGTCTG CCGCTCCCAA CAGCCCGAGT TTCTTGCCCC TTGAGTAAGG GGCAAGAAAC CTAAGCCGCC TG
J23SpTT_14        CGCA GAAGACGGGT TTGATGGGGC GGGGATGTAA GCCGGGAAGGG AAACCGACCC GTTTAGTCTG CCGCTCCCAA CAGCCCGAGT TTCTTGCCCC TTGAGTAAGG GGCAAGAAAC CTAAGCCGCC T
J23SpTT_11      CCCGCA GAAGACGGGT TTGATGGGGC GGGGATGTAA GCCGGGAAGGG AAACCGACCC GTTTAGTCTG CCGCTCCCAA CAGCCCGAGT TTCTTGCCCC TTGAGTAAGG GGCAAGAAAC CTAAGCCGCC
J23SpTT_11        CGCA GAAGACGGGT TTGATGGGGC GGGGATGTAA GCCGGGAAGGG AAACCGACCC GTTTAGTCTG CCGCTCCCAA CAGCCCGAGT TTCTTGCCCC TTGAGTAAGG GGCAAGAAAC CTAAGCCGCC
J23SpTT_24          CA GAAGACGGGT TTGATGGGGC GGGGATGTAA GCCGGGAAGGG AAACCGACCC GTTTAGTCTG CCGCTCCCAA CAGCCCGAGT TTCTTGCCCC TTGAGTAAGG GGCAAGAAAC CTAAGCCGCC TG
J23SpTT_20                                                  GGAAGGG AAACCGACCC GTTTAGTCTG CCGCTCCCAA CAGCCCGAGT TTCTTGCCCC TTGAGTAAGG GGCAAGAAAC CTAAGCCGCC TGGTGGATGG CTCGGCTCGG GGCGCTGACG AAGGGC
J23SpTT_10                                              GCCGGGAAGGG AAACCGACCC GTTTAGTCTG CCGCTCCCAA CAGCCCGAGT TTCTTGCCCC TTGAGTAAGG GGCAAGAAAC CTAAGCCGCC TGGTGGATGG CTCGGCTCGG GGCGCTGACG AAG
J23SpTT_33                                                CGGGAAGGG AAACCGACCC GTTTAGTCTG CCGCTCCCAA CAGCCCGAGT TTCTTGCCCC TTGAGTAAGG GGCAAGAAAC CTAAGCCGCC TGGTGGATGG CTCGGCTCGG GGCGCTGACG AAGGG
J23SpTT_185                                              CCGGGAAGGG AAACCGACCC GTTTAGTCTG CCGCTCCCAA CAGCCCGAGT TTCTTGCCCC TTGAGTAAGG GGCAAGAAAC CTAAGCCGCC TGGTGGATGG CTCGGCTCGG GGCGCTGACG AAGG
J23SpTT_167                                              CCGGGAAGGG AAACCGACCC GTTTAGTCTG CCGCTCCCAA CAGCCCGAGT TTCTTGCCCC TTGAGTAAGG GGCAAGAAAC CTAAGCCGCC TGGTGGATGG CTCGGCTCGG GGCGCTGACG AAG
J23SpTT_15                                                  GGAAGGG AAACCGACCC GTTTAGTCTG CCGCTCCCAA CAGCCCGAGT TTCTTGCCCC TTGAGTAAGG GGCAAGAAAC CTAAGCCGCC TGGTGGATGG CTCGGCTCGG GGCGCTGACG AAGGG
J23SpTT_27                   CGGGT TTGATGGGGC GGGGATGTAA GCCGGGAAGGG AAACCGACCC GTTTAGTCTG CCGCTCCCAA CAGCCCGAGT TTCTTGCCCC TTGAGTAAGG GGCAAGAAAC CTAAGCCGCC TGGTGGAT
J23SpTT_11                  GACGGGT TTGATGGGGC GGGGATGTAA GCCGGGAAGGG AAACCGACCC GTTTAGTCTG CCGCTCCCAA CAGCCCGAGT TTCTTGCCCC TTGAGTAAGG GGCAAGAAAC CTAAGCCGCC TGGTGGAT
J23SpTT_24                   CGGGT TTGATGGGGC GGGGATGTAA GCCGGGAAGGG AAACCGACCC GTTTAGTCTG CCGCTCCCAA CAGCCCGAGT TTCTTGCCCC TTGAGTAAGG GGCAAGAAAC CTAAGCCGCC TGGTGGATG
J23SpTT_24                                                CGGGAAGGG AAACCGACCC GTTTAGTCTG CCGCTCCCAA CAGCCCGAGT TTCTTGCCCC TTGAGTAAGG GGCAAGAAAC CTAAGCCGCC TGGTGGATGG CTCGGCTCGG GGCGCTGACG AAGG
J23SpTT_56                                  C GGGGATGTAA GCCGGGAAGGG AAACCGACCC GTTTAGTCTG CCGCTCCCAA CAGCCCGAGT TTCTTGCCCC TTGAGTAAGG GGCAAGAAAC CTAAGCCGCC TGGTGGATGG CTCGGCTCGG G
J23SpTT_45                  GACGGGT TTGATGGGGC GGGGATGTAA GCCGGGAAGGG AAACCGACCC GTTTAGTCTG CCGCTCCCAA CAGCCCGAGT TTCTTGCCCC TTGAGTAAGG GGCAAGAAAC CTAAGCCGCC TGGTG
J23SpTT_13                 AGACGGGT TTGATGGGGC GGGGATGTAA GCCGGGAAGGG AAACCGACCC GTTTAGTCTG CCGCTCCCAA CAGCCCGAGT TTCTTGCCCC TTGAGTAAGG GGCAAGAAAC CTAAGCCGCC TGGTG
J23SpTT_9                 AAGACGGGT TTGATGGGGC GGGGATGTAA GCCGGGAAGGG AAACCGACCC GTTTAGTCTG CCGCTCCCAA CAGCCCGAGT TTCTTGCCCC TTGAGTAAGG GGCAAGAAAC CTAAGCCGCC TGGTG
J23SpTT_12                  ACGGGT TTGATGGGGC GGGGATGTAA GCCGGGAAGGG AAACCGACCC GTTTAGTCTG CCGCTCCCAA CAGCCCGAGT TTCTTGCCCC TTGAGTAAGG GGCAAGAAAC CTAAGCCGCC TGGTG
J23SpTT_14                                                      GGG AAACCGACCC GTTTAGTCTG CCGCTCCCAA CAGCCCGAGT TTCTTGCCCC TTGAGTAAGG GGCAAGAAAC CTAAGCCGCC TGGTGGATGG CTCGGCTCGG GGCGCTGACG AAGGGCGTGG
J23SpTT_9                                                       GGG AAACCGACCC GTTTAGTCTG CCGCTCCCAA CAGCCCGAGT TTCTTGCCCC TTGAGTAAGG GGCAAGAAAC CTAAGCCGCC TGGTGGATGG CTCGGCTCGG GGCGCTGACG AAGGGCGTG
J23SpTT_11           A GAAGACGGGT TTGATGGGGC GGGGATGTAA GCCGGGAAGGG AAACCGACCC GTTTAGTCTG CCGCTCCCAA CAGCCCGAGT TTCTTGCCCC TTGAGTAAGG GGCAAGAAAC CTAAGCCGCC TG
J23SpTT_13                                                  GGAAGGG AAACCGACCC GTTTAGTCTG CCGCTCCCAA CAGCCCGAGT TTCTTGCCCC TTGAGTAAGG GGCAAGAAAC CTAAGCCGCC TGGTGGATGG CTCGGCTCGG GGCGCTGACG AAGG
J23SpTT_25                                                CGGGAAGGG AAACCGACCC GTTTAGTCTG CCGCTCCCAA CAGCCCGAGT TTCTTGCCCC TTGAGTAAGG GGCAAGAAAC CTAAGCCGCC TGGTGGATGG CTCGGCTCGG GGCGCTGACG AAG
J23SpTT_26                                                      GGG AAACCGACCC GTTTAGTCTG CCGCTCCCAA CAGCCCGAGT TTCTTGCCCC TTGAGTAAGG GGCAAGAAAC CTAAGCCGCC TGGTGGATGG CTCGGCTCGG GGCGCTGACG AAGGGCG
J23SpTT_25                                                  GGAAGGG AAACCGACCC GTTTAGTCTG CCGCTCCCAA CAGCCCGAGT TTCTTGCCCC TTGAGTAAGG GGCAAGAAAC CTAAGCCGCC TGGTGGATGG CTCGGCTCGG GGCGCTGACG AAG
J23SpTT_38                                                      GGG AAACCGACCC GTTTAGTCTG CCGCTCCCAA CAGCCCGAGT TTCTTGCCCC TTGAGTAAGG GGCAAGAAAC CTAAGCCGCC TGGTGGATGG CTCGGCTCGG GGCGCTGACG AAGGGC
J23SpTT_9                       GT TTGATGGGGC GGGGATGTAA GCCGGGAAGGG AAACCGACCC GTTTAGTCTG CCGCTCCCAA CAGCCCGAGT TTCTTGCCCC TTGAGTAAGG GGCAAGAAAC CTAAGCCGCC TGGTGGAT
J23SpTT_8                       GT TTGATGGGGC GGGGATGTAA GCCGGGAAGGG AAACCGACCC GTTTAGTCTG CCGCTCCCAA CAGCCCGAGT TTCTTGCCCC TTGAGTAAGG GGCAAGAAAC CTAAGCCGCC TGGTGGATG
J23SpTT_49                   CGGGT TTGATGGGGC GGGGATGTAA GCCGGGAAGGG AAACCGACCC GTTTAGTCTG CCGCTCCCAA CAGCCCGAGT TTCTTGCCCC TTGAGTAAGG GGCAAGAAAC CTAAGCCGCC TGGTG
J23SpTT_9                    CGGGT TTGATGGGGC GGGGATGTAA GCCGGGAAGGG AAACCGACCC GTTTAGTCTG CCGCTCCCAA CAGCCCGAGT TTCTTGCCCC TTGAGTAAGG GGCAAGAAAC CTAAGCCGCC TGGTGG
J23SpTT_9                                                        GG AAACCGACCC GTTTAGTCTG CCGCTCCCAA CAGCCCGAGT TTCTTGCCCC TTGAGTAAGG GGCAAGAAAC CTAAGCCGCC TGGTGGATGG CTCGGCTCGG GGCGCTGACG AAGGGC
J23SpTT_64                                                      GGG AAACCGACCC GTTTAGTCTG CCGCTCCCAA CAGCCCGAGT TTCTTGCCCC TTGAGTAAGG GGCAAGAAAC CTAAGCCGCC TGGTGGATGG CTCGGCTCGG GGCGCTGACG AAGGG
J23SpTT_45                                                      GGG AAACCGACCC GTTTAGTCTG CCGCTCCCAA CAGCCCGAGT TTCTTGCCCC TTGAGTAAGG GGCAAGAAAC CTAAGCCGCC TGGTGGATGG CTCGGCTCGG GGCGCTGACG AAGG
J23SpTT_26                                                       GG AAACCGACCC GTTTAGTCTG CCGCTCCCAA CAGCCCGAGT TTCTTGCCCC TTGAGTAAGG GGCAAGAAAC CTAAGCCGCC TGGTGGATGG CTCGGCTCGG GGCGCTGACG AAGGG
J23SpTT_15                                                        G AAACCGACCC GTTTAGTCTG CCGCTCCCAA CAGCCCGAGT TTCTTGCCCC TTGAGTAAGG GGCAAGAAAC CTAAGCCGCC TGGTGGATGG CTCGGCTCGG GGCGCTGACG AAGGG
J23SpTT_47                                                      GGG AAACCGACCC GTTTAGTCTG CCGCTCCCAA CAGCCCGAGT TTCTTGCCCC TTGAGTAAGG GGCAAGAAAC CTAAGCCGCC TGGTGGATGG CTCGGCTCGG GGCGCTGACG AAG
J23SpTT_11                                  C GGGGATGTAA GCCGGGAAGGG AAACCGACCC GTTTAGTCTG CCGCTCCCAA CAGCCCGAGT TTCTTGCCCC TTGAGTAAGG GGCAAGAAAC CTAAGCCGCC TGGTGGATGG CTCGGC
J23SpTT_9                                                        GG AAACCGACCC GTTTAGTCTG CCGCTCCCAA CAGCCCGAGT TTCTTGCCCC TTGAGTAAGG GGCAAGAAAC CTAAGCCGCC TGGTGGATGG CTCGGCTCGG GGCGCTGACG AAGG
J23SpTT_8                                                        GG AAACCGACCC GTTTAGTCTG CCGCTCCCAA CAGCCCGAGT TTCTTGCCCC TTGAGTAAGG GGCAAGAAAC CTAAGCCGCC TGGTGGATGG CTCGGCTCGG GGCGCTGACG AAG
J23SpTT_15                      GT TTGATGGGGC GGGGATGTAA GCCGGGAAGGG AAACCGACCC GTTTAGTCTG CCGCTCCCAA CAGCCCGAGT TTCTTGCCCC TTGAGTAAGG GGCAAGAAAC CTAAGCCGCC TGGTG
J23SpTT_9                                                           AAACCGACCC GTTTAGTCTG CCGCTCCCAA CAGCCCGAGT TTCTTGCCCC TTGAGTAAGG GGCAAGAAAC CTAAGCCGCC TGGTGGATGG CTCGGCTCGG GGCGCTGACG AAGGG
J23SpTT_15                   CGGGT TTGATGGGGC GGGGATGTAA GCCGGGAAGGG AAACCGACCC GTTTAGTCTG CCGCTCCCAA CAGCCCGAGT TTCTTGCCCC TTGAGTAAGG GGCAAGAAAC CTAAGCCGCC TG
J23SpTT_11                                                             CGACCC GTTTAGTCTG CCGCTCCCAA CAGCCCGAGT TTCTTGCCCC TTGAGTAAGG GGCAAGAAAC CTAAGCCGCC TGGTGGATGG CTCGGCTCGG GGCGCTGACG AAGGGCG
J23SpTT_13                                                          AACCGACCC GTTTAGTCTG CCGCTCCCAA CAGCCCGAGT TTCTTGCCCC TTGAGTAAGG GGCAAGAAAC CTAAGCCGCC TGGTGGATGG CTCGGCTCGG GGCGCTGACG AAGGG
J23SpTT_23                                                             CGACCC GTTTAGTCTG CCGCTCCCAA CAGCCCGAGT TTCTTGCCCC TTGAGTAAGG GGCAAGAAAC CTAAGCCGCC TGGTGGATGG CTCGGCTCGG GGCGCTGACG AAGGGC
J23SpTT_15                                                            CCGACCC GTTTAGTCTG CCGCTCCCAA CAGCCCGAGT TTCTTGCCCC TTGAGTAAGG GGCAAGAAAC CTAAGCCGCC TGGTGGATGG CTCGGCTCGG GGCGCTGACG AAGGG
```

```
J23SpTT_10                                         GACCC GTTTAGTCTG CCGCTCCCAA CAGCCCGAGT TTCTTGCCCC TTGAGTAAGG GGCAAGAAAC CTAAGCCGCC TGGTGGATGG CTCGGCTCGG GGCGCTGACG AAGGGC
J23SpTT_38                                        CGACCC GTTTAGTCTG CCGCTCCCAA CAGCCCGAGT TTCTTGCCCC TTGAGTAAGG GGCAAGAAAC CTAAGCCGCC TGGTGGATGG CTCGGCTCGG GGCGCTGACG AAGGG
J23SpTT_13                                       CCGACCC GTTTAGTCTG CCGCTCCCAA CAGCCCGAGT TTCTTGCCCC TTGAGTAAGG GGCAAGAAAC CTAAGCCGCC TGGTGGATGG CTCGGCTCGG GGCGCTGACG AAGG
J23SpTT_15                                        CGACCC GTTTAGTCTG CCGCTCCCAA CAGCCCGAGT TTCTTGCCCC TTGAGTAAGG GGCAAGAAAC CTAAGCCGCC TGGTGGATGG CTCGGCTCGG GGCGCTGACG AAGG
J23SpTT_12                                           CCC GTTTAGTCTG CCGCTCCCAA CAGCCCGAGT TTCTTGCCCC TTGAGTAAGG GGCAAGAAAC CTAAGCCGCC TGGTGGATGG CTCGGCTCGG GGCGCTGACG AAGGGC
J23SpTT_18                                        CGACCC GTTTAGTCTG CCGCTCCCAA CAGCCCGAGT TTCTTGCCCC TTGAGTAAGG GGCAAGAAAC CTAAGCCGCC TGGTGGATGG CTCGGCTCGG GGCGCTGACG AAG
J23SpTT_25                                           CCC GTTTAGTCTG CCGCTCCCAA CAGCCCGAGT TTCTTGCCCC TTGAGTAAGG GGCAAGAAAC CTAAGCCGCC TGGTGGATGG CTCGGCTCGG GGCGCTGACG AAGGG
J23SpTT_16              C GGGGATGTAA GCCGGGAAGGG AAACCGACCC GTTTAGTCTG CCGCTCCCAA CAGCCCGAGT TTCTTGCCCC TTGAGTAAGG GGCAAGAAAC CTAAGCCGCC TGGTGGATG
J23SpTT_11                                           CCC GTTTAGTCTG CCGCTCCCAA CAGCCCGAGT TTCTTGCCCC TTGAGTAAGG GGCAAGAAAC CTAAGCCGCC TGGTGGATGG CTCGGCTCGG GGCGCTGACG AAGG
J23SpTT_8                                                GTTTAGTCTG CCGCTCCCAA CAGCCCGAGT TTCTTGCCCC TTGAGTAAGG GGCAAGAAAC CTAAGCCGCC TGGTGGATGG CTCGGCTCGG GGCGCTGACG AAGGGCG
J23SpTT_11                                               GTTTAGTCTG CCGCTCCCAA CAGCCCGAGT TTCTTGCCCC TTGAGTAAGG GGCAAGAAAC CTAAGCCGCC TGGTGGATGG CTCGGCTCGG GGCGCTGACG AAGGGC
J23SpTT_16                                           CCC GTTTAGTCTG CCGCTCCCAA CAGCCCGAGT TTCTTGCCCC TTGAGTAAGG GGCAAGAAAC CTAAGCCGCC TGGTGGATGG CTCGGCTCGG GGCGCTGACG AAG
J23SpTT_15                                               GTTTAGTCTG CCGCTCCCAA CAGCCCGAGT TTCTTGCCCC TTGAGTAAGG GGCAAGAAAC CTAAGCCGCC TGGTGGATGG CTCGGCTCGG GGCGCTGACG AAGGG
J23SpTT_22              C GGGGATGTAA GCCGGGAAGGG AAACCGACCC GTTTAGTCTG CCGCTCCCAA CAGCCCGAGT TTCTTGCCCC TTGAGTAAGG GGCAAGAAAC CTAAGCCGCC TGGTG
J23SpTT_13                                               GTTTAGTCTG CCGCTCCCAA CAGCCCGAGT TTCTTGCCCC TTGAGTAAGG GGCAAGAAAC CTAAGCCGCC TGGTGGATGG CTCGGCTCGG GGCGCTGACG AAGG
J23SpTT_10                                                                 CTCCCAA CAGCCCGAGT TTCTTGCCCC TTGAGTAAGG GGCAAGAAAC CTAAGCCGCC TGGTGGATGG CTCGGCTCGG GGCGCTGACG AAGG
```
