## Supplementary Figure S2d for "Permuted 23S rRNA is integrated in 50S ribosome particles in *Thermococcus barophilus*"

```
            ______________________________23S : H99-101_____________________________________________|_23S : TH3p__|_Junction|_23S : TH5p_|____23S____________________________________________
                                                                                                   ..<<<< <<.<<<<<          >>>> >.>>>>>>..
J23SpTA_41        CGCAGA AGACGGGTTT GATGGGGCGG GGATGTAAGC GGGAAGGGAA ACCGACCCGT TTAGTCTGCC GCTCCCAACA GCCCGAGTTT CTAGCCCCTT GAGTAAGGGG CAAGAAACCT AAGCCGCCTG GTGGATGGCT CGGCTCGGGG CGCTGACGAA GGACGCGTGG ACG
J23SpTA_21        CGCAGA AGACGGGTTT GATGGGGCGG GGATGTAAGC GGGAAGGGAA ACCGACCCGT TTAGTCTGCC GCTCCCAACA GCCCGAGTTT CTAGCCCCTT GAGTAAGGGG CAAGAAACCT AAGCCGCCTG GTGGATGGCT CGGCTCGGGG CGCTGACGAA GGACGCGTGG ACGC
J23SpTA_16      CCCGCAGA AGACGGGTTT GATGGGGCGG GGATGTAAGC GGGAAGGGAA ACCGACCCGT TTAGTCTGCC GCTCCCAACA GCCCGAGTTT CTAGCCCCTT GAGTAAGGGG CAAGAAACCT AAGCCGCCTG GTGGATGGCT CGGCTCGGGG CGCTGACGAA GGACGCGTGG ACG
J23SpTA_47      CCCGCAGA AGACGGGTTT GATGGGGCGG GGATGTAAGC GGGAAGGGAA ACCGACCCGT TTAGTCTGCC GCTCCCAACA GCCCGAGTTT CTAGCCCCTT GAGTAAGGGG CAAGAAACCT AAGCCGCCTG GTGGATGGCT CGGCTCGGGG CGCTGACGAA GGACGCGTGG ACGG
J23SpTA_62       CCGCAGA AGACGGGTTT GATGGGGCGG GGATGTAAGC GGGAAGGGAA ACCGACCCGT TTAGTCTGCC GCTCCCAACA GCCCGAGTTT CTAGCCCCTT GAGTAAGGGG CAAGAAACCT AAGCCGCCTG GTGGATGGCT CGGCTCGGGG CGCTGACGAA GGACGCGTGG ACGG
J23SpTA_21     GCCCGCAGA AGACGGGTTT GATGGGGCGG GGATGTAAGC GGGAAGGGAA ACCGACCCGT TTAGTCTGCC GCTCCCAACA GCCCGAGTTT CTAGCCCCTT GAGTAAGGGG CAAGAAACCT AAGCCGCCTG GTGGATGGCT CGGCTCGGGG CGCTGACGAA GGACGCGTGG ACGC
J23SpTA_21     GCCCGCAGA AGACGGGTTT GATGGGGCGG GGATGTAAGC GGGAAGGGAA ACCGACCCGT TTAGTCTGCC GCTCCCAACA GCCCGAGTTT CTAGCCCCTT GAGTAAGGGG CAAGAAACCT AAGCCGCCTG GTGGATGGCT CGGCTCGGGG CGCTGACGAA GGACGCGTGG ACG
J23SpTA_19      CCCGCAGA AGACGGGTTT GATGGGGCGG GGATGTAAGC GGGAAGGGAA ACCGACCCGT TTAGTCTGCC GCTCCCAACA GCCCGAGTTT CTAGCCCCTT GAGTAAGGGG CAAGAAACCT AAGCCGCCTG GTGGATGGCT CGGCTCGGGG CGCTGACGAA GGACGCGTGG ACGCC
J23SpTA_17    CGCCCGCAGA AGACGGGTTT GATGGGGCGG GGATGTAAGC GGGAAGGGAA ACCGACCCGT TTAGTCTGCC GCTCCCAACA GCCCGAGTTT CTAGCCCCTT GAGTAAGGGG CAAGAAACCT AAGCCGCCTG GTGGATGGCT CGGCTCGGGG CGCTGACGAA GGACGCGTGG ACG
J23SpTA_10       CCGCAGA AGACGGGTTT GATGGGGCGG GGATGTAAGC GGGAAGGGAA ACCGACCCGT TTAGTCTGCC GCTCCCAACA GCCCGAGTTT CTAGCCCCTT GAGTAAGGGG CAAGAAACCT AAGCCGCCTG GTGGATGGCT CGGCTCGGGG CGCTGACGAA GGACGCGTGG ACGC
J23SpTA_5_       CCGCAGA AGACGGGTTT GATGGGGCGG GGATGTAAGC GGGAAGGGAA ACCGACCCGT TTAGTCTGCC GCTCCCAACA GCCCGAGTTT CTAGCCCCTT GAGTAAGGGG CAAGAAACCT AAGCCGCCTG GTGGATGGCT CGGCTCGGGG CGCTGACGAA GGACGCGTGG ACGCA
J23SpTA_17       CCGCAGA AGACGGGTTT GATGGGGCGG GGATGTAAGC GGGAAGGGAA ACCGACCCGT TTAGTCTGCC GCTCCCAACA GCCCGAGTTT CTAGCCCCTT GAGTAAGGGG CAAGAAACCT AAGCCGCCTG GTGGATGGCT CGGCTCGGGG CGCTGACGAA GGACGCGTGG ACGCC
J23SpTA_10      CCCGCAGA AGACGGGTTT GATGGGGCGG GGATGTAAGC GGGAAGGGAA ACCGACCCGT TTAGTCTGCC GCTCCCAACA GCCCGAGTTT CTAGCCCCTT GAGTAAGGGG CAAGAAACCT AAGCCGCCTG GTGGATGGCT CGGCTCGGGG CGCTGACGAA GGACGCGTGG ACAC
J23SpTA_13     GCCCGCAGA AGACGGGTTT GATGGGGCGG GGATGTAAGC GGGAAGGGAA ACCGACCCGT TTAGTCTGCC GCTCCCAACA GCCCGAGTTT CTAGCCCCTT GAGTAAGGGG CAAGAAACCT AAGCCGCCTG GTGGATGGCT CGGCTCGGGG CGCTGACGAA GGACGCGTGG AC
J23SpTA_10    CGCCCGCAGA AGACGGGTTT GATGGGGCGG GGATGTAAGC GGGAAGGGAA ACCGACCCGT TTAGTCTGCC GCTCCCAACA GCCCGAGTTT CTAGCCCCTT GAGTAAGGGG CAAGAAACCT AAGCCGCCTG GTGGATGGCT CGGCTCGGGG CGCTGACGAA GGACGCGTGG AC
J23SpTA_18       CCGCAGA AGACGGGTTT GATGGGGCGG GGATGTAAGC GGGAAGGGAA ACCGACCCGT TTAGTCTGCC GCTCCCAACA GCCCGAGTTT CTAGCCCCTT GAGTAAGGGG CAAGAAACCT AAGCCGCCTG GTGGATGGCT CGGCTCGGGG CGCTGACGAA GGACGCGTGG ACG
J23SpTA_11        CGCAGA AGACGGGTTT GATGGGGCGG GGATGTAAGC GGGAAGGGAA ACCGACCCGT TTAGTCTGCC GCTCCCAACA GCCCGAGTTT CTAGCCCCTT GAGTAAGGGG CAAGAAACCT AAGCCGCCTG GTGGATGGCT CGGCTCGGGG CGCTGACGAA GGACGCGTGG ACGCG
J23SpTA_9_        CGCAGA AGACGGGTTT GATGGGGCGG GGATGTAAGC GGGAAGGGAA ACCGACCCGT TTAGTCTGCC GCTCCCAACA GCCCGAGTTT CTAGCCCCTT GAGTAAGGGG CAAGAAACCT AAGCCGCCTG GTGGATGGCT CGGCTCGGGG CGCTGACGAA GGACGCGTGG ACGCGGG
J23SpTA_66         GCAGA AGACGGGTTT GATGGGGCGG GGATGTAAGC GGGAAGGGAA ACCGACCCGT TTAGTCTGCC GCTCCCAACA GCCCGAGTTT CTAGCCCCTT GAGTAAGGGG CAAGAAACCT AAGCCGCCTG GTGGATGGCT CGGCTCGGGG CGCTGACGAA GGACGCGTGG ACGCGTG
J23SpTA_25        CGCAGA AGACGGGTTT GATGGGGCGG GGATGTAAGC GGGAAGGGAA ACCGACCCGT TTAGTCTGCC GCTCCCAACA GCCCGAGTTT CTAGCCCCTT GAGTAAGGGG CAAGAAACCT AAGCCGCCTG GTGGATGGCT CGGCTCGGGG CGCTGACGAA GGACGCGTGG ACGCGT
J23SpTA_10      CCCGCAGA AGACGGGTTT GATGGGGCGG GGATGTAAGC GGGAAGGGAA ACCGACCCGT TTAGTCTGCC GCTCCCAACA GCCCGAGTTT CTAGCCCCTT GAGTAAGGGG CAAGAAACCT AAGCCGCCTG GTGGATGGCT CGGCTCGGGG CGCTGACGAA GGACGCGTGG AC
J23SpTA_5_      CTCGCAGA AGACGGGTTT GATGGGGCGG GGATGTAAGC GGGAAGGGAA ACCGACCCGT TTAGTCTGCC GCTCCCAACA GCCCGAGTTT CTAGCCCCTT GAGTAAGGGG CAAGAAACCT AAGCCGCCTG GTGGATGGCT CGGCTCGGGG CGCTGACGAA GGACGCGTGG ACG
J23SpTA_6_      CCCGCAGA AGACGGGTTT GATGGGGCGG GGATGTAAGC GGGAAGGGAA ACCGACCCGT TTAGTCTGCC GCTCCCAACA GCCCGAGTTT CTAGCCCCTT GAGTAAGGGG CAAGAAACCT AAGCCGCCTG GTGGATGGCT CGGCTCGGGG CGCTGACGAA GGACGCGTGG ACC
J23SpTA_5_       NCGCAGA AGACGGGTTT GATGGGGCGG GGATGTAAGC GGGAAGGGAA ACCGACCCGT TTAGTCTGCC GCTCCCAACA GCCCGAGTTT CTAGCCCCTT GAGTAAGGGG CAAGAAACCT AAGCCGCCTG GTGGATGGCT CGGCTCGGGG CGCTGACGAA GGACGCGTGG ACGCN
J23SpTA_29        CGCAGA AGACGGGTTT GATGGGGCGG GGATGTAAGC GGGAAGGGAA ACCGACCCGT TTAGTCTGCC GCTCCCAACA GCCCGAGTTT CTAGCCCCTT GAGTAAGGGG CAAGAAACCT AAGCCGCCTG GTGGATGGCT CGGCTCGGGG CGCTGACGAA GGACGCGTGG ACGCC
J23SpTA_6_       CCGCAGA AGACGGGTTT GATGGGGCGG GGATGTAAGC GGGAAGGGAA ACCGACCCGT TTAGTCTGCC GCTCCCAACA GCCCGAGTTT CTAGCCCCTT GAGTAAGGGG CAAGAAACCT AAGCCGCCTG GTGGATGGCT CGGCTCGGGG CGCTGACGAA GGACGCGTGG ACAC
J23SpTA_18      CCCGCAGA AGACGGGTTT GATGGGGCGG GGATGTAAGC GGGAAGGGAA ACCGACCCGT TTAGTCTGCC GCTCCCAACA GCCCGAGTTT CTAGCCCCTT GAGTAAGGGG CAAGAAACCT AAGCCGCCTG GTGGATGGCT CGGCTCGGGG CGCTGACGAA GGACGCGTGA G
J23SpTA_16        CGCAGA AGACGGGTTT GATGGGGCGG GGATGTAAGC GGGAAGGGAA ACCGACCCGT TTAGTCTGCC GCTCCCAACA GCCCGAGTTT CTAGCCCCTT GAGTAAGGGG CAAGAAACCT AAGCCGCCTG GTGGATGGCT CGGCTCGGGG CGCTGACGAA GGACGCGTGG ACGCA
J23SpTA_5_        CGCAGA AGACGGGTTT GATGGGGCGG GGATGTAAGC GGGAAGGGAA ACCGACCCGT TTAGTCTGCC GCTCCCAACA GCCCGAGTTT CTAGCCCCTT GAGTAAGGGG CAAGAAACCT AAGCCGCCTG GTGGATGGCT CGGCTCGGGG CGCTGACGAA GGACGCGTGG ACGCAT
J23SpTA_5_      NCCGCAGA AGACGGGTTT GATGGGGCGG GGATGTAAGC GGGAAGGGAA ACCGACCCGT TTAGTCTGCC GCTCCCAACA GCCCGAGTTT CTAGCCCCTT GAGTAAGGGG CAAGAAACCT AAGCCGCCTG GTGGATGGCT CGGCTCGGGG CGCTGACGAA GGACGCGTGG ACGN
J23SpTA_5_      CCCGCAGA AGACGGGTTT GATGGGGCGG GGATGTAAGC GGGAAGGGAA ACCGACCCGT TTAGTCTGCC GCTCCCAACA GCCCGAGTTT CTAGCCCCTT GAGTAAGGGG CAAGAAACCT AAGCCGCCTG GTGGATGGCT CGGCTCGGGG CGCTGACAAA GGG
J23SpTA_11        CGCAGA AGACGGGTTT GATGGGGCGG GGATGTAAGC GGGAAGGGAA ACCGACCCGT TTAGTCTGCC GCTCCCAACA GCCCGAGTTT CTAGCCCCTT GAGTAAGGGG CAAGAAACCT AAGCCGCCTG GTGGATGGCT CGGCTCGGGG CGCTGACGAA GGACGCGTGG ACGCT
J23SpTA_11      CCCGCAGA AGACGGGTTT GATGGGGCGG GGATGTAAGC GGGAAGGGAA ACCGACCCGT TTAGTCTGCC GCTCCCAACA GCCCGAGTTT CTAGCCCCTT GAGTAAGGGG CAAGAAACCT AAGCCGCCTG GTGGATGGCT CGGCTCGGGG CGCTGACGAA GGACGCGTG
J23SpTA_11       CCGCAGA AGACGGGTTT GATGGGGCGG GGATGTAAGC GGGAAGGGAA ACCGACCCGT TTAGTCTGCC GCTCCCAACA GCCCGAGTTT CTAGCCCCTT GAGTAAGGGG CAAGAAACCT AAGCCGCCTG GTGGATGGCT CGGCTCGGGG CGCTGACGAA GGACGCGTGG AC
J23SpTA_25         GCAGA AGACGGGTTT GATGGGGCGG GGATGTAAGC GGGAAGGGAA ACCGACCCGT TTAGTCTGCC GCTCCCAACA GCCCGAGTTT CTAGCCCCTT GAGTAAGGGG CAAGAAACCT AAGCCGCCTG GTGGATGGCT CGGCTCGGGG CGCTGACGAA GGACGCGTGG ACGC
J23SpTA_15         GCAGA AGACGGGTTT GATGGGGCGG GGATGTAAGC GGGAAGGGAA ACCGACCCGT TTAGTCTGCC GCTCCCAACA GCCCGAGTTT CTAGCCCCTT GAGTAAGGGG CAAGAAACCT AAGCCGCCTG GTGGATGGCT CGGCTCGGGG CGCTGACGAA GGACGCGTGG ACGCG
J23SpTA_6_      CCCGCAGA AGACGGGTTT GATGGGGCGG GGATGTAAGC GGGAAGGGAA ACCGACCCGT TTAGTCTGCC GCTCCCAACA GCCCGAGTTT CTAGCCCCTT GAGTAAGGGG CAAGAAACCT AAGCCGCCTG GTGGATGGCT CGGCTCGGGG CGCTGACGAA GGACGCGTGC
J23SpTA_10        CGCAGA AGACGGGTTT GATGGGGCGG GGATGTAAGC GGGAAGGGAA ACCGACCCGT TTAGTCTGCC GCTCCCAACA GCCCGAGTTT CTAGCCCCTT GAGTAAGGGG CAAGAAACCT AAGCCGCCTG GTGGATGGCT CGGCTCGGGG CGCTGACGAA GGACGCGTGG ACGG
J23SpTA_5_        CGCAGA AGACGGGTTT GATGGGGCGG GGATGTAAGC GGGAAGGGAA ACCGACCCGT TTAGTCTGCC GCTCCCAACA GCCCGAGTTT CTAGCCCCTT GAGTAAGGGG CAAGAAACCT AAGCCGCCTG GTGGATGGCT CGGCTCGGGG CGCTGACGAA GGACGCGTGG ACGGG
J23SpTA_9_        CGCAGA AGACGGGTTT GATGGGGCGG GGATGTAAGC GGGAAGGGAA ACCGACCCGT TTAGTCTGCC GCTCCCAACA GCCCGAGTTT CTAGCCCCTT GAGTAAGGGG CAAGAAACCT AAGCCGCCTG GTGGATGGCT CGGCTCGGGG CGCTGACGAA GGACGCGTGG ACGAC
J23SpTA_6_        CGCAGA AGACGGGTTT GATGGGGCGG GGATGTAAGC GGGAAGGGAA ACCGACCCGT TTAGTCTGCC GCTCCCAACA GCCCGAGTTT CTAGCCCCTT GAGTAAGGGG CAAGAAACCT AAGCCGCCTG GTGGATGGCT CGGCTCGGGG CGCTGACGAA GGACGCGTGG ACGA
J23SpTA_9_        NGCAGA AGACGGGTTT GATGGGGCGG GGATGTAAGC GGGAAGGGAA ACCGACCCGT TTAGTCTGCC GCTCCCAACA GCCCGAGTTT CTAGCCCCTT GAGTAAGGGG CAAGAAACCT AAGCCGCCTG GTGGATGGCT CGGCTCGGGG CGCTGACGAA GGACGCGTGG ACGCN
J23SpTA_6_      CCCGCAGA AGACGGGTTT GATGGGGCGG GGATGTAAGC GGGAAGGGAA ACCGACCCGT TTAGTCTGCC GCTCCCAACA GCCCGAGTTT CTAGCCCCTT GAGTAAGGGG CAAGAAACCT AAGCCGCCTG GTGGATGGCT CGGCTCGGGG CGCTGACAAA GG
J23SpTA_6_       CCCCAGA AGACGGGTTT GATGGGGCGG GGATGTAAGC GGGAAGGGAA ACCGACCCGT TTAGTCTGCC GCTCCCAACA GCCCGAGTTT CTAGCCCCTT GAGTAAGGGG CAAGAAACCT AAGCCGCCTG GTGGATGGCT CGGCTCGGGG CGCTGACGAA GGACGCGTGG ACG
J23SpTA_5_       CCGCAGA AGACGGGTTT GATGGGGCGG GGATGTAAGC GGGAAGGGAA ACCGACCCGT TTAGTCTGCC GCTCCCAACA GCCCGAGTTT CTAGCCCCTT GAGTAAGGGG CAAGAAACCT AAGCCGCCTG GTGGATGGCT CGGCTCGGGG CGCTGACGAG GGG
J23SpTA_5_        CGCAGA AGGCGGGTTT GATGGGGCGG GGATGTAAGC GGGAAGGGAA ACCGACCCGT TTAGTCTGCC GCTCCCAACA GCCCGAGTTT CTAGCCCCTT GAGTAAGGGG CAAGAAACCT AAGCCGCCTG GTGGATGGCT CGGCTCGGGG CGCTGACGAA GGACGCGTGG ACGC
J23SpTA_5_      CCGGCAGA AGACGGGTTT GATGGGGCGG GGATGTAAGC GGGAAGGGAA ACCGACCCGT TTAGTCTGCC GCTCCCAACA GCCCGAGTTT CTAGCCCCTT GAGTAAGGGG CAAGAAACCT AAGCCGCCTG GTGGATGGCT CGGCTCGGGG CGCTGACGAA GGACGCGTGG AC
J23SpTA_30        CGCAGA AGACGGGTTT GATGGGGCGG GGATGTAAGC GGGAAGGGAA ACCGACCCGT TTAGTCTGCC GCTCCCAACA GCCCGAGTTT CTAGCCCCTT GAGTAAGGGG CAAGAAACCT AAGCCGCCTG GTGGATGGCT CGGCTCGGGG CGCTGACGAA GGACGCGTGG ACAC
J23SpTA_6_        CCCAGA AGACGGGTTT GATGGGGCGG GGATGTAAGC GGGAAGGGAA ACCGACCCGT TTAGTCTGCC GCTCCCAACA GCCCGAGTTT CTAGCCCCTT GAGTAAGGGG CAAGAAACCT AAGCCGCCTG GTGGATGGCT CGGCTCGGGG CGCTGACGAA GGACGCGTGG ACGC
J23SpTA_6_        GGCAGA AGACGGGTTT GATGGGGCGG GGATGTAAGC GGGAAGGGAA ACCGACCCGT TTAGTCTGCC GCTCCCAACA GCCCGAGTTT CTAGCCCCTT GAGTAAGGGG CAAGAAACCT AAGCCGCCTG GTGGATGGCT CGGCTCGGGG CGCTGACGAA GGACGCGTGG ACGC
J23SpTA_7_        CGCAGA AGACGGGTTT GATGGGGCGG GGATGTAAGC GGGAAGGGAA ACCGACCCGT TTAGTCTGCC GCTCCCAACA GCCCGAGTTT CTAGCCCCTT GAGTAAGGGG CAAGAAACCT AAGCCGCCTG GTGGATGGCT CGGCTCGGGG CGCTGACGAA GGACGCGTGA GCG
J23SpTA_10      NCCGCAGA AGACGGGTTT GATGGGGCGG GGATGTAAGC GGGAAGGGAA ACCGACCCGT TTAGTCTGCC GCTCCCAACA GCCCGAGTTT CTAGCCCCTT GAGTAAGGGG CAAGAAACCT AAGCCGCCTG GTGGATGGCT CGGCTCGGGG CGCTGACGAA GGACGCGTGG ACN
J23SpTA_5_        CGCAGA AGACGGGTTT GATGGGGCGG GGATGTAAGC GGGAAGGGAA ACCGACCCGT TTAGTCTGCC GCTCCCAACA GCCCGAGTTT CTAGCCCCTT GAGTAAGGGG CAAGAAACCT AAGCCGCCTG GTGGATGGCT CGGCTCGGGG CGCTGACGAA GGACGCGTGG ACGT
J23SpTA_14    CGCCCGCAGA AGACGGGTTT GATGGGGCGG GGATGTAAGC GGGAAGGGAA ACCGACCCGT TTAGTCTGCC GCTCCCAACA GCCCGAGTTT CTAGCCCCTT GAGTAAGGGG CAAGAAACCT AAGCCGCCTG GTGGATGGCT CGGCTCGGGG CGCTGACGAA GGACGCGTG
J23SpTA_9_     GCCCGCAGA AGACGGGTTT GATGGGGCGG GGATGTAAGC GGGAAGGGAA ACCGACCCGT TTAGTCTGCC GCTCCCAACA GCCCGAGTTT CTAGCCCCTT GAGTAAGGGG CAAGAAACCT AAGCCGCCTG GTGGATGGCT CGGCTCGGGG CGCTGACGAA GGACGCGTG
J23SpTA_6_        CGCAGA AGACGGGTTT GATGGGGCGG GGATGTAAGC GGGAAGGGAA ACCGACCCGT TTAGTCTGCC GCTCCCAACA GCCCGAGTTT CTAGCCCCTT GAGTAAGGGG CAAGAAACCT AAGCCGCCTG GTGGATGGCT CGGCTCGGGG CGCTGACGAA AGGCG
J23SpTA_6_        CGCAGA AGACGGGTTT GATGGGGCGG GGATGTAAGC GGGAAGGGAA ACCGACCCGT TTAGTCTGCC GCTCCCAACA GCCCGAGTTT CTAGCCCCTT GAGTAAGGGG CAAGAAACCT AAGCCGCCTG GTGGATGGCT CGGCTCGGGG CGCTGACGAA AGGC
J23SpTA_10       CCGCAGA AGACGGGTTT GATGGGGCGG GGATGTAAGC GGGAAGGGAA ACCGACCCGT TTAGTCTGCC GCTCCCAACA GCCCGAGTTT CTAGCCCCTT GAGTAAGGGG CAAGAAACCT AAGCCGCCTG GTGGATGGCT CGGCTCGGGG CGCTGACAAA GGG
J23SpTA_13       CCGCAGA AGACGGGTTT GATGGGGCGG GGATGTAAGC GGGAAGGGAA ACCGACCCGT TTAGTCTGCC GCTCCCAACA GCCCGAGTTT CTAGCCCCTT GAGTAAGGGG CAAGAAACCT AAGCCGCCTG GTGGATGGCT CGGCTCGGGG CGCTGACGAA GGACGCGTGA G
J23SpTA_6_        CGCAGA AGACGGGTTT GATGGGGCGG GGATGTAAGC GGGAAGGGAA ACCGACCCGT TTAGTCTGCC GCTCCCAACA GCCCGAGTTT CTAGCCCCTT GAGTAAGGGG CAAGAAACCT AAGCCGCCTG GTGGATGGCT CGGCTCGGGG CGCTGACGAA GGACGCGTGG ACGN
J23SpTA_25        CGCAGA AGACGGGTTT GATGGGGCGG GGATGTAAGC GGGAAGGGAA ACCGACCCGT TTAGTCTGCC GCTCCCAACA GCCCGAGTTT CTAGCCCCTT GAGTAAGGGG CAAGAAACCT AAGCCGCCTG GTGGATGGCT CGGCTCGGGG CGCTGACGAA GGACGCGTGG AC
J23SpTA_12       CCGCAGA AGACGGGTTT GATGGGGCGG GGATGTAAGC GGGAAGGGAA ACCGACCCGT TTAGTCTGCC GCTCCCAACA GCCCGAGTTT CTAGCCCCTT GAGTAAGGGG CAAGAAACCT AAGCCGCCTG GTGGATGGCT CGGCTCGGGG CGCTGACGAA GGACGCGTG
J23SpTA_77          CAGA AGACGGGTTT GATGGGGCGG GGATGTAAGC GGGAAGGGAA ACCGACCCGT TTAGTCTGCC GCTCCCAACA GCCCGAGTTT CTAGCCCCTT GAGTAAGGGG CAAGAAACCT AAGCCGCCTG GTGGATGGCT CGGCTCGGGG CGCTGACGAA GGACGCGTGG ACGC
J23SpTA_49          CAGA AGACGGGTTT GATGGGGCGG GGATGTAAGC GGGAAGGGAA ACCGACCCGT TTAGTCTGCC GCTCCCAACA GCCCGAGTTT CTAGCCCCTT GAGTAAGGGG CAAGAAACCT AAGCCGCCTG GTGGATGGCT CGGCTCGGGG CGCTGACGAA GGACGCGTGG ACGCG
J23SpTA_31          CAGA AGACGGGTTT GATGGGGCGG GGATGTAAGC GGGAAGGGAA ACCGACCCGT TTAGTCTGCC GCTCCCAACA GCCCGAGTTT CTAGCCCCTT GAGTAAGGGG CAAGAAACCT AAGCCGCCTG GTGGATGGCT CGGCTCGGGG CGCTGACGAA GGACGCGTGG ACGCGTGG
J23SpTA_18          CAGA AGACGGGTTT GATGGGGCGG GGATGTAAGC GGGAAGGGAA ACCGACCCGT TTAGTCTGCC GCTCCCAACA GCCCGAGTTT CTAGCCCCTT GAGTAAGGGG CAAGAAACCT AAGCCGCCTG GTGGATGGCT CGGCTCGGGG CGCTGACGAA GGACGCGTGG ACGCGTG
J23SpTA_13           AGA AGACGGGTTT GATGGGGCGG GGATGTAAGC GGGAAGGGAA ACCGACCCGT TTAGTCTGCC GCTCCCAACA GCCCGAGTTT CTAGCCCCTT GAGTAAGGGG CAAGAAACCT AAGCCGCCTG GTGGATGGCT CGGCTCGGGG CGCTGACGAA GGACGCGTGG ACGCGTGG
J23SpTA_17           AGA AGACGGGTTT GATGGGGCGG GGATGTAAGC GGGAAGGGAA ACCGACCCGT TTAGTCTGCC GCTCCCAACA GCCCGAGTTT CTAGCCCCTT GAGTAAGGGG CAAGAAACCT AAGCCGCCTG GTGGATGGCT CGGCTCGGGG CGCTGACGAA GGACGCGTGG ACGCGTGGC
J23SpTA_6_          CAGA AGACGGGTTT GATGGGGCGG GGATGTAAGC GGGAAGGGAA ACCGACCCGT TTAGTCTGCC GCTCCCAACA GCCCGAGTTT CTAGCCCCTT GAGTAAGGGG CAAGAAACCT AAGCCGCCTG GTGGATGGCT CGGCTCGGGG CGCTGACGAA GGACGCGTGG ACGCGGGG
J23SpTA_11          CAGA AGACGGGTTT GATGGGGCGG GGATGTAAGC GGGAAGGGAA ACCGACCCGT TTAGTCTGCC GCTCCCAACA GCCCGAGTTT CTAGCCCCTT GAGTAAGGGG CAAGAAACCT AAGCCGCCTG GTGGATGGCT CGGCTCGGGG CGCTGACGAA GGACGCGTGG ACGCGTGAC
J23SpTA_10           AGA AGACGGGTTT GATGGGGCGG GGATGTAAGC GGGAAGGGAA ACCGACCCGT TTAGTCTGCC GCTCCCAACA GCCCGAGTTT CTAGCCCCTT GAGTAAGGGG CAAGAAACCT AAGCCGCCTG GTGGATGGCT CGGCTCGGGG CGCTGACGAA GGACGCGTGG ACGCGTG
J23SpTA_18            GA AGACGGGTTT GATGGGGCGG GGATGTAAGC GGGAAGGGAA ACCGACCCGT TTAGTCTGCC GCTCCCAACA GCCCGAGTTT CTAGCCCCTT GAGTAAGGGG CAAGAAACCT AAGCCGCCTG GTGGATGGCT CGGCTCGGGG CGCTGACGAA GGACGCGTGG ACGCGTGG
J23SpTA_9_           AGA AGACGGGTTT GATGGGGCGG GGATGTAAGC GGGAAGGGAA ACCGACCCGT TTAGTCTGCC GCTCCCAACA GCCCGAGTTT CTAGCCCCTT GAGTAAGGGG CAAGAAACCT AAGCCGCCTG GTGGATGGCT CGGCTCGGGG CGCTGACGAA GGACGCGTGG ACGCGTAG
J23SpTA_6_             A AGACGGGTTT GATGGGGCGG GGATGTAAGC GGGAAGGGAA ACCGACCCGT TTAGTCTGCC GCTCCCAACA GCCCGAGTTT CTAGCCCCTT GAGTAAGGGG CAAGAAACCT AAGCCGCCTG GTGGATGGCT CGGCTCGGGG CGCTGACGAA GGACGCGTGG ACGCGTGGC
J23SpTA_24             A AGACGGGTTT GATGGGGCGG GGATGTAAGC GGGAAGGGAA ACCGACCCGT TTAGTCTGCC GCTCCCAACA GCCCGAGTTT CTAGCCCCTT GAGTAAGGGG CAAGAAACCT AAGCCGCCTG GTGGATGGCT CGGCTCGGGG CGCTGACGAA GGACGCGTGG ACGCGTGG
J23SpTA_12            GA AGACGGGTTT GATGGGGCGG GGATGTAAGC GGGAAGGGAA ACCGACCCGT TTAGTCTGCC GCTCCCAACA GCCCGAGTTT CTAGCCCCTT GAGTAAGGGG CAAGAAACCT AAGCCGCCTG GTGGATGGCT CGGCTCGGGG CGCTGACGAA GGACGCGTGG ACGCGTG
J23SpTA_45         GCAGA AGACGGGTTT GATGGGGCGG GGATGTAAGC GGGAAGGGAA ACCGACCCGT TTAGTCTGCC GCTCCCAACA GCCCGAGTTT CTAGCCCCTT GAGTAAGGGG CAAGAAACCT AAGCCGCCTG GTGGATGGCT CGGCTCGGGG CGCTGACGAA GGACGCGTGG ACG
J23SpTA_5_        CGCAGA AGACGGGTTT GATGGGGCGG GGATGTAAGC GGGAAGGGAA ACCGACCCGT TTAGTCTGCC GCTCCCAACA GCCCGAGTTT CTAGCCCCTT GAGTAAGGGG CAAGAAACCT AAGCCGCCTG GTGGATGGCT CGGCTCGGGG CGCTGACGAA AGG
J23SpTA_8_         GCAGA AGACGGGTTT GATGGGGCGG GGATGTAAGC GGGAAGGGAA ACCGACCCGT TTAGTCTGCC GCTCCCAACA GCCCGAGTTT CTAGCCCCTT GAGTAAGGGG CAAGAAACCT AAGCCGCCTG GTGGATGGCT CGGCTCGGGG CGCTGACGAA GGACGCGTGG ACGG
J23SpTA_9_        NGCAGA AGACGGGTTT GATGGGGCGG GGATGTAAGC GGGAAGGGAA ACCGACCCGT TTAGTCTGCC GCTCCCAACA GCCCGAGTTT CTAGCCCCTT GAGTAAGGGG CAAGAAACCT AAGCCGCCTG GTGGATGGCT CGGCTCGGGG CGCTGACGAA GGACGCGTGG ACGN
J23SpTA_40        CGCAGA AGACGGGTTT GATGGGGCGG GGATGTAAGC GGGAAGGGAA ACCGACCCGT TTAGTCTGCC GCTCCCAACA GCCCGAGTTT CTAGCCCCTT GAGTAAGGGG CAAGAAACCT AAGCCGCCTG GTGGATGGCT CGGCTCGGGG CGCTGACGAA GGACGCGTGA G
J23SpTA_11        CGCAGA AGACGGGTTT GATGGGGCGG GGATGTAAGC GGGAAGGGAA ACCGACCCGT TTAGTCTGCC GCTCCCAACA GCCCGAGTTT CTAGCCCCTT GAGTAAGGGG CAAGAAACCT AAGCCGCCTG GTGGATGGCT CGGCTCGGGG CGCTGACGAG GGG
J23SpTA_29        CGCAGA AGACGGGTTT GATGGGGCGG GGATGTAAGC GGGAAGGGAA ACCGACCCGT TTAGTCTGCC GCTCCCAACA GCCCGAGTTT CTAGCCCCTT GAGTAAGGGG CAAGAAACCT AAGCCGCCTG GTGGATGGCT CGGCTCGGGG CGCTGACAAA GGG
J23SpTA_6_        CGTAGA AGACGGGTTT GATGGGGCGG GGATGTAAGC GGGAAGGGAA ACCGACCCGT TTAGTCTGCC GCTCCCAACA GCCCGAGTTT CTAGCCCCTT GAGTAAGGGG CAAGAAACCT AAGCCGCCTG GTGGATGGCT CGGCTCGGGG CGCTGACGAA GGACGCGTGG ACG
J23SpTA_18        CGCAGA AGACGGGTTT GATGGGGCGG GGATGTAAGC GGGAAGGGAA ACCGACCCGT TTAGTCTGCC GCTCCCAACA GCCCGAGTTT CTAGCCCCTT GAGTAAGGGG CAAGAAACCT AAGCCGCCTG GTGGATGGCT CGGCTCGGGG CGCTGACGAA GGACGCGTGG ACC
J23SpTA_9_        GGCAGA AGACGGGTTT GATGGGGCGG GGATGTAAGC GGGAAGGGAA ACCGACCCGT TTAGTCTGCC GCTCCCAACA GCCCGAGTTT CTAGCCCCTT GAGTAAGGGG CAAGAAACCT AAGCCGCCTG GTGGATGGCT CGGCTCGGGG CGCTGACGAA GGACGCGTGG ACG
J23SpTA_17          CAGA AGACGGGTTT GATGGGGCGG GGATGTAAGC GGGAAGGGAA ACCGACCCGT TTAGTCTGCC GCTCCCAACA GCCCGAGTTT CTAGCCCCTT GAGTAAGGGG CAAGAAACCT AAGCCGCCTG GTGGATGGCT CGGCTCGGGG CGCTGACGAA GGACGCGTGG ACGCC
J23SpTA_7_          CAGA AGACGGGTTT GATGGGGCGG GGATGTAAGC GGGAAGGGAA ACCGACCCGT TTAGTCTGCC GCTCCCAACA GCCCGAGTTT CTAGCCCCTT GAGTAAGGGG CAAGAAACCT AAGCCGCCTG GTGGATGGCT CGGCTCGGGG CGCTGACGAA GGACGCGTGG ACGCT
J23SpTA_6_        CGCAGA AGACGGGTTT GATGGGGCGG GGATGTAAGC GGGAAGGGAA ACCGACCCGT TTAGTCTGCC GCTCCCAACA GCCCGAGTTT CTAGCCCCTT GAGTAAGGGG CAAGAAACCT AAGCTGCCTG GTGGATGGCT CGGCTCGGGG CGCTGACGAA GGG
J23SpTA_8_        CGCAGA AGACGGGTTT GATGGGGCGG GGATGTAAGC GGGAAGGGAA ACCGACCCGT TTAGTCTGCC GCTCCCAACA GCCCGAGTTT CTAGCCCCTT GAGTAAGGGG CAAGAAACCT AAGCCGCCTG GTGGATGGCT CGGCTCGGGG CGCTGACGAA GGACGCGTGG ACN
J23SpTA_6_        CCCAGA AGACGGGTTT GATGGGGCGG GGATGTAAGC GGGAAGGGAA ACCGACCCGT TTAGTCTGCC GCTCCCAACA GCCCGAGTTT CTAGCCCCTT GAGTAAGGGG CAAGAAACCT AAGCCGCCTG GTGGATGGCT CGGCTCGGGG CGCTGACGAA GGACGCGTGG ACG
J23SpTA_29        CGCAGA AGACGGGTTT GATGGGGCGG GGATGTAAGC GGGAAGGGAA ACCGACCCGT TTAGTCTGCC GCTCCCAACA GCCCGAGTTT CTAGCCCCTT GAGTAAGGGG CAAGAAACCT AAGCCGCCTG GTGGATGGCT CGGCTCGGGG CGCTGACGAA GGACGCGTG
J23SpTA_15          CAGA AGACGGGTTT GATGGGGCGG GGATGTAAGC GGGAAGGGAA ACCGACCCGT TTAGTCTGCC GCTCCCAACA GCCCGAGTTT CTAGCCCCTT GAGTAAGGGG CAAGAAACCT AAGCCGCCTG GTGGATGGCT CGGCTCGGGG CGCTGACGAA GGACGCGTGG ACG
J23SpTA_5_        CGCAGA AGACGGGTTT GATGGGGCGG GGATGTAAGC GGGAAGGGAA ACCGACCCGT TTAGTCTGCC GCTCCCAACA GCCCGAGTTT CTAGCCCCTT GAGTAAGGGG CAAGAAACCT AAGCCGCCTG GTGGATGGCT CGGCTCGGGG CGCTGACGAC GG
J23SpTA_9_         GCAGA AGACGGGTTT GATGGGGCGG GGATGTAAGC GGGAAGGGAA ACCGACCCGT TTAGTCTGCC GCTCCCAACA GCCCGAGTTT CTAGCCCCTT GAGTAAGGGG CAAGAAACCT AAGCCGCCTG GTGGATGGCT CGGCTCGGGG CGCTGACGAA GGACGCGTGA G
J23SpTA_10        CGCAGA AGACGGGTTT GATGGGGCGG GGATGTAAGC GGGAAGGGAA ACCGACCCGT TTAGTCTGCC GCTCCCAACA GCCCGAGTTT CTAGCCCCTT GAGTAAGGGG CAAGAAACCT AAGCCGCCTG GTGGATGGCT CGGCTCGGGG CGCTGACGGA GG
J23SpTA_6_       CCGCAGA AGACGGGTTT GATGGGGCGG GGATGTAAGC GGGAAGGGAA ACCGACCCGT TTAGTCTGCC GCTCCCAACA GCCCGAGTTT CTAGCCCCTT GAGTAAGGGG CAAGAAACCT AAGCCGCCTG GTGGATGGCT CGGCTCGGGG CGCTGACGGA GG
J23SpTA_93      CCCGCAGA AGACGGGTTT GATGGGGCGG GGATGTAAGC GGGAAGGGAA ACCGACCCGT TTAGTCTGCC GCTCCCAACA GCCCGAGTTT CTAGCCCCTT GAGTAAGGGG CAAGAAACCT AAGCCGCCTG GTGGATGGCT CGGCTCGGGG CGCTGACGAA GGACGCGTGG ACGC
J23SpTA_51      CCCGCAGA AGACGGGTTT GATGGGGCGG GGATGTAAGC GGGAAGGGAA ACCGACCCGT TTAGTCTGCC GCTCCCAACA GCCCGAGTTT CTAGCCCCTT GAGTAAGGGG CAAGAAACCT AAGCCGCCTG GTGGATGGCT CGGCTCGGGG CGCTGACGAA GGACGCGTGG ACGCG
J23SpTA_91        CGCAGA AGACGGGTTT GATGGGGCGG GGATGTAAGC GGGAAGGGAA ACCGACCCGT TTAGTCTGCC GCTCCCAACA GCCCGAGTTT CTAGCCCCTT GAGTAAGGGG CAAGAAACCT AAGCCGCCTG GTGGATGGCT CGGCTCGGGG CGCTGACGAA GGACGCGTGG ACGCGTGG
J23SpTA_38       CCGCAGA AGACGGGTTT GATGGGGCGG GGATGTAAGC GGGAAGGGAA ACCGACCCGT TTAGTCTGCC GCTCCCAACA GCCCGAGTTT CTAGCCCCTT GAGTAAGGGG CAAGAAACCT AAGCCGCCTG GTGGATGGCT CGGCTCGGGG CGCTGACGAA GGACGCGTGG ACGCGTGG
J23SpTA_34      CCCGCAGA AGACGGGTTT GATGGGGCGG GGATGTAAGC GGGAAGGGAA ACCGACCCGT TTAGTCTGCC GCTCCCAACA GCCCGAGTTT CTAGCCCCTT GAGTAAGGGG CAAGAAACCT AAGCCGCCTG GTGGATGGCT CGGCTCGGGG CGCTGACGAA GGACGCGTGG ACGCGTGG
J23SpTA_86      CCCGCAGA AGACGGGTTT GATGGGGCGG GGATGTAAGC GGGAAGGGAA ACCGACCCGT TTAGTCTGCC GCTCCCAACA GCCCGAGTTT CTAGCCCCTT GAGTAAGGGG CAAGAAACCT AAGCCGCCTG GTGGATGGCT CGGCTCGGGG CGCTGACGAA GGACGCGTGG ACGCGTGGC
J23SpTA_11      CCCGCAGA AGACGGGTTT GATGGGGCGG GGATGTAAGC GGGAAGGGAA ACCGACCCGT TTAGTCTGCC GCTCCCAACA GCCCGAGTTT CTAGCCCCTT GAGTAAGGGG CAAGAAACCT AAGCCGCCTG GTGGATGGCT CGGCTCGGGG CGCTGACGAA GGACGCGTGG ACGCGTGGCG
J23SpTA_9_       CCGCAGA AGACGGGTTT GATGGGGCGG GGATGTAAGC GGGAAGGGAA ACCGACCCGT TTAGTCTGCC GCTCCCAACA GCCCGAGTTT CTAGCCCCTT GAGTAAGGGG CAAGAAACCT AAGCCGCCTG GTGGATGGCT CGGCTCGGGG CGCTGACGAA GGACGCGTGG ACGCGTGGCG
J23SpTA_11       CCGCAGA AGACGGGTTT GATGGGGCGG GGATGTAAGC GGGAAGGGAA ACCGACCCGT TTAGTCTGCC GCTCCCAACA GCCCGAGTTT CTAGCCCCTT GAGTAAGGGG CAAGAAACCT AAGCCGCCTG GTGGATGGCT CGGCTCGGGG CGCTGACGAA GGACGCGTGG ACGCGTGGC
J23SpTA_23        CGCAGA AGACGGGTTT GATGGGGCGG GGATGTAAGC GGGAAGGGAA ACCGACCCGT TTAGTCTGCC GCTCCCAACA GCCCGAGTTT CTAGCCCCTT GAGTAAGGGG CAAGAAACCT AAGCCGCCTG GTGGATGGCT CGGCTCGGGG CGCTGACGAA GGACGCGTGG ACGCGTGGCG
J23SpTA_6_     GCCCGCAGA AGACGGGTTT GATGGGGCGG GGATGTAAGC GGGAAGGGAA ACCGACCCGT TTAGTCTGCC GCTCCCAACA GCCCGAGTTT CTAGCCCCTT GAGTAAGGGG CAAGAAACCT AAGCCGCCTG GTGGATGGCT CGGCTCGGGG CGCTGACGAA GGACGCGTGG ACGCGTGG
J23SpTA_6_    CGCCCGCAGA AGACGGGTTT GATGGGGCGG GGATGTAAGC GGGAAGGGAA ACCGACCCGT TTAGTCTGCC GCTCCCAACA GCCCGAGTTT CTAGCCCCTT GAGTAAGGGG CAAGAAACCT AAGCCGCCTG GTGGATGGCT CGGCTCGGGG CGCTGACGAA GGACGCGTGG ACGCGTGG
J23SpTA_6_    CGCCCGCAGA AGACGGGTTT GATGGGGCGG GGATGTAAGC GGGAAGGGAA ACCGACCCGT TTAGTCTGCC GCTCCCAACA GCCCGAGTTT CTAGCCCCTT GAGTAAGGGG CAAGAAACCT AAGCCGCCTG GTGGATGGCT CGGCTCGGGG CGCTGACGAA GGACGCGTGG ACGCGTG
J23SpTA_10       CCGCAGA AGACGGGTTT GATGGGGCGG GGATGTAAGC GGGAAGGGAA ACCGACCCGT TTAGTCTGCC GCTCCCAACA GCCCGAGTTT CTAGCCCCTT GAGTAAGGGG CAAGAAACCT AAGCCGCCTG GTGGATGGCT CGGCTCGGGG CGCTGACGAA GGACGCGTGG ACGCGTGGCA
J23SpTA_20        CGCAGA AGACGGGTTT GATGGGGCGG GGATGTAAGC GGGAAGGGAA ACCGACCCGT TTAGTCTGCC GCTCCCAACA GCCCGAGTTT CTAGCCCCTT GAGTAAGGGG CAAGAAACCT AAGCCGCCTG GTGGATGGCT CGGCTCGGGG CGCTGACGAA GGACGCGTGG ACGCGTGGC
J23SpTA_20      CCCGCAGA AGACGGGTTT GATGGGGCGG GGATGTAAGC GGGAAGGGAA ACCGACCCGT TTAGTCTGCC GCTCCCAACA GCCCGAGTTT CTAGCCCCTT GAGTAAGGGG CAAGAAACCT AAGCCGCCTG GTGGATGGCT CGGCTCGGGG CGCTGACGAA GGACGCGTGG ACGCGTG
J23SpTA_83        CGCAGA AGACGGGTTT GATGGGGCGG GGATGTAAGC GGGAAGGGAA ACCGACCCGT TTAGTCTGCC GCTCCCAACA GCCCGAGTTT CTAGCCCCTT GAGTAAGGGG CAAGAAACCT AAGCCGCCTG GTGGATGGCT CGGCTCGGGG CGCTGACGAA GGACGCGTGG ACGCGTGGCA AG
J23SpTA_36       CCGCAGA AGACGGGTTT GATGGGGCGG GGATGTAAGC GGGAAGGGAA ACCGACCCGT TTAGTCTGCC GCTCCCAACA GCCCGAGTTT CTAGCCCCTT GAGTAAGGGG CAAGAAACCT AAGCCGCCTG GTGGATGGCT CGGCTCGGGG CGCTGACGAA GGACGCGTGG ACGCGTGGCA AG
J23SpTA_12        CGCAGA AGACGGGTTT GATGGGGCGG GGATGTAAGC GGGAAGGGAA ACCGACCCGT TTAGTCTGCC GCTCCCAACA GCCCGAGTTT CTAGCCCCTT GAGTAAGGGG CAAGAAACCT AAGCCGCCTG GTGGATGGCT CGGCTCGGGG CGCTGACGAA GGACGCGTGG ACGCGTGGCA AGC
J23SpTA_5_        CGCAGA AGACGGGTTT GATGGGGCGG GGATGTAAGC GGGAAGGGAA ACCGACCCGT TTAGTCTGCC GCTCCCAACA GCCCGAGTTT CTAGCCCCTT GAGTAAGGGG CAAGAAACCT AAGCCGCCTG GTGGATGGCT CGGCTCGGGG CGCTGACGAA GGACGCGTGG ACGCGTGGCA AGCG
J23SpTA_17          CAGA AGACGGGTTT GATGGGGCGG GGATGTAAGC GGGAAGGGAA ACCGACCCGT TTAGTCTGCC GCTCCCAACA GCCCGAGTTT CTAGCCCCTT GAGTAAGGGG CAAGAAACCT AAGCCGCCTG GTGGATGGCT CGGCTCGGGG CGCTGACGAA GGACGCGTGG ACGCGTGGCA AGCTG
J23SpTA_5_          CAGA AGACGGGTTT GATGGGGCGG GGATGTAAGC GGGAAGGGAA ACCGACCCGT TTAGTCTGCC GCTCCCAACA GCCCGAGTTT CTAGCCCCTT GAGTAAGGGG CAAGAAACCT AAGCCGCCTG GTGGATGGCT CGGCTCGGGG CGCTGATGAA GGGCGTGGCA AGCTGCT
J23SpTA_5_          CAGA AGACGGGTTT GATGGGGCGG GGATGTAAGC GGGAAGGGAA ACCGACCCGT TTAGTCTGCC GCTCCCAACA GCCCGAGTTT CTAGCCCCTT GAGTAAGGGG CAAGAAACCT AAGCCGCCTG GTGGATGGCT CGGCTCGGGG CGCTGACGAA GGACGCGTGG ACGCGTGGCA AGCT
J23SpTA_10         GCAGA AGACGGGTTT GATGGGGCGG GGATGTAAGC GGGAAGGGAA ACCGACCCGT TTAGTCTGCC GCTCCCAACA GCCCGAGTTT CTAGCCCCTT GAGTAAGGGG CAAGAAACCT AAGCCGCCTG GTGGATGGCT CGGCTCGGGG CGCTGACGAA GGACGCGTGG ACGCGTGGCA AG
J23SpTA_38          CAGA AGACGGGTTT GATGGGGCGG GGATGTAAGC GGGAAGGGAA ACCGACCCGT TTAGTCTGCC GCTCCCAACA GCCCGAGTTT CTAGCCCCTT GAGTAAGGGG CAAGAAACCT AAGCCGCCTG GTGGATGGCT CGGCTCGGGG CGCTGACGAA GGACGCGTGG ACGCGTGGCA AG
J23SpTA_5_       CCGCAGA AGACGGGTTT GATGGGGCGG GGATGTAAGC GGGAAGGGAA ACCGACCCGT TTAGTCTGCC GCTCCCAACA GCCCGAGTTT CTAGCCCCTT GAGTAAGGGG CAAGAAACCT AAGCCGCCTG GTGGATGGCT CGGCTCGGGG CGCTGACGAA GGACGCGTGG ACGCGTGAC
J23SpTA_10        CGCAGA AGACGGGTTT GATGGGGCGG GGATGTAAGC GGGAAGGGAA ACCGACCCGT TTAGTCTGCC GCTCCCAACA GCCCGAGTTT CTAGCCCCTT GAGTAAGGGG CAAGAAACCT AAGCCGCCTG GTGGATGGCT CGGCTCGGGG CGCTGACGAA GGACGCGTGG ACGCGTGGCA
J23SpTA_23       CCGCAGA AGACGGGTTT GATGGGGCGG GGATGTAAGC GGGAAGGGAA ACCGACCCGT TTAGTCTGCC GCTCCCAACA GCCCGAGTTT CTAGCCCCTT GAGTAAGGGG CAAGAAACCT AAGCCGCCTG GTGGATGGCT CGGCTCGGGG CGCTGACGAA GGACGCGTGG ACGCGTG
J23SpTA_8_      CCCGCAGA AGACGGGTTT GATGGGGCGG GGATGTAAGC GGGAAGGGAA ACCGACCCGT TTAGTCTGCC GCTCCCAACA GCCCGAGTTT CTAGCCCCTT GAGTAAGGGG CAAGAAACCT AAGCCGCCTG GTGGATGGCT CGGCTCGGGG CGCTGACGAA GGACGCGTGG ACGCGTAG
J23SpTA_6_       CCGCAGA AGACGGGTTT GATGGGGCGG GGATGTAAGC GGGAAGGGAA ACCGACCCGT TTAGTCTGCC GCTCCCAACA GCCCGAGTTT CTAGCCCCTT GAGTAAGGGG CAAGAAACCT AAGCCGCCTG GTGGATGGCT CGGCTCGGGG CGCTGACGAA GGACGCGTGG ACGCGTAG
J23SpTA_5_        CGCAGA AGACGGGTTT GATGGGGCGG GGATGTAAGC GGGAAGGGAA ACCGACCCGT TTAGTCTGCC GCTCCCAACA GCCCGAGTTT CTAGCCCCTT GAGTAAGGGG CAAGAAACCT AAGCCGCCTG GTGGATGGCT CGGCTCGGGG CGCTGACGAA GGACGCGTGG ACGCGTGGG
J23SpTA_24         GCAGA AGACGGGTTT GATGGGGCGG GGATGTAAGC GGGAAGGGAA ACCGACCCGT TTAGTCTGCC GCTCCCAACA GCCCGAGTTT CTAGCCCCTT GAGTAAGGGG CAAGAAACCT AAGCCGCCTG GTGGATGGCT CGGCTCGGGG CGCTGACGAA GGACGCGTGG ACGCGTGGC
J23SpTA_6_       CCGCAGA AGACGGGTTT GATGGGGCGG GGATGTAAGC GGGAAGGGAA ACCGACCCGT TTAGTCTGCC GCTCCCAACA GCCCGAGTTT CTAGCCCCTT GAGTAAGGGG CAAGAAACCT AAGCCGCCTG GTGGATGGCT CGGCTCGGGG CGCTGACGAA GGACGCGTGG ACGCGAGG
J23SpTA_13        CGCAGA AGACGGGTTT GATGGGGCGG GGATGTAAGC GGGAAGGGAA ACCGACCCGT TTAGTCTGCC GCTCCCAACA GCCCGAGTTT CTAGCCCCTT GAGTAAGGGG CAAGAAACCT AAGCCGCCTG GTGGATGGCT CGGCTCGGGG CGCTGACGAA GGACGCGTGG ACGCGTGAC
J23SpTA_5_      CCCGCAGA AGACGGGTTT GATGGGGCGG GGATGTAAGC GGGAAGGGAA ACCGACCCGT TTAGTCTGCC GCTCCCAACA GCCCGAGTTT CTAGCCCCTT GAGTAAGGGG CAAGAAACCT AAGCCGCCTG GTGGATGGCT CGGCTCGGGG CGCTGACGAA GGACGCGTGG ACGCATGG
J23SpTA_12      CCCGCAGA AGACGGGTTT GATGGGGCGG GGATGTAAGC GGGAAGGGAA ACCGACCCGT TTAGTCTGCC GCTCCCAACA GCCCGAGTTT CTAGCCCCTT GAGTAAGGGG CAAGAAACCT AAGCCGCCTG GTGGATGGCT CGGCTCGGGG CGCTGACGAA GGACGCGTGG ACGCGGGG
J23SpTA_8_       CCGCAGA AGACGGGTTT GATGGGGCGG GGATGTAAGC GGGAAGGGAA ACCGACCCGT TTAGTCTGCC GCTCCCAACA GCCCGAGTTT CTAGCCCCTT GAGTAAGGGG CAAGAAACCT AAGCCGCCTG GTGGATGGCT CGGCTCGGGG CGCTGACGAA GGACGCGTGG ACGCGGGG
J23SpTA_48        CGCAGA AGACGGGTTT GATGGGGCGG GGATGTAAGC GGGAAGGGAA ACCGACCCGT TTAGTCTGCC GCTCCCAACA GCCCGAGTTT CTAGCCCCTT GAGTAAGGGG CAAGAAACCT AAGCCGCCTG GTGGATGGCT CGGCTCGGGG CGCTGACGAA GGACGCGTGG ACGCGTG
J23SpTA_5_       CCGCAGA AGACGGGTTT GATGGGGCGG GGATGTAAGC GGGAAGGGAA ACCGACCCGT TTAGTCTGCC GCTCCCAACA GCCCGAGTTT CTAGCCCCTT GAGTAAGGGG CAAGAAACCT AAGCCGCCTG GTGGATGGCT CGGCTCGGGG CGCTGACGAA GGACGCGTGG ACACGTG
J23SpTA_5_        CGCAGA AGACGGGTTT GATGGGGCGG GGATGTAAGC GGGAAGGGAA ACCGACCCGT TTAGTCTGCC GCTCCCAACA GCCCGAGTTT CTAGCCCCTT GAGTAAGGGG CAAGAAACCT AAGCCGCCTG GTGGATGGCT CGGCTCGGGG CGCTGACGAA GGACGCGTGG ACGCATGG
J23SpTA_98         GCAGA AGACGGGTTT GATGGGGCGG GGATGTAAGC GGGAAGGGAA ACCGACCCGT TTAGTCTGCC GCTCCCAACA GCCCGAGTTT CTAGCCCCTT GAGTAAGGGG CAAGAAACCT AAGCCGCCTG GTGGATGGCT CGGCTCGGGG CGCTGACGAA GGACGCGTGG ACGCGTGG
J23SpTA_5_        CGCAGA AGACGGGTTT GATGGGGCGG GGATGTAAGC GGGAAGGGAA ACCGACCCGT TTAGTCTGCC GCTCCCAACA GCCCGAGTTT CTAGCCCCTT GAGTAAGGGG CAAGAAACCT AAGCCGCCTG GTGGATGGCT CGGCTCGGGG CGCTGACGAA GGACGCGTGG ACGCGCGG
```

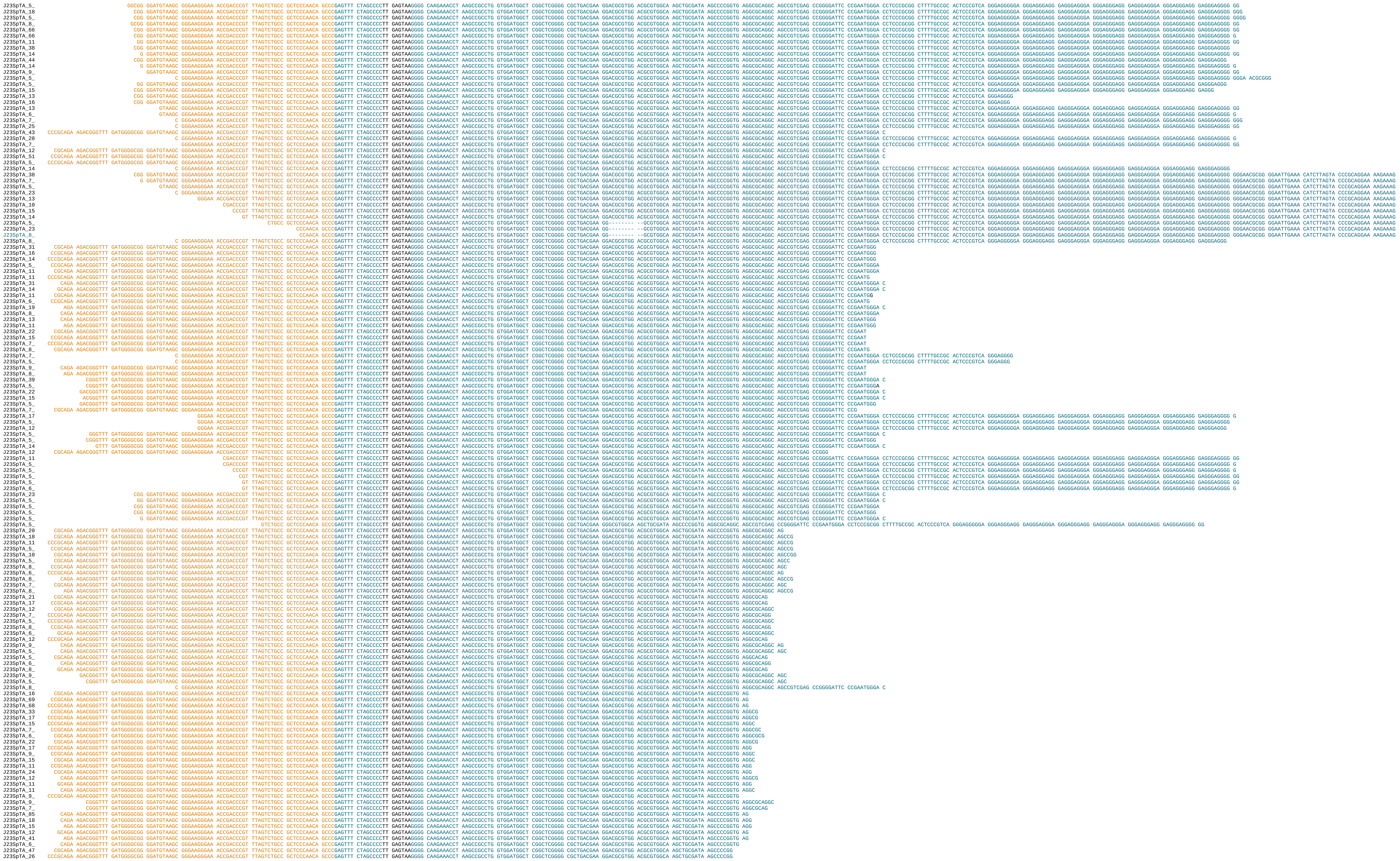

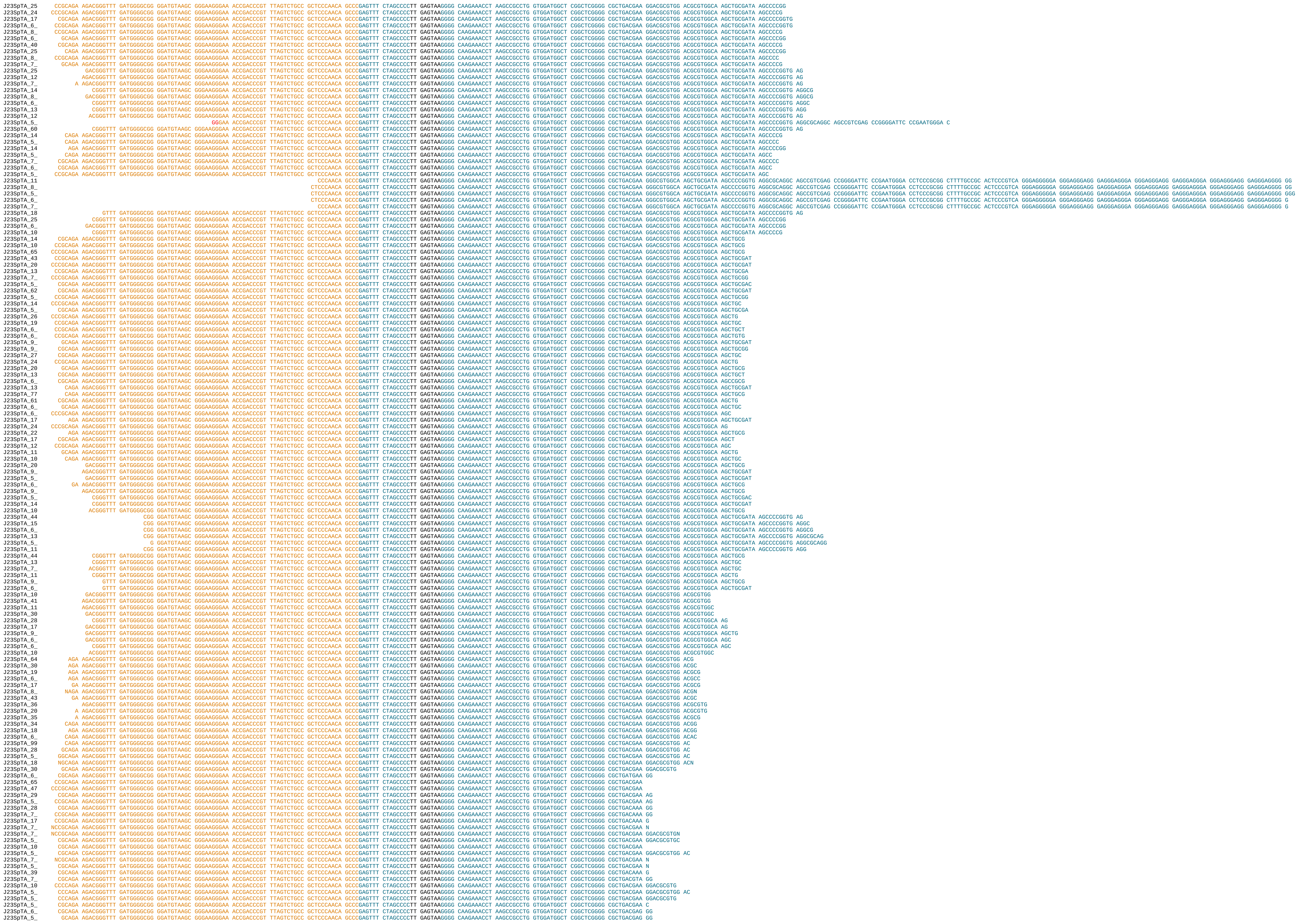

```
J23SpTA_13    CGCAGA AGACGGGTTT GATGGGGCGG GGATGTAAGC GGGAAGGGAA ACCGACCCGT TTAGTCTGCC GCTCCCAACA GCCCGAGTTT CTAGCCCCTT GAGTAAGGGG CAAGAAACCT AAGCCGCCTG GTGGATGGCT CGGCTCGGGG CGCTGACGAG
J23SpTA_8_  CCCGCAGA AGACGGGTTT GATGGGGCGG GGATGTAAGC GGGAAGGGAA ACCGACCCGT TTAGTCTGCC GCTCCCAACA GCCCGAGTTT CTAGCCCCTT GAGTAAGGGG CAAGAAACCT AAGCCGCCTG GTGGATGGCT CGGCTCGGGG CGCTGACGAG
J23SpTA_14    CGCAGA AGACGGGTTT GATGGGGCGG GGATGTAAGC GGGAAGGGAA ACCGACCCGT TTAGTCTGCC GCTCCCAACA GCCCGAGTTT CTAGCCCCTT GAGTAAGGGG CAAGAAACCT AAGCCGCCTG GTGGATGGCT CGGCTCGGGG CGCTGACGA
J23SpTA_12      CAGA AGACGGGTTT GATGGGGCGG GGATGTAAGC GGGAAGGGAA ACCGACCCGT TTAGTCTGCC GCTCCCAACA GCCCGAGTTT CTAGCCCCTT GAGTAAGGGG CAAGAAACCT AAGCCGCCTG GTGGATGGCT CGGCTCGGGG CGCTGACAAA GG
J23SpTA_6_    CGCAGA AGACGGGTTT GATGGGGCGG GGATGTAAGC GGGAAGGGAA ACCGACCCGT TTAGTCTGCC GCTCCCAACA GCCCGAGTTT CTAGCCCCTT GAGTAAGGGG CAAGAAACCT AAGCCGCCTG GTGGATGGCT CGGCTCGGGG CGCTGACGGA G
J23SpTA_10      CAGA AGACGGGTTT GATGGGGCGG GGATGTAAGC GGGAAGGGAA ACCGACCCGT TTAGTCTGCC GCTCCCAACA GCCCGAGTTT CTAGCCCCTT GAGTAAGGGG CAAGAAACCT AAGCCGCCTG GTGGATGGCT CGGCTCGGGG CGCTGACGAA AG
J23SpTA_21     GCAGA AGACGGGTTT GATGGGGCGG GGATGTAAGC GGGAAGGGAA ACCGACCCGT TTAGTCTGCC GCTCCCAACA GCCCGAGTTT CTAGCCCCTT GAGTAAGGGG CAAGAAACCT AAGCCGCCTG GTGGATGGCT CGGCTCGGGG CGCTGACGAA
J23SpTA_11      CAGA AGACGGGTTT GATGGGGCGG GGATGTAAGC GGGAAGGGAA ACCGACCCGT TTAGTCTGCC GCTCCCAACA GCCCGAGTTT CTAGCCCCTT GAGTAAGGGG CAAGAAACCT AAGCCGCCTG GTGGATGGCT CGGCTCGGGG CGCTGACGAA GGACGCGTG
J23SpTA_5_     CCAGA AGACGGGTTT GATGGGGCGG GGATGTAAGC GGGAAGGGAA ACCGACCCGT TTAGTCTGCC GCTCCCAACA GCCCGAGTTT CTAGCCCCTT GAGTAAGGGG CAAGAAACCT AAGCCGCCTG GTGGATGGCT CGGCTCGGGG CGCTGACGAA GGACGCGTG
J23SpTA_14       AGA AGACGGGTTT GATGGGGCGG GGATGTAAGC GGGAAGGGAA ACCGACCCGT TTAGTCTGCC GCTCCCAACA GCCCGAGTTT CTAGCCCCTT GAGTAAGGGG CAAGAAACCT AAGCCGCCTG GTGGATGGCT CGGCTCGGGG CGCTGACGAA GGACGCGTGA G
J23SpTA_13      CAGA AGACGGGTTT GATGGGGCGG GGATGTAAGC GGGAAGGGAA ACCGACCCGT TTAGTCTGCC GCTCCCAACA GCCCGAGTTT CTAGCCCCTT GAGTAAGGGG CAAGAAACCT AAGCCGCCTG GTGGATGGCT CGGCTCGGGG CGCTGACGAA GGACGCGTGA G
J23SpTA_5_      CAGA AGACGGGTTT GATGGGGCGG GGATGTAAGC GGGAAGGGAA ACCGACCCGT TTAGTCTGCC GCTCCCAACA GCCCGAGTTT CTAGCCCCTT GAGTAAGGGG CAAGAAACCT AAGCCGCCTG GTGGATGGCT CGGCTCGGGG CGCTGACGAG GGG
J23SpTA_5_      CAGA AGACGGGTTT GATGGGGCGG GGATGTAAGC GGGAAGGGAA ACCGACCCGT TTAGTCTGCC GCTCCCAACA GCCCGAGTTT CTAGCCCCTT GAGTAAGGGG CAAGAAACCT AAGCCGCCTG GTGGATGGCT CGGCTCGGGG CGCTGACGAG GG
J23SpTA_15    NGCAGA AGACGGGTTT GATGGGGCGG GGATGTAAGC GGGAAGGGAA ACCGACCCGT TTAGTCTGCC GCTCCCAACA GCCCGAGTTT CTAGCCCCTT GAGTAAGGGG CAAGAAACCT AAGCCGCCTG GTGGATGGCT CGGCTCGGGG CGCTGACGAA GGACGCGTGN
J23SpTA_9_    NGCAGA AGACGGGTTT GATGGGGCGG GGATGTAAGC GGGAAGGGAA ACCGACCCGT TTAGTCTGCC GCTCCCAACA GCCCGAGTTT CTAGCCCCTT GAGTAAGGGG CAAGAAACCT AAGCCGCCTG GTGGATGGCT CGGCTCGGGG CGCTGACGAA N
J23SpTA_6_      CAGA AGACGGGTTT GATGGGGCGG GGATGTAAGC GGGAAGGGAA ACCGACCCGT TTAGTCTGCC GCTCCCAACA GCCCGAGTTT CTAGCCCCTT GAGTAAGGGG CAAGAAACCT AAGCCGCCTG GTGGATGGCT CGGCTCGGGG CGCTGACGGA GGG
J23SpTA_6_      CAGA AGACGGGTTT GATGGGGCGG GGATGTAAGC GGGAAGGGAA ACCGACCCGT TTAGTCTGCC GCTCCCAACA GCCCGAGTTT CTAGCCCCTT GAGTAAGGGG CAAGAAACCT AAGCCGCCTG GTGGATGGCT CGGCTCGGGG CGCTGACGAA C
J23SpTA_47       AGA AGACGGGTTT GATGGGGCGG GGATGTAAGC GGGAAGGGAA ACCGACCCGT TTAGTCTGCC GCTCCCAACA GCCCGAGTTT CTAGCCCCTT GAGTAAGGGG CAAGAAACCT AAGCCGCCTG GTGGATGGCT CGGCTCGGGG CGCTGACGAA GGACGCGTG
J23SpTA_46       AGA AGACGGGTTT GATGGGGCGG GGATGTAAGC GGGAAGGGAA ACCGACCCGT TTAGTCTGCC GCTCCCAACA GCCCGAGTTT CTAGCCCCTT GAGTAAGGGG CAAGAAACCT AAGCCGCCTG GTGGATGGCT CGGCTCGGGG CGCTGACGAA GGACGCGTGG AC
J23SpTA_5_      NAGA AGACGGGTTT GATGGGGCGG GGATGTAAGC GGGAAGGGAA ACCGACCCGT TTAGTCTGCC GCTCCCAACA GCCCGAGTTT CTAGCCCCTT GAGTAAGGGG CAAGAAACCT AAGCCGCCTG GTGGATGGCT CGGCTCGGGG CGCTGACGAA GGACGCGTGG ACN
J23SpTA_6_      NAGA AGACGGGTTT GATGGGGCGG GGATGTAAGC GGGAAGGGAA ACCGACCCGT TTAGTCTGCC GCTCCCAACA GCCCGAGTTT CTAGCCCCTT GAGTAAGGGG CAAGAAACCT AAGCCGCCTG GTGGATGGCT CGGCTCGGGG CGCTGACGAA GGACGCGTGN
J23SpTA_97        GA AGACGGGTTT GATGGGGCGG GGATGTAAGC GGGAAGGGAA ACCGACCCGT TTAGTCTGCC GCTCCCAACA GCCCGAGTTT CTAGCCCCTT GAGTAAGGGG CAAGAAACCT AAGCCGCCTG GTGGATGGCT CGGCTCGGGG CGCTGACGAA GGACGCGTGG ACG
J23SpTA_31      CAGA AGACGGGTTT GATGGGGCGG GGATGTAAGC GGGAAGGGAA ACCGACCCGT TTAGTCTGCC GCTCCCAACA GCCCGAGTTT CTAGCCCCTT GAGTAAGGGG CAAGAAACCT AAGCCGCCTG GTGGATGGCT CGGCTCGGGG CGCTGACGAA
J23SpTA_11      CAGA AGACGGGTTT GATGGGGCGG GGATGTAAGC GGGAAGGGAA ACCGACCCGT TTAGTCTGCC GCTCCCAACA GCCCGAGTTT CTAGCCCCTT GAGTAAGGGG CAAGAAACCT AAGCCGCCTG GTGGATGGCT CGGCTCGGGG CGCTGACAAA G
J23SpTA_5_      CAGA AGACGGGTTT GATGGGGCGG GGATGTAAGC GGGAAGGGAA ACCGACCCGT TTAGTCTGCC GCTCCCAACA GCCCGAGTTT CTAGCCCCTT GAGTAAGGGG CAAGAAACCT AAGCCGCCTG GTGGATGGCT CGGCTCGGGG CGCTGATGAA GG
J23SpTA_21            CGGGTTT GATGGGGCGG GGATGTAAGC GGGAAGGGAA ACCGACCCGT TTAGTCTGCC GCTCCCAACA GCCCGAGTTT CTAGCCCCTT GAGTAAGGGG CAAGAAACCT AAGCCGCCTG GTGGATGGCT CGGCTCGGGG CGCTGACGAA GGACGCGTGG ACGCGTGG
J23SpTA_55           ACGGGTTT GATGGGGCGG GGATGTAAGC GGGAAGGGAA ACCGACCCGT TTAGTCTGCC GCTCCCAACA GCCCGAGTTT CTAGCCCCTT GAGTAAGGGG CAAGAAACCT AAGCCGCCTG GTGGATGGCT CGGCTCGGGG CGCTGACGAA GGACGCGTGG ACGCGTGG
J23SpTA_50          GACGGGTTT GATGGGGCGG GGATGTAAGC GGGAAGGGAA ACCGACCCGT TTAGTCTGCC GCTCCCAACA GCCCGAGTTT CTAGCCCCTT GAGTAAGGGG CAAGAAACCT AAGCCGCCTG GTGGATGGCT CGGCTCGGGG CGCTGACGAA GGACGCGTGG ACGCGTG
J23SpTA_48            CGGGTTT GATGGGGCGG GGATGTAAGC GGGAAGGGAA ACCGACCCGT TTAGTCTGCC GCTCCCAACA GCCCGAGTTT CTAGCCCCTT GAGTAAGGGG CAAGAAACCT AAGCCGCCTG GTGGATGGCT CGGCTCGGGG CGCTGACGAA GGACGCGTGG ACGCGTGGC
J23SpTA_7_            CGGGTTT GATGGGGCGG GGATGTAAGC GGGAAGGGAA ACCGACCCGT TTAGTCTGCC GCTCCCAACA GCCCGAGTTT CTAGCCCCTT GAGTAAGGGG CAAGAAACCT AAGCCGCCTG GTGGATGGCT CGGCTCGGGG CGCTGACGAA GGACGCGTGG ACGCGTGGCG
J23SpTA_40           ACGGGTTT GATGGGGCGG GGATGTAAGC GGGAAGGGAA ACCGACCCGT TTAGTCTGCC GCTCCCAACA GCCCGAGTTT CTAGCCCCTT GAGTAAGGGG CAAGAAACCT AAGCCGCCTG GTGGATGGCT CGGCTCGGGG CGCTGACGAA GGACGCGTGG ACGCGTG
J23SpTA_13            CGGGTTT GATGGGGCGG GGATGTAAGC GGGAAGGGAA ACCGACCCGT TTAGTCTGCC GCTCCCAACA GCCCGAGTTT CTAGCCCCTT GAGTAAGGGG CAAGAAACCT AAGCCGCCTG GTGGATGGCT CGGCTCGGGG CGCTGACGAA GGACGCGTGG ACGCGTG
J23SpTA_14             GGGTTT GATGGGGCGG GGATGTAAGC GGGAAGGGAA ACCGACCCGT TTAGTCTGCC GCTCCCAACA GCCCGAGTTT CTAGCCCCTT GAGTAAGGGG CAAGAAACCT AAGCCGCCTG GTGGATGGCT CGGCTCGGGG CGCTGACGAA GGACGCGTGG ACGCGTGG
J23SpTA_9_             GGGTTT GATGGGGCGG GGATGTAAGC GGGAAGGGAA ACCGACCCGT TTAGTCTGCC GCTCCCAACA GCCCGAGTTT CTAGCCCCTT GAGTAAGGGG CAAGAAACCT AAGCCGCCTG GTGGATGGCT CGGCTCGGGG CGCTGACGAA GGACGCGTGG ACGCGTGGC
J23SpTA_5_          GACGGGTTT GATGGGGCGG GGATGTAAGC GGGAAGGGAA ACCGACCCGT TTAGTCTGCC GCTCCCAACA GCCCGAGTTT CTAGCCCCTT GAGTAAGGGG CAAGAAACCT AAGCCGCCTG GTGGATGGCT CGGCTCGGGG CGCTGACGAA GGACGCGTGG ACGCGTA
J23SpTA_5_          GACGGGTTT GATGGGGCGG GGATGTAAGC GGGAAGGGAA ACCGACCCGT TTAGTCTGCC GCTCCCAACA GCCCGAGTTT CTAGCCCCTT GAGTAAGGGG CAAGAAACCT AAGCCGCCTG GTGGATGGCT CGGCTCGGGG CGCTGACGAA GGACGCGTGG ACGCGT
J23SpTA_32          GACGGGTTT GATGGGGCGG GGATGTAAGC GGGAAGGGAA ACCGACCCGT TTAGTCTGCC GCTCCCAACA GCCCGAGTTT CTAGCCCCTT GAGTAAGGGG CAAGAAACCT AAGCCGCCTG GTGGATGGCT CGGCTCGGGG CGCTGACGAA GGACGCGTGG ACGC
J23SpTA_6_          GACGGGTTT GATGGGGCGG GGATGTAAGC GGGAAGGGAA ACCGACCCGT TTAGTCTGCC GCTCCCAACA GCCCGAGTTT CTAGCCCCTT GAGTAAGGGG CAAGAAACCT AAGCCGCCTG GTGGATGGCT CGGCTCGGGG CGCTGACGAA GGACGCGTGG ACGCC
J23SpTA_15          GACGGGTTT GATGGGGCGG GGATGTAAGC GGGAAGGGAA ACCGACCCGT TTAGTCTGCC GCTCCCAACA GCCCGAGTTT CTAGCCCCTT GAGTAAGGGG CAAGAAACCT AAGCCGCCTG GTGGATGGCT CGGCTCGGGG CGCTGACGAA GGACGCGTGG ACGCG
J23SpTA_59         AGACGGGTTT GATGGGGCGG GGATGTAAGC GGGAAGGGAA ACCGACCCGT TTAGTCTGCC GCTCCCAACA GCCCGAGTTT CTAGCCCCTT GAGTAAGGGG CAAGAAACCT AAGCCGCCTG GTGGATGGCT CGGCTCGGGG CGCTGACGAA GGACGCGTGG ACGCG
J23SpTA_10         AGACGGGTTT GATGGGGCGG GGATGTAAGC GGGAAGGGAA ACCGACCCGT TTAGTCTGCC GCTCCCAACA GCCCGAGTTT CTAGCCCCTT GAGTAAGGGG CAAGAAACCT AAGCCGCCTG GTGGATGGCT CGGCTCGGGG CGCTGACGAA GGACGCGTGG ACGC
J23SpTA_45         A AGACGGGTTT GATGGGGCGG GGATGTAAGC GGGAAGGGAA ACCGACCCGT TTAGTCTGCC GCTCCCAACA GCCCGAGTTT CTAGCCCCTT GAGTAAGGGG CAAGAAACCT AAGCCGCCTG GTGGATGGCT CGGCTCGGGG CGCTGACGAA GGACGCGTGG ACGC
J23SpTA_21         AGACGGGTTT GATGGGGCGG GGATGTAAGC GGGAAGGGAA ACCGACCCGT TTAGTCTGCC GCTCCCAACA GCCCGAGTTT CTAGCCCCTT GAGTAAGGGG CAAGAAACCT AAGCCGCCTG GTGGATGGCT CGGCTCGGGG CGCTGACGAA GGACGCGTGG ACG
J23SpTA_14         A AGACGGGTTT GATGGGGCGG GGATGTAAGC GGGAAGGGAA ACCGACCCGT TTAGTCTGCC GCTCCCAACA GCCCGAGTTT CTAGCCCCTT GAGTAAGGGG CAAGAAACCT AAGCCGCCTG GTGGATGGCT CGGCTCGGGG CGCTGACGAA GGACGCGTGG ACG
J23SpTA_78           ACGGGTTT GATGGGGCGG GGATGTAAGC GGGAAGGGAA ACCGACCCGT TTAGTCTGCC GCTCCCAACA GCCCGAGTTT CTAGCCCCTT GAGTAAGGGG CAAGAAACCT AAGCCGCCTG GTGGATGGCT CGGCTCGGGG CGCTGACGAA GGACGCGTGG ACGCG
J23SpTA_14    CGCAGA AGACGGGTTT GATGGGGCGG GGATGTAAGC GGGAAGGGAA ACCGACCCGT TTAGTCTGCC GCTCCCAACA GCCCGAGTTT CTAGCCCCTT GAGTAAGGGG CAAGAAACCT AAGCCGCCTG GTGGATGGCT CGGCTCGGGG CGCTGACG
J23SpTA_70   CCGCAGA AGACGGGTTT GATGGGGCGG GGATGTAAGC GGGAAGGGAA ACCGACCCGT TTAGTCTGCC GCTCCCAACA GCCCGAGTTT CTAGCCCCTT GAGTAAGGGG CAAGAAACCT AAGCCGCCTG GTGGATGGCT CGGCTCGGGG CGCTGACG
J23SpTA_46  CCCGCAGA AGACGGGTTT GATGGGGCGG GGATGTAAGC GGGAAGGGAA ACCGACCCGT TTAGTCTGCC GCTCCCAACA GCCCGAGTTT CTAGCCCCTT GAGTAAGGGG CAAGAAACCT AAGCCGCCTG GTGGATGGCT CGGCTCGGGG CGCTGACG
J23SpTA_37  CCCGCAGA AGACGGGTTT GATGGGGCGG GGATGTAAGC GGGAAGGGAA ACCGACCCGT TTAGTCTGCC GCTCCCAACA GCCCGAGTTT CTAGCCCCTT GAGTAAGGGG CAAGAAACCT AAGCCGCCTG GTGGATGGCT CGGCTCGGGG CGCTGAC
J23SpTA_39   CCGCAGA AGACGGGTTT GATGGGGCGG GGATGTAAGC GGGAAGGGAA ACCGACCCGT TTAGTCTGCC GCTCCCAACA GCCCGAGTTT CTAGCCCCTT GAGTAAGGGG CAAGAAACCT AAGCCGCCTG GTGGATGGCT CGGCTCGGGG CGCTGAC
J23SpTA_5_    CGCAGA AGACGGGTTT GATGGGGCGG GGATGTAAGC GGGAAGGGAA ACCGACCCGT TTAGTCTGCC GCTCCCAACA GCCCGAGTTT CTAGCCCCTT GAGTAAGGGG CAAGAAACCT AAGCCGCCTG GTGGATGGCT CGGCTCGGGG CGCTGACGA
J23SpTA_74    CGCAGA AGACGGGTTT GATGGGGCGG GGATGTAAGC GGGAAGGGAA ACCGACCCGT TTAGTCTGCC GCTCCCAACA GCCCGAGTTT CTAGCCCCTT GAGTAAGGGG CAAGAAACCT AAGCCGCCTG GTGGATGGCT CGGCTCGGGG CGCTGAC
J23SpTA_13     GCAGA AGACGGGTTT GATGGGGCGG GGATGTAAGC GGGAAGGGAA ACCGACCCGT TTAGTCTGCC GCTCCCAACA GCCCGAGTTT CTAGCCCCTT GAGTAAGGGG CAAGAAACCT AAGCCGCCTG GTGGATGGCT CGGCTCGGGG CGCTGACG
J23SpTA_12         AGACGGGTTT GATGGGGCGG GGATGTAAGC GGGAAGGGAA ACCGACCCGT TTAGTCTGCC GCTCCCAACA GCCCGAGTTT CTAGCCCCTT GAGTAAGGGG CAAGAAACCT AAGCCGCCTG GTGGATGGCT CGGCTCGGGG CGCTGACGAA GGACGCGTGG AC
J23SpTA_94         A AGACGGGTTT GATGGGGCGG GGATGTAAGC GGGAAGGGAA ACCGACCCGT TTAGTCTGCC GCTCCCAACA GCCCGAGTTT CTAGCCCCTT GAGTAAGGGG CAAGAAACCT AAGCCGCCTG GTGGATGGCT CGGCTCGGGG CGCTGACGAA GGACGCGTGG AC
J23SpTA_54        GA AGACGGGTTT GATGGGGCGG GGATGTAAGC GGGAAGGGAA ACCGACCCGT TTAGTCTGCC GCTCCCAACA GCCCGAGTTT CTAGCCCCTT GAGTAAGGGG CAAGAAACCT AAGCCGCCTG GTGGATGGCT CGGCTCGGGG CGCTGACGAA GGACGCGTGG AC
J23SpTA_78        GA AGACGGGTTT GATGGGGCGG GGATGTAAGC GGGAAGGGAA ACCGACCCGT TTAGTCTGCC GCTCCCAACA GCCCGAGTTT CTAGCCCCTT GAGTAAGGGG CAAGAAACCT AAGCCGCCTG GTGGATGGCT CGGCTCGGGG CGCTGACGAA GGACGCGTG
J23SpTA_17       AGA AGACGGGTTT GATGGGGCGG GGATGTAAGC GGGAAGGGAA ACCGACCCGT TTAGTCTGCC GCTCCCAACA GCCCGAGTTT CTAGCCCCTT GAGTAAGGGG CAAGAAACCT AAGCCGCCTG GTGGATGGCT CGGCTCGGGG CGCTGACGAA
J23SpTA_76         A AGACGGGTTT GATGGGGCGG GGATGTAAGC GGGAAGGGAA ACCGACCCGT TTAGTCTGCC GCTCCCAACA GCCCGAGTTT CTAGCCCCTT GAGTAAGGGG CAAGAAACCT AAGCCGCCTG GTGGATGGCT CGGCTCGGGG CGCTGACGAA GGACGCGTG
J23SpTA_56               GTTT GATGGGGCGG GGATGTAAGC GGGAAGGGAA ACCGACCCGT TTAGTCTGCC GCTCCCAACA GCCCGAGTTT CTAGCCCCTT GAGTAAGGGG CAAGAAACCT AAGCCGCCTG GTGGATGGCT CGGCTCGGGG CGCTGACGAA GGACGCGTGG ACGCGTGG
J23SpTA_14              GGTTT GATGGGGCGG GGATGTAAGC GGGAAGGGAA ACCGACCCGT TTAGTCTGCC GCTCCCAACA GCCCGAGTTT CTAGCCCCTT GAGTAAGGGG CAAGAAACCT AAGCCGCCTG GTGGATGGCT CGGCTCGGGG CGCTGACGAA GGACGCGTGG  ACGCGTGG
J23SpTA_12               GTTT GATGGGGCGG GGATGTAAGC GGGAAGGGAA ACCGACCCGT TTAGTCTGCC GCTCCCAACA GCCCGAGTTT CTAGCCCCTT GAGTAAGGGG CAAGAAACCT AAGCCGCCTG GTGGATGGCT CGGCTCGGGG CGCTGACGAA GGACGCGTGG ACGCGTGGC
J23SpTA_6_              GGTTT GATGGGGCGG GGATGTAAGC GGGAAGGGAA ACCGACCCGT TTAGTCTGCC GCTCCCAACA GCCCGAGTTT CTAGCCCCTT GAGTAAGGGG CAAGAAACCT AAGCCGCCTG GTGGATGGCT CGGCTCGGGG CGCTGACGAA GGACGCGTGG ACGCGTGGCA AG
J23SpTA_11             GGGTTT GATGGGGCGG GGATGTAAGC GGGAAGGGAA ACCGACCCGT TTAGTCTGCC GCTCCCAACA GCCCGAGTTT CTAGCCCCTT GAGTAAGGGG CAAGAAACCT AAGCCGCCTG GTGGATGGCT CGGCTCGGGG CGCTGACGAA GGACGCGTGG ACGCGTG
J23SpTA_60          GACGGGTTT GATGGGGCGG GGATGTAAGC GGGAAGGGAA ACCGACCCGT TTAGTCTGCC GCTCCCAACA GCCCGAGTTT CTAGCCCCTT GAGTAAGGGG CAAGAAACCT AAGCCGCCTG GTGGATGGCT CGGCTCGGGG CGCTGACGAA GGACGCGTGG ACG
J23SpTA_26          GACGGGTTT GATGGGGCGG GGATGTAAGC GGGAAGGGAA ACCGACCCGT TTAGTCTGCC GCTCCCAACA GCCCGAGTTT CTAGCCCCTT GAGTAAGGGG CAAGAAACCT AAGCCGCCTG GTGGATGGCT CGGCTCGGGG CGCTGACGAA GGACGCGTGG ACGG
J23SpTA_12           ACGGGTTT GATGGGGCGG GGATGTAAGC GGGAAGGGAA ACCGACCCGT TTAGTCTGCC GCTCCCAACA GCCCGAGTTT CTAGCCCCTT GAGTAAGGGG CAAGAAACCT AAGCCGCCTG GTGGATGGCT CGGCTCGGGG CGCTGACGAA GGACGCGTGG ACGG
J23SpTA_36          GACGGGTTT GATGGGGCGG GGATGTAAGC GGGAAGGGAA ACCGACCCGT TTAGTCTGCC GCTCCCAACA GCCCGAGTTT CTAGCCCCTT GAGTAAGGGG CAAGAAACCT AAGCCGCCTG GTGGATGGCT CGGCTCGGGG CGCTGACGAA GGACGCGTGG AC
J23SpTA_19           ACGGGTTT GATGGGGCGG GGATGTAAGC GGGAAGGGAA ACCGACCCGT TTAGTCTGCC GCTCCCAACA GCCCGAGTTT CTAGCCCCTT GAGTAAGGGG CAAGAAACCT AAGCCGCCTG GTGGATGGCT CGGCTCGGGG CGCTGACGAA GGACGCGTGG ACG
J23SpTA_12           ACGGGTTT GATGGGGCGG GGATGTAAGC GGGAAGGGAA ACCGACCCGT TTAGTCTGCC GCTCCCAACA GCCCGAGTTT CTAGCCCCTT GAGTAAGGGG CAAGAAACCT AAGCCGCCTG GTGGATGGCT CGGCTCGGGG CGCTGACGAA GGACGCGTGG ACGC
J23SpTA_39            CGGGTTT GATGGGGCGG GGATGTAAGC GGGAAGGGAA ACCGACCCGT TTAGTCTGCC GCTCCCAACA GCCCGAGTTT CTAGCCCCTT GAGTAAGGGG CAAGAAACCT AAGCCGCCTG GTGGATGGCT CGGCTCGGGG CGCTGACGAA GGACGCGTGG ACGG
J23SpTA_5_          GACGGGTTT GATGGGGCGG GGATGTAAGC GGGAAGGGAA ACCGACCCGT TTAGTCTGCC GCTCCCAACA GCCCGAGTTT CTAGCCCCTT GAGTAAGGGG CAAGAAACCT AAGCCGCCTG GTGGATGGCT CGGCTCGGGG CGCTGACGGA GGG
J23SpTA_5_          GACGGGTTT GATGGGGCGG GGATGTAAGC GGGAAGGGAA ACCGACCCGT TTAGTCTGCC GCTCCCAACA GCCCGAGTTT CTAGCCCCTT GAGTAAGGGG CAAGAAACCT AAGCCGCCTG GTGGATGGCT CGGCTCGGGG CGCTGACGAA GGACGCGTGA G
J23SpTA_50            CGGGTTT GATGGGGCGG GGATGTAAGC GGGAAGGGAA ACCGACCCGT TTAGTCTGCC GCTCCCAACA GCCCGAGTTT CTAGCCCCTT GAGTAAGGGG CAAGAAACCT AAGCCGCCTG GTGGATGGCT CGGCTCGGGG CGCTGACGAA GGACGCGTGG ACGC
J23SpTA_30            CGGGTTT GATGGGGCGG GGATGTAAGC GGGAAGGGAA ACCGACCCGT TTAGTCTGCC GCTCCCAACA GCCCGAGTTT CTAGCCCCTT GAGTAAGGGG CAAGAAACCT AAGCCGCCTG GTGGATGGCT CGGCTCGGGG CGCTGACGAA GGACGCGTGG ACGCG
J23SpTA_6_            CGGGTTT GATGGGGCGG GGATGTAAGC GGGAAGGGAA ACCGACCCGT TTAGTCTGCC GCTCCCAACA GCCCGAGTTT CTAGCCCCTT GAGTAAGGGG CAAGAAACCT AAGCCGCCTG GTGGATGGCT CGGCTCGGGG CGCTGACGAA GGACGCGTGG ACGCGT
J23SpTA_25             GGGTTT GATGGGGCGG GGATGTAAGC GGGAAGGGAA ACCGACCCGT TTAGTCTGCC GCTCCCAACA GCCCGAGTTT CTAGCCCCTT GAGTAAGGGG CAAGAAACCT AAGCCGCCTG GTGGATGGCT CGGCTCGGGG CGCTGACGAA GGACGCGTGG ACGCG
J23SpTA_11            CGGGTTT GATGGGGCGG GGATGTAAGC GGGAAGGGAA ACCGACCCGT TTAGTCTGCC GCTCCCAACA GCCCGAGTTT CTAGCCCCTT GAGTAAGGGG CAAGAAACCT AAGCCGCCTG GTGGATGGCT CGGCTCGGGG CGCTGACGAA GGACGCGTGG ACGCC
J23SpTA_37          GACGGGTTT GATGGGGCGG GGATGTAAGC GGGAAGGGAA ACCGACCCGT TTAGTCTGCC GCTCCCAACA GCCCGAGTTT CTAGCCCCTT GAGTAAGGGG CAAGAAACCT AAGCCGCCTG GTGGATGGCT CGGCTCGGGG CGCTGACGAA GGACGCGTG
J23SpTA_12         AGACGGGTTT GATGGGGCGG GGATGTAAGC GGGAAGGGAA ACCGACCCGT TTAGTCTGCC GCTCCCAACA GCCCGAGTTT CTAGCCCCTT GAGTAAGGGG CAAGAAACCT AAGCCGCCTG GTGGATGGCT CGGCTCGGGG CGCTGACGAA GGACGCGTG
J23SpTA_6_         AGACGGGTTT GATGGGGCGG GGATGTAAGC GGGAAGGGAA ACCGACCCGT TTAGTCTGCC GCTCCCAACA GCCCGAGTTT CTAGCCCCTT GAGTAAGGGG CAAGAAACCT AAGCCGCCTG GTGGATGGCT CGGCTCGGGG CGCTGACGAA
J23SpTA_52      CAGA AGACGGGTTT GATGGGGCGG GGATGTAAGC GGGAAGGGAA ACCGACCCGT TTAGTCTGCC GCTCCCAACA GCCCGAGTTT CTAGCCCCTT GAGTAAGGGG CAAGAAACCT AAGCCGCCTG GTGGATGGCT CGGCTCGGGG CGCTGACG
J23SpTA_6_      CAGA AGACGGGTTT GATGGGGCGG GGATGTAAGC GGGAAGGGAA ACCGACCCGT TTAGTCTGCC GCTCCCAACA GCCCGAGTTT CTAGCCCCTT GAGTAAGGGG CAAGAAACCT AAGCCGCCTG GTGGATGGCT CGGCTCGGGG CGCTGACGAG
J23SpTA_15       AGA AGACGGGTTT GATGGGGCGG GGATGTAAGC GGGAAGGGAA ACCGACCCGT TTAGTCTGCC GCTCCCAACA GCCCGAGTTT CTAGCCCCTT GAGTAAGGGG CAAGAAACCT AAGCCGCCTG GTGGATGGCT CGGCTCGGGG CGCTGACG
J23SpTA_27      CAGA AGACGGGTTT GATGGGGCGG GGATGTAAGC GGGAAGGGAA ACCGACCCGT TTAGTCTGCC GCTCCCAACA GCCCGAGTTT CTAGCCCCTT GAGTAAGGGG CAAGAAACCT AAGCCGCCTG GTGGATGGCT CGGCTCGGGG CGCTGAC
J23SpTA_15       AGA AGACGGGTTT GATGGGGCGG GGATGTAAGC GGGAAGGGAA ACCGACCCGT TTAGTCTGCC GCTCCCAACA GCCCGAGTTT CTAGCCCCTT GAGTAAGGGG CAAGAAACCT AAGCCGCCTG GTGGATGGCT CGGCTCGGGG CGCTGAC
J23SpTA_85            CGGGTTT GATGGGGCGG GGATGTAAGC GGGAAGGGAA ACCGACCCGT TTAGTCTGCC GCTCCCAACA GCCCGAGTTT CTAGCCCCTT GAGTAAGGGG CAAGAAACCT AAGCCGCCTG GTGGATGGCT CGGCTCGGGG CGCTGACGAA GGACGCGTGG ACG
J23SpTA_6_            CGGGTTT GATGGGGCGG GGATGTAAGC GGGAAGGGAA ACCGACCCGT TTAGTCTGCC GCTCCCAACA GCCCGAGTTT CTAGCCCCTT GAGTAAGGGG CAAGAAACCT AAGCCGCCTG GTGGATGGCT CGGCTCGGGG CGCTGACGAA GGACGCGTGG ACAC
J23SpTA_55            CGGGTTT GATGGGGCGG GGATGTAAGC GGGAAGGGAA ACCGACCCGT TTAGTCTGCC GCTCCCAACA GCCCGAGTTT CTAGCCCCTT GAGTAAGGGG CAAGAAACCT AAGCCGCCTG GTGGATGGCT CGGCTCGGGG CGCTGACGAA GGACGCGTGG AC
J23SpTA_14           ACGGGTTT GATGGGGCGG GGATGTAAGC GGGAAGGGAA ACCGACCCGT TTAGTCTGCC GCTCCCAACA GCCCGAGTTT CTAGCCCCTT GAGTAAGGGG CAAGAAACCT AAGCCGCCTG GTGGATGGCT CGGCTCGGGG CGCTGACGAA GGACGCGTGG AC
J23SpTA_13            CGGGTTT GATGGGGCGG GGATGTAAGC GGGAAGGGAA ACCGACCCGT TTAGTCTGCC GCTCCCAACA GCCCGAGTTT CTAGCCCCTT GAGTAAGGGG CAAGAAACCT AAGCCGCCTG GTGGATGGCT CGGCTCGGGG CGCTGACGAA GGACGCGTGA G
J23SpTA_82             GGGTTT GATGGGGCGG GGATGTAAGC GGGAAGGGAA ACCGACCCGT TTAGTCTGCC GCTCCCAACA GCCCGAGTTT CTAGCCCCTT GAGTAAGGGG CAAGAAACCT AAGCCGCCTG GTGGATGGCT CGGCTCGGGG CGCTGACGAA GGACGCGTGG ACG
J23SpTA_5_            CGGGTTT GATGGGGCGG GGATGTAAGC GGGAAGGGAA ACCGACCCGT TTAGTCTGCC GCTCCCAACA GCCCGAGTTT CTAGCCCCTT GAGTAAGGGG CAAGAAACCT AAGCCGCCTG GTGGATGGCT CGGCTCGGGG CGCTGACAAA GGG
J23SpTA_46             GGGTTT GATGGGGCGG GGATGTAAGC GGGAAGGGAA ACCGACCCGT TTAGTCTGCC GCTCCCAACA GCCCGAGTTT CTAGCCCCTT GAGTAAGGGG CAAGAAACCT AAGCCGCCTG GTGGATGGCT CGGCTCGGGG CGCTGACGAA GGACGCGTGG ACGC
J23SpTA_8_    CGCAGA AGACGGGTTT GATGGGGCGG GGATGTAAGC GGGAAGGGAA ACCGACCCGT TTAGTCTGCC GCTCCCAACA GCCCGAGTTT CTAGCCCCTT GAGTAAGGGG CAAGAAACCT AAGCCGCCTG GTGGATGGCT CGGCTCGGGG CGCTG
J23SpTA_6_  CCCGCAGA AGACGGGTTT GATGGGGCGG GGATGTAAGC GGGAAGGGAA ACCGACCCGT TTAGTCTGCC GCTCCCAACA GCCCGAGTTT CTAGCCCCTT GAGTAAGGGG CAAGAAACCT AAGCCGCCTG GTGGATGGCT CGGCTCGGGG CGCT
J23SpTA_63            CGGGTTT GATGGGGCGG GGATGTAAGC GGGAAGGGAA ACCGACCCGT TTAGTCTGCC GCTCCCAACA GCCCGAGTTT CTAGCCCCTT GAGTAAGGGG CAAGAAACCT AAGCCGCCTG GTGGATGGCT CGGCTCGGGG CGCTGACGAA GGACGCGTG
J23SpTA_16           ACGGGTTT GATGGGGCGG GGATGTAAGC GGGAAGGGAA ACCGACCCGT TTAGTCTGCC GCTCCCAACA GCCCGAGTTT CTAGCCCCTT GAGTAAGGGG CAAGAAACCT AAGCCGCCTG GTGGATGGCT CGGCTCGGGG CGCTGACGAA GGACGCGTG
J23SpTA_8_          GACGGGTTT GATGGGGCGG GGATGTAAGC GGGAAGGGAA ACCGACCCGT TTAGTCTGCC GCTCCCAACA GCCCGAGTTT CTAGCCCCTT GAGTAAGGGG CAAGAAACCT AAGCCGCCTG GTGGATGGCT CGGCTCGGGG CGCTGACAAA G
J23SpTA_6_          GACGGGTTT GATGGGGCGG GGATGTAAGC GGGAAGGGAA ACCGACCCGT TTAGTCTGCC GCTCCCAACA GCCCGAGTTT CTAGCCCCTT GAGTAAGGGG CAAGAAACCT AAGCCGCCTG GTGGATGGCT CGGCTCGGGG CGCTGACAAA GG
J23SpTA_6_           ACGGGTTT GATGGGGCGG GGATGTAAGC GGGAAGGGAA ACCGACCCGT TTAGTCTGCC GCTCCCAACA GCCCGAGTTT CTAGCCCCTT GAGTAAGGGG CAAGAAACCT AAGCCGCCTG GTGGATGGCT CGGCTCGGGG CGCTGACAAA G
J23SpTA_6_            CGGGTTT GATGGGGCGG GGATGTAAGC GGGAAGGGAA ACCGACCCGT TTAGTCTGCC GCTCCCAACA GCCCGAGTTT CTAGCCCCTT GAGTAAGGGG CAAGAAACCT AAGCCGCCTG GTGGATGGCT CGGCTCGGGG CGCTGACAAA GG
J23SpTA_21          GACGGGTTT GATGGGGCGG GGATGTAAGC GGGAAGGGAA ACCGACCCGT TTAGTCTGCC GCTCCCAACA GCCCGAGTTT CTAGCCCCTT GAGTAAGGGG CAAGAAACCT AAGCCGCCTG GTGGATGGCT CGGCTCGGGG CGCTGACGAA
J23SpTA_14               GTTT GATGGGGCGG GGATGTAAGC GGGAAGGGAA ACCGACCCGT TTAGTCTGCC GCTCCCAACA GCCCGAGTTT CTAGCCCCTT GAGTAAGGGG CAAGAAACCT AAGCCGCCTG GTGGATGGCT CGGCTCGGGG CGCTGACGAA GGACGCGTGG ACGC
J23SpTA_81               GTTT GATGGGGCGG GGATGTAAGC GGGAAGGGAA ACCGACCCGT TTAGTCTGCC GCTCCCAACA GCCCGAGTTT CTAGCCCCTT GAGTAAGGGG CAAGAAACCT AAGCCGCCTG GTGGATGGCT CGGCTCGGGG CGCTGACGAA GGACGCGTGG ACGCG
J23SpTA_26               GTTT GATGGGGCGG GGATGTAAGC GGGAAGGGAA ACCGACCCGT TTAGTCTGCC GCTCCCAACA GCCCGAGTTT CTAGCCCCTT GAGTAAGGGG CAAGAAACCT AAGCCGCCTG GTGGATGGCT CGGCTCGGGG CGCTGACGAA GGACGCGTGG ACGCGTG
J23SpTA_6_               GTTT GATGGGGCGG GGATGTAAGC GGGAAGGGAA ACCGACCCGT TTAGTCTGCC GCTCCCAACA GCCCGAGTTT CTAGCCCCTT GAGTAAGGGG CAAGAAACCT AAGCCGCCTG GTGGATGGCT CGGCTCGGGG CGCTGACGAA GGACGCGTGG ACGCGT
J23SpTA_23              GGTTT GATGGGGCGG GGATGTAAGC GGGAAGGGAA ACCGACCCGT TTAGTCTGCC GCTCCCAACA GCCCGAGTTT CTAGCCCCTT GAGTAAGGGG CAAGAAACCT AAGCCGCCTG GTGGATGGCT CGGCTCGGGG CGCTGACGAA GGACGCGTGG ACGCG
J23SpTA_31              GGTTT GATGGGGCGG GGATGTAAGC GGGAAGGGAA ACCGACCCGT TTAGTCTGCC GCTCCCAACA GCCCGAGTTT CTAGCCCCTT GAGTAAGGGG CAAGAAACCT AAGCCGCCTG GTGGATGGCT CGGCTCGGGG CGCTGACGAA GGACGCGTGG ACGC
J23SpTA_50             GGGTTT GATGGGGCGG GGATGTAAGC GGGAAGGGAA ACCGACCCGT TTAGTCTGCC GCTCCCAACA GCCCGAGTTT CTAGCCCCTT GAGTAAGGGG CAAGAAACCT AAGCCGCCTG GTGGATGGCT CGGCTCGGGG CGCTGACGAA GGACGCGTG
J23SpTA_43             GGGTTT GATGGGGCGG GGATGTAAGC GGGAAGGGAA ACCGACCCGT TTAGTCTGCC GCTCCCAACA GCCCGAGTTT CTAGCCCCTT GAGTAAGGGG CAAGAAACCT AAGCCGCCTG GTGGATGGCT CGGCTCGGGG CGCTGACGAA GGACGCGTGG AC
J23SpTA_36              GGTTT GATGGGGCGG GGATGTAAGC GGGAAGGGAA ACCGACCCGT TTAGTCTGCC GCTCCCAACA GCCCGAGTTT CTAGCCCCTT GAGTAAGGGG CAAGAAACCT AAGCCGCCTG GTGGATGGCT CGGCTCGGGG CGCTGACGAA GGACGCGTGG AC
J23SpTA_5_             GGGTTT GATGGGGCGG GGATGTAAGC GGGAAGGGAA ACCGACCCGT TTAGTCTGCC GCTCCCAACA GCCCGAGTTT CTAGCCCCTT GAGTAAGGGG CAAGAAACCT AAGCCGCCTG GTGGATGGCT CGGCTCGGGG CGCTGACGAA AG
J23SpTA_33                          CGG GGATGTAAGC GGGAAGGGAA ACCGACCCGT TTAGTCTGCC GCTCCCAACA GCCCGAGTTT CTAGCCCCTT GAGTAAGGGG CAAGAAACCT AAGCCGCCTG GTGGATGGCT CGGCTCGGGG CGCTGACGAA GGACGCGTGG ACGCGTGGCA AGCTGCG
J23SpTA_16                          CGG GGATGTAAGC GGGAAGGGAA ACCGACCCGT TTAGTCTGCC GCTCCCAACA GCCCGAGTTT CTAGCCCCTT GAGTAAGGGG CAAGAAACCT AAGCCGCCTG GTGGATGGCT CGGCTCGGGG CGCTGACGAA GGACGCGTGG ACGCGTGGCA AGCTGCGAT
J23SpTA_10                          CGG GGATGTAAGC GGGAAGGGAA ACCGACCCGT TTAGTCTGCC GCTCCCAACA GCCCGAGTTT CTAGCCCCTT GAGTAAGGGG CAAGAAACCT AAGCCGCCTG GTGGATGGCT CGGCTCGGGG CGCTGACGAA GGACGCGTGG ACGCGTGGCA AGCTGCGATA AGCCCCGG
J23SpTA_6_                          CGG GGATGTAAGC GGGAAGGGAA ACCGACCCGT TTAGTCTGCC GCTCCCAACA GCCCGAGTTT CTAGCCCCTT GAGTAAGGGG CAAGAAACCT AAGCCGCCTG GTGGATGGCT CGGCTCGGGG CGCTGACGAA GGACGCGTGG ACGCGTGGCA AGCTGCGATA AGCCCCGGTG
J23SpTA_8_                          CGG GGATGTAAGC GGGAAGGGAA ACCGACCCGT TTAGTCTGCC GCTCCCAACA GCCCGAGTTT CTAGCCCCTT GAGTAAGGGG CAAGAAACCT AAGCCGCCTG GTGGATGGCT CGGCTCGGGG CGCTGACGAA GGACGCGTGG ACGCGTGGCA AGCTGCGATA AGCCCCG
J23SpTA_5_                           GG GGATGTAAGC GGGAAGGGAA ACCGACCCGT TTAGTCTGCC GCTCCCAACA GCCCGAGTTT CTAGCCCCTT GAGTAAGGGG CAAGAAACCT AAGCCGCCTG GTGGATGGCT CGGCTCGGGG CGCTGACGAA GGACGCGTGG ACGCGTGGCA AGCTGCG
J23SpTA_5_                          CGG GGATGTAAGC GGGAAGGGAA ACCGACCCGT TTAGTCTGCC GCTCCCAACA GCCCGAGTTT CTAGCCCCTT GAGTAAGGGG CAAGAAACCT AAGCCGCCTG GTGGATGGCT CGGCTCGGGG CGCTGACGAA GGACGCGTGG ACGCGTGGCA AGCTGC
J23SpTA_17                          CGG GGATGTAAGC GGGAAGGGAA ACCGACCCGT TTAGTCTGCC GCTCCCAACA GCCCGAGTTT CTAGCCCCTT GAGTAAGGGG CAAGAAACCT AAGCCGCCTG GTGGATGGCT CGGCTCGGGG CGCTGACGAA GGACGCGTGG ACGCGTGGCA AGCTG
J23SpTA_30            CGGGTTT GATGGGGCGG GGATGTAAGC GGGAAGGGAA ACCGACCCGT TTAGTCTGCC GCTCCCAACA GCCCGAGTTT CTAGCCCCTT GAGTAAGGGG CAAGAAACCT AAGCCGCCTG GTGGATGGCT CGGCTCGGGG CGCTGACGAA
J23SpTA_11           ACGGGTTT GATGGGGCGG GGATGTAAGC GGGAAGGGAA ACCGACCCGT TTAGTCTGCC GCTCCCAACA GCCCGAGTTT CTAGCCCCTT GAGTAAGGGG CAAGAAACCT AAGCCGCCTG GTGGATGGCT CGGCTCGGGG CGCTGACGAA
J23SpTA_24               GTTT GATGGGGCGG GGATGTAAGC GGGAAGGGAA ACCGACCCGT TTAGTCTGCC GCTCCCAACA GCCCGAGTTT CTAGCCCCTT GAGTAAGGGG CAAGAAACCT AAGCCGCCTG GTGGATGGCT CGGCTCGGGG CGCTGACGAA GGACGCGTGG ACG
J23SpTA_7_               GTTT GATGGGGCGG GGATGTAAGC GGGAAGGGAA ACCGACCCGT TTAGTCTGCC GCTCCCAACA GCCCGAGTTT CTAGCCCCTT GAGTAAGGGG CAAGAAACCT AAGCCGCCTG GTGGATGGCT CGGCTCGGGG CGCTGACGAA GGACGCGTGG ACGG
J23SpTA_75              GGTTT GATGGGGCGG GGATGTAAGC GGGAAGGGAA ACCGACCCGT TTAGTCTGCC GCTCCCAACA GCCCGAGTTT CTAGCCCCTT GAGTAAGGGG CAAGAAACCT AAGCCGCCTG GTGGATGGCT CGGCTCGGGG CGCTGACGAA GGACGCGTGG ACG
J23SpTA_5_                TTT GATGGGGCGG GGATGTAAGC GGGAAGGGAA ACCGACCCGT TTAGTCTGCC GCTCCCAACA GCCCGAGTTT CTAGCCCCTT GAGTAAGGGG CAAGAAACCT AAGCCGCCTG GTGGATGGCT CGGCTCGGGG CGCTGACGAA GGACGCGTGG ACG
J23SpTA_19               GTTT GATGGGGCGG GGATGTAAGC GGGAAGGGAA ACCGACCCGT TTAGTCTGCC GCTCCCAACA GCCCGAGTTT CTAGCCCCTT GAGTAAGGGG CAAGAAACCT AAGCCGCCTG GTGGATGGCT CGGCTCGGGG CGCTGACGAA GGACGCGTGG AC
J23SpTA_28          GACGGGTTT GATGGGGCGG GGATGTAAGC GGGAAGGGAA ACCGACCCGT TTAGTCTGCC GCTCCCAACA GCCCGAGTTT CTAGCCCCTT GAGTAAGGGG CAAGAAACCT AAGCCGCCTG GTGGATGGCT CGGCTCGGGG CGCTGACG
J23SpTA_7_          GACGGGTTT GATGGGGCGG GGATGTAAGC GGGAAGGGAA ACCGACCCGT TTAGTCTGCC GCTCCCAACA GCCCGAGTTT CTAGCCCCTT GAGTAAGGGG CAAGAAACCT AAGCCGCCTG GTGGATGGCT CGGCTCGGGG CGCTGAC
J23SpTA_7_           ACGGGTTT GATGGGGCGG GGATGTAAGC GGGAAGGGAA ACCGACCCGT TTAGTCTGCC GCTCCCAACA GCCCGAGTTT CTAGCCCCTT GAGTAAGGGG CAAGAAACCT AAGCCGCCTG GTGGATGGCT CGGCTCGGGG CGCTGACG
J23SpTA_5_                    GATGGGGCGG GGATGTAAGC GGGAAGGGAA ACCGACCCGT TTAGTCTGCC GCTCCCAACA GCCCGAGTTT CTAGCCCCTT GAGTAAGGGG CAAGAAACCT AAGCCGCCTG GTGGATGGCT CGGCTCGGGG CGCTGACGAA GGACGCGTGG ACGCGTGG
J23SpTA_16               GTTT GATGGGGCGG GGATGTAAGC GGGAAGGGAA ACCGACCCGT TTAGTCTGCC GCTCCCAACA GCCCGAGTTT CTAGCCCCTT GAGTAAGGGG CAAGAAACCT AAGCCGCCTG GTGGATGGCT CGGCTCGGGG CGCTGACGAA GGACGCGTG
J23SpTA_44              GGTTT GATGGGGCGG GGATGTAAGC GGGAAGGGAA ACCGACCCGT TTAGTCTGCC GCTCCCAACA GCCCGAGTTT CTAGCCCCTT GAGTAAGGGG CAAGAAACCT AAGCCGCCTG GTGGATGGCT CGGCTCGGGG CGCTGACGAA GGACGCGTG
J23SpTA_16    CGCAGA AGACGGGTTT GATGGGGCGG GGATGTAAGC GGGAAGGGAA ACCGACCCGT TTAGTCTGCC GCTCCCAACA GCCCGAGTTT CTAGCCCCTT GAGTAAGGGG CAAGAAACCT AAGCCGCCTG GTGGATGGCT CGGCTCGGG
J23SpTA_11  CCCGCAGA AGACGGGTTT GATGGGGCGG GGATGTAAGC GGGAAGGGAA ACCGACCCGT TTAGTCTGCC GCTCCCAACA GCCCGAGTTT CTAGCCCCTT GAGTAAGGGG CAAGAAACCT AAGCCGCCTG GTGGATGGCT CGGCTCGGG
J23SpTA_10  CCCGCAGA AGACGGGTTT GATGGGGCGG GGATGTAAGC GGGAAGGGAA ACCGACCCGT TTAGTCTGCC GCTCCCAACA GCCCGAGTTT CTAGCCCCTT GAGTAAGGGG CAAGAAACCT AAGCCGCCTG GTGGATGGCT CGGCTCGGGG
J23SpTA_11   CCGCAGA AGACGGGTTT GATGGGGCGG GGATGTAAGC GGGAAGGGAA ACCGACCCGT TTAGTCTGCC GCTCCCAACA GCCCGAGTTT CTAGCCCCTT GAGTAAGGGG CAAGAAACCT AAGCCGCCTG GTGGATGGCT CGGCTCGGGG
J23SpTA_10    CGCAGA AGACGGGTTT GATGGGGCGG GGATGTAAGC GGGAAGGGAA ACCGACCCGT TTAGTCTGCC GCTCCCAACA GCCCGAGTTT CTAGCCCCTT GAGTAAGGGG CAAGAAACCT AAGCCGCCTG GTGGATGGCT CGGCTCGGGG
J23SpTA_5_  CCCGCAGA AGACGGGTTT GATGGGGCGG GGATGTAAGC GGGAAGGGAA ACCGACCCGT TTAGTCTGCC GCTCCCAACA GCCCGAGTTT CTAGCCCCTT GAGTAAGGGG CAAGAAACCT AAGCCGCCTG GTGGATGGCT CGGCTCGG
J23SpTA_74   CCGCAGA AGACGGGTTT GATGGGGCGG GGATGTAAGC GGGAAGGGAA ACCGACCCGT TTAGTCTGCC GCTCCCAACA GCCCGAGTTT CTAGCCCCTT GAGTAAGGGG CAAGAAACCT AAGCCGCCTG GTGGATGGCT CGGCTCGGG
J23SpTA_6_    CGCAGA AGACGGGTTT GATGGGGCGG GGATGTAAGC GGGAAGGGAA ACCGACCCGT TTAGTCTGCC GCTCCCAACA GCCCGAGTTT CTAGCCCCTT GAGTAAGGGG CAAGAAACCT AAGCCGCCTG GTGGATGGCT CGGCTCGGGG CG
```

```
J23SpTA_16   CCGCAGA AGACGGGTTT GATGGGGCGG GGATGTAAGC GGGAAGGGAA ACCGACCCGT TTAGTCTGCC GCTCCCAACA GCCCGAGTTT CTAGCCCCTT GAGTAAGGGG CAAGAAACCT AAGCCGCCTG GTGGATGGCT CGGCTCGG
J23SpTA_5_     GCAGA AGACGGGTTT GATGGGGCGG GGATGTAAGC GGGAAGGGAA ACCGACCCGT TTAGTCTGCC GCTCCCAACA GCCCGAGTTT CTAGCCCCTT GAGTAAGGGG CAAGAAACCT AAGCCGCCTG GTGGATGGCT CGGCTCGGGG
J23SpTA_13  CCCGCAGA AGACGGGTTT GATGGGGCGG GGATGTAAGC GGGAAGGGAA ACCGACCCGT TTAGTCTGCC GCTCCCAACA GCCCGAGTTT CTAGCCCCTT GAGTAAGGGG CAAGAAACCT AAGCCGCCTG GTGGATGGCT CGGCTCG
J23SpTA_36                CGGGTTT GATGGGGCGG GGATGTAAGC GGGAAGGGAA ACCGACCCGT TTAGTCTGCC GCTCCCAACA GCCCGAGTTT CTAGCCCCTT GAGTAAGGGG CAAGAAACCT AAGCCGCCTG GTGGATGGCT CGGCTCGGGG CGCTGACG
J23SpTA_8_                CGGGTTT GATGGGGCGG GGATGTAAGC GGGAAGGGAA ACCGACCCGT TTAGTCTGCC GCTCCCAACA GCCCGAGTTT CTAGCCCCTT GAGTAAGGGG CAAGAAACCT AAGCCGCCTG GTGGATGGCT CGGCTCGGGG CGCTGACGAG
J23SpTA_8_                   GTTT GATGGGGCGG GGATGTAAGC GGGAAGGGAA ACCGACCCGT TTAGTCTGCC GCTCCCAACA GCCCGAGTTT CTAGCCCCTT GAGTAAGGGG CAAGAAACCT AAGCCGCCTG GTGGATGGCT CGGCTCGGGG CGCTGACGAA
J23SpTA_17                        GATGGGGCGG GGATGTAAGC GGGAAGGGAA ACCGACCCGT TTAGTCTGCC GCTCCCAACA GCCCGAGTTT CTAGCCCCTT GAGTAAGGGG CAAGAAACCT AAGCCGCCTG GTGGATGGCT CGGCTCGGGG CGCTGACGAA GGACGCGTGG ACG
J23SpTA_5_                      T GATGGGGCGG GGATGTAAGC GGGAAGGGAA ACCGACCCGT TTAGTCTGCC GCTCCCAACA GCCCGAGTTT CTAGCCCCTT GAGTAAGGGG CAAGAAACCT AAGCCGCCTG GTGGATGGCT CGGCTCGGGG CGCTGACGAA GGACGCGTGG ACG
J23SpTA_6_                      T GATGGGGCGG GGATGTAAGC GGGAAGGGAA ACCGACCCGT TTAGTCTGCC GCTCCCAACA GCCCGAGTTT CTAGCCCCTT GAGTAAGGGG CAAGAAACCT AAGCCGCCTG GTGGATGGCT CGGCTCGGGG CGCTGACGAA GGACGCGTGG AC
J23SpTA_19                CGGGTTT GATGGGGCGG GGATGTAAGC GGGAAGGGAA ACCGACCCGT TTAGTCTGCC GCTCCCAACA GCCCGAGTTT CTAGCCCCTT GAGTAAGGGG CAAGAAACCT AAGCCGCCTG GTGGATGGCT CGGCTCGGGG CGCTGAC
J23SpTA_16                                            C GGGAAGGGAA ACCGACCCGT TTAGTCTGCC GCTCCCAACA GCCCGAGTTT CTAGCCCCTT GAGTAAGGGG CAAGAAACCT AAGCCGCCTG GTGGATGGCT CGGCTCGGGG CGCTGACGAA GGACGCGTGG ACGCGTGGCA AGCTGCGATA AGCCCCGGTG AG
J23SpTA_6_                                            C GGGAAGGGAA ACCGACCCGT TTAGTCTGCC GCTCCCAACA GCCCGAGTTT CTAGCCCCTT GAGTAAGGGG CAAGAAACCT AAGCCGCCTG GTGGATGGCT CGGCTCGGGG CGCTGACGAA GGACGCGTGG ACGCGTGGCA AGCTGCGATA AGCCCCGGTG AGGCGCAG
J23SpTA_5_                                                 GGGAA ACCGACCCGT TTAGTCTGCC GCTCCCAACA GCCCGAGTTT CTAGCCCCTT GAGTAAGGGG CAAGAAACCT AAGCCGCCTG GTGGATGGCT CGGCTCGGGG CGCTGACGAA GGACGCGTGG ACGCGTGGCA AGCTGCGATA AGCCCCGGTG AG
J23SpTA_64      CAGA AGACGGGTTT GATGGGGCGG GGATGTAAGC GGGAAGGGAA ACCGACCCGT TTAGTCTGCC GCTCCCAACA GCCCGAGTTT CTAGCCCCTT GAGTAAGGGG CAAGAAACCT AAGCCGCCTG GTGGATGGCT CGGCTCGGG
J23SpTA_22     GCAGA AGACGGGTTT GATGGGGCGG GGATGTAAGC GGGAAGGGAA ACCGACCCGT TTAGTCTGCC GCTCCCAACA GCCCGAGTTT CTAGCCCCTT GAGTAAGGGG CAAGAAACCT AAGCCGCCTG GTGGATGGCT CGGCTCGGG
J23SpTA_7_     GCAGA AGACGGGTTT GATGGGGCGG GGATGTAAGC GGGAAGGGAA ACCGACCCGT TTAGTCTGCC GCTCCCAACA GCCCGAGTTT CTAGCCCCTT GAGTAAGGGG CAAGAAACCT AAGCCGCCTG GTGGATGGCT CGGCTCGG
J23SpTA_7_    CGCAGA AGACGGGTTT GATGGGGCGG GGATGTAAGC GGGAAGGGAA ACCGACCCGT TTAGTCTGCC GCTCCCAACA GCCCGAGTTT CTAGCCCCTT GAGTAAGGGG CAAGAAACCT AAGCCGCCTG GTGGATGGCT CGGCCCGGG
J23SpTA_6_      CAGA AGACGGGTTT GATGGGGCGG GGATGTAAGC GGGAAGGGAA ACCGACCCGT TTAGTCTGCC GCTCCCAACA GCCCGAGTTT CTAGCCCCTT GAGTAAGGGG CAAGAAACCT AAGCCGCCTG GTGGATGGCT CGGCTCGGGG
J23SpTA_7_    CGCAGA AGACGGGTTT GATGGGGCGG GGATGTAAGC GGGAAGGGAA ACCGACCCGT TTAGTCTGCC GCTCCCAACA GCCCGAGTTT CTAGCCCCTT GAGTAAGGGG CAAGAAACCT AAGCCGCCTG GTGGATGGCT CGGCTTGGG
J23SpTA_30    CGCAGA AGACGGGTTT GATGGGGCGG GGATGTAAGC GGGAAGGGAA ACCGACCCGT TTAGTCTGCC GCTCCCAACA GCCCGAGTTT CTAGCCCCTT GAGTAAGGGG CAAGAAACCT AAGCCGCCTG GTGGATGGCT CGGCTCGG
J23SpTA_12      CAGA AGACGGGTTT GATGGGGCGG GGATGTAAGC GGGAAGGGAA ACCGACCCGT TTAGTCTGCC GCTCCCAACA GCCCGAGTTT CTAGCCCCTT GAGTAAGGGG CAAGAAACCT AAGCCGCCTG GTGGATGGCT CGGCTCGG
J23SpTA_20       AGA AGACGGGTTT GATGGGGCGG GGATGTAAGC GGGAAGGGAA ACCGACCCGT TTAGTCTGCC GCTCCCAACA GCCCGAGTTT CTAGCCCCTT GAGTAAGGGG CAAGAAACCT AAGCCGCCTG GTGGATGGCT CGGCTCGGG
J23SpTA_35                               CGG GGATGTAAGC GGGAAGGGAA ACCGACCCGT TTAGTCTGCC GCTCCCAACA GCCCGAGTTT CTAGCCCCTT GAGTAAGGGG CAAGAAACCT AAGCCGCCTG GTGGATGGCT CGGCTCGGGG CGCTGACGAA GGACGCGTGG ACGCGTGGC
J23SpTA_16                             GGCGG GGATGTAAGC GGGAAGGGAA ACCGACCCGT TTAGTCTGCC GCTCCCAACA GCCCGAGTTT CTAGCCCCTT GAGTAAGGGG CAAGAAACCT AAGCCGCCTG GTGGATGGCT CGGCTCGGGG CGCTGACGAA GGACGCGTGG ACGCGTGG
J23SpTA_5_                             GGCGG GGATGTAAGC GGGAAGGGAA ACCGACCCGT TTAGTCTGCC GCTCCCAACA GCCCGAGTTT CTAGCCCCTT GAGTAAGGGG CAAGAAACCT AAGCCGCCTG GTGGATGGCT CGGCTCGGGG CGCTGACGAA GGACGCGTGG ACGCGTGGC
J23SpTA_6_                           GGGGCGG GGATGTAAGC GGGAAGGGAA ACCGACCCGT TTAGTCTGCC GCTCCCAACA GCCCGAGTTT CTAGCCCCTT GAGTAAGGGG CAAGAAACCT AAGCCGCCTG GTGGATGGCT CGGCTCGGGG CGCTGACGAA GGACGCGTGG ACGCGTGG
J23SpTA_11                               CGG GGATGTAAGC GGGAAGGGAA ACCGACCCGT TTAGTCTGCC GCTCCCAACA GCCCGAGTTT CTAGCCCCTT GAGTAAGGGG CAAGAAACCT AAGCCGCCTG GTGGATGGCT CGGCTCGGGG CGCTGACGAA GGACGCGTGG ACGCGTGGCA AG
J23SpTA_6_                               CGG GGATGTAAGC GGGAAGGGAA ACCGACCCGT TTAGTCTGCC GCTCCCAACA GCCCGAGTTT CTAGCCCCTT GAGTAAGGGG CAAGAAACCT AAGCCGCCTG GTGGATGGCT CGGCTCGGGG CGCTGACGAA GGACGCGTGG ACGCGTGGCA AGC
J23SpTA_6_                                GG GGATGTAAGC GGGAAGGGAA ACCGACCCGT TTAGTCTGCC GCTCCCAACA GCCCGAGTTT CTAGCCCCTT GAGTAAGGGG CAAGAAACCT AAGCCGCCTG GTGGATGGCT CGGCTCGGGG CGCTGACGAA GGACGCGTGG ACGCGTGGCA AG
J23SpTA_11                   GTTT GATGGGGCGG GGATGTAAGC GGGAAGGGAA ACCGACCCGT TTAGTCTGCC GCTCCCAACA GCCCGAGTTT CTAGCCCCTT GAGTAAGGGG CAAGAAACCT AAGCCGCCTG GTGGATGGCT CGGCTCGGGG CGCTGACG
J23SpTA_6_                   GTTT GATGGGGCGG GGATGTAAGC GGGAAGGGAA ACCGACCCGT TTAGTCTGCC GCTCCCAACA GCCCGAGTTT CTAGCCCCTT GAGTAAGGGG CAAGAAACCT AAGCCGCCTG GTGGATGGCT CGGCTCGGGG CGCTGAC
J23SpTA_17                        GATGGGGCGG GGATGTAAGC GGGAAGGGAA ACCGACCCGT TTAGTCTGCC GCTCCCAACA GCCCGAGTTT CTAGCCCCTT GAGTAAGGGG CAAGAAACCT AAGCCGCCTG GTGGATGGCT CGGCTCGGGG CGCTGACGAA GGACGCGTG
J23SpTA_11                      T GATGGGGCGG GGATGTAAGC GGGAAGGGAA ACCGACCCGT TTAGTCTGCC GCTCCCAACA GCCCGAGTTT CTAGCCCCTT GAGTAAGGGG CAAGAAACCT AAGCCGCCTG GTGGATGGCT CGGCTCGGGG CGCTGACGAA GGACGCGTG
J23SpTA_20                           GGGGCGG GGATGTAAGC GGGAAGGGAA ACCGACCCGT TTAGTCTGCC GCTCCCAACA GCCCGAGTTT CTAGCCCCTT GAGTAAGGGG CAAGAAACCT AAGCCGCCTG GTGGATGGCT CGGCTCGGGG CGCTGACGAA GGACGCGTGGACGC
J23SpTA_7_                          TGGGGCGG GGATGTAAGC GGGAAGGGAA ACCGACCCGT TTAGTCTGCC GCTCCCAACA GCCCGAGTTT CTAGCCCCTT GAGTAAGGGG CAAGAAACCT AAGCCGCCTG GTGGATGGCT CGGCTCGGGG CGCTGACGAA GGACGCGTGGACGC
J23SpTA_6_                          TGGGGCGG GGATGTAAGC GGGAAGGGAA ACCGACCCGT TTAGTCTGCC GCTCCCAACA GCCCGAGTTT CTAGCCCCTT GAGTAAGGGG CAAGAAACCT AAGCCGCCTG GTGGATGGCT CGGCTCGGGG CGCTGACGAA GGACGCGTGG ACG
J23SpTA_10                           GGGGCGG GGATGTAAGC GGGAAGGGAA ACCGACCCGT TTAGTCTGCC GCTCCCAACA GCCCGAGTTT CTAGCCCCTT GAGTAAGGGG CAAGAAACCT AAGCCGCCTG GTGGATGGCT CGGCTCGGGG CGCTGACGAA GGACGCGTGG ACGCG
J23SpTA_15                               CGG GGATGTAAGC GGGAAGGGAA ACCGACCCGT TTAGTCTGCC GCTCCCAACA GCCCGAGTTT CTAGCCCCTT GAGTAAGGGG CAAGAAACCT AAGCCGCCTG GTGGATGGCT CGGCTCGGGG CGCTGACGAA GGACGCGTGG ACGCGTGG
J23SpTA_10                               CGG GGATGTAAGC GGGAAGGGAA ACCGACCCGT TTAGTCTGCC GCTCCCAACA GCCCGAGTTT CTAGCCCCTT GAGTAAGGGG CAAGAAACCT AAGCCGCCTG GTGGATGGCT CGGCTCGGGG CGCTGACGAA GGACGCGTGG ACGCGTG
J23SpTA_6_                             GGCGG GGATGTAAGC GGGAAGGGAA ACCGACCCGT TTAGTCTGCC GCTCCCAACA GCCCGAGTTT CTAGCCCCTT GAGTAAGGGG CAAGAAACCT AAGCCGCCTG GTGGATGGCT CGGCTCGGGG CGCTGACGAA GGACGCGTGG ACGCGTG
J23SpTA_5_                              GCGG GGATGTAAGC GGGAAGGGAA ACCGACCCGT TTAGTCTGCC GCTCCCAACA GCCCGAGTTT CTAGCCCCTT GAGTAAGGGG CAAGAAACCT AAGCCGCCTG GTGGATGGCT CGGCTCGGGG CGCTGACGAA GGACGCGTGG ACGCGTG
J23SpTA_16                                GG GGATGTAAGC GGGAAGGGAA ACCGACCCGT TTAGTCTGCC GCTCCCAACA GCCCGAGTTT CTAGCCCCTT GAGTAAGGGG CAAGAAACCT AAGCCGCCTG GTGGATGGCT CGGCTCGGGG CGCTGACGAA GGACGCGTGG ACGCGTGG
J23SpTA_5_                               CGG GGATGTAAGC GGGAAGGGAA ACCGACCCGT TTAGTCTGCC GCTCCCAACA GCCCGAGTTT CTAGCCCCTT GAGTAAGGGG CAAGAAACCT AAGCCGCCTG GTGGATGGCT CGGCTCGGGG CGCTGACGAA GGACGCGTGG ACGCATGG
J23SpTA_23    CGCAGA AGACGGGTTT GATGGGGCGG GGATGTAAGC GGGAAGGGAA ACCGACCCGT TTAGTCTGCC GCTCCCAACA GCCCGAGTTT CTAGCCCCTT GAGTAAGGGG CAAGAAACCT AAGCCGCCTG GTGGATGGCT CGGCTCG
J23SpTA_10   CCGCAGA AGACGGGTTT GATGGGGCGG GGATGTAAGC GGGAAGGGAA ACCGACCCGT TTAGTCTGCC GCTCCCAACA GCCCGAGTTT CTAGCCCCTT GAGTAAGGGG CAAGAAACCT AAGCCGCCTG GTGGATGGCT CGGCTCG
J23SpTA_5_  CCCGCAGA AGACGGGTTT GATGGGGCGG GGATGTAAGC GGGAAGGGAA ACCGACCCGT TTAGTCTGCC GCTCCCAACA GCCCGAGTTT CTAGCCCCTT GAGTAAGGGG CAAGAAACCT AAGCCGCCTG GTGGATGGCT CGGCT
J23SpTA_9_   CCGCAGA AGACGGGTTT GATGGGGCGG GGATGTAAGC GGGAAGGGAA ACCGACCCGT TTAGTCTGCC GCTCCCAACA GCCCGAGTTT CTAGCCCCTT GAGTAAGGGG CAAGAAACCT AAGCCGCCTG GTGGATGGCT CGGC
J23SpTA_20    CGCAGA AGACGGGTTT GATGGGGCGG GGATGTAAGC GGGAAGGGAA ACCGACCCGT TTAGTCTGCC GCTCCCAACA GCCCGAGTTT CTAGCCCCTT GAGTAAGGGG CAAGAAACCT AAGCCGCCTG GTGGATGGCT CGGCT
J23SpTA_10      CAGA AGACGGGTTT GATGGGGCGG GGATGTAAGC GGGAAGGGAA ACCGACCCGT TTAGTCTGCC GCTCCCAACA GCCCGAGTTT CTAGCCCCTT GAGTAAGGGG CAAGAAACCT AAGCCGCCTG GTGGATGGCT CGGCTCG
J23SpTA_12    CGCAGA AGACGGGTTT GATGGGGCGG GGATGTAAGC GGGAAGGGAA ACCGACCCGT TTAGTCTGCC GCTCCCAACA GCCCGAGTTT CTAGCCCCTT GAGTAAGGGG CAAGAAACCT AAGCCGCCTG GTGGATGGCT CGGC
J23SpTA_12   CCGCAGA AGACGGGTTT GATGGGGCGG GGATGTAAGC GGGAAGGGAA ACCGACCCGT TTAGTCTGCC GCTCCCAACA GCCCGAGTTT CTAGCCCCTT GAGTAAGGGG CAAGAAACCT AAGCCGCCTG GTGGATGGCT CGG
J23SpTA_9_  CCCGCAGA AGACGGGTTT GATGGGGCGG GGATGTAAGC GGGAAGGGAA ACCGACCCGT TTAGTCTGCC GCTCCCAACA GCCCGAGTTT CTAGCCCCTT GAGTAAGGGG CAAGAAACCT AAGCCGCCTG GTGGATGGCT CGG
J23SpTA_16           AGACGGGTTT GATGGGGCGG GGATGTAAGC GGGAAGGGAA ACCGACCCGT TTAGTCTGCC GCTCCCAACA GCCCGAGTTT CTAGCCCCTT GAGTAAGGGG CAAGAAACCT AAGCCGCCTG GTGGATGGCT CGGCTCGGG
J23SpTA_7_        GA AGACGGGTTT GATGGGGCGG GGATGTAAGC GGGAAGGGAA ACCGACCCGT TTAGTCTGCC GCTCCCAACA GCCCGAGTTT CTAGCCCCTT GAGTAAGGGG CAAGAAACCT AAGCCGCCTG GTGGATGGCT CGGCTCGGG
J23SpTA_16                          TGGGGCGG GGATGTAAGC GGGAAGGGAA ACCGACCCGT TTAGTCTGCC GCTCCCAACA GCCCGAGTTT CTAGCCCCTT GAGTAAGGGG CAAGAAACCT AAGCCGCCTG GTGGATGGCT CGGCTCGGGG CGCTGACGAA GGACGCGTG
J23SpTA_12                          TGGGGCGG GGATGTAAGC GGGAAGGGAA ACCGACCCGT TTAGTCTGCC GCTCCCAACA GCCCGAGTTT CTAGCCCCTT GAGTAAGGGG CAAGAAACCT AAGCCGCCTG GTGGATGGCT CGGCTCGGGG CGCTGACGAA GGACGCGTGG AC
J23SpTA_10                           GGGGCGG GGATGTAAGC GGGAAGGGAA ACCGACCCGT TTAGTCTGCC GCTCCCAACA GCCCGAGTTT CTAGCCCCTT GAGTAAGGGG CAAGAAACCT AAGCCGCCTG GTGGATGGCT CGGCTCGGGG CGCTGACGAA GGACGCGTGG AC
J23SpTA_31                           GGGGCGG GGATGTAAGC GGGAAGGGAA ACCGACCCGT TTAGTCTGCC GCTCCCAACA GCCCGAGTTT CTAGCCCCTT GAGTAAGGGG CAAGAAACCT AAGCCGCCTG GTGGATGGCT CGGCTCGGGG CGCTGACGAA GGACGCGTGG ACG
J23SpTA_17                             GGCGG GGATGTAAGC GGGAAGGGAA ACCGACCCGT TTAGTCTGCC GCTCCCAACA GCCCGAGTTT CTAGCCCCTT GAGTAAGGGG CAAGAAACCT AAGCCGCCTG GTGGATGGCT CGGCTCGGGG CGCTGACGAA GGACGCGTGG ACGC
J23SpTA_16                             GGCGG GGATGTAAGC GGGAAGGGAA ACCGACCCGT TTAGTCTGCC GCTCCCAACA GCCCGAGTTT CTAGCCCCTT GAGTAAGGGG CAAGAAACCT AAGCCGCCTG GTGGATGGCT CGGCTCGGGG CGCTGACGAA GGACGCGTGG ACGCG
J23SpTA_8_                            GGGCGG GGATGTAAGC GGGAAGGGAA ACCGACCCGT TTAGTCTGCC GCTCCCAACA GCCCGAGTTT CTAGCCCCTT GAGTAAGGGG CAAGAAACCT AAGCCGCCTG GTGGATGGCT CGGCTCGGGG CGCTGACGAA GGACGCGTGG ACGCG
J23SpTA_8_                            GGGCGG GGATGTAAGC GGGAAGGGAA ACCGACCCGT TTAGTCTGCC GCTCCCAACA GCCCGAGTTT CTAGCCCCTT GAGTAAGGGG CAAGAAACCT AAGCCGCCTG GTGGATGGCT CGGCTCGGGG CGCTGACGAA GGACGCGTGG ACGC
J23SpTA_8_                            GGGCGG GGATGTAAGC GGGAAGGGAA ACCGACCCGT TTAGTCTGCC GCTCCCAACA GCCCGAGTTT CTAGCCCCTT GAGTAAGGGG CAAGAAACCT AAGCCGCCTG GTGGATGGCT CGGCTCGGGG CGCTGACGAA GGACGCGTGG ACG
J23SpTA_8_                              GCGG GGATGTAAGC GGGAAGGGAA ACCGACCCGT TTAGTCTGCC GCTCCCAACA GCCCGAGTTT CTAGCCCCTT GAGTAAGGGG CAAGAAACCT AAGCCGCCTG GTGGATGGCT CGGCTCGGGG CGCTGACGAA GGACGCGTGG ACGCG
J23SpTA_40                               CGG GGATGTAAGC GGGAAGGGAA ACCGACCCGT TTAGTCTGCC GCTCCCAACA GCCCGAGTTT CTAGCCCCTT GAGTAAGGGG CAAGAAACCT AAGCCGCCTG GTGGATGGCT CGGCTCGGGG CGCTGACGAA GGACGCGTGG ACGC
J23SpTA_23                               CGG GGATGTAAGC GGGAAGGGAA ACCGACCCGT TTAGTCTGCC GCTCCCAACA GCCCGAGTTT CTAGCCCCTT GAGTAAGGGG CAAGAAACCT AAGCCGCCTG GTGGATGGCT CGGCTCGGGG CGCTGACGAA GGACGCGTGG ACGCG
J23SpTA_6_                               CGG GGATGTAAGC GGGAAGGGAA ACCGACCCGT TTAGTCTGCC GCTCCCAACA GCCCGAGTTT CTAGCCCCTT GAGTAAGGGG CAAGAAACCT AAGCCGCCTG GTGGATGGCT CGGCTCGGGG CGCTGACGAA GGACGCGTGG ACGCGT
J23SpTA_5_                               CGG GGATGTAAGC GGGAAGGGAA ACCGACCCGT TTAGTCTGCC GCTCCCAACA GCCCGAGTTT CTAGCCCCTT GAGTAAGGGG CAAGAAACCT AAGCCGCCTG GTGGATGGCT CGGCTCGGGG CGCTGACGAA GGACGCGTGG ACGCATG
J23SpTA_31                                GG GGATGTAAGC GGGAAGGGAA ACCGACCCGT TTAGTCTGCC GCTCCCAACA GCCCGAGTTT CTAGCCCCTT GAGTAAGGGG CAAGAAACCT AAGCCGCCTG GTGGATGGCT CGGCTCGGGG CGCTGACGAA GGACGCGTGG ACGCG
J23SpTA_21                              GCGG GGATGTAAGC GGGAAGGGAA ACCGACCCGT TTAGTCTGCC GCTCCCAACA GCCCGAGTTT CTAGCCCCTT GAGTAAGGGG CAAGAAACCT AAGCCGCCTG GTGGATGGCT CGGCTCGGGG CGCTGACGAA GGACGCGTGG ACGC
J23SpTA_11                               CGG GGATGTAAGC GGGAAGGGAA ACCGACCCGT TTAGTCTGCC GCTCCCAACA GCCCGAGTTT CTAGCCCCTT GAGTAAGGGG CAAGAAACCT AAGCCGCCTG GTGGATGGCT CGGCTCGGGG CGCTGACGAA GGACGCGTGG ACGCC
J23SpTA_13                                GG GGATGTAAGC GGGAAGGGAA ACCGACCCGT TTAGTCTGCC GCTCCCAACA GCCCGAGTTT CTAGCCCCTT GAGTAAGGGG CAAGAAACCT AAGCCGCCTG GTGGATGGCT CGGCTCGGGG CGCTGACGAA GGACGCGTGG ACGCGTG
J23SpTA_13                                 G GGATGTAAGC GGGAAGGGAA ACCGACCCGT TTAGTCTGCC GCTCCCAACA GCCCGAGTTT CTAGCCCCTT GAGTAAGGGG CAAGAAACCT AAGCCGCCTG GTGGATGGCT CGGCTCGGGG CGCTGACGAA GGACGCGTGG ACGCGTGG
J23SpTA_5_                                 G GGATGTAAGC GGGAAGGGAA ACCGACCCGT TTAGTCTGCC GCTCCCAACA GCCCGAGTTT CTAGCCCCTT GAGTAAGGGG CAAGAAACCT AAGCCGCCTG GTGGATGGCT CGGCTCGGGG CGCTGACGAA GGACGCGTGG ACGCGTGGC
J23SpTA_9_                                 G GGATGTAAGC GGGAAGGGAA ACCGACCCGT TTAGTCTGCC GCTCCCAACA GCCCGAGTTT CTAGCCCCTT GAGTAAGGGG CAAGAAACCT AAGCCGCCTG GTGGATGGCT CGGCTCGGGG CGCTGACGAA GGACGCGTGG ACGCGTG
J23SpTA_9_                                   GGATGTAAGC GGGAAGGGAA ACCGACCCGT TTAGTCTGCC GCTCCCAACA GCCCGAGTTT CTAGCCCCTT GAGTAAGGGG CAAGAAACCT AAGCCGCCTG GTGGATGGCT CGGCTCGGGG CGCTGACGAA GGACGCGTGG ACGCGTGG
J23SpTA_22            GACGGGTTT GATGGGGCGG GGATGTAAGC GGGAAGGGAA ACCGACCCGT TTAGTCTGCC GCTCCCAACA GCCCGAGTTT CTAGCCCCTT GAGTAAGGGG CAAGAAACCT AAGCCGCCTG GTGGATGGCT CGGCTCGGG
J23SpTA_12             ACGGGTTT GATGGGGCGG GGATGTAAGC GGGAAGGGAA ACCGACCCGT TTAGTCTGCC GCTCCCAACA GCCCGAGTTT CTAGCCCCTT GAGTAAGGGG CAAGAAACCT AAGCCGCCTG GTGGATGGCT CGGCTCGGG
J23SpTA_38                             GGCGG GGATGTAAGC GGGAAGGGAA ACCGACCCGT TTAGTCTGCC GCTCCCAACA GCCCGAGTTT CTAGCCCCTT GAGTAAGGGG CAAGAAACCT AAGCCGCCTG GTGGATGGCT CGGCTCGGGG CGCTGACGAA GGACGCGTGG ACG
J23SpTA_24                           GGGGCGG GGATGTAAGC GGGAAGGGAA ACCGACCCGT TTAGTCTGCC GCTCCCAACA GCCCGAGTTT CTAGCCCCTT GAGTAAGGGG CAAGAAACCT AAGCCGCCTG GTGGATGGCT CGGCTCGGGG CGCTGACGAA GGACGCGTG
J23SpTA_11                            GGGCGG GGATGTAAGC GGGAAGGGAA ACCGACCCGT TTAGTCTGCC GCTCCCAACA GCCCGAGTTT CTAGCCCCTT GAGTAAGGGG CAAGAAACCT AAGCCGCCTG GTGGATGGCT CGGCTCGGGG CGCTGACGAA GGACGCGTGG AC
J23SpTA_8_                            GGGCGG GGATGTAAGC GGGAAGGGAA ACCGACCCGT TTAGTCTGCC GCTCCCAACA GCCCGAGTTT CTAGCCCCTT GAGTAAGGGG CAAGAAACCT AAGCCGCCTG GTGGATGGCT CGGCTCGGGG CGCTGACGAA GGACGCGTG
J23SpTA_19                             GGCGG GGATGTAAGC GGGAAGGGAA ACCGACCCGT TTAGTCTGCC GCTCCCAACA GCCCGAGTTT CTAGCCCCTT GAGTAAGGGG CAAGAAACCT AAGCCGCCTG GTGGATGGCT CGGCTCGGGG CGCTGACGAA GGACGCGTGG AC
J23SpTA_62                               CGG GGATGTAAGC GGGAAGGGAA ACCGACCCGT TTAGTCTGCC GCTCCCAACA GCCCGAGTTT CTAGCCCCTT GAGTAAGGGG CAAGAAACCT AAGCCGCCTG GTGGATGGCT CGGCTCGGGG CGCTGACGAA GGACGCGTGG ACG
J23SpTA_35                              GCGG GGATGTAAGC GGGAAGGGAA ACCGACCCGT TTAGTCTGCC GCTCCCAACA GCCCGAGTTT CTAGCCCCTT GAGTAAGGGG CAAGAAACCT AAGCCGCCTG GTGGATGGCT CGGCTCGGGG CGCTGACGAA GGACGCGTGG ACG
J23SpTA_28                               CGG GGATGTAAGC GGGAAGGGAA ACCGACCCGT TTAGTCTGCC GCTCCCAACA GCCCGAGTTT CTAGCCCCTT GAGTAAGGGG CAAGAAACCT AAGCCGCCTG GTGGATGGCT CGGCTCGGGG CGCTGACGAA GGACGCGTGG ACGG
J23SpTA_24                              GCGG GGATGTAAGC GGGAAGGGAA ACCGACCCGT TTAGTCTGCC GCTCCCAACA GCCCGAGTTT CTAGCCCCTT GAGTAAGGGG CAAGAAACCT AAGCCGCCTG GTGGATGGCT CGGCTCGGGG CGCTGACGAA GGACGCGTGGAC
J23SpTA_10    CGCAGA AGACGGGTTT GATGGGGCGG GGATGTAAGC GGGAAGGGAA ACCGACCCGT TTAGTCTGCC GCTCCCAACA GCCCGAGTTT CTAGCCCCTT GAGTAAGGGG CAAGAAACCT AAGCCGCCTG GTGGATGGCT CG
J23SpTA_7_    CGCAGA AGACGGGTTT GATGGGGCGG GGATGTAAGC GGGAAGGGAA ACCGACCCGT TTAGTCTGCC GCTCCCAACA GCCCGAGTTT CTAGCCCCTT GAGTAAGGGG CAAGAAACCT AAGCCGCCTG GTGGATGGCT CGG
J23SpTA_7_      CAGA AGACGGGTTT GATGGGGCGG GGATGTAAGC GGGAAGGGAA ACCGACCCGT TTAGTCTGCC GCTCCCAACA GCCCGAGTTT CTAGCCCCTT GAGTAAGGGG CAAGAAACCT AAGCCGCCTG GTGGATGGCT CGG
J23SpTA_5_    CGCAGA AGACGGGTTT GATGGGGCGG GGATGTAAGC GGGAAGGGAA ACCGACCCGT TTAGTCTGCC GCTCCCAACA GCCCGAGTTT CTAGCCCCTT GAGTAAGGGG CAAGAAACCT AAGCCGCCTG GTGGATGGCT C
J23SpTA_24                              GCGG GGATGTAAGC GGGAAGGGAA ACCGACCCGT TTAGTCTGCC GCTCCCAACA GCCCGAGTTT CTAGCCCCTT GAGTAAGGGG CAAGAAACCT AAGCCGCCTG GTGGATGGCT CGGCTCGGGG CGCTGACGAA GGACGCGTG
J23SpTA_16                             GGCGG GGATGTAAGC GGGAAGGGAA ACCGACCCGT TTAGTCTGCC GCTCCCAACA GCCCGAGTTT CTAGCCCCTT GAGTAAGGGG CAAGAAACCT AAGCCGCCTG GTGGATGGCT CGGCTCGGGG CGCTGACGAA GGACGCGTG
J23SpTA_43                               CGG GGATGTAAGC GGGAAGGGAA ACCGACCCGT TTAGTCTGCC GCTCCCAACA GCCCGAGTTT CTAGCCCCTT GAGTAAGGGG CAAGAAACCT AAGCCGCCTG GTGGATGGCT CGGCTCGGGG CGCTGACGAA GGACGCGTGG AC
J23SpTA_6_                               CGG GGATGTAAGC GGGAAGGGAA ACCGACCCGT TTAGTCTGCC GCTCCCAACA GCCCGAGTTT CTAGCCCCTT GAGTAAGGGG CAAGAAACCT AAGCCGCCTG GTGGATGGCT CGGCTCGGGG CGCTGACGAA GGACGCGTGG ACAC
J23SpTA_44                                GG GGATGTAAGC GGGAAGGGAA ACCGACCCGT TTAGTCTGCC GCTCCCAACA GCCCGAGTTT CTAGCCCCTT GAGTAAGGGG CAAGAAACCT AAGCCGCCTG GTGGATGGCT CGGCTCGGGG CGCTGACGAA GGACGCGTGG ACGC
J23SpTA_42                CGGGTTT GATGGGGCGG GGATGTAAGC GGGAAGGGAA ACCGACCCGT TTAGTCTGCC GCTCCCAACA GCCCGAGTTT CTAGCCCCTT GAGTAAGGGG CAAGAAACCT AAGCCGCCTG GTGGATGGCT CGGCTCGGG
J23SpTA_7_                CGGGTTT GATGGGGCGG GGATGTAAGC GGGAAGGGAA ACCGACCCGT TTAGTCTGCC GCTCCCAACA GCCCGAGTTT CTAGCCCCTT GAGTAAGGGG CAAGAAACCT AAGCCGCCTG GTGGATGGCT CGGCTCGG
J23SpTA_6_              GACGGGTTT GATGGGGCGG GGATGTAAGC GGGAAGGGAA ACCGACCCGT TTAGTCTGCC GCTCCCAACA GCCCGAGTTT CTAGCCCCTT GAGTAAGGGG CAAGAAACCT AAGCCGCCTG GTGGATGGCT CGGCTCG
J23SpTA_41  CCCGCAGA AGACGGGTTT GATGGGGCGG GGATGTAAGC GGGAAGGGAA ACCGACCCGT TTAGTCTGCC GCTCCCAACA GCCCGAGTTT CTAGCCCCTT GAGTAAGGGG CAAGAAACCT AAGCCGCCTG GTGGAT
J23SpTA_36  CCCGCAGA AGACGGGTTT GATGGGGCGG GGATGTAAGC GGGAAGGGAA ACCGACCCGT TTAGTCTGCC GCTCCCAACA GCCCGAGTTT CTAGCCCCTT GAGTAAGGGG CAAGAAACCT AAGCCGCCTG GTGGATG
J23SpTA_11   CCGCAGA AGACGGGTTT GATGGGGCGG GGATGTAAGC GGGAAGGGAA ACCGACCCGT TTAGTCTGCC GCTCCCAACA GCCCGAGTTT CTAGCCCCTT GAGTAAGGGG CAAGAAACCT AAGCCGCCTG GTGGATGG
J23SpTA_9_   CCGCAGA AGACGGGTTT GATGGGGCGG GGATGTAAGC GGGAAGGGAA ACCGACCCGT TTAGTCTGCC GCTCCCAACA GCCCGAGTTT CTAGCCCCTT GAGTAAGGGG CAAGAAACCT AAGCCGCCTG GTGGATGGC
J23SpTA_9_  CCCGCAGA AGACGGGTTT GATGGGGCGG GGATGTAAGC GGGAAGGGAA ACCGACCCGT TTAGTCTGCC GCTCCCAACA GCCCGAGTTT CTAGCCCCTT GAGTAAGGGG CAAGAAACCT AAGCCGCCTG GTGGATGGC
J23SpTA_9_  CCCGCAGA AGACGGGTTT GATGGGGCGG GGATGTAAGC GGGAAGGGAA ACCGACCCGT TTAGTCTGCC GCTCCCAACA GCCCGAGTTT CTAGCCCCTT GAGTAAGGGG CAAGAAACCT AAGCCGCCTG GTGGATGG
J23SpTA_22    CGCAGA AGACGGGTTT GATGGGGCGG GGATGTAAGC GGGAAGGGAA ACCGACCCGT TTAGTCTGCC GCTCCCAACA GCCCGAGTTT CTAGCCCCTT GAGTAAGGGG CAAGAAACCT AAGCCGCCTG GTGGATGGC
J23SpTA_24    CGCAGA AGACGGGTTT GATGGGGCGG GGATGTAAGC GGGAAGGGAA ACCGACCCGT TTAGTCTGCC GCTCCCAACA GCCCGAGTTT CTAGCCCCTT GAGTAAGGGG CAAGAAACCT AAGCCGCCTG GTGGATGG
J23SpTA_19   CCGCAGA AGACGGGTTT GATGGGGCGG GGATGTAAGC GGGAAGGGAA ACCGACCCGT TTAGTCTGCC GCTCCCAACA GCCCGAGTTT CTAGCCCCTT GAGTAAGGGG CAAGAAACCT AAGCCGCCTG GTGGATG
J23SpTA_52                               CGG GGATGTAAGC GGGAAGGGAA ACCGACCCGT TTAGTCTGCC GCTCCCAACA GCCCGAGTTT CTAGCCCCTT GAGTAAGGGG CAAGAAACCT AAGCCGCCTG GTGGATGGCT CGGCTCGGGG CGCTGACGAA GGACGCGTG
J23SpTA_7_                               CGG GGATGTAAGC GGGAAGGGAA ACCGACCCGT TTAGTCTGCC GCTCCCAACA GCCCGAGTTT CTAGCCCCTT GAGTAAGGGG CAAGAAACCT AAGCCGCCTG GTGGATGGCT CGGCTCGGGG CGCTGACGAA GGACGCGTGA G
J23SpTA_57                                GG GGATGTAAGC GGGAAGGGAA ACCGACCCGT TTAGTCTGCC GCTCCCAACA GCCCGAGTTT CTAGCCCCTT GAGTAAGGGG CAAGAAACCT AAGCCGCCTG GTGGATGGCT CGGCTCGGGG CGCTGACGAA GGACGCGTGG ACG
J23SpTA_9_                               CGG GGATGTAAGC GGGAAGGGAA ACCGACCCGT TTAGTCTGCC GCTCCCAACA GCCCGAGTTT CTAGCCCCTT GAGTAAGGGG CAAGAAACCT AAGCCGCCTG GTGGATGGCT CGGCTCGGGG CGCTGACAAA GGG
J23SpTA_6_                               CGG GGATGTAAGC GGGAAGGGAA ACCGACCCGT TTAGTCTGCC GCTCCCAACA GCCCGAGTTT CTAGCCCCTT GAGTAAGGGG CAAGAAACCT AAGCCGCCTG GTGGATGGCT CGGCTCGGGG CGCTGACAAA GG
J23SpTA_11                                            C GGGAAGGGAA ACCGACCCGT TTAGTCTGCC GCTCCCAACA GCCCGAGTTT CTAGCCCCTT GAGTAAGGGG CAAGAAACCT AAGCCGCCTG GTGGATGGCT CGGCTCGGGG CGCTGACGAA GGACGCGTGG ACGCGTGGCA AGCTGCGAT
J23SpTA_8_                                            C GGGAAGGGAA ACCGACCCGT TTAGTCTGCC GCTCCCAACA GCCCGAGTTT CTAGCCCCTT GAGTAAGGGG CAAGAAACCT AAGCCGCCTG GTGGATGGCT CGGCTCGGGG CGCTGACGAA GGACGCGTGG ACGCGTGGCA AGCTGCG
J23SpTA_5_                                            C GGGAAGGGAA ACCGACCCGT TTAGTCTGCC GCTCCCAACA GCCCGAGTTT CTAGCCCCTT GAGTAAGGGG CAAGAAACCT AAGCCGCCTG GTGGATGGCT CGGCTCGGGG CGCTGACGAA GGACGCGTGG ACGCGTGGCA AGCTGC
J23SpTA_8_                                              GGGAAGGGAA ACCGACCCGT TTAGTCTGCC GCTCCCAACA GCCCGAGTTT CTAGCCCCTT GAGTAAGGGG CAAGAAACCT AAGCCGCCTG GTGGATGGCT CGGCTCGGGG CGCTGACGAA GGACGCGTGG ACGCGTGGCA AGCTGCG
J23SpTA_61                                 G GGATGTAAGC GGGAAGGGAA ACCGACCCGT TTAGTCTGCC GCTCCCAACA GCCCGAGTTT CTAGCCCCTT GAGTAAGGGG CAAGAAACCT AAGCCGCCTG GTGGATGGCT CGGCTCGGGG CGCTGACGAA GGACGCGTGG ACG
J23SpTA_36                                 G GGATGTAAGC GGGAAGGGAA ACCGACCCGT TTAGTCTGCC GCTCCCAACA GCCCGAGTTT CTAGCCCCTT GAGTAAGGGG CAAGAAACCT AAGCCGCCTG GTGGATGGCT CGGCTCGGGG CGCTGACGAA GGACGCGTGG ACGC
J23SpTA_22                                 G GGATGTAAGC GGGAAGGGAA ACCGACCCGT TTAGTCTGCC GCTCCCAACA GCCCGAGTTT CTAGCCCCTT GAGTAAGGGG CAAGAAACCT AAGCCGCCTG GTGGATGGCT CGGCTCGGGG CGCTGACGAA GGACGCGTGG ACGCG
J23SpTA_6_                                   GGATGTAAGC GGGAAGGGAA ACCGACCCGT TTAGTCTGCC GCTCCCAACA GCCCGAGTTT CTAGCCCCTT GAGTAAGGGG CAAGAAACCT AAGCCGCCTG GTGGATGGCT CGGCTCGGGG CGCTGACGAA GGACGCGTGG ACGCG
J23SpTA_5_                                   GGATGTAAGC GGGAAGGGAA ACCGACCCGT TTAGTCTGCC GCTCCCAACA GCCCGAGTTT CTAGCCCCTT GAGTAAGGGG CAAGAAACCT AAGCCGCCTG GTGGATGGCT CGGCTCGGGG CGCTGACGAA GGACGCGTGG ACGCGTG
J23SpTA_5_                                    GATGTAAGC GGGAAGGGAA ACCGACCCGT TTAGTCTGCC GCTCCCAACA GCCCGAGTTT CTAGCCCCTT GAGTAAGGGG CAAGAAACCT AAGCCGCCTG GTGGATGGCT CGGCTCGGGG CGCTGACGAA GGACGCGTGG ACGCG
J23SpTA_7_                                   GGATGTAAGC GGGAAGGGAA ACCGACCCGT TTAGTCTGCC GCTCCCAACA GCCCGAGTTT CTAGCCCCTT GAGTAAGGGG CAAGAAACCT AAGCCGCCTG GTGGATGGCT CGGCTCGGGG CGCTGACGAA GGACGCGTGG ACGC
J23SpTA_7_                CGGGTTT GATGGGGCGG GGATGTAAGC GGGAAGGGAA ACCGACCCGT TTAGTCTGCC GCTCCCAACA GCCCGAGTTT CTAGCCCCTT GAGTAAGGGG CAAGAAACCT AAGCCGCCTG GTGGATGGCT CGGCGCGG
J23SpTA_6_                CGGGTTT GATGGGGCGG GGATGTAAGC GGGAAGGGAA ACCGACCCGT TTAGTCTGCC GCTCCCAACA GCCCGAGTTT CTAGCCCCTT GAGTAAGGGG CAAGAAACCT AAGCCGCCTG GTGGATGGCT CGGCTCG
J23SpTA_51                                GG GGATGTAAGC GGGAAGGGAA ACCGACCCGT TTAGTCTGCC GCTCCCAACA GCCCGAGTTT CTAGCCCCTT GAGTAAGGGG CAAGAAACCT AAGCCGCCTG GTGGATGGCT CGGCTCGGGG CGCTGACGAA GGACGCGTGG AC
J23SpTA_5_                               CGG GGATGTAAGC GGGAAGGGAA ACCGACCCGT TTAGTCTGCC GCTCCCAACA GCCCGAGTTT CTAGCCCCTT GAGTAAGGGG CAAGAAACCT AAGCCGCCTG GTGGATGGCT CGGCTCGGGG CGCTGATGAA GG
J23SpTA_5_                               CGG GGATGTAAGC GGGAAGGGAA ACCGACCCGT TTAGTCTGCC GCTCCCAACA GCCCGAGTTT CTAGCCCCTT GAGTAAGGGG CAAGAAACCT AAGCCGCCTG GTGGATGGCT CGGCTCGGGG CGCTGACGGA GG
J23SpTA_9_                                       GTAAGC GGGAAGGGAA ACCGACCCGT TTAGTCTGCC GCTCCCAACA GCCCGAGTTT CTAGCCCCTT GAGTAAGGGG CAAGAAACCT AAGCCGCCTG GTGGATGGCT CGGCTCGGGG CGCTGACGAA GGACGCGTGG ACGCGTGG
J23SpTA_51                                GG GGATGTAAGC GGGAAGGGAA ACCGACCCGT TTAGTCTGCC GCTCCCAACA GCCCGAGTTT CTAGCCCCTT GAGTAAGGGG CAAGAAACCT AAGCCGCCTG GTGGATGGCT CGGCTCGGGG CGCTGACGAA GGACGCGTG
J23SpTA_33                               CGG GGATGTAAGC GGGAAGGGAA ACCGACCCGT TTAGTCTGCC GCTCCCAACA GCCCGAGTTT CTAGCCCCTT GAGTAAGGGG CAAGAAACCT AAGCCGCCTG GTGGATGGCT CGGCTCGGGG CGCTGACGAA
J23SpTA_6_                               CGG GGATGTAAGC GGGAAGGGAA ACCGACCCGT TTAGTCTGCC GCTCCCAACA GCCCGAGTTT CTAGCCCCTT GAGTAAGGGG CAAGAAACCT AAGCCGCCTG GTGGATGGCT CGGCTCGGGG CGCTGACAAA G
J23SpTA_5_                                GG GGATGTAAGC GGGAAGGGAA ACCGACCCGT TTAGTCTGCC GCTCCCAACA GCCCGAGTTT CTAGCCCCTT GAGTAAGGGG CAAGAAACCT AAGCCGCCTG GTGGATGGCT CGGCTCGGGG CGCTGACGAA
J23SpTA_45                                 G GGATGTAAGC GGGAAGGGAA ACCGACCCGT TTAGTCTGCC GCTCCCAACA GCCCGAGTTT CTAGCCCCTT GAGTAAGGGG CAAGAAACCT AAGCCGCCTG GTGGATGGCT CGGCTCGGGG CGCTGACGAA GGACGCGTGG AC
J23SpTA_42                                 G GGATGTAAGC GGGAAGGGAA ACCGACCCGT TTAGTCTGCC GCTCCCAACA GCCCGAGTTT CTAGCCCCTT GAGTAAGGGG CAAGAAACCT AAGCCGCCTG GTGGATGGCT CGGCTCGGGG CGCTGACGAA GGACGCGTG
J23SpTA_27                                   GGATGTAAGC GGGAAGGGAA ACCGACCCGT TTAGTCTGCC GCTCCCAACA GCCCGAGTTT CTAGCCCCTT GAGTAAGGGG CAAGAAACCT AAGCCGCCTG GTGGATGGCT CGGCTCGGGG CGCTGACGAA GGACGCGTGG ACG
J23SpTA_9_                                    GATGTAAGC GGGAAGGGAA ACCGACCCGT TTAGTCTGCC GCTCCCAACA GCCCGAGTTT CTAGCCCCTT GAGTAAGGGG CAAGAAACCT AAGCCGCCTG GTGGATGGCT CGGCTCGGGG CGCTGACGAA GGACGCGTGG ACGC
J23SpTA_22                                   GGATGTAAGC GGGAAGGGAA ACCGACCCGT TTAGTCTGCC GCTCCCAACA GCCCGAGTTT CTAGCCCCTT GAGTAAGGGG CAAGAAACCT AAGCCGCCTG GTGGATGGCT CGGCTCGGGG CGCTGACGAA GGACGCGTGG AC
J23SpTA_16                   GTTT GATGGGGCGG GGATGTAAGC GGGAAGGGAA ACCGACCCGT TTAGTCTGCC GCTCCCAACA GCCCGAGTTT CTAGCCCCTT GAGTAAGGGG CAAGAAACCT AAGCCGCCTG GTGGATGGCT CGGCTCGGG
J23SpTA_7_                                                                                CCCAACA GCCCGAGTTT CTAGCCCCTT GAGTAAGGGG CAAGAAACCT AAGCCGCCTG GTGGATGGCT CGGCTCGGGG CGCTGACGAA GGGCGTGGCA AGCTGCGATA AGCCCCGGTG AGGCGCAGGC AGCCGTCGAG CCGGGGATTC CCGAATGGGA C
J23SpTA_5_                                                                      CCCGT TTAGTCTGCC GCTCCCAACA GCCCGAGTTT CTAGCCCCTT GAGTAAGGGG CAAGAAACCT AAGCCGCCTG GTGGATGGCT CGGCTCGGGG CGCTGACGAA GGACGCGTGG ACGCGTGGCA AGCTGCGATA AGCCCCGGTG AGGCGCAGGC AGCCGTCGAG CCGGGGATTC CCGAATGGGA C
```

```
J23SpTA_5_                                                            CCCGT TTAGTCTGCC GCTCCCAACA GCCCGAGTTT CTAGCCCCTT GAGTAAGGGG CAAGAAACCT AAGCCGCCTG GTGGATGGCT CGGCTCGGGG CGCTGACGAA GGACGCGTGG ACGCGTGGCA AGCTGCGATA AGCCCCGGTG AGGCGCAGGC AGCCGTCGAG CCGGGGATTC CCGAATGGG
J23SpTA_6_              CGGGTTT GATGGGGCGG GGATGTAAGC GGGAAGGGAA ACCGACCCGT TTAGTCTGCC GCTCCCAACA GCCCGAGTTT CTAGCCCCTT GAGTAAGGGG CAAGAAACCT AAGCCGCCTG GTGGATGGCT CGGCT
J23SpTA_14                                     ATGTAAGC GGGAAGGGAA ACCGACCCGT TTAGTCTGCC GCTCCCAACA GCCCGAGTTT CTAGCCCCTT GAGTAAGGGG CAAGAAACCT AAGCCGCCTG GTGGATGGCT CGGCTCGGGG CGCTGACGAA GGACGCGTGG ACG
J23SpTA_8_                                     ATGTAAGC GGGAAGGGAA ACCGACCCGT TTAGTCTGCC GCTCCCAACA GCCCGAGTTT CTAGCCCCTT GAGTAAGGGG CAAGAAACCT AAGCCGCCTG GTGGATGGCT CGGCTCGGGG CGCTGACGAA GGACGCGTGG AC
J23SpTA_8_                                      TGTAAGC GGGAAGGGAA ACCGACCCGT TTAGTCTGCC GCTCCCAACA GCCCGAGTTT CTAGCCCCTT GAGTAAGGGG CAAGAAACCT AAGCCGCCTG GTGGATGGCT CGGCTCGGGG CGCTGACGAA GGACGCGTGG ACG
J23SpTA_9_                                    GATGTAAGC GGGAAGGGAA ACCGACCCGT TTAGTCTGCC GCTCCCAACA GCCCGAGTTT CTAGCCCCTT GAGTAAGGGG CAAGAAACCT AAGCCGCCTG GTGGATGGCT CGGCTCGGGG CGCTGACGAA GGACGCGTGG AC
J23SpTA_30                             CGG GGATGTAAGC GGGAAGGGAA ACCGACCCGT TTAGTCTGCC GCTCCCAACA GCCCGAGTTT CTAGCCCCTT GAGTAAGGGG CAAGAAACCT AAGCCGCCTG GTGGATGGCT CGGCTCGGGG CGCTGACG
J23SpTA_13                             CGG GGATGTAAGC GGGAAGGGAA ACCGACCCGT TTAGTCTGCC GCTCCCAACA GCCCGAGTTT CTAGCCCCTT GAGTAAGGGG CAAGAAACCT AAGCCGCCTG GTGGATGGCT CGGCTCGGGG CGCTGAC
J23SpTA_16                                   GGATGTAAGC GGGAAGGGAA ACCGACCCGT TTAGTCTGCC GCTCCCAACA GCCCGAGTTT CTAGCCCCTT GAGTAAGGGG CAAGAAACCT AAGCCGCCTG GTGGATGGCT CGGCTCGGGG CGCTGACGAA GGACGCGTG
J23SpTA_14                                    GATGTAAGC GGGAAGGGAA ACCGACCCGT TTAGTCTGCC GCTCCCAACA GCCCGAGTTT CTAGCCCCTT GAGTAAGGGG CAAGAAACCT AAGCCGCCTG GTGGATGGCT CGGCTCGGGG CGCTGACGAA GGACGCGTG
J23SpTA_7_                                          AGC GGGAAGGGAA ACCGACCCGT TTAGTCTGCC GCTCCCAACA GCCCGAGTTT CTAGCCCCTT GAGTAAGGGG CAAGAAACCT AAGCCGCCTG GTGGATGGCT CGGCTCGGGG CGCTGACGAA GGACGCGTGG ACGCGTGG
J23SpTA_89  CCCGCAGA AGACGGGTTT GATGGGGCGG GGATGTAAGC GGGAAGGGAA ACCGACCCGT TTAGTCTGCC GCTCCCAACA GCCCGAGTTT CTAGCCCCTT GAGTAAGGGG CAAGAAACCT AAGCCGCCTG GTG
J23SpTA_7_  CCCGCAGA AGACGGGTTT GATGGGGCGG GGATGTAAGC GGGAAGGGAA ACCGACCCGT TTAGTCTGCC GCTCCCAACA GCCCGAGTTT CTAGCCCCTT GAGTAAGGGG CAAGAAACCT AAGCCGCCTG GTGG
J23SpTA_5_   CCGCAGA AGACGGGTTT GATGGGGCGG GGATGTAAGC GGGAAGGGAA ACCGACCCGT TTAGTCTGCC GCTCCCAACA GCCCGAGTTT CTAGCCCCTT GAGTAAGGGG CAAGAAACCT AAGCCGCCTG GTGG
J23SpTA_73    CGCAGA AGACGGGTTT GATGGGGCGG GGATGTAAGC GGGAAGGGAA ACCGACCCGT TTAGTCTGCC GCTCCCAACA GCCCGAGTTT CTAGCCCCTT GAGTAAGGGG CAAGAAACCT AAGCCGCCTG GTGGAT
J23SpTA_32   CCGCAGA AGACGGGTTT GATGGGGCGG GGATGTAAGC GGGAAGGGAA ACCGACCCGT TTAGTCTGCC GCTCCCAACA GCCCGAGTTT CTAGCCCCTT GAGTAAGGGG CAAGAAACCT AAGCCGCCTG GTGGAT
J23SpTA_51    CGCAGA AGACGGGTTT GATGGGGCGG GGATGTAAGC GGGAAGGGAA ACCGACCCGT TTAGTCTGCC GCTCCCAACA GCCCGAGTTT CTAGCCCCTT GAGTAAGGGG CAAGAAACCT AAGCCGCCTG GTGGATG
J23SpTA_7_     GCAGA AGACGGGTTT GATGGGGCGG GGATGTAAGC GGGAAGGGAA ACCGACCCGT TTAGTCTGCC GCTCCCAACA GCCCGAGTTT CTAGCCCCTT GAGTAAGGGG CAAGAAACCT AAGCCGCCTG GTGGATG
J23SpTA_12     GCAGA AGACGGGTTT GATGGGGCGG GGATGTAAGC GGGAAGGGAA ACCGACCCGT TTAGTCTGCC GCTCCCAACA GCCCGAGTTT CTAGCCCCTT GAGTAAGGGG CAAGAAACCT AAGCCGCCTG GTGGAT
J23SpTA_18    CGCAGA AGACGGGTTT GATGGGGCGG GGATGTAAGC GGGAAGGGAA ACCGACCCGT TTAGTCTGCC GCTCCCAACA GCCCGAGTTT CTAGCCCCTT GAGTAAGGGG CAAGAAACCT AAGCCGCCTG GTGG
J23SpTA_88   CCGCAGA AGACGGGTTT GATGGGGCGG GGATGTAAGC GGGAAGGGAA ACCGACCCGT TTAGTCTGCC GCTCCCAACA GCCCGAGTTT CTAGCCCCTT GAGTAAGGGG CAAGAAACCT AAGCCGCCTG GTG
J23SpTA_24      CAGA AGACGGGTTT GATGGGGCGG GGATGTAAGC GGGAAGGGAA ACCGACCCGT TTAGTCTGCC GCTCCCAACA GCCCGAGTTT CTAGCCCCTT GAGTAAGGGG CAAGAAACCT AAGCCGCCTG GTGGAT
J23SpTA_15       AGA AGACGGGTTT GATGGGGCGG GGATGTAAGC GGGAAGGGAA ACCGACCCGT TTAGTCTGCC GCTCCCAACA GCCCGAGTTT CTAGCCCCTT GAGTAAGGGG CAAGAAACCT AAGCCGCCTG GTGGATG
J23SpTA_13      CAGA AGACGGGTTT GATGGGGCGG GGATGTAAGC GGGAAGGGAA ACCGACCCGT TTAGTCTGCC GCTCCCAACA GCCCGAGTTT CTAGCCCCTT GAGTAAGGGG CAAGAAACCT AAGCCGCCTG GTGGATGG
J23SpTA_6_       AGA AGACGGGTTT GATGGGGCGG GGATGTAAGC GGGAAGGGAA ACCGACCCGT TTAGTCTGCC GCTCCCAACA GCCCGAGTTT CTAGCCCCTT GAGTAAGGGG CAAGAAACCT AAGCCGCCTG GTGGATGGC
J23SpTA_13      CAGA AGACGGGTTT GATGGGGCGG GGATGTAAGC GGGAAGGGAA ACCGACCCGT TTAGTCTGCC GCTCCCAACA GCCCGAGTTT CTAGCCCCTT GAGTAAGGGG CAAGAAACCT AAGCCGCCTG GTGGATG
J23SpTA_12                                     ATGTAAGC GGGAAGGGAA ACCGACCCGT TTAGTCTGCC GCTCCCAACA GCCCGAGTTT CTAGCCCCTT GAGTAAGGGG CAAGAAACCT AAGCCGCCTG GTGGATGGCT CGGCTCGGGG CGCTGACGAA GGACGCGTG
J23SpTA_35                                       GTAAGC GGGAAGGGAA ACCGACCCGT TTAGTCTGCC GCTCCCAACA GCCCGAGTTT CTAGCCCCTT GAGTAAGGGG CAAGAAACCT AAGCCGCCTG GTGGATGGCT CGGCTCGGGG CGCTGACGAA GGACGCGTGG AC
J23SpTA_30                                       GTAAGC GGGAAGGGAA ACCGACCCGT TTAGTCTGCC GCTCCCAACA GCCCGAGTTT CTAGCCCCTT GAGTAAGGGG CAAGAAACCT AAGCCGCCTG GTGGATGGCT CGGCTCGGGG CGCTGACGAA GGACGCGTGG ACG
J23SpTA_21                                       GTAAGC GGGAAGGGAA ACCGACCCGT TTAGTCTGCC GCTCCCAACA GCCCGAGTTT CTAGCCCCTT GAGTAAGGGG CAAGAAACCT AAGCCGCCTG GTGGATGGCT CGGCTCGGGG CGCTGACGAA GGACGCGTGG ACGC
J23SpTA_11                                       GTAAGC GGGAAGGGAA ACCGACCCGT TTAGTCTGCC GCTCCCAACA GCCCGAGTTT CTAGCCCCTT GAGTAAGGGG CAAGAAACCT AAGCCGCCTG GTGGATGGCT CGGCTCGGGG CGCTGACGAA GGACGCGTGG ACGCG
J23SpTA_7_                                         AAGC GGGAAGGGAA ACCGACCCGT TTAGTCTGCC GCTCCCAACA GCCCGAGTTT CTAGCCCCTT GAGTAAGGGG CAAGAAACCT AAGCCGCCTG GTGGATGGCT CGGCTCGGGG CGCTGACGAA GGACGCGTGG ACGCG
J23SpTA_5_                                        TAAGC GGGAAGGGAA ACCGACCCGT TTAGTCTGCC GCTCCCAACA GCCCGAGTTT CTAGCCCCTT GAGTAAGGGG CAAGAAACCT AAGCCGCCTG GTGGATGGCT CGGCTCGGGG CGCTGACGAA GGACGCGTGG ACGC
J23SpTA_9_                                        TAAGC GGGAAGGGAA ACCGACCCGT TTAGTCTGCC GCTCCCAACA GCCCGAGTTT CTAGCCCCTT GAGTAAGGGG CAAGAAACCT AAGCCGCCTG GTGGATGGCT CGGCTCGGGG CGCTGACGAA GGACGCGTGG ACG
J23SpTA_7_                                         AAGC GGGAAGGGAA ACCGACCCGT TTAGTCTGCC GCTCCCAACA GCCCGAGTTT CTAGCCCCTT GAGTAAGGGG CAAGAAACCT AAGCCGCCTG GTGGATGGCT CGGCTCGGGG CGCTGACGAA GGACGCGTGG ACGC
J23SpTA_30                                            C GGGAAGGGAA ACCGACCCGT TTAGTCTGCC GCTCCCAACA GCCCGAGTTT CTAGCCCCTT GAGTAAGGGG CAAGAAACCT AAGCCGCCTG GTGGATGGCT CGGCTCGGGG CGCTGACGAA GGACGCGTGG ACGCGTGG
J23SpTA_12                                            C GGGAAGGGAA ACCGACCCGT TTAGTCTGCC GCTCCCAACA GCCCGAGTTT CTAGCCCCTT GAGTAAGGGG CAAGAAACCT AAGCCGCCTG GTGGATGGCT CGGCTCGGGG CGCTGACGAA GGACGCGTGG ACGCGTG
J23SpTA_7_                                              GGGAAGGGAA ACCGACCCGT TTAGTCTGCC GCTCCCAACA GCCCGAGTTT CTAGCCCCTT GAGTAAGGGG CAAGAAACCT AAGCCGCCTG GTGGATGGCT CGGCTCGGGG CGCTGACGAA GGACGCGTGG ACGCGTGG
J23SpTA_10                                                            CCCGT TTAGTCTGCC GCTCCCAACA GCCCGAGTTT CTAGCCCCTT GAGTAAGGGG CAAGAAACCT AAGCCGCCTG GTGGATGGCT CGGCTCGGGG CGCTGACGAA GGACGCGTGG ACGCGTGGCA AGCTGCGATA AGCCCCGGTG AG
J23SpTA_21    CGCAGA AGACGGGTTT GATGGGGCGG GGATGTAAGC GGGAAGGGAA ACCGACCCGT TTAGTCTGCC GCTCCCAACA GCCCGAGTTT CTAGCCCCTT GAGTAAGGGG CAAGAAACCT AAGCCGCCTG GTG
J23SpTA_19     GCAGA AGACGGGTTT GATGGGGCGG GGATGTAAGC GGGAAGGGAA ACCGACCCGT TTAGTCTGCC GCTCCCAACA GCCCGAGTTT CTAGCCCCTT GAGTAAGGGG CAAGAAACCT AAGCCGCCTG GTG
J23SpTA_6_    CGCAGA AGACGGGTTT GATGGGGCGG GGATGTAAGC GGGAAGGGAA ACCGACCCGT TTAGTCTGCC GCTCCCAACA GCCCGAGTTT CTAGCCCCTT GAGTAAGGGG CAAGAAACCT AAGCCGCCTG GT
J23SpTA_43      CAGA AGACGGGTTT GATGGGGCGG GGATGTAAGC GGGAAGGGAA ACCGACCCGT TTAGTCTGCC GCTCCCAACA GCCCGAGTTT CTAGCCCCTT GAGTAAGGGG CAAGAAACCT AAGCCGCCTG GTG
J23SpTA_5_     CCAGA AGACGGGTTT GATGGGGCGG GGATGTAAGC GGGAAGGGAA ACCGACCCGT TTAGTCTGCC GCTCCCAACA GCCCGAGTTT CTAGCCCCTT GAGTAAGGGG CAAGAAACCT AAGCCGCCTG GTG
J23SpTA_11    CGCAGA AGACGGGTTT GATGGGGCGG GGATGTAAGC GGGAAGGGAA ACCGACCCGT TTAGTCTGCC GCTCCCAACA GCCCGAGTTT CTAGCCCCTT GAGTAAGGGG CAAGAAACCT AAGCCGCCTG G
J23SpTA_6_                               G GGATGTAAGC GGGAAGGGAA ACCGACCCGT TTAGTCTGCC GCTCCCAACA GCCCGAGTTT CTAGCCCCTT GAGTAAGGGG CAAGAAACCT AAGCCGCCTG GTGGATGGCT CGGCTCGGGG CGCTGAC
J23SpTA_8_                                              GGGAAGGGAA ACCGACCCGT TTAGTCTGCC GCTCCCAACA GCCCGAGTTT CTAGCCCCTT GAGTAAGGGG CAAGAAACCT AAGCCGCCTG GTGGATGGCT CGGCTCGGGG CGCTGACGAA GGACGCGTGG ACGCGTG
J23SpTA_19                                       GTAAGC GGGAAGGGAA ACCGACCCGT TTAGTCTGCC GCTCCCAACA GCCCGAGTTT CTAGCCCCTT GAGTAAGGGG CAAGAAACCT AAGCCGCCTG GTGGATGGCT CGGCTCGGGG CGCTGACGAA GGACGCGTG
J23SpTA_10                                        TAAGC GGGAAGGGAA ACCGACCCGT TTAGTCTGCC GCTCCCAACA GCCCGAGTTT CTAGCCCCTT GAGTAAGGGG CAAGAAACCT AAGCCGCCTG GTGGATGGCT CGGCTCGGGG CGCTGACGAA GGACGCGTGG AC
J23SpTA_17                                         AAGC GGGAAGGGAA ACCGACCCGT TTAGTCTGCC GCTCCCAACA GCCCGAGTTT CTAGCCCCTT GAGTAAGGGG CAAGAAACCT AAGCCGCCTG GTGGATGGCT CGGCTCGGGG CGCTGACGAA GGACGCGTGG ACG
J23SpTA_14                                          AGC GGGAAGGGAA ACCGACCCGT TTAGTCTGCC GCTCCCAACA GCCCGAGTTT CTAGCCCCTT GAGTAAGGGG CAAGAAACCT AAGCCGCCTG GTGGATGGCT CGGCTCGGGG CGCTGACGAA GGACGCGTGG ACG
J23SpTA_13                                          AGC GGGAAGGGAA ACCGACCCGT TTAGTCTGCC GCTCCCAACA GCCCGAGTTT CTAGCCCCTT GAGTAAGGGG CAAGAAACCT AAGCCGCCTG GTGGATGGCT CGGCTCGGGG CGCTGACGAA GGACGCGTGG ACGC
J23SpTA_5_                                           GC GGGAAGGGAA ACCGACCCGT TTAGTCTGCC GCTCCCAACA GCCCGAGTTT CTAGCCCCTT GAGTAAGGGG CAAGAAACCT AAGCCGCCTG GTGGATGGCT CGGCTCGGGG CGCTGACGAA GGACGCGTGG ACGC
J23SpTA_5_                                           GC GGGAAGGGAA ACCGACCCGT TTAGTCTGCC GCTCCCAACA GCCCGAGTTT CTAGCCCCTT GAGTAAGGGG CAAGAAACCT AAGCCGCCTG GTGGATGGCT CGGCTCGGGG CGCTGACGAA GGACGCGTGG ACGCG
J23SpTA_11            GACGGGTTT GATGGGGCGG GGATGTAAGC GGGAAGGGAA ACCGACCCGT TTAGTCTGCC GCTCCCAACA GCCCGAGTTT CTAGCCCCTT GAGTAAGGGG CAAGAAACCT AAGCCGCCTG GTGGAT
J23SpTA_6_            GACGGGTTT GATGGGGCGG GGATGTAAGC GGGAAGGGAA ACCGACCCGT TTAGTCTGCC GCTCCCAACA GCCCGAGTTT CTAGCCCCTT GAGTAAGGGG CAAGAAACCT AAGCCGCCTG GTGGATG
J23SpTA_12              CGGGTTT GATGGGGCGG GGATGTAAGC GGGAAGGGAA ACCGACCCGT TTAGTCTGCC GCTCCCAACA GCCCGAGTTT CTAGCCCCTT GAGTAAGGGG CAAGAAACCT AAGCCGCCTG GTGGATGGC
J23SpTA_9_              CGGGTTT GATGGGGCGG GGATGTAAGC GGGAAGGGAA ACCGACCCGT TTAGTCTGCC GCTCCCAACA GCCCGAGTTT CTAGCCCCTT GAGTAAGGGG CAAGAAACCT AAGCCGCCTG GTGGATGG
J23SpTA_55    CGCAGA AGACGGGTTT GATGGGGCGG GGATGTAAGC GGGAAGGGAA ACCGACCCGT TTAGTCTGCC GCTCCCAACA GCCCGAGTTT CTAGCCCCTT GAGTAAGGGG CAAGAAACCT AAGCCGCCTG
J23SpTA_22   CCGCAGA AGACGGGTTT GATGGGGCGG GGATGTAAGC GGGAAGGGAA ACCGACCCGT TTAGTCTGCC GCTCCCAACA GCCCGAGTTT CTAGCCCCTT GAGTAAGGGG CAAGAAACCT AAGCCGCCTG
J23SpTA_12  CCCGCAGA AGACGGGTTT GATGGGGCGG GGATGTAAGC GGGAAGGGAA ACCGACCCGT TTAGTCTGCC GCTCCCAACA GCCCGAGTTT CTAGCCCCTT GAGTAAGGGG CAAGAAACCT AAGCCGCCTG
J23SpTA_6_   CCGCAGA AGACGGGTTT GATGGGGCGG GGATGTAAGC GGGAAGGGAA ACCGACCCGT TTAGTCTGCC GCTCCCAACA GCCCGAGTTT CTAGCCCCTT GAGTAAGGGG CAAGAAACCT AAGCCGCCT
J23SpTA_5_     GCAGA AGACGGGTTT GATGGGGCGG GGATGTAAGC GGGAAGGGAA ACCGACCCGT TTAGTCTGCC GCTCCCAACA GCCCGAGTTT CTAGCCCCTT GAGTAAGGGG CAAGAAACCT AAGCCGCCTG
J23SpTA_5_   CCGCAGA AGACGGGTTT GATGGGGCGG GGATGTAAGC GGGAAGGGAA ACCGACCCGT TTAGTCTGCC GCTCCCAACA GCCCGAGTTT CTAGCCCCTT GAGTAAGGGG CAAGAAACCT AAGCCGCC
J23SpTA_12    CGCAGA AGACGGGTTT GATGGGGCGG GGATGTAAGC GGGAAGGGAA ACCGACCCGT TTAGTCTGCC GCTCCCAACA GCCCGAGTTT CTAGCCCCTT GAGTAAGGGG CAAGAAACCT AAGCCGCCT
J23SpTA_36       AGA AGACGGGTTT GATGGGGCGG GGATGTAAGC GGGAAGGGAA ACCGACCCGT TTAGTCTGCC GCTCCCAACA GCCCGAGTTT CTAGCCCCTT GAGTAAGGGG CAAGAAACCT AAGCCGCCTG GTG
J23SpTA_16       AGA AGACGGGTTT GATGGGGCGG GGATGTAAGC GGGAAGGGAA ACCGACCCGT TTAGTCTGCC GCTCCCAACA GCCCGAGTTT CTAGCCCCTT GAGTAAGGGG CAAGAAACCT AAGCCGCCTG GTGGAT
J23SpTA_6_       AGA AGACGGGTTT GATGGGGCGG GGATGTAAGC GGGAAGGGAA ACCGACCCGT TTAGTCTGCC GCTCCCAACA GCCCGAGTTT CTAGCCCCTT GAGTAAGGGG CAAGAAACCT AAGCCGCCTG GTGG
J23SpTA_71                                            C GGGAAGGGAA ACCGACCCGT TTAGTCTGCC GCTCCCAACA GCCCGAGTTT CTAGCCCCTT GAGTAAGGGG CAAGAAACCT AAGCCGCCTG GTGGATGGCT CGGCTCGGGG CGCTGACGAA GGACGCGTGG ACGC
J23SpTA_47                                            C GGGAAGGGAA ACCGACCCGT TTAGTCTGCC GCTCCCAACA GCCCGAGTTT CTAGCCCCTT GAGTAAGGGG CAAGAAACCT AAGCCGCCTG GTGGATGGCT CGGCTCGGGG CGCTGACGAA GGACGCGTGG ACGCG
J23SpTA_11                                              GGGAAGGGAA ACCGACCCGT TTAGTCTGCC GCTCCCAACA GCCCGAGTTT CTAGCCCCTT GAGTAAGGGG CAAGAAACCT AAGCCGCCTG GTGGATGGCT CGGCTCGGGG CGCTGACGAA GGACGCGTGG ACGCG
J23SpTA_6_                                               GGAAGGGAA ACCGACCCGT TTAGTCTGCC GCTCCCAACA GCCCGAGTTT CTAGCCCCTT GAGTAAGGGG CAAGAAACCT AAGCCGCCTG GTGGATGGCT CGGCTCGGGG CGCTGACGAA GGACGCGTGG ACGCG
J23SpTA_6_                                              GGGAAGGGAA ACCGACCCGT TTAGTCTGCC GCTCCCAACA GCCCGAGTTT CTAGCCCCTT GAGTAAGGGG CAAGAAACCT AAGCCGCCTG GTGGATGGCT CGGCTCGGGG CGCTGACGAA GGACGCGTGG ACGC
J23SpTA_9_                                           GC GGGAAGGGAA ACCGACCCGT TTAGTCTGCC GCTCCCAACA GCCCGAGTTT CTAGCCCCTT GAGTAAGGGG CAAGAAACCT AAGCCGCCTG GTGGATGGCT CGGCTCGGGG CGCTGACGAA GGACGCGTGG ACG
J23SpTA_9_                                         AAGC GGGAAGGGAA ACCGACCCGT TTAGTCTGCC GCTCCCAACA GCCCGAGTTT CTAGCCCCTT GAGTAAGGGG CAAGAAACCT AAGCCGCCTG GTGGATGGCT CGGCTCGGGG CGCTGACGAA GGACGCGTG
J23SpTA_7_                                          AGC GGGAAGGGAA ACCGACCCGT TTAGTCTGCC GCTCCCAACA GCCCGAGTTT CTAGCCCCTT GAGTAAGGGG CAAGAAACCT AAGCCGCCTG GTGGATGGCT CGGCTCGGGG CGCTGACGAA GGACGCGTGG AC
J23SpTA_5_                                           GC GGGAAGGGAA ACCGACCCGT TTAGTCTGCC GCTCCCAACA GCCCGAGTTT CTAGCCCCTT GAGTAAGGGG CAAGAAACCT AAGCCGCCTG GTGGATGGCT CGGCTCGGGG CGCTGACGAA GGACGCGTGG AC
J23SpTA_6_                                          AGC GGGAAGGGAA ACCGACCCGT TTAGTCTGCC GCTCCCAACA GCCCGAGTTT CTAGCCCCTT GAGTAAGGGG CAAGAAACCT AAGCCGCCTG GTGGATGGCT CGGCTCGGGG CGCTGACGAA GGACGCGTG
J23SpTA_6_                                               GGAAGGGAA ACCGACCCGT TTAGTCTGCC GCTCCCAACA GCCCGAGTTT CTAGCCCCTT GAGTAAGGGG CAAGAAACCT AAGCCGCCTG GTGGATGGCT CGGCTCGGGG CGCTGACGAA GGACGCGTGG ACGC
J23SpTA_15                                            C GGGAAGGGAA ACCGACCCGT TTAGTCTGCC GCTCCCAACA GCCCGAGTTT CTAGCCCCTT GAGTAAGGGG CAAGAAACCT AAGCCGCCTG GTGGATGGCT CGGCTCGGGG CGCTGACGAA GGACGCGTGG ACG
J23SpTA_7_                                            C GGGAAGGGAA ACCGACCCGT TTAGTCTGCC GCTCCCAACA GCCCGAGTTT CTAGCCCCTT GAGTAAGGGG CAAGAAACCT AAGCCGCCTG GTGGATGGCT CGGCTCGGGG CGCTGACGAA GGACGCGTGA G
J23SpTA_18                                              GGGAAGGGAA ACCGACCCGT TTAGTCTGCC GCTCCCAACA GCCCGAGTTT CTAGCCCCTT GAGTAAGGGG CAAGAAACCT AAGCCGCCTG GTGGATGGCT CGGCTCGGGG CGCTGACGAA GGACGCGTGG ACG
J23SpTA_88                                            C GGGAAGGGAA ACCGACCCGT TTAGTCTGCC GCTCCCAACA GCCCGAGTTT CTAGCCCCTT GAGTAAGGGG CAAGAAACCT AAGCCGCCTG GTGGATGGCT CGGCTCGGGG CGCTGACGAA GGACGCGTGG AC
J23SpTA_13              CGGGTTT GATGGGGCGG GGATGTAAGC GGGAAGGGAA ACCGACCCGT TTAGTCTGCC GCTCCCAACA GCCCGAGTTT CTAGCCCCTT GAGTAAGGGG CAAGAAACCT AAGCCGCCTG GTGGATG
J23SpTA_13              CGGGTTT GATGGGGCGG GGATGTAAGC GGGAAGGGAA ACCGACCCGT TTAGTCTGCC GCTCCCAACA GCCCGAGTTT CTAGCCCCTT GAGTAAGGGG CAAGAAACCT AAGCCGCCTG GTGGAT
J23SpTA_11                                               GGAAGGGAA ACCGACCCGT TTAGTCTGCC GCTCCCAACA GCCCGAGTTT CTAGCCCCTT GAGTAAGGGG CAAGAAACCT AAGCCGCCTG GTGGATGGCT CGGCTCGGGG CGCTGACGAA GGACGCGTGG ACG
J23SpTA_13                                              GGGAAGGGAA ACCGACCCGT TTAGTCTGCC GCTCCCAACA GCCCGAGTTT CTAGCCCCTT GAGTAAGGGG CAAGAAACCT AAGCCGCCTG GTGGATGGCT CGGCTCGGGG CGCTGACGAA GGACGCGTGG AC
J23SpTA_6_                                                   GGGAA ACCGACCCGT TTAGTCTGCC GCTCCCAACA GCCCGAGTTT CTAGCCCCTT GAGTAAGGGG CAAGAAACCT AAGCCGCCTG GTGGATGGCT CGGCTCGGGG CGCTGACGAA GGACGCGTGG ACGCGTG
J23SpTA_22      CAGA AGACGGGTTT GATGGGGCGG GGATGTAAGC GGGAAGGGAA ACCGACCCGT TTAGTCTGCC GCTCCCAACA GCCCGAGTTT CTAGCCCCTT GAGTAAGGGG CAAGAAACCT AAGCCGCCTG
J23SpTA_6_    CGCAGA AGACGGGTTT GATGGGGCGG GGATGTAAGC GGGAAGGGAA ACCGACCCGT TTAGTCTGCC GCTCCCAACA GCCCGAGTTT CTAGCCCCTT GAGTAAGGGG CAAGAAACCT AAGCCGCC
J23SpTA_33                             CGG GGATGTAAGC GGGAAGGGAA ACCGACCCGT TTAGTCTGCC GCTCCCAACA GCCCGAGTTT CTAGCCCCTT GAGTAAGGGG CAAGAAACCT AAGCCGCCTG GTGGATGGCT CGGCTCGGG
J23SpTA_5_                             CGG GGATGTAAGC GGGAAGGGAA ACCGACCCGT TTAGTCTGCC GCTCCCAACA GCCCGAGTTT CTAGCCCCTT GAGTAAGGGG CAAGAAACCT AAGCCGCCTG GTGGATGGCT CGGCTCGGGG
J23SpTA_37            GACGGGTTT GATGGGGCGG GGATGTAAGC GGGAAGGGAA ACCGACCCGT TTAGTCTGCC GCTCCCAACA GCCCGAGTTT CTAGCCCCTT GAGTAAGGGG CAAGAAACCT AAGCCGCCTG GTG
J23SpTA_10            GACGGGTTT GATGGGGCGG GGATGTAAGC GGGAAGGGAA ACCGACCCGT TTAGTCTGCC GCTCCCAACA GCCCGAGTTT CTAGCCCCTT GAGTAAGGGG CAAGAAACCT AAGCCGCCTG GTGG
J23SpTA_11                                            C GGGAAGGGAA ACCGACCCGT TTAGTCTGCC GCTCCCAACA GCCCGAGTTT CTAGCCCCTT GAGTAAGGGG CAAGAAACCT AAGCCGCCTG GTGGATGGCT CGGCTCGGGG CGCTGACGAA GGACGCGTG
J23SpTA_19                                              GGGAAGGGAA ACCGACCCGT TTAGTCTGCC GCTCCCAACA GCCCGAGTTT CTAGCCCCTT GAGTAAGGGG CAAGAAACCT AAGCCGCCTG GTGGATGGCT CGGCTCGGGG CGCTGACGAA GGACGCGTG
J23SpTA_7_                                                    GGAA ACCGACCCGT TTAGTCTGCC GCTCCCAACA GCCCGAGTTT CTAGCCCCTT GAGTAAGGGG CAAGAAACCT AAGCCGCCTG GTGGATGGCT CGGCTCGGGG CGCTGACGAA GGACGCGTGG ACGCGTGG
J23SpTA_12                                               GGAAGGGAA ACCGACCCGT TTAGTCTGCC GCTCCCAACA GCCCGAGTTT CTAGCCCCTT GAGTAAGGGG CAAGAAACCT AAGCCGCCTG GTGGATGGCT CGGCTCGGGG CGCTGACGAA GGACGCGTG
J23SpTA_10                               G GGATGTAAGC GGGAAGGGAA ACCGACCCGT TTAGTCTGCC GCTCCCAACA GCCCGAGTTT CTAGCCCCTT GAGTAAGGGG CAAGAAACCT AAGCCGCCTG GTGGATGGCT CGGCTCGGG
J23SpTA_5_                              GG GGATGTAAGC GGGAAGGGAA ACCGACCCGT TTAGTCTGCC GCTCCCAACA GCCCGAGTTT CTAGCCCCTT GAGTAAGGGG CAAGAAACCT AAGCCGCCTG GTGGATGGCT CGGCTCGGG
J23SpTA_40              CGGGTTT GATGGGGCGG GGATGTAAGC GGGAAGGGAA ACCGACCCGT TTAGTCTGCC GCTCCCAACA GCCCGAGTTT CTAGCCCCTT GAGTAAGGGG CAAGAAACCT AAGCCGCCTG GTG
J23SpTA_12             ACGGGTTT GATGGGGCGG GGATGTAAGC GGGAAGGGAA ACCGACCCGT TTAGTCTGCC GCTCCCAACA GCCCGAGTTT CTAGCCCCTT GAGTAAGGGG CAAGAAACCT AAGCCGCCTG GTG
J23SpTA_7_                                                   GGGAA ACCGACCCGT TTAGTCTGCC GCTCCCAACA GCCCGAGTTT CTAGCCCCTT GAGTAAGGGG CAAGAAACCT AAGCCGCCTG GTGGATGGCT CGGCTCGGGG CGCTGACGAA GGACGCGTGG ACGCG
J23SpTA_5_                 GTTT GATGGGGCGG GGATGTAAGC GGGAAGGGAA ACCGACCCGT TTAGTCTGCC GCTCCCAACA GCCCGAGTTT CTAGCCCCTT GAGTAAGGGG CAAGAAACCT AAGCCGCCTG GTGGAT
J23SpTA_31                                                   GGGAA ACCGACCCGT TTAGTCTGCC GCTCCCAACA GCCCGAGTTT CTAGCCCCTT GAGTAAGGGG CAAGAAACCT AAGCCGCCTG GTGGATGGCT CGGCTCGGGG CGCTGACGAA GGACGCGTGG ACGC
J23SpTA_7_                                                    GGAA ACCGACCCGT TTAGTCTGCC GCTCCCAACA GCCCGAGTTT CTAGCCCCTT GAGTAAGGGG CAAGAAACCT AAGCCGCCTG GTGGATGGCT CGGCTCGGGG CGCTGACGAA GGACGCGTGG ACGCG
J23SpTA_5_               GGGTTT GATGGGGCGG GGATGTAAGC GGGAAGGGAA ACCGACCCGT TTAGTCTGCC GCTCCCAACA GCCCGAGTTT CTAGCCCCTT GAGTAAGGGG CAAGAAACCT AAGCCGCCTG GTG
J23SpTA_26                                                   GGGAA ACCGACCCGT TTAGTCTGCC GCTCCCAACA GCCCGAGTTT CTAGCCCCTT GAGTAAGGGG CAAGAAACCT AAGCCGCCTG GTGGATGGCT CGGCTCGGGG CGCTGACGAA GGACGCGTGG ACG
J23SpTA_9_            GACGGGTTT GATGGGGCGG GGATGTAAGC GGGAAGGGAA ACCGACCCGT TTAGTCTGCC GCTCCCAACA GCCCGAGTTT CTAGCCCCTT GAGTAAGGGG CAAGAAACCT AAGCCGCCTG
J23SpTA_6_                                                    GGAA ACCGACCCGT TTAGTCTGCC GCTCCCAACA GCCCGAGTTT CTAGCCCCTT GAGTAAGGGG CAAGAAACCT AAGCCGCCTG GTGGATGGCT CGGCTCGGGG CGCTGACGAA GGACGCGTGG ACGC
J23SpTA_6_                                                     GAA ACCGACCCGT TTAGTCTGCC GCTCCCAACA GCCCGAGTTT CTAGCCCCTT GAGTAAGGGG CAAGAAACCT AAGCCGCCTG GTGGATGGCT CGGCTCGGGG CGCTGACGAA GGACGCGTGG ACGC
J23SpTA_17                                                   GGGAA ACCGACCCGT TTAGTCTGCC GCTCCCAACA GCCCGAGTTT CTAGCCCCTT GAGTAAGGGG CAAGAAACCT AAGCCGCCTG GTGGATGGCT CGGCTCGGGG CGCTGACGAA GGACGCGTGG AC
J23SpTA_6_                                                  AGGGAA ACCGACCCGT TTAGTCTGCC GCTCCCAACA GCCCGAGTTT CTAGCCCCTT GAGTAAGGGG CAAGAAACCT AAGCCGCCTG GTGGATGGCT CGGCTCGGGG CGCTGACGAA GGACGCGTG
J23SpTA_13                                                    GGAA ACCGACCCGT TTAGTCTGCC GCTCCCAACA GCCCGAGTTT CTAGCCCCTT GAGTAAGGGG CAAGAAACCT AAGCCGCCTG GTGGATGGCT CGGCTCGGGG CGCTGACGAA GGACGCGTGG ACG
J23SpTA_17                 GTTT GATGGGGCGG GGATGTAAGC GGGAAGGGAA ACCGACCCGT TTAGTCTGCC GCTCCCAACA GCCCGAGTTT CTAGCCCCTT GAGTAAGGGG CAAGAAACCT AAGCCGCCTG GTG
J23SpTA_5_                 GTTT GATGGGGCGG GGATGTAAGC GGGAAGGGAA ACCGACCCGT TTAGTCTGCC GCTCCCAACA GCCCGAGTTT CTAGCCCCTT GAGTAAGGGG CAAGAAACCT AAGCCGCCTG GTGG
J23SpTA_16                                                   GGGAA ACCGACCCGT TTAGTCTGCC GCTCCCAACA GCCCGAGTTT CTAGCCCCTT GAGTAAGGGG CAAGAAACCT AAGCCGCCTG GTGGATGGCT CGGCTCGGGG CGCTGACGAA GGACGCGTG
J23SpTA_5_                                                      CCGACCCGT TTAGTCTGCC GCTCCCAACA GCCCGAGTTT CTAGCCCCTT GAGTAAGGGG CAAGAAACCT AAGCCGCCTG GTGGATGGCT CGGCTCGGGG CGCTGACGAA GGACGCGTGG ACGCGTGG
J23SpTA_6_                                                    GGAA ACCGACCCGT TTAGTCTGCC GCTCCCAACA GCCCGAGTTT CTAGCCCCTT GAGTAAGGGG CAAGAAACCT AAGCCGCCTG GTGGATGGCT CGGCTCGGGG CGCTGACGAA GGACGCGTGG AC
J23SpTA_8_                                                     GAA ACCGACCCGT TTAGTCTGCC GCTCCCAACA GCCCGAGTTT CTAGCCCCTT GAGTAAGGGG CAAGAAACCT AAGCCGCCTG GTGGATGGCT CGGCTCGGGG CGCTGACGAA GGACGCGTGG ACG
J23SpTA_14              CGGGTTT GATGGGGCGG GGATGTAAGC GGGAAGGGAA ACCGACCCGT TTAGTCTGCC GCTCCCAACA GCCCGAGTTT CTAGCCCCTT GAGTAAGGGG CAAGAAACCT AAGCCGCCTG
J23SpTA_5_                                                               GT TTAGTCTGCC GCTCCCAACA GCCCGAGTTT CTAGCCCCTT GAGTAAGGGG CAAGAAACCT AAGCCGCCTG GTGGATGGCT CGGCTCGGGG CGCTGACGAA GGACGCGTGG ACGCGTGGCA AGCTGCG
J23SpTA_16                                                    GGAA ACCGACCCGT TTAGTCTGCC GCTCCCAACA GCCCGAGTTT CTAGCCCCTT GAGTAAGGGG CAAGAAACCT AAGCCGCCTG GTGGATGGCT CGGCTCGGGG CGCTGACGAA GGACGCGTG
J23SpTA_5_                                                      CCGACCCGT TTAGTCTGCC GCTCCCAACA GCCCGAGTTT CTAGCCCCTT GAGTAAGGGG CAAGAAACCT AAGCCGCCTG GTGGATGGCT CGGCTCGGGG CGCTGACGAA GGACGCGTGG ACGC
J23SpTA_7_                                                       CGACCCGT TTAGTCTGCC GCTCCCAACA GCCCGAGTTT CTAGCCCCTT GAGTAAGGGG CAAGAAACCT AAGCCGCCTG GTGGATGGCT CGGCTCGGGG CGCTGACGAA GGACGCGTGG ACGCG
J23SpTA_6_                                                    A ACCGACCCGT TTAGTCTGCC GCTCCCAACA GCCCGAGTTT CTAGCCCCTT GAGTAAGGGG CAAGAAACCT AAGCCGCCTG GTGGATGGCT CGGCTCGGGG CGCTGACGAA GGACGCGTGG ACG
J23SpTA_6_                                                     GAA ACCGACCCGT TTAGTCTGCC GCTCCCAACA GCCCGAGTTT CTAGCCCCTT GAGTAAGGGG CAAGAAACCT AAGCCGCCTG GTGGATGGCT CGGCTCGGGG CGCTGACGAA GGACGCGTG
J23SpTA_7_                                                      CCGACCCGT TTAGTCTGCC GCTCCCAACA GCCCGAGTTT CTAGCCCCTT GAGTAAGGGG CAAGAAACCT AAGCCGCCTG GTGGATGGCT CGGCTCGGGG CGCTGACGAA GGACGCGTGG ACG
J23SpTA_12                                                       CGACCCGT TTAGTCTGCC GCTCCCAACA GCCCGAGTTT CTAGCCCCTT GAGTAAGGGG CAAGAAACCT AAGCCGCCTG GTGGATGGCT CGGCTCGGGG CGCTGACGAA GGACGCGTGG ACGC
J23SpTA_15                                                       CGACCCGT TTAGTCTGCC GCTCCCAACA GCCCGAGTTT CTAGCCCCTT GAGTAAGGGG CAAGAAACCT AAGCCGCCTG GTGGATGGCT CGGCTCGGGG CGCTGACGAA GGACGCGTGG ACG
J23SpTA_6_                                                          CCCGT TTAGTCTGCC GCTCCCAACA GCCCGAGTTT CTAGCCCCTT GAGTAAGGGG CAAGAAACCT AAGCCGCCTG GTGGATGGCT CGGCTCGGGG CGCTGACGAA GGACGCGTGG ACGCG
J23SpTA_5_                                                      CCGACCCGT TTAGTCTGCC GCTCCCAACA GCCCGAGTTT CTAGCCCCTT GAGTAAGGGG CAAGAAACCT AAGCCGCCTG GTGGATGGCT CGGCTCGGGG CGCTGACGAA GGACGCGTG
J23SpTA_6_                                                        GACCCGT TTAGTCTGCC GCTCCCAACA GCCCGAGTTT CTAGCCCCTT GAGTAAGGGG CAAGAAACCT AAGCCGCCTG GTGGATGGCT CGGCTCGGGG CGCTGACGAA GGACGCGTGG ACG
J23SpTA_5_                                                          CCCGT TTAGTCTGCC GCTCCCAACA GCCCGAGTTT CTAGCCCCTT GAGTAAGGGG CAAGAAACCT AAGCCGCCTG GTGGATGGCT CGGCTCGGGG CGCTGACGAA GGACGCGTGG ACGC
J23SpTA_12                                                       CGACCCGT TTAGTCTGCC GCTCCCAACA GCCCGAGTTT CTAGCCCCTT GAGTAAGGGG CAAGAAACCT AAGCCGCCTG GTGGATGGCT CGGCTCGGGG CGCTGACGAA GGACGCGTG
J23SpTA_8_                                            C GGGAAGGGAA ACCGACCCGT TTAGTCTGCC GCTCCCAACA GCCCGAGTTT CTAGCCCCTT GAGTAAGGGG CAAGAAACCT AAGCCGCCTG GTGGATGGCT CGGCTCGGG
J23SpTA_6_                             CGG GGATGTAAGC GGGAAGGGAA ACCGACCCGT TTAGTCTGCC GCTCCCAACA GCCCGAGTTT CTAGCCCCTT GAGTAAGGGG CAAGAAACCT AAGCCGCCTG GTGGAT
J23SpTA_5_                             CGG GGATGTAAGC GGGAAGGGAA ACCGACCCGT TTAGTCTGCC GCTCCCAACA GCCCGAGTTT CTAGCCCCTT GAGTAAGGGG CAAGAAACCT AAGCCGCCTG GTGGATG
J23SpTA_26                                                          CCCGT TTAGTCTGCC GCTCCCAACA GCCCGAGTTT CTAGCCCCTT GAGTAAGGGG CAAGAAACCT AAGCCGCCTG GTGGATGGCT CGGCTCGGGG CGCTGACGAA GGACGCGTGG ACG
J23SpTA_7_                                                               GT TTAGTCTGCC GCTCCCAACA GCCCGAGTTT CTAGCCCCTT GAGTAAGGGG CAAGAAACCT AAGCCGCCTG GTGGATGGCT CGGCTCGGGG CGCTGACGAA GGACGCGTGG ACGCG
J23SpTA_12                                                          CCCGT TTAGTCTGCC GCTCCCAACA GCCCGAGTTT CTAGCCCCTT GAGTAAGGGG CAAGAAACCT AAGCCGCCTG GTGGATGGCT CGGCTCGGGG CGCTGACGAA GGACGCGTGG AC
J23SpTA_7_                                                          CCCGT TTAGTCTGCC GCTCCCAACA GCCCGAGTTT CTAGCCCCTT GAGTAAGGGG CAAGAAACCT AAGCCGCCTG GTGGATGGCT CGGCTCGGGG CGCTGACGAA GGACGCGTG
```

```
J23SpTA_8_                                  CGT TTAGTCTGCC GCTCCCAACA GCCCGAGTTT CTAGCCCCTT GAGTAAGGGG CAAGAAACCT AAGCCGCCTG GTGGATGGCT CGGCTCGGGG CGCTGACGAA GGACGCGTGG ACG
J23SpTA_8_                                   GT TTAGTCTGCC GCTCCCAACA GCCCGAGTTT CTAGCCCCTT GAGTAAGGGG CAAGAAACCT AAGCCGCCTG GTGGATGGCT CGGCTCGGGG CGCTGACGAA GGACGCGTGG ACG
J23SpTA_19  CGG GGATGTAAGC GGGAAGGGAA ACCGACCCGT TTAGTCTGCC GCTCCCAACA GCCCGAGTTT CTAGCCCCTT GAGTAAGGGG CAAGAAACCT AAGCCGCCTG GTG
J23SpTA_6_                                   GT TTAGTCTGCC GCTCCCAACA GCCCGAGTTT CTAGCCCCTT GAGTAAGGGG CAAGAAACCT AAGCCGCCTG GTGGATGGCT CGGCTCGGGG CGCTGACGAA GGACGCGTG
J23SpTA_7_  CGG GGATGTAAGC GGGAAGGGAA ACCGACCCGT TTAGTCTGCC GCTCCCAACA GCCCGAGTTT CTAGCCCCTT GAGTAAGGGG CAAGAAACCT AAGCCGCCTG
J23SpTA_11                                           CTGCC GCTCCCAACA GCCCGAGTTT CTAGCCCCTT GAGTAAGGGG CAAGAAACCT AAGCCGCCTG GTGGATGGCT CGGCTCGGGG CGCTGACGAA GGG
J23SpTA_7_                                                  CTCCCAACA GCCCGAGTTT CTAGCCCCTT GAGTAAGGGG CAAGAAACCT AAGCCGCCTG GTGGATGGCT CGGCTCGGGG CGCTGACGAA GGGC
J23SpTA_7_                                                  CTCCCAACA GCCCGAGTTT CTAGCCCCTT GAGTAAGGGG CAAGAAACCT AAGCCGCCTG GTGGATGGCT CGGCTCGGGG CGCTGACGAA GGG
J23SpTA_5_                                                    CCCAACA GCCCGAGTTT CTAGCCCCTT GAGTAAGGGG CAAGAAACCT AAGCCGCCTG GTGGATGGCT CGGCTCGGGG CGCTGACGAA GGGC
```
