## Supplementary Figure S2e for "Permuted 23S rRNA is integrated in 50S ribosome particles in *Thermococcus barophilus*"

```
               ____________23S : H98________________________|__________________________________23S : H99-101___________________________________________|_23S : TH3p__|_Junction|_23S : TH5p_|____23S____
                                                                                                                                                  <<<<<.. <<<<< ........>>>>>.. >>>>>
J23SpCA_1658                                                       CGCAGAAGA CGGGTTTGAT GGGGCGGGGA TGTAAGCGGG AAGGGAAACC GACCCGTTTA GTCTGCCGCT CCCAACAGCC CGAGTTTCCA GCCCCTTGAG TAAGGGGCAA GAAACTAAG CCGCCTGGTG GATGGCTCGG CTCGGGGCGC TGACGAAGGG
J23SpCA_847                                                        CGCAGAAGA CGGGTTTGAT GGGGCGGGGA TGTAAGCGGG AAGGGAAACC GACCCGTTTA GTCTGCCGCT CCCAACAGCC CGAGTTTCCA GCCCCTTGAG TAAGGGGCAA GAAACTAAG CCGCCTGGTG GATGGCTCGG CTCGGGGCGC TGACGAAGGG C
J23SpCA_652                                                     C CCGCAGAAGA CGGGTTTGAT GGGGCGGGGA TGTAAGCGGG AAGGGAAACC GACCCGTTTA GTCTGCCGCT CCCAACAGCC CGAGTTTCCA GCCCCTTGAG TAAGGGGCAA GAAACTAAG CCGCCTGGTG GATGGCTCGG CTCGGGGCGC TGACGAAGGG
J23SpCA_375                                                     C CCGCAGAAGA CGGGTTTGAT GGGGCGGGGA TGTAAGCGGG AAGGGAAACC GACCCGTTTA GTCTGCCGCT CCCAACAGCC CGAGTTTCCA GCCCCTTGAG TAAGGGGCAA GAAACTAAG CCGCCTGGTG GATGGCTCGG CTCGGGGCGC TGACGAAGGG C
J23SpCA_7                                                       C CCGCAGAAGA CGGGTTTGAT GGGGCGGGGA TGTAAGCGGG AAGGGAAACC GACCCGTTTA GTCTGCCGCT CCCAACAGCC CGAGTTTCCA GCCCCTTGAG TAAGGGGCAA GAAACTAAG CCGCCTGGTG GATGGCTCGG CTCGGGGCGC TGACGAAGGG CC
J23SpCA_9                                                         CCGCAGAAGA CGGGTTTGAT GGGGCGGGGA TGTAAGCGGG AAGGGAAACC GACCCGTTTA GTCTGCCGCT CCCAACAGCC CGAGTTTCCA GCCCCTTGAG TAAGGGGCAA GAAACTAAG CCGCCTGGTG GATGGCTCGG CTCGGGGCGC TGACGAAGGG CC
J23SpCA_3                                                       C CCGCAGAAGA CGGGTTTGAT GGGGCGGGGA TGTAAGCGGG AAGGGAAACC GACCCGTTTA GTCTGCCGCT CCCAACAGCC CGAGTTTCCA GCCCCTTGAG TAAGGGGCAA GAAACTAAG CCGCCTGGTG GATGGCTCGG CTCGGGGCGC TGACGAAGGG CA
J23SpCA_3                                                       C CCGCAGAAGA CGGGTTTGAT GGGGCGGGGA TGTAAGCGGG AAGGGAAACC GACCCGTTTA GTCTGCCGCT CCCAACAGCC CGAGTTTCCA GCCCCTTGAG TAAGGGGCAA GAAACTAAG CCGCCTGGTG GATGGCTCGG CTCGGGGCGC TGACGAAGGG CT
J23SpCA_405                                                       CCGCAGAAGA CGGGTTTGAT GGGGCGGGGA TGTAAGCGGG AAGGGAAACC GACCCGTTTA GTCTGCCGCT CCCAACAGCC CGAGTTTCCA GCCCCTTGAG TAAGGGGCAA GAAACTAAG CCGCCTGGTG GATGGCTCGG CTCGGGGCGC TGACGAAGGG C
J23SpCA_4                                                       C CCGCAGAAGA CGGGTTTGAT GGGGCGGGGA TGTAAGCGGG AAGGGAAACC GACCCGTTTA GTCTGCCGCT CCCAACAGCC CGAGTTTCCA GCCCCTTGAG TAAGGGGCAA GAAACTAAG CCGCCTGGTG GATGGCTCGG CTCGGGGCGC TGACGAAGGA C
J23SpCA_24                                                      C CCGCAGAAGA CGGGTTTGAT GGGGCGGGGA TGTAAGCGGG AAGGGAAACC GACCCGTTTA GTCTGCCGCT CCCAACAGCC CGAGTTTCCA GCCCCTTGAG TAAGGGGCAA GAAACTAAG CCGCCTGGTG GATGGCTCGG CTCGGGGCGC TGACGAAGGG G
J23SpCA_12                                                         CGCAGAAGA CGGGTTTGAT GGGGCGGGGA TGTAAGCGGG AAGGGAAACC GACCCGTTTA GTCTGCCGCT CCCAACAGCC CGAGTTTCCA GCCCCTTGAG TAAGGGGCAA GAAACTAAG CCGCCTGGTG GATGGCTCGG CTCGGGGCGC TGACGAAGGG CC
J23SpCA_3                                                       C CCGCAGAAGA CGGGTTTGAT GGGGCGGGGA TGTAAGCGGG AAGGGAAACC GACCCGTTTA GTCTGCCGCT CCCAACAGCC CGAGTTTCCA GCCCCTTGAG TAAGGGGCAA GAAACTAAG CCGCCTGGTG GATGGCTCGG CTCGGGGCGC TGACGAAGGG N
J23SpCA_9                                                         CCGCAGAAGA CGGGTTTGAT GGGGCGGGGA TGTAAGCGGG AAGGGAAACC GACCCGTTTA GTCTGCCGCT CCCAACAGCC CGAGTTTCCA GCCCCTTGAG TAAGGGGCAA GAAACTAAG CCGCCTGGTG GATGGCTCGG CTCGGGGCGC TGACGAAGGG CA
J23SpCA_8                                                     CGC CCGCAGAAGA CGGGTTTGAT GGGGCGGGGA TGTAAGCGGG AAGGGAAACC GACCCGTTTA GTCTGCCGCT CCCAACAGCC CGAGTTTCCA GCCCCTTGAG TAAGGGGCAA GAAACTAAG CCGCCTGGTG GATGGCTCGG CTCGGGGCGC TGACGAAGG
J23SpCA_4                                                     CGC CCGCAGAAGA CGGGTTTGAT GGGGCGGGGA TGTAAGCGGG AAGGGAAACC GACCCGTTTA GTCTGCCGCT CCCAACAGCC CGAGTTTCCA GCCCCTTGAG TAAGGGGCAA GAAACTAAG CCGCCTGGTG GATGGCTCGG CTCGGGGCGC TGACGAAGGG
J23SpCA_7                                                      GC CCGCAGAAGA CGGGTTTGAT GGGGCGGGGA TGTAAGCGGG AAGGGAAACC GACCCGTTTA GTCTGCCGCT CCCAACAGCC CGAGTTTCCA GCCCCTTGAG TAAGGGGCAA GAAACTAAG CCGCCTGGTG GATGGCTCGG CTCGGGGCGC TGACGAAGGG
J23SpCA_4               CG CAGGGCCTGG GTGACCGGCC CATGCCAGGG ACGAGGGCGC CCGCAGAAGA CGGGTTTGAT GGGGCGGGGA TGTAAGCGGG AAGGGAAACC GACCCGTTTA GTCTGCCGCT CCCAACAGCC CGAGTTTCCA GCCCCTTGAG TAAGGGGCAA GAAACTAAG CCGCCTGGTG GATGGCTCGG CTCGGGGCGC TGACGAAG
J23SpCA_3       CTCCCTGGCG CAGGGCCTGG GTGACCGGCC CATGCCAGGG ACGAGGGCGC CCGCAGAAGA CGGGTTTGAT GGGGCGGGGA TGTAAGCGGG AAGGGAAACC GACCCGTTTA GTCTGCCGCT CCCAACAGCC CGAGTTTCCA GCCCCTTGAG TAAGGGGCAA GAAACTAAG CCGCCTGGTG GATGGCTCGG CTCGGGGCGC TGACGAAGG
J23SpCA_3                                                         CCGCAGAAGA CGGGTTTGAT GGGGCGGGGA TGTAAGCGGG AAGGGAAACC GACCCGTTTA GTCTGCCGCT CCCAACAGCC CGAGTTTCCA GCCCCTTGAG TAAGGGGCAA GAAACTAAG CCGCCTGGTG GATGGCTCGG CTCGGGGCGC TGACGAAGGG CT
J23SpCA_5                                                       C CCGCAGAAGA CGGGTTTGAT GGGGCGGGGA TGTAAGCGGG AAGGGAAACC GACCCGTTTA GTCTGCCGCT CCCAACAGCC CGAGTTTCCA GCCCCTTGAG TAAGGGGCAA GAAACTAAG CCGCCTGGTG GATGGCTCGG CTCGGGGCGC TGACGAAGGG T
J23SpCA_685                                                       CCGCAGAAGA CGGGTTTGAT GGGGCGGGGA TGTAAGCGGG AAGGGAAACC GACCCGTTTA GTCTGCCGCT CCCAACAGCC CGAGTTTCCA GCCCCTTGAG TAAGGGGCAA GAAACTAAG CCGCCTGGTG GATGGCTCGG CTCGGGGCGC TGACGAAGGG
J23SpCA_526                                                        CGCAGAAGA CGGGTTTGAT GGGGCGGGGA TGTAAGCGGG AAGGGAAACC GACCCGTTTA GTCTGCCGCT CCCAACAGCC CGAGTTTCCA GCCCCTTGAG TAAGGGGCAA GAAACTAAG CCGCCTGGTG GATGGCTCGG CTCGGGGCGC TGACGAAGGG CG
J23SpCA_4                                                          CGCAGAAGA CGGGTTTGAT GGGGCGGGGA TGTAAGCGGG AAGGGAAACC GACCCGTTTA GTCTGCCGCT CCCAACAGCC CGAGTTTCCA GCCCCTTGAG TAAGGGGCAA GAAACTAAG CCGCCTGGTG GATGGCTCGG CTCGGGGCGC TGACGAAGGG CGGG
J23SpCA_8                                                          CGCAGAAGA CGGGTTTGAT GGGGCGGGGA TGTAAGCGGG AAGGGAAACC GACCCGTTTA GTCTGCCGCT CCCAACAGCC CGAGTTTCCA GCCCCTTGAG TAAGGGGCAA GAAACTAAG CCGCCTGGTG GATGGCTCGG CTCGGGGCGC TGACGAAGGG CA
J23SpCA_4                                                          CGCAGAAGA CGGGTTTGAT GGGGCGGGGA TGTAAGCGGG AAGGGAAACC GACCCGTTTA GTCTGCCGCT CCCAACAGCC CGAGTTTCCA GCCCCTTGAG TAAGGGGCAA GAAACTAAG CCGCCTGGTG GATGGCTCGG CTCGGGGCGC TGACGAAGGG CATG
J23SpCA_6                                                          CGCAGAAGA CGGGTTTGAT GGGGCGGGGA TGTAAGCGGG AAGGGAAACC GACCCGTTTA GTCTGCCGCT CCCAACAGCC CGAGTTTCCA GCCCCTTGAG TAAGGGGCAA GAAACTAAG CCGCCTGGTG GATGGCTCGG CTCGGGGCGC TGACGAAGGG CGT
J23SpCA_450                                                     C CCGCAGAAGA CGGGTTTGAT GGGGCGGGGA TGTAAGCGGG AAGGGAAACC GACCCGTTTA GTCTGCCGCT CCCAACAGCC CGAGTTTCCA GCCCCTTGAG TAAGGGGCAA GAAACTAAG CCGCCTGGTG GATGGCTCGG CTCGGGGCGC TGACGAAGG
J23SpCA_4                                                         CCGCAGAAGA CGGGTTTGAT GGGGCGGGGA TGTAAGCGGG AAGGGAAACC GACCCGTTTA GTCTGCCGCT CCCAACAGCC CGAGTTTCCA GCCCCTTGAG TAAGGGGCAA GAAACTAAG CCGCCTGGTG GATGGCTCGG CTCGGGGCGC TGACGAAGGA CG
J23SpCA_3                                                         CCGCAGAAGA CGGGTTTGAT GGGGCGGGGA TGTAAGCGGG AAGGGAAACC GACCCGTTTA GTCTGCCGCT CCCAACAGCC CGAGTTTCCA GCCCCTTGAG TAAGGGGCAA GAAACTAAG CCGCCTGGTG GATGGCTCGG CTCGGGGCGC TGACGAAGGA C
J23SpCA_3                                                          CGCAGAAGA CGGGTTTGAT GGGGCGGGGA TGTAAGCGGG AAGGGAAACC GACCCGTTTA GTCTGCCGCT CCCAACAGCC CGAGTTTCCA GCCCCTTGAG TAAGGGGCAA GAAACTAAG CCGCCTGGTG GATGGCTCGG CTCGGGGCGC TGACGAAGGA CG
J23SpCA_3                                                         CCGCAGAAGA CGGGTTTGAT GGGGCGGGGA TGTAAGCGGG AAGGGAAACC GACCCGTTTA GTCTGCCGCT CCCAACAGCC CGAGTTTCCA GCCCCTTGAG TAAGGGGCAA GAAACTAAG CCGCCTGGTG GATGGCTCGG CTCGGGGCGC TGACGAAGGG T
J23SpCA_16                                                        CCGCAGAAGA CGGGTTTGAT GGGGCGGGGA TGTAAGCGGG AAGGGAAACC GACCCGTTTA GTCTGCCGCT CCCAACAGCC CGAGTTTCCA GCCCCTTGAG TAAGGGGCAA GAAACTAAG CCGCCTGGTG GATGGCTCGG CTCGGGGCGC TGACGAAGGG G
J23SpCA_13                                                      C CCGCAGAAGA CGGGTTTGAT GGGGCGGGGA TGTAAGCGGG AAGGGAAACC GACCCGTTTA GTCTGCCGCT CCCAACAGCC CGAGTTTCCA GCCCCTTGAG TAAGGGGCAA GAAACTAAG CCGCCTGGTG GATGGCTCGG CTCGGGGCGC TGACGAAGAG
J23SpCA_7                                                          CGCAGAAGA CGGGTTTGAT GGGGCGGGGA TGTAAGCGGG AAGGGAAACC GACCCGTTTA GTCTGCCGCT CCCAACAGCC CGAGTTTCCA GCCCCTTGAG TAAGGGGCAA GAAACTAAG CCGCCTGGTG GATGGCTCGG CTCGGGGCGC TGACGAAGGG AC
J23SpCA_7                                                       C CCGCAGAAGA CGGGTTTGAT GGGGCGGGGA TGTAAGCGGG AAGGGAAACC GACCCGTTTA GTCTGCCGCT CCCAACAGCC CGAGTTTCCA GCCCCTTGAG TAAGGGGCAA GAAACTAAG CCGCCTGGTG GATGGCTCGG CTCGGGGCGC TGACAAAGGG
J23SpCA_5                                                      GC CCGCAGAAGA CGGGTTTGAT GGGGCGGGGA TGTAAGCGGG AAGGGAAACC GACCCGTTTA GTCTGCCGCT CCCAACAGCC CGAGTTTCCA GCCCCTTGAG TAAGGGGCAA GAAACTAAG CCGCCTGGTG GATGGCTCGG CTCGGGGCGC TGACGAAGG
J23SpCA_3                                                       C CCGCAGAAGA CGGGTTTGAT GGGGCGGGGA TGTAAGCGGG AAGGGAAACC GACCCGTTTA GTCTGCCGCT CCCAACAGCC CGAGTTTCCA GCCCCTTGAG TAAGGGGCAA GAAACTAAG CCGCCTGGTG GATGGCTCGG CTCGGGGCGC TGACGAAGGA G
J23SpCA_2                                                      GC CCGCAGAAGA CGGGTTTGAT GGGGCGGGGA TGTAAGCGGG AAGGGAAACC GACCCGTTTA GTCTGCCGCT CCCAACAGCC CGAGTTTCCA GCCCCTTGAG TAAGGGGCAA GAAACTAAG CCGCCTGGTG GATGGCTCGG CTCGGGGCGC TGACGAAGGA G
J23SpCA_5                                                          CGCAGAAGA CGGGTTTGAT GGGGCGGGGA TGTAAGCGGG AAGGGAAACC GACCCGTTTA GTCTGCCGCT CCCAACAGCC CGAGTTTCCA GCCCCTTGAG TAAGGGGCAA GAAACTAAG CCGCCTGGTG GATGGCTCGG CTCGGGGCGC TGACGAAGGG CT
J23SpCA_2                                                       G CCGCAGAAGA CGGGTTTGAT GGGGCGGGGA TGTAAGCGGG AAGGGAAACC GACCCGTTTA GTCTGCCGCT CCCAACAGCC CGAGTTTCCA GCCCCTTGAG TAAGGGGCAA GAAACTAAG CCGCCTGGTG GATGGCTCGG CTCGGGGCGC TGACGAAGGG
J23SpCA_486                                                     C CCGCAGAAGA CGGGTTTGAT GGGGCGGGGA TGTAAGCGGG AAGGGAAACC GACCCGTTTA GTCTGCCGCT CCCAACAGCC CGAGTTTCCA GCCCCTTGAG TAAGGGGCAA GAAACTAAG CCGCCTGGTG GATGGCTCGG CTCGGGGCGC TGACGAAG
J23SpCA_466                                                       CCGCAGAAGA CGGGTTTGAT GGGGCGGGGA TGTAAGCGGG AAGGGAAACC GACCCGTTTA GTCTGCCGCT CCCAACAGCC CGAGTTTCCA GCCCCTTGAG TAAGGGGCAA GAAACTAAG CCGCCTGGTG GATGGCTCGG CTCGGGGCGC TGACGAAGG
J23SpCA_104                                                         GCAGAAGA CGGGTTTGAT GGGGCGGGGA TGTAAGCGGG AAGGGAAACC GACCCGTTTA GTCTGCCGCT CCCAACAGCC CGAGTTTCCA GCCCCTTGAG TAAGGGGCAA GAAACTAAG CCGCCTGGTG GATGGCTCGG CTCGGGGCGC TGACGAAGGG C
J23SpCA_57                                                         CGCAGAAGA CGGGTTTGAT GGGGCGGGGA TGTAAGCGGG AAGGGAAACC GACCCGTTTA GTCTGCCGCT CCCAACAGCC CGAGTTTCCA GCCCCTTGAG TAAGGGGCAA GAAACTAAG CCGCCTGGTG GATGGCTCGG CTCGGGGCGC TGACGAAGGG G
J23SpCA_47                                                          GCAGAAGA CGGGTTTGAT GGGGCGGGGA TGTAAGCGGG AAGGGAAACC GACCCGTTTA GTCTGCCGCT CCCAACAGCC CGAGTTTCCA GCCCCTTGAG TAAGGGGCAA GAAACTAAG CCGCCTGGTG GATGGCTCGG CTCGGGGCGC TGACGAAGGG CG
J23SpCA_3                                                          GGCAGAAGA CGGGTTTGAT GGGGCGGGGA TGTAAGCGGG AAGGGAAACC GACCCGTTTA GTCTGCCGCT CCCAACAGCC CGAGTTTCCA GCCCCTTGAG TAAGGGGCAA GAAACTAAG CCGCCTGGTG GATGGCTCGG CTCGGGGCGC TGACGAAGGG CG
J23SpCA_2                                                          GGCAGAAGA CGGGTTTGAT GGGGCGGGGA TGTAAGCGGG AAGGGAAACC GACCCGTTTA GTCTGCCGCT CCCAACAGCC CGAGTTTCCA GCCCCTTGAG TAAGGGGCAA GAAACTAAG CCGCCTGGTG GATGGCTCGG CTCGGGGCGC TGACGAAGGG C
J23SpCA_2                                                           GCAGAAGA CGGGTTTGAT GGGGCGGGGA TGTAAGCGGG AAGGGAAACC GACCCGTTTA GTCTGCCGCT CCCAACAGCC CGAGTTTCCA GCCCCTTGAG TAAGGGGCAA GAAACTAAG CCGCCTGGTG GATGGCTCGG CTCGGGGCGC TGACGAAGGG CGC
J23SpCA_3                                                          AGCAGAAGA CGGGTTTGAT GGGGCGGGGA TGTAAGCGGG AAGGGAAACC GACCCGTTTA GTCTGCCGCT CCCAACAGCC CGAGTTTCCA GCCCCTTGAG TAAGGGGCAA GAAACTAAG CCGCCTGGTG GATGGCTCGG CTCGGGGCGC TGACGAAGGG C
J23SpCA_2                                                         NCGCAGAAGA CGGGTTTGAT GGGGCGGGGA TGTAAGCGGG AAGGGAAACC GACCCGTTTA GTCTGCCGCT CCCAACAGCC CGAGTTTCCA GCCCCTTGAG TAAGGGGCAA GAAACTAAG CCGCCTGGTG GATGGCTCGG CTCGGGGCGC TGACGAAGGG N
J23SpCA_14                                                         CGCAGAAGA CGGGTTTGAT GGGGCGGGGA TGTAAGCGGG AAGGGAAACC GACCCGTTTA GTCTGCCGCT CCCAACAGCC CGAGTTTCCA GCCCCTTGAG TAAGGGGCAA GAAACTAAG CCGCCTGGTG GATGGCTCGG CTCGGGGCGC TGACGAAGGA C
J23SpCA_2                                                          NGCAGAAGA CGGGTTTGAT GGGGCGGGGA TGTAAGCGGG AAGGGAAACC GACCCGTTTA GTCTGCCGCT CCCAACAGCC CGAGTTTCCA GCCCCTTGAG TAAGGGGCAA GAAACTAAG CCGCCTGGTG GATGGCTCGG CTCGGGGCGC TGACGAAGGG CN
J23SpCA_3                                                         CCGCAGAAGA CGGGTTTGAT GGGGCGGGGA TGTAAGCGGG AAGGGAAACC GACCCGTTTA GTCTGCCGCT CCCAACAGCC CGAGTTTCCA GCCCCTTGAG TAAGGGGCAA GAAACTAAG CCGCCTGGTG GATGGCTCGG CTCGGGGCGC TGACGAAAGG
J23SpCA_6                                                          CGCAGAAGA CGGGTTTGAT GGGGCGGGGA TGTAAGCGGG AAGGGAAACC GACCCGTTTA GTCTGCCGCT CCCAACAGCC CGAGTTTCCA GCCCCTTGAG TAAGGGGCAA GAAACTAAG CCGCCTGGTG GATGGCTCGG CTCGGGGCGC TGACGAAGAG C
J23SpCA_6                                                         CCGCAGAAGA CGGGTTTGAT GGGGCGGGGA TGTAAGCGGG AAGGGAAACC GACCCGTTTA GTCTGCCGCT CCCAACAGCC CGAGTTTCCA GCCCCTTGAG TAAGGGGCAA GAAACTAAG CCGCCTGGTG GATGGCTCGG CTCGGGGCGC TGACGAAGAG
J23SpCA_6                                                         CCGCAGAAGA CGGGTTTGAT GGGGCGGGGA TGTAAGCGGG AAGGGAAACC GACCCGTTTA GTCTGCCGCT CCCAACAGCC CGAGTTTCCA GCCCCTTGAG TAAGGGGCAA GAAACTAAG CCGCCTGGTG GATGGCTCGG CTCGGGGCGC TGACAAAGGG
J23SpCA_4                                                       N CCGCAGAAGA CGGGTTTGAT GGGGCGGGGA TGTAAGCGGG AAGGGAAACC GACCCGTTTA GTCTGCCGCT CCCAACAGCC CGAGTTTCCA GCCCCTTGAG TAAGGGGCAA GAAACTAAG CCGCCTGGTG GATGGCTCGG CTCGGGGCGC TGACGAAGGN
J23SpCA_6                                                       C CCGCAGAAGA CGGGTTTGAT GGGGCGGGGA TGTAAGCGGG AAGGGAAACC GACCCGTTTA GTCTGCCGCT CCCAACAGCC CGAGTTTCCA GCCCCTTGAG TAAGGGGCAA GAAACTAAG CCGCCTGGTG GATGGCTCGG CTCGGGGCGC TGACAAAGG
J23SpCA_5                                                     CGC CCGCAGAAGA CGGGTTTGAT GGGGCGGGGA TGTAAGCGGG AAGGGAAACC GACCCGTTTA GTCTGCCGCT CCCAACAGCC CGAGTTTCCA GCCCCTTGAG TAAGGGGCAA GAAACTAAG CCGCCTGGTG GATGGCTCGG CTCGGGGCGC TGACGAAG
J23SpCA_3                                                       C CCGCAGAAGA CGGGTTTGAT GGGGCGGGGA TGTAAGCGGG AAGGGAAACC GACCCGTTTA GTCTGCCGCT CCCAACAGCC CGAGTTTCCA GCCCCTTGAG TAAGGGGCAA GAAACTAAG CCGCCTGGTG GATGGCTCGG CTCGGGGCGC TGACGAAAG
J23SpCA_3                                                           GCAGAAGA CGGGTTTGAT GGGGCGGGGA TGTAAGCGGG AAGGGAAACC GACCCGTTTA GTCTGCCGCT CCCAACAGCC CGAGTTTCCA GCCCCTTGAG TAAGGGGCAA GAAACTAAG CCGCCTGGTG GATGGCTCGG CTCGGGGCGC TGACGAAGGG CC
J23SpCA_1070                                                       CGCAGAAGA CGGGTTTGAT GGGGCGGGGA TGTAAGCGGG AAGGGAAACC GACCCGTTTA GTCTGCCGCT CCCAACAGCC CGAGTTTCCA GCCCCTTGAG TAAGGGGCAA GAAACTAAG CCGCCTGGTG GATGGCTCGG CTCGGGGCGC TGACGAAGG
J23SpCA_520                                                       CCGCAGAAGA CGGGTTTGAT GGGGCGGGGA TGTAAGCGGG AAGGGAAACC GACCCGTTTA GTCTGCCGCT CCCAACAGCC CGAGTTTCCA GCCCCTTGAG TAAGGGGCAA GAAACTAAG CCGCCTGGTG GATGGCTCGG CTCGGGGCGC TGACGAAG
J23SpCA_296                                                          CAGAAGA CGGGTTTGAT GGGGCGGGGA TGTAAGCGGG AAGGGAAACC GACCCGTTTA GTCTGCCGCT CCCAACAGCC CGAGTTTCCA GCCCCTTGAG TAAGGGGCAA GAAACTAAG CCGCCTGGTG GATGGCTCGG CTCGGGGCGC TGACGAAGGG C
J23SpCA_4                                                         CCCCAGAAGA CGGGTTTGAT GGGGCGGGGA TGTAAGCGGG AAGGGAAACC GACCCGTTTA GTCTGCCGCT CCCAACAGCC CGAGTTTCCA GCCCCTTGAG TAAGGGGCAA GAAACTAAG CCGCCTGGTG GATGGCTCGG CTCGGGGCGC TGACGAAGG
J23SpCA_172                                                          CAGAAGA CGGGTTTGAT GGGGCGGGGA TGTAAGCGGG AAGGGAAACC GACCCGTTTA GTCTGCCGCT CCCAACAGCC CGAGTTTCCA GCCCCTTGAG TAAGGGGCAA GAAACTAAG CCGCCTGGTG GATGGCTCGG CTCGGGGCGC TGACGAAGGG CG
J23SpCA_134                                                          CAGAAGA CGGGTTTGAT GGGGCGGGGA TGTAAGCGGG AAGGGAAACC GACCCGTTTA GTCTGCCGCT CCCAACAGCC CGAGTTTCCA GCCCCTTGAG TAAGGGGCAA GAAACTAAG CCGCCTGGTG GATGGCTCGG CTCGGGGCGC TGACGAAGGG CGTGG
J23SpCA_78                                                           CAGAAGA CGGGTTTGAT GGGGCGGGGA TGTAAGCGGG AAGGGAAACC GACCCGTTTA GTCTGCCGCT CCCAACAGCC CGAGTTTCCA GCCCCTTGAG TAAGGGGCAA GAAACTAAG CCGCCTGGTG GATGGCTCGG CTCGGGGCGC TGACGAAGGG CGTG
J23SpCA_49                                                            AGAAGA CGGGTTTGAT GGGGCGGGGA TGTAAGCGGG AAGGGAAACC GACCCGTTTA GTCTGCCGCT CCCAACAGCC CGAGTTTCCA GCCCCTTGAG TAAGGGGCAA GAAACTAAG CCGCCTGGTG GATGGCTCGG CTCGGGGCGC TGACGAAGGG CGTGG
J23SpCA_16                                                            AGAAGA CGGGTTTGAT GGGGCGGGGA TGTAAGCGGG AAGGGAAACC GACCCGTTTA GTCTGCCGCT CCCAACAGCC CGAGTTTCCA GCCCCTTGAG TAAGGGGCAA GAAACTAAG CCGCCTGGTG GATGGCTCGG CTCGGGGCGC TGACGAAGGG CGTGGC
J23SpCA_3                                                             AGAAGA CGGGTTTGAT GGGGCGGGGA TGTAAGCGGG AAGGGAAACC GACCCGTTTA GTCTGCCGCT CCCAACAGCC CGAGTTTCCA GCCCCTTGAG TAAGGGGCAA GAAACTAAG CCGCCTGGTG GATGGCTCGG CTCGGGGCGC TGACGAAGGG CGTGGCG
J23SpCA_13                                                          GCAGAAGA CGGGTTTGAT GGGGCGGGGA TGTAAGCGGG AAGGGAAACC GACCCGTTTA GTCTGCCGCT CCCAACAGCC CGAGTTTCCA GCCCCTTGAG TAAGGGGCAA GAAACTAAG CCGCCTGGTG GATGGCTCGG CTCGGGGCGC TGACGAAGGG CGTG
J23SpCA_6                                                            CAGAAGA CGGGTTTGAT GGGGCGGGGA TGTAAGCGGG AAGGGAAACC GACCCGTTTA GTCTGCCGCT CCCAACAGCC CGAGTTTCCA GCCCCTTGAG TAAGGGGCAA GAAACTAAG CCGCCTGGTG GATGGCTCGG CTCGGGGCGC TGACGAAGGG CGGGG
J23SpCA_5                                                            CAGAAGA CGGGTTTGAT GGGGCGGGGA TGTAAGCGGG AAGGGAAACC GACCCGTTTA GTCTGCCGCT CCCAACAGCC CGAGTTTCCA GCCCCTTGAG TAAGGGGCAA GAAACTAAG CCGCCTGGTG GATGGCTCGG CTCGGGGCGC TGACGAAGGG CGTAG
J23SpCA_2                                                           GCAGAAGA CGGGTTTGAT GGGGCGGGGA TGTAAGCGGG AAGGGAAACC GACCCGTTTA GTCTGCCGCT CCCAACAGCC CGAGTTTCCA GCCCCTTGAG TAAGGGGCAA GAAACTAAG CCGCCTGGTG GATGGCTCGG CTCGGGGCGC TGACGAAGGG CGTAG
J23SpCA_4                                                              GAAGA CGGGTTTGAT GGGGCGGGGA TGTAAGCGGG AAGGGAAACC GACCCGTTTA GTCTGCCGCT CCCAACAGCC CGAGTTTCCA GCCCCTTGAG TAAGGGGCAA GAAACTAAG CCGCCTGGTG GATGGCTCGG CTCGGGGCGC TGACGAAGGG CGTGG
J23SpCA_35                                                            AGAAGA CGGGTTTGAT GGGGCGGGGA TGTAAGCGGG AAGGGAAACC GACCCGTTTA GTCTGCCGCT CCCAACAGCC CGAGTTTCCA GCCCCTTGAG TAAGGGGCAA GAAACTAAG CCGCCTGGTG GATGGCTCGG CTCGGGGCGC TGACGAAGGG CGTG
J23SpCA_3                                                            CAGAAGA CGGGTTTGAT GGGGCGGGGA TGTAAGCGGG AAGGGAAACC GACCCGTTTA GTCTGCCGCT CCCAACAGCC CGAGTTTCCA GCCCCTTGAG TAAGGGGCAA GAAACTAAG CCGCCTGGTG GATGGCTCGG CTCGGGGCGC TGACGAAGGG CGT
J23SpCA_2                                                              GAAGA CGGGTTTGAT GGGGCGGGGA TGTAAGCGGG AAGGGAAACC GACCCGTTTA GTCTGCCGCT CCCAACAGCC CGAGTTTCCA GCCCCTTGAG TAAGGGGCAA GAAACTAAG CCGCCTGGTG GATGGCTCGG CTCGGGGCGC TGACGAAGGG CGTGGC
J23SpCA_3                                                             AGAAGA CGGGTTTGAT GGGGCGGGGA TGTAAGCGGG AAGGGAAACC GACCCGTTTA GTCTGCCGCT CCCAACAGCC CGAGTTTCCA GCCCCTTGAG TAAGGGGCAA GAAACTAAG CCGCCTGGTG GATGGCTCGG CTCGGGGCGC TGACGAAGGG CGCGG
J23SpCA_10                                                             GAAGA CGGGTTTGAT GGGGCGGGGA TGTAAGCGGG AAGGGAAACC GACCCGTTTA GTCTGCCGCT CCCAACAGCC CGAGTTTCCA GCCCCTTGAG TAAGGGGCAA GAAACTAAG CCGCCTGGTG GATGGCTCGG CTCGGGGCGC TGACGAAGGG CGTG
J23SpCA_2                                                              GAAGA CGGGTTTGAT GGGGCGGGGA TGTAAGCGGG AAGGGAAACC GACCCGTTTA GTCTGCCGCT CCCAACAGCC CGAGTTTCCA GCCCCTTGAG TAAGGGGCAA GAAACTAAG CCGCCTGGTG GATGGCTCGG CTCGGGGCGC TGACGAAGGG CGGGG
J23SpCA_3                                                            CAGAAGA CGGGTTTGAT GGGGCGGGGA TGTAAGCGGG AAGGGAAACC GACCCGTTTA GTCTGCCGCT CCCAACAGCC CGAGTTTCCA GCCCCTTGAG TAAGGGGCAA GAAACTAAG CCGCCTGGTG GATGGCTCGG CTCGGGGCGC TGACGAAGGG CAT
J23SpCA_6                                                             AGAAGA CGGGTTTGAT GGGGCGGGGA TGTAAGCGGG AAGGGAAACC GACCCGTTTA GTCTGCCGCT CCCAACAGCC CGAGTTTCCA GCCCCTTGAG TAAGGGGCAA GAAACTAAG CCGCCTGGTG GATGGCTCGG CTCGGGGCGC TGACGAAGGG CGT
J23SpCA_6                                                               AAGA CGGGTTTGAT GGGGCGGGGA TGTAAGCGGG AAGGGAAACC GACCCGTTTA GTCTGCCGCT CCCAACAGCC CGAGTTTCCA GCCCCTTGAG TAAGGGGCAA GAAACTAAG CCGCCTGGTG GATGGCTCGG CTCGGGGCGC TGACGAAGGG CGTGG
J23SpCA_5                                                              GAAGA CGGGTTTGAT GGGGCGGGGA TGTAAGCGGG AAGGGAAACC GACCCGTTTA GTCTGCCGCT CCCAACAGCC CGAGTTTCCA GCCCCTTGAG TAAGGGGCAA GAAACTAAG CCGCCTGGTG GATGGCTCGG CTCGGGGCGC TGACGAAGGG CATGG
J23SpCA_161                                                         GCAGAAGA CGGGTTTGAT GGGGCGGGGA TGTAAGCGGG AAGGGAAACC GACCCGTTTA GTCTGCCGCT CCCAACAGCC CGAGTTTCCA GCCCCTTGAG TAAGGGGCAA GAAACTAAG CCGCCTGGTG GATGGCTCGG CTCGGGGCGC TGACGAAGGG
J23SpCA_20                                                         CGCAGAAGA CGGGTTTGAT GGGGCGGGGA TGTAAGCGGG AAGGGAAACC GACCCGTTTA GTCTGCCGCT CCCAACAGCC CGAGTTTCCA GCCCCTTGAG TAAGGGGCAA GAAACTAAG CCGCCTGGTG GATGGCTCGG CTCGGGGCGC TGACGAAGAG
J23SpCA_3                                                            CAGAAGA CGGGTTTGAT GGGGCGGGGA TGTAAGCGGG AAGGGAAACC GACCCGTTTA GTCTGCCGCT CCCAACAGCC CGAGTTTCCA GCCCCTTGAG TAAGGGGCAA GAAACTAAG CCGCCTGGTG GATGGCTCGG CTCGGGGCGC TGACGAAGGG CC
J23SpCA_3                                                          CGCAGAAGA CGGGTTTGAT GGGGCGGGGA TGTAAGCGGG CAGGGAAACC GACCCGTTTA GTCTGCCGCT CCCAACAGCC CGAGTTTCCA GCCCCTTGAG TAAGGGGCAA GAAACTAAG CCGCCTGGTG GATGGCTCGG CTCGGGGCGC TGACGAAGGG
J23SpCA_3                                                          CGCAGAAGA CGGGTTTGAT GGGGCGGGGA TGTAAGCGGG AAGGGAAACC GACCCGTTTA GTCTGCCGCT CCCAACAGCC CGAGTTTCCA GCCCCTTGAG TAAGGGGCAA GAAACTAAG CCGCCTGGTG GATGGCTCGG CTCGGGGCGC TGACGAAAGG
J23SpCA_8                                                          CGCAGAAGA CGGGTTTGAT GGGGCGGGGA TGTAAGCGGG AAGGGAAACC GACCCGTTTA GTCTGCCGCT CCCAACAGCC CGAGTTTCCA GCCCCTTGAG TAAGGGGCAA GAAACTAAG CCGCCTGGTG GATGGCTCGG CTCGGGGCGC TGACGAAGGC
J23SpCA_8                                                          CGCAGAAGA CGGGTTTGAT GGGGCGGGGA TGTAAGCGGG AAGGGAAACC GACCCGTTTA GTCTGCCGCT CCCAACAGCC CGAGTTTCCA GCCCCTTGAG TAAGGGGCAA GAAACTAAG CCGCCTGGTG GATGGCTCGG CTCGGGGCGC TGACAAAGGG
J23SpCA_3                                                          GGCAGAAGA CGGGTTTGAT GGGGCGGGGA TGTAAGCGGG AAGGGAAACC GACCCGTTTA GTCTGCCGCT CCCAACAGCC CGAGTTTCCA GCCCCTTGAG TAAGGGGCAA GAAACTAAG CCGCCTGGTG GATGGCTCGG CTCGGGGCGC TGACGAAGGG
J23SpCA_7                                                          CGCAGAAGA CGGGTTTGAT GGGGCGGGGA TGTAAGCGGG AAGGGAAACC GACCCGTTTA GTCTGCCGCT CCCAACAGCC CGAGTTTCCA GCCCCTTGAG TAAGGGGCAA GAAACTAAG CCGCCTGGTG GATGGCTCGG CTCGGGGCGC TGACGAGGGG
J23SpCA_5                                                          NGCAGAAGA CGGGTTTGAT GGGGCGGGGA TGTAAGCGGG AAGGGAAACC GACCCGTTTA GTCTGCCGCT CCCAACAGCC CGAGTTTCCA GCCCCTTGAG TAAGGGGCAA GAAACTAAG CCGCCTGGTG GATGGCTCGG CTCGGGGCGC TGACGAAGGG N
J23SpCA_5                                                           GCAGAAGA CGGGTTTGAT GGGGCGGGGA TGTAAGCGGG AAGGGAAACC GACCCGTTTA GTCTGCCGCT CCCAACAGCC CGAGTTTCCA GCCCCTTGAG TAAGGGGCAA GAAACTAAG CCGCCTGGTG GATGGCTCGG CTCGGGGCGC TGACGAAGGG G
J23SpCA_5                                                          CGCAGAAGA CGGGTTTGAT GGGGCGGGGA TGTAAGCGGG AAGGGAAACC GACCCGTTTA GTCTGCCGCT CCCAACAGCC CGAGTTTCCA GCCCCTTGAG TAAGGGGCAA GAAACTAAG CCGCCTGGTG GATGGCTCGG CTCGGGGCGC TGACGGAGGG
J23SpCA_1295                                                       CGCAGAAGA CGGGTTTGAT GGGGCGGGGA TGTAAGCGGG AAGGGAAACC GACCCGTTTA GTCTGCCGCT CCCAACAGCC CGAGTTTCCA GCCCCTTGAG TAAGGGGCAA GAAACTAAG CCGCCTGGTG GATGGCTCGG CTCGGGGCGC TGACGAAG
J23SpCA_584                                                          CAGAAGA CGGGTTTGAT GGGGCGGGGA TGTAAGCGGG AAGGGAAACC GACCCGTTTA GTCTGCCGCT CCCAACAGCC CGAGTTTCCA GCCCCTTGAG TAAGGGGCAA GAAACTAAG CCGCCTGGTG GATGGCTCGG CTCGGGGCGC TGACGAAGGG
J23SpCA_111                                                           AGAAGA CGGGTTTGAT GGGGCGGGGA TGTAAGCGGG AAGGGAAACC GACCCGTTTA GTCTGCCGCT CCCAACAGCC CGAGTTTCCA GCCCCTTGAG TAAGGGGCAA GAAACTAAG CCGCCTGGTG GATGGCTCGG CTCGGGGCGC TGACGAAGGG C
J23SpCA_91                                                          GCAGAAGA CGGGTTTGAT GGGGCGGGGA TGTAAGCGGG AAGGGAAACC GACCCGTTTA GTCTGCCGCT CCCAACAGCC CGAGTTTCCA GCCCCTTGAG TAAGGGGCAA GAAACTAAG CCGCCTGGTG GATGGCTCGG CTCGGGGCGC TGACGAAGG
J23SpCA_2                                                       G CCGCAGAAGA CGGGTTTGAT GGGGCGGGGA TGTAAGCGGG AAGGGAAACC GACCCGTTTA GTCTGCCGCT CCCAACAGCC CGAGTTTCCA GCCCCTTGAG TAAGGGGCAA GAAACTAAG CCGCCTGGTG GATGGCTCGG CTCGGGGCGC TGACGAAG
J23SpCA_58                                                            AGAAGA CGGGTTTGAT GGGGCGGGGA TGTAAGCGGG AAGGGAAACC GACCCGTTTA GTCTGCCGCT CCCAACAGCC CGAGTTTCCA GCCCCTTGAG TAAGGGGCAA GAAACTAAG CCGCCTGGTG GATGGCTCGG CTCGGGGCGC TGACGAAGGG CG
J23SpCA_4                                                          CGCAGAAGA CGGGTTTGAT GGGGCGGGGA TGTAAGCGGG AAGGGAAACC GACCCGTTTA GTCTGCCGCT CCCAACAGCC CGAGTTTCCA GCCCCTTGAG TAAGGGGCAA GAAACTAAG CCGCCTGGTG GATGGCTCGG CTCGGGGCGC TGACGAAGN
J23SpCA_2                                                         NCGCAGAAGA CGGGTTTGAT GGGGCGGGGA TGTAAGCGGG AAGGGAAACC GACCCGTTTA GTCTGCCGCT CCCAACAGCC CGAGTTTCCA GCCCCTTGAG TAAGGGGCAA GAAACTAAG CCGCCTGGTG GATGGCTCGG CTCGGGGCGC TGACGAAGN
J23SpCA_18                                                           CAGAAGA CGGGTTTGAT GGGGCGGGGA TGTAAGCGGG AAGGGAAACC GACCCGTTTA GTCTGCCGCT CCCAACAGCC CGAGTTTCCA GCCCCTTGAG TAAGGGGCAA GAAACTAAG CCGCCTGGTG GATGGCTCGG CTCGGGGCGC TGACGAAGGG G
J23SpCA_16                                                         CGCAGAAGA CGGGTTTGAT GGGGCGGGGA TGTAAGCGGG AAGGGAAACC GACCCGTTTA GTCTGCCGCT CCCAACAGCC CGAGTTTCCA GCCCCTTGAG TAAGGGGCAA GAAACTAAG CCGCCTGGTG GATGGCTCGG CTCGGGGCGC TGACAAAGG
J23SpCA_7                                                         CCGCAGAAGA CGGGTTTGAT GGGGCGGGGA TGTAAGCGGG AAGGGAAACC GACCCGTTTA GTCTGCCGCT CCCAACAGCC CGAGTTTCCA GCCCCTTGAG TAAGGGGCAA GAAACTAAG CCGCCTGGTG GATGGCTCGG CTCGGGGCGC TGACAAAGG
J23SpCA_6                                                         CCGCAGAAGA CGGGTTTGAT GGGGCGGGGA TGTAAGCGGG AAGGGAAACC GACCCGTTTA GTCTGCCGCT CCCAACAGCC CGAGTTTCCA GCCCCTTGAG TAAGGGGCAA GAAACTAAG CCGCCTGGTG GATGGCTCGG CTCGGGGCGC TGACAAAG
J23SpCA_15                                                      C CCGCAGAAGA CGGGTTTGAT GGGGCGGGGA TGTAAGCGGG AAGGGAAACC GACCCGTTTA GTCTGCCGCT CCCAACAGCC CGAGTTTCCA GCCCCTTGAG TAAGGGGCAA GAAACTAAG CCGCCTGGTG GATGGCTCGG CTCGGGGCGC TGACGAA
J23SpCA_4                                                           CCAGAAGA CGGGTTTGAT GGGGCGGGGA TGTAAGCGGG AAGGGAAACC GACCCGTTTA GTCTGCCGCT CCCAACAGCC CGAGTTTCCA GCCCCTTGAG TAAGGGGCAA GAAACTAAG CCGCCTGGTG GATGGCTCGG CTCGGGGCGC TGACGAAGGG
J23SpCA_13                                                         CGCAGAAGA CGGGTTTGAT GGGGCGGGGA TGTAAGCGGG AAGGGAAACC GACCCGTTTA GTCTGCCGCT CCCAACAGCC CGAGTTTCCA GCCCCTTGAG TAAGGGGCAA GAAACTAAG CCGCCTGGTG GATGGCTCGG CTCGGGGCGC TGACGAAAG
J23SpCA_3                                                         CCGCAGAAGA CGGGTTTGAT GGGGCGGGGA TGTAAGCGGG AAGGGAAACC GACCCGTTTA GTCTGCCGCT CCCAACAGCC CGAGTTTCCA GCCCCTTGAG TAAGGGGCAA GAAACTAAG CCGCCTGGTG GATGGCTCGG CTCGGGGCGC TGACGAAAG
J23SpCA_10                                                         CGCAGAAGA CGGGTTTGAT GGGGCGGGGA TGTAAGCGGG AAGGGAAACC GACCCGTTTA GTCTGCCGCT CCCAACAGCC CGAGTTTCCA GCCCCTTGAG TAAGGGGCAA GAAACTAAG CCGCCTGGTG GATGGCTCGG CTCGGGGCGC TGACGAGGG
J23SpCA_2                                                           GCAGAAGA CGGGTTTGAT GGGGCGGGGA TGTAAGCGGG AAGGGAAACC GACCCGTTTA GTCTGCCGCT CCCAACAGCC CGAGTTTCCA GCCCCTTGAG TAAGGGGCAA GAAACTAAG CCGCCTGGTG GATGGCTCGG CTCGGGGCGC TGACAAAGGG
J23SpCA_8                                                          NGCAGAAGA CGGGTTTGAT GGGGCGGGGA TGTAAGCGGG AAGGGAAACC GACCCGTTTA GTCTGCCGCT CCCAACAGCC CGAGTTTCCA GCCCCTTGAG TAAGGGGCAA GAAACTAAG CCGCCTGGTG GATGGCTCGG CTCGGGGCGC TGACGAAGGN
J23SpCA_2                                                           GCAGAAGA CGGGTTTGAT GGGGCGGGGA TGTAAGCGGG AAGGGAAACC GACCCGTTTA GTCTGCCGCT CCCAACAGCC CGAGTTTCCA GCCCCTTGAG TAAGGGGCAA GAAACTAAG CCGCCTGGTG GATGGCTCGG CTCGGGGCGC TGACGAAGGN
J23SpCA_3                                                          CGCAGAAGA CGGGTTTGAT GGGGCGGGGA TGTAAGCGGG AAGGGAAACC GACCCGTTTA GTCTGCCGCT CCCAACAGCC CGAGTTTCCA GCCCCTTGAG TAAGGGGCAA GAAACTAAG CCGCCTGGTG GATGGCTCGG CTCGGGGCGC TGACGGAGG
J23SpCA_3                                                            CAGAAGA CGGGTTTGAT GGGGCGGGGA TGTAAGCGGG AAGGGAAACC GACCCGTTTA GTCTGCCGCT CCCAACAGCC CGAGTTTCCA GCCCCTTGAG TAAGGGGCAA GAAACTAAG CCGCCTGGTG GATGGCTCGG CTCGGGGCGC TGACGAAGGG A
J23SpCA_3                                                            CAGAAGA CGGGTTTGAT GGGGCGGGGA TGTAAGCGGG AAGGGAAACC GACCCGTTTA GTCTGCCGCT CCCAACAGCC CGAGTTTCCA GCCCCTTGAG TAAGGGGCAA GAAACTAAG CCGCCTGGTG GATGGCTCGG CTCGGGGCGC TGACGAAGGA C
J23SpCA_426                                                        CGCAGAAGA CGGGTTTGAT GGGGCGGGGA TGTAAGCGGG AAGGGAAACC GACCCGTTTA GTCTGCCGCT CCCAACAGCC CGAGTTTCCA GCCCCTTGAG TAAGGGGCAA GAAACTAAG CCGCCTGGTG GATGGCTCGG CTCGGGGCGC TGACGAAGGG CGTGG
J23SpCA_196                                                       CCGCAGAAGA CGGGTTTGAT GGGGCGGGGA TGTAAGCGGG AAGGGAAACC GACCCGTTTA GTCTGCCGCT CCCAACAGCC CGAGTTTCCA GCCCCTTGAG TAAGGGGCAA GAAACTAAG CCGCCTGGTG GATGGCTCGG CTCGGGGCGC TGACGAAGGG CGTGG
J23SpCA_165                                                     C CCGCAGAAGA CGGGTTTGAT GGGGCGGGGA TGTAAGCGGG AAGGGAAACC GACCCGTTTA GTCTGCCGCT CCCAACAGCC CGAGTTTCCA GCCCCTTGAG TAAGGGGCAA GAAACTAAG CCGCCTGGTG GATGGCTCGG CTCGGGGCGC TGACGAAGGG CGTGG
J23SpCA_41                                                      C CCGCAGAAGA CGGGTTTGAT GGGGCGGGGA TGTAAGCGGG AAGGGAAACC GACCCGTTTA GTCTGCCGCT CCCAACAGCC CGAGTTTCCA GCCCCTTGAG TAAGGGGCAA GAAACTAAG CCGCCTGGTG GATGGCTCGG CTCGGGGCGC TGACGAAGGG CGTGGC
J23SpCA_3                                                       C CCGCAGAAGA CGGGTTTGAT GGGGCGGGGA TGTAAGCGGG AAGGGAAACC GACCCGTTTA GTCTGCCGCT CCCAACAGCC CGAGTTTCCA GCCCCTTGAG TAAGGGGCAA GAAACTAAG CCGCCTGGTG GATGGCTCGG CTCGGGGCGC TGACGAAGGG CGTGGCG
J23SpCA_7                                                         CCGCAGAAGA CGGGTTTGAT GGGGCGGGGA TGTAAGCGGG AAGGGAAACC GACCCGTTTA GTCTGCCGCT CCCAACAGCC CGAGTTTCCA GCCCCTTGAG TAAGGGGCAA GAAACTAAG CCGCCTGGTG GATGGCTCGG CTCGGGGCGC TGACGAAGGG CGTGGCG
J23SpCA_48                                                        CCGCAGAAGA CGGGTTTGAT GGGGCGGGGA TGTAAGCGGG AAGGGAAACC GACCCGTTTA GTCTGCCGCT CCCAACAGCC CGAGTTTCCA GCCCCTTGAG TAAGGGGCAA GAAACTAAG CCGCCTGGTG GATGGCTCGG CTCGGGGCGC TGACGAAGGG CGTGGC
J23SpCA_36                                                         CGCAGAAGA CGGGTTTGAT GGGGCGGGGA TGTAAGCGGG AAGGGAAACC GACCCGTTTA GTCTGCCGCT CCCAACAGCC CGAGTTTCCA GCCCCTTGAG TAAGGGGCAA GAAACTAAG CCGCCTGGTG GATGGCTCGG CTCGGGGCGC TGACGAAGGG CGTGGCAAG
J23SpCA_11                                                        CCGCAGAAGA CGGGTTTGAT GGGGCGGGGA TGTAAGCGGG AAGGGAAACC GACCCGTTTA GTCTGCCGCT CCCAACAGCC CGAGTTTCCA GCCCCTTGAG TAAGGGGCAA GAAACTAAG CCGCCTGGTG GATGGCTCGG CTCGGGGCGC TGACGAAGGG CGTGGCAAG
J23SpCA_5                                                          CGCAGAAGA CGGGTTTGAT GGGGCGGGGA TGTAAGCGGG AAGGGAAACC GACCCGTTTA GTCTGCCGCT CCCAACAGCC CGAGTTTCCA GCCCCTTGAG TAAGGGGCAA GAAACTAAG CCGCCTGGTG GATGGCTCGG CTCGGGGCGC TGACGAAGGG CGTGGCAAGC
J23SpCA_3                                                          CGCAGAAGA CGGGTTTGAT GGGGCGGGGA TGTAAGCGGG AAGGGAAACC GACCCGTTTA GTCTGCCGCT CCCAACAGCC CGAGTTTCCA GCCCCTTGAG TAAGGGGCAA GAAACTAAG CCGCCTGGTG GATGGCTCGG CTCGGGGCGC TGACGAAGGG CGTGGCAAGC C
J23SpCA_4                                                         CCGCAGAAGA CGGGTTTGAT GGGGCGGGGA TGTAAGCGGG AAGGGAAACC GACCCGTTTA GTCTGCCGCT CCCAACAGCC CGAGTTTCCA GCCCCTTGAG TAAGGGGCAA GAAACTAAG CCGCCTGGTG GATGGCTCGG CTCGGGGCGC TGACGAAGGG CGTGGCAA
J23SpCA_7                                                            CAGAAGA CGGGTTTGAT GGGGCGGGGA TGTAAGCGGG AAGGGAAACC GACCCGTTTA GTCTGCCGCT CCCAACAGCC CGAGTTTCCA GCCCCTTGAG TAAGGGGCAA GAAACTAAG CCGCCTGGTG GATGGCTCGG CTCGGGGCGC TGACGAAGGG CGTGGCAAGC
J23SpCA_5                                                            CAGAAGA CGGGTTTGAT GGGGCGGGGA TGTAAGCGGG AAGGGAAACC GACCCGTTTA GTCTGCCGCT CCCAACAGCC CGAGTTTCCA GCCCCTTGAG TAAGGGGCAA GAAACTAAG CCGCCTGGTG GATGGCTCGG CTCGGGGCGC TGACGAAGGG CGTGGCAAGC TG
J23SpCA_4                                                         CCGCAGAAGA CGGGTTTGAT GGGGCGGGGA TGTAAGCGGG AAGGGAAACC GACCCGTTTA GTCTGCCGCT CCCAACAGCC CGAGTTTCCA GCCCCTTGAG TAAGGGGCAA GAAACTAAG CCGCCTGGTG GATGGCTCGG CTCGGGGCGC TGACGAAGGG CGTGGCA
J23SpCA_96                                                         CGCAGAAGA CGGGTTTGAT GGGGCGGGGA TGTAAGCGGG AAGGGAAACC GACCCGTTTA GTCTGCCGCT CCCAACAGCC CGAGTTTCCA GCCCCTTGAG TAAGGGGCAA GAAACTAAG CCGCCTGGTG GATGGCTCGG CTCGGGGCGC TGACGAAGGG CGTGGC
J23SpCA_14                                                           CAGAAGA CGGGTTTGAT GGGGCGGGGA TGTAAGCGGG AAGGGAAACC GACCCGTTTA GTCTGCCGCT CCCAACAGCC CGAGTTTCCA GCCCCTTGAG TAAGGGGCAA GAAACTAAG CCGCCTGGTG GATGGCTCGG CTCGGGGCGC TGACGAAGGG CGTGGCAAG
J23SpCA_9                                                          CGCAGAAGA CGGGTTTGAT GGGGCGGGGA TGTAAGCGGG AAGGGAAACC GACCCGTTTA GTCTGCCGCT CCCAACAGCC CGAGTTTCCA GCCCCTTGAG TAAGGGGCAA GAAACTAAG CCGCCTGGTG GATGGCTCGG CTCGGGGCGC TGACGAAGGG CGTGGCG
```

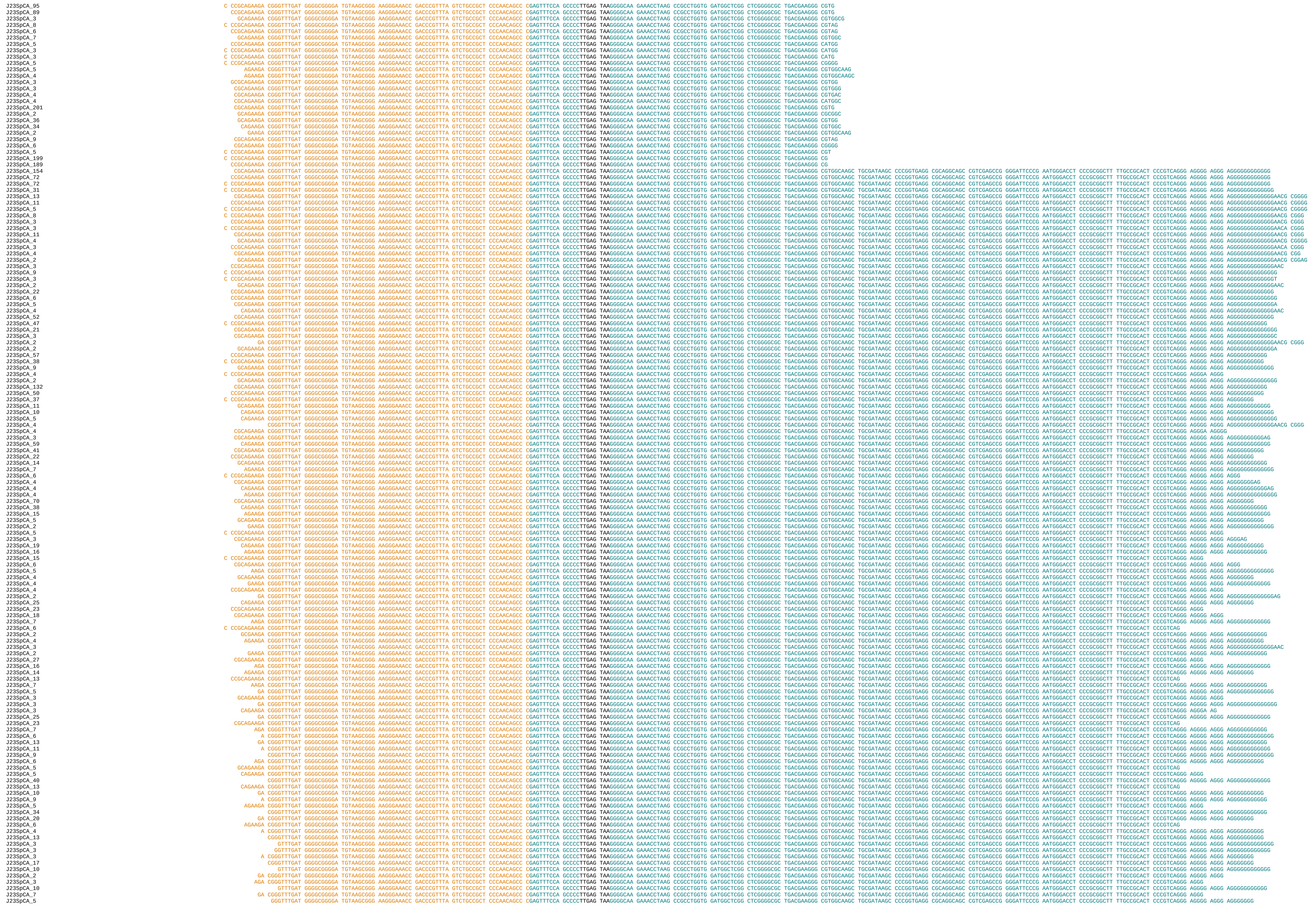

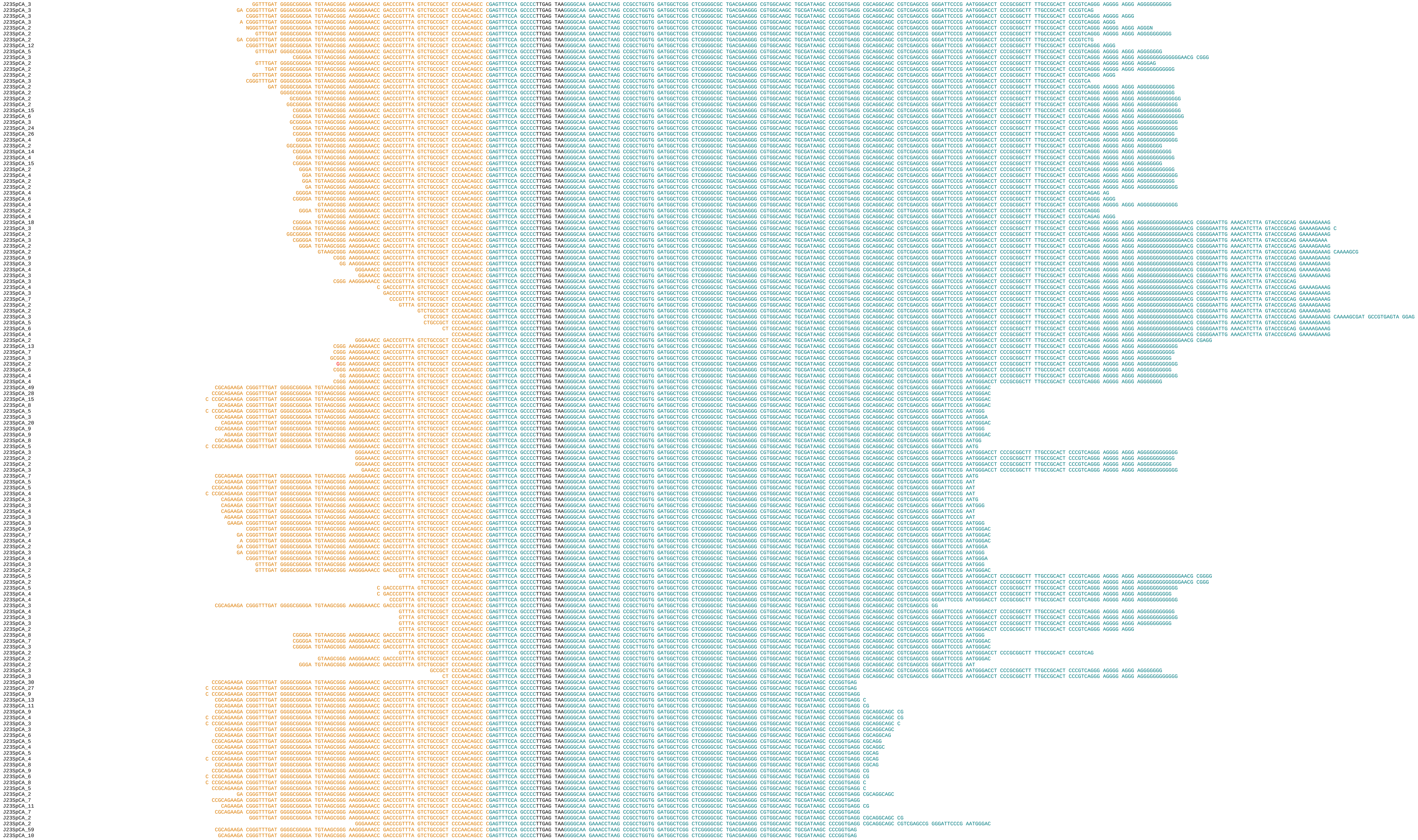

```
J23SpCA_23 CAGAAGA CGGGTTTGAT GGGGCGGGGA TGTAAGCGGG AAGGGAAACC GACCCGTTTA GTCTGCCGCT CCCAACAGCC CGAGTTTCCA GCCCCTTGAG TAAGGGGCAA GAAACTAAGC CCGCCTGGTG GATGGCTCGG CTCGGGGCGC TGACGAAGGG CGTGGCAAGC TGCGATAAGC CCCGGTGAG
J23SpCA_10 AGAAGA CGGGTTTGAT GGGGCGGGGA TGTAAGCGGG AAGGGAAACC GACCCGTTTA GTCTGCCGCT CCCAACAGCC CGAGTTTCCA GCCCCTTGAG TAAGGGGCAA GAAACTAAGC CCGCCTGGTG GATGGCTCGG CTCGGGGCGC TGACGAAGGG CGTGGCAAGC TGCGATAAGC CCCGGTGAG
J23SpCA_3 CGCAGAAGA CGGGTTTGAT GGGGCGGGGA TGTAAGCGGG AAGGGAAACC GACCCGTTTA GTCTGCCGCT CCCAACAGCC CGAGTTTCCA GCCCCTTGAG TAAGGGGCAA GAAACTAAGC CCGCCTGGTG GATGGCTCGG CTCGGGGCGC TGACGAAGGG CGTGGCAAGC TGCGATAAGC CCCGGTG
J23SpCA_3 CCGCAGAAGA CGGGTTTGAT GGGGCGGGGA TGTAAGCGGG AAGGGAAACC GACCCGTTTA GTCTGCCGCT CCCAACAGCC CGAGTTTCCA GCCCCTTGAG TAAGGGGCAA GAAACTAAGC CCGCCTGGTG GATGGCTCGG CTCGGGGCGC TGACGAAGGG CGTGGCAAGC TGCGATAAGC CCCGGTG
J23SpCA_2 GCAGAAGA CGGGTTTGAT GGGGCGGGGA TGTAAGCGGG AAGGGAAACC GACCCGTTTA GTCTGCCGCT CCCAACAGCC CGAGTTTCCA GCCCCTTGAG TAAGGGGCAA GAAACTAAGC CCGCCTGGTG GATGGCTCGG CTCGGGGCGC TGACGAAGGG CGTGGCAAGC TGCGATAAGC CCCGGTG
J23SpCA_11 CCGCAGAAGA CGGGTTTGAT GGGGCGGGGA TGTAAGCGGG AAGGGAAACC GACCCGTTTA GTCTGCCGCT CCCAACAGCC CGAGTTTCCA GCCCCTTGAG TAAGGGGCAA GAAACTAAGC CCGCCTGGTG GATGGCTCGG CTCGGGGCGC TGACGAAGGG CGTGGCAAGC TGCGATAAGC CCCGG
J23SpCA_9 C CCGCAGAAGA CGGGTTTGAT GGGGCGGGGA TGTAAGCGGG AAGGGAAACC GACCCGTTTA GTCTGCCGCT CCCAACAGCC CGAGTTTCCA GCCCCTTGAG TAAGGGGCAA GAAACTAAGC CCGCCTGGTG GATGGCTCGG CTCGGGGCGC TGACGAAGGG CGTGGCAAGC TGCGATAAGC CCCGG
J23SpCA_6 C CCGCAGAAGA CGGGTTTGAT GGGGCGGGGA TGTAAGCGGG AAGGGAAACC GACCCGTTTA GTCTGCCGCT CCCAACAGCC CGAGTTTCCA GCCCCTTGAG TAAGGGGCAA GAAACTAAGC CCGCCTGGTG GATGGCTCGG CTCGGGGCGC TGACGAAGGG CGTGGCAAGC TGCGATAAGC CCCG
J23SpCA_23 CGCAGAAGA CGGGTTTGAT GGGGCGGGGA TGTAAGCGGG AAGGGAAACC GACCCGTTTA GTCTGCCGCT CCCAACAGCC CGAGTTTCCA GCCCCTTGAG TAAGGGGCAA GAAACTAAGC CCGCCTGGTG GATGGCTCGG CTCGGGGCGC TGACGAAGGG CGTGGCAAGC TGCGATAAGC CCCGG
J23SpCA_8 CCGCAGAAGA CGGGTTTGAT GGGGCGGGGA TGTAAGCGGG AAGGGAAACC GACCCGTTTA GTCTGCCGCT CCCAACAGCC CGAGTTTCCA GCCCCTTGAG TAAGGGGCAA GAAACTAAGC CCGCCTGGTG GATGGCTCGG CTCGGGGCGC TGACGAAGGG CGTGGCAAGC TGCGATAAGC CCCG
J23SpCA_4 CAGAAGA CGGGTTTGAT GGGGCGGGGA TGTAAGCGGG AAGGGAAACC GACCCGTTTA GTCTGCCGCT CCCAACAGCC CGAGTTTCCA GCCCCTTGAG TAAGGGGCAA GAAACTAAGC CCGCCTGGTG GATGGCTCGG CTCGGGGCGC TGACGAAGGG CGTGGCAAGC TGCGATAAGC CCCGGTG
J23SpCA_3 GCAGAAGA CGGGTTTGAT GGGGCGGGGA TGTAAGCGGG AAGGGAAACC GACCCGTTTA GTCTGCCGCT CCCAACAGCC CGAGTTTCCA GCCCCTTGAG TAAGGGGCAA GAAACTAAGC CCGCCTGGTG GATGGCTCGG CTCGGGGCGC TGACGAAGGG CGTGGCAAGC TGCGATAAGC CCCGG
J23SpCA_15 CGCAGAAGA CGGGTTTGAT GGGGCGGGGA TGTAAGCGGG AAGGGAAACC GACCCGTTTA GTCTGCCGCT CCCAACAGCC CGAGTTTCCA GCCCCTTGAG TAAGGGGCAA GAAACTAAGC CCGCCTGGTG GATGGCTCGG CTCGGGGCGC TGACGAAGGG CGTGGCAAGC TGCGATAAGC CCCG
J23SpCA_2 GCAGAAGA CGGGTTTGAT GGGGCGGGGA TGTAAGCGGG AAGGGAAACC GACCCGTTTA GTCTGCCGCT CCCAACAGCC CGAGTTTCCA GCCCCTTGAG TAAGGGGCAA GAAACTAAGC CCGCCTGGTG GATGGCTCGG CTCGGGGCGC TGACGAAGGG CGTGGCAAGC TGCGATAAGC CCCG
J23SpCA_8 CAGAAGA CGGGTTTGAT GGGGCGGGGA TGTAAGCGGG AAGGGAAACC GACCCGTTTA GTCTGCCGCT CCCAACAGCC CGAGTTTCCA GCCCCTTGAG TAAGGGGCAA GAAACTAAGC CCGCCTGGTG GATGGCTCGG CTCGGGGCGC TGACGAAGGG CGTGGCAAGC TGCGATAAGC CCCGG
J23SpCA_4 AGAAGA CGGGTTTGAT GGGGCGGGGA TGTAAGCGGG AAGGGAAACC GACCCGTTTA GTCTGCCGCT CCCAACAGCC CGAGTTTCCA GCCCCTTGAG TAAGGGGCAA GAAACTAAGC CCGCCTGGTG GATGGCTCGG CTCGGGGCGC TGACGAAGGG CGTGGCAAGC TGCGATAAGC CCCGGCGAG
J23SpCA_3 GAAGA CGGGTTTGAT GGGGCGGGGA TGTAAGCGGG AAGGGAAACC GACCCGTTTA GTCTGCCGCT CCCAACAGCC CGAGTTTCCA GCCCCTTGAG TAAGGGGCAA GAAACTAAGC CCGCCTGGTG GATGGCTCGG CTCGGGGCGC TGACGAAGGG CGTGGCAAGC TGCGATAAGC CCCGGTGAG
J23SpCA_3 AAGA CGGGTTTGAT GGGGCGGGGA TGTAAGCGGG AAGGGAAACC GACCCGTTTA GTCTGCCGCT CCCAACAGCC CGAGTTTCCA GCCCCTTGAG TAAGGGGCAA GAAACTAAGC CCGCCTGGTG GATGGCTCGG CTCGGGGCGC TGACGAAGGG CGTGGCAAGC TGCGATAAGC CCCGGTGAG
J23SpCA_4 CAGAAGA CGGGTTTGAT GGGGCGGGGA TGTAAGCGGG AAGGGAAACC GACCCGTTTA GTCTGCCGCT CCCAACAGCC CGAGTTTCCA GCCCCTTGAG TAAGGGGCAA GAAACTAAGC CCGCCTGGTG GATGGCTCGG CTCGGGGCGC TGACGAAGGG CGTGGCAAGC TGCGATAAGC CCCG
J23SpCA_8 GA CGGGTTTGAT GGGGCGGGGA TGTAAGCGGG AAGGGAAACC GACCCGTTTA GTCTGCCGCT CCCAACAGCC CGAGTTTCCA GCCCCTTGAG TAAGGGGCAA GAAACTAAGC CCGCCTGGTG GATGGCTCGG CTCGGGGCGC TGACGAAGGG CGTGGCAAGC TGCGATAAGC CCCGGTGAG
J23SpCA_5 AGA CGGGTTTGAT GGGGCGGGGA TGTAAGCGGG AAGGGAAACC GACCCGTTTA GTCTGCCGCT CCCAACAGCC CGAGTTTCCA GCCCCTTGAG TAAGGGGCAA GAAACTAAGC CCGCCTGGTG GATGGCTCGG CTCGGGGCGC TGACGAAGGG CGTGGCAAGC TGCGATAAGC CCCGGTGAG
J23SpCA_14 CGGGTTTGAT GGGGCGGGGA TGTAAGCGGG AAGGGAAACC GACCCGTTTA GTCTGCCGCT CCCAACAGCC CGAGTTTCCA GCCCCTTGAG TAAGGGGCAA GAAACTAAGC CCGCCTGGTG GATGGCTCGG CTCGGGGCGC TGACGAAGGG CGTGGCAAGC TGCGATAAGC CCCGGTGAG
J23SpCA_6 A CGGGTTTGAT GGGGCGGGGA TGTAAGCGGG AAGGGAAACC GACCCGTTTA GTCTGCCGCT CCCAACAGCC CGAGTTTCCA GCCCCTTGAG TAAGGGGCAA GAAACTAAGC CCGCCTGGTG GATGGCTCGG CTCGGGGCGC TGACGAAGGG CGTGGCAAGC TGCGATAAGC CCCGGTGAGG
J23SpCA_3 CGGGTTTGAT GGGGCGGGGA TGTAAGCGGG AAGGGAAACC GACCCGTTTA GTCTGCCGCT CCCAACAGCC CGAGTTTCCA GCCCCTTGAG TAAGGGGCAA GAAACTAAGC CCGCCTGGTG GATGGCTCGG CTCGGGGCGC TGACGAAGGG CGTGGCAAGC TGCGATAAGC CCCGGTGAGG CGCAG
J23SpCA_3 A CGGGTTTGAT GGGGCGGGGA TGTAAGCGGG AAGGGAAACC GACCCGTTTA GTCTGCCGCT CCCAACAGCC CGAGTTTCCA GCCCCTTGAG TAAGGGGCAA GAAACTAAGC CCGCCTGGTG GATGGCTCGG CTCGGGGCGC TGACGAAGGG CGTGGCAAGC TGCGATAAGC CCCGGTGAGG CGCAGG
J23SpCA_3 CGGGTTTGAT GGGGCGGGGA TGTAAGCGGG AAGGGAAACC GACCCGTTTA GTCTGCCGCT CCCAACAGCC CGAGTTTCCA GCCCCTTGAG TAAGGGGCAA GAAACTAAGC CCGCCTGGTG GATGGCTCGG CTCGGGGCGC TGACGAAGGG CGTGGCAAGC TGCGATAAGC CCCGGTGAGG C
J23SpCA_3 CGGGTTTGAT GGGGCGGGGA TGTAAGCGGG AAGGGAAACC GACCCGTTTA GTCTGCCGCT CCCAACAGCC CGAGTTTCCA GCCCCTTGAG TAAGGGGCAA GAAACTAAGC CCGCCTGGTG GATGGCTCGG CTCGGGGCGC TGACGAAGGG CGTGGCAAGC TGCGATAAGC CCCGGTGAGG
J23SpCA_4 GTTTGAT GGGGCGGGGA TGTAAGCGGG AAGGGAAACC GACCCGTTTA GTCTGCCGCT CCCAACAGCC CGAGTTTCCA GCCCCTTGAG TAAGGGGCAA GAAACTAAGC CCGCCTGGTG GATGGCTCGG CTCGGGGCGC TGACGAAGGG CGTGGCAAGC TGCGATAAGC CCCGGTGAGG CG
J23SpCA_2 GTTTGAT GGGGCGGGGA TGTAAGCGGG AAGGGAAACC GACCCGTTTA GTCTGCCGCT CCCAACAGCC CGAGTTTCCA GCCCCTTGAG TAAGGGGCAA GAAACTAAGC CCGCCTGGTG GATGGCTCGG CTCGGGGCGC TGACGAAGGG CGTGGCAAGC TGCGATAAGC CCCGGTGAGG C
J23SpCA_7 GTTTGAT GGGGCGGGGA TGTAAGCGGG AAGGGAAACC GACCCGTTTA GTCTGCCGCT CCCAACAGCC CGAGTTTCCA GCCCCTTGAG TAAGGGGCAA GAAACTAAGC CCGCCTGGTG GATGGCTCGG CTCGGGGCGC TGACGAAGGG CGTGGCAAGC TGCGATAAGC CCCGGTGAG
J23SpCA_2 GTTTGAT GGGGCGGGGA TGTAAGCGGG AAGGGAAACC GACCCGTTTA GTCTGCCGCT CCCAACAGCC CGAGTTTCCA GCCCCTTGAG TAAGGGGCAA GAAACTAAGC CCGCCTGGTG GATGGCTCGG CTCGGGGCGC TGACGAAGGG CGTGGCAAGC TGCGATAAGC CCCGGTGAGG
J23SpCA_4 CGGGTTTGAT GGGGCGGGGA TGTAAGCGGG AAGGGAAACC GACCCGTTTA GTCTGCCGCT CCCAACAGCC CGAGTTTCCA GCCCCTTGAG TAAGGGGCAA GAAACTAAGC CCGCCTGGTG GATGGCTCGG CTCGGGGCGC TGACGAAGGG CGTGGCAAGC TGCGATAAGC CCCGG
J23SpCA_3 CGGGTTTGAT GGGGCGGGGA TGTAAGCGGG AAGGGAAACC GACCCGTTTA GTCTGCCGCT CCCAACAGCC CGAGTTTCCA GCCCCTTGAG TAAGGGGCAA GAAACTAAGC CCGCCTGGTG GATGGCTCGG CTCGGGGCGC TGACGAAGGG CGTGGCAAGC TGCGATAAGC CCCGGTG
J23SpCA_3 GA CGGGTTTGAT GGGGCGGGGA TGTAAGCGGG AAGGGAAACC GACCCGTTTA GTCTGCCGCT CCCAACAGCC CGAGTTTCCA GCCCCTTGAG TAAGGGGCAA GAAACTAAGC CCGCCTGGTG GATGGCTCGG CTCGGGGCGC TGACGAAGGG CGTGGCAAGC TGCGATAAGC CCCGG
J23SpCA_3 GA CGGGTTTGAT GGGGCGGGGA TGTAAGCGGG AAGGGAAACC GACCCGTTTA GTCTGCCGCT CCCAACAGCC CGAGTTTCCA GCCCCTTGAG TAAGGGGCAA GAAACTAAGC CCGCCTGGTG GATGGCTCGG CTCGGGGCGC TGACGAAGGG CGTGGCAAGC TGCGATAAGC CCCG
J23SpCA_4 CGGGTTTGAT GGGGCGGGGA TGTAAGCGGG AAGGGAAACC GACCCGTTTA GTCTGCCGCT CCCAACAGCC CGAGTTTCCA GCCCCTTGAG TAAGGGGCAA GAAACTAAGC CCGCCTGGTG GATGGCTCGG CTCGGGGCGC TGACGAAGGG CGTGGCAAGC TGCGATAAGC CCCG
J23SpCA_4 GTTTGAT GGGGCGGGGA TGTAAGCGGG AAGGGAAACC GACCCGTTTA GTCTGCCGCT CCCAACAGCC CGAGTTTCCA GCCCCTTGAG TAAGGGGCAA GAAACTAAGC CCGCCTGGTG GATGGCTCGG CTCGGGGCGC TGACGAAGGG CGTGGCAAGC TGCGATAAGC CCCGG
J23SpCA_2 GGCGGGGA TGTAAGCGGG AAGGGAAACC GACCCGTTTA GTCTGCCGCT CCCAACAGCC CGAGTTTCCA GCCCCTTGAG TAAGGGGCAA GAAACTAAGC CCGCCTGGTG GATGGCTCGG CTCGGGGCGC TGACGAAGGG CGTGGCAAGC TGCGATAAGC CCCGGTGAGG CGCAG
J23SpCA_58 CGCAGAAGA CGGGTTTGAT GGGGCGGGGA TGTAAGCGGG AAGGGAAACC GACCCGTTTA GTCTGCCGCT CCCAACAGCC CGAGTTTCCA GCCCCTTGAG TAAGGGGCAA GAAACTAAGC CCGCCTGGTG GATGGCTCGG CTCGGGGCGC TGACGAAGGG CGTGGCAAGC TGCG
J23SpCA_26 CGCAGAAGA CGGGTTTGAT GGGGCGGGGA TGTAAGCGGG AAGGGAAACC GACCCGTTTA GTCTGCCGCT CCCAACAGCC CGAGTTTCCA GCCCCTTGAG TAAGGGGCAA GAAACTAAGC CCGCCTGGTG GATGGCTCGG CTCGGGGCGC TGACGAAGGG CGTGGCAAGC TGCGAT
J23SpCA_20 C CCGCAGAAGA CGGGTTTGAT GGGGCGGGGA TGTAAGCGGG AAGGGAAACC GACCCGTTTA GTCTGCCGCT CCCAACAGCC CGAGTTTCCA GCCCCTTGAG TAAGGGGCAA GAAACTAAGC CCGCCTGGTG GATGGCTCGG CTCGGGGCGC TGACGAAGGG CGTGGCAAGC TGCGAT
J23SpCA_7 CCGCAGAAGA CGGGTTTGAT GGGGCGGGGA TGTAAGCGGG AAGGGAAACC GACCCGTTTA GTCTGCCGCT CCCAACAGCC CGAGTTTCCA GCCCCTTGAG TAAGGGGCAA GAAACTAAGC CCGCCTGGTG GATGGCTCGG CTCGGGGCGC TGACGAAGGG CGTGGCAAGC TGCGAT
J23SpCA_5 C CCGCAGAAGA CGGGTTTGAT GGGGCGGGGA TGTAAGCGGG AAGGGAAACC GACCCGTTTA GTCTGCCGCT CCCAACAGCC CGAGTTTCCA GCCCCTTGAG TAAGGGGCAA GAAACTAAGC CCGCCTGGTG GATGGCTCGG CTCGGGGCGC TGACGAAGGG CGTGGCAAGC TGCGAG
J23SpCA_21 C CCGCAGAAGA CGGGTTTGAT GGGGCGGGGA TGTAAGCGGG AAGGGAAACC GACCCGTTTA GTCTGCCGCT CCCAACAGCC CGAGTTTCCA GCCCCTTGAG TAAGGGGCAA GAAACTAAGC CCGCCTGGTG GATGGCTCGG CTCGGGGCGC TGACGAAGGG CGTGGCAAGC TGCG
J23SpCA_21 CCGCAGAAGA CGGGTTTGAT GGGGCGGGGA TGTAAGCGGG AAGGGAAACC GACCCGTTTA GTCTGCCGCT CCCAACAGCC CGAGTTTCCA GCCCCTTGAG TAAGGGGCAA GAAACTAAGC CCGCCTGGTG GATGGCTCGG CTCGGGGCGC TGACGAAGGG CGTGGCAAGC TGCG
J23SpCA_7 GCAGAAGA CGGGTTTGAT GGGGCGGGGA TGTAAGCGGG AAGGGAAACC GACCCGTTTA GTCTGCCGCT CCCAACAGCC CGAGTTTCCA GCCCCTTGAG TAAGGGGCAA GAAACTAAGC CCGCCTGGTG GATGGCTCGG CTCGGGGCGC TGACGAAGGG CGTGGCAAGC TGCGAT
J23SpCA_3 CGCAGAAGA CGGGTTTGAT GGGGCGGGGA TGTAAGCGGG AAGGGAAACC GACCCGTTTA GTCTGCCGCT CCCAACAGCC CGAGTTTCCA GCCCCTTGAG TAAGGGGCAA GAAACTAAGC CCGCCTGGTG GATGGCTCGG CTCGGGGCGC TGACGAAGGG CGTGGCAAGC TGCGAG
J23SpCA_6 C CCGCAGAAGA CGGGTTTGAT GGGGCGGGGA TGTAAGCGGG AAGGGAAACC GACCCGTTTA GTCTGCCGCT CCCAACAGCC CGAGTTTCCA GCCCCTTGAG TAAGGGGCAA GAAACTAAGC CCGCCTGGTG GATGGCTCGG CTCGGGGCGC TGACGAAGGG CGTGGCAAGC TGC
J23SpCA_3 C CCGCAGAAGA CGGGTTTGAT GGGGCGGGGA TGTAAGCGGG AAGGGAAACC GACCCGTTTA GTCTGCCGCT CCCAACAGCC CGAGTTTCCA GCCCCTTGAG TAAGGGGCAA GAAACTAAGC CCGCCTGGTG GATGGCTCGG CTCGGGGCGC TGACGAAGGG CGTGGCAAGC TGCT
J23SpCA_3 CGCAGAAGA CGGGTTTGAT GGGGCGGGGA TGTAAGCGGG AAGGGAAACC GACCCGTTTA GTCTGCCGCT CCCAACAGCC CGAGTTTCCA GCCCCTTGAG TAAGGGGCAA GAAACTAAGC CCGCCTGGTG GATGGCTCGG CTCGGGGCGC TGACGAAGGG CGTGGCAAGC TGCGA
J23SpCA_11 CCGCAGAAGA CGGGTTTGAT GGGGCGGGGA TGTAAGCGGG AAGGGAAACC GACCCGTTTA GTCTGCCGCT CCCAACAGCC CGAGTTTCCA GCCCCTTGAG TAAGGGGCAA GAAACTAAGC CCGCCTGGTG GATGGCTCGG CTCGGGGCGC TGACGAAGGG CGTGGCAAGC TGC
J23SpCA_11 CAGAAGA CGGGTTTGAT GGGGCGGGGA TGTAAGCGGG AAGGGAAACC GACCCGTTTA GTCTGCCGCT CCCAACAGCC CGAGTTTCCA GCCCCTTGAG TAAGGGGCAA GAAACTAAGC CCGCCTGGTG GATGGCTCGG CTCGGGGCGC TGACGAAGGG CGTGGCAAGC TGCGAT
J23SpCA_10 C CCGCAGAAGA CGGGTTTGAT GGGGCGGGGA TGTAAGCGGG AAGGGAAACC GACCCGTTTA GTCTGCCGCT CCCAACAGCC CGAGTTTCCA GCCCCTTGAG TAAGGGGCAA GAAACTAAGC CCGCCTGGTG GATGGCTCGG CTCGGGGCGC TGACGAAGGG CGTGGCAAGC TG
J23SpCA_5 CGCAGAAGA CGGGTTTGAT GGGGCGGGGA TGTAAGCGGG AAGGGAAACC GACCCGTTTA GTCTGCCGCT CCCAACAGCC CGAGTTTCCA GCCCCTTGAG TAAGGGGCAA GAAACTAAGC CCGCCTGGTG GATGGCTCGG CTCGGGGCGC TGACGAAGGG CGTGGCAAGC TGCGG
J23SpCA_4 CCGCAGAAGA CGGGTTTGAT GGGGCGGGGA TGTAAGCGGG AAGGGAAACC GACCCGTTTA GTCTGCCGCT CCCAACAGCC CGAGTTTCCA GCCCCTTGAG TAAGGGGCAA GAAACTAAGC CCGCCTGGTG GATGGCTCGG CTCGGGGCGC TGACGAAGGG CGTGGCAAGC TGCAG
J23SpCA_15 CGCAGAAGA CGGGTTTGAT GGGGCGGGGA TGTAAGCGGG AAGGGAAACC GACCCGTTTA GTCTGCCGCT CCCAACAGCC CGAGTTTCCA GCCCCTTGAG TAAGGGGCAA GAAACTAAGC CCGCCTGGTG GATGGCTCGG CTCGGGGCGC TGACGAAGGG CGTGGCAAGC TGC
J23SpCA_11 CCGCAGAAGA CGGGTTTGAT GGGGCGGGGA TGTAAGCGGG AAGGGAAACC GACCCGTTTA GTCTGCCGCT CCCAACAGCC CGAGTTTCCA GCCCCTTGAG TAAGGGGCAA GAAACTAAGC CCGCCTGGTG GATGGCTCGG CTCGGGGCGC TGACGAAGGG CGTGGCAAGC TG
J23SpCA_8 GCAGAAGA CGGGTTTGAT GGGGCGGGGA TGTAAGCGGG AAGGGAAACC GACCCGTTTA GTCTGCCGCT CCCAACAGCC CGAGTTTCCA GCCCCTTGAG TAAGGGGCAA GAAACTAAGC CCGCCTGGTG GATGGCTCGG CTCGGGGCGC TGACGAAGGG CGTGGCAAGC TGCG
J23SpCA_4 CGCAGAAGA CGGGTTTGAT GGGGCGGGGA TGTAAGCGGG AAGGGAAACC GACCCGTTTA GTCTGCCGCT CCCAACAGCC CGAGTTTCCA GCCCCTTGAG TAAGGGGCAA GAAACTAAGC CCGCCTGGTG GATGGCTCGG CTCGGGGCGC TGACGAAGGG CGTGGCAAGC TGCT
J23SpCA_4 C CCGCAGAAGA CGGGTTTGAT GGGGCGGGGA TGTAAGCGGG AAGGGAAACC GACCCGTTTA GTCTGCCGCT CCCAACAGCC CGAGTTTCCA GCCCCTTGAG TAAGGGGCAA GAAACTAAGC CCGCCTGGTG GATGGCTCGG CTCGGGGCGC TGACGAAGGG CGTGGCAAGC AG
J23SpCA_2 GCAGAAGA CGGGTTTGAT GGGGCGGGGA TGTAAGCGGG AAGGGAAACC GACCCGTTTA GTCTGCCGCT CCCAACAGCC CGAGTTTCCA GCCCCTTGAG TAAGGGGCAA GAAACTAAGC CCGCCTGGTG GATGGCTCGG CTCGGGGCGC TGACGAAGGG CGTGGCAAGC TGC
J23SpCA_26 CGCAGAAGA CGGGTTTGAT GGGGCGGGGA TGTAAGCGGG AAGGGAAACC GACCCGTTTA GTCTGCCGCT CCCAACAGCC CGAGTTTCCA GCCCCTTGAG TAAGGGGCAA GAAACTAAGC CCGCCTGGTG GATGGCTCGG CTCGGGGCGC TGACGAAGGG CGTGGCAAGC TG
J23SpCA_25 CAGAAGA CGGGTTTGAT GGGGCGGGGA TGTAAGCGGG AAGGGAAACC GACCCGTTTA GTCTGCCGCT CCCAACAGCC CGAGTTTCCA GCCCCTTGAG TAAGGGGCAA GAAACTAAGC CCGCCTGGTG GATGGCTCGG CTCGGGGCGC TGACGAAGGG CGTGGCAAGC TGCG
J23SpCA_3 GAAGA CGGGTTTGAT GGGGCGGGGA TGTAAGCGGG AAGGGAAACC GACCCGTTTA GTCTGCCGCT CCCAACAGCC CGAGTTTCCA GCCCCTTGAG TAAGGGGCAA GAAACTAAGC CCGCCTGGTG GATGGCTCGG CTCGGGGCGC TGACGAAGGG CGTGGCAAGC TGCGAT
J23SpCA_2 GCAGAAGA CGGGTTTGAT GGGGCGGGGA TGTAAGCGGG AAGGGAAACC GACCCGTTTA GTCTGCCGCT CCCAACAGCC CGAGTTTCCA GCCCCTTGAG TAAGGGGCAA GAAACTAAGC CCGCCTGGTG GATGGCTCGG CTCGGGGCGC TGACGAAGGG CGTGGCAAGC TGCA
J23SpCA_3 AGAAGA CGGGTTTGAT GGGGCGGGGA TGTAAGCGGG AAGGGAAACC GACCCGTTTA GTCTGCCGCT CCCAACAGCC CGAGTTTCCA GCCCCTTGAG TAAGGGGCAA GAAACTAAGC CCGCCTGGTG GATGGCTCGG CTCGGGGCGC TGACGAAGGG CGTGGCAAGC TGCGAG
J23SpCA_3 CCGCAGAAGA CGGGTTTGAT GGGGCGGGGA TGTAAGCGGG AAGGGAAACC GACCCGTTTA GTCTGCCGCT CCCAACAGCC CGAGTTTCCA GCCCCTTGAG TAAGGGGCAA GAAACTAAGC CCGCCTGGTG GATGGCTCGG CTCGGGGCGC TGACGAAGGG CGTGGCAAGC T
J23SpCA_5 C CCGCAGAAGA CGGGTTTGAT GGGGCGGGGA TGTAAGCGGG AAGGGAAACC GACCCGTTTA GTCTGCCGCT CCCAACAGCC CGAGTTTCCA GCCCCTTGAG TAAGGGGCAA GAAACTAAGC CCGCCTGGTG GATGGCTCGG CTCGGGGCGC TGACGAAGGG CGTGGCAAGC
J23SpCA_2 GCAGAAGA CGGGTTTGAT GGGGCGGGGA TGTAAGCGGG AAGGGAAACC GACCCGTTTA GTCTGCCGCT CCCAACAGCC CGAGTTTCCA GCCCCTTGAG TAAGGGGCAA GAAACTAAGC CCGCCTGGTG GATGGCTCGG CTCGGGGCGC TGACGAAGGG CGTGGCAAGC TG
J23SpCA_13 CCGCAGAAGA CGGGTTTGAT GGGGCGGGGA TGTAAGCGGG AAGGGAAACC GACCCGTTTA GTCTGCCGCT CCCAACAGCC CGAGTTTCCA GCCCCTTGAG TAAGGGGCAA GAAACTAAGC CCGCCTGGTG GATGGCTCGG CTCGGGGCGC TGACGAAGGG CGTGGCAAGC
J23SpCA_13 C CCGCAGAAGA CGGGTTTGAT GGGGCGGGGA TGTAAGCGGG AAGGGAAACC GACCCGTTTA GTCTGCCGCT CCCAACAGCC CGAGTTTCCA GCCCCTTGAG TAAGGGGCAA GAAACTAAGC CCGCCTGGTG GATGGCTCGG CTCGGGGCGC TGACGAAGGG CGTGGCAAG
J23SpCA_8 AGAAGA CGGGTTTGAT GGGGCGGGGA TGTAAGCGGG AAGGGAAACC GACCCGTTTA GTCTGCCGCT CCCAACAGCC CGAGTTTCCA GCCCCTTGAG TAAGGGGCAA GAAACTAAGC CCGCCTGGTG GATGGCTCGG CTCGGGGCGC TGACGAAGGG CGTGGCAAGC TGCG
J23SpCA_6 CAGAAGA CGGGTTTGAT GGGGCGGGGA TGTAAGCGGG AAGGGAAACC GACCCGTTTA GTCTGCCGCT CCCAACAGCC CGAGTTTCCA GCCCCTTGAG TAAGGGGCAA GAAACTAAGC CCGCCTGGTG GATGGCTCGG CTCGGGGCGC TGACGAAGGG CGTGGCAAGC TGC
J23SpCA_5 CGCAGAAGA CGGGTTTGAT GGGGCGGGGA TGTAAGCGGG AAGGGAAACC GACCCGTTTA GTCTGCCGCT CCCAACAGCC CGAGTTTCCA GCCCCTTGAG TAAGGGGCAA GAAACTAAGC CCGCCTGGTG GATGGCTCGG CTCGGGGCGC TGACGAAGGG CGTGGCAAGC T
J23SpCA_5 CGGGTTTGAT GGGGCGGGGA TGTAAGCGGG AAGGGAAACC GACCCGTTTA GTCTGCCGCT CCCAACAGCC CGAGTTTCCA GCCCCTTGAG TAAGGGGCAA GAAACTAAGC CCGCCTGGTG GATGGCTCGG CTCGGGGCGC TGACGAAGGG CGTGGCAAGC TGCGATAAGC C
J23SpCA_2 GTTTGAT GGGGCGGGGA TGTAAGCGGG AAGGGAAACC GACCCGTTTA GTCTGCCGCT CCCAACAGCC CGAGTTTCCA GCCCCTTGAG TAAGGGGCAA GAAACTAAGC CCGCCTGGTG GATGGCTCGG CTCGGGGCGC TGACGAAGGG CGTGGCAAGC TGCGATAAGC CCCG
J23SpCA_11 CGGGTTTGAT GGGGCGGGGA TGTAAGCGGG AAGGGAAACC GACCCGTTTA GTCTGCCGCT CCCAACAGCC CGAGTTTCCA GCCCCTTGAG TAAGGGGCAA GAAACTAAGC CCGCCTGGTG GATGGCTCGG CTCGGGGCGC TGACGAAGGG CGTGGCAAGC TGCG
J23SpCA_8 GA CGGGTTTGAT GGGGCGGGGA TGTAAGCGGG AAGGGAAACC GACCCGTTTA GTCTGCCGCT CCCAACAGCC CGAGTTTCCA GCCCCTTGAG TAAGGGGCAA GAAACTAAGC CCGCCTGGTG GATGGCTCGG CTCGGGGCGC TGACGAAGGG CGTGGCAAGC TGCGAT
J23SpCA_3 AGA CGGGTTTGAT GGGGCGGGGA TGTAAGCGGG AAGGGAAACC GACCCGTTTA GTCTGCCGCT CCCAACAGCC CGAGTTTCCA GCCCCTTGAG TAAGGGGCAA GAAACTAAGC CCGCCTGGTG GATGGCTCGG CTCGGGGCGC TGACGAAGGG CGTGGCAAGC TGCGG
J23SpCA_6 GA CGGGTTTGAT GGGGCGGGGA TGTAAGCGGG AAGGGAAACC GACCCGTTTA GTCTGCCGCT CCCAACAGCC CGAGTTTCCA GCCCCTTGAG TAAGGGGCAA GAAACTAAGC CCGCCTGGTG GATGGCTCGG CTCGGGGCGC TGACGAAGGG CGTGGCAAGC TGCG
J23SpCA_5 CGGGTTTGAT GGGGCGGGGA TGTAAGCGGG AAGGGAAACC GACCCGTTTA GTCTGCCGCT CCCAACAGCC CGAGTTTCCA GCCCCTTGAG TAAGGGGCAA GAAACTAAGC CCGCCTGGTG GATGGCTCGG CTCGGGGCGC TGACGAAGGG CGTGGCAAGC TGCGAT
J23SpCA_5 GA CGGGTTTGAT GGGGCGGGGA TGTAAGCGGG AAGGGAAACC GACCCGTTTA GTCTGCCGCT CCCAACAGCC CGAGTTTCCA GCCCCTTGAG TAAGGGGCAA GAAACTAAGC CCGCCTGGTG GATGGCTCGG CTCGGGGCGC TGACGAAGGG CGTGGCAAGC TG
J23SpCA_3 CGGGTTTGAT GGGGCGGGGA TGTAAGCGGG AAGGGAAACC GACCCGTTTA GTCTGCCGCT CCCAACAGCC CGAGTTTCCA GCCCCTTGAG TAAGGGGCAA GAAACTAAGC CCGCCTGGTG GATGGCTCGG CTCGGGGCGC TGACGAAGGG CGTGGCAAGC TGCT
J23SpCA_4 GGGTTTGAT GGGGCGGGGA TGTAAGCGGG AAGGGAAACC GACCCGTTTA GTCTGCCGCT CCCAACAGCC CGAGTTTCCA GCCCCTTGAG TAAGGGGCAA GAAACTAAGC CCGCCTGGTG GATGGCTCGG CTCGGGGCGC TGACGAAGGG CGTGGCAAGC TGCG
J23SpCA_13 CGGGGA TGTAAGCGGG AAGGGAAACC GACCCGTTTA GTCTGCCGCT CCCAACAGCC CGAGTTTCCA GCCCCTTGAG TAAGGGGCAA GAAACTAAGC CCGCCTGGTG GATGGCTCGG CTCGGGGCGC TGACGAAGGG CGTGGCAAGC TGCGATAAGC CCCGGTGAG
J23SpCA_5 CGGGGA TGTAAGCGGG AAGGGAAACC GACCCGTTTA GTCTGCCGCT CCCAACAGCC CGAGTTTCCA GCCCCTTGAG TAAGGGGCAA GAAACTAAGC CCGCCTGGTG GATGGCTCGG CTCGGGGCGC TGACGAAGGG CGTGGCAAGC TGCGATAAGC CCCGGTGAGG
J23SpCA_3 CGGGGA TGTAAGCGGG AAGGGAAACC GACCCGTTTA GTCTGCCGCT CCCAACAGCC CGAGTTTCCA GCCCCTTGAG TAAGGGGCAA GAAACTAAGC CCGCCTGGTG GATGGCTCGG CTCGGGGCGC TGACGAAGGG CGTGGCAAGC TGCGATAAGC CCCGGTGAGG CG
J23SpCA_2 GGGCGGGGA TGTAAGCGGG AAGGGAAACC GACCCGTTTA GTCTGCCGCT CCCAACAGCC CGAGTTTCCA GCCCCTTGAG TAAGGGGCAA GAAACTAAGC CCGCCTGGTG GATGGCTCGG CTCGGGGCGC TGACGAAGGG CGTGGCAAGC TGCGATAAGC CCCGGTGAG
J23SpCA_2 GCGGGGA TGTAAGCGGG AAGGGAAACC GACCCGTTTA GTCTGCCGCT CCCAACAGCC CGAGTTTCCA GCCCCTTGAG TAAGGGGCAA GAAACTAAGC CCGCCTGGTG GATGGCTCGG CTCGGGGCGC TGACGAAGGG CGTGGCAAGC TGCGATAAGC CCCGGTGAG
J23SpCA_3 GGGGA TGTAAGCGGG AAGGGAAACC GACCCGTTTA GTCTGCCGCT CCCAACAGCC CGAGTTTCCA GCCCCTTGAG TAAGGGGCAA GAAACTAAGC CCGCCTGGTG GATGGCTCGG CTCGGGGCGC TGACGAAGGG CGTGGCAAGC TGCGATAAGC CCCGGTGAG
J23SpCA_2 GGA TGTAAGCGGG AAGGGAAACC GACCCGTTTA GTCTGCCGCT CCCAACAGCC CGAGTTTCCA GCCCCTTGAG TAAGGGGCAA GAAACTAAGC CCGCCTGGTG GATGGCTCGG CTCGGGGCGC TGACGAAGGG CGTGGCAAGC TGCGATAAGC CCCGGTGAGG C
J23SpCA_3 CGGGGA TGTAAGCGGG AAGGGAAACC GACCCGTTTA GTCTGCCGCT CCCAACAGCC CGAGTTTCCA GCCCCTTGAG TAAGGGGCAA GAAACTAAGC CCGCCTGGTG GATGGCTCGG CTCGGGGCGC TGACGAAGGG CGTGGCAAGC TGCGATAAGC CCCGG
J23SpCA_3 T GGGGCGGGGA TGTAAGCGGG AAGGGAAACC GACCCGTTTA GTCTGCCGCT CCCAACAGCC CGAGTTTCCA GCCCCTTGAG TAAGGGGCAA GAAACTAAGC CCGCCTGGTG GATGGCTCGG CTCGGGGCGC TGACGAAGGG CGTGGCAAGC TGCGATAAGC CCCGG
J23SpCA_2 GGGGCGGGGA TGTAAGCGGG AAGGGAAACC GACCCGTTTA GTCTGCCGCT CCCAACAGCC CGAGTTTCCA GCCCCTTGAG TAAGGGGCAA GAAACTAAGC CCGCCTGGTG GATGGCTCGG CTCGGGGCGC TGACGAAGGG CGTGGCAAGC TGCGATAAGC CCC
J23SpCA_4 GA CGGGTTTGAT GGGGCGGGGA TGTAAGCGGG AAGGGAAACC GACCCGTTTA GTCTGCCGCT CCCAACAGCC CGAGTTTCCA GCCCCTTGAG TAAGGGGCAA GAAACTAAGC CCGCCTGGTG GATGGCTCGG CTCGGGGCGC TGACGAAGGG CGTGGCAAG
J23SpCA_2 GA CGGGTTTGAT GGGGCGGGGA TGTAAGCGGG AAGGGAAACC GACCCGTTTA GTCTGCCGCT CCCAACAGCC CGAGTTTCCA GCCCCTTGAG TAAGGGGCAA GAAACTAAGC CCGCCTGGTG GATGGCTCGG CTCGGGGCGC TGACGAAGGG CGTGGCAAGC
J23SpCA_4 CGGGTTTGAT GGGGCGGGGA TGTAAGCGGG AAGGGAAACC GACCCGTTTA GTCTGCCGCT CCCAACAGCC CGAGTTTCCA GCCCCTTGAG TAAGGGGCAA GAAACTAAGC CCGCCTGGTG GATGGCTCGG CTCGGGGCGC TGACGAAGGG CGTGGCAAGC TG
J23SpCA_3 CGGGTTTGAT GGGGCGGGGA TGTAAGCGGG AAGGGAAACC GACCCGTTTA GTCTGCCGCT CCCAACAGCC CGAGTTTCCA GCCCCTTGAG TAAGGGGCAA GAAACTAAGC CCGCCTGGTG GATGGCTCGG CTCGGGGCGC TGACGAAGGG CGTGGCAAG
J23SpCA_3 GTTTGAT GGGGCGGGGA TGTAAGCGGG AAGGGAAACC GACCCGTTTA GTCTGCCGCT CCCAACAGCC CGAGTTTCCA GCCCCTTGAG TAAGGGGCAA GAAACTAAGC CCGCCTGGTG GATGGCTCGG CTCGGGGCGC TGACGAAGGG CGTGGCAAGC TGCG
J23SpCA_2 GTTTGAT GGGGCGGGGA TGTAAGCGGG AAGGGAAACC GACCCGTTTA GTCTGCCGCT CCCAACAGCC CGAGTTTCCA GCCCCTTGAG TAAGGGGCAA GAAACTAAGC CCGCCTGGTG GATGGCTCGG CTCGGGGCGC TGACGAAGGG CGTGGCAAGC TGC
J23SpCA_58 GA CGGGTTTGAT GGGGCGGGGA TGTAAGCGGG AAGGGAAACC GACCCGTTTA GTCTGCCGCT CCCAACAGCC CGAGTTTCCA GCCCCTTGAG TAAGGGGCAA GAAACTAAGC CCGCCTGGTG GATGGCTCGG CTCGGGGCGC TGACGAAGGG CGTGG
J23SpCA_18 AGA CGGGTTTGAT GGGGCGGGGA TGTAAGCGGG AAGGGAAACC GACCCGTTTA GTCTGCCGCT CCCAACAGCC CGAGTTTCCA GCCCCTTGAG TAAGGGGCAA GAAACTAAGC CCGCCTGGTG GATGGCTCGG CTCGGGGCGC TGACGAAGGG CGTGG
J23SpCA_6 AGA CGGGTTTGAT GGGGCGGGGA TGTAAGCGGG AAGGGAAACC GACCCGTTTA GTCTGCCGCT CCCAACAGCC CGAGTTTCCA GCCCCTTGAG TAAGGGGCAA GAAACTAAGC CCGCCTGGTG GATGGCTCGG CTCGGGGCGC TGACGAAGGG CGTGGC
J23SpCA_8 GA CGGGTTTGAT GGGGCGGGGA TGTAAGCGGG AAGGGAAACC GACCCGTTTA GTCTGCCGCT CCCAACAGCC CGAGTTTCCA GCCCCTTGAG TAAGGGGCAA GAAACTAAGC CCGCCTGGTG GATGGCTCGG CTCGGGGCGC TGACGAAGGG CGTGGC
J23SpCA_3 GA CGGGTTTGAT GGGGCGGGGA TGTAAGCGGG AAGGGAAACC GACCCGTTTA GTCTGCCGCT CCCAACAGCC CGAGTTTCCA GCCCCTTGAG TAAGGGGCAA GAAACTAAGC CCGCCTGGTG GATGGCTCGG CTCGGGGCGC TGACGAAGGG CGTGGCA
J23SpCA_3 A CGGGTTTGAT GGGGCGGGGA TGTAAGCGGG AAGGGAAACC GACCCGTTTA GTCTGCCGCT CCCAACAGCC CGAGTTTCCA GCCCCTTGAG TAAGGGGCAA GAAACTAAGC CCGCCTGGTG GATGGCTCGG CTCGGGGCGC TGACGAAGGG CGTGGC
J23SpCA_12 AGA CGGGTTTGAT GGGGCGGGGA TGTAAGCGGG AAGGGAAACC GACCCGTTTA GTCTGCCGCT CCCAACAGCC CGAGTTTCCA GCCCCTTGAG TAAGGGGCAA GAAACTAAGC CCGCCTGGTG GATGGCTCGG CTCGGGGCGC TGACGAAGGG CGTG
J23SpCA_5 AAGA CGGGTTTGAT GGGGCGGGGA TGTAAGCGGG AAGGGAAACC GACCCGTTTA GTCTGCCGCT CCCAACAGCC CGAGTTTCCA GCCCCTTGAG TAAGGGGCAA GAAACTAAGC CCGCCTGGTG GATGGCTCGG CTCGGGGCGC TGACGAAGGG CGTG
J23SpCA_4 GA CGGGTTTGAT GGGGCGGGGA TGTAAGCGGG AAGGGAAACC GACCCGTTTA GTCTGCCGCT CCCAACAGCC CGAGTTTCCA GCCCCTTGAG TAAGGGGCAA GAAACTAAGC CCGCCTGGTG GATGGCTCGG CTCGGGGCGC TGACGAAGGG CGGGG
J23SpCA_37 GA CGGGTTTGAT GGGGCGGGGA TGTAAGCGGG AAGGGAAACC GACCCGTTTA GTCTGCCGCT CCCAACAGCC CGAGTTTCCA GCCCCTTGAG TAAGGGGCAA GAAACTAAGC CCGCCTGGTG GATGGCTCGG CTCGGGGCGC TGACGAAGGG CGTG
J23SpCA_27 CGGGTTTGAT GGGGCGGGGA TGTAAGCGGG AAGGGAAACC GACCCGTTTA GTCTGCCGCT CCCAACAGCC CGAGTTTCCA GCCCCTTGAG TAAGGGGCAA GAAACTAAGC CCGCCTGGTG GATGGCTCGG CTCGGGGCGC TGACGAAGGG CGTGGC
J23SpCA_23 A CGGGTTTGAT GGGGCGGGGA TGTAAGCGGG AAGGGAAACC GACCCGTTTA GTCTGCCGCT CCCAACAGCC CGAGTTTCCA GCCCCTTGAG TAAGGGGCAA GAAACTAAGC CCGCCTGGTG GATGGCTCGG CTCGGGGCGC TGACGAAGGG CGTGG
J23SpCA_2 GGGTTTGAT GGGGCGGGGA TGTAAGCGGG AAGGGAAACC GACCCGTTTA GTCTGCCGCT CCCAACAGCC CGAGTTTCCA GCCCCTTGAG TAAGGGGCAA GAAACTAAGC CCGCCTGGTG GATGGCTCGG CTCGGGGCGC TGACGAAGGG CGTGGCAAG
J23SpCA_35 AGA CGGGTTTGAT GGGGCGGGGA TGTAAGCGGG AAGGGAAACC GACCCGTTTA GTCTGCCGCT CCCAACAGCC CGAGTTTCCA GCCCCTTGAG TAAGGGGCAA GAAACTAAGC CCGCCTGGTG GATGGCTCGG CTCGGGGCGC TGACGAAGGG CG
J23SpCA_21 AAGA CGGGTTTGAT GGGGCGGGGA TGTAAGCGGG AAGGGAAACC GACCCGTTTA GTCTGCCGCT CCCAACAGCC CGAGTTTCCA GCCCCTTGAG TAAGGGGCAA GAAACTAAGC CCGCCTGGTG GATGGCTCGG CTCGGGGCGC TGACGAAGGG CG
J23SpCA_12 GAAGA CGGGTTTGAT GGGGCGGGGA TGTAAGCGGG AAGGGAAACC GACCCGTTTA GTCTGCCGCT CCCAACAGCC CGAGTTTCCA GCCCCTTGAG TAAGGGGCAA GAAACTAAGC CCGCCTGGTG GATGGCTCGG CTCGGGGCGC TGACGAAGGG CG
J23SpCA_17 GAAGA CGGGTTTGAT GGGGCGGGGA TGTAAGCGGG AAGGGAAACC GACCCGTTTA GTCTGCCGCT CCCAACAGCC CGAGTTTCCA GCCCCTTGAG TAAGGGGCAA GAAACTAAGC CCGCCTGGTG GATGGCTCGG CTCGGGGCGC TGACGAAGGG C
J23SpCA_26 GAAGA CGGGTTTGAT GGGGCGGGGA TGTAAGCGGG AAGGGAAACC GACCCGTTTA GTCTGCCGCT CCCAACAGCC CGAGTTTCCA GCCCCTTGAG TAAGGGGCAA GAAACTAAGC CCGCCTGGTG GATGGCTCGG CTCGGGGCGC TGACGAAGGG
J23SpCA_25 AAGA CGGGTTTGAT GGGGCGGGGA TGTAAGCGGG AAGGGAAACC GACCCGTTTA GTCTGCCGCT CCCAACAGCC CGAGTTTCCA GCCCCTTGAG TAAGGGGCAA GAAACTAAGC CCGCCTGGTG GATGGCTCGG CTCGGGGCGC TGACGAAGGG C
J23SpCA_2 NGAAGA CGGGTTTGAT GGGGCGGGGA TGTAAGCGGG AAGGGAAACC GACCCGTTTA GTCTGCCGCT CCCAACAGCC CGAGTTTCCA GCCCCTTGAG TAAGGGGCAA GAAACTAAGC CCGCCTGGTG GATGGCTCGG CTCGGGGCGC TGACGAAGGG N
J23SpCA_2 GAAGA CGGGTTTGAT GGGGCGGGGA TGTAAGCGGG AAGGGAAACC GACCCGTTTA GTCTGCCGCT CCCAACAGCC CGAGTTTCCA GCCCCTTGAG TAAGGGGCAA GAAACTAAGC CCGCCTGGTG GATGGCTCGG CTCGGGGCGC TGACGAAGGA C
J23SpCA_434 CAGAAGA CGGGTTTGAT GGGGCGGGGA TGTAAGCGGG AAGGGAAACC GACCCGTTTA GTCTGCCGCT CCCAACAGCC CGAGTTTCCA GCCCCTTGAG TAAGGGGCAA GAAACTAAGC CCGCCTGGTG GATGGCTCGG CTCGGGGCGC TGACGAAG
J23SpCA_376 CAGAAGA CGGGTTTGAT GGGGCGGGGA TGTAAGCGGG AAGGGAAACC GACCCGTTTA GTCTGCCGCT CCCAACAGCC CGAGTTTCCA GCCCCTTGAG TAAGGGGCAA GAAACTAAGC CCGCCTGGTG GATGGCTCGG CTCGGGGCGC TGACGAAGG
J23SpCA_3 NCAGAAGA CGGGTTTGAT GGGGCGGGGA TGTAAGCGGG AAGGGAAACC GACCCGTTTA GTCTGCCGCT CCCAACAGCC CGAGTTTCCA GCCCCTTGAG TAAGGGGCAA GAAACTAAGC CCGCCTGGTG GATGGCTCGG CTCGGGGCGC TGACGAAGGN
J23SpCA_3 CAGAAGA CGGGTTTGAT GGGGCGGGGA TGTAAGCGGG AAGGGAAACC GACCCGTTTA GTCTGCCGCT CCCAACAGCC CGAGTTTCCA GCCCCTTGAG TAAGGGGCAA GAAACTAAGC CCGCCTGGTG GATGGCTCGG CTCGGGGCGC TGACGAAGGN
J23SpCA_4 NAGAAGA CGGGTTTGAT GGGGCGGGGA TGTAAGCGGG AAGGGAAACC GACCCGTTTA GTCTGCCGCT CCCAACAGCC CGAGTTTCCA GCCCCTTGAG TAAGGGGCAA GAAACTAAGC CCGCCTGGTG GATGGCTCGG CTCGGGGCGC TGACGAAGGN
J23SpCA_248 AGAAGA CGGGTTTGAT GGGGCGGGGA TGTAAGCGGG AAGGGAAACC GACCCGTTTA GTCTGCCGCT CCCAACAGCC CGAGTTTCCA GCCCCTTGAG TAAGGGGCAA GAAACTAAGC CCGCCTGGTG GATGGCTCGG CTCGGGGCGC TGACGAAGGG
J23SpCA_5 AGAAGA CGGGTTTGAT GGGGCGGGGA TGTAAGCGGG AAGGGAAACC GACCCGTTTA GTCTGCCGCT CCCAACAGCC CGAGTTTCCA GCCCCTTGAG TAAGGGGCAA GAAACTAAGC CCGCCTGGTG GATGGCTCGG CTCGGGGCGC TGACGAAGGG G
J23SpCA_4 NAGAAGA CGGGTTTGAT GGGGCGGGGA TGTAAGCGGG AAGGGAAACC GACCCGTTTA GTCTGCCGCT CCCAACAGCC CGAGTTTCCA GCCCCTTGAG TAAGGGGCAA GAAACTAAGC CCGCCTGGTG GATGGCTCGG CTCGGGGCGC TGACGAAGGG N
J23SpCA_150 AGAAGA CGGGTTTGAT GGGGCGGGGA TGTAAGCGGG AAGGGAAACC GACCCGTTTA GTCTGCCGCT CCCAACAGCC CGAGTTTCCA GCCCCTTGAG TAAGGGGCAA GAAACTAAGC CCGCCTGGTG GATGGCTCGG CTCGGGGCGC TGACGAAGG
J23SpCA_110 GCAGAAGA CGGGTTTGAT GGGGCGGGGA TGTAAGCGGG AAGGGAAACC GACCCGTTTA GTCTGCCGCT CCCAACAGCC CGAGTTTCCA GCCCCTTGAG TAAGGGGCAA GAAACTAAGC CCGCCTGGTG GATGGCTCGG CTCGGGGCGC TGACGAAG
J23SpCA_3 NGCAGAAGA CGGGTTTGAT GGGGCGGGGA TGTAAGCGGG AAGGGAAACC GACCCGTTTA GTCTGCCGCT CCCAACAGCC CGAGTTTCCA GCCCCTTGAG TAAGGGGCAA GAAACTAAGC CCGCCTGGTG GATGGCTCGG CTCGGGGCGC TGACGAAGN
J23SpCA_2 GGCAGAAGA CGGGTTTGAT GGGGCGGGGA TGTAAGCGGG AAGGGAAACC GACCCGTTTA GTCTGCCGCT CCCAACAGCC CGAGTTTCCA GCCCCTTGAG TAAGGGGCAA GAAACTAAGC CCGCCTGGTG GATGGCTCGG CTCGGGGCGC TGACGAAG
J23SpCA_8 NGCAGAAGA CGGGTTTGAT GGGGCGGGGA TGTAAGCGGG AAGGGAAACC GACCCGTTTA GTCTGCCGCT CCCAACAGCC CGAGTTTCCA GCCCCTTGAG TAAGGGGCAA GAAACTAAGC CCGCCTGGTG GATGGCTCGG CTCGGGGCGC TGACGAAN
J23SpCA_5 CAGAAGA CGGGTTTGAT GGGGCGGGGA TGTAAGCGGG AAGGGAAACC GACCCGTTTA GTCTGCCGCT CCCAACAGCC CGAGTTTCCA GCCCCTTGAG TAAGGGGCAA GAAACTAAGC CCGCCTGGTG GATGGCTCGG CTCGGGGCGC TGACGAAAG
J23SpCA_3 CAGAAGA CGGGTTTGAT GGGGCGGGGA TGTAAGCGGG AAGGGAAACC GACCCGTTTA GTCTGCCGCT CCCAACAGCC CGAGTTTCCA GCCCCTTGAG TAAGGGGCAA GAAACTAAGC CCGCCTGGTG GATGGCTCGG CTCGGGGCGC TGACGAAAGG
J23SpCA_3 CGGAAGA CGGGTTTGAT GGGGCGGGGA TGTAAGCGGG AAGGGAAACC GACCCGTTTA GTCTGCCGCT CCCAACAGCC CGAGTTTCCA GCCCCTTGAG TAAGGGGCAA GAAACTAAGC CCGCCTGGTG GATGGCTCGG CTCGGGGCGC TGACGAAGGG
J23SpCA_6 CAGAAGA CGGGTTTGAT GGGGCGGGGA TGTAAGCGGG AAGGGAAACC GACCCGTTTA GTCTGCCGCT CCCAACAGCC CGAGTTTCCA GCCCCTTGAG TAAGGGGCAA GAAACTAAGC CCGCCTGGTG GATGGCTCGG CTCGGGGCGC TGACGAGGG
J23SpCA_162 AGAAGA CGGGTTTGAT GGGGCGGGGA TGTAAGCGGG AAGGGAAACC GACCCGTTTA GTCTGCCGCT CCCAACAGCC CGAGTTTCCA GCCCCTTGAG TAAGGGGCAA GAAACTAAGC CCGCCTGGTG GATGGCTCGG CTCGGGGCGC TGACGAAG
J23SpCA_55 CGCAGAAGA CGGGTTTGAT GGGGCGGGGA TGTAAGCGGG AAGGGAAACC GACCCGTTTA GTCTGCCGCT CCCAACAGCC CGAGTTTCCA GCCCCTTGAG TAAGGGGCAA GAAACTAAGC CCGCCTGGTG GATGGCTCGG CTCGGGGCGC TGACGAA
J23SpCA_23 CCGCAGAAGA CGGGTTTGAT GGGGCGGGGA TGTAAGCGGG AAGGGAAACC GACCCGTTTA GTCTGCCGCT CCCAACAGCC CGAGTTTCCA GCCCCTTGAG TAAGGGGCAA GAAACTAAGC CCGCCTGGTG GATGGCTCGG CTCGGGGCGC TGACGAA
J23SpCA_2 N CCGCAGAAGA CGGGTTTGAT GGGGCGGGGA TGTAAGCGGG AAGGGAAACC GACCCGTTTA GTCTGCCGCT CCCAACAGCC CGAGTTTCCA GCCCCTTGAG TAAGGGGCAA GAAACTAAGC CCGCCTGGTG GATGGCTCGG CTCGGGGCGC TGACGAAN
```

```
J23SpCA_4      CGCAGAAGA CGGGTTTGAT GGGGCGGGGA TGTAAGCGGG AAGGGAAACC GACCCGTTTA GTCTGCCGCT CCCAACAGCC CGAGTTTCCA GCCCCTTGAG TAAGGGGCAA GAAACCTAAG CCGCCTGGTG GATGGCTCGG CTCGGGGCGC TGACGAAN
J23SpCA_6     NCGCAGAAGA CGGGTTTGAT GGGGCGGGGA TGTAAGCGGG AAGGGAAACC GACCCGTTTA GTCTGCCGCT CCCAACAGCC CGAGTTTCCA GCCCCTTGAG TAAGGGGCAA GAAACCTAAG CCGCCTGGTG GATGGCTCGG CTCGGGGCGC TGACGAAN
J23SpCA_5     CCGCAGAAGA CGGGTTTGAT GGGGCGGGGA TGTAAGCGGG AAGGGAAACC GACCCGTTTA GTCTGCCGCT CCCAACAGCC CGAGTTTCCA GCCCCTTGAG TAAGGGGCAA GAAACCTAAG CCGCCTGGTG GATGGCTCGG CTCGGGGCGC TGACGA
J23SpCA_26  C CCGCAGAAGA CGGGTTTGAT GGGGCGGGGA TGTAAGCGGG AAGGGAAACC GACCCGTTTA GTCTGCCGCT CCCAACAGCC CGAGTTTCCA GCCCCTTGAG TAAGGGGCAA GAAACCTAAG CCGCCTGGTG GATGGCTCGG CTCGGGGCGC TGACG
J23SpCA_24     CGCAGAAGA CGGGTTTGAT GGGGCGGGGA TGTAAGCGGG AAGGGAAACC GACCCGTTTA GTCTGCCGCT CCCAACAGCC CGAGTTTCCA GCCCCTTGAG TAAGGGGCAA GAAACCTAAG CCGCCTGGTG GATGGCTCGG CTCGGGGCGC TGACAAAG
J23SpCA_2       GCAGAAGA CGGGTTTGAT GGGGCGGGGA TGTAAGCGGG AAGGGAAACC GACCCGTTTA GTCTGCCGCT CCCAACAGCC CGAGTTTCCA GCCCCTTGAG TAAGGGGCAA GAAACCTAAG CCGCCTGGTG GATGGCTCGG CTCGGGGCGC TGACAAAG
J23SpCA_2      GGCAGAAGA CGGGTTTGAT GGGGCGGGGA TGTAAGCGGG AAGGGAAACC GACCCGTTTA GTCTGCCGCT CCCAACAGCC CGAGTTTCCA GCCCCTTGAG TAAGGGGCAA GAAACCTAAG CCGCCTGGTG GATGGCTCGG CTCGGGGCGC TGACAAAGG
J23SpCA_5      CGCAGAAGA CGGGTTTGAT GGGGCGGGGA TGTAAGCGGG AAGGGAAACC GACCCGTTTA GTCTGCCGCT CCCAACAGCC CGAGTTTCCA GCCCCTTGAG TAAGGGGCAA GAAACCTAAG CCGCCTGGTG GATGGCTCGG CTCGGGGCGC TGACGGAG
J23SpCA_2       GCAGAAGA CGGGTTTGAT GGGGCGGGGA TGTAAGCGGG AAGGGAAACC GACCCGTTTA GTCTGCCGCT CCCAACAGCC CGAGTTTCCA GCCCCTTGAG TAAGGGGCAA GAAACCTAAG CCGCCTGGTG GATGGCTCGG CTCGGGGCGC TGACGGAG
J23SpCA_10     CGCAGAAGA CGGGTTTGAT GGGGCGGGGA TGTAAGCGGG AAGGGAAACC GACCCGTTTA GTCTGCCGCT CCCAACAGCC CGAGTTTCCA GCCCCTTGAG TAAGGGGCAA GAAACCTAAG CCGCCTGGTG GATGGCTCGG CTCGGGGCGC TGACGA
J23SpCA_5      CGCAGAAGA TGGGTTTGAT GGGGCGGGGA TGTAAGCGGG AAGGGAAACC GACCCGTTTA GTCTGCCGCT CCCAACAGCC CGAGTTTCCA GCCCCTTGAG TAAGGGGCAA GAAACCTAAG CCGCCTGGTG GATGGCTCGG CTCGGGGCGC TGACGAAG
J23SpCA_6       GCAGAAGA CGGGTTTGAT GGGGCGGGGA TGTAAGCGGG AAGGGAAACC GACCCGTTTA GTCTGCCGCT CCCAACAGCC CGAGTTTCCA GCCCCTTGAG TAAGGGGCAA GAAACCTAAG CCGCCTGGTG GATGGCTCGG CTCGGGGCGC TGACGAA
J23SpCA_6      CGCAGAAGA CGGGTTTGAT GGGGCGGGGA TGTAAGCGGG AAGGGAAACC GACCCGTTTA GTCTGCCGCT CCCAACAGCC CGAGTTTCCA GCCCCTTGAG TAAGGGGCAA GAAACCTAAG CCGCCTGGTG GATGGCTCGG CTCGGGGCGC TGACGAGG
J23SpCA_5      CGCAGAAGA CGGGTTTGAT GGGGCGGGGA TGTAAGCGGG AAGGGAAACC GACCCGTTTA GTCTGCCGCT CCCAACAGCC CGAGTTTCCA GCCCCTTGAG TAAGGGGCAA GAAACCTAAG CCGCCTGGTG GATGGCTCGG CTCGGGGCGC TGACGAG
J23SpCA_13       CAGAAGA CGGGTTTGAT GGGGCGGGGA TGTAAGCGGG AAGGGAAACC GACCCGTTTA GTCTGCCGCT CCCAACAGCC CGAGTTTCCA GCCCCTTGAG TAAGGGGCAA GAAACCTAAG CCGCCTGGTG GATGGCTCGG CTCGGGGCGC TGACGAA
J23SpCA_10       CAGAAGA CGGGTTTGAT GGGGCGGGGA TGTAAGCGGG AAGGGAAACC GACCCGTTTA GTCTGCCGCT CCCAACAGCC CGAGTTTCCA GCCCCTTGAG TAAGGGGCAA GAAACCTAAG CCGCCTGGTG GATGGCTCGG CTCGGGGCGC TGACGAAGAG
J23SpCA_4        CAGAAGA CGGGTTTGAT GGGGCGGGGA TGTAAGCGGG AAGGGAAACC GACCCGTTTA GTCTGCCGCT CCCAACAGCC CGAGTTTCCA GCCCCTTGAG TAAGGGGCAA GAAACCTAAG CCGCCTGGTG GATGGCTCGG CTCGGGGCGC TGACGAAGA
J23SpCA_4          GAAGA CGGGTTTGAT GGGGCGGGGA TGTAAGCGGG AAGGGAAACC GACCCGTTTA GTCTGCCGCT CCCAACAGCC CGAGTTTCCA GCCCCTTGAG TAAGGGGCAA GAAACCTAAG CCGCCTGGTG GATGGCTCGG CTCGGGGCGC TGACGAAGAG
J23SpCA_8        CAGAAGA CGGGTTTGAT GGGGCGGGGA TGTAAGCGGG AAGGGAAACC GACCCGTTTA GTCTGCCGCT CCCAACAGCC CGAGTTTCCA GCCCCTTGAG TAAGGGGCAA GAAACCTAAG CCGCCTGGTG GATGGCTCGG CTCGGGGCGC TGACAAAG
J23SpCA_216           GA CGGGTTTGAT GGGGCGGGGA TGTAAGCGGG AAGGGAAACC GACCCGTTTA GTCTGCCGCT CCCAACAGCC CGAGTTTCCA GCCCCTTGAG TAAGGGGCAA GAAACCTAAG CCGCCTGGTG GATGGCTCGG CTCGGGGCGC TGACGAAGGG
J23SpCA_107           GA CGGGTTTGAT GGGGCGGGGA TGTAAGCGGG AAGGGAAACC GACCCGTTTA GTCTGCCGCT CCCAACAGCC CGAGTTTCCA GCCCCTTGAG TAAGGGGCAA GAAACCTAAG CCGCCTGGTG GATGGCTCGG CTCGGGGCGC TGACGAAGGG C
J23SpCA_60            GA CGGGTTTGAT GGGGCGGGGA TGTAAGCGGG AAGGGAAACC GACCCGTTTA GTCTGCCGCT CCCAACAGCC CGAGTTTCCA GCCCCTTGAG TAAGGGGCAA GAAACCTAAG CCGCCTGGTG GATGGCTCGG CTCGGGGCGC TGACGAAGGG CG
J23SpCA_2             GA CGGGTTTGAT GGGGCGGGGA TGTAAGCGGG AAGGGAAACC GACCCGTTTA GTCTGCCGCT CCCAACAGCC CGAGTTTCCA GCCCCTTGAG TAAGGGGCAA GAAACCTAAG CCGCCTGGTG GATGGCTCGG CTCGGGGCGC TGACGAAGGG CGGG
J23SpCA_2             GA CGGGTTTGAT GGGGCGGGGA TGTAAGCGGG AAGGGAAACC GACCCGTTTA GTCTGCCGCT CCCAACAGCC CGAGTTTCCA GCCCCTTGAG TAAGGGGCAA GAAACCTAAG CCGCCTGGTG GATGGCTCGG CTCGGGGCGC TGACGAAGGG CT
J23SpCA_37           AGA CGGGTTTGAT GGGGCGGGGA TGTAAGCGGG AAGGGAAACC GACCCGTTTA GTCTGCCGCT CCCAACAGCC CGAGTTTCCA GCCCCTTGAG TAAGGGGCAA GAAACCTAAG CCGCCTGGTG GATGGCTCGG CTCGGGGCGC TGACGAAGGG C
J23SpCA_28             A CGGGTTTGAT GGGGCGGGGA TGTAAGCGGG AAGGGAAACC GACCCGTTTA GTCTGCCGCT CCCAACAGCC CGAGTTTCCA GCCCCTTGAG TAAGGGGCAA GAAACCTAAG CCGCCTGGTG GATGGCTCGG CTCGGGGCGC TGACGAAGGG CG
J23SpCA_2             GA CGGGTTTGAT GGGGCGGGGA TGTAAGCGGG AAGGGAAACC GACCCGTTTA GTCTGCCGCT CCCAACAGCC CGAGTTTCCA GCCCCTTGAG TAAGGGGCAA GAAACCTAAG CCGCCTGGTG GATGGCTCGG CTCGGGGCGC TGACGAAGGG CC
J23SpCA_118              CGGGTTTGAT GGGGCGGGGA TGTAAGCGGG AAGGGAAACC GACCCGTTTA GTCTGCCGCT CCCAACAGCC CGAGTTTCCA GCCCCTTGAG TAAGGGGCAA GAAACCTAAG CCGCCTGGTG GATGGCTCGG CTCGGGGCGC TGACGAAGGG CG
J23SpCA_107              CGGGTTTGAT GGGGCGGGGA TGTAAGCGGG AAGGGAAACC GACCCGTTTA GTCTGCCGCT CCCAACAGCC CGAGTTTCCA GCCCCTTGAG TAAGGGGCAA GAAACCTAAG CCGCCTGGTG GATGGCTCGG CTCGGGGCGC TGACGAAGGG CGTGG
J23SpCA_70               CGGGTTTGAT GGGGCGGGGA TGTAAGCGGG AAGGGAAACC GACCCGTTTA GTCTGCCGCT CCCAACAGCC CGAGTTTCCA GCCCCTTGAG TAAGGGGCAA GAAACCTAAG CCGCCTGGTG GATGGCTCGG CTCGGGGCGC TGACGAAGGG CGTG
J23SpCA_21             A CGGGTTTGAT GGGGCGGGGA TGTAAGCGGG AAGGGAAACC GACCCGTTTA GTCTGCCGCT CCCAACAGCC CGAGTTTCCA GCCCCTTGAG TAAGGGGCAA GAAACCTAAG CCGCCTGGTG GATGGCTCGG CTCGGGGCGC TGACGAAGGG CGTG
J23SpCA_7                 GGGTTTGAT GGGGCGGGGA TGTAAGCGGG AAGGGAAACC GACCCGTTTA GTCTGCCGCT CCCAACAGCC CGAGTTTCCA GCCCCTTGAG TAAGGGGCAA GAAACCTAAG CCGCCTGGTG GATGGCTCGG CTCGGGGCGC TGACGAAGGG CGTGG
J23SpCA_4                  GGTTTGAT GGGGCGGGGA TGTAAGCGGG AAGGGAAACC GACCCGTTTA GTCTGCCGCT CCCAACAGCC CGAGTTTCCA GCCCCTTGAG TAAGGGGCAA GAAACCTAAG CCGCCTGGTG GATGGCTCGG CTCGGGGCGC TGACGAAGGG CGTGGC
J23SpCA_3                CGGGTTTGAT GGGGCGGGGA TGTAAGCGGG AAGGGAAACC GACCCGTTTA GTCTGCCGCT CCCAACAGCC CGAGTTTCCA GCCCCTTGAG TAAGGGGCAA GAAACCTAAG CCGCCTGGTG GATGGCTCGG CTCGGGGCGC TGACGAAGGG CGT
J23SpCA_2                 GGGTTTGAT GGGGCGGGGA TGTAAGCGGG AAGGGAAACC GACCCGTTTA GTCTGCCGCT CCCAACAGCC CGAGTTTCCA GCCCCTTGAG TAAGGGGCAA GAAACCTAAG CCGCCTGGTG GATGGCTCGG CTCGGGGCGC TGACGAAGGG CGTG
J23SpCA_7                  GGTTTGAT GGGGCGGGGA TGTAAGCGGG AAGGGAAACC GACCCGTTTA GTCTGCCGCT CCCAACAGCC CGAGTTTCCA GCCCCTTGAG TAAGGGGCAA GAAACCTAAG CCGCCTGGTG GATGGCTCGG CTCGGGGCGC TGACGAAGGG CGTGG
J23SpCA_58             A CGGGTTTGAT GGGGCGGGGA TGTAAGCGGG AAGGGAAACC GACCCGTTTA GTCTGCCGCT CCCAACAGCC CGAGTTTCCA GCCCCTTGAG TAAGGGGCAA GAAACCTAAG CCGCCTGGTG GATGGCTCGG CTCGGGGCGC TGACGAAGGG C
J23SpCA_71           AGA CGGGTTTGAT GGGGCGGGGA TGTAAGCGGG AAGGGAAACC GACCCGTTTA GTCTGCCGCT CCCAACAGCC CGAGTTTCCA GCCCCTTGAG TAAGGGGCAA GAAACCTAAG CCGCCTGGTG GATGGCTCGG CTCGGGGCGC TGACGAAGGG
J23SpCA_61          AAGA CGGGTTTGAT GGGGCGGGGA TGTAAGCGGG AAGGGAAACC GACCCGTTTA GTCTGCCGCT CCCAACAGCC CGAGTTTCCA GCCCCTTGAG TAAGGGGCAA GAAACCTAAG CCGCCTGGTG GATGGCTCGG CTCGGGGCGC TGACGAAGGG
J23SpCA_23          AAGA CGGGTTTGAT GGGGCGGGGA TGTAAGCGGG AAGGGAAACC GACCCGTTTA GTCTGCCGCT CCCAACAGCC CGAGTTTCCA GCCCCTTGAG TAAGGGGCAA GAAACCTAAG CCGCCTGGTG GATGGCTCGG CTCGGGGCGC TGACGAAGG
J23SpCA_56           AGA CGGGTTTGAT GGGGCGGGGA TGTAAGCGGG AAGGGAAACC GACCCGTTTA GTCTGCCGCT CCCAACAGCC CGAGTTTCCA GCCCCTTGAG TAAGGGGCAA GAAACCTAAG CCGCCTGGTG GATGGCTCGG CTCGGGGCGC TGACGAAGG
J23SpCA_36          AAGA CGGGTTTGAT GGGGCGGGGA TGTAAGCGGG AAGGGAAACC GACCCGTTTA GTCTGCCGCT CCCAACAGCC CGAGTTTCCA GCCCCTTGAG TAAGGGGCAA GAAACCTAAG CCGCCTGGTG GATGGCTCGG CTCGGGGCGC TGACGAAG
J23SpCA_26         GAAGA CGGGTTTGAT GGGGCGGGGA TGTAAGCGGG AAGGGAAACC GACCCGTTTA GTCTGCCGCT CCCAACAGCC CGAGTTTCCA GCCCCTTGAG TAAGGGGCAA GAAACCTAAG CCGCCTGGTG GATGGCTCGG CTCGGGGCGC TGACGAAG
J23SpCA_24         GAAGA CGGGTTTGAT GGGGCGGGGA TGTAAGCGGG AAGGGAAACC GACCCGTTTA GTCTGCCGCT CCCAACAGCC CGAGTTTCCA GCCCCTTGAG TAAGGGGCAA GAAACCTAAG CCGCCTGGTG GATGGCTCGG CTCGGGGCGC TGACGAAGG
J23SpCA_4         AGAAGA CGGGTTTGAT GGGGCGGGGA TGTAAGCGGG AAGGGAAACC GACCCGTTTA GTCTGCCGCT CCCAACAGCC CGAGTTTCCA GCCCCTTGAG TAAGGGGCAA GAAACCTAAG CCGCCTGGTG GATGGCTCGG CTCGGGGCGC TGACGAGG
J23SpCA_3         AGAAGA CGGGTTTGAT GGGGCGGGGA TGTAAGCGGG AAGGGAAACC GACCCGTTTA GTCTGCCGCT CCCAACAGCC CGAGTTTCCA GCCCCTTGAG TAAGGGGCAA GAAACCTAAG CCGCCTGGTG GATGGCTCGG CTCGGGGCGC TGACGAGGGG
J23SpCA_62     CGCAGAAGA CGGGTTTGAT GGGGCGGGGA TGTAAGCGGG AAGGGAAACC GACCCGTTTA GTCTGCCGCT CCCAACAGCC CGAGTTTCCA GCCCCTTGAG TAAGGGGCAA GAAACCTAAG CCGCCTGGTG GATGGCTCGG CTCGGGGCGC TGACG
J23SpCA_20    CCGCAGAAGA CGGGTTTGAT GGGGCGGGGA TGTAAGCGGG AAGGGAAACC GACCCGTTTA GTCTGCCGCT CCCAACAGCC CGAGTTTCCA GCCCCTTGAG TAAGGGGCAA GAAACCTAAG CCGCCTGGTG GATGGCTCGG CTCGGGGCGC TGACG
J23SpCA_21  C CCGCAGAAGA CGGGTTTGAT GGGGCGGGGA TGTAAGCGGG AAGGGAAACC GACCCGTTTA GTCTGCCGCT CCCAACAGCC CGAGTTTCCA GCCCCTTGAG TAAGGGGCAA GAAACCTAAG CCGCCTGGTG GATGGCTCGG CTCGGGGCGC TGAC
J23SpCA_15    CCGCAGAAGA CGGGTTTGAT GGGGCGGGGA TGTAAGCGGG AAGGGAAACC GACCCGTTTA GTCTGCCGCT CCCAACAGCC CGAGTTTCCA GCCCCTTGAG TAAGGGGCAA GAAACCTAAG CCGCCTGGTG GATGGCTCGG CTCGGGGCGC TGAC
J23SpCA_37     CGCAGAAGA CGGGTTTGAT GGGGCGGGGA TGTAAGCGGG AAGGGAAACC GACCCGTTTA GTCTGCCGCT CCCAACAGCC CGAGTTTCCA GCCCCTTGAG TAAGGGGCAA GAAACCTAAG CCGCCTGGTG GATGGCTCGG CTCGGGGCGC TGAC
J23SpCA_7       GCAGAAGA CGGGTTTGAT GGGGCGGGGA TGTAAGCGGG AAGGGAAACC GACCCGTTTA GTCTGCCGCT CCCAACAGCC CGAGTTTCCA GCCCCTTGAG TAAGGGGCAA GAAACCTAAG CCGCCTGGTG GATGGCTCGG CTCGGGGCGC TGACG
J23SpCA_21       CAGAAGA CGGGTTTGAT GGGGCGGGGA TGTAAGCGGG AAGGGAAACC GACCCGTTTA GTCTGCCGCT CCCAACAGCC CGAGTTTCCA GCCCCTTGAG TAAGGGGCAA GAAACCTAAG CCGCCTGGTG GATGGCTCGG CTCGGGGCGC TGACG
J23SpCA_3       GCAGAAGA CGGGTTTGAT GGGGCGGGGA TGTAAGCGGG AAGGGAAACC GACCCGTTTA GTCTGCCGCT CCCAACAGCC CGAGTTTCCA GCCCCTTGAG TAAGGGGCAA GAAACCTAAG CCGCCTGGTG GATGGCTCGG CTCGGGGCGC TGAC
J23SpCA_5     CCGCAGAAGA CGGGTTTGAT GGGGCGGGGA TGTAAGCGGG AAGGGAAACC GACCCGTTTA GTCTGCCGCT CCCAACAGCC CGAGTTTCCA GCCCCTTGAG TAAGGGGCAA GAAACCTAAG CCGCCTGGTG GATGGCTCGG CTCGGGGCGC TG
J23SpCA_3        CAGAAGA CGGGTTTGAT GGGGCGGGGA TGTAAGCGGG AAGGGAAACC GACCCGTTTA GTCTGCCGCT CCCAACAGCC CGAGTTTCCA GCCCCTTGAG TAAGGGGCAA GAAACCTAAG CCGCCTGGTG GATGGCTCGG CTCGGGGCGC TGACGAG
J23SpCA_2      NGCAGAAGA CGGGTTTGAT GGGGCGGGGA TGTAAGCGGG AAGGGAAACC GACCCGTTTA GTCTGCCGCT CCCAACAGCC CGAGTTTCCA GCCCCTTGAG TAAGGGGCAA GAAACCTAAG CCGCCTGGTG GATGGCTCGG CTCGGGGCGC TGAN
J23SpCA_19       CAGAAGA CGGGTTTGAT GGGGCGGGGA TGTAAGCGGG AAGGGAAACC GACCCGTTTA GTCTGCCGCT CCCAACAGCC CGAGTTTCCA GCCCCTTGAG TAAGGGGCAA GAAACCTAAG CCGCCTGGTG GATGGCTCGG CTCGGGGCGC TGAC
J23SpCA_14        AGAAGA CGGGTTTGAT GGGGCGGGGA TGTAAGCGGG AAGGGAAACC GACCCGTTTA GTCTGCCGCT CCCAACAGCC CGAGTTTCCA GCCCCTTGAG TAAGGGGCAA GAAACCTAAG CCGCCTGGTG GATGGCTCGG CTCGGGGCGC TGACGAA
J23SpCA_3         NGAAGA CGGGTTTGAT GGGGCGGGGA TGTAAGCGGG AAGGGAAACC GACCCGTTTA GTCTGCCGCT CCCAACAGCC CGAGTTTCCA GCCCCTTGAG TAAGGGGCAA GAAACCTAAG CCGCCTGGTG GATGGCTCGG CTCGGGGCGC TGACGAAN
J23SpCA_8         AGAAGA CGGGTTTGAT GGGGCGGGGA TGTAAGCGGG AAGGGAAACC GACCCGTTTA GTCTGCCGCT CCCAACAGCC CGAGTTTCCA GCCCCTTGAG TAAGGGGCAA GAAACCTAAG CCGCCTGGTG GATGGCTCGG CTCGGGGCGC TGACAAAG
J23SpCA_8         AGAAGA CGGGTTTGAT GGGGCGGGGA TGTAAGCGGG AAGGGAAACC GACCCGTTTA GTCTGCCGCT CCCAACAGCC CGAGTTTCCA GCCCCTTGAG TAAGGGGCAA GAAACCTAAG CCGCCTGGTG GATGGCTCGG CTCGGGGCGC TGACG
J23SpCA_4        CAGAAGA CGGGTTTGAT GGGGCGGGGA TGTAAGCGGG AAGGGAA-CC GACCCGTTTA GTCTGCCGCT CCCAACAGCC CGAGTTTCCA GCCCCTTGAG TAAGGGGCAA GAAACCTAAG CCGCCTGGTG GATGGCTCGG CTCGGGGCGC TGACGAAGGG C
J23SpCA_341              CGGGTTTGAT GGGGCGGGGA TGTAAGCGGG AAGGGAAACC GACCCGTTTA GTCTGCCGCT CCCAACAGCC CGAGTTTCCA GCCCCTTGAG TAAGGGGCAA GAAACCTAAG CCGCCTGGTG GATGGCTCGG CTCGGGGCGC TGACGAAGGG
J23SpCA_181              CGGGTTTGAT GGGGCGGGGA TGTAAGCGGG AAGGGAAACC GACCCGTTTA GTCTGCCGCT CCCAACAGCC CGAGTTTCCA GCCCCTTGAG TAAGGGGCAA GAAACCTAAG CCGCCTGGTG GATGGCTCGG CTCGGGGCGC TGACGAAGGG C
J23SpCA_5                CGGGTTTGAT GGGGCGGGGA TGTAAGCGGG AAGGGAAACC GACCCGTTTA GTCTGCCGCT CCCAACAGCC CGAGTTTCCA GCCCCTTGAG TAAGGGGCAA GAAACCTAAG CCGCCTGGTG GATGGCTCGG CTCGGGGCGC TGACGAAGGG CA
J23SpCA_4                CGGGTTTGAT GGGGCGGGGA TGTAAGCGGG AAGGGAAACC GACCCGTTTA GTCTGCCGCT CCCAACAGCC CGAGTTTCCA GCCCCTTGAG TAAGGGGCAA GAAACCTAAG CCGCCTGGTG GATGGCTCGG CTCGGGGCGC TGACGAAGGG CC
J23SpCA_156           GA CGGGTTTGAT GGGGCGGGGA TGTAAGCGGG AAGGGAAACC GACCCGTTTA GTCTGCCGCT CCCAACAGCC CGAGTTTCCA GCCCCTTGAG TAAGGGGCAA GAAACCTAAG CCGCCTGGTG GATGGCTCGG CTCGGGGCGC TGACGAAGG
J23SpCA_11            GA CGGGTTTGAT GGGGCGGGGA TGTAAGCGGG AAGGGAAACC GACCCGTTTA GTCTGCCGCT CCCAACAGCC CGAGTTTCCA GCCCCTTGAG TAAGGGGCAA GAAACCTAAG CCGCCTGGTG GATGGCTCGG CTCGGGGCGC TGACGAAGGG G
J23SpCA_2             GA CGGGTTTGAT GGGGCGGGGA TGTAAGCGGG AAGGGAAACC GACCCGTTTA GTCTGCCGCT CCCAACAGCC CGAGTTTCCA GCCCCTTGAG TAAGGGGCAA GAAACCTAAG CCGCCTGGTG GATGGCTCGG CTCGGGGCGC TGACGAAGGG A
J23SpCA_4             GA CGGGTTTGAT GGGGCGGGGA TGTAAGCGGG AAGGGAAACC GACCCGTTTA GTCTGCCGCT CCCAACAGCC CGAGTTTCCA GCCCCTTGAG TAAGGGGCAA GAAACCTAAG CCGCCTGGTG GATGGCTCGG CTCGGGGCGC TGACAAAGGG
J23SpCA_3             GA CGGGTTTGAT GGGGCGGGGA TGTAAGCGGG AAGGGAAACC GACCCGTTTA GTCTGCCGCT CCCAACAGCC CGAGTTTCCA GCCCCTTGAG TAAGGGGCAA GAAACCTAAG CCGCCTGGTG GATGGCTCGG CTCGGGGCGC TGACAAAGG
J23SpCA_81             A CGGGTTTGAT GGGGCGGGGA TGTAAGCGGG AAGGGAAACC GACCCGTTTA GTCTGCCGCT CCCAACAGCC CGAGTTTCCA GCCCCTTGAG TAAGGGGCAA GAAACCTAAG CCGCCTGGTG GATGGCTCGG CTCGGGGCGC TGACGAAGGG
J23SpCA_2             GA CGGGTTTGAT GGGGCGGGGA TGTAAGCGGG AAGGGAAACC GCCCCGTTTA GTCTGCCGCT CCCAACAGCC CGAGTTTCCA GCCCCTTGAG TAAGGGGCAA GAAACCTAAG CCGCCTGGTG GATGGCTCGG CTCGGGGCGC TGACGAAGGG
J23SpCA_8             GA CGGGTTTGAT GGGGCGGGGA TGTAAGCGGG AAGGGAAACC GACCCGTTTA GTCTGCCGCT CCCAACAGCC CGAGTTTCCA GCCCCTTGAG TAAGGGGCAA GAAACCTAAG CCGCCTGGTG GATGGCTCGG CTCGGGGCGC TGACGAAGAG
J23SpCA_6                CGGGTTTGAT GGGGCGGGGA TGTAAGCGGG AAGGGAAACC GACCCGTTTA GTCTGCCGCT CCCAACAGCC CGAGTTTCCA GCCCCTTGAG TAAGGGGCAA GAAACCTAAG CCGCCTGGTG GATGGCTCGG CTCGGGGCGC TGACGAAGGG G
J23SpCA_67             A CGGGTTTGAT GGGGCGGGGA TGTAAGCGGG AAGGGAAACC GACCCGTTTA GTCTGCCGCT CCCAACAGCC CGAGTTTCCA GCCCCTTGAG TAAGGGGCAA GAAACCTAAG CCGCCTGGTG GATGGCTCGG CTCGGGGCGC TGACGAAGG
J23SpCA_2             NA CGGGTTTGAT GGGGCGGGGA TGTAAGCGGG AAGGGAAACC GACCCGTTTA GTCTGCCGCT CCCAACAGCC CGAGTTTCCA GCCCCTTGAG TAAGGGGCAA GAAACCTAAG CCGCCTGGTG GATGGCTCGG CTCGGGGCGC TGACGAAGGN
J23SpCA_3              A CGGGTTTGAT GGGGCGGGGA TGTAAGCGGG AAGGGAAACC GACCCGTTTA GTCTGCCGCT CCCAACAGCC CGAGTTTCCA GCCCCTTGAG TAAGGGGCAA GAAACCTAAG CCGCCTGGTG GATGGCTCGG CTCGGGGCGC TGACGAAGAG
J23SpCA_3                CGGGTTTGAT GGGGCGGGGA TGTAAGCGGG AAGGGAAACC GACCCGTTTA GTCTGCCGCT CCCAACAGCC CGAGTTTCCA GCCCCTTGAG TAAGGGGCAA GAAACCTAAG CCGCCTGGTG GATGGCTCGG CTCGGGGCGC TGACGAAGGG T
J23SpCA_201           GA CGGGTTTGAT GGGGCGGGGA TGTAAGCGGG AAGGGAAACC GACCCGTTTA GTCTGCCGCT CCCAACAGCC CGAGTTTCCA GCCCCTTGAG TAAGGGGCAA GAAACCTAAG CCGCCTGGTG GATGGCTCGG CTCGGGGCGC TGACGAAG
J23SpCA_64           AGA CGGGTTTGAT GGGGCGGGGA TGTAAGCGGG AAGGGAAACC GACCCGTTTA GTCTGCCGCT CCCAACAGCC CGAGTTTCCA GCCCCTTGAG TAAGGGGCAA GAAACCTAAG CCGCCTGGTG GATGGCTCGG CTCGGGGCGC TGACGAAG
J23SpCA_3            AGA CGGGTTTGAT GGGGCGGGGA TGTAAGCGGG AAGGGAAACC GACCCGTTTA GTCTGCCGCT CCCAACAGCC CGAGTTTCCA GCCCCTTGAG TAAGGGGCAA GAAACCTAAG CCGCCTGGTG GATGGCTCGG CTCGGGGCGC TGACGAA
J23SpCA_3             GA CGGGTTTGAT GGGGCGGGGA TGTAAGCGGG AAGGGAAACC GACCCGTTTA GTCTGCCGCC CCCAACAGCC CGAGTTTCCA GCCCCTTGAG TAAGGGGCAA GAAACCTAAG CCGCCTGGTG GATGGCTCGG CTCGGGGCGC TGACGAAGG
J23SpCA_74             A CGGGTTTGAT GGGGCGGGGA TGTAAGCGGG AAGGGAAACC GACCCGTTTA GTCTGCCGCT CCCAACAGCC CGAGTTTCCA GCCCCTTGAG TAAGGGGCAA GAAACCTAAG CCGCCTGGTG GATGGCTCGG CTCGGGGCGC TGACGAAG
J23SpCA_2             GA CGGGTTTGAT GGGGCGGGGA TGTAAGCGGG AAGGGAAACC GACCCGTTTA GTCTGCCGCT CCCAACAGCC CGAGTTTCCA GCCCCTTGAG TAAGGGGCAA GAAACCTAAG CCGCCTGGTG GATGGCTCGG CTCGGGGCGC TGATGAAGG
J23SpCA_15            GA CGGGTTTGAT GGGGCGGGGA TGTAAGCGGG AAGGGAAACC GACCCGTTTA GTCTGCCGCT CCCAACAGCC CGAGTTTCCA GCCCCTTGAG TAAGGGGCAA GAAACCTAAG CCGCCTGGTG GATGGCTCGG CTCGGGGCGC TGACGAA
J23SpCA_2             GA CGGGTTTGAT GGGGCGGGGA TGTAAGCGGG AAGGGAAACC GACCCGTTTA GTCTGCCGCT CCCAACAGCC CGAGTTTCCA GCCCCTTGAG TAAGGGGCAA GAAACCTAAG CCGCCTGGTG GATGGCTCGG CTCGGGGCGC TGACGTAGG
J23SpCA_2             GA CGGGTTTGAT GGGGCGGGGA TGTAAGCGGG AAGGGAAACC GACCCGTTTA GTCTGCCGCT CCCAACAGCC CGAGTTTCCA GCCCCTTGAG TAAGGGGCAA GAAACCTAAG CCGCCTGGTG GATGGCTCGG CTCGGGGCGC TGACGAAAG
J23SpCA_8             GA CGGGTTTGAT GGGGCGGGGA TGTAAGCGGG AAGGGAAACC GACCCGTTTA GTCTGCCGCT CCCAACAGCC CGAGTTTCCA GCCCCTTGAG TAAGGGGCAA GAAACCTAAG CCGCCTGGTG GATGGCTCGG CTCGGGGCGC TGACAAAG
J23SpCA_33                  GTTTGAT GGGGCGGGGA TGTAAGCGGG AAGGGAAACC GACCCGTTTA GTCTGCCGCT CCCAACAGCC CGAGTTTCCA GCCCCTTGAG TAAGGGGCAA GAAACCTAAG CCGCCTGGTG GATGGCTCGG CTCGGGGCGC TGACGAAGGG CG
J23SpCA_24                GGGTTTGAT GGGGCGGGGA TGTAAGCGGG AAGGGAAACC GACCCGTTTA GTCTGCCGCT CCCAACAGCC CGAGTTTCCA GCCCCTTGAG TAAGGGGCAA GAAACCTAAG CCGCCTGGTG GATGGCTCGG CTCGGGGCGC TGACGAAGGG C
J23SpCA_11                GGGTTTGAT GGGGCGGGGA TGTAAGCGGG AAGGGAAACC GACCCGTTTA GTCTGCCGCT CCCAACAGCC CGAGTTTCCA GCCCCTTGAG TAAGGGGCAA GAAACCTAAG CCGCCTGGTG GATGGCTCGG CTCGGGGCGC TGACGAAGGG CG
J23SpCA_4                  GGTTTGAT GGGGCGGGGA TGTAAGCGGG AAGGGAAACC GACCCGTTTA GTCTGCCGCT CCCAACAGCC CGAGTTTCCA GCCCCTTGAG TAAGGGGCAA GAAACCTAAG CCGCCTGGTG GATGGCTCGG CTCGGGGCGC TGACGAAGGG CG
J23SpCA_2                NGGGTTTGAT GGGGCGGGGA TGTAAGCGGG AAGGGAAACC GACCCGTTTA GTCTGCCGCT CCCAACAGCC CGAGTTTCCA GCCCCTTGAG TAAGGGGCAA GAAACCTAAG CCGCCTGGTG GATGGCTCGG CTCGGGGCGC TGACGAAGGG CN
J23SpCA_23                GGGTTTGAT GGGGCGGGGA TGTAAGCGGG AAGGGAAACC GACCCGTTTA GTCTGCCGCT CCCAACAGCC CGAGTTTCCA GCCCCTTGAG TAAGGGGCAA GAAACCTAAG CCGCCTGGTG GATGGCTCGG CTCGGGGCGC TGACGAAGGG
J23SpCA_19                  GTTTGAT GGGGCGGGGA TGTAAGCGGG AAGGGAAACC GACCCGTTTA GTCTGCCGCT CCCAACAGCC CGAGTTTCCA GCCCCTTGAG TAAGGGGCAA GAAACCTAAG CCGCCTGGTG GATGGCTCGG CTCGGGGCGC TGACGAAGGG CGTGG
J23SpCA_6                   GTTTGAT GGGGCGGGGA TGTAAGCGGG AAGGGAAACC GACCCGTTTA GTCTGCCGCT CCCAACAGCC CGAGTTTCCA GCCCCTTGAG TAAGGGGCAA GAAACCTAAG CCGCCTGGTG GATGGCTCGG CTCGGGGCGC TGACGAAGGG CGTGGC
J23SpCA_5                   GTTTGAT GGGGCGGGGA TGTAAGCGGG AAGGGAAACC GACCCGTTTA GTCTGCCGCT CCCAACAGCC CGAGTTTCCA GCCCCTTGAG TAAGGGGCAA GAAACCTAAG CCGCCTGGTG GATGGCTCGG CTCGGGGCGC TGACGAAGGG CGTG
J23SpCA_2                  GGTTTGAT GGGGCGGGGA TGTAAGCGGG AAGGGAAACC GACCCGTTTA GTCTGCCGCT CCCAACAGCC CGAGTTTCCA GCCCCTTGAG TAAGGGGCAA GAAACCTAAG CCGCCTGGTG GATGGCTCGG CTCGGGGCGC TGACGAAGGG CGTG
J23SpCA_2                   GTTTGAT GGGGCGGGGA TGTAAGCGGG AAGGGAAACC GACCCGTTTA GTCTGCCGCT CCCAACAGCC CGAGTTTCCA GCCCCTTGAG TAAGGGGCAA GAAACCTAAG CCGCCTGGTG GATGGCTCGG CTCGGGGCGC TGACGAAGGG CGGGG
J23SpCA_4                      TGAT GGGGCGGGGA TGTAAGCGGG AAGGGAAACC GACCCGTTTA GTCTGCCGCT CCCAACAGCC CGAGTTTCCA GCCCCTTGAG TAAGGGGCAA GAAACCTAAG CCGCCTGGTG GATGGCTCGG CTCGGGGCGC TGACGAAGGG CGTGG
J23SpCA_11                 GGTTTGAT GGGGCGGGGA TGTAAGCGGG AAGGGAAACC GACCCGTTTA GTCTGCCGCT CCCAACAGCC CGAGTTTCCA GCCCCTTGAG TAAGGGGCAA GAAACCTAAG CCGCCTGGTG GATGGCTCGG CTCGGGGCGC TGACGAAGGG C
J23SpCA_4      CGCAGAAGA CGGGTTTGAT GGGGCGGGGA TGTAAGCGGG AAGGGAAACC GACCCGTTTA GTCTGCCGCT CCCAACAGCC CGAGTTTCCA GCCCCTTGAG TAAGGGGCAA GAAACCTAAG CCGCCTGGTG GATGGCTCGG CTCGGGGCGC TG
J23SpCA_3       GCAGAAGA CGGGTTTGAT GGGGCGGGGA TGTAAGCGGG AAGGGAAACC GACCCGTTTA GTCTGCCGCT CCCAACAGCC CGAGTTTCCA GCCCCTTGAG TAAGGGGCAA GAAACCTAAG CCGCCTGGTG GATGGCTCGG CTCGGGGCGC TG
J23SpCA_263              CGGGTTTGAT GGGGCGGGGA TGTAAGCGGG AAGGGAAACC GACCCGTTTA GTCTGCCGCT CCCAACAGCC CGAGTTTCCA GCCCCTTGAG TAAGGGGCAA GAAACCTAAG CCGCCTGGTG GATGGCTCGG CTCGGGGCGC TGACGAAGG
J23SpCA_9                CGGGTTTGAT GGGGCGGGGA TGTAAGCGGG AAGGGAAACC GACCCGTTTA GTCTGCCGCT CCCAACAGCC CGAGTTTCCA GCCCCTTGAG TAAGGGGCAA GAAACCTAAG CCGCCTGGTG GATGGCTCGG CTCGGGGCGC TGACGAAGGA C
J23SpCA_257              CGGGTTTGAT GGGGCGGGGA TGTAAGCGGG AAGGGAAACC GACCCGTTTA GTCTGCCGCT CCCAACAGCC CGAGTTTCCA GCCCCTTGAG TAAGGGGCAA GAAACCTAAG CCGCCTGGTG GATGGCTCGG CTCGGGGCGC TGACGAAG
J23SpCA_3                CGGGTTTGAT GGGGCGGGGA TGTAAGCGGG AGGGGAAACC GACCCGTTTA GTCTGCCGCT CCCAACAGCC CGAGTTTCCA GCCCCTTGAG TAAGGGGCAA GAAACCTAAG CCGCCTGGTG GATGGCTCGG CTCGGGGCGC TGACGAAGGG
J23SpCA_5                CGGGTTTGAT GGGGCGGGGA TGTAAGCGGG AAGGGAAACC GACCCGTTTA GTCTGCCGCT CCCAACAGCC CGAGTTTCCA GCCCCTTGAG TAAGGGGCAA GAAACCTAAG CCGCCTGGTG GATGGCTCGG CTCGGGGCGC TGACGAAGAG
J23SpCA_27                GGGTTTGAT GGGGCGGGGA TGTAAGCGGG AAGGGAAACC GACCCGTTTA GTCTGCCGCT CCCAACAGCC CGAGTTTCCA GCCCCTTGAG TAAGGGGCAA GAAACCTAAG CCGCCTGGTG GATGGCTCGG CTCGGGGCGC TGACGAAGG
J23SpCA_3                CGGGTTTGAT GGGGCGGGGA TGTAAGCGGG AAGGGAAACC GACCCGTTTA GTCTGCCGCT CCCAACAGCC CGAGTTTCCA GCCCCTTGAG TAAGGGGCAA GAAACCTAAG CCGCCTGGTG GATGGCTCGG CTCGGGGCGC TGACAAAG
J23SpCA_21                 GGTTTGAT GGGGCGGGGA TGTAAGCGGG AAGGGAAACC GACCCGTTTA GTCTGCCGCT CCCAACAGCC CGAGTTTCCA GCCCCTTGAG TAAGGGGCAA GAAACCTAAG CCGCCTGGTG GATGGCTCGG CTCGGGGCGC TGACGAAGGG
J23SpCA_21                GGGTTTGAT GGGGCGGGGA TGTAAGCGGG AAGGGAAACC GACCCGTTTA GTCTGCCGCT CCCAACAGCC CGAGTTTCCA GCCCCTTGAG TAAGGGGCAA GAAACCTAAG CCGCCTGGTG GATGGCTCGG CTCGGGGCGC TGACGAAG
J23SpCA_17                 GGTTTGAT GGGGCGGGGA TGTAAGCGGG AAGGGAAACC GACCCGTTTA GTCTGCCGCT CCCAACAGCC CGAGTTTCCA GCCCCTTGAG TAAGGGGCAA GAAACCTAAG CCGCCTGGTG GATGGCTCGG CTCGGGGCGC TGACGAAGG
J23SpCA_10               CGGGTTTGAT GGGGCGGGGA TGTAAGCGGG AAGGGAAACC GACCCGTTTA GTCTGCCGCT CCCAACAGCC CGAGTTTCCA GCCCCTTGAG TAAGGGGCAA GAAACCTAAG CCGCCTGGTG GATGGCTCGG CTCGGGGCGC TGACGAA
J23SpCA_9                                CGGGGA TGTAAGCGGG AAGGGAAACC GACCCGTTTA GTCTGCCGCT CCCAACAGCC CGAGTTTCCA GCCCCTTGAG TAAGGGGCAA GAAACCTAAG CCGCCTGGTG GATGGCTCGG CTCGGGGCGC TGACGAAGGG CGTGGCAAGC TG
J23SpCA_8                                CGGGGA TGTAAGCGGG AAGGGAAACC GACCCGTTTA GTCTGCCGCT CCCAACAGCC CGAGTTTCCA GCCCCTTGAG TAAGGGGCAA GAAACCTAAG CCGCCTGGTG GATGGCTCGG CTCGGGGCGC TGACGAAGGG CGTGGCAAGC TGCGAT
J23SpCA_5                                CGGGGA TGTAAGCGGG AAGGGAAACC GACCCGTTTA GTCTGCCGCT CCCAACAGCC CGAGTTTCCA GCCCCTTGAG TAAGGGGCAA GAAACCTAAG CCGCCTGGTG GATGGCTCGG CTCGGGGCGC TGACGAAGGG CGTGGCAAGC TGCG
J23SpCA_4                                CGGGGA TGTAAGCGGG AAGGGAAACC GACCCGTTTA GTCTGCCGCT CCCAACAGCC CGAGTTTCCA GCCCCTTGAG TAAGGGGCAA GAAACCTAAG CCGCCTGGTG GATGGCTCGG CTCGGGGCGC TGACGAAGGG CGTGGCAAGC TGC
J23SpCA_2                            GGGGCGGGGA TGTAAGCGGG AAGGGAAACC GACCCGTTTA GTCTGCCGCT CCCAACAGCC CGAGTTTCCA GCCCCTTGAG TAAGGGGCAA GAAACCTAAG CCGCCTGGTG GATGGCTCGG CTCGGGGCGC TGACGAAGGG CGTGGCAAGC TGCG
J23SpCA_2                             GGGCGGGGA TGTAAGCGGG AAGGGAAACC GACCCGTTTA GTCTGCCGCT CCCAACAGCC CGAGTTTCCA GCCCCTTGAG TAAGGGGCAA GAAACCTAAG CCGCCTGGTG GATGGCTCGG CTCGGGGCGC TGACGAAGGG CGTGGCAAGC TG
J23SpCA_2                              GGCGGGGA TGTAAGCGGG AAGGGAAACC GACCCGTTTA GTCTGCCGCT CCCAACAGCC CGAGTTTCCA GCCCCTTGAG TAAGGGGCAA GAAACCTAAG CCGCCTGGTG GATGGCTCGG CTCGGGGCGC TGACGAAGGG CGTGGCAAGC TAC
J23SpCA_4                                CGGGGA TGTAAGCGGG AAGGGAAACC GACCCGTTTA GTCTGCCGCT CCCAACAGCC CGAGTTTCCA GCCCCTTGAG TAAGGGGCAA GAAACCTAAG CCGCCTGGTG GATGGCTCGG CTCGGGGCGC TGACGAAGGG CGTGGCAAGC
J23SpCA_102                 GTTTGAT GGGGCGGGGA TGTAAGCGGG AAGGGAAACC GACCCGTTTA GTCTGCCGCT CCCAACAGCC CGAGTTTCCA GCCCCTTGAG TAAGGGGCAA GAAACCTAAG CCGCCTGGTG GATGGCTCGG CTCGGGGCGC TGACGAAGGG
J23SpCA_4                   GTTTGAT GGGGCGGGGA TGTAAGCGGG AAGGGAAACC GACCCGTTTA GTCTGCCGCT CCCAACAGCC CGAGTTTCCA GCCCCTTGAG TAAGGGGCAA GAAACCTAAG CCGCCTGGTG GATGGCTCGG CTCGGGGCGC TGACGAAGGG G
J23SpCA_42                  GTTTGAT GGGGCGGGGA TGTAAGCGGG AAGGGAAACC GACCCGTTTA GTCTGCCGCT CCCAACAGCC CGAGTTTCCA GCCCCTTGAG TAAGGGGCAA GAAACCTAAG CCGCCTGGTG GATGGCTCGG CTCGGGGCGC TGACGAAGGG C
J23SpCA_62                  GTTTGAT GGGGCGGGGA TGTAAGCGGG AAGGGAAACC GACCCGTTTA GTCTGCCGCT CCCAACAGCC CGAGTTTCCA GCCCCTTGAG TAAGGGGCAA GAAACCTAAG CCGCCTGGTG GATGGCTCGG CTCGGGGCGC TGACGAAGG
J23SpCA_2                  GGTTTGAT GGGGCGGGGA TGTAAGCGGG AAGGGAAACC GACCCGTTTA GTCTGCCGCT CCCAACAGCC CGAGTTTCCA GCCCCTTGAG TAAGGGGCAA GAAACCTAAG CCGCCTGGTG GATGGCTCGG CTCGGGGCGC TGACGAAGAG
J23SpCA_2                   GTTTGAT GGGGCGGGGA TGTAAGCGGG AAGGGAAACC GACCCGTTTA GTCTGCCGCT CCCAACAGCC CGAGTTTCCA GCCCCTTGAG TAAGGGGCAA GAAACCTAAG CCGCCTGGTG GATGGCTCGG CTCGGGGCGC TGATGAAGGG CG
J23SpCA_2                   GTTTGAT GGGGCGGGGA TGTAAGCGGG AAGGGAAACC GACCCGTTTA GTCTGCCGCT CCCAACAGCC CGAGTTTCCA GCCCCTTGAG TAAGGGGCAA GAAACCTAAG CCGCCTGGTG GATGGCTCGG CTCGGGGCGC TGATGAAGGG
J23SpCA_2                  GGTTTGAT GGGGCGGGGA TGTAAGCGGG AAGGGAAACC GACCCGTTTA GTCTGCCGCT CCCAACAGCC CGAGTTTCCA GCCCCTTGAG TAAGGGGCAA GAAACCTAAG CCGCCTGGTG GATGGCTTGG CTCGGGGCGC TGACGAAGGG
J23SpCA_53                  GTTTGAT GGGGCGGGGA TGTAAGCGGG AAGGGAAACC GACCCGTTTA GTCTGCCGCT CCCAACAGCC CGAGTTTCCA GCCCCTTGAG TAAGGGGCAA GAAACCTAAG CCGCCTGGTG GATGGCTCGG CTCGGGGCGC TGACGAAG
J23SpCA_3                   GTTTGAT GGGGCGGGGA TGTAAGCGGG AAGGGAAACC GACCCGTTTA GTCTGCCGCT CCCAACAGCC CGAGTTTCCA GCCCCTTGAG TAAGGGGCAA GAAACCTAAG CCGCCTGGTG GATGGCTCGG CTCGGGGCGC TGACGAAAG
J23SpCA_25                 GGTTTGAT GGGGCGGGGA TGTAAGCGGG AAGGGAAACC GACCCGTTTA GTCTGCCGCT CCCAACAGCC CGAGTTTCCA GCCCCTTGAG TAAGGGGCAA GAAACCTAAG CCGCCTGGTG GATGGCTCGG CTCGGGGCGC TGACGAAG
J23SpCA_18            GA CGGGTTTGAT GGGGCGGGGA TGTAAGCGGG AAGGGAAACC GACCCGTTTA GTCTGCCGCT CCCAACAGCC CGAGTTTCCA GCCCCTTGAG TAAGGGGCAA GAAACCTAAG CCGCCTGGTG GATGGCTCGG CTCGGGGCGC TGACG
J23SpCA_5            AGA CGGGTTTGAT GGGGCGGGGA TGTAAGCGGG AAGGGAAACC GACCCGTTTA GTCTGCCGCT CCCAACAGCC CGAGTTTCCA GCCCCTTGAG TAAGGGGCAA GAAACCTAAG CCGCCTGGTG GATGGCTCGG CTCGGGGCGC TGACG
J23SpCA_6              A CGGGTTTGAT GGGGCGGGGA TGTAAGCGGG AAGGGAAACC GACCCGTTTA GTCTGCCGCT CCCAACAGCC CGAGTTTCCA GCCCCTTGAG TAAGGGGCAA GAAACCTAAG CCGCCTGGTG GATGGCTCGG CTCGGGGCGC TGACG
J23SpCA_3                CGGGTTTGAT GGGGCGGGGA TGTAAGCGGG AAGGGAAACC GACCCGTTTA GTCTGCCGCT CCCAACAGCC CGAGTTTCCA GCCCCTTGAG TAAGGGGCAA GAAACCTAAG CCGCCTGGTG GATGGCTCGG CTCGGGGCGC TGACGAG
J23SpCA_2             GA CGGGTTTGAT GGGGCGGGGA TGTAAGCGGG AAGGGAAACC GACCCGTTTA GTCTGCCGCT CCCAACAGCC CGAGTTTCCA GCCCCTTGAG TAAGGGGCAA GAAACCTAAG CCGCCTGGTG GATGGCTCGG CTCGGGGCGC TGACGAG
J23SpCA_2             GA CGGGTTTGAT GGGGCGGGGA TGTAAGCGGG AAGGGAAACC GACCCGTTTA GTCTGCCGCT CCCAACAGCC CGAGTTTCCA GCCCCTTGAG TAAGGGGCAA GAAACCTAAG CCGCCTGGTG GATGGCTCGG CTCGGGGCGC TGACGA
J23SpCA_3             GA CGGGTTTGAT GGGGCGGGGA TGTAAGCGGG AAGGGAAACC GACCCGTTTA GTCTGCCGCT CCCAACAGCC CGAGTTTCCA GCCCCTTGAG TAAGGGGCAA GAAACCTAAG CCGCCTGGTG GATGGCTCGG CTCGGGGCGC TGAC
```

```
J23SpCA_12 CGGGTTTGAT GGGGCGGGGA TGTAAGCGGG AAGGGAAACC GACCCGTTTA GTCTGCCGCT CCCAACAGCC CGAGTTTCCA GCCCCTTGAG TAAGGGGCAA GAAACCTAAG CCGCCTGGTG GATGGCTCGG CTCGGGGCGC TGACG
J23SpCA_46 CGCAGAAGA CGGGTTTGAT GGGGCGGGGA TGTAAGCGGG AAGGGAAACC GACCCGTTTA GTCTGCCGCT CCCAACAGCC CGAGTTTCCA GCCCCTTGAG TAAGGGGCAA GAAACCTAAG CCGCCTGGTG GATGGCTCGG CTCGGG
J23SpCA_32 CCGCAGAAGA CGGGTTTGAT GGGGCGGGGA TGTAAGCGGG AAGGGAAACC GACCCGTTTA GTCTGCCGCT CCCAACAGCC CGAGTTTCCA GCCCCTTGAG TAAGGGGCAA GAAACCTAAG CCGCCTGGTG GATGGCTCGG CTCGGG
J23SpCA_4 CCGCAGAAGA CGGGTTTGAT GGGGCGGGGA TGTAAGCGGG AAGGGAAACC GACCCGTTTA GTCTGCCGCT CCCAACAGCC CGAGTTTCCA GCCCCTTGAG TAAGGGGCAA GAAACCTAAG CCGCCTGGTG GATGGCTCGG CTCGGGG
J23SpCA_19 C CCGCAGAAGA CGGGTTTGAT GGGGCGGGGA TGTAAGCGGG AAGGGAAACC GACCCGTTTA GTCTGCCGCT CCCAACAGCC CGAGTTTCCA GCCCCTTGAG TAAGGGGCAA GAAACCTAAG CCGCCTGGTG GATGGCTCGG CTCGGG
J23SpCA_9 CGCAGAAGA CGGGTTTGAT GGGGCGGGGA TGTAAGCGGG AAGGGAAACC GACCCGTTTA GTCTGCCGCT CCCAACAGCC CGAGTTTCCA GCCCCTTGAG TAAGGGGCAA GAAACCTAAG CCGCCTGGTG GATGGCTCGG CTCGGGG
J23SpCA_2 GCAGAAGA CGGGTTTGAT GGGGCGGGGA TGTAAGCGGG AAGGGAAACC GACCCGTTTA GTCTGCCGCT CCCAACAGCC CGAGTTTCCA GCCCCTTGAG TAAGGGGCAA GAAACCTAAG CCGCCTGGTG GATGGCTCGG CTCGGGG
J23SpCA_6 CCGCAGAAGA CGGGTTTGAT GGGGCGGGGA TGTAAGCGGG AAGGGAAACC GACCCGTTTA GTCTGCCGCT CCCAACAGCC CGAGTTTCCA GCCCCTTGAG TAAGGGGCAA GAAACCTAAG CCGCCTGGTG GATGGCTCGG CTCGG
J23SpCA_13 CGCAGAAGA CGGGTTTGAT GGGGCGGGGA TGTAAGCGGG AAGGGAAACC GACCCGTTTA GTCTGCCGCT CCCAACAGCC CGAGTTTCCA GCCCCTTGAG TAAGGGGCAA GAAACCTAAG CCGCCTGGTG GATGGCTCGG CTCGG
J23SpCA_3 GCAGAAGA CGGGTTTGAT GGGGCGGGGA TGTAAGCGGG AAGGGAAACC GACCCGTTTA GTCTGCCGCT CCCAACAGCC CGAGTTTCCA GCCCCTTGAG TAAGGGGCAA GAAACCTAAG CCGCCTGGTG GATGGCTCGG CTCGGG
J23SpCA_3 CGCAGAAGA CGGGTTTGAT GGGGCGGGGA TGTAAGCGGG AAGGGAAACC GACCCGTTTA GTCTGCCGCT CCCAACAGCC CGAGTTTCCA GCCCCTTGAG TAAGGGGCAA GAAACCTAAG CCGCCTGGTG GATGGCTCGG ATCGGG
J23SpCA_21 CAGAAGA CGGGTTTGAT GGGGCGGGGA TGTAAGCGGG AAGGGAAACC GACCCGTTTA GTCTGCCGCT CCCAACAGCC CGAGTTTCCA GCCCCTTGAG TAAGGGGCAA GAAACCTAAG CCGCCTGGTG GATGGCTCGG CTCGGG
J23SpCA_7 CGCAGAAGA CGGGTTTGAT GGGGCGGGGA TGTAAGCGGG AAGGGAAACC GACCCGTTTA GTCTGCCGCT CCCAACAGCC CGAGTTTCCA GCCCCTTGAG TAAGGGGCAA GAAACCTAAG CCGCCTGGTG GATGGCTCGG CTCG
J23SpCA_4 GGGAAACC GACCCGTTTA GTCTGCCGCT CCCAACAGCC CGAGTTTCCA GCCCCTTGAG TAAGGGGCAA GAAACCTAAG CCGCCTGGTG GATGGCTCGG CTCGGGGCGC TGACGAAGGG CGTGGCAAGC TGCGATAAGC CCCGGTGAGG CGCAGG
J23SpCA_4 CGGG AAGGGAAACC GACCCGTTTA GTCTGCCGCT CCCAACAGCC CGAGTTTCCA GCCCCTTGAG TAAGGGGCAA GAAACCTAAG CCGCCTGGTG GATGGCTCGG CTCGGGGCGC TGACGAAGGG CGTGGCAAGC TGCGATAAGC CCCGGTGAG
J23SpCA_4 AGCGGG AAGGGAAACC GACCCGTTTA GTCTGCCGCT CCCAACAGCC CGAGTTTCCA GCCCCTTGAG TAAGGGGCAA GAAACCTAAG CCGCCTGGTG GATGGCTCGG CTCGGGGCGC TGACGAAGGG CGTGGCAAGC TGCGATAAGC CCCGGTGAG
J23SpCA_2 GGG AAGGGAAACC GACCCGTTTA GTCTGCCGCT CCCAACAGCC CGAGTTTCCA GCCCCTTGAG TAAGGGGCAA GAAACCTAAG CCGCCTGGTG GATGGCTCGG CTCGGGGCGC TGACGAAGGG CGTGGCAAGC TGCGATAAGC CCCGGTGAGG CG
J23SpCA_2 GCGGG AAGGGAAACC GACCCGTTTA GTCTGCCGCT CCCAACAGCC CGAGTTTCCA GCCCCTTGAG TAAGGGGCAA GAAACCTAAG CCGCCTGGTG GATGGCTCGG CTCGGGGCGC TGACGAAGGG CGTGGCAAGC TGCGATAAGC CCCGGTGAGG CG
J23SpCA_2 GGA TGTAAGCGGG AAGGGAAACC GACCCGTTTA GTCTGCCGCT CCCAACAGCC CGAGTTTCCA GCCCCTTGAG TAAGGGGCAA GAAACCTAAG CCGCCTGGTG GATGGCTCGG CTCGGGGCGC TGACGAAGGG CGTGGCAAGC TGCGATAAGC CCCG
J23SpCA_3 TGAT GGGGCGGGGA TGTAAGCGGG AAGGGAAACC GACCCGTTTA GTCTGCCGCT CCCAACAGCC CGAGTTTCCA GCCCCTTGAG TAAGGGGCAA GAAACCTAAG CCGCCTGGTG GATGGCTCGG CTCGGGGCGC TGACGAAGGG CG
J23SpCA_3 GAT GGGGCGGGGA TGTAAGCGGG AAGGGAAACC GACCCGTTTA GTCTGCCGCT CCCAACAGCC CGAGTTTCCA GCCCCTTGAG TAAGGGGCAA GAAACCTAAG CCGCCTGGTG GATGGCTCGG CTCGGGGCGC TGACGAAGGG C
J23SpCA_4 GTTTGAT GGGGCGGGGA TGTAAGCGGG AAGGGAAACC GACCCGTTTA GTCTGCCGCT CCCAACAGCC CGAGTTTCCA GCCCCTTGAG TAAGGGGCAA GAAACCTAAG CCGCCTGGTG GATGGCTCGG CTCGGGGCGC TGACGAA
J23SpCA_2 GGGTTTGAT GGGGCGGGGA TGTAAGCGGG AAGGGAAACC GACCCGTTTA GTCTGCCGCT CCCAACAGCC CGAGTTTCCA GCCCCTTGAG TAAGGGGCAA GAAACCTAAG CCGCCTGGTG GATGGCTCGG CTCGGGGCGC TGACGGAG
J23SpCA_2 GGGTTTGAT GGGGCGGGGA TGTAAGCGGG AAGGGAAACC GACCCGTTTA GTCTGCCGCT CCCAACAGCC CGAGTTTCCA GCCCCTTGAG TAAGGGGCAA GAAACCTAAG CCGCCTGGTG GATGGCTCGG CTCGGGGCGC TGACG
J23SpCA_3 GTTTGAT GGGGCGGGGA TGTAAGCGGG AAGGGAAACC GACCCGTTTA GTCTGCCGCT CCCAACAGCC CGAGTTTCCA GCCCCTTGAG TAAGGGGCAA GAAACCTAAG CCGCCTGGTG GATGGCTCGG CTCGGGGCGC TGACAAAG
J23SpCA_2 TTTGAT GGGGCGGGGA TGTAAGCGGG AAGGGAAACC GACCCGTTTA GTCTGCCGCT CCCAACAGCC CGAGTTTCCA GCCCCTTGAG TAAGGGGCAA GAAACCTAAG CCGCCTGGTG GATGGCTCGG CTCGGGGCGC TGACGAAG
J23SpCA_2 GTTTGAT GGGGCGGGGA TGTAAGCGGG AAGGGAAACC GACCCGTTTA GTTTGCCGCT CCCAACAGCC CGAGTTTCCA GCCCCTTGAG TAAGGGGCAA GAAACCTAAG CCGCCTGGTG GATGGCTCGG CTCGGGGCGC TGACGAAG
J23SpCA_2 TGAT GGGGCGGGGA TGTAAGCGGG AAGGGAAACC GACCCGTTTA GTCTGCCGCT CCCAACAGCC CGAGTTTCCA GCCCCTTGAG TAAGGGGCAA GAAACCTAAG CCGCCTGGTG GATGGCTCGG CTCGGGGCGC TGACGAAG
J23SpCA_11 CGGGTTTGAT GGGGCGGGGA TGTAAGCGGG AAGGGAAACC GACCCGTTTA GTCTGCCGCT CCCAACAGCC CGAGTTTCCA GCCCCTTGAG TAAGGGGCAA GAAACCTAAG CCGCCTGGTG GATGGCTCGG CTCGGGGCGC TGAC
J23SpCA_70 CGGGGA TGTAAGCGGG AAGGGAAACC GACCCGTTTA GTCTGCCGCT CCCAACAGCC CGAGTTTCCA GCCCCTTGAG TAAGGGGCAA GAAACCTAAG CCGCCTGGTG GATGGCTCGG CTCGGGGCGC TGACGAAGGG CGTGG
J23SpCA_5 GCGGGGA TGTAAGCGGG AAGGGAAACC GACCCGTTTA GTCTGCCGCT CCCAACAGCC CGAGTTTCCA GCCCCTTGAG TAAGGGGCAA GAAACCTAAG CCGCCTGGTG GATGGCTCGG CTCGGGGCGC TGACGAAGGG CGTGG
J23SpCA_3 GGCGGGGA TGTAAGCGGG AAGGGAAACC GACCCGTTTA GTCTGCCGCT CCCAACAGCC CGAGTTTCCA GCCCCTTGAG TAAGGGGCAA GAAACCTAAG CCGCCTGGTG GATGGCTCGG CTCGGGGCGC TGACGAAGGG CGTGG
J23SpCA_2 GGGGCGGGGA TGTAAGCGGG AAGGGAAACC GACCCGTTTA GTCTGCCGCT CCCAACAGCC CGAGTTTCCA GCCCCTTGAG TAAGGGGCAA GAAACCTAAG CCGCCTGGTG GATGGCTCGG CTCGGGGCGC TGACGAAGGG CGTG
J23SpCA_17 CGGGGA TGTAAGCGGG AAGGGAAACC GACCCGTTTA GTCTGCCGCT CCCAACAGCC CGAGTTTCCA GCCCCTTGAG TAAGGGGCAA GAAACCTAAG CCGCCTGGTG GATGGCTCGG CTCGGGGCGC TGACGAAGGG CGTGGC
J23SpCA_5 CGGGGA TGTAAGCGGG AAGGGAAACC GACCCGTTTA GTCTGCCGCT CCCAACAGCC CGAGTTTCCA GCCCCTTGAG TAAGGGGCAA GAAACCTAAG CCGCCTGGTG GATGGCTCGG CTCGGGGCGC TGACGAAGGG CGTGGCAAG
J23SpCA_4 CGGGGA TGTAAGCGGG AAGGGAAACC GACCCGTTTA GTCTGCCGCT CCCAACAGCC CGAGTTTCCA GCCCCTTGAG TAAGGGGCAA GAAACCTAAG CCGCCTGGTG GATGGCTCGG CTCGGGGCGC TGACGAAGGG CGTGGCG
J23SpCA_2 GA CGGGTTTGAT GGGGCGGGGA TGTAAGCGGG AAGGGAAACC GACCCGTTTA GTCTGCCGCT CCCAACAGCC CGAGTTTCCA GCCCCTTGAG TAAGGGGCAA GAAACCTAAG CCGCCTGGTG GATGGCTCGG CTCGGGGCGC T
J23SpCA_4 AGAAGA CGGGTTTGAT GGGGCGGGGA TGTAAGCGGG AAGGGAAACC GACCCGTTTA GTCTGCCGCT CCCAACAGCC CGAGTTTCCA GCCCCTTGAG TAAGGGGCAA GAAACCTAAG CCGCCTGGTG GATGGCTCGG CTCGGG
J23SpCA_2 GTTTGAT GGGGCGGGGA TGTAAGCGGG AAGGGAAACC GACCCGTTTA GTCTGCCGCT CCCAACAGCC CGAGTTTCCA GCCCCTTGAG TAAGGGGCAA GAAACCTAAG CCGCCTGGTG GATGGCTCGG CTCGGGGCGC TGACG
J23SpCA_2 GTTTGAT GGGGCGGGGA TGTAAGCGGG AAGGGAAACC GACCCGTTTA GTCTGCCGCT CCCAACAGCC CGAGTTTCCA GCCCCTTGAG TAAGGGGCAA GAAACCTAAG CCGCCTGGTG GATGGCTCGG CTCGGGGCGC TGAC
J23SpCA_11 GGGCGGGGA TGTAAGCGGG AAGGGAAACC GACCCGTTTA GTCTGCCGCT CCCAACAGCC CGAGTTTCCA GCCCCTTGAG TAAGGGGCAA GAAACCTAAG CCGCCTGGTG GATGGCTCGG CTCGGGGCGC TGACGAAGGG
J23SpCA_8 GGGGCGGGGA TGTAAGCGGG AAGGGAAACC GACCCGTTTA GTCTGCCGCT CCCAACAGCC CGAGTTTCCA GCCCCTTGAG TAAGGGGCAA GAAACCTAAG CCGCCTGGTG GATGGCTCGG CTCGGGGCGC TGACGAAGGG C
J23SpCA_4 T GGGGCGGGGA TGTAAGCGGG AAGGGAAACC GACCCGTTTA GTCTGCCGCT CCCAACAGCC CGAGTTTCCA GCCCCTTGAG TAAGGGGCAA GAAACCTAAG CCGCCTGGTG GATGGCTCGG CTCGGGGCGC TGACGAAGGG C
J23SpCA_3 T GGGGCGGGGA TGTAAGCGGG AAGGGAAACC GACCCGTTTA GTCTGCCGCT CCCAACAGCC CGAGTTTCCA GCCCCTTGAG TAAGGGGCAA GAAACCTAAG CCGCCTGGTG GATGGCTCGG CTCGGGGCGC TGACGAAGG
J23SpCA_7 GGGGCGGGGA TGTAAGCGGG AAGGGAAACC GACCCGTTTA GTCTGCCGCT CCCAACAGCC CGAGTTTCCA GCCCCTTGAG TAAGGGGCAA GAAACCTAAG CCGCCTGGTG GATGGCTCGG CTCGGGGCGC TGACGAAGGG
J23SpCA_2 T GGGGCGGGGA TGTAAGCGGG AAGGGAAACC GACCCGTTTA GTCTGCCGCT CCCAACAGCC CGAGTTTCCA GCCCCTTGAG TAAGGGGCAA GAAACCTAAG CCGCCTGGTG GATGGCTCGG CTCGGGGCGC TGACGAAG
J23SpCA_2 GGGGCGGGGA TGTAAGCGGG AAGGGAAACC GACCCGTTTA GTCTGCCGCT CCCAACAGCC CGAGTTTCCA GCCCCTTGAG TAAGGGGCAA GAAACCTAAG CCGCCTGGTG GATGGCTCGG CTCGGGGCGC TGACGAAGG
J23SpCA_7 GA CGGGTTTGAT GGGGCGGGGA TGTAAGCGGG AAGGGAAACC GACCCGTTTA GTCTGCCGCT CCCAACAGCC CGAGTTTCCA GCCCCTTGAG TAAGGGGCAA GAAACCTAAG CCGCCTGGTG GATGGCTCGG CTCGGG
J23SpCA_7 AGA CGGGTTTGAT GGGGCGGGGA TGTAAGCGGG AAGGGAAACC GACCCGTTTA GTCTGCCGCT CCCAACAGCC CGAGTTTCCA GCCCCTTGAG TAAGGGGCAA GAAACCTAAG CCGCCTGGTG GATGGCTCGG CTCGGG
J23SpCA_3 AGA CGGGTTTGAT GGGGCGGGGA TGTAAGCGGG AAGGGAAACC GACCCGTTTA GTCTGCCGCT CCCAACAGCC CGAGTTTCCA GCCCCTTGAG TAAGGGGCAA GAAACCTAAG CCGCCTGGTG GATGGCTCGG CTCGGGG
J23SpCA_3 GA CGGGTTTGAT GGGGCGGGGA TGTAAGCGGG AAGGGAAACC GACCCGTTTA GTCTGCCGCT CCCAACAGCC CGAGTTTCCA GCCCCTTGAG TAAGGGGCAA GAAACCTAAG CCGCCTGGTG GATGGCTCGG CTCGGGG
J23SpCA_82 CGGGGA TGTAAGCGGG AAGGGAAACC GACCCGTTTA GTCTGCCGCT CCCAACAGCC CGAGTTTCCA GCCCCTTGAG TAAGGGGCAA GAAACCTAAG CCGCCTGGTG GATGGCTCGG CTCGGGGCGC TGACGAAGGG CG
J23SpCA_40 CGGGGA TGTAAGCGGG AAGGGAAACC GACCCGTTTA GTCTGCCGCT CCCAACAGCC CGAGTTTCCA GCCCCTTGAG TAAGGGGCAA GAAACCTAAG CCGCCTGGTG GATGGCTCGG CTCGGGGCGC TGACGAAGGG CGTG
J23SpCA_3 CGGGGA TGTAAGCGGG AAGGGAAACC GACCCGTTTA GTCTGCCGCT CCCAACAGCC CGAGTTTCCA GCCCCTTGAG TAAGGGGCAA GAAACCTAAG CCGCCTGGTG GATGGCTCGG CTCGGGGCGC TGACGAAGGG CGT
J23SpCA_3 CGGGGA TGTAAGCGGG AAGGGAAACC GACCCGTTTA GTCTGCCGCT CCCAACAGCC CGAGTTTCCA GCCCCTTGAG TAAGGGGCAA GAAACCTAAG CCGCCTGGTG GATGGCTCGG CTCGGGGCGC TGACGAAGGG CGTAG
J23SpCA_3 CGGGGA TGTAAGCGGG AAGGGAAACC GACCCGTTTA GTCTGCCGCT CCCAACAGCC CGAGTTTCCA GCCCCTTGAG TAAGGGGCAA GAAACCTAAG CCGCCTGGTG GATGGCTCGG CTCGGGGCGC TGACGAAGGG CGGGG
J23SpCA_7 GGGGA TGTAAGCGGG AAGGGAAACC GACCCGTTTA GTCTGCCGCT CCCAACAGCC CGAGTTTCCA GCCCCTTGAG TAAGGGGCAA GAAACCTAAG CCGCCTGGTG GATGGCTCGG CTCGGGGCGC TGACGAAGGG CGTG
J23SpCA_8 GGGGA TGTAAGCGGG AAGGGAAACC GACCCGTTTA GTCTGCCGCT CCCAACAGCC CGAGTTTCCA GCCCCTTGAG TAAGGGGCAA GAAACCTAAG CCGCCTGGTG GATGGCTCGG CTCGGGGCGC TGACGAAGGG CGTGG
J23SpCA_5 GGGA TGTAAGCGGG AAGGGAAACC GACCCGTTTA GTCTGCCGCT CCCAACAGCC CGAGTTTCCA GCCCCTTGAG TAAGGGGCAA GAAACCTAAG CCGCCTGGTG GATGGCTCGG CTCGGGGCGC TGACGAAGGG CGTGG
J23SpCA_6 GGCGGGGA TGTAAGCGGG AAGGGAAACC GACCCGTTTA GTCTGCCGCT CCCAACAGCC CGAGTTTCCA GCCCCTTGAG TAAGGGGCAA GAAACCTAAG CCGCCTGGTG GATGGCTCGG CTCGGGGCGC TGACGAAGGG CG
J23SpCA_4 GCGGGGA TGTAAGCGGG AAGGGAAACC GACCCGTTTA GTCTGCCGCT CCCAACAGCC CGAGTTTCCA GCCCCTTGAG TAAGGGGCAA GAAACCTAAG CCGCCTGGTG GATGGCTCGG CTCGGGGCGC TGACGAAGGG CG
J23SpCA_2 GGGCGGGGA TGTAAGCGGG AAGGGAAACC GACCCGTTTA GTCTGCCGCT CCCAACAGCC CGAGTTTCCA GCCCCTTGAG TAAGGGGCAA GAAACCTAAG CCGCCTGGTG GATGGCTCGG CTCGGGGCGC TGACGAAGGG C
J23SpCA_3 GGCGGGGA TGTAAGCGGG AAGGGAAACC GACCCGTTTA GTCTGCCGCT CCCAACAGCC CGAGTTTCCA GCCCCTTGAG TAAGGGGCAA GAAACCTAAG CCGCCTGGTG GATGGCTCGG CTCGGGGCGC TGACGAAGGG C
J23SpCA_130 CGGGGA TGTAAGCGGG AAGGGAAACC GACCCGTTTA GTCTGCCGCT CCCAACAGCC CGAGTTTCCA GCCCCTTGAG TAAGGGGCAA GAAACCTAAG CCGCCTGGTG GATGGCTCGG CTCGGGGCGC TGACGAAGGG C
J23SpCA_3 GCGGGGA TGTAAGCGGG AAGGGAAACC GACCCGTTTA GTCTGCCGCT CCCAACAGCC CGAGTTTCCA GCCCCTTGAG TAAGGGGCAA GAAACCTAAG CCGCCTGGTG GATGGCTCGG CTCGGGGCGC TGACGAAGGG C
J23SpCA_9 GCGGGGA TGTAAGCGGG AAGGGAAACC GACCCGTTTA GTCTGCCGCT CCCAACAGCC CGAGTTTCCA GCCCCTTGAG TAAGGGGCAA GAAACCTAAG CCGCCTGGTG GATGGCTCGG CTCGGGGCGC TGACGAAGGG
J23SpCA_6 CAGAAGA CGGGTTTGAT GGGGCGGGGA TGTAAGCGGG AAGGGAAACC GACCCGTTTA GTCTGCCGCT CCCAACAGCC CGAGTTTCCA GCCCCTTGAG TAAGGGGCAA GAAACCTAAG CCGCCTGGTG GATGGCTCGG CT
J23SpCA_5 CGCAGAAGA CGGGTTTGAT GGGGCGGGGA TGTAAGCGGG AAGGGAAACC GACCCGTTTA GTCTGCCGCT CCCAACAGCC CGAGTTTCCA GCCCCTTGAG TAAGGGGCAA GAAACCTAAG CCGCCTGGTG GATGGCTCGG C
J23SpCA_4 GCAGAAGA CGGGTTTGAT GGGGCGGGGA TGTAAGCGGG AAGGGAAACC GACCCGTTTA GTCTGCCGCT CCCAACAGCC CGAGTTTCCA GCCCCTTGAG TAAGGGGCAA GAAACCTAAG CCGCCTGGTG GATGGCTCGG CTCA
J23SpCA_3 CGCAGAAGA CGGGTTTGAT GGGGCGGGGA TGTAAGCGGG AAGGGAAACC GACCCGTTTA GTCTGCCGCT CCCAACAGCC CGAGTTTCCA GCCCCTTGAG TAAGGGGCAA GAAACCTAAG CCGCCTGGTG GATGGCTCGG CT
J23SpCA_4 CCGCAGAAGA CGGGTTTGAT GGGGCGGGGA TGTAAGCGGG AAGGGAAACC GACCCGTTTA GTCTGCCGCT CCCAACAGCC CGAGTTTCCA GCCCCTTGAG TAAGGGGCAA GAAACCTAAG CCGCCTGGTG GATGGCTCGG
J23SpCA_3 CGCAGAAGA CGGGTTTGAT GGGGCGGGGA TGTAAGCGGG AAGGGAAACC GACCCGTTTA GTCTGCCGCT CCCAACAGCC CGAGTTTCCA GCCCCTTGAG TAAGGGGCAA GAAACCTAAG CCGCCTGGTG GATGGCTCGG
J23SpCA_3 CGCAGAAGA CGGGTTTGAT GGGGCGGGGA TGTAAGCGGG AAGGGAAACC GACCCGTTTA GTCTGCCGCT CCCAACAGCC CGAGTTTCCA GCCCCTTGAG TAAGGGGCAA GAAACCTAAG CCGCCTGGTG GATGGCTCG
J23SpCA_10 GGCGGGGA TGTAAGCGGG AAGGGAAACC GACCCGTTTA GTCTGCCGCT CCCAACAGCC CGAGTTTCCA GCCCCTTGAG TAAGGGGCAA GAAACCTAAG CCGCCTGGTG GATGGCTCGG CTCGGGGCGC TGACGAAGGG
J23SpCA_8 GGCGGGGA TGTAAGCGGG AAGGGAAACC GACCCGTTTA GTCTGCCGCT CCCAACAGCC CGAGTTTCCA GCCCCTTGAG TAAGGGGCAA GAAACCTAAG CCGCCTGGTG GATGGCTCGG CTCGGGGCGC TGACGAAGG
J23SpCA_5 GGGCGGGGA TGTAAGCGGG AAGGGAAACC GACCCGTTTA GTCTGCCGCT CCCAACAGCC CGAGTTTCCA GCCCCTTGAG TAAGGGGCAA GAAACCTAAG CCGCCTGGTG GATGGCTCGG CTCGGGGCGC TGACGAAGG
J23SpCA_2 GGGGCGGGGA TGTAAGCGGG AAGGGAAACC GACCCGTTTA GTCTGCCGCT CCCAACAGCC CGAGTTTCCA GCCCCTTGAG TAAGGGGCAA GAAACCTAAG CCGCCTGGTG GATGGCTCGG CTCGGGGCGC TGACGAAG
J23SpCA_7 GGGCGGGGA TGTAAGCGGG AAGGGAAACC GACCCGTTTA GTCTGCCGCT CCCAACAGCC CGAGTTTCCA GCCCCTTGAG TAAGGGGCAA GAAACCTAAG CCGCCTGGTG GATGGCTCGG CTCGGGGCGC TGACGAAG
J23SpCA_8 GGA TGTAAGCGGG AAGGGAAACC GACCCGTTTA GTCTGCCGCT CCCAACAGCC CGAGTTTCCA GCCCCTTGAG TAAGGGGCAA GAAACCTAAG CCGCCTGGTG GATGGCTCGG CTCGGGGCGC TGACGAAGGG CGTG
J23SpCA_289 CGGGGA TGTAAGCGGG AAGGGAAACC GACCCGTTTA GTCTGCCGCT CCCAACAGCC CGAGTTTCCA GCCCCTTGAG TAAGGGGCAA GAAACCTAAG CCGCCTGGTG GATGGCTCGG CTCGGGGCGC TGACGAAGGG
J23SpCA_5 CGGGGA TGTAAGCGGG AAGGGAAACC GACCCGTTTA GTCTGCCGCT CCCAACAGCC CGAGTTTCCA GCCCCTTGAG TAAGGGGCAA GAAACCTAAG CCGCCTGGTG GATGGCTCGG CTCGGGGCGC TGACGAAGGG G
J23SpCA_4 GGGGA TGTAAGCGGG AAGGGAAACC GACCCGTTTA GTCTGCCGCT CCCAACAGCC CGAGTTTCCA GCCCCTTGAG TAAGGGGCAA GAAACCTAAG CCGCCTGGTG GATGGCTCGG CTCGGGGCGC TGACGAAGGG T
J23SpCA_4 CGGGGA TGTAAGCGGG AAGGGAAACC GACCCGTTTA GTCTGCCGCT CCCAACAGCC CGAGTTTCCA GCCCCTTGAG TAAGGGGCAA GAAACCTAAG CCGCCTGGTG GATGGCTCGG CTCGGGGCGC TGACGAAGGG TC
J23SpCA_5 CGGGGA TGTAAGCGGG AAGGGAAACT GACCCGTTTA GTCTGCCGCT CCCAACAGCC CGAGTTTCCA GCCCCTTGAG TAAGGGGCAA GAAACCTAAG CCGCCTGGTG GATGGCTCGG CTCGGGGCGC TGACGAAGGG C
J23SpCA_180 CGGGGA TGTAAGCGGG AAGGGAAACC GACCCGTTTA GTCTGCCGCT CCCAACAGCC CGAGTTTCCA GCCCCTTGAG TAAGGGGCAA GAAACCTAAG CCGCCTGGTG GATGGCTCGG CTCGGGGCGC TGACGAAGG
J23SpCA_11 GGGA TGTAAGCGGG AAGGGAAACC GACCCGTTTA GTCTGCCGCT CCCAACAGCC CGAGTTTCCA GCCCCTTGAG TAAGGGGCAA GAAACCTAAG CCGCCTGGTG GATGGCTCGG CTCGGGGCGC TGACGAAGGG CG
J23SpCA_10 GGGGA TGTAAGCGGG AAGGGAAACC GACCCGTTTA GTCTGCCGCT CCCAACAGCC CGAGTTTCCA GCCCCTTGAG TAAGGGGCAA GAAACCTAAG CCGCCTGGTG GATGGCTCGG CTCGGGGCGC TGACGAAGGG CG
J23SpCA_7 GGGGA TGTAAGCGGG AAGGGAAACC GACCCGTTTA GTCTGCCGCT CCCAACAGCC CGAGTTTCCA GCCCCTTGAG TAAGGGGCAA GAAACCTAAG CCGCCTGGTG GATGGCTCGG CTCGGGGCGC TGACGAAGGG C
J23SpCA_5 GCGGGGA TGTAAGCGGG AAGGGAAACC GACCCGTTTA GTCTGCCGCT CCCAACAGCC CGAGTTTCCA GCCCCTTGAG TAAGGGGCAA GAAACCTAAG CCGCCTGGTG GATGGCTCGG CTCGGGGCGC TGACGAAGG
J23SpCA_3 CGGGGA TGTAAGCGGG AAGGGAAACC GACCCGTTTA GTCTGCCGCT CCCAACAGCC CGAGTTTCCA GCCCCTTGAG TAAGGGGCAA GAAACCTAAG CCGCCTGGTG GATGGCTCGG CTCGGGGCGC TGACGAAGAG
J23SpCA_25 GGGGA TGTAAGCGGG AAGGGAAACC GACCCGTTTA GTCTGCCGCT CCCAACAGCC CGAGTTTCCA GCCCCTTGAG TAAGGGGCAA GAAACCTAAG CCGCCTGGTG GATGGCTCGG CTCGGGGCGC TGACGAAGGG
J23SpCA_4 A TGTAAGCGGG AAGGGAAACC GACCCGTTTA GTCTGCCGCT CCCAACAGCC CGAGTTTCCA GCCCCTTGAG TAAGGGGCAA GAAACCTAAG CCGCCTGGTG GATGGCTCGG CTCGGGGCGC TGACGAAGGG CGTGG
J23SpCA_29 CGGGTTTGAT GGGGCGGGGA TGTAAGCGGG AAGGGAAACC GACCCGTTTA GTCTGCCGCT CCCAACAGCC CGAGTTTCCA GCCCCTTGAG TAAGGGGCAA GAAACCTAAG CCGCCTGGTG GATGGCTCGG CTCGGG
J23SpCA_4 A CGGGTTTGAT GGGGCGGGGA TGTAAGCGGG AAGGGAAACC GACCCGTTTA GTCTGCCGCT CCCAACAGCC CGAGTTTCCA GCCCCTTGAG TAAGGGGCAA GAAACCTAAG CCGCCTGGTG GATGGCTCGG CTCGGG
J23SpCA_5 CGGGTTTGAT GGGGCGGGGA TGTAAGCGGG AAGGGAAACC GACCCGTTTA GTCTGCCGCT CCCAACAGCC CGAGTTTCCA GCCCCTTGAG TAAGGGGCAA GAAACCTAAG CCGCCTGGTG GATGGCTCGG CTCGGGG
J23SpCA_2 GA CGGGTTTGAT GGGGCGGGGA TGTAAGCGGG AAGGGAAACC GACCCGTTTA GTCTGCCGCT CCCAACAGCC CGAGTTTCCA GCCCCTTGAG TAAGGGGCAA GAAACCTAAG CCGCCTGGTG GATGGCTCGG CTGGG
J23SpCA_2 GA CGGGTTTGAT GGGGCGGGGA TGTAAGCGGG AAGGGAAACC GACCCGTTTA GTCTGCCGCT CCCAACAGCC CGAGTTTCCA GCCCCTTGAG TAAGGGGCAA GAAACCTAAG CCGCCTGGTG GATGGCTCGG CTCGG
J23SpCA_3 A CGGGTTTGAT GGGGCGGGGA TGTAAGCGGG AAGGGAAACC GACCCGTTTA GTCTGCCGCT CCCAACAGCC CGAGTTTCCA GCCCCTTGAG TAAGGGGCAA GAAACCTAAG CCGCCTGGTG GATGGCTCGG CTCGG
J23SpCA_2 GGGTTTGAT GGGGCGGGGA TGTAAGCGGG AAGGGAAACC GACCCGTTTA GTCTGCCGCT CCCAACAGCC CGAGTTTCCA GCCCCTTGAG TAAGGGGCAA GAAACCTAAG CCGCCTGGTG GATGGCTCGG CTCGGG
J23SpCA_6 CGGG AAGGGAAACC GACCCGTTTA GTCTGCCGCT CCCAACAGCC CGAGTTTCCA GCCCCTTGAG TAAGGGGCAA GAAACCTAAG CCGCCTGGTG GATGGCTCGG CTCGGGGCGC TGACGAAGGG CGTGGCAAGC TGCGAT
J23SpCA_3 CGGG AAGGGAAACC GACCCGTTTA GTCTGCCGCT CCCAACAGCC CGAGTTTCCA GCCCCTTGAG TAAGGGGCAA GAAACCTAAG CCGCCTGGTG GATGGCTCGG CTCGGGGCGC TGACGAAGGG CGTGGCAAGC TGCG
J23SpCA_6 GCGGGGA TGTAAGCGGG AAGGGAAACC GACCCGTTTA GTCTGCCGCT CCCAACAGCC CGAGTTTCCA GCCCCTTGAG TAAGGGGCAA GAAACCTAAG CCGCCTGGTG GATGGCTCGG CTCGGGGCGC TGACGAAG
J23SpCA_5 GGCGGGGA TGTAAGCGGG AAGGGAAACC GACCCGTTTA GTCTGCCGCT CCCAACAGCC CGAGTTTCCA GCCCCTTGAG TAAGGGGCAA GAAACCTAAG CCGCCTGGTG GATGGCTCGG CTCGGGGCGC TGACGAAG
J23SpCA_14 GGGA TGTAAGCGGG AAGGGAAACC GACCCGTTTA GTCTGCCGCT CCCAACAGCC CGAGTTTCCA GCCCCTTGAG TAAGGGGCAA GAAACCTAAG CCGCCTGGTG GATGGCTCGG CTCGGGGCGC TGACGAAGGG C
J23SpCA_4 GGA TGTAAGCGGG AAGGGAAACC GACCCGTTTA GTCTGCCGCT CCCAACAGCC CGAGTTTCCA GCCCCTTGAG TAAGGGGCAA GAAACCTAAG CCGCCTGGTG GATGGCTCGG CTCGGGGCGC TGACGAAGGG CG
J23SpCA_2 GA TGTAAGCGGG AAGGGAAACC GACCCGTTTA GTCTGCCGCT CCCAACAGCC CGAGTTTCCA GCCCCTTGAG TAAGGGGCAA GAAACCTAAG CCGCCTGGTG GATGGCTCGG CTCGGGGCGC TGACGAAGGG CG
J23SpCA_9 GGA TGTAAGCGGG AAGGGAAACC GACCCGTTTA GTCTGCCGCT CCCAACAGCC CGAGTTTCCA GCCCCTTGAG TAAGGGGCAA GAAACCTAAG CCGCCTGGTG GATGGCTCGG CTCGGGGCGC TGACGAAGGG C
J23SpCA_174 CGGGGA TGTAAGCGGG AAGGGAAACC GACCCGTTTA GTCTGCCGCT CCCAACAGCC CGAGTTTCCA GCCCCTTGAG TAAGGGGCAA GAAACCTAAG CCGCCTGGTG GATGGCTCGG CTCGGGGCGC TGACGAAG
J23SpCA_3 CGGGGA TGTAAGCGGG AAGGGAAACC GACCCGTTTA GTCTGCCGCT CCCAACAGCC CGAGTTTCCA GCCCCTTGAG TAAGGGGCAA GAAACCTAAG CCGCCTGGTG GATGGCTCGG CTCGGGGCGC TGACAAAGG
J23SpCA_8 CGGGGA TGTAAGCGGG AAGGGAAACC GACCCGTTTA GTCTGCCGCT CCCAACAGCC CGAGTTTCCA GCCCCTTGAG TAAGGGGCAA GAAACCTAAG CCGCCTGGTG GATGGCTCGG CTCGGGGCGC TGACGAA
J23SpCA_3 GGGGA TGTAAGCGGG AAGGGAAACC GACCCGTTTA GTCTGCCGCT CCCAACAGCC CGAGTTTCCA GCCCCTTGAG TAAGGGGCAA GAAACCTAAG CCGCCTGGTG GATGGCTCGG CTCGGGGCGC TGACGAAGAG
J23SpCA_15 GGGGA TGTAAGCGGG AAGGGAAACC GACCCGTTTA GTCTGCCGCT CCCAACAGCC CGAGTTTCCA GCCCCTTGAG TAAGGGGCAA GAAACCTAAG CCGCCTGGTG GATGGCTCGG CTCGGGGCGC TGACGAAGG
J23SpCA_5 CGGGGA TGTAAGCGGG AAGGGAAACC GACCCGTTTA GTCTGCCGCT CCCAACAGCC CGAGTTTCCA GCCCCTTGAG TAAGGGGCAA GAAACCTAAG CCGCCTGGTG GATGGCTCGG CTCGGGGCGC TGACGAAAG
J23SpCA_31 GGGGA TGTAAGCGGG AAGGGAAACC GACCCGTTTA GTCTGCCGCT CCCAACAGCC CGAGTTTCCA GCCCCTTGAG TAAGGGGCAA GAAACCTAAG CCGCCTGGTG GATGGCTCGG CTCGGGGCGC TGACGAAG
J23SpCA_2 GAT GGGGCGGGGA TGTAAGCGGG AAGGGAAACC GACCCGTTTA GTCTGCCGCT CCCAACAGCC CGAGTTTCCA GCCCCTTGAG TAAGGGGCAA GAAACCTAAG CCGCCTGGTG GATGGCTCGG CTCGGGGCGC TG
J23SpCA_5 GTAAGCGGG AAGGGAAACC GACCCGTTTA GTCTGCCGCT CCCAACAGCC CGAGTTTCCA GCCCCTTGAG TAAGGGGCAA GAAACCTAAG CCGCCTGGTG GATGGCTCGG CTCGGGGCGC TGACGAAGGG CGTGG
J23SpCA_25 GGGA TGTAAGCGGG AAGGGAAACC GACCCGTTTA GTCTGCCGCT CCCAACAGCC CGAGTTTCCA GCCCCTTGAG TAAGGGGCAA GAAACCTAAG CCGCCTGGTG GATGGCTCGG CTCGGGGCGC TGACGAAGGG
J23SpCA_12 GGA TGTAAGCGGG AAGGGAAACC GACCCGTTTA GTCTGCCGCT CCCAACAGCC CGAGTTTCCA GCCCCTTGAG TAAGGGGCAA GAAACCTAAG CCGCCTGGTG GATGGCTCGG CTCGGGGCGC TGACGAAGGG
J23SpCA_16 GGGA TGTAAGCGGG AAGGGAAACC GACCCGTTTA GTCTGCCGCT CCCAACAGCC CGAGTTTCCA GCCCCTTGAG TAAGGGGCAA GAAACCTAAG CCGCCTGGTG GATGGCTCGG CTCGGGGCGC TGACGAAGG
J23SpCA_27 CGCAGAAGA CGGGTTTGAT GGGGCGGGGA TGTAAGCGGG AAGGGAAACC GACCCGTTTA GTCTGCCGCT CCCAACAGCC CGAGTTTCCA GCCCCTTGAG TAAGGGGCAA GAAACCTAAG CCGCCTGGTG GAT
J23SpCA_17 C CCGCAGAAGA CGGGTTTGAT GGGGCGGGGA TGTAAGCGGG AAGGGAAACC GACCCGTTTA GTCTGCCGCT CCCAACAGCC CGAGTTTCCA GCCCCTTGAG TAAGGGGCAA GAAACCTAAG CCGCCTGGTG GAT
J23SpCA_11 CGCAGAAGA CGGGTTTGAT GGGGCGGGGA TGTAAGCGGG AAGGGAAACC GACCCGTTTA GTCTGCCGCT CCCAACAGCC CGAGTTTCCA GCCCCTTGAG TAAGGGGCAA GAAACCTAAG CCGCCTGGTG GATGG
J23SpCA_6 CGCAGAAGA CGGGTTTGAT GGGGCGGGGA TGTAAGCGGG AAGGGAAACC GACCCGTTTA GTCTGCCGCT CCCAACAGCC CGAGTTTCCA GCCCCTTGAG TAAGGGGCAA GAAACCTAAG CCGCCTGGTG GATGGC
J23SpCA_3 CGCAGAAGA CGGGTTTGAT GGGGCGGGGA TGTAAGCGGG AAGGGAAACC GACCCGTTTA GTCTGCCGCT CCCAACAGCC CGAGTTTCCA GCCCCTTGAG TAAGGGGCAA GAAACCTAAG CCGCCTGGTG GATGGCC
J23SpCA_7 CCGCAGAAGA CGGGTTTGAT GGGGCGGGGA TGTAAGCGGG AAGGGAAACC GACCCGTTTA GTCTGCCGCT CCCAACAGCC CGAGTTTCCA GCCCCTTGAG TAAGGGGCAA GAAACCTAAG CCGCCTGGTG GATGG
J23SpCA_5 C CCGCAGAAGA CGGGTTTGAT GGGGCGGGGA TGTAAGCGGG AAGGGAAACC GACCCGTTTA GTCTGCCGCT CCCAACAGCC CGAGTTTCCA GCCCCTTGAG TAAGGGGCAA GAAACCTAAG CCGCCTGGTG GATG
J23SpCA_4 C CCGCAGAAGA CGGGTTTGAT GGGGCGGGGA TGTAAGCGGG AAGGGAAACC GACCCGTTTA GTCTGCCGCT CCCAACAGCC CGAGTTTCCA GCCCCTTGAG TAAGGGGCAA GAAACCTAAG CCGCCTGGTG GATGGC
J23SpCA_4 C CCGCAGAAGA CGGGTTTGAT GGGGCGGGGA TGTAAGCGGG AAGGGAAACC GACCCGTTTA GTCTGCCGCT CCCAACAGCC CGAGTTTCCA GCCCCTTGAG TAAGGGGCAA GAAACCTAAG CCGCCTGGTG GATGG
J23SpCA_2 GC CCGCAGAAGA CGGGTTTGAT GGGGCGGGGA TGTAAGCGGG AAGGGAAACC GACCCGTTTA GTCTGCCGCT CCCAACAGCC CGAGTTTCCA GCCCCTTGAG TAAGGGGCAA GAAACCTAAG CCGCCTGGTG GATGG
J23SpCA_3 CCGCAGAAGA CGGGTTTGAT GGGGCGGGGA TGTAAGCGGG AAGGGAAACC GACCCGTTTA GTCTGCCGCT CCCAACAGCC CGAGTTTCCA GCCCCTTGAG TAAGGGGCAA GAAACCTAAG CCGCCTGGTG GATGGC
J23SpCA_6 CCGCAGAAGA CGGGTTTGAT GGGGCGGGGA TGTAAGCGGG AAGGGAAACC GACCCGTTTA GTCTGCCGCT CCCAACAGCC CGAGTTTCCA GCCCCTTGAG TAAGGGGCAA GAAACCTAAG CCGCCTGGTG GATG
J23SpCA_9 CCGCAGAAGA CGGGTTTGAT GGGGCGGGGA TGTAAGCGGG AAGGGAAACC GACCCGTTTA GTCTGCCGCT CCCAACAGCC CGAGTTTCCA GCCCCTTGAG TAAGGGGCAA GAAACCTAAG CCGCCTGGTG GAT
J23SpCA_9 CGCAGAAGA CGGGTTTGAT GGGGCGGGGA TGTAAGCGGG AAGGGAAACC GACCCGTTTA GTCTGCCGCT CCCAACAGCC CGAGTTTCCA GCCCCTTGAG TAAGGGGCAA GAAACCTAAG CCGCCTGGTG GATG
J23SpCA_6 CAGAAGA CGGGTTTGAT GGGGCGGGGA TGTAAGCGGG AAGGGAAACC GACCCGTTTA GTCTGCCGCT CCCAACAGCC CGAGTTTCCA GCCCCTTGAG TAAGGGGCAA GAAACCTAAG CCGCCTGGTG GATGGC
J23SpCA_8 GGA TGTAAGCGGG AAGGGAAACC GACCCGTTTA GTCTGCCGCT CCCAACAGCC CGAGTTTCCA GCCCCTTGAG TAAGGGGCAA GAAACCTAAG CCGCCTGGTG GATGGCTCGG CTCGGGGCGC TGACGAAGG
J23SpCA_8 GA TGTAAGCGGG AAGGGAAACC GACCCGTTTA GTCTGCCGCT CCCAACAGCC CGAGTTTCCA GCCCCTTGAG TAAGGGGCAA GAAACCTAAG CCGCCTGGTG GATGGCTCGG CTCGGGGCGC TGACGAAGGG
J23SpCA_4 A TGTAAGCGGG AAGGGAAACC GACCCGTTTA GTCTGCCGCT CCCAACAGCC CGAGTTTCCA GCCCCTTGAG TAAGGGGCAA GAAACCTAAG CCGCCTGGTG GATGGCTCGG CTCGGGGCGC TGACGAAGGG C
J23SpCA_4 A TGTAAGCGGG AAGGGAAACC GACCCGTTTA GTCTGCCGCT CCCAACAGCC CGAGTTTCCA GCCCCTTGAG TAAGGGGCAA GAAACCTAAG CCGCCTGGTG GATGGCTCGG CTCGGGGCGC TGACGAAGGG
J23SpCA_6 GA TGTAAGCGGG AAGGGAAACC GACCCGTTTA GTCTGCCGCT CCCAACAGCC CGAGTTTCCA GCCCCTTGAG TAAGGGGCAA GAAACCTAAG CCGCCTGGTG GATGGCTCGG CTCGGGGCGC TGACGAAGG
J23SpCA_23 GGGA TGTAAGCGGG AAGGGAAACC GACCCGTTTA GTCTGCCGCT CCCAACAGCC CGAGTTTCCA GCCCCTTGAG TAAGGGGCAA GAAACCTAAG CCGCCTGGTG GATGGCTCGG CTCGGGGCGC TGACGAAG
J23SpCA_10 GGA TGTAAGCGGG AAGGGAAACC GACCCGTTTA GTCTGCCGCT CCCAACAGCC CGAGTTTCCA GCCCCTTGAG TAAGGGGCAA GAAACCTAAG CCGCCTGGTG GATGGCTCGG CTCGGGGCGC TGACGAAG
J23SpCA_2 TAAGCGGG AAGGGAAACC GACCCGTTTA GTCTGCCGCT CCCAACAGCC CGAGTTTCCA GCCCCTTGAG TAAGGGGCAA GAAACCTAAG CCGCCTGGTG GATGGCTCGG CTCGGGGCGC TGACGAAGGG CGTG
```

```
J23SpCA_5 CGGGTTTGAT GGGGCGGGGA TGTAAGCGGG AAGGGAAACC GACCCGTTTA GTCTGCCGCT CCCAACAGCC CGAGTTTCCA GCCCCTTGAG TAAGGGGCAA GAAACCTAAG CCGCCTGGTG GATGGCTCGG CTCG
J23SpCA_3 GA CGGGTTTGAT GGGGCGGGGA TGTAAGCGGG AAGGGAAACC GACCCGTTTA GTCTGCCGCT CCCAACAGCC CGAGTTTCCA GCCCCTTGAG TAAGGGGCAA GAAACCTAAG CCGCCTGGTG GATGGCTCGG CC
J23SpCA_2 GA CGGGTTTGAT GGGGCGGGGA TGTAAGCGGG AAGGGAAACC GACCCGTTTA GTCTGCCGCT CCCAACAGCC CGAGTTTCCA GCCCCTTGAG TAAGGGGCAA GAAACCTAAG CCGCCTGGTG GATGGCTCGG CT
J23SpCA_3 CGGGTTTGAT GGGGCGGGGA TGTAAGCGGG AAGGGAAACC GACCCGTTTA GTCTGCCGCT CCCAACAGCC CGAGTTTCCA GCCCCTTGAG TAAGGGGCAA GAAACCTAAG CCGCCTGGTG GATGGCTCGG CT
J23SpCA_13 GTTTGAT GGGGCGGGGA TGTAAGCGGG AAGGGAAACC GACCCGTTTA GTCTGCCGCT CCCAACAGCC CGAGTTTCCA GCCCCTTGAG TAAGGGGCAA GAAACCTAAG CCGCCTGGTG GATGGCTCGG CTCGGG
J23SpCA_4 GGTTTGAT GGGGCGGGGA TGTAAGCGGG AAGGGAAACC GACCCGTTTA GTCTGCCGCT CCCAACAGCC CGAGTTTCCA GCCCCTTGAG TAAGGGGCAA GAAACCTAAG CCGCCTGGTG GATGGCTCGG CTCGGG
J23SpCA_2 GTTTGAT GGGGCGGGGA TGTAAGCGGG AAGGGAAACC GACCCGTTTA GTCTGCCGCT CCCAACAGCC CGAGTTTCCA GCCCCTTGAG TAAGGGGCAA GAAACCTAAG CCGCCTGGTG GATGGCTTGG CTCGGG
J23SpCA_3 GTTTGAT GGGGCGGGGA TGTAAGCGGG AAGGGAAACC GACCCGTTTA GTCTGCCGCT CCCAACAGCC CGAGTTTCCA GCCCCTTGAG TAAGGGGCAA GAAACCTAAG CCGCCTGGTG GATGGCTCGG CTCGG
J23SpCA_12 CGGGGA TGTAAGCGGG AAGGGAAACC GACCCGTTTA GTCTGCCGCT CCCAACAGCC CGAGTTTCCA GCCCCTTGAG TAAGGGGCAA GAAACCTAAG CCGCCTGGTG GATGGCTCGG CTCGGGGCGC TGACG
J23SpCA_4 GCGGGGA TGTAAGCGGG AAGGGAAACC GACCCGTTTA GTCTGCCGCT CCCAACAGCC CGAGTTTCCA GCCCCTTGAG TAAGGGGCAA GAAACCTAAG CCGCCTGGTG GATGGCTCGG CTCGGGGCGC TGAC
J23SpCA_9 CGGGGA TGTAAGCGGG AAGGGAAACC GACCCGTTTA GTCTGCCGCT CCCAACAGCC CGAGTTTCCA GCCCCTTGAG TAAGGGGCAA GAAACCTAAG CCGCCTGGTG GATGGCTCGG CTCGGGGCGC TGAC
J23SpCA_7 GTAAGCGGG AAGGGAAACC GACCCGTTTA GTCTGCCGCT CCCAACAGCC CGAGTTTCCA GCCCCTTGAG TAAGGGGCAA GAAACCTAAG CCGCCTGGTG GATGGCTCGG CTCGGGGCGC TGACGAAGGG C
J23SpCA_5 GTAAGCGGG AAGGGAAACC GACCCGTTTA GTCTGCCGCT CCCAACAGCC CGAGTTTCCA GCCCCTTGAG TAAGGGGCAA GAAACCTAAG CCGCCTGGTG GATGGCTCGG CTCGGGGCGC TGACGAAGGG CG
J23SpCA_20 GTAAGCGGG AAGGGAAACC GACCCGTTTA GTCTGCCGCT CCCAACAGCC CGAGTTTCCA GCCCCTTGAG TAAGGGGCAA GAAACCTAAG CCGCCTGGTG GATGGCTCGG CTCGGGGCGC TGACGAAGGG
J23SpCA_4 TGTAAGCGGG AAGGGAAACC GACCCGTTTA GTCTGCCGCT CCCAACAGCC CGAGTTTCCA GCCCCTTGAG TAAGGGGCAA GAAACCTAAG CCGCCTGGTG GATGGCTCGG CTCGGGGCGC TGACGAAGGG
J23SpCA_8 A TGTAAGCGGG AAGGGAAACC GACCCGTTTA GTCTGCCGCT CCCAACAGCC CGAGTTTCCA GCCCCTTGAG TAAGGGGCAA GAAACCTAAG CCGCCTGGTG GATGGCTCGG CTCGGGGCGC TGACGAAG
J23SpCA_3 GA TGTAAGCGGG AAGGGAAACC GACCCGTTTA GTCTGCCGCT CCCAACAGCC CGAGTTTCCA GCCCCTTGAG TAAGGGGCAA GAAACCTAAG CCGCCTGGTG GATGGCTCGG CTCGGGGCGC TGACGAAG
J23SpCA_5 AAGCGGG AAGGGAAACC GACCCGTTTA GTCTGCCGCT CCCAACAGCC CGAGTTTCCA GCCCCTTGAG TAAGGGGCAA GAAACCTAAG CCGCCTGGTG GATGGCTCGG CTCGGGGCGC TGACGAAGGG C
J23SpCA_5 GGGAAACC GACCCGTTTA GTCTGCCGCT CCCAACAGCC CGAGTTTCCA GCCCCTTGAG TAAGGGGCAA GAAACCTAAG CCGCCTGGTG GATGGCTCGG CTCGGGGCGC TGACGAAGGG CGTGGCAAGC TGCGAT
J23SpCA_2 GGGAAACC GACCCGTTTA GTCTGCCGCT CCCAACAGCC CGAGTTTCCA GCCCCTTGAG TAAGGGGCAA GAAACCTAAG CCGCCTGGTG GATGGCTCGG CTCGGGGCGC TGACGAAGGG CGTGGCAAGC TGCGATAAGC C
J23SpCA_2 GGAAACC GACCCGTTTA GTCTGCCGCT CCCAACAGCC CGAGTTTCCA GCCCCTTGAG TAAGGGGCAA GAAACCTAAG CCGCCTGGTG GATGGCTCGG CTCGGGGCGC TGACGAAGGG CGTGGCAAGC TGCGATAAGC C
J23SpCA_2 GGGAAACC GACCCGTTTA GTCTGCCGCT CCCAACAGCC CGAGTTTCCA GCCCCTTGAG TAAGGGGCAA GAAACCTAAG CCGCCTGGTG GATGGCTCGG CTCGGGGCGC TGACGAAGGG CGTGGCAAGC TGCGATAAG
J23SpCA_2 GGGAAACC GACCCGTTTA GTCTGCCGCT CCCAACAGCC CGAGTTTCCA GCCCCTTGAG TAAGGGGCAA GAAACCTAAG CCGCCTGGTG GATGGCTCGG CTCGGGGCGC TGACGAAGGG CGTGGCAAGC TGCGATA
J23SpCA_4 GAAACC GACCCGTTTA GTCTGCCGCT CCCAACAGCC CGAGTTTCCA GCCCCTTGAG TAAGGGGCAA GAAACCTAAG CCGCCTGGTG GATGGCTCGG CTCGGGGCGC TGACGAAGGG CGTGGCAAGC TGCGAT
J23SpCA_3 GGGAAACC GACCCGTTTA GTCTGCCGCT CCCAACAGCC CGAGTTTCCA GCCCCTTGAG TAAGGGGCAA GAAACCTAAG CCGCCTGGTG GATGGCTCGG CTCGGGGCGC TGACGAAGGG CGTGGCAAGC TGCG
J23SpCA_3 AACC GACCCGTTTA GTCTGCCGCT CCCAACAGCC CGAGTTTCCA GCCCCTTGAG TAAGGGGCAA GAAACCTAAG CCGCCTGGTG GATGGCTCGG CTCGGGGCGC TGACGAAGGG CGTGGCAAGC TGCGAT
J23SpCA_14 CGGG AAGGGAAACC GACCCGTTTA GTCTGCCGCT CCCAACAGCC CGAGTTTCCA GCCCCTTGAG TAAGGGGCAA GAAACCTAAG CCGCCTGGTG GATGGCTCGG CTCGGGGCGC TGACGAAGGG CGTG
J23SpCA_8 CGGG AAGGGAAACC GACCCGTTTA GTCTGCCGCT CCCAACAGCC CGAGTTTCCA GCCCCTTGAG TAAGGGGCAA GAAACCTAAG CCGCCTGGTG GATGGCTCGG CTCGGGGCGC TGACGAAGGG CGTGG
J23SpCA_5 CGGG AAGGGAAACC GACCCGTTTA GTCTGCCGCT CCCAACAGCC CGAGTTTCCA GCCCCTTGAG TAAGGGGCAA GAAACCTAAG CCGCCTGGTG GATGGCTCGG CTCGGGGCGC TGACGAAGGG CGTGGC
J23SpCA_3 CGGG AAGGGAAACC GACCCGTTTA GTCTGCCGCT CCCAACAGCC CGAGTTTCCA GCCCCTTGAG TAAGGGGCAA GAAACCTAAG CCGCCTGGTG GATGGCTCGG CTCGGGGCGC TGACGAAGGG CGTGGCA
J23SpCA_2 GGG AAGGGAAACC GACCCGTTTA GTCTGCCGCT CCCAACAGCC CGAGTTTCCA GCCCCTTGAG TAAGGGGCAA GAAACCTAAG CCGCCTGGTG GATGGCTCGG CTCGGGGCGC TGACGAAGGG CGTGGCG
J23SpCA_3 CGGG AAGGGAAACC GACCCGTTTA GTCTGCCGCT CCCAACAGCC CGAGTTTCCA GCCCCTTGAG TAAGGGGCAA GAAACCTAAG CCGCCTGGTG GATGGCTCGG CTCGGGGCGC TGACGAAGGG CGTAG
J23SpCA_3 GGGA TGTAAGCGGG AAGGGAAACC GACCCGTTTA GTCTGCCGCT CCCAACAGCC CGAGTTTCCA GCCCCTTGAG TAAGGGGCAA GAAACCTAAG CCGCCTGGTG GATGGCTCGG CTCGGGGCGC TGACG
J23SpCA_53 CGCAGAAGA CGGGTTTGAT GGGGCGGGGA TGTAAGCGGG AAGGGAAACC GACCCGTTTA GTCTGCCGCT CCCAACAGCC CGAGTTTCCA GCCCCTTGAG TAAGGGGCAA GAAACCTAAG CCGCCTGGTG
J23SpCA_34 CCGCAGAAGA CGGGTTTGAT GGGGCGGGGA TGTAAGCGGG AAGGGAAACC GACCCGTTTA GTCTGCCGCT CCCAACAGCC CGAGTTTCCA GCCCCTTGAG TAAGGGGCAA GAAACCTAAG CCGCCTGGTG
J23SpCA_21 C CCGCAGAAGA CGGGTTTGAT GGGGCGGGGA TGTAAGCGGG AAGGGAAACC GACCCGTTTA GTCTGCCGCT CCCAACAGCC CGAGTTTCCA GCCCCTTGAG TAAGGGGCAA GAAACCTAAG CCGCCTGGTG
J23SpCA_3 C CCGCAGAAGA CGGGTTTGAT GGGGCGGGGA TGTAAGCGGG AAGGGAAACC GACCCGTTTA GTCTGCCGCT CCCAACAGCC CGAGTTTCCA GCCCCTTGAG TAAGGGGCAA GAAACCTAAG CCGCCTGGTG G
J23SpCA_3 CCGCAGAAGA CGGGTTTGAT GGGGCGGGGA TGTAAGCGGG AAGGGAAACC GACCCGTTTA GTCTGCCGCT CCCAACAGCC CGAGTTTCCA GCCCCTTGAG TAAGGGGCAA GAAACCTAAG CCGCCTGGTG G
J23SpCA_3 C CCGCAGAAGA CGGGTTTGAT GGGGCGGGGA TGTAAGCGGG AAGGGAAACC GACCCGTTTA GTCTGCCGCT CCCAACAGCC CGAGTTTCCA GCCCCTTGAG TAAGGGGCAA GAAACCTAAG TCGCCTGGTG
J23SpCA_5 CGCAGAAGA CGGGTTTGAT GGGGCGGGGA TGTAAGCGGG AAGGGAAACC GACCCGTTTA GTCTGCCGCT CCCAACAGCC CGAGTTTCCA GCCCCTTGAG TAAGGGGCAA GAAACCTAAG CCGCCTGGTG G
J23SpCA_7 CAGAAGA CGGGTTTGAT GGGGCGGGGA TGTAAGCGGG AAGGGAAACC GACCCGTTTA GTCTGCCGCT CCCAACAGCC CGAGTTTCCA GCCCCTTGAG TAAGGGGCAA GAAACCTAAG CCGCCTGGTG GAT
J23SpCA_2 GCAGAAGA CGGGTTTGAT GGGGCGGGGA TGTAAGCGGG AAGGGAAACC GACCCGTTTA GTCTGCCGCT CCCAACAGCC CGAGTTTCCA GCCCCTTGAG TAAGGGGCAA GAAACCTAAG CCGCCTGGTG GAT
J23SpCA_3 CAGAAGA CGGGTTTGAT GGGGCGGGGA TGTAAGCGGG AAGGGAAACC GACCCGTTTA GTCTGCCGCT CCCAACAGCC CGAGTTTCCA GCCCCTTGAG TAAGGGGCAA GAAACCTAAG CCGCCTGGTG GATG
J23SpCA_4 AGAAGA CGGGTTTGAT GGGGCGGGGA TGTAAGCGGG AAGGGAAACC GACCCGTTTA GTCTGCCGCT CCCAACAGCC CGAGTTTCCA GCCCCTTGAG TAAGGGGCAA GAAACCTAAG CCGCCTGGTG GAT
J23SpCA_5 CGCAGAAGA CGGGTTTGAT GGGGCGGGGA TGTAAGCGGG AAGGGAAACC GACCCGTTTA GTCTGCCGCT CCCAACAGCC CGAGTTTCCA GCCCCTTGAG TAAGGGGCAA GAAACCTAAG CCGCCTGGT
J23SpCA_8 GCAGAAGA CGGGTTTGAT GGGGCGGGGA TGTAAGCGGG AAGGGAAACC GACCCGTTTA GTCTGCCGCT CCCAACAGCC CGAGTTTCCA GCCCCTTGAG TAAGGGGCAA GAAACCTAAG CCGCCTGGTG
J23SpCA_3 CCGCAGAAGA CGGGTTTGAT GGGGCGGGGA TGTAAGCGGG AAGGGAAACC GACCCGTTTA GTCTGCCGCT CCCAACAGCC CGAGTTTCCA GCCCCTTGAG TAAGGGGCAA GAAACCTAAG CCGCCTGG
J23SpCA_3 C CCGCAGAAGA CGGGTTTGAT GGGGCGGGGA TGTAAGCGGG AAGGGAAACC GACCCGTTTA GTCTGCCGCT CCCAACAGCC CGAGTTTCCA GCCCCTTGAG TAAGGGGCAA GAAACCTAAG CCGCCTGG
J23SpCA_13 AAGCGGG AAGGGAAACC GACCCGTTTA GTCTGCCGCT CCCAACAGCC CGAGTTTCCA GCCCCTTGAG TAAGGGGCAA GAAACCTAAG CCGCCTGGTG GATGGCTCGG CTCGGGGCGC TGACGAAGGG
J23SpCA_13 GTAAGCGGG AAGGGAAACC GACCCGTTTA GTCTGCCGCT CCCAACAGCC CGAGTTTCCA GCCCCTTGAG TAAGGGGCAA GAAACCTAAG CCGCCTGGTG GATGGCTCGG CTCGGGGCGC TGACGAAGG
J23SpCA_13 GTAAGCGGG AAGGGAAACC GACCCGTTTA GTCTGCCGCT CCCAACAGCC CGAGTTTCCA GCCCCTTGAG TAAGGGGCAA GAAACCTAAG CCGCCTGGTG GATGGCTCGG CTCGGGGCGC TGACGAAG
J23SpCA_3 TGTAAGCGGG AAGGGAAACC GACCCGTTTA GTCTGCCGCT CCCAACAGCC CGAGTTTCCA GCCCCTTGAG TAAGGGGCAA GAAACCTAAG CCGCCTGGTG GATGGCTCGG CTCGGGGCGC TGACGAAG
J23SpCA_5 AGCGGG AAGGGAAACC GACCCGTTTA GTCTGCCGCT CCCAACAGCC CGAGTTTCCA GCCCCTTGAG TAAGGGGCAA GAAACCTAAG CCGCCTGGTG GATGGCTCGG CTCGGGGCGC TGACGAAGGG CG
J23SpCA_6 AAGCGGG AAGGGAAACC GACCCGTTTA GTCTGCCGCT CCCAACAGCC CGAGTTTCCA GCCCCTTGAG TAAGGGGCAA GAAACCTAAG CCGCCTGGTG GATGGCTCGG CTCGGGGCGC TGACGAAGG
J23SpCA_5 GGG AAGGGAAACC GACCCGTTTA GTCTGCCGCT CCCAACAGCC CGAGTTTCCA GCCCCTTGAG TAAGGGGCAA GAAACCTAAG CCGCCTGGTG GATGGCTCGG CTCGGGGCGC TGACGAAGGG CGTG
J23SpCA_3 GG AAGGGAAACC GACCCGTTTA GTCTGCCGCT CCCAACAGCC CGAGTTTCCA GCCCCTTGAG TAAGGGGCAA GAAACCTAAG CCGCCTGGTG GATGGCTCGG CTCGGGGCGC TGACGAAGGG CGTGG
J23SpCA_4 T GGGGCGGGGA TGTAAGCGGG AAGGGAAACC GACCCGTTTA GTCTGCCGCT CCCAACAGCC CGAGTTTCCA GCCCCTTGAG TAAGGGGCAA GAAACCTAAG CCGCCTGGTG GATGGCTCGG CTCGGG
J23SpCA_3 GGGGCGGGGA TGTAAGCGGG AAGGGAAACC GACCCGTTTA GTCTGCCGCT CCCAACAGCC CGAGTTTCCA GCCCCTTGAG TAAGGGGCAA GAAACCTAAG CCGCCTGGTG GATGGCTCGG CTCGGG
J23SpCA_2 GTAAGCGGG AAGGGAAACC GACCCGTTTA GTCTGCCGCT CCCAACAGCC CGAGTTTCCA GCCCCTTGAG TAAGGGGCAA GAAACCTAAG CCGCCTGGTG GATGGCTCGG CTCGGGGCGC TGACGAA
J23SpCA_3 CTGCCGCT CCCAACAGCC CGAGTTTCCA GCCCCTTGAG TAAGGGGCAA GAAACCTAAG CCGCCTGGTG GATGGCTCGG CTCGGGGCGC TGACGAAGGG CGTGGCAAGC TGCGATAAGC CCCGGTGAGG CGCAGGCAGC CGTCGAGCCG GGGATTCCCG AATG
J23SpCA_2 GCCCCGTTTA GTCTGCCGCT CCCAACAGCC CGAGTTTCCA GCCCCTTGAG TAAGGGGCAA GAAACCTAAG CCGCCTGGTG GATGGCTCGG CTCGGGGCGC TGACGAAGGG CGTGGCAAGC TGCGATAAGC CCCGGTGAGG CGCAGGCAGC
J23SpCA_2 GTTTA GTCTGCCGCT CCCAACAGCC CGAGTTTCCA GCCCCTTGAG TAAGGGGCAA GAAACCTAAG CCGCCTGGTG GATGGCTCGG CTCGGGGCGC TGACGAAGGG CGTGGCAAGC TGCGATAAGC CCCGGTGAG
J23SpCA_3 GA CGGGTTTGAT GGGGCGGGGA TGTAAGCGGG AAGGGAAACC GACCCGTTTA GTCTGCCGCT CCCAACAGCC CGAGTTTCCA GCCCCTTGAG TAAGGGGCAA GAAACCTAAG CCGCCTGGTG GAT
J23SpCA_2 GA CGGGTTTGAT GGGGCGGGGA TGTAAGCGGG AAGGGAAACC GACCCGTTTA GTCTGCCGCT CCCAACAGCC CGAGTTTCCA GCCCCTTGAG TAAGGGGCAA GAAACCTAAG CCGCCTGGTG GATG
J23SpCA_3 GAAGA CGGGTTTGAT GGGGCGGGGA TGTAAGCGGG AAGGGAAACC GACCCGTTTA GTCTGCCGCT CCCAACAGCC CGAGTTTCCA GCCCCTTGAG TAAGGGGCAA GAAACCTAAG CCGCCTGGTG GA
J23SpCA_3 AAGA CGGGTTTGAT GGGGCGGGGA TGTAAGCGGG AAGGGAAACC GACCCGTTTA GTCTGCCGCT CCCAACAGCC CGAGTTTCCA GCCCCTTGAG TAAGGGGCAA GAAACCTAAG CCGCCTGGTG GAT
J23SpCA_16 CAGAAGA CGGGTTTGAT GGGGCGGGGA TGTAAGCGGG AAGGGAAACC GACCCGTTTA GTCTGCCGCT CCCAACAGCC CGAGTTTCCA GCCCCTTGAG TAAGGGGCAA GAAACCTAAG CCGCCTGGTG
J23SpCA_11 AGAAGA CGGGTTTGAT GGGGCGGGGA TGTAAGCGGG AAGGGAAACC GACCCGTTTA GTCTGCCGCT CCCAACAGCC CGAGTTTCCA GCCCCTTGAG TAAGGGGCAA GAAACCTAAG CCGCCTGGTG
J23SpCA_5 AAGCGGG AAGGGAAACC GACCCGTTTA GTCTGCCGCT CCCAACAGCC CGAGTTTCCA GCCCCTTGAG TAAGGGGCAA GAAACCTAAG CCGCCTGGTG GATGGCTCGG CTCGGGGCGC TGACGAAG
J23SpCA_11 GCGGG AAGGGAAACC GACCCGTTTA GTCTGCCGCT CCCAACAGCC CGAGTTTCCA GCCCCTTGAG TAAGGGGCAA GAAACCTAAG CCGCCTGGTG GATGGCTCGG CTCGGGGCGC TGACGAAGGG
J23SpCA_26 CGGG AAGGGAAACC GACCCGTTTA GTCTGCCGCT CCCAACAGCC CGAGTTTCCA GCCCCTTGAG TAAGGGGCAA GAAACCTAAG CCGCCTGGTG GATGGCTCGG CTCGGGGCGC TGACGAAGGG CG
J23SpCA_26 CGGG AAGGGAAACC GACCCGTTTA GTCTGCCGCT CCCAACAGCC CGAGTTTCCA GCCCCTTGAG TAAGGGGCAA GAAACCTAAG CCGCCTGGTG GATGGCTCGG CTCGGGGCGC TGACGAAGGG C
J23SpCA_3 GCGGG AAGGGAAACC GACCCGTTTA GTCTGCCGCT CCCAACAGCC CGAGTTTCCA GCCCCTTGAG TAAGGGGCAA GAAACCTAAG CCGCCTGGTG GATGGCTCGG CTCGGGGCGC TGACGAAGG
J23SpCA_2 GGGA TGTAAGCGGG AAGGGAAACC GACCCGTTTA GTCTGCCGCT CCCAACAGCC CGAGTTTCCA GCCCCTTGAG TAAGGGGCAA GAAACCTAAG CCGCCTGGTG GATGGCTCGG CTCGGGCGC T
J23SpCA_4 GGG AAGGGAAACC GACCCGTTTA GTCTGCCGCT CCCAACAGCC CGAGTTTCCA GCCCCTTGAG TAAGGGGCAA GAAACCTAAG CCGCCTGGTG GATGGCTCGG CTCGGGGCGC TGACGAAGGG C
J23SpCA_7 CCGCAGAAGA CGGGTTTGAT GGGGCGGGGA TGTAAGCGGG AAGGGAAACC GACCCGTTTA GTCTGCCGCT CCCAACAGCC CGAGTTTCCA GCCCCTTGAG TAAGGGGCAA GAAACCTAAG CCGCCTG
J23SpCA_5 C CCGCAGAAGA CGGGTTTGAT GGGGCGGGGA TGTAAGCGGG AAGGGAAACC GACCCGTTTA GTCTGCCGCT CCCAACAGCC CGAGTTTCCA GCCCCTTGAG TAAGGGGCAA GAAACCTAAG CCGCCTG
J23SpCA_6 CGCAGAAGA CGGGTTTGAT GGGGCGGGGA TGTAAGCGGG AAGGGAAACC GACCCGTTTA GTCTGCCGCT CCCAACAGCC CGAGTTTCCA GCCCCTTGAG TAAGGGGCAA GAAACCTAAG CCGCCTG
J23SpCA_3 CGCAGAAGA CGGGTTTGAT GGGGCGGGGA TGTAAGCGGG AAGGGAAACC GACCCGTTTA GTCTGCCGCT CCCAACAGCC CGAGTTTCCA GCCCCTTGAG TAAGGGGCAA GAAACCTAAG CCGCCTGG
J23SpCA_6 CGCAGAAGA CGGGTTTGAT GGGGCGGGGA TGTAAGCGGG AAGGGAAACC GACCCGTTTA GTCTGCCGCT CCCAACAGCC CGAGTTTCCA GCCCCTTGAG TAAGGGGCAA GAAACCTAAG CCGCCT
J23SpCA_4 CGCAGAAGA CGGGTTTGAT GGGGCGGGGA TGTAAGCGGG AAGGGAAACC GACCCGTTTA GTCTGCCGCT CCCAACAGCC CGAGTTTCCA GCCCCTTGAG TAAGGGGCAA GAAACCTAAG CCGCC
J23SpCA_3 AGCGGG AAGGGAAACC GACCCGTTTA GTCTGCCGCT CCCAACAGCC CGAGTTTCCA GCCCCTTGAG TAAGGGGCAA GAAACCTAAG CCGCCTGGTG GATGGCTCGG CTCGGGGCGC TGACGAAG
J23SpCA_45 CGGG AAGGGAAACC GACCCGTTTA GTCTGCCGCT CCCAACAGCC CGAGTTTCCA GCCCCTTGAG TAAGGGGCAA GAAACCTAAG CCGCCTGGTG GATGGCTCGG CTCGGGGCGC TGACGAAGGG
J23SpCA_3 CGGG AAGGGAAACC GACCCGTTTA GTCTGCCGCT CCCAACAGCC CGAGTTTCCA GCCCCTTGAG TAAGGGGCAA GAAACCTAAG CCGCCTGGTG GATGGCTCGG CTCGGGGCGC TGACGAAGGG G
J23SpCA_5 GG AAGGGAAACC GACCCGTTTA GTCTGCCGCT CCCAACAGCC CGAGTTTCCA GCCCCTTGAG TAAGGGGCAA GAAACCTAAG CCGCCTGGTG GATGGCTCGG CTCGGGGCGC TGACGAAGGG CG
J23SpCA_4 GGG AAGGGAAACC GACCCGTTTA GTCTGCCGCT CCCAACAGCC CGAGTTTCCA GCCCCTTGAG TAAGGGGCAA GAAACCTAAG CCGCCTGGTG GATGGCTCGG CTCGGGGCGC TGACGAAGGG
J23SpCA_5 GCGGG AAGGGAAACC GACCCGTTTA GTCTGCCGCT CCCAACAGCC CGAGTTTCCA GCCCCTTGAG TAAGGGGCAA GAAACCTAAG CCGCCTGGTG GATGGCTCGG CTCGGGGCGC TGACGAAG
J23SpCA_32 CGGG AAGGGAAACC GACCCGTTTA GTCTGCCGCT CCCAACAGCC CGAGTTTCCA GCCCCTTGAG TAAGGGGCAA GAAACCTAAG CCGCCTGGTG GATGGCTCGG CTCGGGGCGC TGACGAAGG
J23SpCA_3 GG AAGGGAAACC GACCCGTTTA GTCTGCCGCT CCCAACAGCC CGAGTTTCCA GCCCCTTGAG TAAGGGGCAA GAAACCTAAG CCGCCTGGTG GATGGCTCGG CTCGGGGCGC TGACGAAGGG
J23SpCA_17 CGGGGA TGTAAGCGGG AAGGGAAACC GACCCGTTTA GTCTGCCGCT CCCAACAGCC CGAGTTTCCA GCCCCTTGAG TAAGGGGCAA GAAACCTAAG CCGCCTGGTG GATGGCTCGG CTCGGG
J23SpCA_2 GCGGGGA TGTAAGCGGG AAGGGAAACC GACCCGTTTA GTCTGCCGCT CCCAACAGCC CGAGTTTCCA GCCCCTTGAG TAAGGGGCAA GAAACCTAAG CCGCCTGGTG GATGGCTCGG CTCGGG
J23SpCA_5 CGGGTTTGAT GGGGCGGGGA TGTAAGCGGG AAGGGAAACC GACCCGTTTA GTCTGCCGCT CCCAACAGCC CGAGTTTCCA GCCCCTTGAG TAAGGGGCAA GAAACCTAAG CCGCCTGGTG GATGGC
J23SpCA_4 CGGGTTTGAT GGGGCGGGGA TGTAAGCGGG AAGGGAAACC GACCCGTTTA GTCTGCCGCT CCCAACAGCC CGAGTTTCCA GCCCCTTGAG TAAGGGGCAA GAAACCTAAG CCGCCTGGTG GATG
J23SpCA_4 CGGGTTTGAT GGGGCGGGGA TGTAAGCGGG AAGGGAAACC GACCCGTTTA GTCTGCCGCT CCCAACAGCC CGAGTTTCCA GCCCCTTGAG TAAGGGGCAA GAAACCTAAG CCGCCTGGTG GAT
J23SpCA_40 CGGG AAGGGAAACC GACCCGTTTA GTCTGCCGCT CCCAACAGCC CGAGTTTCCA GCCCCTTGAG TAAGGGGCAA GAAACCTAAG CCGCCTGGTG GATGGCTCGG CTCGGGGCGC TGACGAAG
J23SpCA_3 GGG AAGGGAAACC GACCCGTTTA GTCTGCCGCT CCCAACAGCC CGAGTTTCCA GCCCCTTGAG TAAGGGGCAA GAAACCTAAG CCGCCTGGTG GATGGCTCGG CTCGGGGCGC TGACGAAGG
J23SpCA_7 GGG AAGGGAAACC GACCCGTTTA GTCTGCCGCT CCCAACAGCC CGAGTTTCCA GCCCCTTGAG TAAGGGGCAA GAAACCTAAG CCGCCTGGTG GATGGCTCGG CTCGGGGCGC TGACGAAG
J23SpCA_4 GG AAGGGAAACC GACCCGTTTA GTCTGCCGCT CCCAACAGCC CGAGTTTCCA GCCCCTTGAG TAAGGGGCAA GAAACCTAAG CCGCCTGGTG GATGGCTCGG CTCGGGGCGC TGACGAAGG
J23SpCA_3 GGGAAACC GACCCGTTTA GTCTGCCGCT CCCAACAGCC CGAGTTTCCA GCCCCTTGAG TAAGGGGCAA GAAACCTAAG CCGCCTGGTG GATGGCTCGG CTCGGGGCGC TGACGAAGGG CGTG
J23SpCA_2 GCGGG AAGGGAAACC GACCCGTTTA GTCTGCCGCT CCCAACAGCC CGAGTTTCCA GCCCCTTGAG TAAGGGGCAA GAAACCTAAG CCGCCTGGTG GATGGCTCGG CTCGGGGCGC TGACG
J23SpCA_10 GG AAGGGAAACC GACCCGTTTA GTCTGCCGCT CCCAACAGCC CGAGTTTCCA GCCCCTTGAG TAAGGGGCAA GAAACCTAAG CCGCCTGGTG GATGGCTCGG CTCGGGGCGC TGACGAAG
J23SpCA_3 AGGGAAACC GACCCGTTTA GTCTGCCGCT CCCAACAGCC CGAGTTTCCA GCCCCTTGAG TAAGGGGCAA GAAACCTAAG CCGCCTGGTG GATGGCTCGG CTCGGGGCGC TGACGAAGGG
J23SpCA_3 GGGA TGTAAGCGGG AAGGGAAACC GACCCGTTTA GTCTGCCGCT CCCAACAGCC CGAGTTTCCA GCCCCTTGAG TAAGGGGCAA GAAACCTAAG CCGCCTGGTG GATGGCTCGG CTCGGG
J23SpCA_16 CGGGTTTGAT GGGGCGGGGA TGTAAGCGGG AAGGGAAACC GACCCGTTTA GTCTGCCGCT CCCAACAGCC CGAGTTTCCA GCCCCTTGAG TAAGGGGCAA GAAACCTAAG CCGCCTGGTG
J23SpCA_3 CGGGTTTGAT GGGGCGGGGA TGTAAGCGGG AAGGGAAACC GACCCGTTTA GTCTGCCGCT CCCAACAGCC CGAGTTTCCA GCCCCTTGAG TAAGGGGCAA GAAACCTAAG CCGCCTGGTG G
J23SpCA_2 GA CGGGTTTGAT GGGGCGGGGA TGTAAGCGGG AAGGGAAACC GACCCGTTTA GTCTGCCGCT CCCAACAGCC CGAGTTTCCA GCCCCTTGAG TAAGGGGCAA GAAACCTAAG CCGCCTGGTG G
J23SpCA_2 G AAGGGAAACC GACCCGTTTA GTCTGCCGCT CCCAACAGCC CGAGTTTCCA GCCCCTTGAG TAAGGGGCAA GAAACCTAAG CCGCCTGGTG GATGGCTCGG CTCGGGGCGC TGACGAAG
J23SpCA_5 GGGAAACC GACCCGTTTA GTCTGCCGCT CCCAACAGCC CGAGTTTCCA GCCCCTTGAG TAAGGGGCAA GAAACCTAAG CCGCCTGGTG GATGGCTCGG CTCGGGGCGC TGACGAAGGG C
J23SpCA_4 GGGAAACC GACCCGTTTA GTCTGCCGCT CCCAACAGCC CGAGTTTCCA GCCCCTTGAG TAAGGGGCAA GAAACCTAAG CCGCCTGGTG GATGGCTCGG CTCGGGGCGC TGACGAAGGG CG
J23SpCA_4 CGGG AAGGGAAACC GACCCGTTTA GTCTGCCGCT CCCAACAGCC CGAGTTTCCA GCCCCTTGAG TAAGGGGCAA GAAACCTAAG CCGCCTGGTG GATGGCTCGG CTCGGGGCGC TGACG
J23SpCA_3 AAACC GACCCGTTTA GTCTGCCGCT CCCAACAGCC CGAGTTTCCA GCCCCTTGAG TAAGGGGCAA GAAACCTAAG CCGCCTGGTG GATGGCTCGG CTCGGGGCGC TGACGAAGGG CGTGG
J23SpCA_10 GGGAAACC GACCCGTTTA GTCTGCCGCT CCCAACAGCC CGAGTTTCCA GCCCCTTGAG TAAGGGGCAA GAAACCTAAG CCGCCTGGTG GATGGCTCGG CTCGGGGCGC TGACGAAGGG
J23SpCA_5 GGGAAACC GACCCGTTTA GTCTGCCGCT CCCAACAGCC CGAGTTTCCA GCCCCTTGAG TAAGGGGCAA GAAACCTAAG CCGCCTGGTG GATGGCTCGG CTCGGGGCGC TGACGAAGG
J23SpCA_3 GAAACC GACCCGTTTA GTCTGCCGCT CCCAACAGCC CGAGTTTCCA GCCCCTTGAG TAAGGGGCAA GAAACCTAAG CCGCCTGGTG GATGGCTCGG CTCGGGGCGC TGACGAAGGG CG
J23SpCA_4 GGAAACC GACCCGTTTA GTCTGCCGCT CCCAACAGCC CGAGTTTCCA GCCCCTTGAG TAAGGGGCAA GAAACCTAAG CCGCCTGGTG GATGGCTCGG CTCGGGGCGC TGACGAAGGG
J23SpCA_5 GAAACC GACCCGTTTA GTCTGCCGCT CCCAACAGCC CGAGTTTCCA GCCCCTTGAG TAAGGGGCAA GAAACCTAAG CCGCCTGGTG GATGGCTCGG CTCGGGGCGC TGACGAAGGG
J23SpCA_3 GGAAACC GACCCGTTTA GTCTGCCGCT CCCAACAGCC CGAGTTTCCA GCCCCTTGAG TAAGGGGCAA GAAACCTAAG CCGCCTGGTG GATGGCTCGG CTCGGGGCGC TGACGAAGG
J23SpCA_10 GGGAAACC GACCCGTTTA GTCTGCCGCT CCCAACAGCC CGAGTTTCCA GCCCCTTGAG TAAGGGGCAA GAAACCTAAG CCGCCTGGTG GATGGCTCGG CTCGGGGCGC TGACGAAG
J23SpCA_3 GGAAACC GACCCGTTTA GTCTGCCGCT CCCAACAGCC CGAGTTTCCA GCCCCTTGAG TAAGGGGCAA GAAACCTAAG CCGCCTGGTG GATGGCTCGG CTCGGGGCGC TGACGAAG
J23SpCA_3 AAACC GACCCGTTTA GTCTGCCGCT CCCAACAGCC CGAGTTTCCA GCCCCTTGAG TAAGGGGCAA GAAACCTAAG CCGCCTGGTG GATGGCTCGG CTCGGGGCGC TGACGAAGGG
J23SpCA_3 CGGGTTTGAT GGGGCGGGGA TGTAAGCGGG AAGGGAAACC GACCCGTTTA GTCTGCCGCT CCCAACAGCC CGAGTTTCCA GCCCCTTGAG TAAGGGGCAA GAAACCTAAG CCGCCTG
J23SpCA_4 AACC GACCCGTTTA GTCTGCCGCT CCCAACAGCC CGAGTTTCCA GCCCCTTGAG TAAGGGGCAA GAAACCTAAG CCGCCTGGTG GATGGCTCGG CTCGGGGCGC TGACGAAGGG C
J23SpCA_4 CCCGTTTA GTCTGCCGCT CCCAACAGCC CGAGTTTCCA GCCCCTTGAG TAAGGGGCAA GAAACCTAAG CCGCCTGGTG GATGGCTCGG CTCGGGGCGC TGACGAAGGG CGTGGCAAGC TGCG
J23SpCA_2 GTTTA GTCTGCCGCT CCCAACAGCC CGAGTTTCCA GCCCCTTGAG TAAGGGGCAA GAAACCTAAG CCGCCTGGTG GATGGCTCGG CTCGGGGCGC TGACGAAGGG CGTGGCAAGC TGCG
J23SpCA_2 GTTTA GTCTGCCGCT CCCAACAGCC CGAGTTTCCA GCCCCTTGAG TAAGGGGCAA GAAACCTAAG CCGCCTGGTG GATGGCTCGG CTCGGGGCGC TGACGAAGGG CGTGGCAAGC TG
J23SpCA_4 GTTTGAT GGGGCGGGGA TGTAAGCGGG AAGGGAAACC GACCCGTTTA GTCTGCCGCT CCCAACAGCC CGAGTTTCCA GCCCCTTGAG TAAGGGGCAA GAAACCTAAG CCGCCTGGTG
J23SpCA_4 GTTTGAT GGGGCGGGGA TGTAAGCGGG AAGGGAAACC GACCCGTTTA GTCTGCCGCT CCCAACAGCC CGAGTTTCCA GCCCCTTGAG TAAGGGGCAA GAAACCTAAG CCGCCTGG
J23SpCA_5 AACC GACCCGTTTA GTCTGCCGCT CCCAACAGCC CGAGTTTCCA GCCCCTTGAG TAAGGGGCAA GAAACCTAAG CCGCCTGGTG GATGGCTCGG CTCGGGGCGC TGACGAAGGG
J23SpCA_3 AAACC GACCCGTTTA GTCTGCCGCT CCCAACAGCC CGAGTTTCCA GCCCCTTGAG TAAGGGGCAA GAAACCTAAG CCGCCTGGTG GATGGCTCGG CTCGGGGCGC TGACGAAG
J23SpCA_3 AACC GACCCGTTTA GTCTGCCGCT CCCAACAGCC CGAGTTTCCA GCCCCTTGAG TAAGGGGCAA GAAACCTAAG CCGCCTGGTG GATGGCTCGG CTCGGGGCGC TGACGAAG
J23SpCA_5 C GACCCGTTTA GTCTGCCGCT CCCAACAGCC CGAGTTTCCA GCCCCTTGAG TAAGGGGCAA GAAACCTAAG CCGCCTGGTG GATGGCTCGG CTCGGGGCGC TGACGAAGGG C
J23SpCA_3 C GACCCGTTTA GTCTGCCGCT CCCAACAGCC CGAGTTTCCA GCCCCTTGAG TAAGGGGCAA GAAACCTAAG CCGCCTGGTG GATGGCTCGG CTCGGGGCGC TGACGAAGGG
J23SpCA_3 CGGGGA TGTAAGCGGG AAGGGAAACC GACCCGTTTA GTCTGCCGCT CCCAACAGCC CGAGTTTCCA GCCCCTTGAG TAAGGGGCAA GAAACCTAAG CCGCCTGGTG GATGGC
J23SpCA_7 C GACCCGTTTA GTCTGCCGCT CCCAACAGCC CGAGTTTCCA GCCCCTTGAG TAAGGGGCAA GAAACCTAAG CCGCCTGGTG GATGGCTCGG CTCGGGGCGC TGACGAAGG
J23SpCA_2 GACCCGTTTA GTCTGCCGCT CCCAACAGCC CGAGTTTCCA GCCCCTTGAG TAAGGGGCAA GAAACCTAAG CCGCCTGGTG GATGGCTCGG CTCGGGGCGC TGACGAAGGG
J23SpCA_6 CGGG AAGGGAAACC GACCCGTTTA GTCTGCCGCT CCCAACAGCC CGAGTTTCCA GCCCCTTGAG TAAGGGGCAA GAAACCTAAG CCGCCTGGTG GATGGCTCGG CTCGGG
J23SpCA_6 CCCGTTTA GTCTGCCGCT CCCAACAGCC CGAGTTTCCA GCCCCTTGAG TAAGGGGCAA GAAACCTAAG CCGCCTGGTG GATGGCTCGG CTCGGGGCGC TGACGAAGGG
J23SpCA_3 CGGG AAGGGAAACC GACCCGTTTA GTCTGCCGCT CCCAACAGCC CGAGTTTCCA GCCCCTTGAG TAAGGGGCAA GAAACCTAAG CCGCCTGGTG GATGGCTCGG CTCGG
J23SpCA_5 CCCGTTTA GTCTGCCGCT CCCAACAGCC CGAGTTTCCA GCCCCTTGAG TAAGGGGCAA GAAACCTAAG CCGCCTGGTG GATGGCTCGG CTCGGGGCGC TGACGAAGGG C
J23SpCA_5 CCCGTTTA GTCTGCCGCT CCCAACAGCC CGAGTTTCCA GCCCCTTGAG TAAGGGGCAA GAAACCTAAG CCGCCTGGTG GATGGCTCGG CTCGGGGCGC TGACGAAGG
J23SpCA_2 GACCCGTTTA GTCTGCCGCT CCCAACAGCC CGAGTTTCCA GCCCCTTGAG TAAGGGGCAA GAAACCTAAG CCGCCTGGTG GATGGCTCGG CTCGGGGCGC TGACGAAG
J23SpCA_2 GTTTA GTCTGCCGCT CCCAACAGCC CGAGTTTCCA GCCCCTTGAG TAAGGGGCAA GAAACCTAAG CCGCCTGGTG GATGGCTCGG CTCGGGGCGC TGACGAAGGG C
```

```
J23SpCA_4                                                                                        GTTTA GTCTGCCGCT CCCAACAGCC CGAGTTTCCA GCCCCTTGAG TAAGGGGCAA GAAACCTAAG CCGCCTGGTG GATGGCTCGG CTCGGGGCGC TGACGAAGGG
J23SpCA_3                                                                                       CCGTTTA GTCTGCCGCT CCCAACAGCC CGAGTTTCCA GCCCCTTGAG TAAGGGGCAA GAAACCTAAG CCGCCTGGTG GATGGCTCGG CTCGGGGCGC TGACGAAG
J23SpCA_4                                                          CGGGGA TGTAAGCGGG AAGGGAAACC GACCCGTTTA GTCTGCCGCT CCCAACAGCC CGAGTTTCCA GCCCCTTGAG TAAGGGGCAA GAAACCTAAG CCGCCTGGTG
J23SpCA_2                                                                                        GTTTA GTCTGCCGCT CCCAACAGCC CGAGTTTCCA GCCCCTTGAG TAAGGGGCAA GAAACCTAAG CCGCCTGGTG GATGGCTCGG CTCGGGGCGC TGACGAAGG
J23SpCA_2                                                            GGGA TGTAAGCGGG AAGGGAAACC GACCCGTTTA GTCTGCCGCT CCCAACAGCC CGAGTTTCCA GCCCCTTGAG TAAGGGGCAA GAAACCTAAG CCGCCTGGTG
J23SpCA_3                                                                                  GAAACC GACCCGTTTA GTCTGCCGCT CCCAACAGCC CGAGTTTCCA GCCCCTTGAG TAAGGGGCAA GAAACCTAAG CCGCCTGGTG GATGGCTCGG CTCGGG
J23SpCA_4                                                                                             GTCTGCCGCT CCCAACAGCC CGAGTTTCCA GCCCCTTGAG TAAGGGGCAA GAAACCTAAG CCGCCTGGTG GATGGCTCGG CTCGGGGCGC TGACGAAGGG
J23SpCA_3                                                                                               CTGCCGCT CCCAACAGCC CGAGTTTCCA GCCCCTTGAG TAAGGGGCAA GAAACCTAAG CCGCCTGGTG GATGGCTCGG CTCGGGGCGC TGACGAAGGG
```
