## Supplementary figures and images for "Permuted 23S rRNA is integrated in 50S ribosome particles in *Thermococcus barophilus*"

### Supplementary Figure S1

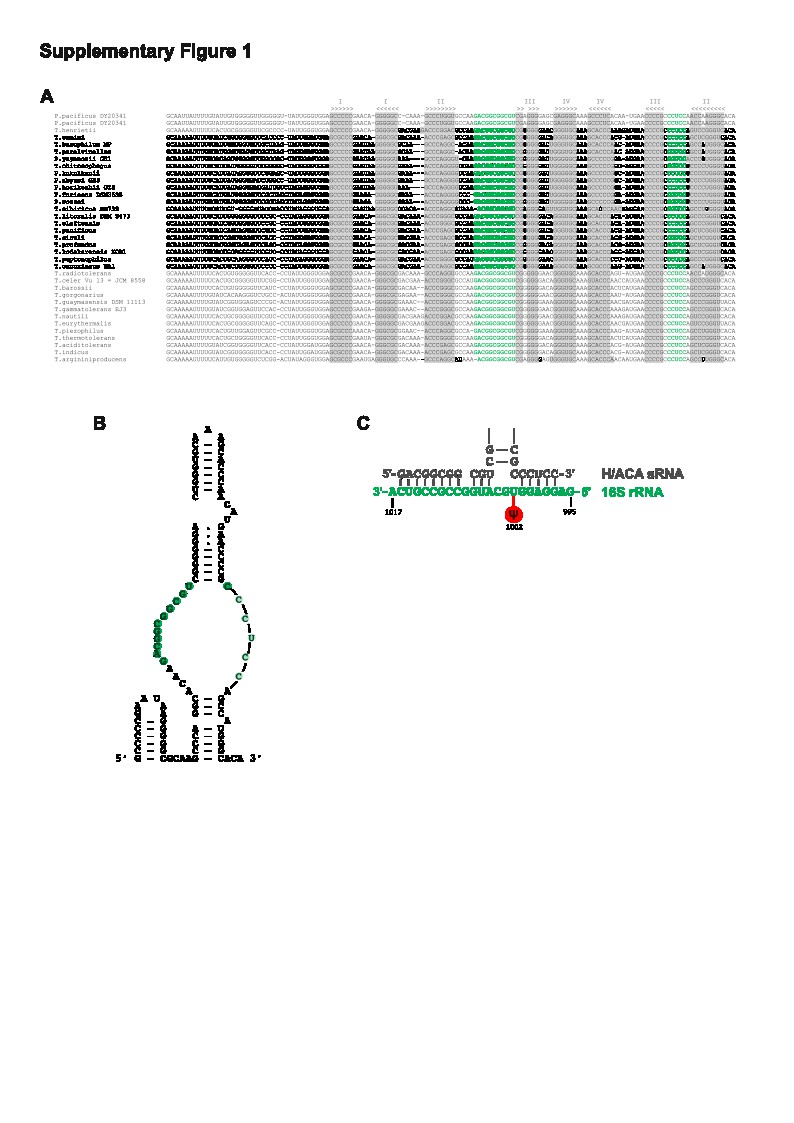

### Supplementary Figure S3

#### Supplementary Figure S3

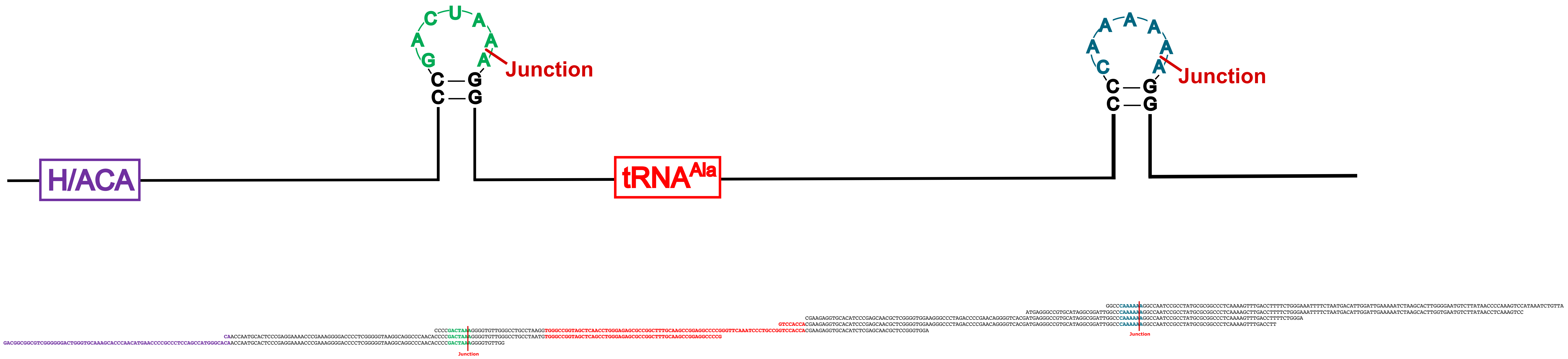
