## Supplementary Figure S4 for "Permuted 23S rRNA is integrated in 50S ribosome particles in *Thermococcus barophilus*"

```
Str1                                                 ...<<<<...<<<<<<<<<<<<<.<<<< <<<<<<<<<<<<<<.............. -------  ......................>>>>>>>>>>>>>>>>>>>>>>>>>>>>>>>>>>>>...
Str2                                                 ...<<<<...<<<<<<<<<<<<<<.....>>>>>>><<<<<<<.............. -------  ......................>>>>>>><<<<<<<.....>>>>>>>>>>>>>>>>>...
Thermococcus barophilus MP           NC_014804.1     GCCCAAAAAATGCCCAGCCCCTTGAGT-AAGGGGCAAGAAACCTAAGCCGCCTGGTG --23S-- CTGCCGCTCCCAACAGCCCGAGTTTCTTGCCCCTGAAAGAGGGGCTGGGCATTTGATA
Thermococcus litoralis DSM 5473      NC_022084.1     GCCCAAAAAATGTCCAGCCCCC-AAGT-AGGGGGTAAGAAACCTAAGCCGCCTGGTG --23S-- CTGCCGCTCCCAACAGCCCGAGTTTCTTGCCCCTGAAAGAGGGGCTGGGCATTTGATA
Thermococcus sibiricus MM 739        NC_012883.1     GCCCAAAAAATGTCCAGCCTCC-AAGT-AGGGGGTAAGAAACTTAAGCCACCTGGTG --23S-- CTGCCGCTTCCAATAGCCCGAGGTTCTTACCCCCGAAA-GGAGGTTGGGCATTTGATA
Thermococcus onnurineus NA1          NC_011529.1     ACCCAAAAAATGCCCAGCCCCCTGAGT-AGGGGGCAGAAAACCTAAGCCGCCTGGTG --23S-- CTGCCGCTCCCAACAGCCCGAGGTTTCTGCCTCTGAAAGGGGGGCTGGGCATTTGATA
Thermococcus piezophilus CDGS        NZ_CP015520.1   ACCCAAAAAATGCCCAGCCCTCTAAGT-AGAGGGTAGAAAACCTAAGCCGCCTGGTG --23S-- CTGCCGCTCCCAACAGCCCGAGGTTTCTGCCCTCGAAAGGAGGGCTGGGCATTTGATA
Thermococcus celer Vu 13             NZ_CP014854.1   ACCCAAAAAATGCCCAGCCCCCTGAGT-AGGGGGCAGAGAACCTAAGCCGTCTGGTG --23S-- CTGCCGCTCCCAATCGCCCGAGGTTTCTGCCTCTGAAAGGGGGGCTGGGCATTTGATA
Thermococcus barossii SHCK-94        NZ_CP015101.1   ACCCAAAAAATGCCCAGCCCCCTGAGT-AGGGGGCAGAAAACCTAAGCCGCCTGGTG --23S-- CTGCCGCTCCCAACAGCCCGAGGTTTCTGCCTCTGAAAGGGGGGCTGGGCATTTGATA
Thermococcus radiotolerans EJ2       NZ_CP015106.1   ACCCAAAAAATGCCCAGCCCCCTGAGT-AGGGGGCAGAAAACCTAAGCCGCCTGGTG --23S-- CTGCCGCTCCCAACAGCCCGAGGTTTCTGCCTCTGAAAGGGGGGCTGGGCATTTGATA
Thermococcus thermotolerans          NZ_CP102602.1   ACCCAAAAAATGCCCAGCCCCCTGAGT-AGGGGGCAGAAAACCTAAGCCGCCTGGTG --23S-- CTGCCGCTCCCAACAGCCCGAGGTTTCTGCCTCTGAAAGGGGGGCTGGGCATTTGATA
Thermococcus camini                  NZ_LR881183.1   ACCCAAAAAATGCCCAGCCCCCTGAGT-AGGGGGCAGAAAACTTAAGCCGTCTGGTG --23S-- CTGCCGCTCCCAACAGCCCGAGGTTTCTGCCTCTGAAAGGGGGGCTGGGCATTTGATA
Thermococcus cleftensis              NC_018015.1     ACCCAAAAAATGCCCAGCCCCCTGAGT-AGGGGGCAGAAAACCTAAGCCGCCTGGTG --23S-- CTGCCGCTCCCAACAGCCCGAGGTTTCTGCCCCTGAAAGGGGGGCTGGGCATTTGATA
Thermococcus pacificus P-4           NZ_CP015102.1   ACCCAAAAAATGCCCAGCCCCCTGAGT-AGGGGGCAGAAAACCTAAGCCGCCTGGTG --23S-- CTGCCGCTCCCAACAGCCCGAGGTTTCTGTCCCTGAAAGGGGGGCTGGGCATTTGATA
Thermococcus siculi RG-20            NZ_CP015103.1   ACCCAAAAAATGCCCAGCCCCCTGAGT-AGGGGGCAGAAAACCTAAGCCGCCTGGTG --23S-- CTGCCGCTCCCAACAGCCCGAGGTTTCTGTCCCTGAAAGGGGGGCTGGGCATTTGATA
Thermococcus indicus IOH1            NZ_CP040846.1   ACCCAAAAAATGCCCAGCCCCCTGAGT-AGGGGGCAGAAAACTTAAGCCGTCTGGTG --23S-- CTGCCGCTCCCAACAGCCCGAGGTTTCTGCCTCTGAAAGGGGGGCTGGGCATTTGATA
Thermococcus aciditolerans SY113     NZ_CP041932.1   ACCCAAAAAATGCCCAGCCCCCTGAGT-AGGGGGCAGAAAACTTAAGCCGTCTGGTG --23S-- CTGCCGCTCCCAATAGCCCGAGGTTTCTGCCTCTGAAAGGGGGGCTGGGCATTTGATA
Pyrococcus yayanosii CH1             NC_015680.1     GCCCAAAAAATGCCCAGCCCCCTGAGT-AGGGGGCAAGAACGCTAAGCCGCCCGGTG --23S-- CTGCCGCTCCCAATCGCCCGAGGTTCTTGCCCCCGAAAAGGGGGCTGGGCATTTGATA
Thermococcus argininiproducens       NZ_CP080572.1   GCCCAAAAAATGTCCAGCCTCC-AAGT-AGGGGGTAAGAAACCTAAGCCACCTGGTG --23S---CTGCCGCTCCCAATAGCCCGAGGTTCTTACCCCCGAAA-GGGGGCTGGGCATTTGATA
Thermococcus chitonophagus GC74      NZ_CP015193.1   GCCCAAGAAGTGCCTGGCCCCTCCGGTAGGGGGGCTAGGACTCTAAGCCGCCCGGTG --23S-- CTGCCGCCCCCAATCGCCCGAGGTCCTACCCCCCGAAGGAGGGGCCAGGCACTTGACA
Pyrococcus kukulkanii NCB100         NZ_CP010835.1   GCCCAAGAAGTGTCTGGCCCCTCCGGTAGGGGGGCTAGGACGCTAAGCCGCCCGGTG --23S-- CTGCCGCCCCCAATCGCCCGAGGTCCTACCCCCCGAAGGAGGGGCCAGGCACTTGACA
Pyrococcus abyssi GE5                NC_000868.1     GCCCAAGAAGTGCCTGGCCCCTCCGGT-AGGGGGTCAGGACACTAAGCCGCCCGGTG --23S-- CTGCCGCCCCCAATCGCCCGAGGTCCTGTCCCCCGAAGGAGGGGCCAGGCACTTGACA
Pyrococcus horikoshii OT3            NC_000961.1     GCCCAAGAAGTGCCTGGCCCCTCCGGT-AGGGGGTCAGGACTCTAAGCCGCCCGGTG --23S-- CTGCCGCTCCCAATCGCCCGAGGTCCTGTCCCCCGAAGGAGGGGCCAGGCACTTGACA
Pyrococcus furiosus DSM 3638         NC_003413.1     GCCCAAGAAGTGCCTGGCCCCTCCGGTAGAGGGGCTAGGACGCTAAGCCGCCCGGTG --23S-- CCGCCGCCCCTAATCGCCCGAGGTCCTGCCCCTCGAAGGAGGGGCCAGGCACTTGACA
Pyrococcus woesei                    NZ_CP179867.1   GCCCAAGAAGTGCCTGGCCCCTCCGGTAGAGGGGCTAGGACGCTAAGCCGCCCGGTG --23S-- CCGCCGCCCCTAATCGCCCGAGGTCCTGCCCCTCGAAGGAGGGGCCAGGCACTTGACA
Palaeococcus pacificus DY20341       NZ_CP006019.1   ACCCAAAAAACTGCCCAGCCTCCATAGTA-GGGGCAAGTACACTAAACCGCCTGGTG --23S-- CTGCCGTTCCCAATTGCCCGAGGTC-TTGCCCTGAATTG-GGGGCTGGGCGTTTGAAA
Thermococcus guaymasensis DSM 11113  NZ_CP007140.1   GTCCAGAAAACGTCCCGCCCGGTGGCA-------------ACTAAGCCGCCTGGTG --23S-- CTGCCGCTCCCAACAGCCC-------------GAGTGCCCCGGGCGGGACGTCTGAGA
Thermococcus gammatolerans EJ3       NC_012804.1     GTCCAGAAAACGCCCCGCCCGGTGGCA-------------ACTAAGCCGCCTGGTG --23S-- CTGCCGCTCCCAACAGCCC-------------GAGTGCCCCGGGCGGGGCGTCTGAGA
Thermococcus henrietii EXT12c        NZ_LT900021.1   GTCCAGAAAACGCCCCGCCCGGTGGCA-------------ACTAAGCCGCCTGGTG --23S-- CTGCCGCTCCCAACAGCCC-------------GAGTGCCCCGGGCGGGGCGTCTGAGA
Thermococcus nautili 30-1            NZ_CP007264.1   GTCCAGAAAACGCCCCGCCCGGTGGCA-------------ACTAAGCCGCCTGGTG --23S-- CTGCCGCTCCCAACAGCCC-------------GAGTGCCCCGGGCGGGGCGTCTGAGA
Thermococcus eurythermalis A501      NZ_CP008887.1   GTCCAGAAAACGCCCCGCCCGGTGGCA-------------ACTAAGCCGCCTGGTG --23S-- CTGCCGCTCCCAACAGCCC-------------GAGTGCCCCGGGCGGGGCGTCTGAGA
Thermococcus kodakarensis KOD1       NC_006624.1     GTCCAGAAAACGTCCCGCCCGGTGGCA-------------ACTAAGCCGCCTGGTG --23S-- CTGCCGCTCCCAACAGCCC------------GAGTGCTACCGGGCGGGACGTCTGAGA
Thermococcus peptonophilus OG-1      NZ_CP014750.1   GTCCAGGAAACGTCCCGCCCGGTGGCA-------------ACTAAGCCGCCTGGTG --23S-- CTGCCGCTCCCAACAGCCC------------GAGTGCTACCGGGCGGGACGTCTGAGA
Thermococcus gorgonarius W-12        NZ_CP014855.1   GTCCAGAAAACGCCCCGCCCGGTGGCA-------------ACTAAGCCGCCTGGTG --23S-- CTGCCGCTCCCAACAGCCC-------------GAGTGCCCCGGGCGGGGCGTCTGATA
Thermococcus profundus DT 5432       NZ_CP014862.1   GTCCAGGAAACGTCCCGCCCGGTGGCA-------------ACTAAGCCGCCTGGTG --23S-- CTGCCGCTCCCAACAGCCC-------------GAGTGCCCCGGGCGGGACGTCTGAGA
Str1                                                 ...<<<<...<<<<<<<<<<<<<.<<<<............................. -------  ....................................>>>>>>>>>>>>>>>>>>>>>...
```
